# Quantitative proteomics of hamster lung tissues infected with SARS-CoV-2 reveal host-factors having implication in the disease pathogenesis and severity

**DOI:** 10.1101/2021.03.09.434371

**Authors:** Voddu Suresh, Varshasnata Mohanty, Kiran Avula, Arup Ghosh, Bharti Singh, R. Rajendra Kumar Reddy, Amol Ratnakar Suryawanshi, Sunil K. Raghav, Soma Chattopadhyay, Punit Prasad, Rajeeb Kumar Swain, Rupesh Dash, Ajay Parida, Gulam Hussain Syed, Shantibhusan Senapati

**Author notes:** These authors contributed equally (Voddu Suresh, Varshasnata Mohanty, Kiran Avula). Address for correspondence: Shantibhusan Senapati, BVSc & AH.; PhD, Scientist, Institute of Life Sciences, Nalco Square, Bhubaneswar-751023, Odisha, India., Contact: And Gulam Hussain Syed, PhD, Scientist, Institute of Life Sciences, Nalco Square, Bhubaneswar-751023, Odisha, India., Contact.

## Abstract

Syrian golden hamsters (*Mesocricetus auratus*) infected by severe acute respiratory syndrome coronavirus 2 (SARS-CoV-2) manifests lung pathology that resembles human COVID-19 patients. In this study, efforts were made to check the infectivity of a local SARS-CoV-2 isolate in hamster model and evaluate the differential expression of lung proteins during acute infection and convalescence. The findings of this study confirm the infectivity of this isolate *in vivo*. Analysis of clinical parameters and tissue samples shows a similar type of pathophysiological manifestation of SARS-CoV-2 infection as reported earlier in COVID-19 patients and hamsters infected with other isolates. The lung-associated pathological changes were very prominent on the 4th day post-infection (dpi), mostly resolved by 14dpi. Here, we carried out quantitative proteomic analysis of the lung tissues from SARS-CoV-2-infected hamsters at day 4 and day 14 post infection. This resulted in the identification of 1,585 differentially expressed proteins of which 68 proteins were significantly altered among both the infected groups. Pathway analysis revealed complement and coagulation cascade, platelet activation, ferroptosis and focal adhesion as the top enriched pathways. In addition, we also identified altered expression of two pulmonary surfactant-associated proteins (Sftpd and Sftpb), known for their protective role in lung function. Together, these findings will aid in the identification of candidate biomarkers and understanding the mechanism(s) involved in SARS-CoV-2 pathogenesis.

**Graphical abstract:** 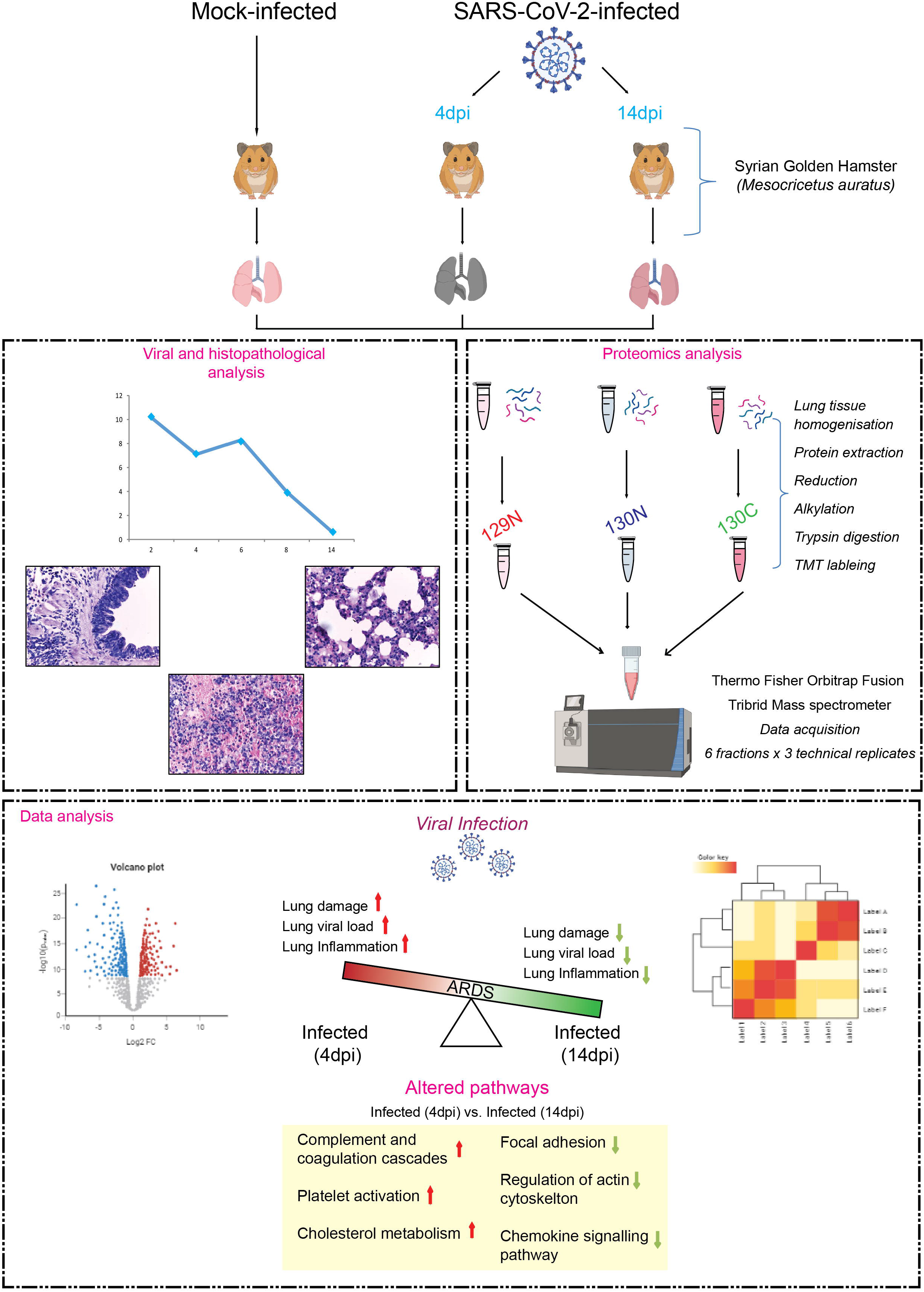

## Introduction

The recent outbreak of severe acute respiratory syndrome coronavirus 2 (SARS-CoV-2) has emerged as a global public health crisis affecting millions of people worldwide. SARS-CoV-2, the causative agent of the coronavirus disease 2019 (COVID-19), primarily infects the respiratory tract resulting in respiratory failure consistent with the acute respiratory distress syndrome (ARDS). The aggressive inflammatory response associated with infection leads to tissue damage and fatal lung injury. The death of the virus affected epithelial cells and endothelial cells, and activation of resident dendritic cells, monocytes, and macrophages result in enhanced production of cytokines and chemokines (cytokine storm). This triggers further activation and recruitment of immune cells leading to tissue damage and exacerbation of respiratory distress and disease severity. Severe clinical manifestations involving multi-organ failure have resulted in high morbidity and mortality, thus demanding immediate therapeutic measures and disease management. In this regard, a better understanding of viral and host factors involved in this disease's pathogenesis has paramount significance. After identifying the SARS-CoV-2 virus, multiple genomic and proteomics-based approaches have been adapted to understand the host response to this viral infection. In the recent past, different studies have been conducted using proteomics technology to understand the virus-host protein interactome, changes in host protein expression upon virus infection, and for diagnosis of SARS-CoV-2 infection. Most of these studies have been carried out with patients’ liquid specimens like serum, plasma, sputum, and bronchoalveolar lavage [1–5]. Clinical manifestations of COVID-19 are primarily related to the lung; hence clinical proteomics of infected lung tissues are essential to understand and combat this disease. The information obtained from such studies will provide the rationale for designing novel diagnostic and therapeutic interventions. Moreover, the identification of differentially expressed proteins signature can serve as markers of COVID-19 disease severity, which will be highly useful for better disease management.

Preclinical animal models that recapitulate multiple sequential events associated with human diseases are precious tools to understand the mechanistic aspects of human disease progression. In human patients, the COVID-19 associated lung pathology is a major clinical concern [6, 7]. The manifestation of similar pulmonary damages in hamsters infected with SARS-CoV-2 virus has emphasized this model’s relevance [8–12]. Further characterization of this model is essential for the development of effective therapeutics and vaccines against this virus. An in-depth understanding of host response to SARS-CoV-2 infection in hamsters will elucidate this model’s similarity or dissimilarity with human patients. Although efforts have been made to identify differentially expressed proteins in diverse body fluids of COVID-19 patients; there is a dearth of evidence related to differentially expressed proteins in human lung tissues at acute and convalescence stages of SARS-CoV-2 infection. Fresh lung tissues of COVID-19 patients during infection or recovery are ethically impossible to obtain. Hence, so far, data obtained from tissues of deceased COVID-19 patients are only available. In this regard, tissues obtained from animals infected with SARS-CoV-2 at different viral infection stages will prove beneficial. In this study, efforts have been made to quantitatively compare host protein levels in lung tissues of hamsters at various stages of SARS-CoV-2 infection. The differentially expressed proteins identified in this study provide information of various host proteins that might have a significant role in the pathogenesis of SARS-CoV-2 infection or disease manifestation. Some of the deregulated proteins identified in this study might be validated in easily assessable body fluids as potential biomarkers for predicting the disease course.

## Materials and methods

### Animal ethics

In this study, attempts were made to evaluate the infectivity of one of the local isolates of SARS-CoV2 (NCBI accession ID- MW559533.2) in the Syrian Golden Hamster model. All the experiments were done with prior approval of Institutional Biosafety Committee (IBSC) and Institutional Animal Ethical Committee (IAEC). The study was carried out adhering to the guidelines of the Committee for the Purpose of Control and Supervision of Experiments on Animals (CPCSEA), Govt. of India.

### Animal studies

For this study, eleven hamsters of the age group 6-7 months were acclimated at the ILS ABSL3 facility for 4-6 days prior to the experiments. As shown in the schematic (**Figure 1A**), on day zero, six animals were infected intranasally with SARS-CoV-2 (10^5^ TCID50) and five animals were mock-infected (only PBS) under intraperitoneal ketamine (200mg/kg) and xylazine (10mg/kg) anaesthesia. The SARS-CoV-2 virus used in this study was isolated from a clinically confirmed local COVID-19 patient (NCBI accession ID- MW559533.2). Virus from the 10th passage was used for this animal challenge study. Out of the six infected animals, three animals were sacrificed on 4dpi (day post-infection), and the other three were sacrificed on 14dpi. Out of the five mock-infected animals, three were sacrificed on 4dpi, and the other two were sacrificed on 14dpi. Throughout the experiment, all the animals were monitored daily, and body weights were recorded on alternate days. On the day of sacrifice, tissues from all the vital organs and other organs like the pancreas, spleen, and gastrointestinal tracts were harvested, preserved, and further processed for histopathological analysis and viral load estimation.

**Figure 1:**
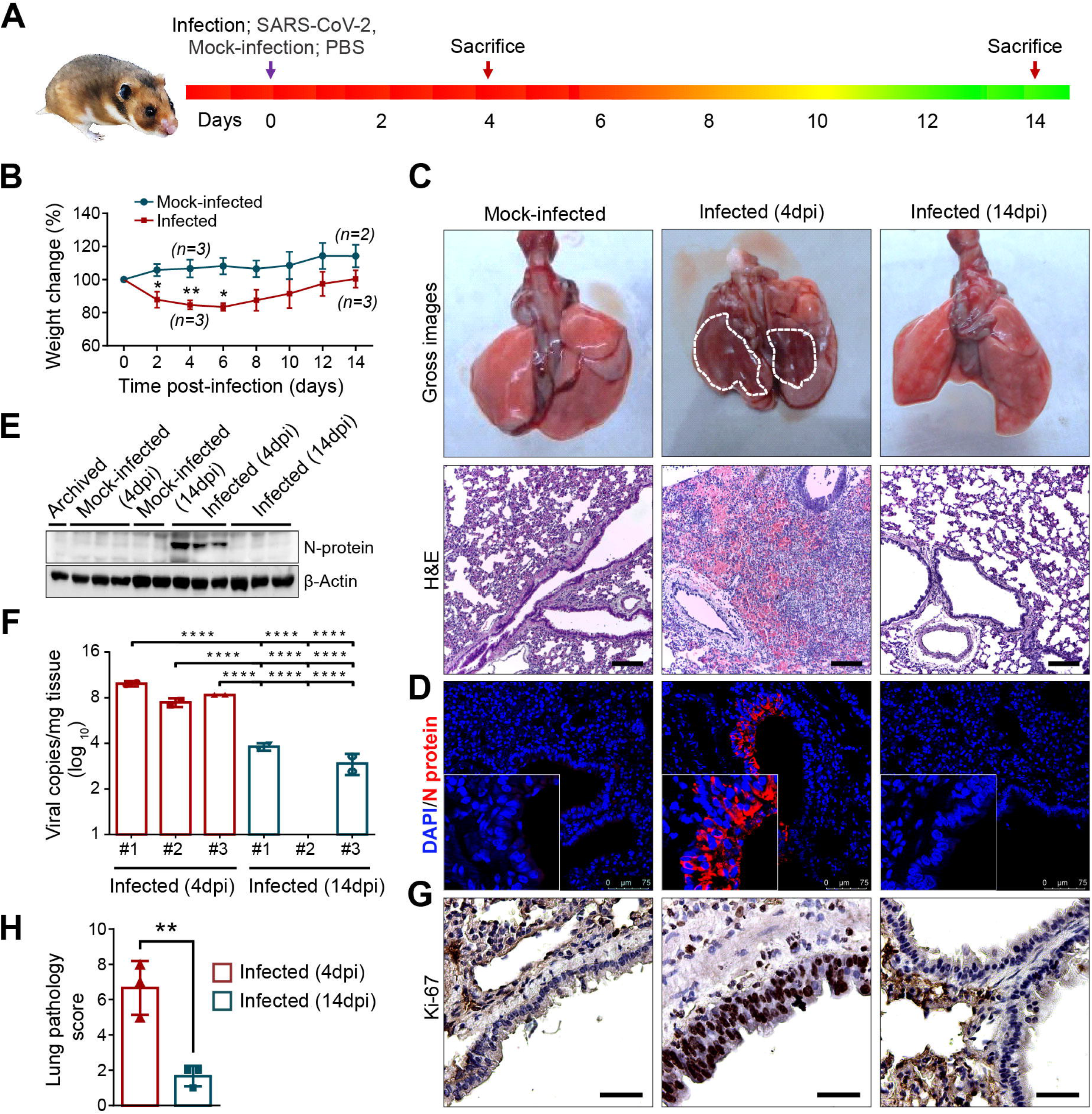
Syrian Golden Hamster model of SARS-CoV-2 infection by using a local SARS-CoV-2 isolate. **(A)** Study design to evaluate the infectivity of local-isolate in Syrian Golden Hamster infection model. **(B)** Graph showing percent body weight change of hamster after mock-infection or SARS-CoV-2 infection. **(C)** Digital images of lungs harvested from mock-infected or infected animals. At 4dpi, infected lungs have massive congestions visible from the surface (highlighted with white border). No gross changes were noticed in mock-infected (4dpi) and infected (14dpi) lungs. Images showing H&E stained lung tissues harvested from mock-infected or infected (4 or 14dpi) hamsters (scale bar = 200 μm). **(D)** Immunofluorescence images of mock-infected or infected lung tissue sections showing presence of Nucleocapsid protein (N) in bronchial and alveolar epithelial cells. **(E)** Immunoblot analysis showing the Nucleocapsid protein expression in the lungs previously archived lysate (normal), mock-infected (4dpi), infected 4dpi or 14dpi lung tissues. **(F)** Graph showing the viral RNA quantification by using RT-qPCR in hamster lung tissues. **(G)** Immunohistochemistry staining with Ki67 showing cell proliferation of bronchial and alveolar cells (scale bar = 50 μm). **(H)** Graph showing pathology score of the lung tissues after the infection (4 and 14dpi).

### Sample collection, storage, and processing

Tissues harvested from different organs, including lungs, were divided into three parts. One part was stored in buffered formalin and further processed for histopathological analysis. The second part was immediately put into Trizol RNA isolation buffer, and the third part was snap-frozen and stored at −80°C until further use. Formalin-fixed tissues were further processed and sectioned for H&E, Immunohistochemistry (IHC), and Immunofluorescence analysis (IF) as reported earlier [13].

### Haemotoxylin and Eosin (H&E) staining and lung pathology scoring

Multiple H&E stained lung sections for each animal were evaluated blindly, and based on the extent of lung epithelial injury and inflammation, a lung pathology score was given for each animal (**Supplementary figure 1**). The scoring was done on a scale of 0-10 based on the severity of lung pathology.

### Immunohistochemistry

Collected tissues were processed and sectioned as reported previously [13, 14]. Sections were deparaffinized, rehydrated, and subjected to antigen retrieval (Vector Laboratories) treatment for 20 min followed by blocking the endogenous peroxidase with 3% hydrogen peroxide in methanol for 20 min. Horse serum (Vector Laboratories) was used for blocking the sections for 30 min at room temperature and incubated with Ki-67 antibody (#VP-RM04; Vector Laboratories, 1:100) overnight at 4°C. Sections were washed twice with 1x PBS and treated with biotinylated anti-rabbit/mouse IgG secondary antibody (Vector Laboratories) for 45 minutes, followed by ABC reagent for 30 min. Diaminobenzidine (Vector Laboratories) was used as a substrate to develop the stain. Hematoxylin was used as a counterstain followed by dehydration with alcohol, clearing with xylene, and mounting with permanent mounting media (Vector Laboratories). Stained sections were observed under the microscope (Leica DM500), and images were taken at different magnifications.

### Immunofluorescence

Sections were deparaffinized, rehydrated, and subjected to antigen retrieval treatment and serum blocking as reported earlier [13]. Sections were incubated with SARS-CoV-2 N protein (Nucleocapsid) (#11-2003; Abgenex, 1:200) or SOD2/Mn-SOD (#NB100-1992; Novus biologicals, 1:100) in a humidified chamber overnight at 4°C. Sections were washed twice with PBST for 5 min each and incubated with anti-Rabbit Alexa Fluor 594 (#A-11037; Life technologies, 1:500) or anti-Mouse Alexa Fluor 594 (#A-11005; Life technologies, 1:500) for 45 min under dark conditions at room temperature. Sections were washed with PBST twice and mounted with ProLong Gold Antifade reagent with DAPI (#P36935; Invitrogen) and visualized using Leica TCS SP8STED confocal microscope.

### Western blot analysis

For Western blot analysis, cells were re-suspended in RIPA buffer (20 mM Tris-HCl [pH 7.5], 150 mM NaCl, 50 mM NaF, 1 mM Na3VO4, 0.1% SDS, and 0.5% TritonX-100) containing the protease inhibitor cocktail (Thermo Scientific). The whole-cell lysates (WCL) were subjected to SDS-PAGE and transferred to nitrocellulose membrane (Thermo Scientific), followed by blocking and immunoblotting with antibodies specific for SARS-CoV-2 Nucleocapsid protein (#11-2003, Abgenex) or β-actin (#4970, CST) [15].

### qRT-PCR

RNA isolation from culture supernatant was performed using QIAamp Viral RNA Mini Kit (#52906, Qiagen), and for hamster tissue samples, TRizol reagent (#10296010, Invitrogen) was used. The isolated RNA was subjected to qRT-PCR for determining the viral load. We performed one-step multiplex real-time PCR using TaqPath™ 1-Step Multiplex Master Mix (#A28526, Thermo Fisher Scientific), targeting three different SARS-CoV-2 genes with primer and probe sets specific for spike (S), nucleocapsid (N), and open reading frame 1 (ORF1). The standard curve for absolute quantification of viral genome copies was generated using log-fold dilutions of plasmid pLVXEF1alpha-nCoV2019-N-2xStrep-IRES-Puro plasmid harboring the SARS-CoV-2 nucleocapsid gene [16].

### Sample preparation for proteomics analysis

The lung tissue samples of SARS-CoV-2-infected on 4 and 14dpi along with mock-infected (4dpi) were processed and homogenized using liquid nitrogen and lysed in lysis buffer containing 4% sodium dodecyl sulfate (SDS) and 50mM triethylammonium bicarbonate (TEABC). The samples were subjected to sonication three times for 10 sec storing on ice to prevent overheating between the sonication and was followed by heating at 90°C for 5 min. The lysates were then incubated at room temperature for cooling and centrifuged at 12,000 rpm for 10 minutes. The protein concentration present in the supernatant was determined using a bicinchoninic acid assay (BCA) kit (Thermo Scientific Pierce) and an equal amount of protein from each group was pooled for further analysis. LC-MS/MS approach was used using isotopomer labels, “tandem mass tags” (TMTs), to determine the relative quantification of proteins [17].

### In-solution digestion and TMT labeling

Protein lysate of 300 μg from pooled samples of each group was reduced by incubating in 10mM Dithiothreitol (DTT) at 60°C for 20 minutes. Alkylation was carried out in the dark with 20 mM iodoacetamide (IAA) at room temperature for 10 minutes. The lysate was further subjected to acetone precipitation, and the pellet was dissolved in 50mM TEABC. Digestion was carried out at 37°C for 16 hours using trypsin (Sciex, #4326682) at a final concentration of 1:20 (w/w). The reaction was acidified using 0.1% formic acid, and the peptides were lyophilized and stored at −80°C until further use.

### LC-MS/MS acquisition

The digested peptides were fractionated into six fractions using StageTip fractionation. Fractionated peptides (6 fractions in triplicate, total 18 runs) were analyzed on Orbitrap Fusion Tribrid mass spectrometer (Thermo Fisher Scientific, Bremen, Germany) interfaced with Easy-nLC 1000 nanoflow liquid chromatography system (Thermo Scientific, Odense, Southern Denmark). Each fraction was reconstituted in solvent A (0.1% formic acid) and loaded onto trap column (75 μm x 2 cm) Thermo Scientific™ Acclaim™ PepMap™ 100 C18 (#164535; Thermo Scientific) (3 μm particle size, pore size 100Å) at a flow rate of 5 μl/min with solvent A (0.1% formic acid in water).

The peptides were resolved on an analytical column (EASY-Spray™ C18 Reversed Phase HPLC Column, 2 μm, 75 μm × 500 mm; Thermo Scientific) using a linear gradient of 7-30% solvent B (0.1% formic acid in 95% acetonitrile) over 100 minutes at a flow rate of 300 nl/min. Data-dependent Mass Spectrometry acquisition was carried out at *top speed* mode with full scans (350-1500 m/z) acquired using an Orbitrap mass analyzer at a mass resolution of 120,000 at 200 m/z. For MS/MS, top intense precursor ions from a 3-second duty cycle were selected and subjected to higher-energy collision dissociation (HCD) with 35% normalized collision energy. The fragment ions were detected at a mass resolution of 30,000 at m/z of 200. Dynamic exclusion was set for 30 seconds. Lock-mass from ambient air (m/z 445.1200025) was enabled for internal calibration as described previously [18].

### Bioinformatics and statistical analysis

The Proteome Discoverer 2.3 (Thermo Scientific, Bremen, Germany) was used to carry out protein identification and quantitation. All raw files were searched against a *Mesocricetus auratus* (Syrian Golden hamster) protein database in Universal Proteins Resource Knowledgebase (UniProt) (32,336 entries) supplemented with common contaminants (116 entries) using SequestHT as a search algorithm. The search parameters included trypsin as the proteolytic enzyme with a maximum allowed missed cleavages to two. Oxidation of methionine and acetylation of protein N-terminus were set as dynamic modifications. In contrast, static modifications included cysteine carbamidomethylation and TMT modification at N-terminus of the peptide and lysine residue. Precursor mass tolerance was set to 10 ppm, and fragment mass tolerance was set to 0.05 Da; all the PSMs were identified with 1% FDR. The differential expression ratios between the groups were calculated. Proteins with differential expression ratios ≥1.5 (upregulated) or ≤ 0.67 (downregulated) were considered differentially expressed. The significance of differences between groups was calculated using *Student’s t-test* (two-tailed), and a p-value ≤0.05 was considered statistically significant. Gene Ontology (GO) and pathway analysis using Kyoto Encyclopedia of Genes and Genomes (KEGG) were performed using the EnrichR online tool http://amp.pharm.mssm.edu/Enrichr/) [19].

Principal component analysis was performed using sample-wise scaled unfiltered normalized protein abundance data using PCAtools. Heatmaps were generated using protein-wise scaled and filtered protein abundance (1.5 and 1.3 up- or down-regulation; p-value ≤0.05 in any comparison) and k-means clustering (k=3).

### Data availability

The raw data files and the MSF files were submitted to the PRIDE partner repository [20] with dataset identifier PXD024547.

## Results and discussion

### Clinical features, viral load and histopathological changes in lungs

Studies using the hamster model of COVID-19 have shown differential infectivity of various SARS-CoV-2 isolates indicating a possible role of viral factors in regulating disease pathogenesis. To investigate the pathogenicity of SARS-CoV-2, Syrian hamsters were infected with the virus, and harvested tissue samples were collected at two different time points (4 & 14dpi) for viral load and pathological analysis (**Figure 1A**). Corroborating earlier reports, a significant weight loss was noticed in all the infected animals at 4dpi. On 4dpi all the animals lost around 15% of their initial body weight (**Figure 1B**). After 6dpi the infected animals started regaining their body weight. During the course of the experiments, no mortality was found in both infected and mock-infected animals. At the time of organ isolation, congestion of lungs was grossly visible only in infected animals at 4dpi (**Figure 1C**). Further histopathological analysis of the 4dpi SARS-CoV-2 infected tissue samples showed the presence of severe pathological lesions in the lungs (**Figure 1C**). Multifocal necrosis and the desquamation of bronchial epithelial cells and infiltration of inflammatory cells were present in the SARS-CoV-2 infected lung tissues (**Figure 1C**). Around 50 % lung area was affected in all the infected animals, and the lesions were patchy throughout the lungs. Necrosuppurative bronchitis and interstitial pneumonia was evident in all the infected animals at 4dpi (**Supplementary figure 1D and 1E)**. Different areas of the infected lung tissues showed consolidation of lungs and hemorrhage (**Supplementary figure 1C)**. These hamsters exhibited severe interstitial pneumonia, as evidenced by the thickening of the alveolar wall, altered alveolar structure, and immune cells’ infiltration (**Supplementary figure 1D)**. Endothelium near the damaged areas was reactive, as evidenced by mononuclear cells' adhesion to the endothelium (**Supplementary figure 1A**). In certain instances, the immune cells have invaded the vessel wall and caused endotheliitis (**Supplementary figure 1B**). In one of the infected animal (4dpi), visceral pleural invasion of immune cells was also noticed (**Supplementary figure 1F**). However, no noticeable histopathological changes were observed in mock-infected hamster lung at any time point. After 14dpi, hamsters infected with SARS-CoV-2 exhibited only mild inflammatory infiltration and tissue damage suggestive of resolution of disease manifestation (**Figure 1C**). Histopathological evaluation of tracheal tissues from all the animals also showed a severe tracheal epithelial and endothelial damage in 4dpi infected tissues compared to 14dpi infected or mock-infected tissues (**Supplementary figure 2)**. The significantly higher lung pathology score at 4dpi infected tissues than 14dpi corroborates the earlier reports and indicates the self-limiting nature of this disease in the hamster (**Figure 1H**).

Immunofluorescence (IF) staining of lung tissue sections with SARS-CoV-2 nucleocapsid (N) protein showed the viral antigen-positive bronchial and alveolar epithelial cells at 4dpi, which were not detected at14dpi tissues (**Figure 1D**). Immunoblot analysis of the lung tissue protein lysates also corroborated the IF findings (**Figure 1E**). The viral genome copy number estimation showed the presence of viral genome in both 4dpi and 14dpi infected lung tissues, however the copy number was significantly low in 14dpi tissues compared to 4dpi tissues (**Figure 1F**). Immunohistochemical staining for Ki67, a marker of cell proliferation, showed marked cellular proliferation (hyperplasia) of bronchial and alveolar cells at 4dpi **(Figure 1G**). This finding corroborates with the earlier report [12]. Together, the findings from this study suggest pathophysiological manifestation of SARS-CoV-2 infection similar to that observed in COVID-19 patients and hamsters infected with other clinical isolates of SARS-CoV-2 [21, 22].

### Quantitative proteomics analysis

#### Molecular alterations in pulmonary pathology induced by SARS-CoV-2 infection

Our data provides an in-depth proteome analysis of the SARS-CoV-2-infected hamster lungs at 4dpi and 14dpi compared to mock-infected (4dpi). The quantitative proteomics data was analyzed using high-resolution LC-MS/MS in triplicates where the raw files were searched using Proteome Discoverer 2.3. The search resulted in the identification of 2,211 proteins (at 1% FDR) and 1,585 proteins were differentially expressed across all the samples. Of these, 50 and 18 proteins were dysregulated (cut off 1.5 fold, p≤0.05) at 4dpi and 14dpi, respectively. A complete list of the proteins identified is provided in **Supplementary Table 1**. All the differentially expressed proteins were further represented as a heat map using hierarchical clustering analysis comparing mock-infected with the infected samples (**Figure 2A**). Interestingly, the principal component analysis (PCA) demonstrated distinct protein expression patterns among the mock-infected 4dpi and 14dpi infected samples, depicting variation among them (**Figure 2B**). In PC1, we observed distinct clusters of data representing the 14dpi and 4dpi lung tissue samples, and separate clusters representing the mock-infected (4dpi) and 14dpi samples in PC2. Technical triplicate samples of the same groups are clustered together.

**Figure 2:**
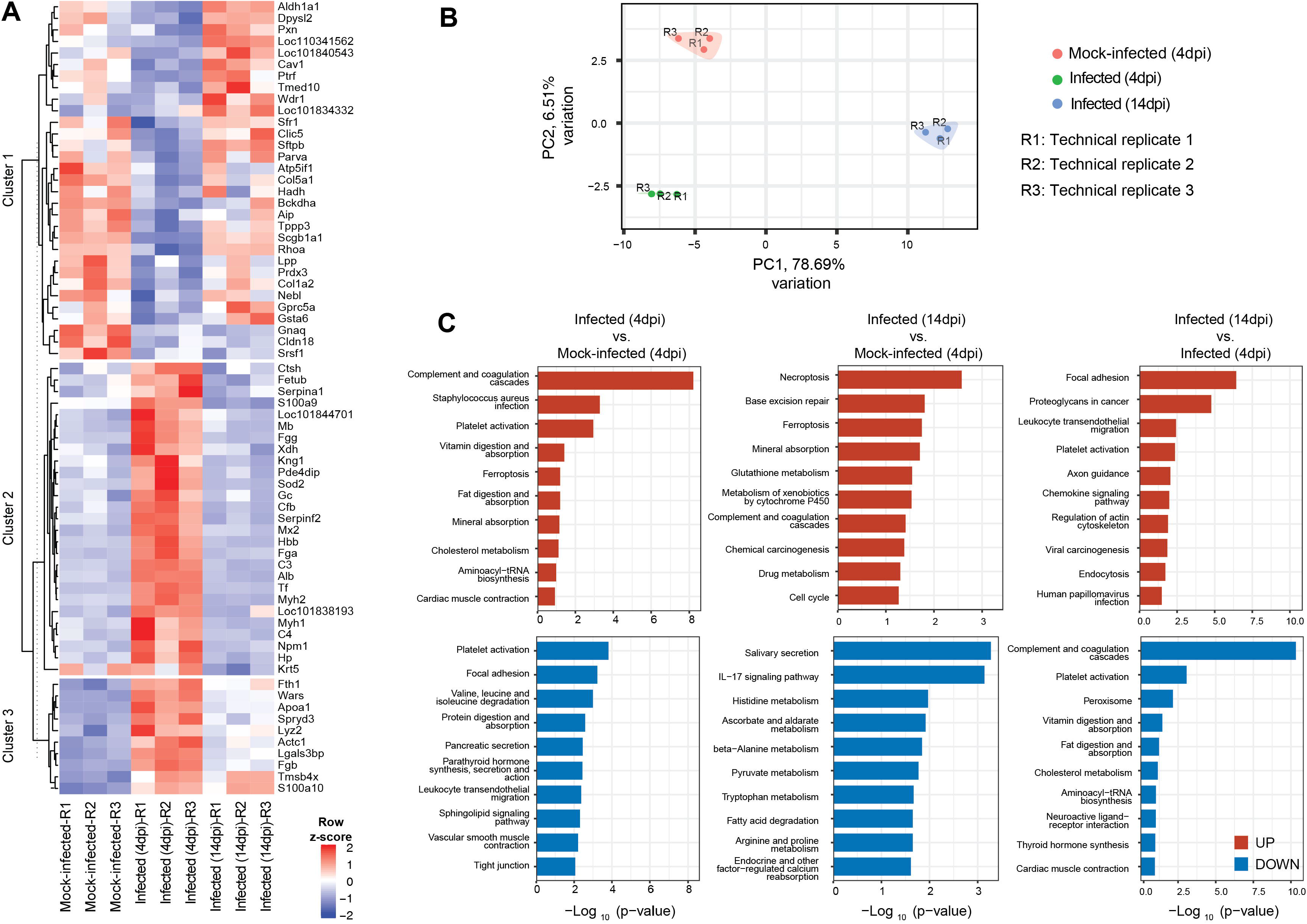
Lung tissue proteome of hamster infected with SARS-CoV-2 at different time points. **(A)** Heatmap of the differential expressed proteins (p ≤ 0.05; fold change cut off of 1.5) were plotted sample wise suggesting the proteomic alterations across the infected samples (4 and 14dpi) compared to mock-infected (4dpi). **(B)** Principal Component Analysis (PCA) was performed and plotted using PCAtools with technical replicates marked by R1, R2 and R3. The PCA plot represents the variance explained by the principal components (denoted by PC) indicating a clear separation of the samples among the three groups i.e mock-infected (4 dpi), Infected (4dpi) and Infected (14dpi). **(C)** Pathway analysis was performed using Enrichr online tool using KEGG database. The enrichment analysis was performed for upregulated and downregulated differentially expressed proteins for all the three groups Infected (4dpi) vs. Mock-infected (4dpi); Infected (14dpi) vs. Mock-infected (4dpi); and Infected (14dpi) vs. Infected (4dpi). The horizontal axis represents the enrichment score −Log_10_ (p-value) of the pathway and the vertical axis represents the pathway category. The red colour represents upregulated proteins and blue bar represents downregulated proteins.

We further compared the dysregulated proteins identified across the two groups (Infected (4dpi) and infected (14dpi)) to identify the number of common and differentially expressed proteins. This comparison was carried out using a Venn diagram (Venny 2.1, https://bioinfogp.cnb.csic.es/tools/venny/). Upon comparison, we identified 41 proteins exclusive to 4dpi tissue samples and 9 exclusive to 14dpi tissue samples when compared with mock-infected (4dpi) tissue samples, while 9 proteins were shared between the groups (**Supplementary Figure 3**). Of the 1,585 proteins dysregulated in the 4dpi lung tissue, 50 were significantly dysregulated (33 upregulated and 17 downregulated; p≤0.05). Among the dysregulated proteins, most proteins were involved in the blood coagulation, integrin signaling pathway, alternative complement activation signaling pathway, and plasminogen activating cascade (PANTHER classification system, version 16.0) [23]. Similarly, of all the identified proteins in the infected (14dpi) set, 18 were significantly dysregulated (9 upregulated and 9 downregulated; p≤0.05). This included proteins belonging to the plasminogen activating cascade and PI3K-Akt signaling pathway. The dysregulated proteins are graphically represented as a heat map and volcano plots representing distinct proteomic patterns between the two groups compared to mock-infected **Supplementary Figure 4A-C**). We also evaluated the altered protein expression during the acute and convalescence stages of infection (14dpi vs. 4dpi) (**Supplementary Figure 4D**).

**Figure 3:**
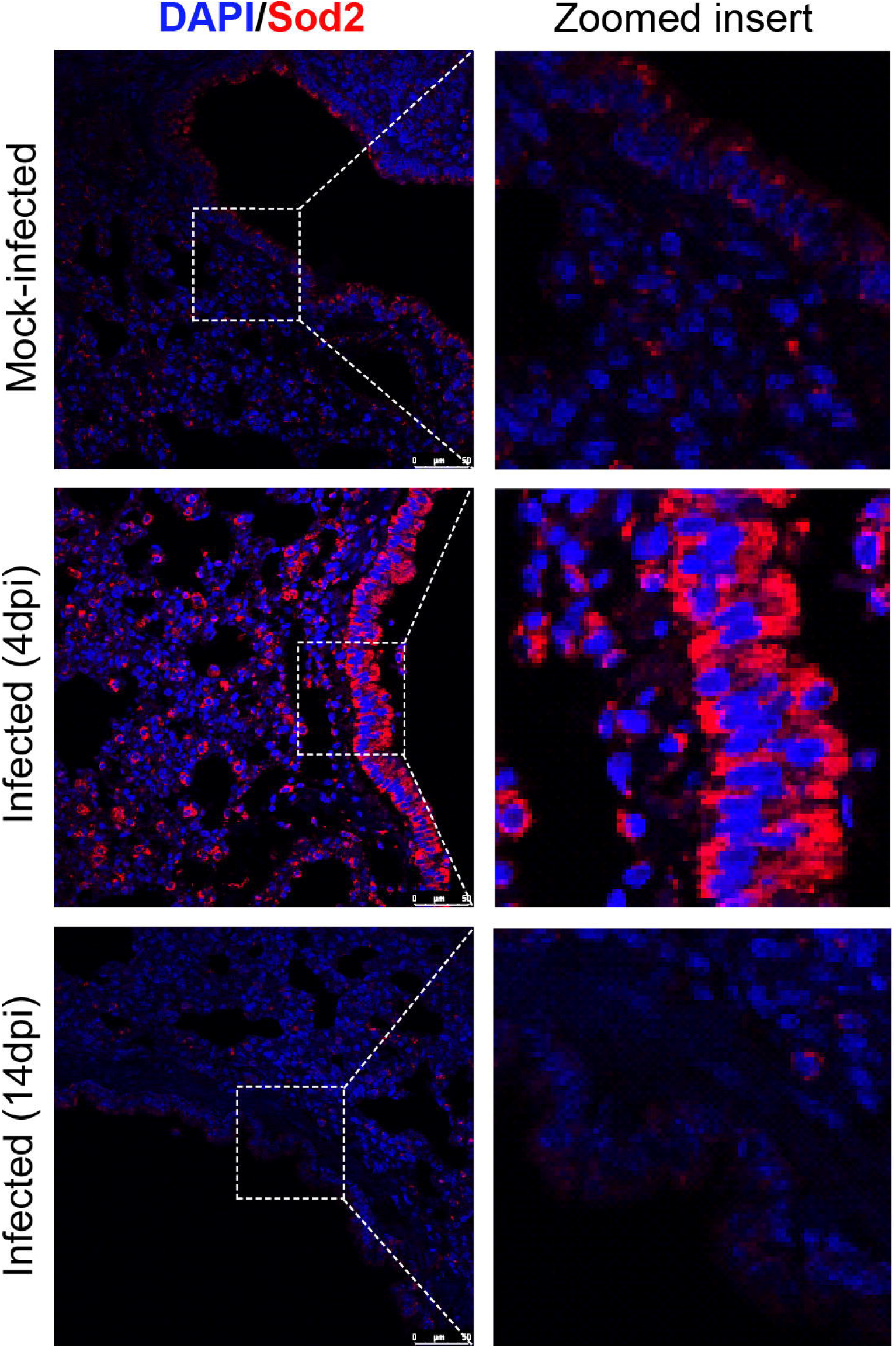
Immunofluorescence images of mock-infected or infected lung tissue sections showing the expression of Sod2 in bronchial and alveolar epithelial cells.

We further compared the significantly dysregulated proteins identified across both the infected (4dpi and 14 dpi) lung tissue compared to mock-infected (4dpi), where a cross-comparison of the down-regulated proteins and upregulated proteins from both the groups did not result in any common protein. However, of the 33 proteins upregulated in the 4dpi tissue samples, 3 proteins such as superoxide dismutase (Sod2), myosin-2 (Myh2), and calgranulin-B (S100a9) were downregulated in the 14dpi tissue samples. These proteins are known to regulate various cellular processes such as cellular localization, cell adhesion, and cellular migration. Similarly, amongst the 33 proteins upregulated in the 4dpi lung tissue, 4 proteins, including Fibrinogen beta chain (Fgb), Ferritin (Fth1), Calpactin I (S100a10), Thymosin (Tmsb4x) were found to be upregulated in a 14dpi sample. These proteins are involved in the regulation of biological processes and exhibit catalytic activity. On further comparison of downregulated proteins identified across both infected (4dpi) and infected (14dpi), two proteins were in common such as serine/arginine-rich splicing factor 1 (Srsf1) and Guanine nucleotide-binding protein G (q) subunit alpha (Gnaq).

#### Functional characteristic of significantly altered proteins

To systematically investigate the molecular differences in the hamster lung owing to SARS-CoV-2 infection, we carried out a Gene Ontology (GO) analysis of the significantly dysregulated proteins. The biological process such as ‘platelet degranulation’, ‘regulated exocytosis’ and ‘fibrinolysis’ were enriched mainly in the proteins upregulated in infected (4dpi) lung tissues, whereas ‘wound healing’, ‘collagen fibril organization’ and ‘actin cytoskeleton reorganization’ were enriched in downregulated proteins (**Supplementary Table 2**). The proteins associated with these pathways include fibrinogen (Fga, Fgb, and Fgg); complement factors (C4a, C3); alpha-2-antiplasmin (Serpinf2); Alpha-1-B Glycoprotein (A1bg), Apolipoprotein A1 (Apoa1) and Alpha-1-acid glycoprotein 1 (Orm1).

Notably, the terms such as ‘regulation of substrate adhesion-dependent cell spreading’ and ‘endothelial cell migration’ were mostly observed in upregulated and downregulated proteins in 14dpi (**Supplementary Table 3**). The reduced expression of the proteins involved in these pathways contributes to dysregulation in actin polymerization. Reports also indicate that any alteration in the cell-matrix adhesion and extracellular matrix during an injury repair can disrupt lung structure, further leading to lung damage. Our results also indicate the dysregulation of collagens in the 14dpi lung tissues. The proteins associated with ‘endothelial cell migration’ include Thymosin beta-4 (TMSB4X) and high mobility group box 1 (HMGB1), both of which have been reported to be potential therapeutic targets in drug discovery. TMSB4X is a small and water-soluble peptide known for its role in angiogenesis, wound healing, and increased metastatic potential of tumor cells [24, 25]. Its ability to induce fibrinolysis makes it an interesting molecular drug target. Recently, HMGB1 has also emerged as a potential target for therapeutic interventions for COVID-19. It is known to play a critical role in various infections, and its elevated expression has been reported in many viral infections, including in serum of COVID-19 patients [26, 27]. This association further results in receptor-dependent responses suggesting its possible role in SARS-CoV-2 infection. Further studies are warranted to uncover the underlying mechanism of HMGB1 and evaluate its inhibitors in the COVID-19 treatment.

#### Pathway analysis

KEGG pathway analyses of the differentially expressed proteins in lung tissue at 4dpi compared to mock-infected (4dpi) tissue revealed that SARS-CoV-2-triggered the activation of complement and coagulation cascade and ferroptosis. The analysis also resulted in the identification of other aberrant pathways such as platelet activation, focal adhesion, and tight junction. Interestingly, we also observed an aberrant expression of proteins associated with necroptosis, cholesterol metabolism, ferroptosis and interleukin-17 signaling pathway in the 14dpi tissue samples suggestive of the underlying mechanisms that lead to tissue damage and lung injury during SARS-CoV-2 infection (**Figure 2C**).

Ferroptosis, is a form of programmed cell death associated with unchecked lipid peroxidation due to the accumulation of lipid reactive oxygen species (ROS) in cells. Its role is well documented in pathophysiological processes of various diseases, such as tumors, nervous system diseases, ischemia-reperfusion injury, kidney injury, and blood diseases [28]. Iron is a pivotal component of the ferroptosis pathway and disruption of iron metabolism has been reported in COVID-19 patients [29]. In our study, we observed increased ferritin (Fth1) levels in 4dpi tissues sample. Its increased expression has also been reported in COVID-19 patients [30, 31].

Platelets are anucleated cell fragments derived from megakaryocytes and essential for physiological hemostasis. Additionally, they are also known for their diverse role during inflammatory and immune response by acting as inflammatory effector cells [32, 33]. They also serve as an indispensable element in coagulation and inflammation and their activated state is associated with cancer progression [34]. Recently, the lung has also been proposed as a platelet biogenesis site, accounting for almost 50% of total platelets [35]. There are reports demonstrating the role of platelets in inflammatory lung syndromes/disorders such as acute respiratory distress syndrome (ARDS), chronic obstructive pulmonary disease (COPD), cystic fibrosis (CF), aspirin exacerbated respiratory disease (AERD), and asthma [33]. The significant enrichment of coagulation proteins in SARS-CoV2 infected tissues indicates disruption of coagulation mechanisms during SARS-CoV-2 infection.

#### Secretory proteins

To investigate the amount of secretory proteins among the pool of differentially expressed proteins in the infected tissue samples, we compared our data with the proteins annotated as ‘secretory proteins (2,640 proteins)’, ‘Secreted in the blood (729 proteins)’, ‘Lung enriched (13 proteins)’, ‘Lung proteome (19,649 proteins)’ and ‘Group enriched (61 proteins expressed in the lung)’ (**Supplementary Figure 4E**). We found four proteins, Ager, secretoglobin family 1A member 1 (Scgb1a1), Sftpb, and Sftpd identified in the current study were specific to lung proteome. Pulmonary surfactant proteins constitute a type of lipoprotein complex comprising of 90% lipids and 10% surfactant proteins (Sftpa, Sftpb, Sftpc, and Sftpd). These surfactant proteins contribute to providing defense against pathogens and play a critical role in efficient gaseous exchange at the air-liquid interface in the alveoli and provide lung stability [36]. We observed aberrant expression of the surfactant proteins (Sftpb, and Sftpd) across the 4 and 14dpi infected samples compared to the mock-infected samples suggesting the decline in the normal functionality of lung respiratory gaseous exchange owing to virus infection. The previous report has suggested the role of viral proteins in modulating the surfactant metabolism and thus, resulting in host immune compromise [37]. Scgb1a1 encodes a member of the secretoglobin family of small secreted proteins, a component of pulmonary surfactant, which is expressed in mucus-secreting cells. This protein is known for its anti-inflammatory/ immunomodulatory and anti-fibrotic functions [38]. We identified severe downregulation of Scgb1a1 in the lung tissue post-infection (4dpi), suggesting respiratory distress owing to virus-mediated lung injury. Reports have also reported its altered expression following lung injury and where its absence is marked with the greater inflammatory response [39–41]. It is also known to regulate alveolar macrophage and respond to inflammation upon virus invasion [41].

As reported earlier, we also noticed distinct damage to the epithelial-endothelial barrier in 4dpi infected lung tissues. This barrier system's damage is believed to be the major mediator of ARDS in different respiratory viral infections, including SARS-CoV-2 [42]. The loss of the epithelial-endothelial barrier allows leakage of blood components into the alveolar lumen and lung interstitium. At the same time, it also allows the leakage of lung proteins into circulation [43]. In the recent past, it is shown that COVID-19 patients who developed ARDS have significantly higher IL-6 and SP-D circulatory levels compared to patients who did not have ARDS [44]. These findings further indicate that identification of pneumoproteins in circulation is an indicator of the severity of lung pathology. In our lung tissue proteome analysis, we noticed significant downregulation of Sftpd protein in tissues with high lung pathology score (4dpi), which suggests that certain proteins might have a differential pattern of expression level in circulation and at the primary site of infection (lungs). Based on several features of pulmonary surfactants, they are believed to have importance in COVID-19 pathogenesis, diagnosis or therapy. It has been proposed that a lower concentration of pulmonary surfactant is a critical risk factor for COVID-19 [45]. Studies have also reported that concentration of SP-A and SP-B were low in BAL of patients before and after onset of ARDS [46]. The low level of pulmonary surfactant proteins detected in 4dpi lung tissues underscores their possible role in the COVID-19 pathology. Altogether, these pulmonary lung surfactants might be considered as potential therapeutic targets which will aid in COVID-19 treatment. However, further studies are required to study its effectiveness and the underlying mechanism(s) associated with virus infection. Further, it suggests the usefulness of the hamster model of SARS-CoV-2 infection in evaluating the therapeutic efficacies of these proteins for COVID-19.

#### Comparison with previous studies

Through an ultra-high-throughput clinical proteomics approach, Messner *et al*., identified protein expression signatures in serum/plasma samples including complement factors, components of coagulation systems, immunomodulators and proinflammatory factors that can classify COVID-19 patients based on WHO grading [1]. The authors have proposed 27 protein groups (23 upregulated and 4 downregulated) as potential biomarkers of disease severity. Out of these upregulated proteins, we noticed six proteins (Cfb, Fga, Fgb, Fgg, Hp and Lgals3bp) are also present in the list of upregulated proteins at 4dpi vs. mock-infected groups of our study. However, Alb, and Tf whose downregulation correlated with COVID-19 severity [1] are found to be upregulated at the 4dpi infection group of our study. There are multiple possible reason for this discrepancy like (a) blood proteome and lung proteome might be different in SARS-CoV-2 infected human or animals, (b) difference in techniques or methods used to analyze samples and (c) possible differential species-dependent (hamster or human) host response to SARS-CoV-2 infection. Similarly, proteomic and metabolomics profiling of serum samples obtained from 46 COVID-19 and 53 control individuals showed deregulation of three major pathways namely complement system, macrophage function and platelet degranulation in severe COVID-19 patients [47]. Study by Park J *et al.,* employed an in-depth proteome profiling of undepleted plasma revealed signatures of proteins involved in neutrophil activation, platelet function, and T cell suppression [3]. Based on the findings, this study's investigators have proposed specific plasma proteins as predictive biomarkers of COVID-19 [3]. In a different study, proteomic analysis of serum samples from early COVID-19 patients also identified differentially expressed proteins known to have a function in SARS-CoV-2 infection-associated inflammation and immune signaling [4]. The findings of our study also corroborate the aforementioned findings.

Efforts have been made to investigate the alteration in bronchoalveolar lavage fluid (BALF) proteome in COVID-19 patients compared with the non-COVID controls. This study's findings demonstrated that SARS-CoV-2 infection induces alteration in BALF proteome with enrichment of proteins involved in proinflammatory cytokine-mediated signaling and oxidative stress response [2]. In this study we found a significantly higher level of Superoxide dismutase 2 (Sod2) in infected 4dpi tissues than 4dpi mock-infected or 14dpi infected tissues (**Figure 3**). Superoxide dismutase (SOD) is vital for human health and upregulation of SOD2 expression upon challenge with human pathogens suggests its role in immune response. Antioxidant enzymes such as SOD2 are pivotal to protect from free superoxide anion, which can damage epithelial cells and impair their function. Thus enhanced expression of Sod2 in SARS-CoV-2 infected hamster lungs, suggests the upregulation of antioxidant response to prevent the oxidative stress-induced tissue damage, lung injury and respiratory distress.

The present study has some limitations of the small sample size considered for the analysis. This demands similar extensive research in a large number of animals for further validation. Taken together, the current study highlights the proteomic alterations caused owing to SARS-CoV-2 infection during the course of infection and provides an insight on the molecular pathogenesis and possible therapeutic targets of COVID-19. Importantly, the current study provides strong molecular evidence that shows the similarities between SARS-CoV-2 infection in human and hamster and advocates the clinical relevance of this model in COVID-19 research.

## Acknowledgements

The study is financially supported by DBT-ILS and DBT-BIRAC (BT/CS0004/CS/02/20). We sincerely acknowledge the technical supports given by Madan Mohan Mallick, Sushanta Kumar Swain and Bhabani S. Sahoo, ILS. Voddu Suresh is a recipient of Council of Scientific and Industrial Research (CSIR) students’ research fellowship, Government of India. Varshasnata Mohanty is a recipient of the Women Scientist-A award from the Department of Science and Technology (DST), Government of India. The graphical abstract has been created with BioRender.com.

## Supplementary figures

**Supplementary Figure 1: Histopathological changes in Lungs.** Histopathological analysis of 4dpi infected lung tissues showed various pathological features associated with lung tissue damage and inflammation (A, B, C, D, and E: scale bar = 50 μm; F: scale bar = 200 μm).

**Supplementary Figure 2: Histopathological changes in the trachea.** Histopathological analysis of tracheal tissues demonstrated desquamation of epithelial cells, mononuclear cell infiltration into epithelium and lamina propria at 4dpi (marked with # sign). In 4dpi infected tracheal tissues, presence of cell debris along with mononuclear cells also noticed in the lumen (marked with empty arrow sign). No visible epithelial damage was noticed in mock-infected and 14dpi infected tissues (marked with star sign) (scale bar = 100 μm).

**Supplementary Figure 3: Differential expression of proteins at infected 4dpi and 14dpi**. Comparison of the proteomic data between the two groups [Infected (4dpi) and infected (14dpi)]. A schematic diagram reflecting the results of the proteomics analysis of the altered proteome observed owing to SARS-CoV-2-mediated lung infection.

**Supplementary Figure 4: Differential expression of proteins in the SARS-CoV-2 mediated pulmonary damage.**

(**A**) Heatmap of the differential expressed proteins (p ≤ 0.05; fold change cut off of 1.3) were plotted protein-wise scaled representing the proteomic alterations across the infected samples (4 and 14dpi) compared to mock-infected (4dpi).

Volcano plots of the significantly altered proteins in (**B**) Infected (4dpi) vs. Mock-Infected (4dpi); (**C**) Infected (14dpi) vs. Mock-Infected (4dpi); (**D**) Infected (14dpi) vs. Infected (4dpi); The red and blue dots represent significantly altered proteins (p-value ≤ 0.05);

(**E**) Venn diagram showing the overlap of the differentially expressed proteins identified in the current study with the proteins termed as ‘secretory proteins’, ‘Secreted in the blood’, ‘Lung enriched’, ‘Lung proteome’ and ‘Group enriched’ as annotated in Human Protein Atlas (http://www.proteinatlas.org/).

## Supplementary Tables

**Supplementary Table 1:** List of differentially expressed proteins identified across the 4dpi infected and 14dpi infected

**Supplementary Table 2:** Top GO-Biological process of proteins significantly upregulated and downregulated in infected (4dpi) lung tissues as compared to mock-infected (4dpi) using Enrichr.

**Supplementary Table 3:** Top GO-Biological process of proteins significantly upregulated and downregulated in infected (14dpi) lung tissues as compared to mock-infected (4dpi) using Enrichr.

