## Supplementary figures and images for "Quantitative proteomics of hamster lung tissues infected with SARS-CoV-2 reveal host-factors having implication in the disease pathogenesis and severity"

### Supplementary figure 1

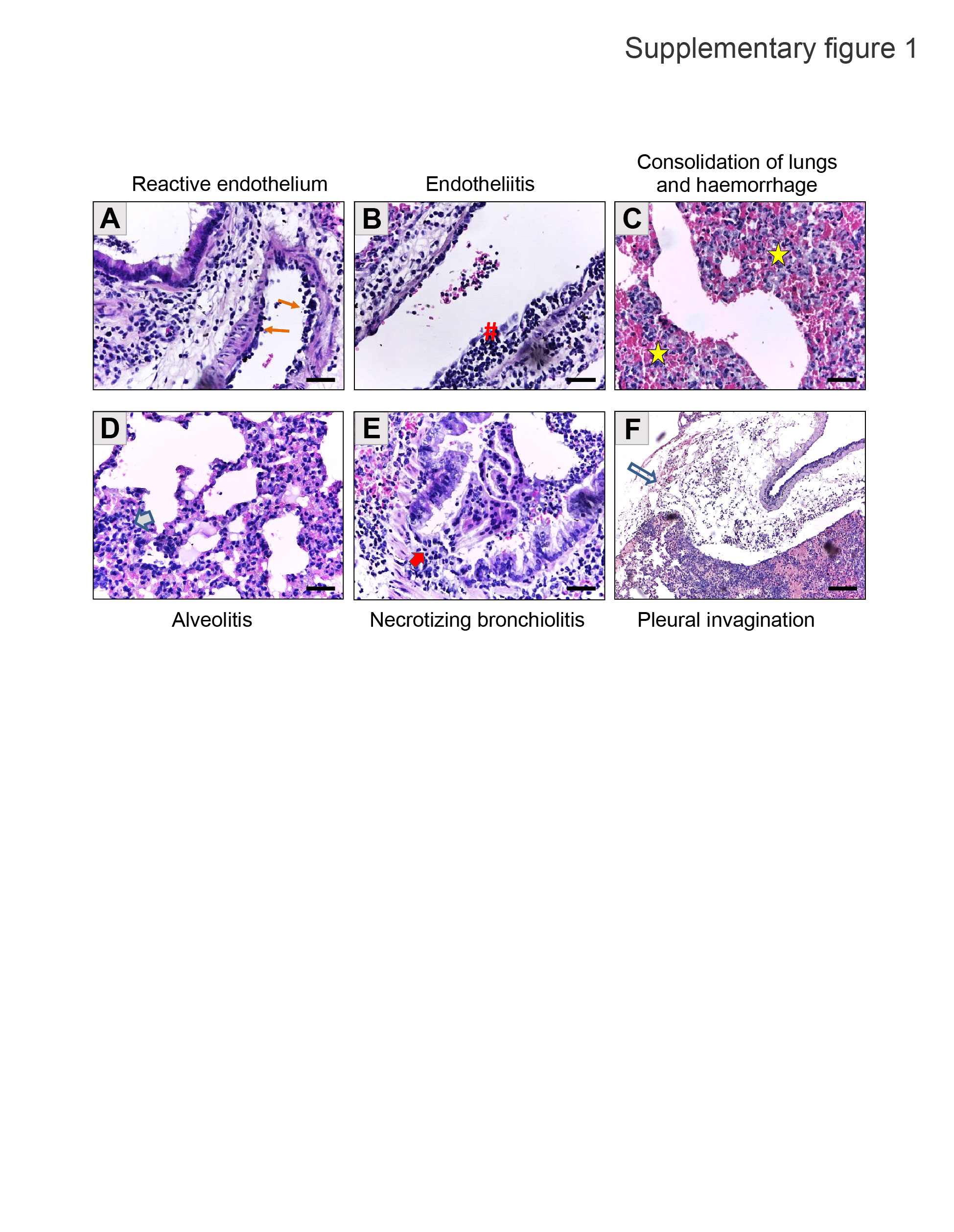

### Supplementary figure 2

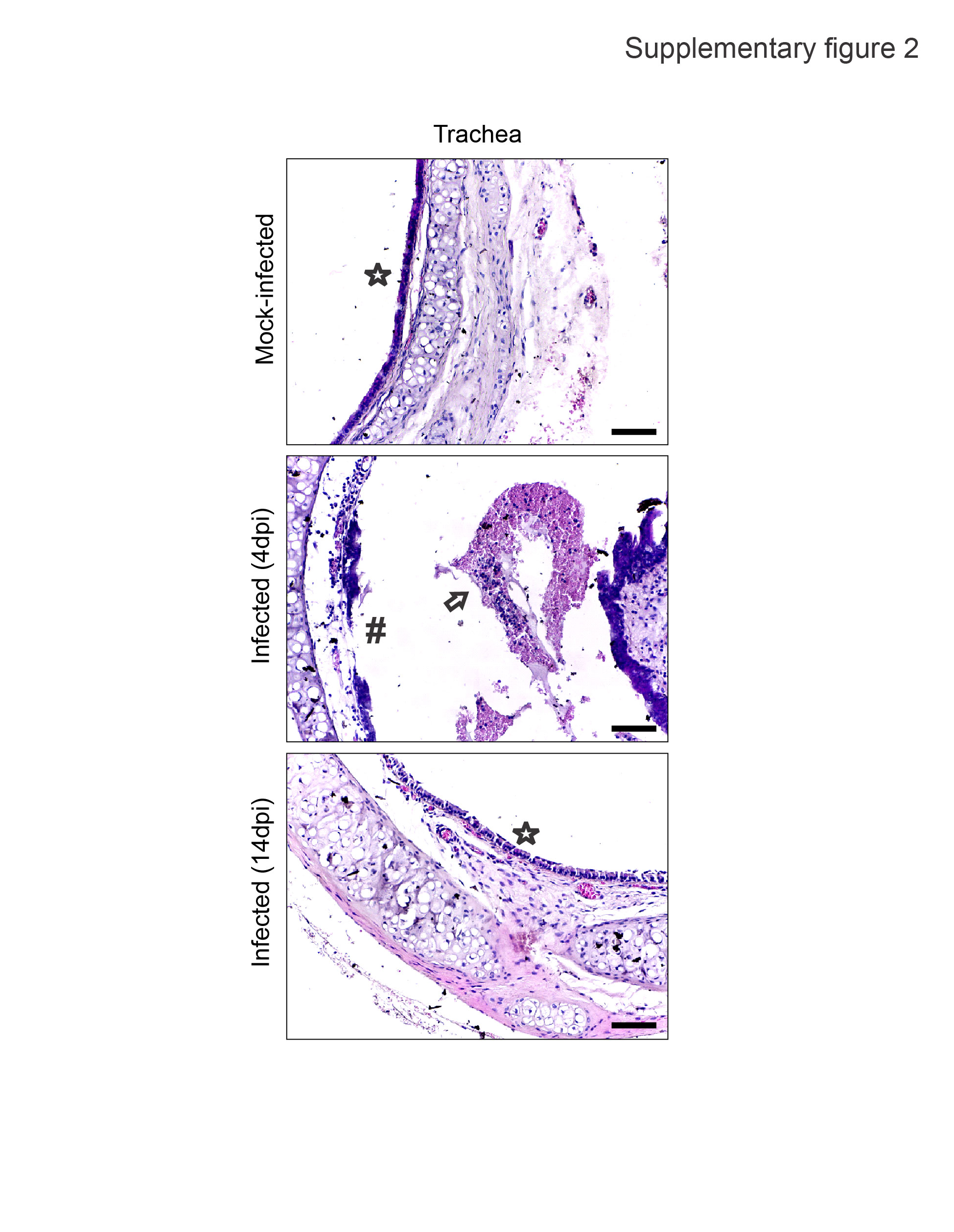

### Supplementary figure 3

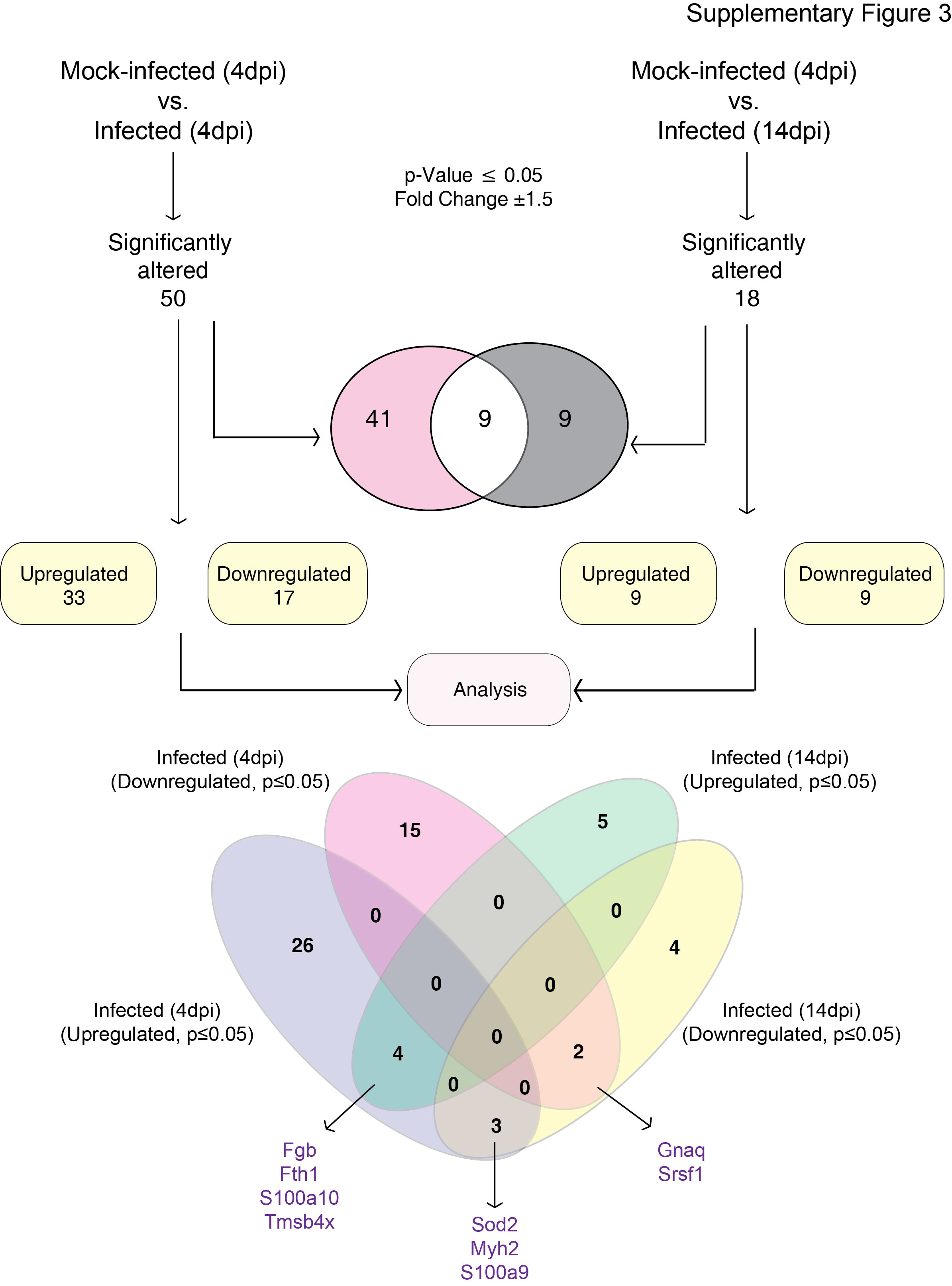

### Supplementary figure 4

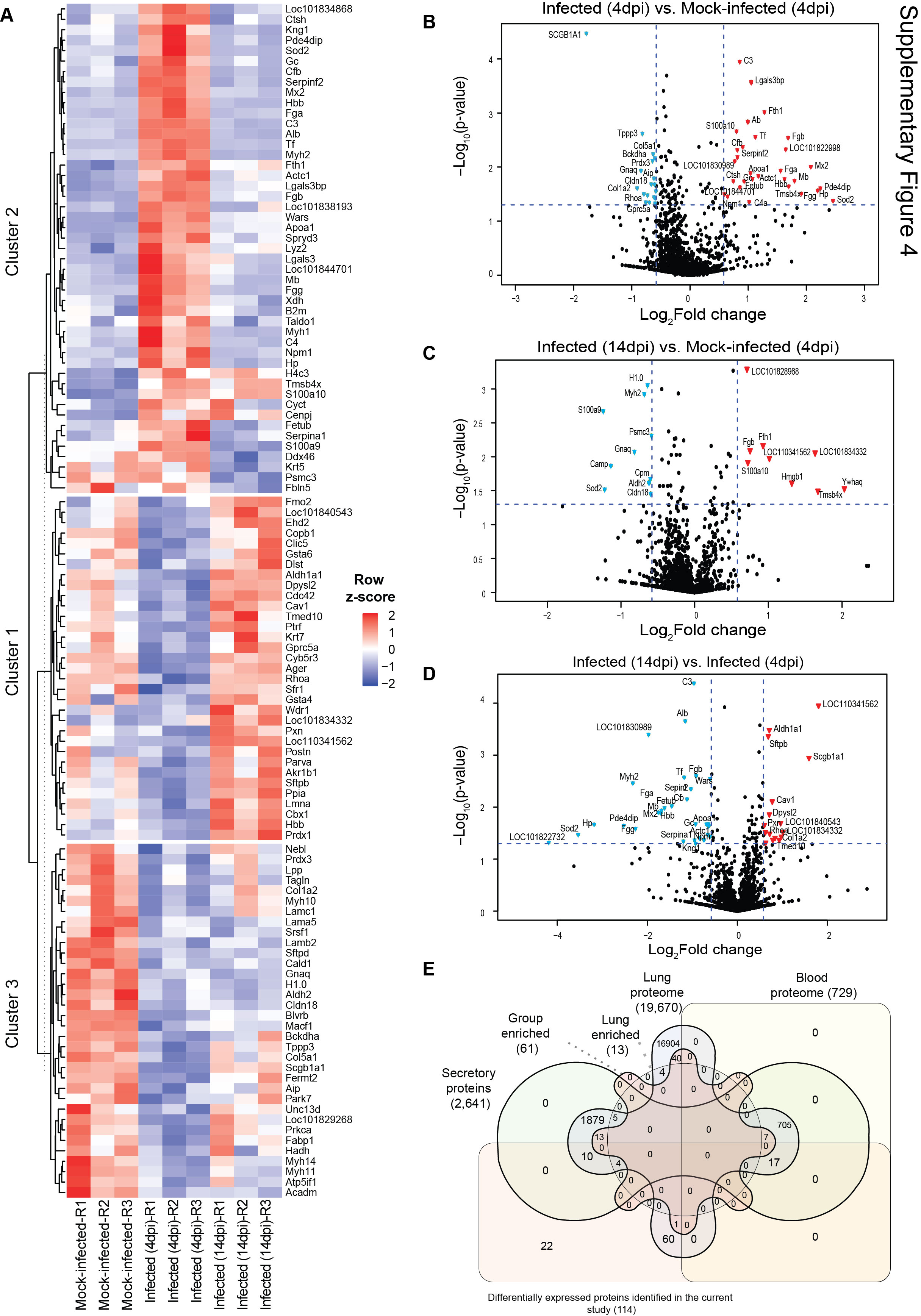
