## Supplementary table 1 for "Quantitative proteomics of hamster lung tissues infected with SARS-CoV-2 reveal host-factors having implication in the disease pathogenesis and severity"

**Supplementary Table 1: List of differentially expressed proteins identified across the 4dpi infected and 14dpi infected**

| Accession | Description | Sum PEP Score | Coverage [%] | # Peptides | # PSMs | # Unique Peptides | # AAs | MW [kDa] | calc. pI | Score Sequest HT: Sequest HT | # Peptides (by Search Engine): Sequest HT | Mock-infected (4dpi) Replicate_R1 | Mock-infected (4dpi) Replicate_R2 | Mock-infected (4dpi) Replicate_R3 | SARS-CoV-2-infected (4dpi) Replicate_R1 | SARS-CoV-2-infected (4dpi) Replicate_R2 | SARS-CoV-2-infected (4dpi) Replicate_R3 | SARS-CoV-2-(14dpi) Replicate_R1 | SARS-CoV-2-infected (14dpi) Replicate_R2 | SARS-CoV-2-infected (14dpi) Replicate_R3 | Abundance Ratio: SARS-CoV-2-infected (4dpi) / Mock-infected (4dpi) | P-Value: SARS-CoV-2-infected (4dpi) / Mock-infected (4dpi) | Abundance Ratio: SARS-CoV-2-infected (14dpi) / Mock-infected (4dpi) | P-Value: SARS-CoV-2-infected (14dpi) / Mock-infected (4dpi) | Abundance Ratio: SARS-CoV-2-infected (14dpi) / SARS-CoV-2-infected (4dpi) | P-Value: SARS-CoV-2-infected (14dpi) / SARS-CoV-2-infected (4dpi) |
|---|---|---|---|---|---|---|---|---|---|---|---|---|---|---|---|---|---|---|---|---|---|---|---|---|---|---|
| A0A1U7QTM9 | Acetoacetyl-CoA synthetase OS=Mesocricetus auratus OX=10036 GN=Aacs PE=3 SV=1 | 4.709 | 3 | 2 | 6 | 2 | 672 | 75 | 6.46 | 11.91 | 2 | 0 | 49.7 | 55.1 | 0 | 39.8 | 42.3 | 0 | 37.2 | 45.6 | 0.784 | 0.751904356 | 0.790076336 | 0.761026353 | 1.008526188 | 0.991070968 |
| Q8CFX8 | Alanine--tRNA ligase, cytoplasmic OS=Mesocricetus auratus OX=10036 GN=Aars1 PE=2 SV=1 | 12.251 | 4 | 4 | 7 | 4 | 968 | 106.6 | 5.53 | 19.43 | 4 | 31.2 | 78.5 | 0 | 27.8 | 96.8 | 0 | 33 | 62.9 | 0 | 1.049 | 0.899276696 | 0.87420237 | 0.882687306 | 0.769662921 | 0.794976864 |
| A0A1U7QPC5 | Lysine ketoglutarate reductase OS=Mesocricetus auratus OX=10036 GN=Aass PE=3 SV=1 | 7.88 | 4 | 3 | 9 | 3 | 926 | 103 | 6.93 | 19.68 | 3 | 687 | 301.3 | 279.8 | 439 | 190.3 | 187.4 | 654 | 309.7 | 268.4 | 0.639 | 0.399759421 | 0.971611072 | 0.950098252 | 1.508632301 | 0.408865367 |
| A0A1U7R5S4 | ATP-binding cassette sub-family B member 7, mitochondrial OS=Mesocricetus auratus OX=10036 GN=Abcb7 PE=4 SV=1 | 3.48 | 1 | 1 | 6 | 1 | 752 | 82.5 | 9.42 | 17.38 | 1 | 129.2 | 0 | 113.4 | 108.1 | 0 | 90.3 | 128.1 | 0 | 96.5 | 0.816 | 0.794092718 | 0.925803792 | 0.919901462 | 1.132056452 | 0.872581334 |
| A0A1U7QWW5 | abl interactor 1 isoform X3 OS=Mesocricetus auratus OX=10036 GN=Abi1 PE=3 SV=1 | 6.933 | 3 | 1 | 4 | 1 | 476 | 51.7 | 7.06 | 12.41 | 1 | 89.7 | 0 | 43.7 | 66.1 | 0 | 31.4 | 99.4 | 0 | 50.7 | 0.727 | 0.730352535 | 1.125187406 | 0.892523796 | 1.539487179 | 0.641475443 |
| A0A1U7QMD8 | Acetyl-CoA carboxylase 1 OS=Mesocricetus auratus OX=10036 GN=Acaca PE=4 SV=1 | 3.826 | 1 | 2 | 2 | 2 | 2346 | 265.2 | 6.44 | 5.26 | 2 | 406.5 | 0 | 0 | 0 | 0 | 0 | 0 | 0 | 0 | 0.01 | 0.422649731 | #DIV/0! | 0.422649731 | #DIV/0! | #DIV/0! |
| A0A1U7QEH5 | Long-chain specific acyl-CoA dehydrogenase, mitochondrial OS=Mesocricetus auratus OX=10036 GN=Acadl PE=3 SV=1 | 8.27 | 7 | 3 | 5 | 3 | 430 | 47.9 | 7.78 | 11.4 | 3 | 0 | 236.9 | 0 | 0 | 267.3 | 0 | 0 | 205.3 | 0 | 1.128 | 0.936317257 | 0.866610384 | 0.924649906 | 0.768050879 | 0.863563227 |
| A0A1U8CGI4 | Medium-chain specific acyl-CoA dehydrogenase, mitochondrial OS=Mesocricetus auratus OX=10036 GN=Acadm PE=3 SV=1 | 8.518 | 5 | 3 | 12 | 3 | 421 | 46.5 | 8.68 | 30.93 | 3 | 674.9 | 576 | 605.2 | 500.2 | 440.9 | 491.8 | 488 | 496.3 | 454.2 | 0.765 | 0.021381173 | 0.775012122 | 0.026982636 | 1.003908158 | 0.938489576 |
| A0A1U7QA30 | very long-chain specific acyl-CoA dehydrogenase, mitochondrial OS=Mesocricetus auratus OX=10036 GN=Acadvl PE=3 SV=1 | 29.414 | 16 | 7 | 23 | 7 | 655 | 70.7 | 8.9 | 81.05 | 7 | 265.1 | 127.8 | 202.8 | 206.1 | 101.5 | 171.1 | 248.3 | 126.5 | 172.4 | 0.844 | 0.483170669 | 0.918583179 | 0.776788689 | 1.143095885 | 0.652810367 |
| A0A1U7QN51 | acetyl-CoA acetyltransferase, mitochondrial OS=Mesocricetus auratus OX=10036 GN=Acat1 PE=3 SV=1 | 17.609 | 15 | 6 | 21 | 6 | 424 | 44.6 | 8.65 | 40.81 | 6 | 171.4 | 246.1 | 155.8 | 141.5 | 201.9 | 135.6 | 153.4 | 219.9 | 159.2 | 0.826 | 0.423338015 | 0.928833072 | 0.719098653 | 1.111691023 | 0.584418965 |
| A0A1U7RF50 | ATP-citrate synthase OS=Mesocricetus auratus OX=10036 GN=Acly PE=3 SV=1 | 16.386 | 8 | 6 | 13 | 6 | 1101 | 120.8 | 7.55 | 34.47 | 6 | 85.1 | 85.3 | 0 | 45.3 | 57.2 | 0 | 93.1 | 55.2 | 0 | 0.598 | 0.541339767 | 0.870305164 | 0.860128943 | 1.446829268 | 0.663705025 |
| A0A1U7QQT0 | Citrate hydro-lyase OS=Mesocricetus auratus OX=10036 GN=Aco1 PE=3 SV=1 | 14.898 | 8 | 5 | 20 | 5 | 889 | 98.2 | 7.36 | 51.03 | 5 | 213.9 | 169 | 95.7 | 184.8 | 152.5 | 97 | 214.9 | 178.1 | 98.1 | 0.864 | 0.749605893 | 1.026117844 | 0.935978765 | 1.130785172 | 0.684005438 |
| A0A1U8BGS0 | Aconitate hydratase, mitochondrial OS=Mesocricetus auratus OX=10036 GN=Aco2 PE=3 SV=1 | 38.56 | 15 | 9 | 37 | 9 | 777 | 85 | 7.83 | 118.29 | 9 | 303 | 298.7 | 425.1 | 228.8 | 253.2 | 353.3 | 302.5 | 306.7 | 370.7 | 0.831 | 0.320504416 | 0.954324114 | 0.760703941 | 1.173111457 | 0.348893892 |
| A0A1U7Q7H6 | long-chain-fatty-acid--CoA ligase 1 isoform X2 OS=Mesocricetus auratus OX=10036 GN=Acsl1 PE=4 SV=1 | 5.264 | 2 | 1 | 8 | 1 | 698 | 77.8 | 8 | 24.31 | 1 | 101.8 | 162 | 130.5 | 105.3 | 165 | 122 | 67.7 | 119.8 | 93.6 | 1.004 | 0.979897878 | 0.712908953 | 0.177514479 | 0.716543462 | 0.188647627 |
| A0A1U7QF48 | Acetyl-coenzyme A synthetase OS=Mesocricetus auratus OX=10036 GN=Acss1 PE=3 SV=1 | 4.019 | 2 | 1 | 3 | 1 | 682 | 74.6 | 6.98 | 11.15 | 1 | 53.5 | 44.9 | 60.9 | 56.3 | 37.4 | 61.9 | 33.1 | 35.2 | 46.5 | 1.016 | 0.895788797 | 0.720652856 | 0.076303776 | 0.737789203 | 0.203682945 |
| A0A1U7Q273 | actin, cytoplasmic 1 OS=Mesocricetus auratus OX=10036 GN=Actb PE=3 SV=1 | 250.873 | 78 | 27 | 501 | 7 | 375 | 41.7 | 5.48 | 1755.34 | 27 | 11241.8 | 11229.9 | 12130 | 9677.2 | 11664.5 | 9349.2 | 11512.1 | 14412.1 | 11460.3 | 0.861 | 0.205719658 | 1.080423794 | 0.445905856 | 1.218097221 | 0.146036013 |
| Q38G72 | Beta-actin (Fragment) OS=Mesocricetus auratus OX=10036 PE=2 SV=1 | 118.615 | 64 | 12 | 289 | 2 | 169 | 19.2 | 5.78 | 773.94 | 12 | 1407.3 | 1898.9 | 1137.2 | 665 | 913.2 | 506.4 | 970.1 | 1372.8 | 1892.4 | 0.481 | 0.051656411 | 0.953166494 | 0.851944654 | 2.031708721 | 0.098778749 |
| A0A1U8C9T4 | actin, alpha cardiac muscle 1 OS=Mesocricetus auratus OX=10036 GN=Actc1 PE=3 SV=1 | 181.703 | 60 | 23 | 472 | 7 | 377 | 42 | 5.39 | 1503.17 | 23 | 607.2 | 718.4 | 800.4 | 1328.7 | 1676.6 | 1805.6 | 876.4 | 1170.6 | 1004.6 | 2.256 | 0.01461249 | 1.43537159 | 0.046986272 | 0.634309589 | 0.03358798 |
| A0A1U7QYX8 | 1,2-dihydroxy-3-keto-5-methylthiopentene dioxygenase OS=Mesocricetus auratus OX=10036 GN=Adi1 PE=3 SV=1 | 6.209 | 17 | 2 | 4 | 2 | 179 | 21.6 | 5.38 | 13.21 | 2 | 51.7 | 56.6 | 0 | 71.1 | 47.3 | 0 | 41 | 49.5 | 0 | 1.072 | 0.909067346 | 0.835641736 | 0.814920917 | 0.764358108 | 0.739142318 |
| C9DFG2 | Adiponectin OS=Mesocricetus auratus OX=10036 GN=Adipoq PE=2 SV=1 | 7.198 | 6 | 1 | 7 | 1 | 244 | 26.3 | 5.4 | 29.39 | 1 | 89.4 | 112.9 | 56.2 | 97.4 | 147.6 | 42.2 | 72.9 | 116 | 61.6 | 0.84 | 0.799695106 | 0.969052224 | 0.914579298 | 0.872214485 | 0.746817823 |
| A0A1U7RBA2 | Adenosine kinase OS=Mesocricetus auratus OX=10036 GN=Adk PE=3 SV=1 | 2.407 | 2 | 1 | 4 | 1 | 361 | 40.2 | 6.62 | 0 | 1 | 64.3 | 0 | 27.4 | 76.3 | 0 | 28.5 | 59.1 | 0 | 34.3 | 1.11 | 0.887915359 | 1.018538713 | 0.983217419 | 0.891221374 | 0.89933178 |
| A0A1U8CP48 | advanced glycosylation end product-specific receptor OS=Mesocricetus auratus OX=10036 GN=Ager PE=4 SV=1 | 61.322 | 33 | 11 | 145 | 11 | 409 | 43 | 6.37 | 439.5 | 11 | 3931.8 | 4178.3 | 4332.1 | 2857.3 | 3061.2 | 3191.1 | 4069.5 | 4263.3 | 4392.3 | 0.737 | 0.002096405 | 1.022737137 | 0.564256505 | 1.396888996 | 0.000876617 |
| A0A1U8BQ65 | 1-acyl-sn-glycerol-3-phosphate acyltransferase gamma OS=Mesocricetus auratus OX=10036 GN=Agpat3 PE=3 SV=1 | 8.055 | 9 | 2 | 3 | 2 | 376 | 43.2 | 8.51 | 9.15 | 2 | 0 | 108.6 | 57.5 | 0 | 75.1 | 41.8 | 0 | 100.5 | 57.8 | 0.709 | 0.692059398 | 0.953040337 | 0.954491355 | 1.354148845 | 0.724858621 |
| A0A1U8C7M0 | Agrin OS=Mesocricetus auratus OX=10036 GN=Agrn PE=4 SV=1 | 104.033 | 16 | 18 | 85 | 18 | 2055 | 219.1 | 5.8 | 366.85 | 18 | 433.1 | 437.1 | 836.7 | 433.8 | 401.1 | 788.9 | 369.1 | 334.8 | 756.5 | 0.918 | 0.886805757 | 0.85558615 | 0.688122925 | 0.899371844 | 0.78157297 |
| A0A1U7RAJ1 | Adenosylhomocysteinase OS=Mesocricetus auratus OX=10036 GN=Ahcy PE=3 SV=1 | 54.085 | 20 | 9 | 66 | 8 | 432 | 47.6 | 6.29 | 203.91 | 9 | 2428.9 | 2354.6 | 1852.6 | 1846.8 | 1766.5 | 1382.6 | 2207.2 | 2303.9 | 1902.2 | 0.75 | 0.080353977 | 0.966426064 | 0.75256291 | 1.283712644 | 0.067120501 |
| A0A1U8C582 | Adenosylhomocysteinase OS=Mesocricetus auratus OX=10036 GN=Ahcyl1 PE=3 SV=1 | 15.781 | 8 | 4 | 17 | 3 | 529 | 58.8 | 6.89 | 31.78 | 4 | 362.9 | 271.7 | 274.6 | 301.8 | 299.6 | 342.1 | 368.2 | 285.6 | 304.1 | 0.98 | 0.752992915 | 1.053563572 | 0.699327413 | 1.015262321 | 0.876964065 |
| A0A1U8BVC5 | Alpha-2-HS-glycoprotein OS=Mesocricetus auratus OX=10036 GN=Ahsg PE=4 SV=1 | 22.894 | 13 | 3 | 20 | 3 | 375 | 39.8 | 6.15 | 78.06 | 3 | 46.8 | 0 | 0 | 72.1 | 0 | 0 | 34.8 | 0 | 0 | 1.542 | 0.785430769 | 0.743589744 | 0.8478147 | 0.482662968 | 0.674203576 |
| A0A1U7QBI1 | AH receptor-interacting protein OS=Mesocricetus auratus OX=10036 GN=Aip PE=4 SV=1 | 9.232 | 7 | 2 | 9 | 2 | 330 | 37.6 | 6.4 | 20.07 | 2 | 438.1 | 389 | 474.3 | 275.7 | 227 | 329 | 353.1 | 305.3 | 411.5 | 0.651 | 0.01613692 | 0.822114646 | 0.125128804 | 1.286401347 | 0.135580368 |
| A0A1U7Q5Z8 | GTP:AMP phosphotransferase AK3, mitochondrial OS=Mesocricetus auratus OX=10036 GN=Ak3 PE=3 SV=1 | 16.245 | 22 | 5 | 26 | 5 | 227 | 25.4 | 8.84 | 94.17 | 5 | 1047.2 | 711.1 | 1013.1 | 838.9 | 509.1 | 791.6 | 998.5 | 767.1 | 1072.4 | 0.781 | 0.228826458 | 1.024031176 | 0.882629954 | 1.326416153 | 0.167904504 |
| A0A1U8BDX2 | A-kinase anchor protein 12 isoform X2 OS=Mesocricetus auratus OX=10036 GN=Akap12 PE=4 SV=1 | 41.962 | 8 | 7 | 32 | 7 | 1622 | 173.1 | 4.37 | 83.69 | 7 | 242.4 | 40.5 | 265.5 | 181.7 | 27.6 | 244.6 | 209 | 32.5 | 246 | 0.735 | 0.760010702 | 0.888949672 | 0.844819645 | 1.074025116 | 0.909140082 |
| A0A1U8C5D0 | A-kinase anchor protein 5 OS=Mesocricetus auratus OX=10036 GN=Akap5 PE=4 SV=1 | 15.333 | 17 | 5 | 10 | 5 | 506 | 54.8 | 5.02 | 22.58 | 5 | 35.2 | 50.5 | 45 | 30.8 | 38.2 | 40.3 | 25.1 | 30.3 | 33.3 | 0.868 | 0.262324085 | 0.678653405 | 0.068717047 | 0.811527905 | 0.143176791 |
| A0A1U7R9F1 | aldose reductase OS=Mesocricetus auratus OX=10036 GN=Akr1b1 PE=3 SV=1 | 36.985 | 28 | 9 | 52 | 9 | 316 | 35.7 | 7.43 | 143.33 | 9 | 1418.1 | 1228.8 | 1375.7 | 1032.9 | 1062.7 | 1119.5 | 1477.9 | 1254.4 | 1575.4 | 0.749 | 0.027572602 | 1.070874559 | 0.44974615 | 1.339833909 | 0.053497939 |
| A0A1U7R7H9 | aflatoxin B1 aldehyde reductase member 2 OS=Mesocricetus auratus OX=10036 GN=LOC101835797 PE=4 SV=1 | 8.355 | 9 | 2 | 7 | 2 | 367 | 40.5 | 8.28 | 14.95 | 2 | 160 | 62.6 | 249.8 | 119 | 52.2 | 198.7 | 174.8 | 57.5 | 260.9 | 0.796 | 0.646306011 | 1.044030483 | 0.935113877 | 1.333333333 | 0.604363159 |
| A0A1U7Q2I1 | serum albumin precursor OS=Mesocricetus auratus OX=10036 GN=Alb PE=4 SV=1 | 666.653 | 82 | 75 | 1332 | 71 | 608 | 68.2 | 6.35 | 4634.85 | 75 | 22547.4 | 22969.3 | 19483.6 | 43481.5 | 42458.1 | 42712.5 | 20657.5 | 18918.2 | 18085.3 | 1.986 | 0.001436692 | 0.887088213 | 0.149640505 | 0.448193228 | 0.000221697 |
| A0A1U8C9T1 | Activated leukocyte cell adhesion molecule OS=Mesocricetus auratus OX=10036 GN=Alcam PE=4 SV=1 | 23.049 | 14 | 7 | 13 | 7 | 570 | 63.7 | 7.3 | 31.76 | 7 | 62.9 | 0 | 85.6 | 48.8 | 0 | 57 | 46.3 | 0 | 85.5 | 0.719 | 0.674413398 | 0.887542088 | 0.883269505 | 1.245746692 | 0.791397364 |
| A0A1U7QDB4 | retinal dehydrogenase 1 OS=Mesocricetus auratus OX=10036 GN=LOC101822890 PE=3 SV=1 | 217.166 | 48 | 29 | 377 | 8 | 501 | 54.6 | 7.58 | 1303.3 | 29 | 10225.6 | 9769.7 | 8584.4 | 7338.1 | 6847.4 | 6446.2 | 11755.6 | 10805.6 | 11176.3 | 0.701 | 0.016875435 | 1.180470754 | 0.051383278 | 1.635226375 | 0.000329003 |
| A0A1U7QVC8 | retinal dehydrogenase 2 OS=Mesocricetus auratus OX=10036 GN=Aldh1a2 PE=3 SV=1 | 14.642 | 5 | 3 | 26 | 1 | 518 | 56.6 | 5.74 | 81.43 | 3 | 39.2 | 0 | 0 | 34.7 | 0 | 0 | 27.8 | 0 | 0 | 0.887 | 0.935689376 | 0.709183673 | 0.825346258 | 0.801152738 | 0.884529829 |
| A0A1U7QWP4 | aldehyde dehydrogenase, mitochondrial OS=Mesocricetus auratus OX=10036 GN=Aldh2 PE=3 SV=1 | 85.006 | 27 | 16 | 137 | 14 | 519 | 56.4 | 7.05 | 415.8 | 16 | 3464.9 | 3216.4 | 4044.5 | 2538.1 | 2171.5 | 2740.4 | 2386.5 | 2138.3 | 2463.7 | 0.733 | 0.026424231 | 0.65155979 | 0.024233817 | 0.938053691 | 0.480270957 |
| A0A1U7RFZ4 | methylmalonate-semialdehyde dehydrogenase [acylating], mitochondrial OS=Mesocricetus auratus OX=10036 GN=Aldh6a1 PE=3 SV=1 | 32.164 | 12 | 7 | 40 | 7 | 534 | 57.8 | 8.24 | 125.31 | 7 | 368.6 | 263.1 | 446.7 | 274.1 | 216.2 | 364.9 | 294 | 237 | 447.6 | 0.753 | 0.341305326 | 0.90745549 | 0.707514696 | 1.144293732 | 0.622030779 |
| A0A1U7QLV2 | AMP deaminase OS=Mesocricetus auratus OX=10036 GN=Ampd2 PE=3 SV=1 | 2.363 | 1 | 1 | 3 | 1 | 798 | 91.8 | 6.3 | 2.73 | 1 | 152.4 | 150.7 | 101.8 | 120.6 | 112.3 | 76.4 | 126.6 | 106.2 | 94 | 0.75 | 0.21393119 | 0.807112867 | 0.261678754 | 1.056579373 | 0.744442663 |
| A0A1U7QGS2 | Anoctamin OS=Mesocricetus auratus OX=10036 GN=Ano6 PE=3 SV=1 | 12.729 | 5 | 2 | 5 | 2 | 911 | 106.2 | 6.55 | 16.76 | 2 | 64 | 0 | 0 | 53.5 | 0 | 0 | 63.5 | 0 | 0 | 0.837 | 0.906085435 | 0.9921875 | 0.995840623 | 1.186915888 | 0.910105335 |
| A0A1U8C3B7 | acidic leucine-rich nuclear phosphoprotein 32 family member A isoform X1 OS=Mesocricetus auratus OX=10036 GN=Anp32a PE=4 SV=1 | 37.487 | 38 | 9 | 52 | 7 | 266 | 30.5 | 4.2 | 106.66 | 9 | 1100.8 | 675.9 | 563.7 | 1456.6 | 711.9 | 841.2 | 1330.6 | 634.6 | 972.3 | 1.323 | 0.477567384 | 1.255127329 | 0.486882969 | 0.976010898 | 0.941002271 |
| A0A1U7QTG8 | acidic leucine-rich nuclear phosphoprotein 32 family member B OS=Mesocricetus auratus OX=10036 GN=Anp32b PE=4 SV=1 | 11.961 | 16 | 4 | 19 | 2 | 272 | 30.8 | 4.01 | 31 | 4 | 289.4 | 330.3 | 377.3 | 318.4 | 386.4 | 428.9 | 344.1 | 418.4 | 446.3 | 1.137 | 0.331745125 | 1.212437312 | 0.152225316 | 1.066243274 | 0.602578278 |
| A0A1U7Q618 | Annexin OS=Mesocricetus auratus OX=10036 GN=Anxa1 PE=3 SV=1 | 116.157 | 49 | 21 | 136 | 21 | 346 | 38.8 | 7.02 | 331.89 | 21 | 3955.7 | 3977.1 | 3472 | 3886.3 | 3845.1 | 3352.6 | 3655.1 | 3910.2 | 3309 | 0.966 | 0.676378589 | 0.953484498 | 0.502051364 | 0.981080837 | 0.789098824 |
| A0A1U7QJZ2 | Annexin OS=Mesocricetus auratus OX=10036 GN=Anxa2 PE=3 SV=1 | 192.193 | 76 | 33 | 290 | 14 | 339 | 38.6 | 7.69 | 985.96 | 33 | 4418 | 5282.8 | 4098.8 | 4050.5 | 4891.2 | 3811.1 | 3858.8 | 5020.4 | 4132.3 | 0.926 | 0.50942794 | 0.942889649 | 0.625760667 | 1.020285741 | 0.866139053 |
| A0A1U7Q8Y5 | Annexin OS=Mesocricetus auratus OX=10036 GN=Anxa3 PE=3 SV=1 | 93.053 | 48 | 23 | 95 | 23 | 323 | 36.3 | 5.9 | 245.04 | 23 | 2985.5 | 2626.9 | 3558.2 | 2550.1 | 2351.8 | 2469 | 3304.2 | 4519.5 | 3639 | 0.854 | 0.152342863 | 1.249940026 | 0.172366532 | 1.555128953 | 0.060587266 |
| A0A1U8C197 | Annexin OS=Mesocricetus auratus OX=10036 GN=Anxa4 PE=3 SV=1 | 35.762 | 25 | 9 | 49 | 9 | 317 | 35.8 | 5.91 | 137.68 | 9 | 133.1 | 299.6 | 247.5 | 119.2 | 301.2 | 293.3 | 145.1 | 310.9 | 231.9 | 0.993 | 0.892089489 | 1.0113202 | 0.971961664 | 0.963850357 | 0.915909012 |
| A0A1U8BR52 | Annexin OS=Mesocricetus auratus OX=10036 GN=Anxa5 PE=3 SV=1 | 83.924 | 57 | 18 | 135 | 18 | 321 | 36 | 5.05 | 407.53 | 18 | 3570.8 | 3296.5 | 3873.3 | 2688.2 | 2847 | 3119.9 | 3016.2 | 3470.4 | 3826.7 | 0.835 | 0.032463143 | 0.960216375 | 0.649128782 | 1.191586463 | 0.127542507 |
| A0A1U7Q485 | Annexin OS=Mesocricetus auratus OX=10036 GN=Anxa6 PE=3 SV=1 | 109.053 | 39 | 23 | 157 | 23 | 667 | 75.3 | 5.5 | 507.67 | 23 | 2783.4 | 3140 | 3130.6 | 2464.5 | 2705.4 | 2823.9 | 2668.4 | 3141.4 | 3166.2 | 0.865 | 0.089592747 | 0.991385023 | 0.903414817 | 1.122870224 | 0.177218225 |
| A0A1U7R8U6 | Annexin OS=Mesocricetus auratus OX=10036 GN=Anxa7 PE=3 SV=1 | 23.677 | 11 | 6 | 33 | 6 | 485 | 52.2 | 5.31 | 82.3 | 6 | 0 | 39.7 | 56 | 0 | 42.2 | 42.3 | 0 | 43.9 | 48.9 | 0.896 | 0.872505404 | 0.96969697 | 0.968161485 | 1.098224852 | 0.901453173 |
| A0A1U7QBA5 | Amine oxidase OS=Mesocricetus auratus OX=10036 GN=LOC101827582 PE=3 SV=1 | 46.316 | 11 | 8 | 64 | 8 | 763 | 84.7 | 6.52 | 259.88 | 8 | 1192.4 | 1162.1 | 1525.7 | 1170.6 | 1151.1 | 1524.9 | 1159.1 | 1147.8 | 1464.8 | 0.982 | 0.950140442 | 0.972037524 | 0.828227926 | 0.980528259 | 0.883595582 |
| A0A1U7QT54 | AP-1 complex subunit mu-2 isoform X1 OS=Mesocricetus auratus OX=10036 GN=Ap1m2 PE=3 SV=1 | 4.083 | 5 | 1 | 1 | 1 | 425 | 48.2 | 7.18 | 5.04 | 1 | 60.9 | 0 | 0 | 41.9 | 0 | 0 | 52.5 | 0 | 0 | 0.688 | 0.81136522 | 0.862068966 | 0.92192862 | 1.252983294 | 0.882611043 |
| A0A1U7Q6J0 | AP-2 complex subunit alpha OS=Mesocricetus auratus OX=10036 GN=Ap2a2 PE=3 SV=1 | 18.015 | 5 | 5 | 19 | 3 | 938 | 103.9 | 6.93 | 53.43 | 5 | 34.9 | 83 | 60.9 | 49.6 | 73.7 | 76 | 49.5 | 71.2 | 61.6 | 1.423 | 0.700253223 | 1.019574944 | 0.94419097 | 0.914701455 | 0.620969215 |
| A0A1U8C2U8 | AP complex subunit beta OS=Mesocricetus auratus OX=10036 GN=Ap2b1 PE=3 SV=1 | 15.401 | 4 | 6 | 13 | 6 | 951 | 105.6 | 5.34 | 15.17 | 6 | 269.4 | 344.8 | 48.7 | 263.4 | 293.7 | 45.9 | 281.5 | 333.2 | 48.5 | 0.965 | 0.874238202 | 1.000452557 | 0.999398761 | 1.099834163 | 0.872570911 |
| A0A1U7Q7D6 | AP-2 complex subunit mu OS=Mesocricetus auratus OX=10036 GN=Ap2m1 PE=3 SV=1 | 8.428 | 10 | 4 | 23 | 4 | 435 | 49.6 | 9.54 | 44.2 | 4 | 1041.7 | 669.4 | 933.9 | 857 | 562.9 | 791 | 664.3 | 499.6 | 652.2 | 0.823 | 0.368303192 | 0.686616257 | 0.113580125 | 0.821430187 | 0.28737377 |
| A0A1U7QXV7 | AP-3 complex subunit delta OS=Mesocricetus auratus OX=10036 GN=Ap3d1 PE=3 SV=1 | 12.669 | 4 | 3 | 15 | 3 | 1199 | 134.9 | 7.28 | 40.33 | 3 | 236.7 | 300.2 | 336.8 | 224.1 | 280.5 | 297.2 | 217.5 | 280.9 | 281.6 | 0.947 | 0.55145926 | 0.89275495 | 0.440617481 | 0.972811175 | 0.824370746 |
| A0A1U7QAU3 | Acyl-peptide hydrolase OS=Mesocricetus auratus OX=10036 GN=Apeh PE=4 SV=1 | 14.066 | 8 | 5 | 12 | 5 | 732 | 81.6 | 5.55 | 26.72 | 5 | 464.9 | 442.8 | 379 | 381.3 | 358.8 | 290.3 | 440.1 | 358.8 | 332.8 | 0.82 | 0.085639436 | 0.8795368 | 0.282470294 | 1.098311335 | 0.471016216 |
| A0A1U7QQL9 | Adipocyte plasma membrane-associated protein OS=Mesocricetus auratus OX=10036 GN=Apmap PE=3 SV=1 | 8.627 | 10 | 3 | 16 | 3 | 415 | 46.4 | 6.6 | 40.09 | 3 | 153.1 | 345.6 | 246.2 | 107.4 | 300.4 | 191 | 149.9 | 293.9 | 235.4 | 0.869 | 0.57008119 | 0.911800242 | 0.769741472 | 1.134268537 | 0.721986106 |
| A0A1U7Q2B6 | apolipoprotein A-I precursor OS=Mesocricetus auratus OX=10036 GN=Apoa1 PE=3 SV=1 | 122.077 | 69 | 22 | 191 | 22 | 264 | 30.7 | 6.2 | 576.18 | 22 | 2492.3 | 2472.5 | 2541 | 6170.3 | 5083.7 | 5195.1 | 3281.8 | 3621.8 | 3590.6 | 2.056 | 0.012908999 | 1.398145434 | 0.009707407 | 0.637980193 | 0.021005687 |
| A0A1U7R7S5 | Apolipoprotein A-II OS=Mesocricetus auratus OX=10036 GN=Apoa2 PE=3 SV=1 | 3.094 | 9 | 1 | 3 | 1 | 101 | 11.4 | 8.69 | 0 | 1 | 455.2 | 210.2 | 0 | 632.8 | 249.1 | 0 | 519.1 | 223.1 | 0 | 1.284 | 0.767259782 | 1.115419297 | 0.904315714 | 0.841592017 | 0.854562886 |
| A0A1U7QMM4 | apolipoprotein A-IV OS=Mesocricetus auratus OX=10036 GN=Apoa4 PE=3 SV=1 | 9.907 | 5 | 1 | 11 | 1 | 387 | 44 | 5.29 | 42.29 | 1 | 42.4 | 26.6 | 69 | 46.5 | 37.2 | 87.9 | 43.2 | 28 | 53.4 | 1.274 | 0.605010485 | 0.902898551 | 0.775266985 | 0.726107226 | 0.433626886 |
| A0A1U7QK70 | Apolipoprotein C-III OS=Mesocricetus auratus OX=10036 GN=Apoc3 PE=3 SV=1 | 12.129 | 13 | 2 | 10 | 2 | 99 | 10.7 | 5.36 | 26.74 | 2 | 47.5 | 50.3 | 92 | 31.5 | 49.2 | 82.9 | 40.7 | 25.2 | 64.8 | 0.73 | 0.696745287 | 0.6886196 | 0.348067032 | 0.798899756 | 0.596260752 |
| A0A1U7QUL7 | apolipoprotein E OS=Mesocricetus auratus OX=10036 GN=Apoe PE=3 SV=1 | 6.436 | 8 | 3 | 7 | 3 | 314 | 35.9 | 5.45 | 2.05 | 3 | 61.8 | 355.8 | 168.6 | 44.1 | 275.5 | 125.5 | 58.9 | 308.2 | 170.4 | 0.744 | 0.690622533 | 0.916922552 | 0.892125927 | 1.207593799 | 0.771202927 |
| A0A1U7R837 | MICOS complex subunit OS=Mesocricetus auratus OX=10036 GN=Apool PE=3 SV=1 | 2.691 | 7 | 1 | 1 | 1 | 265 | 29.4 | 9.47 | 4.34 | 1 | 66.9 | 0 | 0 | 69.2 | 0 | 0 | 51.1 | 0 | 0 | 1.035 | 0.982081478 | 0.763826607 | 0.860858247 | 0.738439306 | 0.844473596 |
| A0A1U8CEP8 | DCC-interacting protein 13-alpha OS=Mesocricetus auratus OX=10036 GN=Appl1 PE=4 SV=1 | 6.322 | 5 | 3 | 9 | 3 | 671 | 75.1 | 5.67 | 11.37 | 3 | 0 | 103.8 | 153.9 | 0 | 111.6 | 165.4 | 0 | 78.7 | 128.8 | 1.075 | 0.927641765 | 0.805199845 | 0.790608966 | 0.749097473 | 0.726564422 |
| A0A1U7QM61 | Aquaporin-1 OS=Mesocricetus auratus OX=10036 GN=Aqp1 PE=3 SV=1 | 26.765 | 10 | 2 | 17 | 2 | 269 | 28.6 | 7.01 | 96 | 2 | 53.5 | 0 | 109.5 | 43.3 | 0 | 84.4 | 37.1 | 0 | 77.2 | 0.79 | 0.783712532 | 0.701226994 | 0.69858028 | 0.895066562 | 0.898995431 |

**Supplementary Table 1: List of differentially expressed proteins identified across the 4dpi infected and 14dpi infected**

| Accession | Description | Sum PEP Score | Coverage [%] | # Peptides | # PSMs | # Unique Peptides | # AAs | MW [kDa] | calc. pI | Score Sequest HT: Sequest HT | # Peptides (by Search Engine): Sequest HT | Mock-infected (4dpi) Replicate_R1 | Mock-infected (4dpi) Replicate_R2 | Mock-infected (4dpi) Replicate_R3 | SARS-CoV-2-infected (4dpi) Replicate_R1 | SARS-CoV-2-infected (4dpi) Replicate_R2 | SARS-CoV-2-infected (4dpi) Replicate_R3 | SARS-CoV-2-(14dpi) Replicate_R1 | SARS-CoV-2-infected (14dpi) Replicate_R2 | SARS-CoV-2-infected (14dpi) Replicate_R3 | Abundance Ratio: SARS-CoV-2-infected (4dpi) / Mock-infected (4dpi) | P-Value: SARS-CoV-2-infected (4dpi) / Mock-infected (4dpi) | Abundance Ratio: SARS-CoV-2-infected (14dpi) / Mock-infected (4dpi) | P-Value: SARS-CoV-2-infected (14dpi) / Mock-infected (4dpi) | Abundance Ratio: SARS-CoV-2-infected (14dpi) / SARS-CoV-2-infected (4dpi) | P-Value: SARS-CoV-2-infected (14dpi) / SARS-CoV-2-infected (4dpi) |
|---|---|---|---|---|---|---|---|---|---|---|---|---|---|---|---|---|---|---|---|---|---|---|---|---|---|
| A0A1U7Q829 | Aquaporin-5 OS=Mesocricetus auratus OX=10036 GN=LOC101823084 PE=3 SV=1 | 9.704 | 6 | 1 | 34 | 1 | 265 | 28.2 | 8.29 | 128.78 | 1 | 236.5 | 260.4 | 203.7 | 196.4 | 198.4 | 174.7 | 244.8 | 301.5 | 219.7 | 0.831 | 0.100296138 | 1.093348558 | 0.50270647 | 1.345039508 | 0.102703041 |
| A0A1U7QJ99 | ADP-ribosylation factor 3 OS=Mesocricetus auratus OX=10036 GN=Arf3 PE=3 SV=1 | 19.921 | 40 | 5 | 23 | 5 | 181 | 20.6 | 7.43 | 56.43 | 5 | 685.6 | 1156 | 902.4 | 513 | 940.6 | 679.9 | 420 | 758.9 | 623.4 | 0.748 | 0.331865807 | 0.656814869 | 0.141730407 | 0.844762128 | 0.526799053 |
| A0A1U7R5X9 | arfaptin-1 isoform X2 OS=Mesocricetus auratus OX=10036 GN=Arfip1 PE=4 SV=1 | 1.769 | 3 | 1 | 3 | 1 | 341 | 38.5 | 6.76 | 9.88 | 1 | 120.8 | 157.3 | 137.8 | 100.7 | 136.4 | 130.7 | 123.6 | 175.2 | 164.4 | 0.867 | 0.353682558 | 1.113729262 | 0.457819437 | 1.259380098 | 0.181402216 |
| A0A1U7Q5G4 | rho GDP-dissociation inhibitor 1 OS=Mesocricetus auratus OX=10036 GN=Arhgdia PE=3 SV=1 | 88.016 | 67 | 12 | 74 | 12 | 204 | 23.4 | 5.2 | 206.98 | 12 | 1137.6 | 1040.6 | 906.7 | 958.6 | 987.7 | 730.2 | 1029.5 | 1043.6 | 877.1 | 0.843 | 0.268489685 | 0.9563357 | 0.628827426 | 1.102260415 | 0.40939669 |
| A0A1U8BXV1 | rho guanine nucleotide exchange factor 12 isoform X1 OS=Mesocricetus auratus OX=10036 GN=Arhgef12 PE=4 SV=1 | 3.829 | 1 | 1 | 5 | 1 | 1544 | 172.7 | 5.72 | 21.79 | 1 | 154.3 | 126.9 | 62.7 | 137.4 | 123.3 | 43.7 | 141.1 | 116.8 | 49.6 | 0.852 | 0.75772838 | 0.894155278 | 0.76865779 | 1.010183968 | 0.980630207 |
| A0A1U8C9H7 | Guanine nucleotide exchange factor H1 OS=Mesocricetus auratus OX=10036 GN=Arhgef2 PE=4 SV=1 | 11.67 | 6 | 4 | 7 | 4 | 971 | 109.7 | 6.98 | 19.14 | 4 | 327.2 | 171.1 | 214.8 | 243.6 | 105.9 | 152.7 | 273.8 | 140.3 | 192.4 | 0.711 | 0.318665341 | 0.85051185 | 0.589987413 | 1.207686181 | 0.568807418 |
| A0A1U8BYM3 | rho guanine nucleotide exchange factor 7 isoform X1 OS=Mesocricetus auratus OX=10036 GN=Arhgef7 PE=4 SV=1 | 12.348 | 5 | 4 | 10 | 4 | 883 | 98.8 | 6.83 | 16.8 | 4 | 243.1 | 306.1 | 422.2 | 201.7 | 256.6 | 318.1 | 192.6 | 237.6 | 369.7 | 0.83 | 0.365130275 | 0.82345069 | 0.486612789 | 1.030267903 | 0.907916628 |
| A0A1U7QNG3 | PRA1 family protein OS=Mesocricetus auratus OX=10036 GN=Arl6ip5 PE=3 SV=1 | 16.86 | 24 | 5 | 21 | 5 | 188 | 21.5 | 9.61 | 49.22 | 5 | 235.8 | 199.2 | 72.1 | 252.2 | 212.9 | 59.7 | 171.7 | 152.7 | 81 | 1.002 | 0.94260383 | 0.799447841 | 0.590872736 | 0.772484756 | 0.585221832 |
| A0A1U7QJA5 | ADP-ribosylation factor-like protein 8A OS=Mesocricetus auratus OX=10036 GN=Arl8a PE=3 SV=1 | 2.943 | 5 | 1 | 3 | 1 | 186 | 21.4 | 7.77 | 7.67 | 1 | 80.5 | 103.9 | 105.1 | 61.4 | 93.3 | 80.6 | 66.1 | 109.4 | 93.3 | 0.767 | 0.215780585 | 0.928497409 | 0.672681242 | 1.142371441 | 0.518829556 |
| A0A1U7R2Z3 | Actin-related protein 2/3 complex subunit OS=Mesocricetus auratus OX=10036 GN=Arpc1b PE=3 SV=1 | 2.781 | 3 | 1 | 2 | 1 | 372 | 41 | 8.35 | 0 | 1 | 82.9 | 60.7 | 0 | 61.8 | 35.6 | 0 | 73.2 | 74.4 | 0 | 0.661 | 0.643372423 | 1.027855153 | 0.971368941 | 1.515400411 | 0.614252481 |
| A0A1U7QBT5 | Arp2/3 complex 34 kDa subunit OS=Mesocricetus auratus OX=10036 GN=Arpc2 PE=3 SV=1 | 22.577 | 21 | 6 | 25 | 6 | 300 | 34.3 | 7.36 | 75.77 | 6 | 645.3 | 1372 | 589.1 | 786.5 | 762.4 | 810 | 718.2 | 1377.5 | 682.3 | 1.219 | 0.774729787 | 1.065837937 | 0.87413463 | 1.177667557 | 0.599692583 |
| A0A1U8CFP1 | Arpin OS=Mesocricetus auratus OX=10036 GN=Arpin PE=3 SV=1 | 3.055 | 4 | 1 | 5 | 1 | 226 | 25.3 | 5.31 | 16.01 | 1 | 101.3 | 293.2 | 290.8 | 101.9 | 246.1 | 305.4 | 74.3 | 223.9 | 238.1 | 1.05 | 0.909363478 | 0.782576974 | 0.580204301 | 0.820783594 | 0.651540533 |
| A0A1U7QFT8 | acid ceramidase OS=Mesocricetus auratus OX=10036 GN=Asah1 PE=3 SV=1 | 3.884 | 5 | 2 | 2 | 2 | 395 | 44.7 | 7.64 | 2.35 | 2 | 82.3 | 0 | 0 | 86.6 | 0 | 0 | 63.9 | 0 | 0 | 1.053 | 0.973017088 | 0.776427704 | 0.868908242 | 0.737875289 | 0.844107211 |
| A0A1U8BW09 | achaete-scute homolog 2 OS=Mesocricetus auratus OX=10036 GN=Ascl2 PE=4 SV=1 | 1.354 | 6 | 1 | 2 | 1 | 193 | 20.3 | 10.29 | 0 | 1 | 118.4 | 139.4 | 0 | 100.6 | 114.1 | 0 | 94.8 | 123.5 | 0 | 0.834 | 0.811829727 | 0.84678045 | 0.829567658 | 1.016767583 | 0.982644248 |
| A0A1U7QIR7 | ubiquitin-like-conjugating enzyme ATG3 OS=Mesocricetus auratus OX=10036 GN=Atg3 PE=4 SV=1 | 8.354 | 10 | 2 | 4 | 2 | 314 | 35.7 | 4.74 | 17.81 | 2 | 0 | 76.5 | 0 | 0 | 85.1 | 0 | 0 | 53.2 | 0 | 1.112 | 0.943738616 | 0.695424837 | 0.816271833 | 0.625146886 | 0.769364021 |
| A0A1U7Q676 | 5-aminoimidazole-4-carboxamide ribonucleotide formyltransferase OS=Mesocricetus auratus OX=10036 GN=Atic PE=3 SV=1 | 20.438 | 6 | 3 | 31 | 3 | 592 | 64.7 | 6.98 | 111.55 | 3 | 188.3 | 159.1 | 110.2 | 163 | 156.1 | 113.9 | 184.6 | 184.8 | 106.2 | 0.981 | 0.782158299 | 1.039335664 | 0.871253061 | 1.098383372 | 0.669499585 |
| A0A1U8BH98 | copper transport protein ATOX1 isoform X1 OS=Mesocricetus auratus OX=10036 GN=Atox1 PE=4 SV=1 | 2.077 | 10 | 1 | 2 | 1 | 90 | 10 | 8 | 2.29 | 1 | 0 | 130.5 | 68.5 | 0 | 115.9 | 46.3 | 0 | 115.3 | 54.1 | 0.774 | 0.820326918 | 0.851256281 | 0.854173837 | 1.044389642 | 0.962020221 |
| A0A1U7QLM8 | Sodium/potassium-transporting ATPase subunit alpha OS=Mesocricetus auratus OX=10036 GN=Atp1a1 PE=3 SV=1 | 39.685 | 13 | 10 | 52 | 6 | 1023 | 113 | 5.45 | 142.19 | 10 | 1266.8 | 1017.8 | 887.1 | 922.2 | 756.9 | 681.4 | 935.9 | 839.1 | 685.5 | 0.728 | 0.122467809 | 0.775766939 | 0.160877369 | 1.042363906 | 0.759848041 |
| F1T2M2 | ATP synthase subunit alpha OS=Mesocricetus auratus OX=10036 GN=atp5a1 PE=2 SV=1 | 75.985 | 38 | 17 | 78 | 17 | 553 | 59.7 | 9.19 | 227.23 | 17 | 1363.1 | 1797.1 | 1668.1 | 1320.8 | 1847.8 | 1583.1 | 1295 | 1748.4 | 1625.2 | 0.969 | 0.904379256 | 0.966924176 | 0.789796461 | 0.982511522 | 0.898452977 |
| A0A1U7QGG6 | ATP synthase subunit gamma OS=Mesocricetus auratus OX=10036 GN=Atp5c1 PE=3 SV=1 | 7.883 | 13 | 3 | 11 | 3 | 298 | 32.9 | 9.13 | 11.78 | 3 | 133.2 | 0 | 0 | 125.5 | 0 | 0 | 147.5 | 0 | 0 | 0.942 | 0.968462684 | 1.107357357 | 0.94612768 | 1.175298805 | 0.915163346 |
| A0A1U7QD82 | ATP synthase F(0) complex subunit B1, mitochondrial OS=Mesocricetus auratus OX=10036 GN=Atp5f1 PE=3 SV=1 | 14.036 | 20 | 5 | 20 | 5 | 256 | 28.9 | 9.09 | 48.81 | 5 | 354 | 242.2 | 422.2 | 319.3 | 188.5 | 376.1 | 308.1 | 217.6 | 360.2 | 0.793 | 0.588963927 | 0.869893951 | 0.547491403 | 1.002262699 | 0.992832712 |
| A0A1U7QK43 | ATP synthase-coupling factor 6, mitochondrial OS=Mesocricetus auratus OX=10036 GN=Atp5j PE=3 SV=1 | 13.453 | 52 | 4 | 20 | 4 | 108 | 12.5 | 9.48 | 35.52 | 4 | 508.8 | 566.8 | 584.2 | 386 | 435.3 | 469.4 | 470.2 | 558 | 632.6 | 0.783 | 0.020965154 | 1.000602482 | 0.995317267 | 1.286743627 | 0.101756993 |
| A0A1U7QCT0 | V-type proton ATPase subunit OS=Mesocricetus auratus OX=10036 GN=Atp6v0d1 PE=3 SV=1 | 2.454 | 5 | 1 | 1 | 1 | 351 | 40.3 | 4.94 | 2.49 | 1 | 37 | 0 | 0 | 41 | 0 | 0 | 25.4 | 0 | 0 | 1.111 | 0.945772433 | 0.686486486 | 0.810353074 | 0.619512195 | 0.765577238 |
| A0A1U8BYP2 | H(+)-transporting two-sector ATPase OS=Mesocricetus auratus OX=10036 GN=Atp6v1a PE=3 SV=1 | 21.115 | 10 | 4 | 29 | 4 | 617 | 68.3 | 5.58 | 85.29 | 4 | 271.8 | 414.5 | 243.3 | 195.3 | 293.3 | 174.4 | 258.5 | 455.4 | 235.6 | 0.708 | 0.247984131 | 1.021407057 | 0.943516973 | 1.432126697 | 0.311643245 |
| A0A1U7QHB6 | V-type proton ATPase subunit E 1 OS=Mesocricetus auratus OX=10036 GN=Atp6v1e1 PE=3 SV=1 | 15.107 | 6 | 2 | 18 | 2 | 226 | 26.1 | 8.43 | 73.19 | 2 | 87.4 | 110.4 | 65 | 82.3 | 84 | 52.4 | 85.5 | 107 | 56.9 | 0.761 | 0.42972425 | 0.949010654 | 0.8306225 | 1.140374943 | 0.598815855 |
| P30442 | Beta-2-microglobulin (Fragment) OS=Mesocricetus auratus OX=10036 GN=B2M PE=2 SV=1 | 13.931 | 67 | 3 | 13 | 3 | 54 | 6.4 | 6.52 | 33.22 | 3 | 465.4 | 421.2 | 300.8 | 663 | 578.4 | 494.7 | 453.7 | 392.9 | 345.6 | 1.424 | 0.057253498 | 1.004042446 | 0.979639511 | 0.686711595 | 0.043386825 |
| A0A1U8C1C4 | Hexosyltransferase OS=Mesocricetus auratus OX=10036 GN=B3gnt9 PE=3 SV=1 | 1.465 | 2 | 1 | 2 | 1 | 395 | 43.8 | 9.94 | 3.92 | 1 | 0 | 120.3 | 134.8 | 0 | 111.4 | 115.4 | 0 | 115.7 | 116.9 | 0.89 | 0.876811895 | 0.911799294 | 0.902894209 | 1.025573192 | 0.973234862 |
| A0A1U8CCP3 | BAI1-associated protein 3 OS=Mesocricetus auratus OX=10036 GN=Baiap3 PE=4 SV=1 | 1.544 | 1 | 1 | 3 | 1 | 1152 | 129.3 | 6.27 | 7.53 | 1 | 467.8 | 163.1 | 0 | 378.6 | 133.3 | 0 | 415.3 | 171.9 | 0 | 0.813 | 0.833480142 | 0.930733872 | 0.940278185 | 1.147099043 | 0.885633362 |
| A0A1U8C000 | barrier-to-autointegration factor OS=Mesocricetus auratus OX=10036 GN=Banf1 PE=4 SV=1 | 23.12 | 61 | 5 | 17 | 5 | 89 | 10.1 | 6.09 | 43.39 | 5 | 193.4 | 81.3 | 73.8 | 207.2 | 81.9 | 284.3 | 172.3 | 60.9 | 256.1 | 1.755 | 0.356452555 | 1.404017217 | 0.535503062 | 0.853331008 | 0.748806063 |
| A0A1U8CNB5 | basal cell adhesion molecule OS=Mesocricetus auratus OX=10036 GN=Bcam PE=4 SV=1 | 57.212 | 26 | 10 | 53 | 10 | 623 | 67.7 | 5.82 | 186.97 | 10 | 696 | 426.7 | 1332 | 448.3 | 311.1 | 785.4 | 609.4 | 382.2 | 1069.8 | 0.624 | 0.39022693 | 0.839776755 | 0.71777258 | 1.334412222 | 0.527623561 |
| A0A1U8CZL7 | B-cell receptor-associated protein OS=Mesocricetus auratus OX=10036 GN=Bcap31 PE=3 SV=1 | 12.08 | 20 | 5 | 20 | 5 | 245 | 27.9 | 8.43 | 47.89 | 5 | 837.2 | 707.9 | 584.9 | 860.3 | 795.6 | 656.1 | 755.6 | 639.3 | 514.5 | 1.028 | 0.557105947 | 0.896431925 | 0.506023097 | 0.825865052 | 0.220091408 |
| A0A1U8BDS0 | bcl-2-associated transcription factor 1 isoform X2 OS=Mesocricetus auratus OX=10036 GN=Bclaf1 PE=3 SV=1 | 3.605 | 2 | 1 | 3 | 1 | 918 | 105.7 | 9.98 | 11.44 | 1 | 139.7 | 60.4 | 0 | 113.3 | 46.1 | 0 | 97.9 | 54.3 | 0 | 0.786 | 0.808055848 | 0.76061969 | 0.764412956 | 0.954830615 | 0.958612203 |
| A0A1U7RGR7 | Biglycan OS=Mesocricetus auratus OX=10036 GN=Bgn PE=3 SV=1 | 35.741 | 25 | 9 | 43 | 8 | 369 | 41.6 | 7.28 | 127.54 | 9 | 1337.4 | 1186.9 | 1169.9 | 1075.3 | 1031 | 1095.2 | 1274.9 | 1320.3 | 1194.1 | 0.827 | 0.077463149 | 1.025743057 | 0.65312757 | 1.183601437 | 0.018155133 |
| A0A1U7Q434 | Bleomycin hydrolase OS=Mesocricetus auratus OX=10036 GN=Blmh PE=3 SV=1 | 3.248 | 4 | 2 | 5 | 2 | 455 | 52.4 | 6.34 | 9.65 | 2 | 506.1 | 87.8 | 100.5 | 476.1 | 107.5 | 81.6 | 437.2 | 78.8 | 104.7 | 1.224 | 0.961071205 | 0.893865207 | 0.89788348 | 0.933102826 | 0.935417819 |
| A0A1U7RG83 | flavin reductase (NADPH) OS=Mesocricetus auratus OX=10036 GN=Blvrb PE=4 SV=1 | 21.137 | 17 | 3 | 31 | 3 | 206 | 22.2 | 7.28 | 142.65 | 3 | 703.2 | 700.3 | 712.8 | 529.4 | 525.1 | 556.3 | 613.9 | 590.3 | 609.9 | 0.75 | 0.001157045 | 0.85720361 | 0.001164636 | 1.126210579 | 0.006414101 |
| A0A1U7R9X4 | bolA-like protein 1 OS=Mesocricetus auratus OX=10036 GN=Bola1 PE=3 SV=1 | 5.537 | 11 | 1 | 11 | 1 | 137 | 14.3 | 8.76 | 41.27 | 1 | 71.5 | 44 | 153.6 | 56.9 | 35.6 | 135.7 | 62.4 | 43.1 | 135.8 | 0.82 | 0.776344528 | 0.896692679 | 0.841498421 | 1.057405784 | 0.921338403 |
| A0A1U7QZF0 | Phosphoglycerate mutase OS=Mesocricetus auratus OX=10036 GN=Bpgm PE=3 SV=1 | 11.217 | 14 | 4 | 14 | 4 | 259 | 29.9 | 7.31 | 36.74 | 4 | 344.4 | 295 | 456.6 | 267.7 | 220.2 | 410.1 | 342.7 | 259.1 | 367.7 | 0.746 | 0.426794094 | 0.884580292 | 0.512316692 | 1.079621381 | 0.739959156 |
| A0A1U7Q3M5 | protein BRICK1 OS=Mesocricetus auratus OX=10036 GN=Brk1 PE=3 SV=1 | 6.887 | 23 | 2 | 18 | 2 | 75 | 8.8 | 5.45 | 64.53 | 2 | 1037.9 | 1119.1 | 962.1 | 916.4 | 952.4 | 848.4 | 792.4 | 869.5 | 826 | 0.882 | 0.079294712 | 0.797633933 | 0.026632049 | 0.915611659 | 0.119548963 |
| A0A1U7QZX0 | BRO1 domain- and CAAX motif-containing protein OS=Mesocricetus auratus OX=10036 GN=Brox PE=3 SV=1 | 6.025 | 8 | 2 | 4 | 2 | 411 | 46.1 | 7.47 | 10.96 | 2 | 0 | 63.9 | 141.6 | 0 | 41.4 | 94.5 | 0 | 73 | 111.5 | 0.657 | 0.665426302 | 0.897810219 | 0.900470085 | 1.357615894 | 0.723805276 |
| A0A1U7QXY5 | basigin isoform X3 OS=Mesocricetus auratus OX=10036 GN=Bsg PE=4 SV=1 | 9.673 | 7 | 3 | 20 | 3 | 386 | 42.2 | 6.16 | 75.77 | 3 | 116.9 | 128.6 | 161 | 90.1 | 96.9 | 136.1 | 140.8 | 132.1 | 191 | 0.771 | 0.227183526 | 1.141205412 | 0.449456977 | 1.435778397 | 0.11824978 |
| A0A1U7R7E8 | Complement C1q subcomponent subunit A OS=Mesocricetus auratus OX=10036 GN=C1qa PE=4 SV=1 | 5.012 | 11 | 2 | 4 | 2 | 245 | 25.7 | 9.14 | 6.87 | 2 | 0 | 0 | 113.8 | 0 | 0 | 141.8 | 0 | 0 | 100.9 | 1.246 | 0.885398426 | 0.886643234 | 0.936536895 | 0.711565585 | 0.826912414 |
| A0A1U7QVF6 | C4b-binding protein alpha chain OS=Mesocricetus auratus OX=10036 GN=C4bpa PE=4 SV=1 | 5.74 | 5 | 2 | 4 | 2 | 584 | 64.8 | 6.61 | 7.46 | 2 | 80.3 | 65.5 | 0 | 54 | 60.8 | 0 | 51.9 | 53.3 | 0 | 0.79 | 0.758677094 | 0.721536351 | 0.680315413 | 0.916376307 | 0.908132773 |
| A0A1U7QIM1 | Carbonic anhydrase OS=Mesocricetus auratus OX=10036 GN=Ca1 PE=3 SV=1 | 32.678 | 29 | 5 | 55 | 5 | 261 | 28.6 | 6.96 | 186.15 | 5 | 1201.8 | 1253.9 | 1157.5 | 1137.2 | 1155.3 | 1068.4 | 1058.4 | 1149.5 | 1042.4 | 0.921 | 0.094032956 | 0.899562714 | 0.05141898 | 0.967092148 | 0.437797114 |
| A0A1U7Q3W6 | Carbonic anhydrase OS=Mesocricetus auratus OX=10036 GN=Ca2 PE=3 SV=1 | 83.576 | 50 | 13 | 96 | 13 | 260 | 29 | 7.43 | 296.02 | 13 | 780.7 | 814 | 744.5 | 827.3 | 763 | 662.8 | 756.8 | 713.4 | 698 | 1.06 | 0.622980832 | 0.926898085 | 0.100793399 | 0.962318583 | 0.62413918 |
| A0A1U7QDT3 | Carbonic anhydrase OS=Mesocricetus auratus OX=10036 GN=Ca4 PE=3 SV=1 | 25.447 | 23 | 8 | 23 | 8 | 307 | 34.7 | 7.05 | 70.68 | 8 | 457.6 | 415.2 | 745.7 | 287.7 | 289.2 | 530.9 | 406 | 497.8 | 765.5 | 0.65 | 0.269237314 | 1.031387087 | 0.915394292 | 1.506860444 | 0.242464951 |
| Q4U471 | Calumenin OS=Mesocricetus auratus OX=10036 GN=CALU PE=2 SV=1 | 48.163 | 40 | 9 | 45 | 9 | 315 | 37 | 4.67 | 163.27 | 9 | 928.9 | 958.6 | 992.4 | 852.2 | 847.1 | 888.8 | 823.8 | 825.9 | 831.8 | 0.896 | 0.015505072 | 0.861661863 | 0.017214164 | 0.958811483 | 0.109325531 |
| A0A1U8C3K2 | calnexin isoform X1 OS=Mesocricetus auratus OX=10036 GN=Canx PE=3 SV=1 | 38.749 | 9 | 5 | 68 | 5 | 592 | 67.3 | 4.61 | 212.76 | 5 | 1072.5 | 1172.6 | 999.7 | 1182.8 | 1036.1 | 794.9 | 1479.5 | 1255.5 | 1047.6 | 0.795 | 0.58126199 | 1.16574211 | 0.286017367 | 1.255093238 | 0.203256419 |
| A0A1U7QR77 | Adenylyl cyclase-associated protein OS=Mesocricetus auratus OX=10036 GN=Cap1 PE=3 SV=1 | 36.354 | 31 | 10 | 38 | 10 | 474 | 51.5 | 7.5 | 120.86 | 10 | 906.8 | 701.5 | 718.8 | 883.6 | 737.6 | 707.2 | 862.7 | 677.4 | 685.7 | 1.051 | 0.99620055 | 0.956469425 | 0.724662222 | 0.955935406 | 0.695977217 |
| A0A1U7QSM1 | macrophage-capping protein isoform X2 OS=Mesocricetus auratus OX=10036 GN=Capg PE=4 SV=1 | 11.458 | 17 | 3 | 10 | 3 | 349 | 38.7 | 6.73 | 29.12 | 3 | 43 | 51.6 | 102.4 | 49.7 | 60.9 | 111.3 | 45.8 | 53.4 | 94.3 | 1.181 | 0.769804771 | 0.982233503 | 0.963471839 | 0.872014421 | 0.716581888 |
| A0A1U7QYM4 | Calcium-activated neutral proteinase small subunit OS=Mesocricetus auratus OX=10036 GN=Capns1 PE=4 SV=1 | 15.14 | 14 | 3 | 16 | 3 | 268 | 28.5 | 5.66 | 68.59 | 3 | 105.8 | 29.4 | 0 | 86.6 | 34.5 | 0 | 80.5 | 26.3 | 0 | 0.981 | 0.91313648 | 0.789940828 | 0.822987779 | 0.881915772 | 0.897022548 |
| A0A1U7QF88 | caprin-1 isoform X1 OS=Mesocricetus auratus OX=10036 GN=Caprin1 PE=3 SV=1 | 1.52 | 2 | 1 | 4 | 1 | 707 | 78.2 | 5.25 | 7.66 | 1 | 106.9 | 0 | 118.1 | 113.9 | 0 | 128.7 | 83.4 | 0 | 99.7 | 1.077 | 0.920799739 | 0.813777778 | 0.788963539 | 0.754740313 | 0.718801607 |
| A0A1U7QD74 | F-actin-capping protein subunit alpha OS=Mesocricetus auratus OX=10036 GN=Capza1 PE=3 SV=1 | 11.998 | 12 | 3 | 20 | 3 | 286 | 32.9 | 5.69 | 54.48 | 3 | 524.9 | 577.6 | 711.5 | 469.9 | 502.2 | 673.6 | 541.4 | 617.8 | 765 | 0.895 | 0.542005753 | 1.060749724 | 0.691731784 | 1.1692289 | 0.365929094 |
| A0A1U7Q360 | Catalase OS=Mesocricetus auratus OX=10036 GN=Cat PE=3 SV=1 | 82.546 | 33 | 16 | 107 | 10 | 527 | 60 | 7.52 | 340.31 | 16 | 1658.2 | 1787.2 | 1529.9 | 1530.8 | 1665.8 | 1657.5 | 1564.9 | 1728 | 1751.7 | 0.932 | 0.668977754 | 1.013928808 | 0.819858633 | 1.039245174 | 0.438278579 |
| A0A1U8BT65 | chromobox protein homolog 5 isoform X3 OS=Mesocricetus auratus OX=10036 GN=Cbx5 PE=4 SV=1 | 8.491 | 15 | 2 | 5 | 2 | 191 | 22.2 | 5.86 | 19.7 | 2 | 154.4 | 65.1 | 69.2 | 168.2 | 72.1 | 78.2 | 143.6 | 48.8 | 83.3 | 1.13 | 0.827005354 | 0.954970558 | 0.919326904 | 0.865620094 | 0.749252581 |
| A0A1U7QBV5 | coiled-coil domain-containing protein 148 isoform X1 OS=Mesocricetus auratus OX=10036 GN=Ccdc148 PE=4 SV=1 | 2.021 | 1 | 1 | 4 | 1 | 599 | 72 | 8.44 | 11.29 | 1 | 63.1 | 146.8 | 95.9 | 40.9 | 104.3 | 63.2 | 47.2 | 136.2 | 88.9 | 0.711 | 0.352671111 | 0.890451275 | 0.768304072 | 1.306621881 | 0.542009825 |
| A0A1U7QNY1 | Coiled-coil domain-containing protein 47 OS=Mesocricetus auratus OX=10036 GN=Ccdc47 PE=4 SV=1 | 3.007 | 2 | 1 | 8 | 1 | 483 | 55.8 | 4.84 | 20.55 | 1 | 689.1 | 493 | 606.7 | 497.7 | 415.3 | 488.7 | 500.5 | 398.7 | 503.4 | 0.82 | 0.137377375 | 0.784101073 | 0.13992488 | 1.000642077 | 0.994814696 |
| A0A1U7QRS4 | coiled-coil domain-containing protein 50 isoform X3 OS=Mesocricetus auratus OX=10036 GN=Ccdc50 PE=4 SV=1 | 3.854 | 5 | 1 | 5 | 1 | 305 | 35.4 | 7.03 | 19.59 | 1 | 97.6 | 42.5 | 85.7 | 82.2 | 33.8 | 76.1 | 72.8 | 49.9 | 83.8 | 0.797 | 0.645790446 | 0.914526129 | 0.761422549 | 1.074960958 | 0.806979692 |
| A0A1U8BUA2 | Superoxide dismutase [Cu-Zn] OS=Mesocricetus auratus OX=10036 GN=Ccs PE=3 SV=1 | 1.768 | 3 | 1 | 4 | 1 | 228 | 24.1 | 5.31 | 9.29 | 1 | 31.9 | 41.4 | 82.7 | 22.3 | 37.7 | 57.5 | 28.6 | 36.1 | 70.6 | 0.739 | 0.534399827 | 0.867307692 | 0.751094661 | 1.151489362 | 0.737721008 |
| A0A1U7Q620 | T-complex protein 1 subunit delta OS=Mesocricetus auratus OX=10036 GN=Cct4 PE=3 SV=1 | 25.171 | 14 | 7 | 33 | 6 | 539 | 58 | 8.18 | 101.21 | 7 | 402.9 | 691.3 | 404.4 | 318.4 | 554 | 355 | 298.7 | 522.6 | 347.1 | 0.79 | 0.497994365 | 0.779661017 | 0.407493758 | 0.951930911 | 0.853614266 |
| A0A1U7Q689 | T-complex protein 1 subunit epsilon isoform X1 OS=Mesocricetus auratus OX=10036 GN=Cct5 PE=3 SV=1 | 31.046 | 21 | 8 | 31 | 7 | 541 | 59.5 | 5.9 | 65.8 | 8 | 773.1 | 786.6 | 740.5 | 732.9 | 674.3 | 600.4 | 747.3 | 776 | 681.1 | 0.883 | 0.113348001 | 0.958351448 | 0.384491424 | 1.098027496 | 0.245672435 |
| A0A1U7QQT8 | T-complex protein 1 subunit eta OS=Mesocricetus auratus OX=10036 GN=Cct7 PE=3 SV=1 | 21.129 | 12 | 6 | 16 | 6 | 543 | 59.4 | 7.39 | 40.06 | 6 | 531.6 | 684.8 | 252.4 | 423.4 | 563.2 | 220.9 | 462.1 | 665.5 | 224.7 | 0.822 | 0.61858899 | 0.920683551 | 0.839369943 | 1.119917184 | 0.780823274 |
| A0A1U7QVF4 | CCT-theta OS=Mesocricetus auratus OX=10036 GN=Cct8 PE=3 SV=1 | 11.648 | 10 | 5 | 6 | 5 | 548 | 59.6 | 5.62 | 15.67 | 5 | 0 | 97.8 | 0 | 0 | 94.4 | 0 | 0 | 59.7 | 0 | 0.965 | 0.981243827 | 0.610429448 | 0.759469707 | 0.632415254 | 0.774245403 |
| A0A1U8BL34 | Monocyte differentiation antigen CD14 OS=Mesocricetus auratus OX=10036 GN=Cd14 PE=4 SV=1 | 11.115 | 15 | 3 | 8 | 3 | 370 | 40 | 4.93 | 31.04 | 3 | 0 | 87 | 0 | 0 | 97.8 | 0 | 0 | 73 | 0 | 1.124 | 0.938257687 | 0.83908046 | 0.908007331 | 0.746421268 | 0.849647923 |
| A0A1U7QHE8 | CD2-associated protein OS=Mesocricetus auratus OX=10036 GN=Cd2ap PE=4 SV=1 | 2.725 | 1 | 1 | 3 | 1 | 637 | 70.7 | 6.19 | 3.72 | 1 | 31.9 | 0 | 0 | 34.5 | 0 | 0 | 21.3 | 0 | 0 | 1.081 | 0.958541809 | 0.667711599 | 0.797867738 | 0.617391304 | 0.76415142 |
| A0A1U8CHZ6 | hematopoietic progenitor cell antigen CD34 isoform X2 OS=Mesocricetus auratus OX=10036 GN=Cd34 PE=4 SV=1 | 4.063 | 6 | 2 | 2 | 2 | 327 | 35.2 | 5.05 | 5.67 | 2 | 62 | 87.8 | 0 | 53.6 | 68.4 | 0 | 53.4 | 76.9 | 0 | 0.821 | 0.795382585 | 0.869826435 | 0.860246656 | 1.068032787 | 0.932800959 |

**Supplementary Table 1: List of differentially expressed proteins identified across the 4dpi infected and 14dpi infected**

| Accession | Description | Sum PEP Score | Coverage [%] | # Peptides | # PSMs | # Unique Peptides | # AAs | MW [kDa] | calc. pI | Score Sequest HT: Sequest HT | # Peptides (by Search Engine): Sequest HT | Mock-infected (4dpi) Replicate_R1 | Mock-infected (4dpi) Replicate_R2 | Mock-infected (4dpi) Replicate_R3 | SARS-CoV-2-infected (4dpi) Replicate_R1 | SARS-CoV-2-infected (4dpi) Replicate_R2 | SARS-CoV-2-infected (4dpi) Replicate_R3 | SARS-CoV-2-(14dpi) Replicate_R1 | SARS-CoV-2-infected (14dpi) Replicate_R2 | SARS-CoV-2-infected (14dpi) Replicate_R3 | Abundance Ratio: SARS-CoV-2-infected (4dpi) / Mock-infected (4dpi) | P-Value: SARS-CoV-2-infected (4dpi) / Mock-infected (4dpi) | Abundance Ratio: SARS-CoV-2-infected (14dpi) / Mock-infected (4dpi) | P-Value: SARS-CoV-2-infected (14dpi) / Mock-infected (4dpi) | Abundance Ratio: SARS-CoV-2-infected (14dpi) / SARS-CoV-2-infected (4dpi) | P-Value: SARS-CoV-2-infected (14dpi) / SARS-CoV-2-infected (4dpi) |
|---|---|---|---|---|---|---|---|---|---|---|---|---|---|---|---|---|---|---|---|---|---|---|---|---|---|
| A0A1U8CM00 | Glycoprotein IIIb OS=Mesocricetus auratus OX=10036 GN=Cd36 PE=3 SV=1 | 18.973 | 4 | 3 | 39 | 3 | 472 | 52.9 | 8.65 | 136.48 | 3 | 304.7 | 319.3 | 329.4 | 273.1 | 257.4 | 290.4 | 296.2 | 288.9 | 331.8 | 0.882 | 0.023661964 | 0.961715964 | 0.477089918 | 1.116944817 | 0.128642791 |
| A0A1U8BYM1 | Integrin-associated protein OS=Mesocricetus auratus OX=10036 GN=Cd47 PE=4 SV=1 | 3.248 | 2 | 1 | 9 | 1 | 321 | 34.8 | 7.74 | 20.67 | 1 | 99.2 | 35.6 | 34.8 | 118.8 | 35.8 | 36.4 | 127 | 36 | 27.7 | 1.007 | 0.848442447 | 1.124410377 | 0.864441869 | 0.998429319 | 0.998220325 |
| A0A1U7QTA0 | complement component C1q receptor OS=Mesocricetus auratus OX=10036 GN=Cd93 PE=4 SV=1 | 5.811 | 2 | 1 | 4 | 1 | 711 | 76.7 | 6.46 | 15.58 | 1 | 0 | 27.1 | 74.3 | 0 | 27.5 | 94.3 | 0 | 40.7 | 83.7 | 1.134 | 0.857693741 | 1.226824458 | 0.825140564 | 1.02134647 | 0.982450164 |
| A0A1U7R0P2 | Hsp90 chaperone protein kinase-targeting subunit OS=Mesocricetus auratus OX=10036 GN=Cdc37 PE=3 SV=1 | 8.354 | 6 | 3 | 15 | 3 | 379 | 44.5 | 5.39 | 15.78 | 3 | 605.6 | 453 | 641.2 | 440 | 367.8 | 459.9 | 590.7 | 407.7 | 642.7 | 0.751 | 0.113762868 | 0.965466525 | 0.841910399 | 1.294549184 | 0.216098095 |
| A0A1U7QUA0 | Cell division control protein 42 homolog OS=Mesocricetus auratus OX=10036 GN=Cdc42 PE=3 SV=1 | 17.65 | 37 | 5 | 25 | 5 | 191 | 21.2 | 6.55 | 86.31 | 5 | 316 | 358.2 | 311.3 | 270.3 | 294.9 | 251.2 | 374 | 390.1 | 373 | 0.855 | 0.046324589 | 1.153830543 | 0.062972756 | 1.392822146 | 0.006128474 |
| A0A1U8C5A0 | Non-specific serine/threonine protein kinase OS=Mesocricetus auratus OX=10036 GN=Cdc42bpa PE=3 SV=1 | 1.655 | 1 | 1 | 1 | 1 | 1644 | 186.5 | 6.54 | 0 | 1 | 0 | 75.9 | 0 | 0 | 54.9 | 0 | 0 | 70.2 | 0 | 0.724 | 0.834619061 | 0.924901186 | 0.958691892 | 1.278688525 | 0.872481951 |
| A0A1U8C835 | Non-specific serine/threonine protein kinase OS=Mesocricetus auratus OX=10036 GN=Cdc42bpg PE=3 SV=1 | 9.452 | 3 | 3 | 14 | 3 | 1537 | 171 | 6.19 | 20.82 | 3 | 180.7 | 158.5 | 95 | 134.5 | 125.6 | 77.1 | 162.6 | 153 | 101.2 | 0.792 | 0.36604377 | 0.959926301 | 0.865620953 | 1.236061684 | 0.367241361 |
| A0A1U7QZI8 | cadherin-1 isoform X1 OS=Mesocricetus auratus OX=10036 GN=Cdh1 PE=4 SV=1 | 9.474 | 6 | 4 | 7 | 4 | 886 | 98.3 | 4.84 | 13.87 | 4 | 158.9 | 138.8 | 75.7 | 140 | 124.2 | 65.8 | 112.1 | 116.3 | 71.1 | 0.869 | 0.690281293 | 0.802088913 | 0.453399548 | 0.907575758 | 0.72669629 |
| A0A1U7QTB8 | Cadherin-5 OS=Mesocricetus auratus OX=10036 GN=Cdh5 PE=4 SV=1 | 19.048 | 10 | 6 | 27 | 6 | 785 | 87.6 | 5.16 | 88.25 | 6 | 669.1 | 651.5 | 499.8 | 499.6 | 491.4 | 370.8 | 688.6 | 681.9 | 492 | 0.747 | 0.09196003 | 1.023126785 | 0.875545985 | 1.367675136 | 0.106748407 |
| A0A1U7QXD8 | centrin-1 OS=Mesocricetus auratus OX=10036 GN=Cetn1 PE=3 SV=1 | 1.833 | 8 | 1 | 1 | 1 | 172 | 19.6 | 4.88 | 0 | 1 | 59.9 | 0 | 0 | 46.6 | 0 | 0 | 54.3 | 0 | 0 | 0.778 | 0.869885249 | 0.906510851 | 0.948133181 | 1.165236052 | 0.919599786 |
| A0A1U7RGK9 | C3/C5 convertase OS=Mesocricetus auratus OX=10036 GN=Cfb PE=4 SV=1 | 31.286 | 17 | 11 | 42 | 11 | 764 | 85.3 | 7.58 | 62.5 | 11 | 868.7 | 1091.3 | 832.2 | 1799.1 | 1916.8 | 1547.2 | 762.1 | 887.5 | 773.4 | 1.756 | 0.004817621 | 0.867774515 | 0.268554623 | 0.460375064 | 0.007027428 |
| A0A1U8CF62 | complement factor I OS=Mesocricetus auratus OX=10036 GN=Cfi PE=4 SV=1 | 4.46 | 4 | 1 | 1 | 1 | 592 | 66.3 | 7.74 | 2.7 | 1 | 34.5 | 0 | 0 | 56.2 | 0 | 0 | 25.1 | 0 | 0 | 1.628 | 0.761789713 | 0.727536232 | 0.837373851 | 0.446619217 | 0.650832984 |
| A0A1U8D196 | cingulin OS=Mesocricetus auratus OX=10036 GN=Cgn PE=4 SV=1 | 27.765 | 9 | 7 | 22 | 7 | 1202 | 137 | 5.54 | 74.69 | 7 | 163.3 | 400.8 | 197.5 | 146.7 | 249 | 144.7 | 186.9 | 360.4 | 199 | 0.715 | 0.437220239 | 0.979910714 | 0.959021775 | 1.381014064 | 0.365946402 |
| A0A1U7R6K6 | calcium homeostasis endoplasmic reticulum protein isoform X1 OS=Mesocricetus auratus OX=10036 GN=Cherp PE=4 SV=1 | 2.148 | 1 | 1 | 1 | 1 | 939 | 106.3 | 9.32 | 2.67 | 1 | 120.6 | 0 | 0 | 56 | 0 | 0 | 95.1 | 0 | 0 | 0.465 | 0.662267286 | 0.788557214 | 0.876601431 | 1.698214286 | 0.744954593 |
| A0A1U8CLT0 | neural cell adhesion molecule L1-like protein isoform X1 OS=Mesocricetus auratus OX=10036 GN=Chl1 PE=3 SV=1 | 2.063 | 1 | 1 | 2 | 1 | 1224 | 136.5 | 5.74 | 6.04 | 1 | 39 | 0 | 78.5 | 55.5 | 0 | 80.9 | 34.5 | 0 | 56.1 | 1.211 | 0.857603349 | 0.77106383 | 0.765817713 | 0.664222874 | 0.629163243 |
| A0A1U8CRG6 | charged multivesicular body protein 2a OS=Mesocricetus auratus OX=10036 GN=Chmp2a PE=3 SV=1 | 1.678 | 3 | 1 | 2 | 1 | 222 | 25.1 | 5.97 | 1.97 | 1 | 0 | 85.9 | 0 | 0 | 81.9 | 0 | 0 | 85.1 | 0 | 0.954 | 0.974732734 | 0.990686845 | 0.995037973 | 1.039072039 | 0.979684622 |
| A0A1U7R011 | charged multivesicular body protein 4b OS=Mesocricetus auratus OX=10036 GN=Chmp4b PE=3 SV=1 | 5.833 | 8 | 2 | 4 | 2 | 224 | 24.9 | 4.82 | 2.83 | 2 | 381.7 | 140.8 | 120.2 | 303.7 | 133.8 | 117.1 | 281.5 | 131 | 100.6 | 0.796 | 0.791078735 | 0.798350708 | 0.693575481 | 0.925171295 | 0.873855543 |
| A0A1U7QAY3 | Creatine kinase OS=Mesocricetus auratus OX=10036 GN=Ckb PE=3 SV=1 | 72.873 | 48 | 15 | 110 | 14 | 381 | 42.7 | 5.67 | 327.22 | 15 | 2214.3 | 2340.3 | 2437.3 | 2725.2 | 2822.4 | 2379 | 2316.1 | 1557.5 | 2155.7 | 1.206 | 0.132023167 | 0.862326406 | 0.297122512 | 0.760641385 | 0.092867481 |
| A0A1U7R5G9 | Creatine kinase OS=Mesocricetus auratus OX=10036 GN=Ckm PE=3 SV=1 | 29.545 | 26 | 8 | 29 | 7 | 381 | 43 | 7.06 | 87.74 | 8 | 167.1 | 139.5 | 26 | 460.5 | 419.5 | 153 | 134.9 | 55 | 40.8 | 3.014 | 0.121407689 | 0.693625977 | 0.555051187 | 0.223330106 | 0.098542813 |
| A0A1U8C574 | Chloride channel CLIC-like protein 1 OS=Mesocricetus auratus OX=10036 GN=Clcc1 PE=3 SV=1 | 7.758 | 5 | 2 | 3 | 2 | 559 | 63 | 6.39 | 7.64 | 2 | 71.3 | 42.6 | 53.2 | 78.6 | 38.1 | 53.7 | 64.4 | 35.8 | 51.5 | 1.011 | 0.94341612 | 0.907839617 | 0.685301702 | 0.890258216 | 0.689945288 |
| A0A1U7QE95 | Claudin OS=Mesocricetus auratus OX=10036 GN=Cldn18 PE=3 SV=1 | 9.675 | 18 | 3 | 9 | 3 | 261 | 27.9 | 7.88 | 31.27 | 3 | 172.9 | 140.4 | 181.6 | 99.5 | 92.1 | 122 | 116.1 | 100.8 | 110.1 | 0.656 | 0.020824276 | 0.660739543 | 0.034936663 | 1.042729592 | 0.687034778 |
| A0A1U8CRR4 | Chloride intracellular channel protein OS=Mesocricetus auratus OX=10036 GN=Clic1 PE=3 SV=1 | 53.078 | 40 | 7 | 57 | 7 | 241 | 26.9 | 5.17 | 235.19 | 7 | 454.7 | 400.2 | 293.2 | 477.2 | 439.3 | 314.6 | 444.8 | 425.1 | 313.8 | 1.049 | 0.706106638 | 1.031007752 | 0.858981195 | 0.961497847 | 0.817104493 |
| A0A1U7QIG5 | Chloride intracellular channel protein OS=Mesocricetus auratus OX=10036 GN=Clic4 PE=3 SV=1 | 61.759 | 42 | 9 | 54 | 9 | 253 | 28.8 | 5.45 | 173.91 | 9 | 511.2 | 598.3 | 486.1 | 538.5 | 611.6 | 518.7 | 456.5 | 592.4 | 473 | 1.022 | 0.611272493 | 0.953810479 | 0.67758363 | 0.911972675 | 0.401543941 |
| A0A1U7R3R0 | Clathrin light chain OS=Mesocricetus auratus OX=10036 GN=Clta PE=3 SV=1 | 16.907 | 25 | 5 | 18 | 5 | 216 | 23.4 | 4.5 | 40.05 | 5 | 110 | 449.5 | 584 | 112.4 | 412.9 | 501.5 | 115.6 | 478.9 | 622.1 | 1.022 | 0.842980359 | 1.063926541 | 0.911752409 | 1.184846124 | 0.758351912 |
| A0A1U7QDR7 | Clathrin heavy chain OS=Mesocricetus auratus OX=10036 GN=Cltc PE=3 SV=1 | 121.347 | 18 | 27 | 168 | 27 | 1675 | 191.5 | 5.69 | 545.09 | 27 | 2546.5 | 2523.7 | 2549.8 | 2340.7 | 2371.7 | 2448.9 | 2328.2 | 2514.3 | 2562.5 | 0.919 | 0.034790884 | 0.971784777 | 0.421608458 | 1.034030134 | 0.381461136 |
| A0A1U8C2W4 | Clusterin OS=Mesocricetus auratus OX=10036 GN=Clu PE=3 SV=1 | 18.723 | 7 | 2 | 14 | 2 | 448 | 51.6 | 5.55 | 55.52 | 2 | 0 | 0 | 24.5 | 0 | 0 | 57.8 | 0 | 0 | 28.8 | 2.356 | 0.636351652 | 1.175510204 | 0.915070695 | 0.498269896 | 0.684457436 |
| A0A1U7QDV0 | Carboxymethylenebutenolidase homolog OS=Mesocricetus auratus OX=10036 GN=Cmbl PE=3 SV=1 | 5.688 | 7 | 1 | 2 | 1 | 245 | 28 | 6.93 | 10.38 | 1 | 0 | 94.3 | 0 | 0 | 51 | 0 | 0 | 64.7 | 0 | 0.54 | 0.712725492 | 0.686108165 | 0.810102177 | 1.268627451 | 0.876409471 |
| A0A1U7QFI7 | cellular nucleic acid-binding protein OS=Mesocricetus auratus OX=10036 GN=Cnbp PE=4 SV=1 | 5.256 | 8 | 1 | 5 | 1 | 177 | 19.5 | 7.71 | 21.93 | 1 | 0 | 0 | 33.1 | 0 | 0 | 31.6 | 0 | 0 | 38.4 | 0.956 | 0.975425049 | 1.160120846 | 0.921874714 | 1.215189873 | 0.898077 |
| A0A1U7R3A2 | Calponin OS=Mesocricetus auratus OX=10036 GN=Cnn1 PE=3 SV=1 | 19.166 | 14 | 4 | 28 | 4 | 297 | 33.3 | 8.97 | 91.11 | 4 | 510.9 | 454.6 | 662 | 377.3 | 323.9 | 485.7 | 321.8 | 282.9 | 429.5 | 0.712 | 0.137917881 | 0.635453149 | 0.066278668 | 0.871345522 | 0.475819779 |
| A0A1U7QQB3 | protein canopy homolog 3 OS=Mesocricetus auratus OX=10036 GN=Cnpy3 PE=4 SV=1 | 2.716 | 3 | 1 | 7 | 1 | 274 | 30.7 | 6.34 | 18.23 | 1 | 96 | 74.2 | 38.2 | 116.2 | 73.4 | 34.6 | 99.2 | 73.6 | 34.5 | 0.907 | 0.865461596 | 0.994721689 | 0.989120273 | 0.924620874 | 0.861315965 |
| A0A1U7QV79 | collagen alpha-1(XII) chain isoform X3 OS=Mesocricetus auratus OX=10036 GN=Col12a1 PE=4 SV=1 | 31.76 | 3 | 8 | 29 | 8 | 3120 | 339.8 | 5.68 | 109.96 | 8 | 470.3 | 375.1 | 670.8 | 377.2 | 265.7 | 524.3 | 422.1 | 324.2 | 639.5 | 0.802 | 0.369790873 | 0.913995515 | 0.750536523 | 1.187285812 | 0.57650596 |
| A0A1U8BTB6 | collagen alpha-1(XVIII) chain isoform X3 OS=Mesocricetus auratus OX=10036 GN=Col18a1 PE=4 SV=1 | 11.824 | 4 | 4 | 7 | 4 | 1517 | 155.1 | 5.99 | 23.47 | 4 | 0 | 101.6 | 0 | 0 | 93.6 | 0 | 0 | 74.8 | 0 | 0.921 | 0.956614789 | 0.736220472 | 0.843031679 | 0.799145299 | 0.88327045 |
| A0A1U7QKK9 | collagen alpha-1(I) chain isoform X1 OS=Mesocricetus auratus OX=10036 GN=Col1a1 PE=4 SV=1 | 19.607 | 4 | 5 | 18 | 5 | 1453 | 137.9 | 5.99 | 41.87 | 5 | 362.2 | 496.6 | 748.2 | 228.5 | 350 | 477.6 | 312.1 | 403.6 | 630.4 | 0.646 | 0.254432556 | 0.837647791 | 0.588026094 | 1.274595209 | 0.464674187 |
| A0A1U7R801 | collagen alpha-2(I) chain OS=Mesocricetus auratus OX=10036 GN=Col1a2 PE=4 SV=1 | 15.964 | 3 | 5 | 23 | 5 | 1378 | 129.6 | 9.04 | 33.53 | 5 | 607.1 | 852.1 | 714.1 | 337.2 | 454.1 | 364.9 | 535.2 | 740.8 | 625.2 | 0.533 | 0.02451614 | 0.874798693 | 0.384223986 | 1.644352188 | 0.032461563 |
| A0A1U7QWN2 | collagen alpha-3(IV) chain OS=Mesocricetus auratus OX=10036 GN=Col4a3 PE=4 SV=1 | 6.418 | 1 | 2 | 10 | 2 | 1622 | 156.5 | 9.44 | 35.81 | 2 | 408.6 | 435.2 | 640.2 | 293 | 288 | 429.7 | 363.3 | 409.2 | 545.4 | 0.662 | 0.15564482 | 0.888072776 | 0.579608494 | 1.303947759 | 0.228369661 |
| A0A1U7QGM7 | collagen type IV alpha-3-binding protein OS=Mesocricetus auratus OX=10036 GN=Col4a3bp PE=4 SV=1 | 2.696 | 1 | 1 | 3 | 1 | 565 | 64.2 | 5.27 | 7.23 | 1 | 196.6 | 368 | 0 | 167.7 | 303.9 | 0 | 162.1 | 320.2 | 0 | 0.839 | 0.833565859 | 0.854233085 | 0.855271138 | 1.022688719 | 0.979034023 |
| D0G6X1 | Collagen type VI alpha 1 subunit OS=Mesocricetus auratus OX=10036 GN=COL6A1 PE=2 SV=1 | 86.367 | 20 | 18 | 123 | 18 | 1026 | 108.8 | 5.27 | 391.39 | 18 | 2552.3 | 2292.9 | 2559.7 | 2152.5 | 1899.8 | 2176.9 | 2274.8 | 2079.7 | 2284.4 | 0.843 | 0.034734186 | 0.896554984 | 0.086111534 | 1.065770886 | 0.2902765 |
| D0G6X2 | Collagen type VI alpha 2 subunit OS=Mesocricetus auratus OX=10036 GN=COL6A2 PE=2 SV=1 | 84.754 | 19 | 18 | 116 | 18 | 1026 | 109.5 | 6.48 | 363.81 | 18 | 3305.4 | 4215.3 | 4192.5 | 2966.5 | 3568.7 | 3674.7 | 3198 | 3268.6 | 3516.2 | 0.848 | 0.255005026 | 0.852269235 | 0.186271256 | 0.977756883 | 0.775570001 |
| D0G7D9 | Collagen type VI alpha 3 subunit OS=Mesocricetus auratus OX=10036 GN=COL6A3 PE=2 SV=1 | 358.382 | 31 | 71 | 518 | 71 | 2675 | 288.5 | 8.34 | 1553.99 | 71 | 9268.4 | 8739.1 | 10309.1 | 7392.4 | 7279.6 | 7863.7 | 8209.5 | 8242.5 | 9541.8 | 0.793 | 0.038962088 | 0.917970378 | 0.290840216 | 1.15344986 | 0.10447834 |
| A0A1U8CEA2 | Catechol O-methyltransferase OS=Mesocricetus auratus OX=10036 GN=Comt PE=3 SV=1 | 8 | 11 | 3 | 10 | 3 | 264 | 29.6 | 6.89 | 27.83 | 3 | 358 | 331.1 | 194.6 | 302.3 | 295.7 | 178.6 | 331.1 | 314.8 | 206.2 | 0.885 | 0.61135464 | 0.964241258 | 0.87771473 | 1.097218645 | 0.677136949 |
| A0A1U7QYD2 | Coatomer subunit beta OS=Mesocricetus auratus OX=10036 GN=Copb1 PE=4 SV=1 | 16.968 | 6 | 4 | 18 | 4 | 953 | 107 | 6 | 55.45 | 4 | 197.4 | 197.7 | 211.8 | 127.4 | 145.1 | 138.8 | 183.9 | 182.3 | 221.4 | 0.734 | 0.000777871 | 0.968199044 | 0.674411661 | 1.428640895 | 0.030448826 |
| A0A1U7QLH6 | COP9 signalosome complex subunit 2 OS=Mesocricetus auratus OX=10036 GN=Cops2 PE=3 SV=1 | 17.215 | 17 | 6 | 12 | 6 | 443 | 51.6 | 5.53 | 31.48 | 6 | 94.5 | 86.6 | 77.6 | 76 | 91.2 | 74.1 | 83.3 | 68.5 | 73.2 | 0.804 | 0.471130729 | 0.869733282 | 0.162356984 | 0.932449233 | 0.480034366 |
| A0A1U7R6Y5 | COP9 signalosome complex subunit 8 OS=Mesocricetus auratus OX=10036 GN=Cops8 PE=3 SV=1 | 10.593 | 23 | 3 | 9 | 3 | 209 | 23.2 | 5.2 | 23.62 | 3 | 51 | 43.4 | 76.8 | 40.7 | 35 | 63 | 37.6 | 33.9 | 56.6 | 0.82 | 0.460187033 | 0.748247664 | 0.315302575 | 0.923576063 | 0.766018676 |
| A0A1U8BRU3 | Coronin OS=Mesocricetus auratus OX=10036 GN=Coro1a PE=3 SV=1 | 48.269 | 21 | 8 | 45 | 8 | 461 | 50.9 | 6.48 | 123.35 | 8 | 486.8 | 335.4 | 286.8 | 548.9 | 358.9 | 302.5 | 455.2 | 273.3 | 282.1 | 1.098 | 0.743075737 | 0.911271416 | 0.717556403 | 0.834999587 | 0.524763016 |
| A0A1U8CAN2 | Coronin OS=Mesocricetus auratus OX=10036 GN=Coro1b PE=3 SV=1 | 14.107 | 8 | 4 | 18 | 4 | 484 | 53.8 | 5.78 | 59.32 | 4 | 270.1 | 227.3 | 141.5 | 263.3 | 221.4 | 148.6 | 225.9 | 215.3 | 118.9 | 0.974 | 0.972321013 | 0.876663015 | 0.633147698 | 0.884414969 | 0.63637139 |
| A0A1U7QIT6 | coactosin-like protein OS=Mesocricetus auratus OX=10036 GN=Cotl1 PE=4 SV=1 | 4.311 | 11 | 2 | 4 | 2 | 142 | 16 | 5.4 | 9.38 | 2 | 85.8 | 49 | 47.6 | 114.2 | 62.1 | 69.6 | 92.7 | 60.2 | 66.3 | 1.269 | 0.364020119 | 1.201754386 | 0.487909999 | 0.891419276 | 0.669768134 |
| C9VZR0 | Cytochrome c oxidase subunit 2 OS=Mesocricetus auratus OX=10036 GN=COX2 PE=3 SV=1 | 10.978 | 15 | 3 | 14 | 3 | 227 | 25.9 | 4.73 | 38.15 | 3 | 184.6 | 223.9 | 67.5 | 159.6 | 226.2 | 75.2 | 146.7 | 194.3 | 57.6 | 0.978 | 0.941637232 | 0.837394958 | 0.69799034 | 0.864642082 | 0.743500935 |
| A0A1U8CUP5 | copine-1 isoform X3 OS=Mesocricetus auratus OX=10036 GN=Cpne1 PE=3 SV=1 | 5.243 | 2 | 2 | 5 | 2 | 1531 | 161.7 | 6.9 | 11.37 | 2 | 0 | 0 | 34.8 | 0 | 0 | 25.1 | 0 | 0 | 37.4 | 0.722 | 0.833273967 | 1.074712644 | 0.961861815 | 1.490039841 | 0.800141211 |
| A0A1U7QEH1 | carnitine O-palmitoyltransferase 2, mitochondrial OS=Mesocricetus auratus OX=10036 GN=Cpt2 PE=3 SV=1 | 4.344 | 2 | 1 | 4 | 1 | 656 | 74 | 7.14 | 12.29 | 1 | 0 | 32.1 | 0 | 0 | 37.7 | 0 | 0 | 34.1 | 0 | 1.175 | 0.915533463 | 1.062305296 | 0.967989759 | 0.904509284 | 0.946973751 |
| A0A1U7QIK1 | cysteine-rich protein 1 OS=Mesocricetus auratus OX=10036 GN=LOC101822555 PE=4 SV=1 | 13.113 | 35 | 4 | 30 | 4 | 77 | 8.5 | 8.57 | 88.8 | 4 | 1413.8 | 1813.7 | 1969 | 1283.8 | 1487.5 | 1752.7 | 1207.3 | 1646.3 | 2103.6 | 0.86 | 0.355974381 | 0.953949774 | 0.810006496 | 1.095755968 | 0.654859519 |
| A0A1U7QKZ6 | cysteine-rich protein 2 isoform X3 OS=Mesocricetus auratus OX=10036 GN=Crip2 PE=4 SV=1 | 24.763 | 39 | 5 | 13 | 5 | 209 | 22.8 | 8.63 | 63.12 | 5 | 204.9 | 161.3 | 0 | 178.9 | 110.3 | 0 | 175.6 | 126.5 | 0 | 0.773 | 0.768301153 | 0.824959039 | 0.806186313 | 1.044605809 | 0.95635168 |
| A0A1U7Q9Y2 | adapter molecule crk isoform X1 OS=Mesocricetus auratus OX=10036 GN=Crk PE=3 SV=1 | 25.87 | 22 | 6 | 30 | 6 | 304 | 33.8 | 5.55 | 104.51 | 6 | 185.4 | 353.2 | 94.5 | 158.1 | 318.3 | 85.8 | 135.6 | 312 | 73.1 | 0.853 | 0.828701219 | 0.822460907 | 0.737411512 | 0.926182853 | 0.89581978 |
| A0A1U7QLW5 | dihydropyrimidinase-related protein 1 isoform X2 OS=Mesocricetus auratus OX=10036 GN=Crmp1 PE=3 SV=1 | 15.35 | 8 | 4 | 16 | 1 | 572 | 62 | 7.12 | 32.07 | 4 | 0 | 127.9 | 109.3 | 0 | 151.2 | 95.4 | 0 | 113.7 | 109.2 | 1.015 | 0.960552215 | 0.939713322 | 0.934563865 | 0.903892944 | 0.897912999 |
| Q6LEK0 | Alpha(B)-crystallin (Fragment) OS=Mesocricetus auratus OX=10036 PE=2 SV=1 | 8.72 | 8 | 1 | 9 | 1 | 172 | 19.7 | 6.81 | 28.34 | 1 | 37.8 | 40.1 | 0 | 37.8 | 38.6 | 0 | 33.1 | 31.7 | 0 | 0.98 | 0.979398096 | 0.831835687 | 0.809330821 | 0.848167539 | 0.828576739 |
| A0A1U7QV33 | Citrate synthase OS=Mesocricetus auratus OX=10036 GN=Cs PE=3 SV=1 | 43.208 | 23 | 10 | 52 | 10 | 466 | 51.7 | 8.06 | 168.75 | 10 | 1543.1 | 1072.3 | 1388.4 | 1539.9 | 1063.7 | 1241 | 1437 | 1165.2 | 1345.7 | 0.952 | 0.800199844 | 0.986038264 | 0.914177524 | 1.026868855 | 0.842888687 |
| A0A1U8C9A2 | Chromosome segregation 1-like protein OS=Mesocricetus auratus OX=10036 GN=Cse1l PE=3 SV=1 | 8.101 | 4 | 2 | 4 | 2 | 971 | 110.3 | 5.77 | 11.56 | 2 | 0 | 0 | 57.5 | 0 | 0 | 35.4 | 0 | 0 | 44.1 | 0.616 | 0.762982081 | 0.766956522 | 0.862863073 | 1.245762712 | 0.885512611 |
| A0A1U7QKS8 | Tyrosine-protein kinase OS=Mesocricetus auratus OX=10036 GN=Csk PE=3 SV=1 | 3.618 | 4 | 2 | 4 | 2 | 450 | 50.7 | 7.06 | 7.7 | 2 | 66.5 | 93.6 | 122.7 | 61.3 | 89.6 | 106 | 52.9 | 89.4 | 84.2 | 0.921 | 0.700674767 | 0.800919378 | 0.40325203 | 0.881666018 | 0.590694901 |
| A0A1U7QJB5 | cysteine and glycine-rich protein 1 OS=Mesocricetus auratus OX=10036 GN=Csrp1 PE=4 SV=1 | 26.207 | 27 | 3 | 19 | 3 | 193 | 20.5 | 8.57 | 68.43 | 3 | 281.4 | 138.7 | 236.5 | 229 | 126.9 | 204 | 346.9 | 145 | 250.3 | 0.814 | 0.572843491 | 1.130368565 | 0.713734128 | 1.325593856 | 0.42385798 |
| A0A1U7QI90 | cysteine and glycine-rich protein 2 OS=Mesocricetus auratus OX=10036 GN=Csrp2 PE=4 SV=1 | 1.879 | 4 | 1 | 1 | 1 | 193 | 20.9 | 8.62 | 0 | 1 | 0 | 39.7 | 0 | 0 | 46.2 | 0 | 0 | 34.5 | 0 | 1.162 | 0.920269105 | 0.869017632 | 0.926088428 | 0.746753247 | 0.849862688 |
| A0A1U8BPJ7 | Protein-serine/threonine phosphatase OS=Mesocricetus auratus OX=10036 GN=Ctdsp1 PE=4 SV=1 | 1.845 | 4 | 1 | 1 | 1 | 261 | 29.4 | 5.81 | 2.58 | 1 | 0 | 152.9 | 0 | 0 | 165.2 | 0 | 0 | 135.5 | 0 | 1.081 | 0.95905832 | 0.886200131 | 0.936276058 | 0.820217918 | 0.89640795 |
| A0A1U7QDJ2 | catenin alpha-1 OS=Mesocricetus auratus OX=10036 GN=Ctnna1 PE=3 SV=1 | 56.08 | 15 | 11 | 66 | 11 | 905 | 100 | 6.29 | 206.49 | 11 | 1247.2 | 967.8 | 1061.3 | 986.8 | 809.4 | 803.9 | 1221.4 | 873.1 | 1046.1 | 0.759 | 0.097265453 | 0.958581326 | 0.745738286 | 1.20787662 | 0.214289074 |
| A0A1U8CE17 | catenin beta-1 OS=Mesocricetus auratus OX=10036 GN=Ctnnb1 PE=3 SV=1 | 17.992 | 10 | 5 | 25 | 4 | 781 | 85.4 | 5.86 | 59.63 | 5 | 163.5 | 274.1 | 309.2 | 120.1 | 209 | 244.3 | 167.9 | 268.8 | 312.9 | 0.776 | 0.372322976 | 1.00374933 | 0.988598928 | 1.30728985 | 0.35944978 |
| A0A1U7QYR0 | CTP synthase OS=Mesocricetus auratus OX=10036 GN=Ctps1 PE=3 SV=1 | 3.216 | 1 | 1 | 5 | 1 | 591 | 66.6 | 6.67 | 14.23 | 1 | 165.3 | 67.7 | 140.4 | 133.7 | 45.9 | 118.1 | 143.7 | 81 | 145.5 | 0.809 | 0.561273558 | 0.991430102 | 0.978005471 | 1.243533759 | 0.522775893 |
| A0A1U8BXC8 | CTP synthase OS=Mesocricetus auratus OX=10036 GN=Ctps2 PE=3 SV=1 | 1.353 | 3 | 1 | 1 | 1 | 430 | 48.5 | 7.47 | 3.22 | 1 | 0 | 34.8 | 0 | 0 | 36.4 | 0 | 0 | 36.6 | 0 | 1.045 | 0.976178888 | 1.051724138 | 0.973280357 | 1.005494505 | 0.997094107 |
| A0A1U7QV28 | Cathepsin B OS=Mesocricetus auratus OX=10036 GN=Ctsb PE=3 SV=1 | 11.305 | 14 | 3 | 11 | 3 | 339 | 37.4 | 6.27 | 22.39 | 3 | 66.3 | 95.6 | 297.1 | 76.8 | 135 | 371.3 | 63.4 | 57.9 | 244 | 1.375 | 0.739394537 | 0.795860566 | 0.758956728 | 0.626479163 | 0.545900756 |
| A0A1U7R496 | pro-cathepsin H OS=Mesocricetus auratus OX=10036 GN=Ctsh PE=3 SV=1 | 5.34 | 3 | 1 | 5 | 1 | 333 | 37.4 | 8.4 | 14.24 | 1 | 69.1 | 100.5 | 82.9 | 115.7 | 137.9 | 137.6 | 62.2 | 97.9 | 82.7 | 1.674 | 0.018176703 | 0.961584158 | 0.826041175 | 0.620654397 | 0.021321694 |
| A0A1U7Q2W5 | src substrate cortactin isoform X1 OS=Mesocricetus auratus OX=10036 GN=Cttn PE=4 SV=1 | 8.55 | 6 | 4 | 18 | 4 | 554 | 61.8 | 5.27 | 18.39 | 4 | 441.2 | 433.6 | 206.8 | 356.5 | 332.7 | 168.7 | 375.7 | 401.4 | 182.4 | 0.816 | 0.487371331 | 0.887111686 | 0.714143592 | 1.118428721 | 0.728795817 |
| A0A1U7QSN5 | cullin-2 OS=Mesocricetus auratus OX=10036 GN=Cul2 PE=3 SV=1 | 4.538 | 2 | 1 | 4 | 1 | 745 | 86.8 | 7.02 | 12.83 | 1 | 31.3 | 0 | 0 | 32.5 | 0 | 0 | 36.8 | 0 | 0 | 1.038 | 0.980058459 | 1.17571885 | 0.914979274 | 1.132307692 | 0.934479266 |

Supplementary Table 1: List of differentially expressed proteins identified across the 4dpi infected and 14dpi infected

| Accession | Description | Sum PEP Score | Coverage [%] | # Peptides | # PSMs | # Unique Peptides | # AAs | MW [kDa] | calc. pI | Score Sequest HT: Sequest HT | # Peptides (by Search Engine): Sequest HT | Mock-infected (4dpi) Replicate_R1 | Mock-infected (4dpi) Replicate_R2 | Mock-infected (4dpi) Replicate_R3 | SARS-CoV-2-infected (4dpi) Replicate_R1 | SARS-CoV-2-infected (4dpi) Replicate_R2 | SARS-CoV-2-infected (4dpi) Replicate_R3 | SARS-CoV-2-(14dpi) Replicate_R1 | SARS-CoV-2-infected (14dpi) Replicate_R2 | SARS-CoV-2-infected (14dpi) Replicate_R3 | Abundance Ratio: SARS-CoV-2-infected (4dpi) / Mock-infected (4dpi) | P-Value: SARS-CoV-2-infected (4dpi) / Mock-infected (4dpi) | Abundance Ratio: SARS-CoV-2-infected (14dpi) / Mock-infected (4dpi) | P-Value: SARS-CoV-2-infected (14dpi) / Mock-infected (4dpi) | Abundance Ratio: SARS-CoV-2-infected (14dpi) / SARS-CoV-2-infected (4dpi) | P-Value: SARS-CoV-2-infected (14dpi) / SARS-CoV-2-infected (4dpi) |
|---|---|---|---|---|---|---|---|---|---|---|---|---|---|---|---|---|---|---|---|---|---|---|---|---|---|
| A0A1U7QAV2 | coxsackievirus and adenovirus receptor isoform X2 OS=Mesocricetus auratus OX=10036 GN=Cxadr PE=4 SV=1 | 6.569 | 9 | 2 | 4 | 2 | 352 | 38.8 | 7.27 | 15.51 | 2 | 0 | 0 | 91.5 | 0 | 0 | 44.8 | 0 | 0 | 78.8 | 0.49 | 0.678763858 | 0.861202186 | 0.921408135 | 1.758928571 | 0.731290268 |
| A0A1U7R5Q6 | cytohesin-3 isoform X1 OS=Mesocricetus auratus OX=10036 GN=Cyth3 PE=4 SV=1 | 5.541 | 6 | 2 | 3 | 2 | 399 | 46.3 | 5.54 | 5.05 | 2 | 0 | 59.9 | 0 | 0 | 50.3 | 0 | 0 | 30.3 | 0 | 0.84 | 0.908405734 | 0.505843072 | 0.689457695 | 0.602385686 | 0.754058563 |
| A0A1U8C202 | Alpha-dystroglycan OS=Mesocricetus auratus OX=10036 GN=Dag1 PE=4 SV=1 | 17.277 | 12 | 4 | 12 | 4 | 895 | 97.1 | 8.66 | 33.15 | 4 | 0 | 56 | 51.4 | 0 | 41.5 | 49.1 | 0 | 48.9 | 34.8 | 0.842 | 0.82407481 | 0.779329609 | 0.750217157 | 0.92384106 | 0.918348975 |
| A0A1U7QBI0 | acyl-CoA-binding protein OS=Mesocricetus auratus OX=10036 GN=Dbi PE=4 SV=1 | 12.83 | 43 | 4 | 23 | 4 | 87 | 10 | 8.81 | 16.34 | 4 | 966.3 | 707.2 | 842.2 | 751 | 570.1 | 618.2 | 1002.3 | 709.7 | 872.1 | 0.806 | 0.112715809 | 1.027189252 | 0.850037489 | 1.332491105 | 0.111204992 |
| A0A1U8BM80 | Decorin OS=Mesocricetus auratus OX=10036 GN=Dcn PE=3 SV=1 | 27.335 | 22 | 10 | 52 | 9 | 362 | 40 | 8.19 | 102.31 | 10 | 2016.9 | 2222.5 | 1835.5 | 1640.1 | 1729.4 | 1419.6 | 1795 | 1918.9 | 1580.1 | 0.781 | 0.043491875 | 0.871454674 | 0.157269094 | 1.105426907 | 0.281392023 |
| A0A1U7QPU2 | dynactin subunit 1 isoform X5 OS=Mesocricetus auratus OX=10036 GN=Dctn1 PE=4 SV=1 | 15.032 | 7 | 7 | 13 | 7 | 1142 | 126.9 | 5.47 | 2.07 | 7 | 263 | 282.6 | 210.6 | 209.2 | 241.7 | 184.3 | 217.5 | 264.6 | 200.1 | 0.796 | 0.216170963 | 0.90214229 | 0.441463985 | 1.073992443 | 0.572046114 |
| A0A1U7QIR5 | dynactin subunit 2 isoform X2 OS=Mesocricetus auratus OX=10036 GN=Dctn2 PE=3 SV=1 | 39.719 | 35 | 11 | 32 | 11 | 404 | 44.5 | 5.08 | 100.22 | 11 | 271.9 | 432.3 | 346.1 | 229.3 | 341.8 | 299.1 | 266.1 | 388.4 | 280.6 | 0.843 | 0.356025559 | 0.890317052 | 0.559945977 | 1.074580556 | 0.691707328 |
| A0A1U7R164 | dynactin subunit 3 OS=Mesocricetus auratus OX=10036 GN=Dctn3 PE=4 SV=1 | 3.399 | 9 | 2 | 2 | 2 | 186 | 21.1 | 5.48 | 2.66 | 2 | 38.9 | 0 | 0 | 33.6 | 0 | 0 | 36.7 | 0 | 0 | 0.864 | 0.922938976 | 0.94344473 | 0.969164503 | 1.092261905 | 0.953333533 |
| A0A1U7Q257 | L-xylulose reductase OS=Mesocricetus auratus OX=10036 GN=Dcxr PE=4 SV=1 | 4.776 | 7 | 2 | 4 | 2 | 244 | 25.7 | 7.02 | 6.52 | 2 | 73.4 | 84 | 95.2 | 62.8 | 86.3 | 60.5 | 46.5 | 80 | 72.3 | 0.84 | 0.243763354 | 0.787015044 | 0.220540201 | 0.948473282 | 0.797012213 |
| A0A1U7R660 | Dimethylargininase OS=Mesocricetus auratus OX=10036 GN=Ddah2 PE=3 SV=1 | 15.628 | 16 | 4 | 22 | 4 | 285 | 29.5 | 5.81 | 90.56 | 4 | 799.1 | 196.9 | 788.4 | 532 | 178.4 | 582.9 | 674.5 | 199.6 | 703.4 | 0.767 | 0.532592832 | 0.884050661 | 0.802493456 | 1.219747932 | 0.672333744 |
| A0A1U8CLX3 | protein DDI1 homolog 2 isoform X2 OS=Mesocricetus auratus OX=10036 GN=LOC101843704 PE=3 SV=1 | 7.784 | 11 | 2 | 7 | 2 | 399 | 44.5 | 5.05 | 26.04 | 2 | 44 | 0 | 0 | 42.8 | 0 | 0 | 38.4 | 0 | 0 | 0.974 | 0.985339744 | 0.872727273 | 0.928299861 | 0.897196262 | 0.942720689 |
| A0A1U7QX96 | Dolichyl-diphosphooligosaccharide--protein glycosyltransferase 48 kDa subunit OS=Mesocricetus auratus OX=10036 GN=Ddost PE=3 SV=1 | 5.152 | 4 | 2 | 13 | 2 | 441 | 48.9 | 5.97 | 25.13 | 2 | 324.1 | 297.2 | 314.1 | 279 | 282 | 302.2 | 315 | 295.8 | 336.4 | 0.93 | 0.088288716 | 1.012614924 | 0.796150876 | 1.097312326 | 0.126122701 |
| A0A1U7R640 | RNA helicase OS=Mesocricetus auratus OX=10036 GN=Ddx39b PE=4 SV=1 | 18.939 | 16 | 5 | 23 | 5 | 428 | 49 | 5.67 | 44.79 | 5 | 629.9 | 632.9 | 601.3 | 648.1 | 583.2 | 568.9 | 513.8 | 518.2 | 508.2 | 0.946 | 0.485041777 | 0.826243227 | 0.005413354 | 0.855571603 | 0.068818106 |
| A0A1U8CKS0 | RNA helicase OS=Mesocricetus auratus OX=10036 GN=Ddx3x PE=3 SV=1 | 46.472 | 17 | 9 | 53 | 8 | 661 | 73.1 | 7.18 | 142.5 | 9 | 364.9 | 190.8 | 190.8 | 313.9 | 185.2 | 270.5 | 311.5 | 173.2 | 173.2 | 0.876 | 0.917656948 | 0.881312793 | 0.711617899 | 0.854859667 | 0.567355065 |
| A0A1U7R308 | RNA helicase OS=Mesocricetus auratus OX=10036 GN=Ddx46 PE=4 SV=1 | 7.482 | 3 | 2 | 7 | 2 | 1031 | 117.1 | 9.29 | 24.99 | 2 | 135.2 | 134 | 155.4 | 143.9 | 157 | 159.5 | 118.6 | 102.1 | 120.9 | 1.065 | 0.239292565 | 0.804521903 | 0.039948899 | 0.74196351 | 0.007344882 |
| A0A1U8BRT1 | DEAD box protein 5 OS=Mesocricetus auratus OX=10036 GN=Ddx5 PE=3 SV=1 | 42.312 | 18 | 10 | 54 | 7 | 615 | 69.2 | 8.92 | 111.54 | 10 | 1591.6 | 920.8 | 1329 | 1432 | 916.2 | 1215.5 | 1390.9 | 911.5 | 1293.1 | 0.957 | 0.726889013 | 0.93598688 | 0.754946301 | 1.00892331 | 0.962013438 |
| A0A1U7QCK1 | RNA helicase OS=Mesocricetus auratus OX=10036 GN=Ddx6 PE=3 SV=1 | 2.348 | 2 | 1 | 6 | 1 | 483 | 54.3 | 8.66 | 6.17 | 1 | 243.6 | 199.6 | 212.7 | 220.7 | 195.1 | 176.7 | 202.9 | 173.7 | 224.3 | 0.906 | 0.311231257 | 0.916145754 | 0.403791224 | 1.014177215 | 0.892554124 |
| Q54A94 | Desmin OS=Mesocricetus auratus OX=10036 GN=Des PE=2 SV=1 | 114.714 | 45 | 24 | 173 | 19 | 469 | 53.4 | 5.27 | 526.68 | 24 | 1940.9 | 2277.3 | 3190.5 | 2239 | 2406.2 | 2908 | 2399.6 | 2680 | 2924.6 | 1.096 | 0.916577497 | 1.080378474 | 0.660263619 | 1.059709792 | 0.584933243 |
| A0A1U8BR45 | RNA helicase OS=Mesocricetus auratus OX=10036 GN=Dhx9 PE=3 SV=1 | 35.512 | 9 | 7 | 40 | 7 | 1387 | 149.8 | 6.83 | 138.72 | 7 | 980.1 | 891.3 | 767.2 | 923.8 | 815.6 | 700.2 | 877.7 | 802.1 | 658.8 | 0.915 | 0.499057329 | 0.886303343 | 0.324445799 | 0.95859977 | 0.730277882 |
| A0A1U7Q518 | protein diaphanous homolog 1 isoform X4 OS=Mesocricetus auratus OX=10036 GN=Diaph1 PE=3 SV=1 | 11.214 | 4 | 3 | 4 | 3 | 1243 | 138 | 5.48 | 13.41 | 3 | 0 | 35.6 | 0 | 0 | 40.2 | 0 | 0 | 36.6 | 0 | 1.13 | 0.935906221 | 1.028089888 | 0.985312683 | 0.910447761 | 0.950407844 |
| Q811C4 | Dihydrolipoyl dehydrogenase, mitochondrial (Fragment) OS=Mesocricetus auratus OX=10036 GN=DLD PE=1 SV=1 | 13.326 | 9 | 4 | 12 | 4 | 479 | 50.7 | 7.28 | 37.2 | 4 | 108 | 80.8 | 0 | 97.1 | 81.8 | 0 | 112.7 | 70.6 | 0 | 0.954 | 0.944183153 | 0.970868644 | 0.970236819 | 1.024594746 | 0.975355517 |
| A0A1U7QDI4 | disks large homolog 1 isoform X8 OS=Mesocricetus auratus OX=10036 GN=Dlg1 PE=3 SV=1 | 6.892 | 3 | 3 | 8 | 3 | 891 | 99.4 | 6.07 | 20.79 | 3 | 460.8 | 217.4 | 169.8 | 372.9 | 177.2 | 158.2 | 425.8 | 181.6 | 152.2 | 0.818 | 0.703473158 | 0.895754717 | 0.825323305 | 1.072426938 | 0.884972121 |
| A0A1U7QUP4 | Dihydrolipoyllysine-residue succinyltransferase component of 2-oxoglutarate dehydrogenase complex, mitochondrial OS=Mesocricetus auratus OX=10036 GN=Dlst PE=3 SV=1 | 2.402 | 2 | 1 | 3 | 1 | 453 | 48.8 | 8.38 | 7.5 | 1 | 60 | 69.5 | 75.2 | 51.1 | 64.6 | 57.1 | 71 | 73.3 | 84.9 | 0.835 | 0.147409171 | 1.119687347 | 0.256739539 | 1.326388889 | 0.032223008 |
| A0A1U7R1Q6 | dnaJ homolog subfamily A member 2 OS=Mesocricetus auratus OX=10036 GN=Dnaja2 PE=3 SV=1 | 2.833 | 4 | 1 | 1 | 1 | 412 | 45.8 | 6.48 | 4.01 | 1 | 0 | 0 | 97.2 | 0 | 0 | 104.5 | 0 | 0 | 99.6 | 1.075 | 0.961670403 | 1.024691358 | 0.98706735 | 0.953110048 | 0.974552847 |
| A0A1U7QTX3 | dnaJ homolog subfamily C member 5 OS=Mesocricetus auratus OX=10036 GN=Dnajc5 PE=4 SV=1 | 3.449 | 9 | 1 | 2 | 1 | 198 | 22.1 | 5.07 | 8.36 | 1 | 0 | 75.9 | 0 | 0 | 70.3 | 0 | 0 | 84.8 | 0 | 0.927 | 0.959441777 | 1.117259552 | 0.941465808 | 1.20625889 | 0.901833555 |
| A0A1U7QEC2 | Aspartyl aminopeptidase OS=Mesocricetus auratus OX=10036 GN=Dnpep PE=3 SV=1 | 16.741 | 7 | 3 | 19 | 3 | 475 | 52.5 | 7.24 | 42.52 | 3 | 77.5 | 39.1 | 31.6 | 52 | 28.5 | 32.8 | 62.9 | 39.4 | 30.5 | 0.795 | 0.518986057 | 0.89608637 | 0.781993593 | 1.172109444 | 0.620828756 |
| A0A1U7QHP0 | 2'-deoxynucleoside 5'-phosphate N-hydrolase 1 OS=Mesocricetus auratus OX=10036 GN=Dnph1 PE=3 SV=1 | 2.358 | 7 | 1 | 5 | 1 | 163 | 17.6 | 5.05 | 9.76 | 1 | 43 | 50.1 | 163.5 | 29.6 | 44.6 | 134.2 | 34.5 | 36.6 | 123.3 | 0.841 | 0.768493364 | 0.757599376 | 0.694323708 | 0.932821497 | 0.920417433 |
| A0A1U8CGQ2 | dedicator of cytokinesis protein 6 isoform X3 OS=Mesocricetus auratus OX=10036 GN=Dock6 PE=3 SV=1 | 5.76 | 1 | 2 | 4 | 2 | 2079 | 232.3 | 6.8 | 12.38 | 2 | 99.9 | 0 | 250.3 | 80.1 | 0 | 208.7 | 110.1 | 0 | 253 | 0.817 | 0.839946675 | 1.036836094 | 0.968769309 | 1.257271468 | 0.807959028 |
| A0A1U7QWB3 | Dolichol-phosphate mannosyltransferase subunit 1 OS=Mesocricetus auratus OX=10036 GN=Dpm1 PE=3 SV=1 | 1.438 | 3 | 1 | 8 | 1 | 260 | 29.1 | 9.57 | 12.92 | 1 | 92.3 | 177.2 | 64.4 | 72.9 | 94.9 | 45.7 | 78.1 | 162.9 | 54.5 | 0.622 | 0.363438837 | 0.884995508 | 0.799939385 | 1.384074941 | 0.506464673 |
| A0A1U8BTS8 | Dipeptidyl peptidase 3 OS=Mesocricetus auratus OX=10036 GN=Dpp3 PE=3 SV=1 | 19.893 | 13 | 7 | 15 | 7 | 740 | 82.9 | 5.19 | 56.74 | 7 | 0 | 76.8 | 225.5 | 0 | 75.1 | 191.5 | 0 | 69.3 | 213.2 | 0.912 | 0.897425052 | 0.93450215 | 0.945811898 | 1.05963991 | 0.952724601 |
| A0A075TJ59 | Dipeptidylpeptidase 4 OS=Mesocricetus auratus OX=10036 GN=DPP4 PE=2 SV=1 | 59.505 | 19 | 14 | 85 | 14 | 764 | 87.9 | 6.4 | 256.68 | 14 | 1478.9 | 1327.6 | 2232.7 | 2089.7 | 1761.1 | 2089.2 | 2407.1 | 1919.1 | 2341 | 1.413 | 0.40154493 | 1.323067154 | 0.184575421 | 1.122424242 | 0.273387724 |
| A0A1U8CMG1 | dipeptidyl peptidase 9 isoform X1 OS=Mesocricetus auratus OX=10036 GN=Dpp9 PE=4 SV=1 | 7.945 | 4 | 2 | 2 | 2 | 923 | 105.2 | 7.03 | 6.07 | 2 | 94.9 | 0 | 0 | 74 | 0 | 0 | 90.5 | 0 | 0 | 0.78 | 0.871032498 | 0.953635406 | 0.974844342 | 1.222972973 | 0.894835257 |
| A0A1U7QXQ0 | Dihydropyrimidinase-related protein 2 OS=Mesocricetus auratus OX=10036 GN=Dpysl2 PE=3 SV=1 | 132.05 | 36 | 19 | 171 | 16 | 673 | 73.1 | 6.4 | 671.88 | 19 | 3781.4 | 4059.3 | 2945.5 | 2551.4 | 2661.7 | 2041.7 | 3926.7 | 4008 | 3933.7 | 0.658 | 0.051230047 | 1.100331906 | 0.393871737 | 1.635937586 | 0.013791769 |
| A0A1U7Q9J6 | Actin-depolymerizing factor OS=Mesocricetus auratus OX=10036 GN=Dstn PE=3 SV=1 | 23.052 | 48 | 6 | 21 | 6 | 165 | 18.6 | 8.05 | 51.46 | 6 | 61.9 | 156.8 | 159.6 | 74.7 | 165.6 | 145.9 | 51.7 | 134.2 | 166.9 | 1.118 | 0.953463211 | 0.93259318 | 0.865219449 | 0.913516313 | 0.813302529 |
| A0A1U7QGE3 | RING-type E3 ubiquitin transferase OS=Mesocricetus auratus OX=10036 GN=Dtx3l PE=3 SV=1 | 3.023 | 1 | 1 | 1 | 1 | 765 | 84.8 | 7.05 | 1.79 | 1 | 34 | 0 | 0 | 42.4 | 0 | 0 | 28.1 | 0 | 0 | 1.248 | 0.884989937 | 0.826470588 | 0.900270159 | 0.662735849 | 0.794548198 |
| A0A1U7QXC8 | Dynein light intermediate chain OS=Mesocricetus auratus OX=10036 GN=Dync1li1 PE=3 SV=1 | 6.126 | 7 | 3 | 5 | 3 | 523 | 56.5 | 6.37 | 7.43 | 3 | 28.9 | 0 | 50.9 | 50 | 0 | 56.7 | 23.9 | 0 | 42.9 | 1.389 | 0.719239701 | 0.837092732 | 0.833384168 | 0.626054358 | 0.57807848 |
| A0A1U8BQ56 | ethylmalonyl-CoA decarboxylase isoform X1 OS=Mesocricetus auratus OX=10036 GN=Echdc1 PE=3 SV=1 | 5.022 | 11 | 3 | 3 | 3 | 299 | 32.6 | 7.46 | 7.89 | 3 | 131.6 | 0 | 0 | 110.3 | 0 | 0 | 112.8 | 0 | 0 | 0.838 | 0.907436312 | 0.857142857 | 0.918965923 | 1.022665458 | 0.988116251 |
| A0A1U8CFA4 | enoyl-CoA delta isomerase 1, mitochondrial OS=Mesocricetus auratus OX=10036 GN=Eci1 PE=3 SV=1 | 6.986 | 8 | 2 | 15 | 2 | 289 | 32.3 | 9.39 | 51.44 | 2 | 109.5 | 72.7 | 92.8 | 84.6 | 64.2 | 105 | 71.1 | 66.8 | 77.7 | 1.132 | 0.679396611 | 0.784 | 0.197164399 | 0.849487786 | 0.394079733 |
| A0A1U8BJP0 | early endosome antigen 1 OS=Mesocricetus auratus OX=10036 GN=Eea1 PE=4 SV=1 | 1.452 | 1 | 1 | 1 | 1 | 1411 | 161.5 | 5.54 | 2.08 | 1 | 58.8 | 0 | 0 | 54 | 0 | 0 | 55.5 | 0 | 0 | 0.918 | 0.954960057 | 0.943877551 | 0.969406831 | 1.027777778 | 0.98547357 |
| A0A1U7Q9F2 | Elongation factor 1-alpha OS=Mesocricetus auratus OX=10036 GN=Eef1a1 PE=3 SV=1 | 59.432 | 33 | 14 | 100 | 14 | 462 | 50.1 | 9.01 | 213.44 | 14 | 3934.9 | 3415.1 | 3738.2 | 4504.1 | 3524.8 | 4185.5 | 4217.1 | 3444.9 | 4799.9 | 1.145 | 0.332030934 | 1.123888458 | 0.367381571 | 1.020262968 | 0.874394348 |
| A0A1U7QME6 | Elongation factor 1-beta OS=Mesocricetus auratus OX=10036 GN=Eef1b2 PE=3 SV=1 | 7.688 | 17 | 3 | 8 | 3 | 225 | 24.7 | 4.69 | 9.84 | 3 | 212.8 | 292.6 | 178 | 184.7 | 304.5 | 161.1 | 203.3 | 309.2 | 176.7 | 0.932 | 0.853707228 | 1.008486977 | 0.972599039 | 1.059818545 | 0.839743474 |
| A0A1U7QFB6 | eukaryotic translation elongation factor 1 epsilon-1 OS=Mesocricetus auratus OX=10036 GN=Eef1e1 PE=4 SV=1 | 1.464 | 5 | 1 | 1 | 1 | 174 | 19.9 | 8.15 | 3.33 | 1 | 184.3 | 0 | 0 | 140.5 | 0 | 0 | 160.4 | 0 | 0 | 0.762 | 0.859907379 | 0.87032013 | 0.926865632 | 1.141637011 | 0.930206902 |
| A0A1U7QXE7 | EF-hand domain-containing protein D2 isoform X1 OS=Mesocricetus auratus OX=10036 GN=Efhd2 PE=4 SV=1 | 10.542 | 12 | 2 | 12 | 2 | 239 | 26.7 | 5.14 | 33.85 | 2 | 89.5 | 0 | 94 | 137.4 | 0 | 132.6 | 114.3 | 0 | 137.4 | 1.472 | 0.627967189 | 1.371662125 | 0.68865403 | 0.932222222 | 0.926250825 |
| A0A1U7QT03 | ephrin-A1 OS=Mesocricetus auratus OX=10036 GN=Efna1 PE=3 SV=1 | 2.065 | 5 | 1 | 1 | 1 | 205 | 23.6 | 6.89 | 0 | 1 | 0 | 106.5 | 0 | 0 | 77.9 | 0 | 0 | 77.5 | 0 | 0.732 | 0.839929999 | 0.727699531 | 0.837480447 | 0.994865212 | 0.997269879 |
| A0A1U7Q4W1 | EH domain-containing protein 1 OS=Mesocricetus auratus OX=10036 GN=Ehd1 PE=4 SV=1 | 43.579 | 21 | 12 | 55 | 9 | 534 | 60.6 | 6.83 | 170.56 | 12 | 1519.1 | 770.9 | 1738.4 | 949 | 533.8 | 1098.1 | 1439.9 | 697.4 | 1659.3 | 0.692 | 0.243433011 | 0.942458544 | 0.860663731 | 1.471037235 | 0.309692792 |
| A0A1U7RDJ6 | EH domain-containing protein 2 OS=Mesocricetus auratus OX=10036 GN=Ehd2 PE=4 SV=1 | 141.385 | 50 | 30 | 244 | 26 | 543 | 61.1 | 6.37 | 740.57 | 30 | 4719.7 | 5202.8 | 6132 | 3962.3 | 4223.9 | 4594.2 | 5473.9 | 6786.1 | 6630.4 | 0.812 | 0.102844894 | 1.176642063 | 0.181839312 | 1.478075804 | 0.024514241 |
| A0A1U7QS99 | EH domain-containing protein 3 OS=Mesocricetus auratus OX=10036 GN=Ehd3 PE=4 SV=1 | 43.856 | 23 | 11 | 69 | 5 | 535 | 60.8 | 6.51 | 196.48 | 11 | 276.4 | 118.2 | 255.7 | 238.6 | 98 | 221.4 | 258.9 | 106.3 | 272.8 | 0.866 | 0.668055992 | 0.981085653 | 0.957838894 | 1.143369176 | 0.720672125 |
| A0A1U7Q5E9 | EH domain-containing protein 4 OS=Mesocricetus auratus OX=10036 GN=Ehd4 PE=4 SV=1 | 43.597 | 17 | 11 | 77 | 7 | 541 | 61.4 | 6.76 | 195.29 | 11 | 881.3 | 1197.3 | 542.6 | 730.7 | 892.1 | 423.4 | 786.1 | 887.1 | 549.3 | 0.829 | 0.462322427 | 0.847894094 | 0.577876996 | 1.086159711 | 0.748623171 |
| A0A1U7QHW3 | Eukaryotic translation initiation factor 4C OS=Mesocricetus auratus OX=10036 GN=Eif1ax PE=3 SV=1 | 12.424 | 26 | 3 | 11 | 3 | 144 | 16.5 | 5.24 | 30.68 | 3 | 164.1 | 154.7 | 121.9 | 138.7 | 161.9 | 114 | 146.2 | 138.5 | 89.9 | 0.935 | 0.668342506 | 0.850011346 | 0.373943635 | 0.903521466 | 0.585469409 |
| A0A1U7Q8G2 | Eukaryotic translation initiation factor 2 subunit 1 OS=Mesocricetus auratus OX=10036 GN=Eif2s1 PE=3 SV=1 | 1.363 | 3 | 1 | 1 | 1 | 315 | 36.1 | 5.14 | 0 | 1 | 0 | 0 | 56 | 0 | 0 | 82.4 | 0 | 0 | 46.6 | 1.47 | 0.805789177 | 0.832142857 | 0.903759208 | 0.565533981 | 0.729279786 |
| A0A1U7R304 | eukaryotic translation initiation factor 2 subunit 2 OS=Mesocricetus auratus OX=10036 GN=Eif2s2 PE=3 SV=1 | 11.04 | 12 | 3 | 8 | 3 | 333 | 38.2 | 5.8 | 25.97 | 3 | 64 | 84.8 | 0 | 88.9 | 129.1 | 0 | 68.5 | 74.4 | 0 | 1.455 | 0.645231166 | 0.960349462 | 0.957831214 | 0.655504587 | 0.612938448 |
| A0A1U8BRK0 | Eukaryotic translation initiation factor 3 subunit C OS=Mesocricetus auratus OX=10036 GN=LOC101843174 PE=3 SV=1 | 7.827 | 2 | 1 | 7 | 1 | 912 | 105.6 | 5.67 | 35.2 | 1 | 38.8 | 0 | 55.8 | 45.1 | 0 | 59.2 | 31.2 | 0 | 45.8 | 1.11 | 0.900686745 | 0.813953488 | 0.797447606 | 0.738255034 | 0.706655166 |
| A0A1U7QIU3 | Eukaryotic translation initiation factor 3 subunit D OS=Mesocricetus auratus OX=10036 GN=Eif3d PE=3 SV=1 | 12.804 | 8 | 3 | 9 | 3 | 548 | 63.9 | 6.05 | 21.27 | 3 | 116.1 | 0 | 163.7 | 85.2 | 0 | 190.7 | 119.5 | 0 | 176.2 | 0.925 | 0.986752191 | 1.056826305 | 0.944199884 | 1.071765132 | 0.934774873 |
| A0A1U7QI12 | Eukaryotic translation initiation factor 3 subunit E OS=Mesocricetus auratus OX=10036 GN=Eif3e PE=3 SV=1 | 8.298 | 7 | 3 | 5 | 3 | 445 | 52.2 | 6.04 | 15.69 | 3 | 51.7 | 0 | 0 | 46.3 | 0 | 0 | 51.8 | 0 | 0 | 0.896 | 0.941760355 | 1.001934236 | 0.998975209 | 1.118790497 | 0.940749629 |
| A0A1U7QQB2 | Eukaryotic translation initiation factor 3 subunit G OS=Mesocricetus auratus OX=10036 GN=Eif3g PE=3 SV=1 | 3.362 | 5 | 1 | 6 | 1 | 320 | 35.6 | 5.9 | 15.99 | 1 | 64.3 | 0 | 0 | 81.8 | 0 | 0 | 67.5 | 0 | 0 | 1.272 | 0.875024403 | 1.049766719 | 0.974265597 | 0.825183374 | 0.899476437 |
| A0A1U7Q9G9 | Eukaryotic translation initiation factor 3 subunit L OS=Mesocricetus auratus OX=10036 GN=Eif3l PE=3 SV=1 | 3.83 | 2 | 1 | 4 | 1 | 564 | 66.6 | 6.34 | 4.28 | 1 | 192.8 | 172.5 | 0 | 177.9 | 162.5 | 0 | 161.2 | 179.9 | 0 | 0.932 | 0.925664533 | 0.93375308 | 0.927860312 | 1.002056404 | 0.997829314 |
| A0A1U8BR82 | RNA helicase OS=Mesocricetus auratus OX=10036 GN=Eif4a3 PE=3 SV=1 | 22.988 | 18 | 6 | 37 | 5 | 411 | 46.8 | 6.73 | 96.83 | 6 | 258.9 | 284.7 | 180.2 | 218.2 | 258.9 | 149.2 | 227.4 | 246.7 | 166.2 | 0.843 | 0.508935434 | 0.88463664 | 0.52426089 | 1.022353505 | 0.913508647 |
| A9QXE6 | Eukaryotic Translation initiation factor 4 gamma 1 OS=Mesocricetus auratus OX=10036 GN=Eif4g1 PE=2 SV=1 | 21.859 | 6 | 7 | 18 | 7 | 1592 | 175.2 | 5.39 | 34.99 | 7 | 406.5 | 109.5 | 166.2 | 383.1 | 91.1 | 174.9 | 364.9 | 105.1 | 149.7 | 0.942 | 0.934328364 | 0.908384638 | 0.872126464 | 0.954706517 | 0.937923283 |
| A0A1U7Q4F4 | Eukaryotic translation initiation factor 5 OS=Mesocricetus auratus OX=10036 GN=Eif5 PE=3 SV=1 | 8.031 | 8 | 3 | 7 | 3 | 430 | 49.1 | 5.58 | 11.74 | 3 | 47.1 | 272.9 | 329.7 | 37.5 | 227.9 | 298.3 | 22.1 | 210.2 | 349.1 | 0.905 | 0.817497278 | 0.894874557 | 0.867714304 | 1.031399681 | 0.96402461 |
| A0A1U7RE46 | ELAV-like protein OS=Mesocricetus auratus OX=10036 GN=Elavl1 PE=3 SV=1 | 2.452 | 2 | 1 | 3 | 1 | 326 | 36.1 | 9.04 | 5.95 | 1 | 36.3 | 0 | 37.7 | 40.9 | 0 | 57.7 | 29.4 | 0 | 32.9 | 1.313 | 0.719417579 | 0.841891892 | 0.821460282 | 0.631845842 | 0.585302039 |
| A0A1U8C2Z7 | engulfment and cell motility protein 1 isoform X1 OS=Mesocricetus auratus OX=10036 GN=Elmo1 PE=4 SV=1 | 23.505 | 11 | 5 | 10 | 2 | 727 | 83.8 | 6.28 | 33.48 | 5 | 0 | 163.8 | 0 | 0 | 167.5 | 0 | 0 | 115.7 | 0 | 1.023 | 0.988156127 | 0.706349206 | 0.82348064 | 0.690746269 | 0.813175993 |
| A0A1U7QF36 | ER membrane protein complex subunit 7 OS=Mesocricetus auratus OX=10036 GN=Emc7 PE=3 SV=1 | 2.744 | 5 | 1 | 2 | 1 | 241 | 26.3 | 9.23 | 5.95 | 1 | 52.8 | 80.8 | 0 | 53.8 | 67.6 | 0 | 69.7 | 88.3 | 0 | 0.923 | 0.903338107 | 1.182634731 | 0.831697338 | 1.301482702 | 0.738103851 |
| A0A1U8CN01 | echinoderm microtubule-associated protein-like 2 isoform X4 OS=Mesocricetus auratus OX=10036 GN=Eml2 PE=3 SV=1 | 10.802 | 5 | 3 | 12 | 3 | 649 | 70.6 | 6.4 | 53.81 | 3 | 270 | 248.6 | 267.5 | 241 | 204.3 | 242.7 | 205.9 | 219.4 | 223 | 0.893 | 0.103130877 | 0.824704236 | 0.006861283 | 0.942296512 | 0.409157885 |

**Supplementary Table 1: List of differentially expressed proteins identified across the 4dpi infected and 14dpi infected**

| Accession | Description | Sum PEP Score | Coverage [%] | # Peptides | # PSMs | # Unique Peptides | # AAs | MW [kDa] | calc. pI | Score Sequest HT: Sequest HT | # Peptides (by Search Engine): Sequest HT | Mock-infected (4dpi) Replicate_R1 | Mock-infected (4dpi) Replicate_R2 | Mock-infected (4dpi) Replicate_R3 | SARS-CoV-2-infected (4dpi) Replicate_R1 | SARS-CoV-2-infected (4dpi) Replicate_R2 | SARS-CoV-2-infected (4dpi) Replicate_R3 | SARS-CoV-2-(14dpi) Replicate_R1 | SARS-CoV-2-infected (14dpi) Replicate_R2 | SARS-CoV-2-infected (14dpi) Replicate_R3 | Abundance Ratio: SARS-CoV-2-infected (4dpi) / Mock-infected (4dpi) | P-Value: SARS-CoV-2-infected (4dpi) / Mock-infected (4dpi) | Abundance Ratio: SARS-CoV-2-infected (14dpi) / Mock-infected (4dpi) | P-Value: SARS-CoV-2-infected (14dpi) / Mock-infected (4dpi) | Abundance Ratio: SARS-CoV-2-infected (14dpi) / SARS-CoV-2-infected (4dpi) | P-Value: SARS-CoV-2-infected (14dpi) / SARS-CoV-2-infected (4dpi) |
|---|---|---|---|---|---|---|---|---|---|---|---|---|---|---|---|---|---|---|---|---|---|---|---|---|---|
| A0A1U7QTU5 | Epithelial membrane protein 1 OS=Mesocricetus auratus OX=10036 GN=Emp1 PE=3 SV=1 | 1.41 | 4 | 1 | 1 | 1 | 160 | 18 | 7.97 | 1.86 | 1 | 0 | 141.2 | 0 | 0 | 122.6 | 0 | 0 | 158.9 | 0 | 0.869 | 0.925643119 | 1.125354108 | 0.93769302 | 1.296084829 | 0.865801857 |
| A0A1U7RC65 | endonuclease domain-containing 1 protein OS=Mesocricetus auratus OX=10036 GN=Endod1 PE=4 SV=1 | 6.977 | 11 | 3 | 10 | 3 | 500 | 54.9 | 7.36 | 15.08 | 3 | 39.7 | 37.7 | 53.3 | 23.4 | 31.5 | 46.6 | 34.5 | 26.9 | 42.8 | 0.827 | 0.315980258 | 0.797245601 | 0.258999012 | 1.026600985 | 0.918618151 |
| A0A1U7Q6W7 | EGF domain-specific O-linked N-acetylglucosamine transferase OS=Mesocricetus auratus OX=10036 GN=Eogt PE=3 SV=1 | 3.06 | 2 | 1 | 2 | 1 | 527 | 61.7 | 6.86 | 6.02 | 1 | 93.3 | 0 | 0 | 67.6 | 0 | 0 | 103.7 | 0 | 0 | 0.725 | 0.835419561 | 1.111468382 | 0.944186011 | 1.534023669 | 0.787292792 |
| A0A1U7QE15 | band 4.1-like protein 5 isoform X3 OS=Mesocricetus auratus OX=10036 GN=Epb41l5 PE=4 SV=1 | 3.985 | 2 | 1 | 3 | 1 | 504 | 57.8 | 9.09 | 12.97 | 1 | 105.3 | 101.1 | 55.6 | 88.1 | 110.6 | 54.6 | 83.5 | 77.9 | 46.9 | 1.019 | 0.904753009 | 0.795038168 | 0.416998909 | 0.822345045 | 0.496622086 |
| A0A1U7QV09 | bifunctional epoxide hydrolase 2 OS=Mesocricetus auratus OX=10036 GN=Ephx2 PE=4 SV=1 | 12.826 | 9 | 3 | 16 | 3 | 554 | 62.2 | 6.28 | 45.67 | 3 | 0 | 98.2 | 105.5 | 0 | 78.4 | 77 | 0 | 79.7 | 89.3 | 0.763 | 0.726901081 | 0.829651448 | 0.807102166 | 1.087516088 | 0.911680907 |
| A0A1U7QUN4 | EPM2A-interacting protein 1 OS=Mesocricetus auratus OX=10036 GN=Epm2aip1 PE=4 SV=1 | 10.87 | 9 | 3 | 6 | 3 | 606 | 69.9 | 5.72 | 25.68 | 3 | 80.8 | 35.3 | 36.7 | 87.4 | 36.9 | 37.9 | 66.9 | 25.7 | 23.1 | 1.044 | 0.89551581 | 0.757198953 | 0.580754903 | 0.713316893 | 0.518887534 |
| A0A1U7R9Q3 | epsin-1 isoform X2 OS=Mesocricetus auratus OX=10036 GN=Epn1 PE=3 SV=1 | 12.786 | 9 | 2 | 6 | 2 | 551 | 57.3 | 4.86 | 26.23 | 2 | 0 | 0 | 74.3 | 0 | 0 | 64.9 | 0 | 0 | 63.5 | 0.873 | 0.928751278 | 0.854643338 | 0.917458337 | 0.978428351 | 0.988436764 |
| A0A1U7QZH9 | Glutamyl-tRNA synthetase OS=Mesocricetus auratus OX=10036 GN=Eprs PE=3 SV=1 | 32.232 | 6 | 7 | 22 | 7 | 1511 | 169.7 | 7.09 | 66.55 | 7 | 36.3 | 31.2 | 0 | 31.7 | 35.8 | 0 | 31.4 | 31.7 | 0 | 1.002 | 1 | 0.934814815 | 0.929054825 | 0.934814815 | 0.928939435 |
| A0A1U7QC16 | eosinophil peroxidase OS=Mesocricetus auratus OX=10036 GN=Epx PE=4 SV=1 | 25.689 | 12 | 9 | 37 | 7 | 716 | 81.2 | 10.23 | 96.83 | 9 | 805.1 | 775.9 | 758.7 | 769.1 | 720.7 | 701 | 831.5 | 810.6 | 791.7 | 0.955 | 0.121257575 | 1.040218831 | 0.154069291 | 1.110918386 | 0.036727989 |
| A0A1U8CQ48 | erbin isoform X4 OS=Mesocricetus auratus OX=10036 GN=Erbin PE=3 SV=1 | 5.554 | 3 | 2 | 4 | 2 | 1383 | 154.6 | 5.41 | 11.16 | 2 | 0 | 0 | 67.6 | 0 | 0 | 56.9 | 0 | 0 | 37.3 | 0.841 | 0.90961553 | 0.551775148 | 0.720051716 | 0.655536028 | 0.789737988 |
| E7CYH3 | Proliferation associated 2G4 (Fragment) OS=Mesocricetus auratus OX=10036 GN=PA2G4 PE=2 SV=1 | 7.93 | 6 | 2 | 5 | 2 | 388 | 43 | 7.5 | 14.12 | 2 | 78 | 87.8 | 93.2 | 92.5 | 92 | 107.1 | 91.9 | 95.6 | 95.5 | 1.149 | 0.178744398 | 1.092664093 | 0.208455441 | 0.970507545 | 0.625196754 |
| A0A1U7QI78 | Enhancer of rudimentary homolog OS=Mesocricetus auratus OX=10036 GN=Erh PE=3 SV=1 | 5.106 | 24 | 2 | 13 | 2 | 104 | 12.3 | 5.92 | 34.45 | 2 | 713.1 | 822 | 713.6 | 680.6 | 716.6 | 682 | 685.4 | 766.3 | 710.6 | 0.951 | 0.255131879 | 0.961577801 | 0.548409816 | 1.039967295 | 0.376890309 |
| A0A1U7QFP2 | erlin-2 OS=Mesocricetus auratus OX=10036 GN=Erlin2 PE=3 SV=1 | 6.897 | 14 | 3 | 5 | 3 | 339 | 37.8 | 5.62 | 2.86 | 3 | 55.9 | 0 | 0 | 32.4 | 0 | 0 | 42.9 | 0 | 0 | 0.579 | 0.738735312 | 0.76744186 | 0.863173638 | 1.324074074 | 0.85534371 |
| A0A1U7QKA1 | Endoplasmic reticulum resident protein 29 OS=Mesocricetus auratus OX=10036 GN=Erp29 PE=4 SV=1 | 19.889 | 26 | 4 | 22 | 4 | 260 | 28.6 | 6.62 | 66.57 | 4 | 110 | 35.8 | 137.5 | 135.2 | 34.6 | 139.7 | 122.8 | 36 | 136.8 | 1.182 | 0.858252343 | 1.043416873 | 0.929891654 | 0.955088853 | 0.9256047 |
| A0A1U7R0Z8 | endoplasmic reticulum resident protein 44 OS=Mesocricetus auratus OX=10036 GN=Erp44 PE=4 SV=1 | 9.995 | 8 | 3 | 5 | 2 | 406 | 46.9 | 5.33 | 18.23 | 3 | 41.9 | 365.9 | 110.5 | 35.5 | 312.2 | 117 | 38.2 | 283.5 | 104 | 1.059 | 0.896173019 | 0.821338993 | 0.814958996 | 0.916074887 | 0.911735825 |
| A0A1U7Q547 | endothelial cell-selective adhesion molecule OS=Mesocricetus auratus OX=10036 GN=Esam PE=4 SV=1 | 15.19 | 15 | 4 | 9 | 4 | 394 | 42 | 9.25 | 20.85 | 4 | 0 | 0 | 45.3 | 0 | 0 | 48.2 | 0 | 0 | 56.9 | 1.064 | 0.967139168 | 1.25607064 | 0.881377991 | 1.180497925 | 0.912891268 |
| A0A1U8C186 | S-formylglutathione hydrolase OS=Mesocricetus auratus OX=10036 GN=Esd PE=3 SV=1 | 9.543 | 14 | 3 | 29 | 3 | 282 | 31.4 | 7.12 | 81.21 | 3 | 847.6 | 1362.8 | 1037.1 | 995 | 1072.2 | 1172.3 | 1249 | 1355.1 | 1603.7 | 1.174 | 0.987880495 | 1.295704388 | 0.16449999 | 1.298904152 | 0.072845271 |
| A0A1U7QS81 | extended synaptotagmin-1 OS=Mesocricetus auratus OX=10036 GN=Esyt1 PE=3 SV=1 | 15.968 | 4 | 3 | 17 | 3 | 1089 | 120.7 | 5.85 | 56.79 | 3 | 159 | 87.4 | 103 | 125.4 | 75.9 | 91.5 | 129.7 | 98.7 | 117.4 | 0.81 | 0.516468676 | 0.989696623 | 0.962903488 | 1.181010929 | 0.372306341 |
| O35588 | C11 protein (Fragment) OS=Mesocricetus auratus OX=10036 PE=2 SV=1 | 3.054 | 17 | 1 | 2 | 1 | 60 | 6.8 | 4.5 | 5.94 | 1 | 0 | 100.4 | 134.3 | 0 | 96.6 | 159.6 | 0 | 92.8 | 147.4 | 1.07 | 0.912925542 | 1.023434171 | 0.976692082 | 0.93754879 | 0.936910401 |
| A0A1U7QS16 | Electron transfer flavoprotein subunit beta OS=Mesocricetus auratus OX=10036 GN=Etfb PE=3 SV=1 | 24.678 | 29 | 5 | 21 | 5 | 255 | 27.7 | 8.1 | 63.7 | 5 | 250.3 | 202.2 | 201.5 | 263.5 | 220.3 | 205.3 | 197.1 | 196.7 | 214.9 | 1.019 | 0.648561582 | 0.930733945 | 0.455726934 | 0.883326077 | 0.260623705 |
| A0A1U7QNQ0 | Exocyst complex component 7 OS=Mesocricetus auratus OX=10036 GN=Exoc7 PE=3 SV=1 | 1.916 | 3 | 1 | 1 | 1 | 684 | 78.1 | 7.23 | 3.66 | 1 | 0 | 39.4 | 0 | 0 | 30 | 0 | 0 | 33.8 | 0 | 0.762 | 0.859315226 | 0.85786802 | 0.919402764 | 1.126666667 | 0.937084467 |
| A0A1U7RI72 | Junctional adhesion molecule 1 OS=Mesocricetus auratus OX=10036 GN=F11r PE=3 SV=1 | 10.027 | 10 | 3 | 5 | 3 | 299 | 32.4 | 7.64 | 15.26 | 3 | 44.9 | 0 | 0 | 25.2 | 0 | 0 | 58.7 | 0 | 0 | 0.56 | 0.726402533 | 1.307349666 | 0.861550224 | 2.329365079 | 0.639785807 |
| A0A1U7QFC8 | Prothrombin OS=Mesocricetus auratus OX=10036 GN=F2 PE=3 SV=1 | 8.356 | 4 | 2 | 6 | 2 | 618 | 70 | 6.13 | 21.34 | 2 | 43.6 | 0 | 0 | 45.8 | 0 | 0 | 32.1 | 0 | 0 | 1.051 | 0.973917057 | 0.736238532 | 0.843043421 | 0.700873362 | 0.819870812 |
| A0A1U7QG37 | fatty acid-binding protein, epidermal isoform X1 OS=Mesocricetus auratus OX=10036 GN=Fabp5 PE=3 SV=1 | 16.116 | 33 | 4 | 8 | 4 | 135 | 15.1 | 7.01 | 18.27 | 4 | 42.5 | 102.1 | 33.2 | 53.4 | 120.3 | 40.3 | 52.8 | 115.1 | 51.9 | 1.178 | 0.732373628 | 1.236220472 | 0.665631389 | 1.027102804 | 0.955389266 |
| A0A1U7R0A8 | FAS-associated factor 2 isoform X2 OS=Mesocricetus auratus OX=10036 GN=Faf2 PE=4 SV=1 | 2.878 | 2 | 1 | 5 | 1 | 426 | 50.2 | 5.27 | 8.45 | 1 | 70.3 | 69.1 | 0 | 47.7 | 42.5 | 0 | 60 | 54.9 | 0 | 0.646 | 0.590755902 | 0.824246772 | 0.80032195 | 1.27383592 | 0.753997469 |
| A0A1U8CLP4 | redox-regulatory protein FAM213A isoform X2 OS=Mesocricetus auratus OX=10036 GN=Fam213a PE=4 SV=1 | 4.132 | 6 | 1 | 5 | 1 | 218 | 24 | 9.13 | 14.63 | 1 | 39.7 | 0 | 0 | 32.6 | 0 | 0 | 44.3 | 0 | 0 | 0.822 | 0.896990128 | 1.115869018 | 0.942117363 | 1.358895706 | 0.842817862 |
| A0A1U7QJY3 | Ubiquitinyl hydrolase 1 OS=Mesocricetus auratus OX=10036 GN=Mindy2 PE=3 SV=1 | 2.774 | 2 | 1 | 3 | 1 | 600 | 65.5 | 4.61 | 8.51 | 1 | 0 | 537.9 | 0 | 0 | 353.7 | 0 | 0 | 499.6 | 0 | 0.658 | 0.791089096 | 0.928797174 | 0.960906785 | 1.412496466 | 0.824545293 |
| A0A1U7QWR4 | protein FAM65A OS=Mesocricetus auratus OX=10036 GN=Ripor1 PE=3 SV=1 | 4.144 | 1 | 1 | 1 | 1 | 1246 | 135.1 | 6.04 | 3.65 | 1 | 161 | 0 | 0 | 205.5 | 0 | 0 | 117.7 | 0 | 0 | 1.277 | 0.87337213 | 0.731055901 | 0.839669648 | 0.572749392 | 0.734126255 |
| A0A1U7RHE0 | Phenylalanine--tRNA ligase OS=Mesocricetus auratus OX=10036 GN=Farsa PE=3 SV=1 | 4.215 | 3 | 1 | 6 | 1 | 508 | 57.5 | 7.68 | 27.69 | 1 | 120.2 | 103.4 | 129 | 112.6 | 102.5 | 130.4 | 134.7 | 100.4 | 108.6 | 0.936 | 0.841461624 | 0.974758934 | 0.828860951 | 0.994790159 | 0.965959362 |
| A0A1U7QAS8 | 3-hydroxyacyl-[acyl-carrier-protein] dehydratase OS=Mesocricetus auratus OX=10036 GN=Fasn PE=4 SV=1 | 104.773 | 16 | 26 | 94 | 26 | 2500 | 271.4 | 6.29 | 315.21 | 26 | 580.5 | 865 | 1211.6 | 487.6 | 725.7 | 965.9 | 486.9 | 706.1 | 1185.6 | 0.839 | 0.527313758 | 0.895186482 | 0.753262855 | 1.091501468 | 0.80395272 |
| A0A1U7QDP0 | 40S ribosomal protein S30 OS=Mesocricetus auratus OX=10036 GN=Fau PE=3 SV=1 | 8.285 | 8 | 2 | 7 | 2 | 133 | 14.4 | 10.17 | 14.14 | 2 | 50.7 | 119 | 75.3 | 41.3 | 99.5 | 76.7 | 40.5 | 104.1 | 66.7 | 0.975 | 0.744370434 | 0.86244898 | 0.700841617 | 0.971494253 | 0.938226487 |
| A0A1U7QUH3 | rRNA 2'-O-methyltransferase fibrillarin OS=Mesocricetus auratus OX=10036 GN=Fbl PE=3 SV=1 | 6.047 | 7 | 2 | 5 | 2 | 328 | 34.3 | 10.24 | 9.93 | 2 | 144.9 | 66 | 102.9 | 151.3 | 78.7 | 98.4 | 176.2 | 74.2 | 118.4 | 1.013 | 0.884549655 | 1.175270873 | 0.650413698 | 1.123020706 | 0.733370298 |
| A0A1U8BK44 | Fibulin-5 OS=Mesocricetus auratus OX=10036 GN=Fbln5 PE=3 SV=1 | 15.757 | 19 | 5 | 21 | 5 | 447 | 50.1 | 4.74 | 85.62 | 5 | 43 | 56.9 | 33.1 | 38.5 | 48.8 | 40.4 | 32.5 | 30 | 24.3 | 1.076 | 0.831960908 | 0.652631579 | 0.144486658 | 0.679718089 | 0.029691918 |
| A0A1U8BKX0 | fibrillin-1 isoform X1 OS=Mesocricetus auratus OX=10036 GN=Fbn1 PE=3 SV=1 | 53.129 | 6 | 13 | 58 | 11 | 2871 | 311.6 | 4.92 | 188.85 | 13 | 1121.5 | 947.6 | 1546.9 | 800.7 | 706.9 | 1156.3 | 1004 | 877.5 | 1426.6 | 0.714 | 0.234878283 | 0.914850664 | 0.695054802 | 1.241825894 | 0.376663991 |
| A0A1U7Q9X5 | fibrillin-2 OS=Mesocricetus auratus OX=10036 GN=Fbn2 PE=3 SV=1 | 4.978 | 1 | 3 | 8 | 1 | 2906 | 312.9 | 4.83 | 10.54 | 3 | 69.9 | 0 | 301.5 | 129 | 0 | 510.2 | 55.7 | 0 | 266.1 | 1.768 | 0.648347627 | 0.866451265 | 0.898775255 | 0.503441802 | 0.584119387 |
| A0A1U8CJT0 | IgG receptor FcRn large subunit p51 OS=Mesocricetus auratus OX=10036 GN=Fcgrt PE=4 SV=1 | 3.492 | 4 | 1 | 5 | 1 | 368 | 40.2 | 5.19 | 6.54 | 1 | 119.5 | 84.8 | 30.8 | 101 | 73.5 | 22 | 118.1 | 89.3 | 40.3 | 0.788 | 0.729584545 | 1.053594215 | 0.908741313 | 1.260559796 | 0.62656403 |
| A0A1U7R208 | fermitin family homolog 2 isoform X5 OS=Mesocricetus auratus OX=10036 GN=Fermt2 PE=3 SV=1 | 75.497 | 20 | 10 | 103 | 10 | 687 | 78.7 | 6.74 | 357.66 | 10 | 637.2 | 636.3 | 663.7 | 487.5 | 494 | 464.8 | 581 | 575.5 | 663.6 | 0.758 | 0.000204013 | 0.939551931 | 0.302762171 | 1.258452603 | 0.038711478 |
| A0A1U7QP51 | fetuin-B OS=Mesocricetus auratus OX=10036 GN=Fetub PE=4 SV=1 | 50.925 | 17 | 7 | 74 | 7 | 421 | 45.3 | 6.89 | 268.64 | 7 | 730.1 | 655.4 | 1009.4 | 1392.9 | 1335 | 1829.5 | 519.7 | 382.8 | 757 | 1.812 | 0.023681275 | 0.692930811 | 0.185470837 | 0.364133058 | 0.009524268 |
| A0A1U7QP17 | Fibrinogen alpha chain OS=Mesocricetus auratus OX=10036 GN=Fga PE=4 SV=1 | 141.183 | 37 | 29 | 144 | 29 | 781 | 86.5 | 6.15 | 456.71 | 29 | 2133.3 | 1920.4 | 2111.5 | 6281.2 | 7520 | 5807.6 | 1927.5 | 2149.5 | 2314.3 | 2.944 | 0.011639817 | 1.036673587 | 0.601837488 | 0.325940394 | 0.01038507 |
| A0A1U7QCL5 | Fibrinogen beta chain OS=Mesocricetus auratus OX=10036 GN=Fgb PE=4 SV=1 | 113.161 | 50 | 20 | 120 | 20 | 479 | 54.4 | 7.08 | 454.38 | 20 | 703.4 | 817.1 | 760.5 | 2201.1 | 2600.1 | 2446.4 | 1195.7 | 1419.9 | 1225.8 | 3.217 | 0.002864564 | 1.684085927 | 0.00807511 | 0.530023732 | 0.002514583 |
| A0A1U7QZA0 | fibrinogen gamma chain isoform X2 OS=Mesocricetus auratus OX=10036 GN=Fgg PE=4 SV=1 | 65.038 | 37 | 15 | 91 | 15 | 436 | 49.5 | 5.9 | 221.39 | 15 | 1833.1 | 1636.6 | 1673.5 | 8151.1 | 6374.2 | 5125.5 | 1566.8 | 1187.6 | 1340 | 3.765 | 0.030901369 | 0.796080261 | 0.066268783 | 0.208357929 | 0.025946539 |
| A0A1U7R004 | Fumarate hydratase, mitochondrial OS=Mesocricetus auratus OX=10036 GN=Fh PE=3 SV=1 | 2.013 | 2 | 1 | 1 | 1 | 507 | 54.3 | 8.79 | 2.97 | 1 | 113.2 | 0 | 0 | 106.5 | 0 | 0 | 100.7 | 0 | 0 | 0.941 | 0.967688807 | 0.889575972 | 0.938260869 | 0.945539906 | 0.970336654 |
| A0A1U7R2T5 | four and a half LIM domains protein 1 isoform X4 OS=Mesocricetus auratus OX=10036 GN=Fhl1 PE=4 SV=1 | 54.031 | 40 | 10 | 80 | 10 | 296 | 33.5 | 8.41 | 294.62 | 10 | 2653.7 | 2341.1 | 2703.4 | 2026.8 | 1870.2 | 2136.7 | 2456 | 2246.5 | 2561.9 | 0.799 | 0.019943239 | 0.943649165 | 0.381400582 | 1.203971029 | 0.028677102 |
| A0A1U7QP81 | FH1/FH2 domain-containing protein 1 isoform X1 OS=Mesocricetus auratus OX=10036 GN=Fhod1 PE=4 SV=1 | 4.23 | 2 | 2 | 9 | 2 | 1182 | 128.4 | 6.21 | 26.16 | 2 | 110.6 | 102.5 | 158.4 | 99.7 | 86.5 | 149.4 | 82.5 | 72.6 | 137.8 | 0.943 | 0.668284788 | 0.788425303 | 0.384015471 | 0.872765197 | 0.636847678 |
| A0A1U8C7W9 | Peptidylprolyl isomerase OS=Mesocricetus auratus OX=10036 GN=Fkbp2 PE=4 SV=1 | 18.399 | 26 | 4 | 27 | 4 | 140 | 15.4 | 8.88 | 80.4 | 4 | 866.2 | 946.9 | 621 | 670.8 | 752.6 | 524 | 769.6 | 879.1 | 587.2 | 0.774 | 0.252038364 | 0.9185736 | 0.638072145 | 1.148146246 | 0.427191458 |
| A0A1U7Q976 | Peptidylprolyl isomerase OS=Mesocricetus auratus OX=10036 GN=Fkbp3 PE=4 SV=1 | 12.002 | 16 | 4 | 14 | 4 | 223 | 25.1 | 9.28 | 42.28 | 4 | 304.1 | 331.5 | 232.6 | 240.7 | 267.6 | 210 | 254.3 | 248.6 | 191.1 | 0.807 | 0.232053631 | 0.799354987 | 0.188652311 | 0.966170124 | 0.772820732 |
| A0A1U7R9P2 | Peptidylprolyl isomerase OS=Mesocricetus auratus OX=10036 GN=Fkbp4 PE=4 SV=1 | 8.447 | 7 | 2 | 4 | 2 | 458 | 51.4 | 5.49 | 11.94 | 2 | 0 | 0 | 190.8 | 0 | 0 | 215.1 | 0 | 0 | 184 | 1.127 | 0.936764072 | 0.964360587 | 0.980763775 | 0.855416086 | 0.917924729 |
| A0A1U8CZP3 | filamin-A OS=Mesocricetus auratus OX=10036 GN=Flna PE=3 SV=1 | 198.664 | 39 | 40 | 217 | 4 | 1076 | 115.6 | 6.8 | 614.53 | 40 | 114.5 | 152.2 | 0 | 105 | 132.1 | 0 | 77.8 | 128.7 | 0 | 0.892 | 0.879401164 | 0.774278215 | 0.751983833 | 0.870940531 | 0.861902159 |
| A0A1U7R1A1 | filamin-B isoform X3 OS=Mesocricetus auratus OX=10036 GN=Flnb PE=3 SV=1 | 54.86 | 5 | 14 | 48 | 4 | 2578 | 275 | 5.81 | 104.98 | 14 | 0 | 0 | 38.7 | 0 | 0 | 37.9 | 0 | 0 | 29 | 0.978 | 0.98892401 | 0.749354005 | 0.851543936 | 0.765171504 | 0.861720128 |
| A0A1U7QRB7 | Dimethylaniline monooxygenase [N-oxide-forming] OS=Mesocricetus auratus OX=10036 GN=Fmo1 PE=3 SV=1 | 13.894 | 12 | 6 | 11 | 5 | 532 | 59.7 | 8.73 | 24.36 | 6 | 318.3 | 290.2 | 255.3 | 295.7 | 327.7 | 228.4 | 254.5 | 236.3 | 233.5 | 1.078 | 0.914241392 | 0.838504283 | 0.11251926 | 0.850316976 | 0.281444856 |
| A0A1U7QFV1 | Dimethylaniline monooxygenase [N-oxide-forming] OS=Mesocricetus auratus OX=10036 GN=Fmo2 PE=3 SV=1 | 109.363 | 35 | 22 | 150 | 21 | 535 | 60.9 | 8.56 | 424.12 | 22 | 2827.4 | 3669.8 | 4106.7 | 2502 | 2981.4 | 3730.3 | 4268.7 | 4703.3 | 4784.4 | 0.812 | 0.421977 | 1.297296278 | 0.09123539 | 1.493037542 | 0.035201669 |
| A0A1U7QEF6 | Fibronectin OS=Mesocricetus auratus OX=10036 GN=Fn1 PE=4 SV=1 | 83.802 | 13 | 20 | 89 | 20 | 2385 | 262.6 | 5.9 | 288.34 | 20 | 433 | 437.7 | 486.4 | 658.1 | 413 | 656.6 | 406.2 | 399.6 | 467.5 | 1.236 | 0.266317287 | 0.938250682 | 0.370743297 | 0.736991376 | 0.198347793 |
| A0A1U7R5R2 | Fascin OS=Mesocricetus auratus OX=10036 GN=Fscn1 PE=3 SV=1 | 12.431 | 12 | 5 | 13 | 5 | 493 | 54.5 | 6.93 | 29.84 | 5 | 470.1 | 600.6 | 67.6 | 408 | 531.6 | 57.3 | 427.7 | 486 | 54 | 0.868 | 0.836807621 | 0.850127383 | 0.800187044 | 0.970709199 | 0.962817508 |
| A0A1U7QB70 | Ferritin OS=Mesocricetus auratus OX=10036 GN=Fth1 PE=3 SV=1 | 17.322 | 35 | 5 | 24 | 5 | 186 | 21.5 | 6.3 | 68.36 | 5 | 231.5 | 170.4 | 270 | 561.6 | 545.8 | 640.5 | 405.8 | 394.6 | 477.9 | 2.426 | 0.000960671 | 1.902515255 | 0.006801313 | 0.731334745 | 0.016689629 |
| A0A1U7R8X3 | far upstream element-binding protein 1 isoform X10 OS=Mesocricetus auratus OX=10036 GN=Fubp1 PE=4 SV=1 | 7.96 | 2 | 3 | 14 | 1 | 655 | 68.8 | 7.93 | 16.35 | 3 | 0 | 0 | 42 | 0 | 0 | 34.9 | 0 | 0 | 32 | 0.831 | 0.903028104 | 0.761904762 | 0.859625523 | 0.916905444 | 0.954122126 |
| A0A1U7Q6P8 | RNA-binding protein FUS isoform X1 OS=Mesocricetus auratus OX=10036 GN=Fus PE=3 SV=1 | 33.209 | 8 | 4 | 39 | 3 | 517 | 52.6 | 9.36 | 159.11 | 4 | 406.4 | 324.3 | 218 | 507.2 | 375.5 | 283.6 | 349.7 | 238.9 | 190.3 | 1.301 | 0.441710727 | 0.821018235 | 0.47720484 | 0.667838464 | 0.189446794 |
| A0A1U7QK87 | ras GTPase-activating protein-binding protein 1 OS=Mesocricetus auratus OX=10036 GN=G3bp1 PE=4 SV=1 | 1.862 | 2 | 1 | 3 | 1 | 465 | 51.8 | 5.59 | 6.31 | 1 | 0 | 39 | 0 | 0 | 40.5 | 0 | 0 | 32.8 | 0 | 1.039 | 0.979995802 | 0.841025641 | 0.909194586 | 0.809876543 | 0.889983724 |
| A0A1U7R6A3 | Glucose-6-phosphate 1-dehydrogenase OS=Mesocricetus auratus OX=10036 GN=G6pd PE=3 SV=1 | 38.734 | 21 | 9 | 43 | 9 | 515 | 59.3 | 6.43 | 123.29 | 9 | 1084.6 | 1183.3 | 1034.2 | 1182.6 | 1260.4 | 1113.4 | 1008.8 | 1112.5 | 1022.6 | 1.09 | 0.237012152 | 0.952091094 | 0.39258022 | 0.884011922 | 0.066038471 |
| A0A1U7QJP2 | Aldose 1-epimerase OS=Mesocricetus auratus OX=10036 GN=Galm PE=3 SV=1 | 32.582 | 25 | 6 | 33 | 6 | 342 | 37.8 | 6.35 | 109.75 | 6 | 649.5 | 760.7 | 551 | 458.3 | 536.5 | 417.2 | 660.1 | 776.3 | 525.1 | 0.705 | 0.074072257 | 1.000152968 | 0.999208201 | 1.389164306 | 0.111224202 |
| A0A1U7Q657 | neutral alpha-glucosidase AB isoform X2 OS=Mesocricetus auratus OX=10036 GN=Ganab PE=3 SV=1 | 41.574 | 15 | 11 | 52 | 11 | 944 | 107.1 | 6.15 | 176.28 | 11 | 832.4 | 790.5 | 1004.8 | 770.4 | 839.2 | 932.7 | 689.6 | 701.4 | 909.9 | 1.062 | 0.743802847 | 0.875632683 | 0.324942701 | 0.905046611 | 0.408240345 |
| A0A1U8BZW0 | Trifunctional purine biosynthetic protein adenosine-3 OS=Mesocricetus auratus OX=10036 GN=Gart PE=3 SV=1 | 4.749 | 2 | 1 | 2 | 1 | 1008 | 107.3 | 6.79 | 7.72 | 1 | 0 | 0 | 49.5 | 0 | 0 | 43.8 | 0 | 0 | 30 | 0.886 | 0.935479809 | 0.606060606 | 0.7565309 | 0.684931507 | 0.809321641 |
| Q6IU03 | Glycine amidinotransferase, mitochondrial (Fragment) OS=Mesocricetus auratus OX=10036 PE=2 SV=1 | 10.606 | 23 | 3 | 15 | 3 | 239 | 27.7 | 7.23 | 42.95 | 3 | 417.4 | 502.4 | 320 | 274.5 | 323.6 | 222.8 | 411.2 | 441.3 | 314.1 | 0.665 | 0.099851324 | 0.940958219 | 0.728873562 | 1.421123158 | 0.079663657 |
| A0A1U8BMS0 | Gc-globulin OS=Mesocricetus auratus OX=10036 GN=Gc PE=4 SV=1 | 57.45 | 34 | 14 | 54 | 14 | 486 | 54.4 | 5.47 | 170.79 | 14 | 574.3 | 645.9 | 346.2 | 1067.1 | 1230.4 | 872 | 516.3 | 681.9 | 468.9 | 1.905 | 0.018356584 | 1.064287538 | 0.779026231 | 0.525982016 | 0.021148979 |
| A0A1U7QH02 | Gamma-ECS OS=Mesocricetus auratus OX=10036 GN=Gclc PE=3 SV=1 | 1.708 | 2 | 1 | 1 | 1 | 637 | 72.8 | 5.83 | 0 | 1 | 47.3 | 0 | 0 | 43.8 | 0 | 0 | 44 | 0 | 0 | 0.927 | 0.959319763 | 0.930232558 | 0.961720858 | 1.00456621 | 0.997583932 |
| A0A1U7Q4Q9 | Guanine deaminase OS=Mesocricetus auratus OX=10036 GN=Gda PE=3 SV=1 | 36.074 | 24 | 10 | 46 | 10 | 454 | 51 | 5.62 | 117.05 | 10 | 651.7 | 330 | 722.1 | 789.3 | 409.9 | 840.5 | 685.9 | 339.5 | 843 | 1.202 | 0.571494302 | 1.096607583 | 0.789329261 | 0.916017061 | 0.790967868 |
| A0A1U7QDS7 | Rab GDP dissociation inhibitor OS=Mesocricetus auratus OX=10036 GN=Gdi2 PE=3 SV=1 | 75.061 | 34 | 13 | 105 | 10 | 445 | 50.5 | 6.25 | 314.6 | 13 | 1235.1 | 1254.7 | 1013.4 | 1106 | 1070.4 | 874.8 | 1134.8 | 1150.6 | 920.5 | 0.895 | 0.227192436 | 0.915134734 | 0.407691262 | 1.050701363 | 0.643895721 |

**Supplementary Table 1: List of differentially expressed proteins identified across the 4dpi infected and 14dpi infected**

| Accession | Description | Sum PEP Score | Coverage [%] | # Peptides | # PSMs | # Unique Peptides | # AAs | MW [kDa] | calc. pI | Score Sequest HT: Sequest HT | # Peptides (by Search Engine): Sequest HT | Mock-infected (4dpi) Replicate_R1 | Mock-infected (4dpi) Replicate_R2 | Mock-infected (4dpi) Replicate_R3 | SARS-CoV-2-infected (4dpi) Replicate_R1 | SARS-CoV-2-infected (4dpi) Replicate_R2 | SARS-CoV-2-infected (4dpi) Replicate_R3 | SARS-CoV-2-(14dpi) Replicate_R1 | SARS-CoV-2-infected (14dpi) Replicate_R2 | SARS-CoV-2-infected (14dpi) Replicate_R3 | Abundance Ratio: SARS-CoV-2-infected (4dpi) / Mock-infected (4dpi) | P-Value: SARS-CoV-2-infected (4dpi) / Mock-infected (4dpi) | Abundance Ratio: SARS-CoV-2-infected (14dpi) / Mock-infected (4dpi) | P-Value: SARS-CoV-2-infected (14dpi) / Mock-infected (4dpi) | Abundance Ratio: SARS-CoV-2-infected (14dpi) / SARS-CoV-2-infected (4dpi) | P-Value: SARS-CoV-2-infected (14dpi) / SARS-CoV-2-infected (4dpi) |
|---|---|---|---|---|---|---|---|---|---|---|---|---|---|---|---|---|---|---|---|---|---|---|---|---|---|
| A0A1U7QQ75 | glycerophosphodiester phosphodiesterase domain-containing protein 1 isoform X1 OS=Mesocricetus auratus OX=10036 GN=Gdpd1 PE=3 SV=1 | 2.276 | 3 | 1 | 1 | 1 | 314 | 35.9 | 8.15 | 0 | 1 | 0 | 0 | 67.9 | 0 | 0 | 171.3 | 0 | 0 | 52.9 | 2.524 | 0.619167392 | 0.779086892 | 0.870599598 | 0.308814945 | 0.56724538 |
| A0A1U7R342 | Golgi to ER traffic protein 4 homolog isoform X1 OS=Mesocricetus auratus OX=10036 GN=Get4 PE=3 SV=1 | 2.915 | 5 | 1 | 2 | 1 | 325 | 36.4 | 5.41 | 6.36 | 1 | 35.7 | 0 | 41.1 | 44.1 | 0 | 41.8 | 32 | 0 | 39.1 | 1.12 | 0.882680642 | 0.92578125 | 0.91940064 | 0.827706636 | 0.805421644 |
| A0A1U7R479 | GTPase IMAP family member 4 isoform X2 OS=Mesocricetus auratus OX=10036 GN=LOC101830572 PE=3 SV=1 | 14.331 | 13 | 4 | 17 | 4 | 327 | 37.7 | 6.58 | 48.76 | 4 | 595.1 | 415 | 289.1 | 490.5 | 341.9 | 236.7 | 469.4 | 347.5 | 252.6 | 0.824 | 0.54363052 | 0.823198892 | 0.524183327 | 1.000374146 | 0.998967803 |
| A0A1U8CPX1 | GTPase IMAP family member 7 OS=Mesocricetus auratus OX=10036 GN=LOC101835218 PE=3 SV=1 | 1.496 | 3 | 1 | 1 | 1 | 292 | 33.6 | 7.06 | 2.7 | 1 | 0 | 188.9 | 0 | 0 | 124.6 | 0 | 0 | 160.9 | 0 | 0.66 | 0.792459613 | 0.851773425 | 0.915723796 | 1.291332263 | 0.867612973 |
| A0A1U7QQ12 | Glyoxalase I OS=Mesocricetus auratus OX=10036 GN=Glo1 PE=3 SV=1 | 26.295 | 49 | 8 | 34 | 8 | 184 | 20.9 | 5.31 | 94.03 | 8 | 1001 | 1127.9 | 1229.9 | 1132.7 | 863.5 | 942 | 1156.3 | 975.1 | 1150 | 0.929 | 0.250400306 | 0.976956056 | 0.78631591 | 1.116806208 | 0.319532797 |
| A0A1U7Q435 | Glyoxalase domain-containing protein 4 OS=Mesocricetus auratus OX=10036 GN=Glod4 PE=4 SV=1 | 7.872 | 8 | 2 | 7 | 2 | 298 | 33.3 | 5.47 | 18.91 | 2 | 59.8 | 116.8 | 146.5 | 81 | 107.1 | 137.2 | 62.9 | 117 | 162.9 | 0.937 | 0.981971822 | 1.060971835 | 0.872976624 | 1.053796496 | 0.871030078 |
| A0A1U7QE94 | glutaredoxin-3 OS=Mesocricetus auratus OX=10036 GN=Glrx3 PE=4 SV=1 | 3.065 | 3 | 1 | 2 | 1 | 337 | 37.8 | 5.72 | 4.1 | 1 | 0 | 57.4 | 38.2 | 0 | 57.1 | 51.3 | 0 | 44.4 | 31.3 | 1.156 | 0.871671284 | 0.791841004 | 0.77295143 | 0.698339483 | 0.654586357 |
| A0A1U8CAE7 | ceramide-1-phosphate transfer protein OS=Mesocricetus auratus OX=10036 GN=Cptp PE=4 SV=1 | 2.524 | 3 | 1 | 4 | 1 | 216 | 24.7 | 6.64 | 10.77 | 1 | 99.4 | 111.8 | 190.2 | 87.1 | 91.6 | 165.3 | 93.2 | 100.4 | 158.1 | 0.819 | 0.642141664 | 0.876183358 | 0.663554402 | 1.022383721 | 0.94122836 |
| A0A1U7QZ08 | Glutamate dehydrogenase (NAD(P)(+)) OS=Mesocricetus auratus OX=10036 GN=Glud1 PE=3 SV=1 | 45.466 | 17 | 7 | 53 | 7 | 558 | 61.3 | 8 | 171.66 | 7 | 1149.6 | 1094.2 | 978.9 | 994 | 925.6 | 851.1 | 1087 | 980.4 | 934.7 | 0.865 | 0.083961374 | 0.931548081 | 0.338233382 | 1.083516801 | 0.276191083 |
| A0A1U7REQ2 | GMP reductase OS=Mesocricetus auratus OX=10036 GN=Gmpr2 PE=3 SV=1 | 1.85 | 2 | 1 | 1 | 1 | 347 | 37.9 | 7.27 | 0 | 1 | 0 | 79 | 0 | 0 | 73 | 0 | 0 | 91.6 | 0 | 0.924 | 0.958207321 | 1.159493671 | 0.922154562 | 1.254794521 | 0.88188711 |
| A0A1U7QNV6 | guanine nucleotide-binding protein subunit alpha-13 OS=Mesocricetus auratus OX=10036 GN=Gna13 PE=3 SV=1 | 27.158 | 15 | 5 | 28 | 4 | 377 | 44 | 8.21 | 90.48 | 5 | 284.2 | 463.3 | 294.8 | 275.5 | 406.5 | 250.5 | 247 | 434.3 | 276.5 | 0.902 | 0.65411653 | 0.918929291 | 0.748917352 | 1.027131367 | 0.916760373 |
| A0A1U7R767 | guanine nucleotide-binding protein G(I) subunit alpha-1 OS=Mesocricetus auratus OX=10036 GN=Gnai1 PE=3 SV=1 | 23.542 | 19 | 5 | 14 | 2 | 354 | 40.3 | 5.97 | 44.97 | 5 | 35.8 | 46.8 | 42 | 24.6 | 37 | 39 | 37 | 47.7 | 39.9 | 0.726 | 0.228113704 | 1 | 1 | 1.238568588 | 0.228407793 |
| A0A1U7QAT2 | guanine nucleotide-binding protein G(I) subunit alpha-2 OS=Mesocricetus auratus OX=10036 GN=Gnai2 PE=3 SV=1 | 42.108 | 30 | 9 | 35 | 6 | 355 | 40.5 | 5.45 | 94.62 | 9 | 1064.6 | 1268.6 | 490.2 | 975 | 1110.1 | 703.3 | 1002.4 | 1224.6 | 922 | 1.056 | 0.967285696 | 1.115321952 | 0.697749704 | 1.129321475 | 0.470914793 |
| A0A1U7QZZ8 | guanine nucleotide-binding protein G(K) subunit alpha OS=Mesocricetus auratus OX=10036 GN=Gnai3 PE=3 SV=1 | 25.576 | 14 | 5 | 18 | 2 | 354 | 40.5 | 5.69 | 60.4 | 5 | 136.5 | 61.8 | 70.5 | 102.4 | 48.3 | 54.6 | 105.5 | 68.7 | 52.2 | 0.75 | 0.511208754 | 0.842261905 | 0.648011689 | 1.102776425 | 0.777309834 |
| A0A1U7QHQ1 | guanine nucleotide-binding protein G(O) subunit alpha isoform X2 OS=Mesocricetus auratus OX=10036 GN=Gnao1 PE=3 SV=1 | 34.55 | 19 | 6 | 18 | 4 | 354 | 40 | 5.87 | 54.68 | 6 | 113.6 | 312.9 | 304.1 | 67.4 | 213.6 | 231.5 | 88.7 | 248.1 | 241.4 | 0.761 | 0.433965756 | 0.791404325 | 0.576332707 | 1.128195122 | 0.780736783 |
| A0A1U7Q621 | guanine nucleotide-binding protein G(Q) subunit alpha OS=Mesocricetus auratus OX=10036 GN=LOC101825355 PE=3 SV=1 | 17.17 | 11 | 3 | 11 | 2 | 359 | 42.1 | 5.68 | 28.82 | 3 | 259.3 | 215 | 257.1 | 144.8 | 151 | 170.9 | 125.9 | 140.3 | 148.7 | 0.558 | 0.01164334 | 0.567268253 | 0.008436942 | 0.889007928 | 0.171269905 |
| A0A1U7R3M4 | Guanine nucleotide-binding protein subunit gamma OS=Mesocricetus auratus OX=10036 GN=Gng10 PE=3 SV=1 | 5.18 | 46 | 2 | 2 | 2 | 68 | 7.2 | 7.85 | 2.28 | 2 | 23.4 | 0 | 39 | 33.6 | 0 | 55.9 | 32.4 | 0 | 38.5 | 1.434 | 0.674687351 | 1.136217949 | 0.871768775 | 0.792178771 | 0.775137159 |
| A0A1U7Q2H8 | Glucosamine-6-phosphate isomerase OS=Mesocricetus auratus OX=10036 GN=Gnpda1 PE=3 SV=1 | 6.133 | 20 | 2 | 5 | 2 | 289 | 32.6 | 6.92 | 12.71 | 2 | 50.3 | 0 | 0 | 43.3 | 0 | 0 | 39.6 | 0 | 0 | 0.86 | 0.921187535 | 0.787276342 | 0.875791683 | 0.914549654 | 0.95276957 |
| A0A1U7R0U8 | Aspartate aminotransferase OS=Mesocricetus auratus OX=10036 GN=Got2 PE=3 SV=1 | 9.959 | 7 | 3 | 10 | 3 | 430 | 47.4 | 8.98 | 21.75 | 3 | 256.2 | 479.1 | 412.1 | 240.3 | 429.4 | 366 | 241.4 | 462.8 | 346.7 | 0.896 | 0.688996501 | 0.91589681 | 0.744029404 | 1.014676064 | 0.955227462 |
| A0A1U7QJ81 | Glycerol-3-phosphate dehydrogenase [NAD(+)] OS=Mesocricetus auratus OX=10036 GN=Gpd1l PE=3 SV=1 | 4.925 | 4 | 1 | 4 | 1 | 351 | 38.2 | 6.77 | 14.55 | 1 | 170.3 | 82.9 | 101.3 | 135.1 | 74.3 | 73.3 | 170.4 | 71.2 | 110.3 | 0.793 | 0.517427892 | 0.992665726 | 0.983444671 | 1.244782455 | 0.553373814 |
| A0A1U8BKK5 | Glycerol-3-phosphate dehydrogenase OS=Mesocricetus auratus OX=10036 GN=Gpd2 PE=3 SV=1 | 14.753 | 8 | 6 | 23 | 6 | 727 | 80.7 | 7.64 | 38.36 | 6 | 408 | 569.8 | 394.2 | 275.8 | 380.1 | 263.7 | 337 | 487.8 | 322.5 | 0.676 | 0.099589408 | 0.83622449 | 0.387500656 | 1.247607656 | 0.311503793 |
| A0A1U7QYI9 | Glucose-6-phosphate isomerase OS=Mesocricetus auratus OX=10036 GN=Gpi PE=3 SV=1 | 28.003 | 15 | 8 | 38 | 8 | 558 | 62.6 | 7.75 | 127.56 | 8 | 397.5 | 333.3 | 265 | 447.7 | 348.9 | 301.7 | 363.7 | 337.6 | 278.9 | 1.047 | 0.585173975 | 0.984334204 | 0.915788403 | 0.892470181 | 0.48328194 |
| A0A1U7R6M4 | transmembrane glycoprotein NMB OS=Mesocricetus auratus OX=10036 GN=Gpnmb PE=4 SV=1 | 1.449 | 1 | 1 | 1 | 1 | 612 | 67 | 7.14 | 0 | 1 | 58.9 | 0 | 0 | 51.1 | 0 | 0 | 23.7 | 0 | 0 | 0.867 | 0.92522497 | 0.40237691 | 0.622882889 | 0.463796477 | 0.661911701 |
| A0A1U7Q8K6 | retinoic acid-induced protein 3 OS=Mesocricetus auratus OX=10036 GN=Gprc5a PE=4 SV=1 | 48.26 | 23 | 5 | 48 | 5 | 354 | 39.6 | 6.87 | 182.16 | 5 | 587.6 | 843.7 | 676.3 | 339.5 | 520.2 | 416.8 | 658.5 | 1011.2 | 844.3 | 0.616 | 0.045043724 | 1.192825963 | 0.349560566 | 1.969447709 | 0.036969177 |
| A0A1U7QY99 | Glutathione peroxidase OS=Mesocricetus auratus OX=10036 GN=Gpx1 PE=3 SV=1 | 8.524 | 11 | 3 | 9 | 3 | 201 | 22.4 | 6.57 | 22.45 | 3 | 206.3 | 183.4 | 106.4 | 187 | 346.7 | 103.1 | 225.8 | 196.1 | 128.7 | 1.124 | 0.592549641 | 1.109856884 | 0.685560497 | 0.864635678 | 0.737054053 |
| A0A1U8BIS9 | Glutathione peroxidase OS=Mesocricetus auratus OX=10036 GN=Gpx3 PE=3 SV=1 | 6.029 | 8 | 2 | 5 | 2 | 226 | 25.5 | 7.3 | 13.57 | 2 | 27 | 0 | 0 | 21.8 | 0 | 0 | 30.3 | 0 | 0 | 0.806 | 0.888443252 | 1.122222222 | 0.939148707 | 1.389908257 | 0.832100228 |
| A0A1U7QXX6 | Glutathione peroxidase OS=Mesocricetus auratus OX=10036 GN=Gpx4 PE=3 SV=1 | 3.771 | 7 | 1 | 1 | 1 | 197 | 22.2 | 8.31 | 4.09 | 1 | 0 | 0 | 60 | 0 | 0 | 49.8 | 0 | 0 | 48.4 | 0.83 | 0.902442774 | 0.806666667 | 0.88798062 | 0.97188755 | 0.984882135 |
| A0A1U7QNR6 | Adapter protein GRB2 OS=Mesocricetus auratus OX=10036 GN=Grb2 PE=3 SV=1 | 5.842 | 9 | 2 | 2 | 2 | 217 | 25.2 | 6.32 | 7.44 | 2 | 0 | 0 | 84.5 | 0 | 0 | 79.9 | 0 | 0 | 64.7 | 0.946 | 0.970349075 | 0.765680473 | 0.862046136 | 0.809762203 | 0.889912444 |
| A0A1U7R3Y0 | G protein-coupled receptor kinase OS=Mesocricetus auratus OX=10036 GN=Grk5 PE=3 SV=1 | 4.382 | 3 | 1 | 1 | 1 | 590 | 67.7 | 8.16 | 3.33 | 1 | 0 | 0 | 60.1 | 0 | 0 | 33.1 | 0 | 0 | 41.3 | 0.551 | 0.719364174 | 0.68718802 | 0.810818242 | 1.247734139 | 0.884717926 |
| A0A1U7R6U4 | Actin-depolymerizing factor OS=Mesocricetus auratus OX=10036 GN=Gsn PE=3 SV=1 | 42.297 | 13 | 11 | 57 | 11 | 778 | 85.4 | 6.05 | 147.04 | 11 | 1173.6 | 1272.6 | 1289.3 | 972.9 | 1088 | 1063.6 | 1011.8 | 1133.9 | 1253 | 0.829 | 0.015514123 | 0.90983804 | 0.24764123 | 1.087758041 | 0.326825415 |
| A0A1U7RD56 | Glutathione synthetase OS=Mesocricetus auratus OX=10036 GN=Gss PE=3 SV=1 | 8.579 | 9 | 3 | 16 | 3 | 405 | 44.8 | 6.1 | 51.86 | 3 | 642.2 | 778 | 842.8 | 569.7 | 686.5 | 717.8 | 587.2 | 685.6 | 721.6 | 0.882 | 0.269168912 | 0.881307998 | 0.286969584 | 1.010334347 | 0.915808504 |
| A0A1U7Q833 | GST class-pi OS=Mesocricetus auratus OX=10036 GN=Gstp1 PE=3 SV=1 | 71.654 | 46 | 10 | 83 | 10 | 210 | 23.4 | 7.36 | 250.34 | 10 | 1627.7 | 1388.9 | 1368.7 | 1440.7 | 1228.9 | 1311.3 | 1747.6 | 1568.9 | 1509.3 | 0.885 | 0.268241066 | 1.100449228 | 0.253361097 | 1.212238439 | 0.041839078 |
| A0A1U7QYH1 | Guanylate cyclase OS=Mesocricetus auratus OX=10036 GN=Gucy1a2 PE=3 SV=1 | 5.874 | 3 | 2 | 2 | 2 | 731 | 81.7 | 7.33 | 7.35 | 2 | 74.5 | 50.3 | 0 | 73.4 | 57.3 | 0 | 63.6 | 42.2 | 0 | 1.06 | 0.952863527 | 0.84775641 | 0.83707582 | 0.809487376 | 0.789846946 |
| A0A1U7QWH7 | Guanylate cyclase OS=Mesocricetus auratus OX=10036 GN=Gucy1b3 PE=3 SV=1 | 6.286 | 2 | 1 | 6 | 1 | 619 | 70.4 | 5.35 | 26.01 | 1 | 40.5 | 120.8 | 132.7 | 85.6 | 102.2 | 112.9 | 0 | 121.8 | 146.6 | 0.901 | 0.946680425 | 1.369387755 | 0.882861468 | 1.338875956 | 0.835570423 |
| A0A1U7QAZ8 | Histone H3 OS=Mesocricetus auratus OX=10036 GN=LOC101840088 PE=3 SV=1 | 37.635 | 40 | 9 | 124 | 1 | 136 | 15.3 | 11.27 | 229.08 | 9 | 0 | 0 | 37.8 | 0 | 0 | 30.4 | 0 | 0 | 27.4 | 0.804 | 0.886458959 | 0.724867725 | 0.835630837 | 0.901315789 | 0.945119758 |
| A0A1U7QS51 | Enoyl-CoA hydratase OS=Mesocricetus auratus OX=10036 GN=Hadha PE=3 SV=1 | 50.52 | 16 | 11 | 74 | 11 | 763 | 82.7 | 9.07 | 224.14 | 11 | 814.6 | 1030.8 | 960.4 | 689.5 | 798.6 | 807.7 | 689.5 | 1005.1 | 874.3 | 0.8 | 0.098520957 | 0.915567753 | 0.522347189 | 1.118956355 | 0.433645082 |
| A0A1U7QUW5 | trifunctional enzyme subunit beta, mitochondrial OS=Mesocricetus auratus OX=10036 GN=Hadhb PE=3 SV=1 | 26.088 | 19 | 7 | 28 | 7 | 475 | 51.3 | 9.47 | 79.7 | 7 | 489.6 | 512.5 | 393.1 | 431.2 | 442 | 270.1 | 461.7 | 462.7 | 338.2 | 0.689 | 0.285406914 | 0.904959862 | 0.468770624 | 1.104347066 | 0.599051525 |
| A0A1U8C284 | hematopoietic lineage cell-specific protein OS=Mesocricetus auratus OX=10036 GN=Hcls1 PE=4 SV=1 | 4.415 | 6 | 2 | 7 | 2 | 465 | 51.8 | 4.97 | 14.56 | 2 | 204.7 | 69 | 227.6 | 222.5 | 81.2 | 259.5 | 197.9 | 106.2 | 249.1 | 1.14 | 0.792933921 | 1.10353082 | 0.802988504 | 0.982244318 | 0.963691728 |
| A0A1U7QSW2 | hepatoma-derived growth factor isoform X1 OS=Mesocricetus auratus OX=10036 GN=Hdgf PE=4 SV=1 | 21.872 | 15 | 4 | 28 | 3 | 236 | 26.4 | 4.79 | 93.59 | 4 | 136.3 | 205.7 | 261 | 98.7 | 164.6 | 234.2 | 148.2 | 210.7 | 252.6 | 0.724 | 0.545052044 | 1.014096186 | 0.95502852 | 1.229145729 | 0.487986175 |
| A0A1U7QRC1 | heme-binding protein 1 OS=Mesocricetus auratus OX=10036 GN=Hebp1 PE=3 SV=1 | 11.077 | 24 | 3 | 6 | 3 | 194 | 21.6 | 5.43 | 17.05 | 3 | 61.4 | 181.5 | 0 | 39.2 | 199.5 | 0 | 42.3 | 213.4 | 0 | 0.837 | 0.987055982 | 1.052696583 | 0.962126021 | 1.071219103 | 0.952476254 |
| A0A1U8BM61 | Beta-hexosaminidase OS=Mesocricetus auratus OX=10036 GN=Hexa PE=3 SV=1 | 3.01 | 2 | 1 | 3 | 1 | 538 | 61.7 | 6.42 | 8.73 | 1 | 36.1 | 0 | 0 | 33 | 0 | 0 | 34.6 | 0 | 0 | 0.914 | 0.952526855 | 0.958448753 | 0.977508341 | 1.048484848 | 0.974911913 |
| A0A1U7Q3G6 | Beta-hexosaminidase OS=Mesocricetus auratus OX=10036 GN=Hexb PE=3 SV=1 | 7.807 | 4 | 2 | 12 | 2 | 539 | 61 | 8.13 | 32.9 | 2 | 162.3 | 220.9 | 221.1 | 112.7 | 152.5 | 170.7 | 152.9 | 182.8 | 179.8 | 0.694 | 0.098228149 | 0.853053119 | 0.26987526 | 1.182610691 | 0.265143919 |
| A0A1U7QL34 | Hepatocyte growth factor-regulated tyrosine kinase substrate OS=Mesocricetus auratus OX=10036 GN=Hgs PE=4 SV=1 | 3.66 | 2 | 1 | 2 | 1 | 774 | 85.8 | 6.16 | 7.42 | 1 | 0 | 0 | 39 | 0 | 0 | 44 | 0 | 0 | 37.4 | 1.128 | 0.936372318 | 0.958974359 | 0.977798512 | 0.85 | 0.91465007 |
| A0A1U7R136 | histidine triad nucleotide-binding protein 2, mitochondrial OS=Mesocricetus auratus OX=10036 GN=Hint2 PE=4 SV=1 | 8.449 | 20 | 2 | 8 | 2 | 163 | 17.4 | 9.7 | 26.1 | 2 | 0 | 35.5 | 0 | 0 | 37.2 | 0 | 0 | 39.3 | 0 | 1.048 | 0.975213482 | 1.107042254 | 0.946276872 | 1.056451613 | 0.970909255 |
| A0A1U7QXL1 | high mobility group protein B2 OS=Mesocricetus auratus OX=10036 GN=Hmgb2 PE=3 SV=1 | 26.148 | 44 | 7 | 28 | 6 | 210 | 24.2 | 7.39 | 71.79 | 7 | 305.6 | 702.3 | 656.2 | 484.1 | 1136.3 | 943.9 | 326.4 | 716.1 | 646.9 | 1.618 | 0.2735748 | 1.015203413 | 0.963564963 | 0.658815271 | 0.281926802 |
| B5ANQ1 | 3-hydroxy-3-methylglutaryl coenzyme A synthase (Fragment) OS=Mesocricetus auratus OX=10036 PE=2 SV=1 | 6.67 | 9 | 1 | 3 | 1 | 282 | 31.3 | 8.62 | 16.56 | 1 | 0 | 0 | 38.1 | 0 | 0 | 49.5 | 0 | 0 | 35.2 | 1.302 | 0.864615563 | 0.923884514 | 0.958112638 | 0.711111111 | 0.826613702 |
| A0A1U7QJZ4 | non-histone chromosomal protein HMG-14 OS=Mesocricetus auratus OX=10036 GN=Hmgn1 PE=3 SV=1 | 2.645 | 9 | 1 | 3 | 1 | 97 | 10.2 | 9.76 | 2.15 | 1 | 153.8 | 0 | 49.7 | 125.5 | 0 | 52 | 132.9 | 0 | 57.8 | 0.923 | 0.889008937 | 0.937100737 | 0.946311656 | 1.074366197 | 0.937798458 |
| A0A1U7QIE7 | non-histone chromosomal protein HMG-17 OS=Mesocricetus auratus OX=10036 GN=Hmgn2 PE=3 SV=1 | 3.775 | 17 | 2 | 4 | 2 | 90 | 9.4 | 9.99 | 6.43 | 2 | 68 | 45.8 | 66.3 | 84.4 | 45 | 53.1 | 67.9 | 37.7 | 38.2 | 0.921 | 0.957675325 | 0.798445308 | 0.385334383 | 0.787945205 | 0.45675608 |
| A0A1U8CJL9 | Histamine N-methyltransferase OS=Mesocricetus auratus OX=10036 GN=Hnmt PE=4 SV=1 | 2.525 | 5 | 1 | 3 | 1 | 291 | 33.4 | 5.35 | 8.63 | 1 | 98.3 | 59.1 | 102.4 | 56.8 | 46.5 | 70.8 | 70.1 | 54.3 | 90.9 | 0.65 | 0.163189989 | 0.828714396 | 0.444973312 | 1.236645606 | 0.349558552 |
| A0A1U8BTE0 | Helix-destabilizing protein OS=Mesocricetus auratus OX=10036 GN=Hnrnpa1 PE=4 SV=1 | 30.974 | 21 | 8 | 24 | 6 | 373 | 38.8 | 9.13 | 68.51 | 8 | 421.5 | 546.3 | 495.6 | 370.5 | 475.3 | 449.5 | 390.2 | 584.2 | 529.2 | 0.87 | 0.309328027 | 1.027470275 | 0.855426709 | 1.160812167 | 0.366463715 |
| A0A1U8CPF6 | heterogeneous nuclear ribonucleoproteins C1/C2 isoform X5 OS=Mesocricetus auratus OX=10036 GN=Hnrnpc PE=3 SV=1 | 6.849 | 10 | 3 | 9 | 3 | 294 | 32.4 | 5.08 | 4.55 | 3 | 150.3 | 253.7 | 94.2 | 144.7 | 222.4 | 73.3 | 149.6 | 288.4 | 94.4 | 0.826 | 0.776893082 | 1.06864713 | 0.88571226 | 1.208900999 | 0.693763233 |
| A0A1U8BJX1 | heterogeneous nuclear ribonucleoprotein D0 isoform X1 OS=Mesocricetus auratus OX=10036 GN=Hnrnpd PE=4 SV=1 | 25.249 | 23 | 6 | 27 | 5 | 352 | 38.1 | 7.81 | 35.12 | 6 | 733.6 | 567.1 | 696.5 | 753 | 832.2 | 610.2 | 736.3 | 945.8 | 695.3 | 0.988 | 0.469451666 | 1.190366513 | 0.253532562 | 1.08290061 | 0.58202471 |
| A0A1U7QKQ3 | heterogeneous nuclear ribonucleoprotein D-like isoform X2 OS=Mesocricetus auratus OX=10036 GN=Hnrnpdl PE=4 SV=1 | 10.227 | 9 | 4 | 13 | 3 | 364 | 40 | 9.96 | 26.43 | 4 | 584.4 | 557.9 | 665.8 | 520.7 | 514.4 | 618.1 | 473 | 463.9 | 580.7 | 0.891 | 0.33094137 | 0.839334108 | 0.124130988 | 0.917977256 | 0.420536735 |
| A0A1U8BEQ4 | heterogeneous nuclear ribonucleoprotein F OS=Mesocricetus auratus OX=10036 GN=Hnrnpf PE=4 SV=1 | 27.118 | 18 | 6 | 29 | 4 | 415 | 45.7 | 5.49 | 121.03 | 6 | 0 | 0 | 130.7 | 0 | 0 | 138.8 | 0 | 0 | 112.1 | 1.062 | 0.968154512 | 0.857689365 | 0.919295164 | 0.807636888 | 0.888586529 |
| A0A1U7QSW4 | Heterogeneous nuclear ribonucleoprotein K OS=Mesocricetus auratus OX=10036 GN=Hnrnpk PE=4 SV=1 | 68.052 | 31 | 15 | 77 | 15 | 464 | 51 | 5.47 | 213.87 | 15 | 1204.8 | 1392.5 | 1371.3 | 1157.1 | 1416.5 | 1369.1 | 1127.1 | 1267.3 | 1210.2 | 0.96 | 0.935293654 | 0.908279998 | 0.176275694 | 0.914246582 | 0.297826374 |
| A0A1U7RFN8 | heterogeneous nuclear ribonucleoprotein L isoform X2 OS=Mesocricetus auratus OX=10036 GN=Hnrnpl PE=4 SV=1 | 22.926 | 13 | 5 | 22 | 5 | 586 | 63.9 | 8.1 | 75.65 | 5 | 168.9 | 365.9 | 188.7 | 154.2 | 329.8 | 161.9 | 145 | 346.2 | 188.1 | 0.901 | 0.775875131 | 0.938908086 | 0.874514414 | 1.051710791 | 0.900754851 |
| A0A1U7QUS6 | heterogeneous nuclear ribonucleoprotein M isoform X1 OS=Mesocricetus auratus OX=10036 GN=Hnrnpm PE=4 SV=1 | 28.247 | 16 | 8 | 31 | 8 | 730 | 77.6 | 8.63 | 89.62 | 8 | 273.8 | 443.2 | 461.2 | 301.9 | 385.2 | 387.6 | 233.4 | 376.4 | 364.3 | 1.035 | 0.639117075 | 0.826769649 | 0.420143517 | 0.906392482 | 0.572973527 |
| A0A1U7R7D6 | heterogeneous nuclear ribonucleoprotein R isoform X2 OS=Mesocricetus auratus OX=10036 GN=Hnrnpr PE=4 SV=1 | 26.818 | 10 | 6 | 25 | 2 | 633 | 70.9 | 8.13 | 71.38 | 6 | 369.9 | 165.3 | 252 | 361.8 | 129.9 | 265.7 | 429.6 | 149.7 | 276.7 | 0.978 | 0.917209337 | 1.087398374 | 0.831341818 | 1.130182202 | 0.770878602 |
| A0A1U7QFZ3 | heterogeneous nuclear ribonucleoprotein U OS=Mesocricetus auratus OX=10036 GN=Hnrnpu PE=4 SV=1 | 48.803 | 18 | 14 | 79 | 14 | 799 | 87.8 | 6.24 | 163.92 | 14 | 1503.1 | 2513 | 1629.5 | 1820.4 | 2279.4 | 1749.1 | 1910.4 | 2643 | 1781 | 1.189 | 0.862069166 | 1.122006518 | 0.611044823 | 1.083007061 | 0.64019976 |
| A0A1U8CR89 | heterogeneous nuclear ribonucleoprotein U-like protein 1 isoform X1 OS=Mesocricetus auratus OX=10036 GN=Hnrnpul1 PE=4 SV=1 | 7.817 | 5 | 3 | 13 | 3 | 869 | 96.6 | 6.4 | 29.16 | 3 | 270.2 | 391.5 | 335.7 | 281.6 | 331.9 | 297.9 | 275.9 | 340 | 299.1 | 0.941 | 0.511670595 | 0.917385202 | 0.538369914 | 1.003949967 | 0.962465727 |
| A0A1U7QKE3 | Homeodomain-only protein OS=Mesocricetus auratus OX=10036 GN=Hopx PE=4 SV=1 | 4.545 | 14 | 1 | 4 | 1 | 73 | 8.3 | 5.1 | 16.81 | 1 | 131.5 | 46.1 | 148.2 | 119.5 | 33.4 | 110.9 | 93.5 | 43.4 | 149 | 0.909 | 0.647635574 | 0.877532228 | 0.777189809 | 1.083775588 | 0.866188327 |

**Supplementary Table 1: List of differentially expressed proteins identified across the 4dpi infected and 14dpi infected**

| Accession | Description | Sum PEP Score | Coverage [%] | # Peptides | # PSMs | # Unique Peptides | # AAs | MW [kDa] | calc. pI | Score Sequest HT: Sequest HT | # Peptides (by Search Engine): Sequest HT | Mock-infected (4dpi) Replicate_R1 | Mock-infected (4dpi) Replicate_R2 | Mock-infected (4dpi) Replicate_R3 | SARS-CoV-2-infected (4dpi) Replicate_R1 | SARS-CoV-2-infected (4dpi) Replicate_R2 | SARS-CoV-2-infected (4dpi) Replicate_R3 | SARS-CoV-2-(14dpi) Replicate_R1 | SARS-CoV-2-infected (14dpi) Replicate_R2 | SARS-CoV-2-infected (14dpi) Replicate_R3 | Abundance Ratio: SARS-CoV-2-infected (4dpi) / Mock-infected (4dpi) | P-Value: SARS-CoV-2-infected (4dpi) / Mock-infected (4dpi) | Abundance Ratio: SARS-CoV-2-infected (14dpi) / Mock-infected (4dpi) | P-Value: SARS-CoV-2-infected (14dpi) / Mock-infected (4dpi) | Abundance Ratio: SARS-CoV-2-infected (14dpi) / SARS-CoV-2-infected (4dpi) | P-Value: SARS-CoV-2-infected (14dpi) / SARS-CoV-2-infected (4dpi) |
|---|---|---|---|---|---|---|---|---|---|---|---|---|---|---|---|---|---|---|---|---|---|---|---|---|---|---|
| A0A1U7QDD1 | homeobox protein Hox-D8 isoform X1 OS=Mesocricetus auratus OX=10036 GN=Hoxd8 PE=4 SV=1 | 1.509 | 3 | 1 | 2 | 1 | 288 | 31.4 | 8.09 | 3.54 | 1 | 0 | 34 | 45.4 | 0 | 57.6 | 69.8 | 0 | 28.8 | 42.6 | 1.613 | 0.56989654 | 0.899244332 | 0.89258646 | 0.56043956 | 0.50471072 |
| A0A1U7Q238 | Haptoglobin OS=Mesocricetus auratus OX=10036 GN=Hp PE=3 SV=1 | 12.812 | 15 | 4 | 18 | 4 | 346 | 38.6 | 6.14 | 50.23 | 4 | 210.6 | 94.3 | 206 | 896.8 | 525.3 | 833 | 112.1 | 18.1 | 118.3 | 4.575 | 0.027129388 | 0.486396555 | 0.15671022 | 0.11019467 | 0.021534259 |
| A0A1U8CMQ8 | hippocalcin-like protein 1 OS=Mesocricetus auratus OX=10036 GN=Hpcal1 PE=4 SV=1 | 10.425 | 15 | 3 | 13 | 3 | 193 | 22.3 | 5.5 | 38.23 | 3 | 317 | 444.6 | 368.1 | 279.6 | 363.1 | 312.8 | 359.7 | 456.9 | 428.3 | 0.817 | 0.270548073 | 1.101973975 | 0.462137858 | 1.302878074 | 0.064472414 |
| A0A1U7QTB7 | Hypoxanthine phosphoribosyltransferase OS=Mesocricetus auratus OX=10036 GN=Hprt1 PE=3 SV=1 | 7.259 | 11 | 2 | 6 | 2 | 218 | 24.6 | 7.42 | 17.67 | 2 | 51.3 | 105.7 | 52.2 | 38.1 | 104.1 | 42.5 | 51.6 | 87.1 | 52 | 0.813 | 0.784545906 | 0.911567878 | 0.790575048 | 1.032485111 | 0.939502811 |
| A0A1U7QU66 | inactive hydroxysteroid dehydrogenase-like protein 1 OS=Mesocricetus auratus OX=10036 GN=Hsdl1 PE=4 SV=1 | 2.47 | 2 | 1 | 6 | 1 | 330 | 37 | 8.63 | 17.25 | 1 | 366.1 | 946.7 | 202.6 | 154.1 | 397.1 | 90.4 | 133.1 | 432.9 | 87.6 | 0.446 | 0.328496434 | 0.431305266 | 0.337761173 | 1.018703242 | 0.979059505 |
| A0A1U7Q8T8 | endoplasmin OS=Mesocricetus auratus OX=10036 GN=Hsp90b1 PE=3 SV=1 | 81.891 | 24 | 18 | 107 | 17 | 802 | 92.4 | 4.83 | 313.7 | 18 | 3106.8 | 4120.6 | 4072.2 | 2537.2 | 3383.3 | 3588.6 | 3269.2 | 3298.6 | 3422.8 | 0.881 | 0.265174455 | 0.884155191 | 0.316558772 | 1.050635707 | 0.668708609 |
| A0A1U7QBD8 | heat shock 70 kDa protein 12B OS=Mesocricetus auratus OX=10036 GN=Hspa12b PE=4 SV=1 | 12.555 | 6 | 4 | 18 | 4 | 685 | 76.1 | 8.6 | 46.44 | 4 | 42 | 46.1 | 190.7 | 32 | 39.8 | 159.6 | 35 | 43.8 | 176.2 | 0.864 | 0.817486191 | 0.914634146 | 0.911357811 | 1.1019879 | 0.904560269 |
| A0A1U8CDJ8 | 78 kDa glucose-regulated protein OS=Mesocricetus auratus OX=10036 GN=Hspa5 PE=3 SV=1 | 88.021 | 33 | 22 | 111 | 21 | 655 | 72.3 | 5.16 | 321.12 | 22 | 2304.8 | 2622.4 | 2153.9 | 2502.5 | 2737.2 | 2344.4 | 2256.6 | 2479.3 | 1976 | 1.044 | 0.403981909 | 0.947861208 | 0.572922996 | 0.884996242 | 0.195158999 |
| A0A1U8BQW6 | heat shock cognate 71 kDa protein OS=Mesocricetus auratus OX=10036 GN=Hspa8 PE=3 SV=1 | 106.036 | 34 | 22 | 143 | 20 | 646 | 70.6 | 5.4 | 376.17 | 22 | 3165.1 | 3063.2 | 3156.4 | 3430.3 | 3373.5 | 3509.8 | 4184.7 | 3256 | 3923.9 | 1.111 | 0.00423885 | 1.210971049 | 0.137857446 | 1.101904282 | 0.332275429 |
| A0A1U7QTC6 | Heat shock 27 kDa protein OS=Mesocricetus auratus OX=10036 GN=Hspb1 PE=3 SV=1 | 43.621 | 58 | 12 | 61 | 12 | 212 | 23.2 | 6.55 | 166.07 | 12 | 2366.6 | 1982.4 | 1534.9 | 1916.7 | 1713.6 | 1290.5 | 1958.6 | 1709.9 | 1238.5 | 0.864 | 0.35271783 | 0.833970666 | 0.367177518 | 0.997195578 | 0.987709328 |
| A0A1U7R8L5 | heat shock protein beta-6 OS=Mesocricetus auratus OX=10036 GN=Hspb6 PE=3 SV=1 | 4.554 | 8 | 1 | 4 | 1 | 162 | 17.3 | 6.4 | 14.56 | 1 | 197 | 58.3 | 0 | 162.4 | 66.4 | 0 | 167.4 | 57 | 0 | 0.968 | 0.912245472 | 0.878965922 | 0.899374295 | 0.980769231 | 0.983848758 |
| A0A1U7QHG7 | Heat shock protein beta-8 OS=Mesocricetus auratus OX=10036 GN=Hspb8 PE=3 SV=1 | 1.453 | 5 | 1 | 1 | 1 | 196 | 21.5 | 5.12 | 1.72 | 1 | 0 | 0 | 53.8 | 0 | 0 | 57.1 | 0 | 0 | 43.4 | 1.062 | 0.968470793 | 0.80669145 | 0.887996102 | 0.760070053 | 0.858447487 |
| A0A1U8BSF0 | LOW QUALITY PROTEIN: 10 kDa heat shock protein, mitochondrial OS=Mesocricetus auratus OX=10036 GN=Hspe1 PE=3 SV=1 | 17.339 | 49 | 6 | 34 | 6 | 102 | 10.9 | 9.2 | 64.25 | 6 | 645 | 529.5 | 403.2 | 514.5 | 452.2 | 329.8 | 530.7 | 457 | 367.9 | 0.798 | 0.352259366 | 0.859225455 | 0.435405577 | 1.045584265 | 0.797702151 |
| A0A1U8CK40 | Putative hydroxypyruvate isomerase OS=Mesocricetus auratus OX=10036 GN=Hyi PE=3 SV=1 | 10.895 | 12 | 2 | 2 | 2 | 263 | 29 | 5.48 | 10.71 | 2 | 0 | 0 | 58.3 | 0 | 0 | 38.3 | 0 | 0 | 59 | 0.657 | 0.790681153 | 1.012006861 | 0.993670661 | 1.540469974 | 0.785466968 |
| A0A1U8BQU1 | hypoxia up-regulated protein 1 OS=Mesocricetus auratus OX=10036 GN=Hyou1 PE=3 SV=1 | 23.354 | 10 | 6 | 30 | 6 | 998 | 111.3 | 5.19 | 84.71 | 6 | 840.4 | 914.4 | 319.2 | 788.5 | 862.5 | 353.4 | 824.2 | 905.5 | 303.8 | 1.024 | 0.929387186 | 0.980472517 | 0.96191549 | 1.01451806 | 0.970544101 |
| A0A1U7QLF2 | huntingtin-interacting protein K isoform X1 OS=Mesocricetus auratus OX=10036 GN=Hypk PE=4 SV=1 | 2.547 | 9 | 1 | 1 | 1 | 129 | 14.7 | 4.93 | 3.52 | 1 | 192.5 | 0 | 0 | 171.9 | 0 | 0 | 149.1 | 0 | 0 | 0.893 | 0.940260724 | 0.774545455 | 0.867709467 | 0.867364747 | 0.925100992 |
| A0A1U7QMD6 | Isocitrate dehydrogenase [NADP] OS=Mesocricetus auratus OX=10036 GN=Idh1 PE=3 SV=1 | 43.577 | 29 | 10 | 58 | 9 | 414 | 46.8 | 6.99 | 154.72 | 10 | 556.8 | 434.5 | 412.5 | 545.9 | 384 | 392.2 | 454.5 | 401.9 | 375.8 | 0.968 | 0.714424197 | 0.877760365 | 0.33986436 | 0.932002118 | 0.641434614 |
| A0A1U7QNX2 | Isocitrate dehydrogenase [NADP] OS=Mesocricetus auratus OX=10036 GN=Idh2 PE=3 SV=1 | 56.356 | 32 | 13 | 75 | 12 | 452 | 50.8 | 8.68 | 207.68 | 13 | 1160.8 | 1413.4 | 1082.3 | 1009.6 | 1158.7 | 926.7 | 1105 | 1354.6 | 1019.2 | 0.87 | 0.205544257 | 0.951401614 | 0.697528673 | 1.124006462 | 0.358903721 |
| A0A1U8BJQ7 | interferon-induced protein with tetratricopeptide repeats 2 OS=Mesocricetus auratus OX=10036 GN=Ifit2 PE=4 SV=1 | 6.059 | 3 | 1 | 1 | 1 | 468 | 54.7 | 6.95 | 5.11 | 1 | 0 | 38.8 | 0 | 0 | 38.6 | 0 | 0 | 44.4 | 0 | 0.995 | 0.997259297 | 1.144329897 | 0.928982062 | 1.150259067 | 0.92629839 |
| A0A1U7R661 | interleukin enhancer-binding factor 2 OS=Mesocricetus auratus OX=10036 GN=Ilf2 PE=4 SV=1 | 4.429 | 3 | 1 | 6 | 1 | 390 | 43 | 5.26 | 19.65 | 1 | 205.9 | 146.8 | 152.9 | 190.7 | 133.3 | 132.6 | 189.3 | 158 | 173.3 | 0.926 | 0.576314551 | 1.029667722 | 0.826350828 | 1.140166448 | 0.393433895 |
| A0A1U7RF83 | Integrin-linked protein kinase OS=Mesocricetus auratus OX=10036 GN=Ilk PE=4 SV=1 | 62.028 | 32 | 14 | 65 | 14 | 452 | 51.4 | 8.07 | 192.33 | 14 | 1264.4 | 1022.5 | 1877.4 | 949.6 | 909.8 | 1587.8 | 1235.1 | 984.9 | 1835.5 | 0.771 | 0.517056831 | 0.97387316 | 0.924262855 | 1.176462056 | 0.577803878 |
| A0A1U7QIL1 | Inositol-1-monophosphatase OS=Mesocricetus auratus OX=10036 GN=Impa1 PE=3 SV=1 | 8.807 | 10 | 2 | 8 | 2 | 277 | 30.4 | 5.54 | 31.88 | 2 | 91.7 | 81.8 | 174.6 | 88.5 | 94.6 | 139 | 90.4 | 56.3 | 181.9 | 0.965 | 0.811932704 | 0.943981614 | 0.898468508 | 1.020180068 | 0.961270719 |
| A0A1U7QAV1 | Inosine-5'-monophosphate dehydrogenase OS=Mesocricetus auratus OX=10036 GN=Impdh2 PE=3 SV=1 | 11 | 8 | 3 | 10 | 3 | 514 | 55.8 | 7.27 | 19.47 | 3 | 281 | 176 | 229.3 | 276.8 | 175.3 | 207.5 | 309.8 | 174.6 | 221.2 | 0.937 | 0.844754235 | 1.028121813 | 0.904066048 | 1.069739236 | 0.774131273 |
| A0A1U8BZ66 | Kinase OS=Mesocricetus auratus OX=10036 GN=Ip6k1 PE=3 SV=1 | 2.089 | 3 | 1 | 1 | 1 | 455 | 51.5 | 7.23 | 3.6 | 1 | 0 | 78.1 | 0 | 0 | 53.6 | 0 | 0 | 65.3 | 0 | 0.687 | 0.810229148 | 0.836107554 | 0.906189534 | 1.218283582 | 0.896784882 |
| A0A1U7QNY2 | ras GTPase-activating-like protein IQGAP1 OS=Mesocricetus auratus OX=10036 GN=Iqgap1 PE=4 SV=1 | 125.337 | 22 | 28 | 120 | 28 | 1657 | 188.9 | 6.4 | 323.2 | 28 | 1705.4 | 1815.8 | 927.2 | 1416.4 | 1489.2 | 727.4 | 1477.4 | 1519.7 | 845.5 | 0.814 | 0.504371733 | 0.863816204 | 0.601174565 | 1.057693366 | 0.841020549 |
| A0A1U8CAF8 | ubiquitin-like protein ISG15 OS=Mesocricetus auratus OX=10036 GN=Isg15 PE=4 SV=1 | 21.692 | 20 | 3 | 30 | 3 | 164 | 18.6 | 7.9 | 61.74 | 3 | 96.7 | 22.6 | 103.4 | 246.9 | 175.6 | 364 | 117.2 | 74 | 195.1 | 3.577 | 0.05733524 | 1.734620566 | 0.287623318 | 0.491163382 | 0.122650695 |
| A0A1U7QRD5 | IST1 homolog OS=Mesocricetus auratus OX=10036 GN=Ist1 PE=3 SV=1 | 8.622 | 6 | 3 | 6 | 3 | 366 | 40 | 5.44 | 6.26 | 3 | 166.1 | 149.4 | 49.5 | 122.9 | 144.3 | 43.2 | 196.8 | 163.4 | 59.2 | 0.74 | 0.722486687 | 1.149041096 | 0.759089883 | 1.351159794 | 0.523273226 |
| A0A1U7QC50 | integrin alpha-3 isoform X2 OS=Mesocricetus auratus OX=10036 GN=Itga3 PE=3 SV=1 | 14.157 | 4 | 4 | 13 | 4 | 1051 | 116.4 | 6.28 | 39 | 4 | 463.1 | 458.2 | 202.5 | 295.6 | 252.4 | 197.3 | 426.2 | 479.6 | 160.3 | 0.706 | 0.277577118 | 0.948656345 | 0.890490468 | 1.430430699 | 0.393836319 |
| A0A1U7Q562 | integrin alpha-M isoform X2 OS=Mesocricetus auratus OX=10036 GN=Itgam PE=3 SV=1 | 5.146 | 3 | 2 | 5 | 2 | 1151 | 127.5 | 6.42 | 14.97 | 2 | 43.6 | 101.2 | 0 | 41.9 | 110.2 | 0 | 38.3 | 113 | 0 | 1.023 | 0.95807253 | 1.044889503 | 0.963345296 | 0.994740302 | 0.995669084 |
| A0A1U7Q8L0 | protein jagunal homolog 1 isoform X1 OS=Mesocricetus auratus OX=10036 GN=Jagn1 PE=3 SV=1 | 3.168 | 7 | 1 | 1 | 1 | 183 | 21.1 | 9.76 | 3.52 | 1 | 46.7 | 0 | 0 | 30.1 | 0 | 0 | 36.7 | 0 | 0 | 0.645 | 0.78237731 | 0.785867238 | 0.874900137 | 1.219269103 | 0.896374256 |
| A0A1U8C8V1 | KN motif and ankyrin repeat domain-containing protein 2 OS=Mesocricetus auratus OX=10036 GN=Kank2 PE=4 SV=1 | 19.913 | 9 | 5 | 24 | 5 | 853 | 91 | 5.66 | 82.26 | 5 | 504.5 | 348.4 | 634.3 | 336.8 | 279.5 | 466.3 | 361.7 | 273.7 | 527.4 | 0.749 | 0.256008915 | 0.781871974 | 0.386413016 | 1.074080916 | 0.788397688 |
| A0A1U8CBY1 | KN motif and ankyrin repeat domain-containing protein 3 isoform X3 OS=Mesocricetus auratus OX=10036 GN=Kank3 PE=4 SV=1 | 26.504 | 12 | 7 | 19 | 7 | 786 | 83.8 | 5.19 | 56.09 | 7 | 594.1 | 201.5 | 392.7 | 562.1 | 244.9 | 321.9 | 406.4 | 214.1 | 307.1 | 1.022 | 0.900363985 | 0.780610957 | 0.542037135 | 0.821684826 | 0.583891283 |
| A0A1U7QBJ7 | BTB/POZ domain-containing protein KCTD12 OS=Mesocricetus auratus OX=10036 GN=Kctd12 PE=4 SV=1 | 3.665 | 6 | 1 | 8 | 1 | 329 | 36 | 5.81 | 26.15 | 1 | 0 | 87.8 | 53.3 | 0 | 63.8 | 41.1 | 0 | 73.8 | 38 | 0.749 | 0.723969309 | 0.792345854 | 0.784078921 | 1.06577693 | 0.939258748 |
| A0A1U8CLC0 | uncharacterized protein KIAA0754 homolog OS=Mesocricetus auratus OX=10036 GN=Kiaa0754 PE=4 SV=1 | 2.16 | 1 | 1 | 6 | 1 | 1077 | 116.6 | 4.51 | 12.3 | 1 | 25.3 | 0 | 132.7 | 79.2 | 0 | 281.5 | 20.7 | 0 | 159 | 2.577 | 0.522521711 | 1.137341772 | 0.916172654 | 0.498197948 | 0.576789101 |
| A0A1U7R0X2 | junctional protein associated with coronary artery disease isoform X1 OS=Mesocricetus auratus OX=10036 GN=Jcad PE=4 SV=1 | 18.775 | 5 | 4 | 22 | 4 | 1320 | 145.1 | 6.9 | 81.78 | 4 | 102 | 37.1 | 44.9 | 84.4 | 28.4 | 41.1 | 83.9 | 35.3 | 24.4 | 0.765 | 0.72550402 | 0.780434783 | 0.649612348 | 0.933073424 | 0.897181154 |
| A0A1U8C0T9 | Krueppel-like factor 3 isoform X1 OS=Mesocricetus auratus OX=10036 GN=Klf3 PE=4 SV=1 | 4.54 | 3 | 1 | 9 | 1 | 343 | 38.6 | 9.38 | 4.63 | 1 | 894.2 | 368.4 | 875.2 | 392 | 147.5 | 331.3 | 954.9 | 402 | 947.6 | 0.438 | 0.118966209 | 1.07797736 | 0.835895336 | 2.646417088 | 0.106102596 |
| A0A1U7QDL3 | Bradykinin OS=Mesocricetus auratus OX=10036 GN=Kng1 PE=4 SV=1 | 19.064 | 8 | 5 | 27 | 5 | 614 | 67.9 | 6.38 | 72.85 | 5 | 544 | 586.7 | 444.3 | 838.1 | 995.7 | 662.6 | 439.1 | 470.4 | 403.3 | 1.541 | 0.068428564 | 0.83352381 | 0.162573695 | 0.525877263 | 0.049696606 |
| A0A1U7QKP8 | importin subunit beta-1 OS=Mesocricetus auratus OX=10036 GN=Kpnb1 PE=3 SV=1 | 28.577 | 10 | 7 | 51 | 7 | 876 | 97.1 | 4.78 | 129.34 | 7 | 355.4 | 540.8 | 413.7 | 359.7 | 542.6 | 421.8 | 323.3 | 438.3 | 329.9 | 1.012 | 0.953739512 | 0.833269715 | 0.34096192 | 0.82433351 | 0.308593919 |
| A0A1U7QR87 | GTPase KRas isoform X3 OS=Mesocricetus auratus OX=10036 GN=Kras PE=4 SV=1 | 4.94 | 9 | 2 | 5 | 1 | 227 | 26.1 | 8.91 | 10.89 | 2 | 249.8 | 404.3 | 0 | 208.7 | 295.8 | 0 | 221.2 | 389.6 | 0 | 0.782 | 0.752638778 | 0.933802171 | 0.933743605 | 1.210703667 | 0.817110728 |
| A0A1U7QJF8 | keratin, type I cytoskeletal 18 OS=Mesocricetus auratus OX=10036 GN=Krt18 PE=3 SV=1 | 32.588 | 18 | 8 | 35 | 7 | 425 | 47.7 | 5.33 | 105.16 | 8 | 370.3 | 405.1 | 686.3 | 419.7 | 429.6 | 713.7 | 458.9 | 506.1 | 913.9 | 1.04 | 0.819928472 | 1.285421085 | 0.478039873 | 1.202111324 | 0.581499222 |
| A0A1U7QWJ5 | keratin, type I cytoskeletal 19 isoform X1 OS=Mesocricetus auratus OX=10036 GN=Krt19 PE=3 SV=1 | 72.333 | 33 | 16 | 106 | 11 | 403 | 44.4 | 5.25 | 296.22 | 16 | 5199.5 | 3941.6 | 4532.9 | 3902.3 | 3073.7 | 4623.6 | 5189 | 4531.1 | 5842.1 | 0.78 | 0.299248784 | 1.13808688 | 0.296562287 | 1.341615228 | 0.089218917 |
| A0A1U7Q9T8 | keratin, type II cytoskeletal 5 OS=Mesocricetus auratus OX=10036 GN=LOC101834526 PE=3 SV=1 | 31.399 | 9 | 7 | 66 | 2 | 591 | 62.5 | 7.81 | 147.12 | 7 | 884.1 | 629.8 | 905.6 | 882.4 | 661.9 | 956 | 505.6 | 412.2 | 577.2 | 0.998 | 0.84007761 | 0.61789626 | 0.053190586 | 0.597928249 | 0.042876886 |
| A0A1U7Q9T5 | keratin, type II cytoskeletal 7 OS=Mesocricetus auratus OX=10036 GN=LOC101833224 PE=3 SV=1 | 92.142 | 33 | 17 | 150 | 10 | 460 | 51 | 5.53 | 388.19 | 17 | 883 | 1042.5 | 870.8 | 718.5 | 784 | 690.8 | 867.3 | 1151.1 | 917.6 | 0.793 | 0.048848753 | 1.049958874 | 0.679963926 | 1.338622168 | 0.094146521 |
| A0A1U7QGZ0 | keratin, type II cytoskeletal 75 isoform X2 OS=Mesocricetus auratus OX=10036 GN=Krt75 PE=3 SV=1 | 34.623 | 8 | 6 | 57 | 1 | 552 | 59.6 | 8.31 | 132 | 6 | 0 | 77.8 | 0 | 0 | 62.4 | 0 | 0 | 38.6 | 0 | 0.802 | 0.885096314 | 0.496143959 | 0.683056533 | 0.618589744 | 0.764957127 |
| A0A1U7Q424 | keratin, type II cytoskeletal 2 oral OS=Mesocricetus auratus OX=10036 GN=LOC101837044 PE=3 SV=1 | 31.366 | 10 | 7 | 55 | 2 | 596 | 62.5 | 7.83 | 112.93 | 7 | 95 | 49.9 | 88.7 | 77.8 | 41.4 | 79 | 68 | 21.4 | 60.8 | 0.831 | 0.563592677 | 0.642979452 | 0.241088468 | 0.757820383 | 0.448827362 |
| A0A1U7QH04 | keratin, type II cytoskeletal 8 OS=Mesocricetus auratus OX=10036 GN=Krt8 PE=3 SV=1 | 82.782 | 36 | 19 | 122 | 12 | 487 | 54.2 | 5.82 | 298.98 | 19 | 1391.6 | 1524.9 | 1513.6 | 1082.1 | 1202.9 | 1183.1 | 1350.6 | 1429.9 | 1545.1 | 0.778 | 0.005084353 | 0.976411368 | 0.650210575 | 1.247253539 | 0.018108986 |
| A0A1U7QLD6 | laminin subunit alpha-5 OS=Mesocricetus auratus OX=10036 GN=Lama5 PE=4 SV=1 | 84.394 | 7 | 19 | 73 | 19 | 3746 | 405.9 | 6.87 | 237.25 | 19 | 1582.7 | 1909.6 | 1943.3 | 1192.4 | 1453 | 1411.5 | 1173.4 | 1301.8 | 1433.9 | 0.726 | 0.036104606 | 0.719166237 | 0.026971658 | 0.963568242 | 0.678652356 |
| A0A1U7QZD4 | laminin subunit beta-1 OS=Mesocricetus auratus OX=10036 GN=Lamb1 PE=4 SV=1 | 46.583 | 9 | 14 | 56 | 14 | 1834 | 202 | 4.94 | 116.39 | 14 | 982.8 | 1095.7 | 794.7 | 906.1 | 1161.8 | 666.6 | 881.5 | 1069.1 | 694.7 | 0.922 | 0.799158711 | 0.920680774 | 0.615507833 | 0.967379777 | 0.876854922 |
| A0A1U8BYW4 | laminin subunit beta-2 isoform X3 OS=Mesocricetus auratus OX=10036 GN=Lamb2 PE=4 SV=1 | 145.748 | 23 | 29 | 144 | 29 | 1776 | 193.5 | 6.74 | 449.6 | 29 | 3478 | 3602.4 | 3165.5 | 2603.2 | 2608.7 | 2235.5 | 2771.2 | 2774.8 | 2464.8 | 0.706 | 0.0065478 | 0.781854205 | 0.012211006 | 1.075650563 | 0.30928074 |
| A0A3Q0DBP2 | laminin subunit beta-3 isoform X1 OS=Mesocricetus auratus OX=10036 GN=Lamb3 PE=4 SV=1 | 8.482 | 3 | 3 | 7 | 3 | 1188 | 130.2 | 7.12 | 14.73 | 3 | 117.6 | 203.7 | 199.5 | 84.7 | 140.7 | 148.2 | 79.5 | 159.6 | 160.5 | 0.71 | 0.234544231 | 0.767281106 | 0.356804977 | 1.069593148 | 0.809617021 |
| A0A1U7QD11 | laminin subunit gamma-1 OS=Mesocricetus auratus OX=10036 GN=Lamc1 PE=4 SV=1 | 83.358 | 13 | 18 | 108 | 18 | 1607 | 177.2 | 5.17 | 248.77 | 18 | 2270.2 | 2868.1 | 2712.9 | 1793.4 | 2103 | 1926.2 | 2053.1 | 2568.6 | 2384.5 | 0.71 | 0.044485069 | 0.89237314 | 0.297141163 | 1.203276887 | 0.103077045 |
| A0A1U7Q4Q8 | Cytosol aminopeptidase OS=Mesocricetus auratus OX=10036 GN=Lap3 PE=3 SV=1 | 33.057 | 23 | 8 | 32 | 8 | 519 | 56.2 | 7.12 | 100 | 8 | 362.2 | 322.5 | 286.4 | 301.4 | 301.8 | 248.9 | 253.1 | 311.8 | 220.3 | 0.869 | 0.2336191 | 0.808567604 | 0.150386889 | 0.921488088 | 0.530085014 |
| A0A1U7QPG2 | 3-beta-hydroxysterol Delta (14)-reductase OS=Mesocricetus auratus OX=10036 GN=Lbr PE=3 SV=1 | 4.94 | 5 | 2 | 6 | 2 | 619 | 70.4 | 9.36 | 10.12 | 2 | 0 | 272.5 | 0 | 0 | 269.4 | 0 | 0 | 287.8 | 0 | 0.989 | 0.993932604 | 1.056146789 | 0.971061783 | 1.068299926 | 0.965018833 |
| A0A1U8C1L0 | plastin-2 OS=Mesocricetus auratus OX=10036 GN=Lcp1 PE=4 SV=1 | 76.814 | 33 | 15 | 76 | 12 | 627 | 70.2 | 5.33 | 236.48 | 15 | 517.4 | 300 | 424.3 | 553.2 | 374.8 | 480.2 | 512 | 248.5 | 413.2 | 1.132 | 0.534711798 | 0.945236369 | 0.831144063 | 0.833475359 | 0.452648761 |
| A0A1U7Q8M1 | L-lactate dehydrogenase OS=Mesocricetus auratus OX=10036 GN=Ldhb PE=3 SV=1 | 42.796 | 24 | 9 | 41 | 6 | 334 | 36.6 | 6.24 | 94.82 | 9 | 906.8 | 1431.9 | 1208.3 | 673.9 | 1128.8 | 1165.9 | 801.9 | 1318.3 | 1089.8 | 0.826 | 0.429356033 | 0.904990133 | 0.62618977 | 1.081317793 | 0.730318738 |
| A0A1U7QXZ6 | inner nuclear membrane protein Man1 OS=Mesocricetus auratus OX=10036 GN=Lemd3 PE=4 SV=1 | 1.611 | 2 | 1 | 1 | 1 | 915 | 99.2 | 7.78 | 4.23 | 1 | 40.1 | 0 | 0 | 29.6 | 0 | 0 | 25.8 | 0 | 0 | 0.737 | 0.844288672 | 0.643391521 | 0.781607942 | 0.871621622 | 0.927641431 |
| A0A1U7Q9G8 | Galectin OS=Mesocricetus auratus OX=10036 GN=Lgals1 PE=4 SV=1 | 14.532 | 31 | 4 | 20 | 4 | 135 | 14.8 | 5.8 | 61.16 | 4 | 219.9 | 472.7 | 330.9 | 341.1 | 605.8 | 451.4 | 223.2 | 485.5 | 350.2 | 1.352 | 0.304180206 | 1.034587201 | 0.916176961 | 0.757276693 | 0.353328256 |
| A0A1U7R1X7 | Galectin OS=Mesocricetus auratus OX=10036 GN=Lgals3 PE=4 SV=1 | 28.887 | 27 | 6 | 50 | 6 | 245 | 25.7 | 8.92 | 196.1 | 6 | 967.1 | 957.5 | 930.5 | 1424.3 | 1256.1 | 1170.7 | 1143.7 | 992.1 | 976.6 | 1.312 | 0.044218365 | 1.090119435 | 0.246160645 | 0.808184675 | 0.060864088 |
| A0A1U7Q3N9 | Galectin-3-binding protein OS=Mesocricetus auratus OX=10036 GN=Lgals3bp PE=4 SV=1 | 9.569 | 7 | 3 | 10 | 3 | 578 | 64.4 | 5.24 | 16.62 | 3 | 137.4 | 149.4 | 165.6 | 292.6 | 311.1 | 301.4 | 187.8 | 202.9 | 220.5 | 2.071 | 0.000263964 | 1.351016799 | 0.013885079 | 0.675284499 | 0.002329728 |
| A0A1U7R0T0 | Galectin OS=Mesocricetus auratus OX=10036 GN=Lgals9 PE=4 SV=1 | 10.941 | 6 | 2 | 25 | 2 | 354 | 39.7 | 9.54 | 87.73 | 2 | 1031.3 | 867.2 | 1017.9 | 935 | 783.9 | 972.1 | 778.1 | 698.8 | 806.5 | 0.955 | 0.390248778 | 0.782951584 | 0.035881873 | 0.848532144 | 0.12740882 |
| A0A1U7Q831 | LIM domain and actin-binding protein 1 OS=Mesocricetus auratus OX=10036 GN=Lima1 PE=4 SV=1 | 4.423 | 4 | 2 | 3 | 2 | 755 | 84.1 | 7.36 | 0 | 2 | 51.3 | 0 | 60.8 | 49.5 | 0 | 46.6 | 68.8 | 0 | 54.2 | 0.859 | 0.840351655 | 1.097234612 | 0.903725198 | 1.279916753 | 0.752009776 |
| A0A1U7QLG1 | LIM domain-containing protein 2 OS=Mesocricetus auratus OX=10036 GN=Limd2 PE=4 SV=1 | 1.959 | 10 | 1 | 1 | 1 | 128 | 14.2 | 9.17 | 3 | 1 | 87 | 0 | 0 | 55 | 0 | 0 | 90.8 | 0 | 0 | 0.632 | 0.774090098 | 1.043678161 | 0.977343222 | 1.650909091 | 0.756306387 |
| A0A1U7Q352 | Protein lin-7 homolog OS=Mesocricetus auratus OX=10036 GN=Lin7c PE=3 SV=1 | 3.783 | 8 | 2 | 4 | 2 | 197 | 21.8 | 8.43 | 4.1 | 2 | 141.7 | 122 | 152.1 | 122.2 | 117.6 | 130.6 | 119.8 | 120.7 | 132 | 0.863 | 0.222624266 | 0.895863396 | 0.23951974 | 1.005669546 | 0.904308039 |
| A0A1U7QU24 | protein ERGIC-53 OS=Mesocricetus auratus OX=10036 GN=Lman1 PE=4 SV=1 | 14.191 | 12 | 3 | 10 | 3 | 517 | 57.9 | 6.49 | 43.82 | 3 | 124.9 | 211.5 | 134.9 | 99.6 | 164.2 | 119.2 | 100.1 | 164.6 | 122.6 | 0.798 | 0.433071785 | 0.821769574 | 0.452602273 | 1.011227154 | 0.960042491 |
| A0A1U7Q3M9 | LIM and cysteine-rich domains protein 1 OS=Mesocricetus auratus OX=10036 GN=Lmcd1 PE=4 SV=1 | 17.54 | 24 | 6 | 24 | 6 | 365 | 41 | 8.02 | 70.74 | 6 | 408.5 | 333.7 | 617.1 | 402.7 | 320.4 | 611.4 | 318.1 | 294.7 | 489.6 | 0.986 | 0.948898168 | 0.811005665 | 0.463536893 | 0.826077182 | 0.510722185 |
| A0A1U7QSX8 | lamin OS=Mesocricetus auratus OX=10036 GN=Lmna PE=3 SV=1 | 173.775 | 46 | 36 | 261 | 34 | 663 | 74 | 7.02 | 688.56 | 36 | 8462.9 | 7363 | 7577.4 | 7063.4 | 6012.8 | 6995.3 | 9620.2 | 8126.6 | 9108.2 | 0.851 | 0.080857675 | 1.147487747 | 0.110380636 | 1.337966769 | 0.01706664 |
| A0A1U7QJV3 | lamin-B1 OS=Mesocricetus auratus OX=10036 GN=Lmnb1 PE=3 SV=1 | 79.958 | 38 | 20 | 110 | 16 | 454 | 51.7 | 5.07 | 285.58 | 20 | 1856.1 | 2286.1 | 2067.3 | 1874.6 | 2529.8 | 2044.9 | 3172 | 2867.2 | 2438.4 | 1.01 | 0.751026601 | 1.365262904 | 0.049743768 | 1.314499248 | 0.080188001 |

Supplementary Table 1: List of differentially expressed proteins identified across the 4dpi infected and 14dpi infected

| Accession | Description | Sum PEP Score | Coverage [%] | # Peptides | # PSMs | # Unique Peptides | # AAs | MW [kDa] | calc. pI | Score Sequest HT: Sequest HT | # Peptides (by Search Engine): Sequest HT | Mock-infected (4dpi) Replicate_R1 | Mock-infected (4dpi) Replicate_R2 | Mock-infected (4dpi) Replicate_R3 | SARS-CoV-2-infected (4dpi) Replicate_R1 | SARS-CoV-2-infected (4dpi) Replicate_R2 | SARS-CoV-2-infected (4dpi) Replicate_R3 | SARS-CoV-2-(14dpi) Replicate_R1 | SARS-CoV-2-infected (14dpi) Replicate_R2 | SARS-CoV-2-infected (14dpi) Replicate_R3 | Abundance Ratio: SARS-CoV-2-infected (4dpi) / Mock-infected (4dpi) | P-Value: SARS-CoV-2-infected (4dpi) / Mock-infected (4dpi) | Abundance Ratio: SARS-CoV-2-infected (14dpi) / Mock-infected (4dpi) | P-Value: SARS-CoV-2-infected (14dpi) / Mock-infected (4dpi) | Abundance Ratio: SARS-CoV-2-infected (14dpi) / SARS-CoV-2-infected (4dpi) | P-Value: SARS-CoV-2-infected (14dpi) / SARS-CoV-2-infected (4dpi) |
|---|---|---|---|---|---|---|---|---|---|---|---|---|---|---|---|---|---|---|---|---|---|---|---|---|---|---|
| A0A1U7RAW2 | lamin-B2 isoform X1 OS=Mesocricetus auratus OX=10036 GN=Lmnb2 PE=3 SV=1 | 38.827 | 14 | 12 | 66 | 8 | 591 | 66.5 | 5.27 | 141.63 | 12 | 1140.2 | 1001.3 | 1238.7 | 818.5 | 857.9 | 1060.2 | 977.3 | 907.6 | 1106 | 0.718 | 0.103084859 | 0.8848293 | 0.22516703 | 1.092925528 | 0.424552119 |
| A0A1U8BPW9 | prosaposin isoform X3 OS=Mesocricetus auratus OX=10036 GN=Psap PE=4 SV=1 | 7.423 | 5 | 2 | 10 | 2 | 543 | 59.8 | 5.48 | 24.51 | 2 | 0 | 45.8 | 48.8 | 0 | 83.5 | 86.9 | 0 | 49.9 | 71.6 | 1.8 | 0.491571666 | 1.28435518 | 0.752814415 | 0.713028169 | 0.671384127 |
| A0A1U7QX31 | N-acyl-L-amino-acid amidohydrolase OS=Mesocricetus auratus OX=10036 GN=Acy1 PE=3 SV=1 | 5.272 | 4 | 2 | 6 | 2 | 408 | 45.8 | 6.35 | 14.63 | 2 | 0 | 336 | 310.6 | 0 | 275.8 | 289.1 | 0 | 293.2 | 272 | 0.874 | 0.858730127 | 0.874110733 | 0.85934529 | 1.000531067 | 0.999437694 |
| A0A1U7QAE1 | Complex III subunit 8 OS=Mesocricetus auratus OX=10036 GN=LOC101822731 PE=3 SV=1 | 2.707 | 10 | 1 | 12 | 1 | 82 | 9.9 | 10.59 | 37.27 | 1 | 532.3 | 524 | 473.4 | 438.1 | 430.7 | 411.5 | 543.5 | 564.8 | 496.8 | 0.869 | 0.030774964 | 1.049290711 | 0.408778404 | 1.253690541 | 0.021161868 |
| A0A1U7QSJ9 | cytochrome P450 4B1 OS=Mesocricetus auratus OX=10036 GN=LOC101822933 PE=3 SV=1 | 44.818 | 26 | 12 | 36 | 12 | 509 | 58.6 | 8.56 | 87.54 | 12 | 169.1 | 238.9 | 211 | 158.4 | 204.9 | 185.7 | 181.7 | 234.4 | 219.9 | 0.858 | 0.399861089 | 1.027463651 | 0.836679574 | 1.158469945 | 0.235634615 |
| A0A1U8CPM3 | alcohol dehydrogenase 1 isoform X2 OS=Mesocricetus auratus OX=10036 GN=LOC101823051 PE=3 SV=1 | 19.681 | 22 | 7 | 21 | 7 | 335 | 35.4 | 7.87 | 47.15 | 7 | 373.7 | 530.7 | 421.9 | 330.5 | 440.1 | 415.3 | 351.7 | 489.5 | 429.9 | 0.899 | 0.462740994 | 0.958380457 | 0.779055535 | 1.071844169 | 0.614210124 |
| A0A1U7R614 | 40S ribosomal protein S5 OS=Mesocricetus auratus OX=10036 GN=Rps5 PE=3 SV=1 | 7.331 | 12 | 2 | 11 | 2 | 204 | 22.9 | 9.72 | 29.14 | 2 | 63.1 | 86.5 | 121.4 | 58.4 | 86.7 | 95 | 59.4 | 90.1 | 102.6 | 1.002 | 0.64164479 | 0.930258303 | 0.782665984 | 1.049979175 | 0.825314941 |
| A0A1U7R1Z2 | 40S ribosomal protein S23 OS=Mesocricetus auratus OX=10036 GN=LOC101824003 PE=3 SV=1 | 5.747 | 10 | 2 | 6 | 2 | 143 | 15.8 | 10.49 | 18.1 | 2 | 330.3 | 308.5 | 325.2 | 332 | 291.1 | 373.5 | 236.8 | 319.4 | 250.5 | 1.106 | 0.697643315 | 0.836825726 | 0.170239515 | 0.809452137 | 0.144412553 |
| A0A1U7QGL7 | Protein odr-4 homolog OS=Mesocricetus auratus OX=10036 GN=CUNH1orf27 PE=3 SV=1 | 5.254 | 4 | 1 | 5 | 1 | 448 | 49.9 | 6.06 | 20.49 | 1 | 35.1 | 33.1 | 43.4 | 36.6 | 39.2 | 36.8 | 39 | 39.8 | 38.4 | 1.072 | 0.926904117 | 1.050179211 | 0.614889059 | 1.040852575 | 0.200585685 |
| A0A1U7QQY0 | prenylcysteine oxidase 1 OS=Mesocricetus auratus OX=10036 GN=Pcyox1 PE=3 SV=1 | 23.151 | 14 | 6 | 25 | 6 | 505 | 56.4 | 6.39 | 63.92 | 6 | 421.4 | 348.6 | 286.4 | 309.8 | 280.6 | 227.2 | 326.8 | 265.7 | 214.7 | 0.793 | 0.171897988 | 0.764104506 | 0.179164603 | 0.987279843 | 0.936111038 |
| A0A1U7R9S7 | 40S ribosomal protein S9 OS=Mesocricetus auratus OX=10036 GN=Rps9 PE=3 SV=1 | 23.358 | 36 | 8 | 60 | 8 | 194 | 22.6 | 10.65 | 155.46 | 8 | 1785.2 | 1710.9 | 1445.5 | 1783.6 | 1675.9 | 1413.9 | 1718.8 | 1710.6 | 1542.7 | 0.98 | 0.887351603 | 1.00617209 | 0.936556553 | 1.020252801 | 0.807663361 |
| A0A1U8C433 | ubiquitin-conjugating enzyme E2 L3 isoform X2 OS=Mesocricetus auratus OX=10036 GN=Ube2l3 PE=3 SV=1 | 14.612 | 20 | 2 | 29 | 2 | 122 | 14.1 | 8.92 | 97.99 | 2 | 606.1 | 311.1 | 379.3 | 457 | 243.2 | 339.7 | 515.6 | 277.2 | 309.4 | 0.86 | 0.479637639 | 0.850134979 | 0.608179636 | 1.059909607 | 0.841207389 |
| A0A1U7QVQ2 | guanylate-binding protein 2-like OS=Mesocricetus auratus OX=10036 GN=LOC101824821 PE=3 SV=1 | 5.06 | 2 | 1 | 6 | 1 | 592 | 67.7 | 5.71 | 28.3 | 1 | 87 | 62.1 | 50.5 | 291.7 | 115.2 | 185.4 | 55.9 | 45.5 | 0 | 3.351 | 0.119802676 | 0.762024048 | 0.194775998 | 0.256795543 | 0.073492882 |
| A0A1U8C0Q7 | 60S ribosomal protein L31 OS=Mesocricetus auratus OX=10036 GN=LOC101825069 PE=3 SV=1 | 10.785 | 34 | 4 | 7 | 4 | 107 | 12.4 | 10.62 | 12.13 | 4 | 108.5 | 132 | 217.6 | 178.9 | 139.4 | 241.2 | 96.2 | 138.2 | 242.2 | 1.108 | 0.489993275 | 1.040384196 | 0.915887516 | 0.851831993 | 0.630234586 |
| A0A1U7QGC7 | GTP-binding nuclear protein Ran OS=Mesocricetus auratus OX=10036 GN=LOC101825141 PE=3 SV=1 | 28.802 | 23 | 5 | 22 | 1 | 216 | 24.4 | 7.91 | 63.42 | 5 | 0 | 130 | 76.4 | 0 | 134.2 | 102.3 | 0 | 105.5 | 60 | 1.176 | 0.864959676 | 0.801841085 | 0.79331071 | 0.699788584 | 0.666737681 |
| A0A1U7RDX9 | Serum amyloid A protein OS=Mesocricetus auratus OX=10036 GN=LOC101825362 PE=3 SV=1 | 8.124 | 20 | 2 | 5 | 2 | 110 | 12.4 | 6.16 | 16.55 | 2 | 49.5 | 0 | 86.4 | 110.9 | 0 | 242.9 | 63.7 | 0 | 75.7 | 2.511 | 0.414432133 | 1.025754231 | 0.974521515 | 0.394007914 | 0.419975343 |
| A0A1U7QBG8 | GlcNAc 2-epimerase OS=Mesocricetus auratus OX=10036 GN=Renbp PE=3 SV=1 | 1.837 | 2 | 1 | 2 | 1 | 415 | 47.8 | 8.05 | 4.08 | 1 | 0 | 106.8 | 118.1 | 0 | 94.9 | 104 | 0 | 137 | 100.6 | 0.884 | 0.871471858 | 1.056469542 | 0.943015891 | 1.194570136 | 0.819378485 |
| A0A1U7Q8D9 | Cytochrome b-c1 complex subunit 7 OS=Mesocricetus auratus OX=10036 GN=LOC101825739 PE=3 SV=1 | 8.229 | 21 | 2 | 16 | 2 | 111 | 13.6 | 9.25 | 53.67 | 2 | 422 | 402.4 | 483.2 | 342.6 | 314.2 | 358.6 | 375.8 | 351.1 | 462.8 | 0.812 | 0.037530215 | 0.909834812 | 0.404451989 | 1.17165649 | 0.222199477 |
| A0A1U7R4Q2 | Caveolin OS=Mesocricetus auratus OX=10036 GN=Cav1 PE=3 SV=1 | 64.822 | 46 | 12 | 87 | 12 | 178 | 20.6 | 6.02 | 272.08 | 12 | 2192.3 | 2685.5 | 2300 | 1487 | 1847.8 | 1626.6 | 3125.3 | 2837 | 2526 | 0.707 | 0.019384067 | 1.182576834 | 0.130356275 | 1.7108679 | 0.007781848 |
| A0A1U7R5Y6 | eukaryotic peptide chain release factor GTP-binding subunit ERF3B OS=Mesocricetus auratus OX=10036 GN=Gspt2 PE=4 SV=1 | 1.991 | 1 | 1 | 1 | 1 | 629 | 68.8 | 5.19 | 0 | 1 | 0 | 131.3 | 0 | 0 | 93.2 | 0 | 0 | 115 | 0 | 0.71 | 0.825767975 | 0.875856816 | 0.930160386 | 1.233905579 | 0.890331655 |
| A0A1U8BZD8 | myosin XVB OS=Mesocricetus auratus OX=10036 GN=Myo15b PE=3 SV=1 | 2.34 | 0 | 1 | 13 | 1 | 3054 | 333.6 | 8.37 | 44.33 | 1 | 141.8 | 237 | 159.7 | 115.2 | 206.7 | 133.4 | 117.8 | 177.4 | 146.6 | 0.812 | 0.530582236 | 0.820427112 | 0.407140751 | 0.97034922 | 0.898878394 |
| A0A1U8CFS1 | Thymosin beta OS=Mesocricetus auratus OX=10036 GN=Tmsb10 PE=3 SV=1 | 8.196 | 34 | 3 | 23 | 1 | 44 | 5 | 5.36 | 16.26 | 3 | 124.5 | 201.3 | 245.2 | 129.4 | 283.6 | 303.5 | 111.6 | 231.7 | 285 | 1.409 | 0.505888529 | 1.100350263 | 0.776093077 | 0.876901605 | 0.715886693 |
| A0A1U8BJJ2 | aspartyl/asparaginyl beta-hydroxylase isoform X1 OS=Mesocricetus auratus OX=10036 GN=Asph PE=3 SV=1 | 1.579 | 2 | 1 | 1 | 1 | 755 | 84.7 | 5.19 | 2.18 | 1 | 0 | 72.7 | 0 | 0 | 60.3 | 0 | 0 | 55.5 | 0 | 0.828 | 0.902095978 | 0.763411279 | 0.860591954 | 0.92039801 | 0.956122197 |
| A0A1U7QYH2 | Cofilin, non-muscle isoform OS=Mesocricetus auratus OX=10036 GN=Cfl1 PE=3 SV=1 | 49.933 | 49 | 8 | 53 | 8 | 166 | 18.5 | 8.09 | 140.82 | 8 | 1004.8 | 705.5 | 1057.9 | 958.2 | 756.4 | 1065.1 | 961.6 | 1395.3 | 1168.2 | 1.007 | 0.97983087 | 1.273426776 | 0.205457863 | 1.268158434 | 0.190064143 |
| A0A1U7Q3S0 | histone H1.2 OS=Mesocricetus auratus OX=10036 GN=LOC101826763 PE=3 SV=1 | 50.121 | 34 | 10 | 57 | 1 | 211 | 21.2 | 11 | 118.5 | 10 | 418.7 | 305.9 | 534.4 | 500.9 | 384.6 | 589.8 | 381.4 | 282.9 | 434.9 | 1.255 | 0.462728353 | 0.873073868 | 0.544710665 | 0.7450688 | 0.17215797 |
| A0A1U7QEP7 | Complex I-9kD OS=Mesocricetus auratus OX=10036 GN=Ndufv3 PE=3 SV=1 | 3.563 | 3 | 1 | 4 | 1 | 470 | 49.8 | 9.29 | 2.47 | 1 | 83.4 | 45.7 | 186.8 | 58.2 | 30.3 | 139.9 | 68.2 | 32.1 | 143.4 | 0.749 | 0.616124048 | 0.77144666 | 0.677023787 | 1.066987741 | 0.91783281 |
| A0A1U7QK33 | protein phosphatase inhibitor 2-like OS=Mesocricetus auratus OX=10036 GN=LOC101826911 PE=3 SV=1 | 3.415 | 6 | 1 | 3 | 1 | 182 | 20.8 | 7.61 | 2.92 | 1 | 0 | 0 | 80.7 | 0 | 0 | 65 | 0 | 0 | 83.3 | 0.805 | 0.887221691 | 1.032218092 | 0.983189555 | 1.281538462 | 0.871378007 |
| Q9QZ50 | Cytochrome P-450 OS=Mesocricetus auratus OX=10036 GN=LOC101826941 PE=2 SV=1 | 16.175 | 10 | 4 | 23 | 3 | 494 | 56.5 | 9.13 | 78.91 | 4 | 0 | 52.1 | 53.3 | 0 | 36.2 | 37.6 | 0 | 36.5 | 53.7 | 0.701 | 0.651949177 | 0.855787476 | 0.840954344 | 1.222222222 | 0.799421131 |
| A0A1U8BK54 | 6.8 kDa mitochondrial proteolipid OS=Mesocricetus auratus OX=10036 GN=CUNH14orf2 PE=4 SV=1 | 2.298 | 12 | 1 | 3 | 1 | 60 | 6.9 | 10.1 | 5.72 | 1 | 33.8 | 58.1 | 40.9 | 31.1 | 54.7 | 42.6 | 32.1 | 53.2 | 39.6 | 0.958 | 0.889669394 | 0.940512048 | 0.795595768 | 0.972741433 | 0.905218766 |
| A0A1U8CWS7 | kynurenine--oxoglutarate transaminase 1 isoform X5 OS=Mesocricetus auratus OX=10036 GN=Kyat1 PE=4 SV=1 | 6.892 | 6 | 2 | 4 | 2 | 424 | 48 | 6.68 | 12.14 | 2 | 0 | 56.6 | 63.5 | 0 | 52.2 | 49.7 | 0 | 56.1 | 57.6 | 0.849 | 0.829415328 | 0.946711074 | 0.942195328 | 1.115799804 | 0.884791138 |
| A0A1U7RHD6 | cystatin-B OS=Mesocricetus auratus OX=10036 GN=Cstb PE=3 SV=1 | 2.741 | 10 | 1 | 3 | 1 | 98 | 11 | 7.39 | 0 | 1 | 22.5 | 0 | 0 | 30.5 | 0 | 0 | 23.5 | 0 | 0 | 1.356 | 0.843996519 | 1.044444444 | 0.976954798 | 0.770491803 | 0.865123346 |
| A0A1U7QYM3 | 40S ribosomal protein S21 OS=Mesocricetus auratus OX=10036 GN=LOC101830140 PE=3 SV=1 | 4.593 | 13 | 1 | 7 | 1 | 83 | 9.1 | 8.32 | 21.74 | 1 | 170.4 | 350.2 | 214.4 | 174.2 | 283.4 | 191.6 | 136.9 | 234 | 158.7 | 0.93 | 0.681500327 | 0.720544218 | 0.345292043 | 0.815773259 | 0.425331886 |
| A0A1U8BYU0 | Protein-tyrosine-phosphatase OS=Mesocricetus auratus OX=10036 GN=LOC101827410 PE=4 SV=1 | 1.299 | 0 | 1 | 4 | 1 | 1713 | 187.6 | 6.54 | 8.46 | 1 | 285.2 | 164.6 | 141.5 | 294.9 | 157.8 | 145.2 | 329.3 | 169.9 | 164.6 | 0.959 | 0.974801798 | 1.122611196 | 0.748027821 | 1.1102191 | 0.776426085 |
| A0A1U7QZZ0 | Glutathione S-transferase OS=Mesocricetus auratus OX=10036 GN=Gstm3 PE=3 SV=1 | 12.588 | 29 | 4 | 11 | 3 | 226 | 26.8 | 6.76 | 28.03 | 4 | 47.3 | 0 | 37 | 22.9 | 0 | 31.4 | 31.4 | 0 | 36.5 | 0.641 | 0.595884094 | 0.805456702 | 0.781244525 | 1.250460405 | 0.774761148 |
| A0A1U8BG52 | histone H1.0 OS=Mesocricetus auratus OX=10036 GN=LOC101827473 PE=3 SV=1 | 7.595 | 12 | 3 | 9 | 3 | 193 | 20.7 | 10.89 | 20.25 | 3 | 613.4 | 566.6 | 621.4 | 418 | 360.8 | 462.9 | 388.5 | 363.8 | 404.7 | 0.713 | 0.010116587 | 0.642278228 | 0.00087773 | 0.931787066 | 0.448844184 |
| A0A1U7RA48 | 60S ribosomal protein L11 OS=Mesocricetus auratus OX=10036 GN=LOC101827706 PE=3 SV=1 | 9.184 | 12 | 2 | 7 | 2 | 178 | 20.2 | 8.7 | 18.8 | 2 | 51.6 | 0 | 378.5 | 53.6 | 0 | 341.5 | 43.8 | 0 | 259.7 | 0.969 | 0.945078011 | 0.705649849 | 0.784694727 | 0.76815996 | 0.830458052 |
| Q4R4G1 | Signal transducer and activator of transcription OS=Mesocricetus auratus OX=10036 GN=STAT1 alpha PE=2 SV=1 | 38.852 | 16 | 9 | 38 | 9 | 749 | 87.2 | 5.83 | 111.88 | 9 | 238.5 | 241.3 | 365.9 | 520.7 | 429.4 | 760.4 | 231.1 | 144.3 | 328.3 | 2.183 | 0.08349594 | 0.832091758 | 0.525138739 | 0.411400175 | 0.056290547 |
| A0A1U7R7I2 | Tubulin beta chain OS=Mesocricetus auratus OX=10036 GN=Tubb PE=3 SV=1 | 131.884 | 52 | 19 | 233 | 4 | 444 | 49.6 | 4.89 | 871.25 | 19 | 106.4 | 0 | 0 | 107.5 | 0 | 0 | 96.1 | 0 | 0 | 1.01 | 0.994545627 | 0.903195489 | 0.946211637 | 0.893953488 | 0.940826321 |
| A0A1U7QDA7 | Glutathione S-transferase OS=Mesocricetus auratus OX=10036 GN=LOC110339199 PE=3 SV=1 | 42.228 | 46 | 10 | 53 | 3 | 218 | 25.7 | 6.8 | 174.13 | 10 | 0 | 135.7 | 113.7 | 0 | 88.4 | 64.1 | 0 | 105.1 | 102.1 | 0.606 | 0.556941162 | 0.830793905 | 0.809241721 | 1.358688525 | 0.697769459 |
| A0A1U8CW05 | LOW QUALITY PROTEIN: paraneoplastic antigen-like protein 6B OS=Mesocricetus auratus OX=10036 GN=LOC101828172 PE=4 SV=1 | 2.398 | 4 | 1 | 3 | 1 | 419 | 47 | 4.67 | 4.85 | 1 | 43.8 | 0 | 52.4 | 154.2 | 0 | 111.2 | 0 | 0 | 0 | 2.733 | 0.345911344 | #DIV/0! | 0.186746647 | #DIV/0! | 0.194013604 |
| A0A1U7Q8Z0 | histone H1.4 OS=Mesocricetus auratus OX=10036 GN=LOC101828346 PE=3 SV=1 | 49.406 | 33 | 12 | 58 | 6 | 219 | 21.9 | 11.03 | 100.44 | 12 | 4471.9 | 4421.1 | 6538.8 | 3237 | 2835.3 | 3487.7 | 3490.7 | 2892.4 | 3695.7 | 0.554 | 0.098096555 | 0.653118884 | 0.112721415 | 1.054267782 | 0.604710614 |
| A0A1U7Q479 | Histone H2A OS=Mesocricetus auratus OX=10036 GN=LOC101828968 PE=3 SV=1 | 42.685 | 46 | 8 | 106 | 1 | 130 | 14.1 | 11.05 | 286.62 | 8 | 3419.8 | 3410.9 | 2968.9 | 3988.1 | 3204.9 | 4646.7 | 5489.8 | 5091.7 | 5450.2 | 1.551 | 0.239274184 | 1.635954529 | 0.000505893 | 1.354063025 | 0.067955767 |
| A0A1U7Q2C9 | Glutathione S-transferase OS=Mesocricetus auratus OX=10036 GN=LOC101829268 PE=3 SV=1 | 60.885 | 54 | 13 | 77 | 7 | 218 | 25.7 | 7.36 | 232.5 | 13 | 1111.3 | 1027.3 | 1010.7 | 787 | 765.6 | 812 | 1100.1 | 990.8 | 904.3 | 0.745 | 0.006363345 | 0.951068491 | 0.4830569 | 1.266683583 | 0.058699437 |
| A0A1U8BXI6 | Histone H2A OS=Mesocricetus auratus OX=10036 GN=LOC101829328 PE=3 SV=1 | 21.451 | 22 | 5 | 60 | 1 | 143 | 15.1 | 10.74 | 163.21 | 5 | 132.6 | 140 | 33.1 | 119.1 | 130.7 | 34.6 | 130.4 | 127.6 | 47.1 | 0.934 | 0.884634241 | 0.998037291 | 0.996599835 | 1.07278481 | 0.873939508 |
| A0A1U7QJA2 | cathelicidin antimicrobial peptide OS=Mesocricetus auratus OX=10036 GN=Camp PE=3 SV=1 | 8.23 | 20 | 3 | 5 | 3 | 175 | 19.9 | 7.83 | 11.22 | 3 | 58.8 | 82.2 | 76.8 | 44.2 | 225.5 | 231.3 | 33 | 0 | 0 | 2.533 | 0.26325359 | 0.454545455 | 0.01349148 | 0.19760479 | 0.122125527 |
| A0A1U7QY13 | 40S ribosomal protein S29 OS=Mesocricetus auratus OX=10036 GN=LOC101829442 PE=3 SV=1 | 1.959 | 14 | 1 | 3 | 1 | 56 | 6.7 | 10.13 | 5.29 | 1 | 75 | 0 | 76 | 79.9 | 0 | 76.8 | 95.6 | 0 | 94.5 | 1.038 | 0.960749255 | 1.258940397 | 0.764295504 | 1.213146139 | 0.800207292 |
| A0A1U8BLE6 | Histone H2B OS=Mesocricetus auratus OX=10036 GN=LOC101829502 PE=3 SV=1 | 33.974 | 51 | 8 | 105 | 2 | 126 | 14 | 10.37 | 263.58 | 8 | 0 | 51.7 | 258 | 0 | 40.1 | 193.8 | 0 | 38.7 | 219.6 | 0.763 | 0.81112192 | 0.834032935 | 0.877170484 | 1.104318085 | 0.932282451 |
| A0A1U8D0E0 | plastin-3 isoform X1 OS=Mesocricetus auratus OX=10036 GN=Pls3 PE=4 SV=1 | 33.761 | 14 | 8 | 43 | 5 | 643 | 72.3 | 5.68 | 103.78 | 8 | 1145.6 | 546 | 580.9 | 991.4 | 451.4 | 497.4 | 1104.4 | 588.4 | 572.8 | 0.827 | 0.692363329 | 0.996963696 | 0.99340228 | 1.167714669 | 0.681726215 |
| A0A1U7QX94 | Glutathione S-transferase OS=Mesocricetus auratus OX=10036 GN=LOC101829715 PE=3 SV=1 | 39.57 | 41 | 9 | 71 | 6 | 218 | 26 | 7.37 | 230.6 | 9 | 2566.6 | 2781.4 | 1825.8 | 1854.1 | 1915.2 | 1408.6 | 2242.1 | 2483.3 | 1875.8 | 0.72 | 0.134284056 | 0.920181773 | 0.609421897 | 1.274879778 | 0.117792576 |
| A0A1U7QBI9 | myocardial zonula adherens protein isoform X1 OS=Mesocricetus auratus OX=10036 GN=Myzap PE=4 SV=1 | 3.661 | 6 | 2 | 3 | 2 | 466 | 53.9 | 6.18 | 2.15 | 2 | 80.9 | 0 | 0 | 76.7 | 0 | 0 | 67.2 | 0 | 0 | 0.948 | 0.971756932 | 0.83065513 | 0.902845457 | 0.876140808 | 0.930328988 |
| A0A1U7RB39 | cytochrome P450 4F5 isoform X1 OS=Mesocricetus auratus OX=10036 GN=LOC101830103 PE=3 SV=1 | 18.507 | 10 | 4 | 17 | 4 | 524 | 60.1 | 6.98 | 54.77 | 4 | 164.5 | 355.2 | 117.9 | 167.3 | 302.7 | 133 | 158.4 | 341 | 99.5 | 1.017 | 0.903966101 | 0.939303639 | 0.90613446 | 0.993200663 | 0.988592329 |
| A0A1U7QZZ4 | Glutathione transferase OS=Mesocricetus auratus OX=10036 GN=LOC101830172 PE=3 SV=1 | 22.634 | 38 | 6 | 18 | 4 | 218 | 25.5 | 6.8 | 37.2 | 6 | 0 | 96.2 | 0 | 0 | 80 | 0 | 0 | 69.8 | 0 | 0.832 | 0.903426412 | 0.725571726 | 0.836090875 | 0.8725 | 0.928164565 |
| A0A1U7QTV0 | Histone H4 OS=Mesocricetus auratus OX=10036 GN=LOC101825928 PE=3 SV=1 | 59.261 | 57 | 10 | 143 | 10 | 103 | 11.4 | 11.36 | 345.87 | 10 | 3853 | 3973.9 | 3527.5 | 5221.5 | 5998.2 | 4778.5 | 5204.9 | 5570.1 | 4813.9 | 1.355 | 0.03629207 | 1.372939125 | 0.009010337 | 0.974415872 | 0.76366803 |
| A0A1U7RIT4 | MICOS complex subunit MIC13 OS=Mesocricetus auratus OX=10036 GN=CUNH19orf70 PE=3 SV=1 | 1.383 | 6 | 1 | 1 | 1 | 119 | 13.2 | 8.1 | 1.65 | 1 | 0 | 0 | 43.6 | 0 | 0 | 28.8 | 0 | 0 | 36.5 | 0.66 | 0.793088968 | 0.837155963 | 0.906831037 | 1.267361111 | 0.876907168 |
| A0A1U8CS54 | C4a anaphylatoxin OS=Mesocricetus auratus OX=10036 GN=LOC101830930 PE=4 SV=1 | 69.393 | 13 | 16 | 78 | 16 | 1750 | 192.7 | 7.59 | 267.98 | 16 | 677.8 | 504.2 | 546 | 1509.9 | 1001.2 | 1105 | 549.2 | 549.3 | 476.9 | 2.024 | 0.044167806 | 0.911689815 | 0.446125805 | 0.435662731 | 0.0453653 |
| A0A1U7R6L4 | zinc-alpha-2-glycoprotein OS=Mesocricetus auratus OX=10036 GN=Azgp1 PE=3 SV=1 | 10.835 | 12 | 2 | 10 | 2 | 312 | 35 | 6.09 | 30.37 | 2 | 103.9 | 38.7 | 82.8 | 151.1 | 48.4 | 114.8 | 93.1 | 13.5 | 44.5 | 1.386 | 0.46055932 | 0.670363798 | 0.458214994 | 0.480750875 | 0.229502919 |
| O35587 | Transmembrane emp24 domain-containing protein 10 OS=Mesocricetus auratus OX=10036 GN=TMED10 PE=1 SV=1 | 11.849 | 17 | 3 | 25 | 3 | 219 | 24.8 | 6.68 | 51 | 3 | 586.6 | 659.9 | 532.8 | 459.4 | 519.5 | 417.9 | 705.5 | 844.9 | 610.3 | 0.787 | 0.056751466 | 1.214353959 | 0.196921456 | 1.546892898 | 0.048083822 |
| A0A1U7R6B6 | heterogeneous nuclear ribonucleoprotein H2 OS=Mesocricetus auratus OX=10036 GN=Hnrnph2 PE=4 SV=1 | 32.755 | 17 | 5 | 30 | 2 | 449 | 49.2 | 6.3 | 123.03 | 5 | 30.2 | 0 | 0 | 31.5 | 0 | 0 | 38.4 | 0 | 0 | 1.042 | 0.977663714 | 1.271523179 | 0.875274219 | 1.219047619 | 0.896466497 |
| O35534 | Carboxylic ester hydrolase OS=Mesocricetus auratus OX=10036 GN=LOC101832386 PE=2 SV=1 | 82.774 | 33 | 15 | 110 | 15 | 565 | 62.1 | 6.58 | 293.6 | 15 | 497.4 | 465.8 | 271.3 | 352.6 | 332.7 | 205.5 | 460.7 | 433 | 251.6 | 0.709 | 0.256775259 | 0.927744026 | 0.773275073 | 1.285698249 | 0.355808833 |
| A0A1U7QZE2 | 40S ribosomal protein S17 OS=Mesocricetus auratus OX=10036 GN=Rps17 PE=3 SV=1 | 10.406 | 31 | 3 | 11 | 3 | 135 | 15.5 | 9.85 | 21.31 | 3 | 280.4 | 177.7 | 413 | 197.7 | 163.8 | 344.2 | 235.1 | 149.7 | 407.4 | 0.705 | 0.56530487 | 0.909424865 | 0.809195694 | 1.122573331 | 0.775344868 |

**Supplementary Table 1: List of differentially expressed proteins identified across the 4dpi infected and 14dpi infected**

| Accession | Description | Sum PEP Score | Coverage [%] | # Peptides | # PSMs | # Unique Peptides | # AAs | MW [kDa] | calc. pI | Score Sequest HT: Sequest HT | # Peptides (by Search Engine): Sequest HT | Mock-infected (4dpi) Replicate_R1 | Mock-infected (4dpi) Replicate_R2 | Mock-infected (4dpi) Replicate_R3 | SARS-CoV-2-infected (4dpi) Replicate_R1 | SARS-CoV-2-infected (4dpi) Replicate_R2 | SARS-CoV-2-infected (4dpi) Replicate_R3 | SARS-CoV-2-(14dpi) Replicate_R1 | SARS-CoV-2-infected (14dpi) Replicate_R2 | SARS-CoV-2-infected (14dpi) Replicate_R3 | Abundance Ratio: SARS-CoV-2-infected (4dpi) / Mock-infected (4dpi) | P-Value: SARS-CoV-2-infected (4dpi) / Mock-infected (4dpi) | Abundance Ratio: SARS-CoV-2-infected (14dpi) / Mock-infected (4dpi) | P-Value: SARS-CoV-2-infected (14dpi) / Mock-infected (4dpi) | Abundance Ratio: SARS-CoV-2-infected (14dpi) / SARS-CoV-2-infected (4dpi) | P-Value: SARS-CoV-2-infected (14dpi) / SARS-CoV-2-infected (4dpi) |
|---|---|---|---|---|---|---|---|---|---|---|---|---|---|---|---|---|---|---|---|---|---|---|---|---|---|---|
| A0A1U7QSU9 | Core histone macro-H2A OS=Mesocricetus auratus OX=10036 GN=LOC101833610 PE=4 SV=1 | 15.772 | 11 | 5 | 15 | 5 | 369 | 39.3 | 9.83 | 29.13 | 5 | 127.9 | 155.5 | 157.6 | 118.1 | 153 | 145.5 | 128.2 | 169.8 | 152.7 | 0.923 | 0.599870961 | 1.021995465 | 0.844515368 | 1.081853096 | 0.518998261 |
| A0A1U8D127 | 40S ribosomal protein S2 OS=Mesocricetus auratus OX=10036 GN=LOC101834330 PE=3 SV=1 | 27.301 | 21 | 6 | 27 | 6 | 293 | 31.3 | 10.13 | 70.15 | 6 | 635.6 | 554.6 | 584 | 571.5 | 500 | 509.3 | 571.7 | 525.1 | 526.2 | 0.899 | 0.119523878 | 0.914778492 | 0.160419061 | 1.026695344 | 0.635613079 |
| A0A1U7R3R6 | H-2 class II histocompatibility antigen, E-Q beta chain-like OS=Mesocricetus auratus OX=10036 GN=LOC101834334 PE=4 SV=1 | 5.092 | 18 | 2 | 4 | 1 | 181 | 21.1 | 8.47 | 8.65 | 2 | 0 | 0 | 60.8 | 0 | 0 | 116.4 | 0 | 0 | 45.1 | 1.916 | 0.700396316 | 0.741776316 | 0.846639278 | 0.387457045 | 0.61367298 |
| A0A1U7QC43 | cytochrome b-c1 complex subunit 2, mitochondrial OS=Mesocricetus auratus OX=10036 GN=LOC101834409 PE=3 SV=1 | 13.285 | 9 | 4 | 6 | 4 | 453 | 48.5 | 9.06 | 20.41 | 4 | 46 | 143.8 | 99 | 33 | 150.9 | 93.4 | 40.1 | 121.8 | 83.7 | 1.05 | 0.935254447 | 0.850415512 | 0.716298241 | 0.885683375 | 0.812681133 |
| A0A1U7QV99 | Cytochrome c oxidase polypeptide Vb OS=Mesocricetus auratus OX=10036 GN=LOC101834557 PE=3 SV=1 | 10.994 | 29 | 3 | 17 | 3 | 129 | 13.8 | 7.77 | 43.59 | 3 | 297 | 262.1 | 654.5 | 287 | 262.7 | 604.5 | 250.3 | 261.4 | 533.1 | 0.967 | 0.911359898 | 0.86090969 | 0.737761497 | 0.905215734 | 0.812648761 |
| A0A1U7RE76 | H-2 class II histocompatibility antigen, A-U alpha chain OS=Mesocricetus auratus OX=10036 GN=LOC101834602 PE=3 SV=1 | 5.198 | 5 | 1 | 5 | 1 | 256 | 28.2 | 4.79 | 24.2 | 1 | 90.6 | 30.8 | 88.2 | 135.5 | 39.4 | 126.8 | 100.8 | 27.9 | 118.1 | 1.437 | 0.454121626 | 1.177480916 | 0.734630577 | 0.818031157 | 0.680824566 |
| A0A1U7Q2G3 | guanine nucleotide-binding protein G(S) subunit alpha OS=Mesocricetus auratus OX=10036 GN=Gnas PE=3 SV=1 | 16.551 | 8 | 3 | 16 | 2 | 394 | 45.6 | 5.96 | 39.86 | 3 | 175 | 243.3 | 209.2 | 156.2 | 213.3 | 181 | 140 | 249.5 | 162.9 | 0.877 | 0.376543632 | 0.880318725 | 0.560934213 | 1.003451408 | 0.987516276 |
| A0A1U7RGM3 | rano class II histocompatibility antigen, A beta chain-like OS=Mesocricetus auratus OX=10036 GN=LOC101834868 PE=3 SV=1 | 11.736 | 16 | 3 | 18 | 2 | 263 | 29.6 | 7.21 | 41.21 | 3 | 246 | 282.6 | 282.6 | 353.4 | 392.7 | 367.8 | 234 | 265.4 | 282.4 | 1.375 | 0.003873255 | 0.963757396 | 0.62857068 | 0.701858336 | 0.004266341 |
| A0A1U7QXQ3 | 40S ribosomal protein S7 OS=Mesocricetus auratus OX=10036 GN=LOC101835445 PE=3 SV=1 | 29.609 | 26 | 6 | 32 | 6 | 194 | 22.1 | 10.1 | 80.58 | 6 | 24.4 | 0 | 0 | 27.4 | 0 | 0 | 30.7 | 0 | 0 | 1.123 | 0.938809602 | 1.258196721 | 0.880531492 | 1.120437956 | 0.939980312 |
| A0A1U8BR26 | leukocyte elastase inhibitor OS=Mesocricetus auratus OX=10036 GN=Serpinb1 PE=3 SV=1 | 54.511 | 30 | 12 | 76 | 12 | 379 | 42.7 | 6.21 | 203.84 | 12 | 1621 | 2949.3 | 2163.9 | 2241.9 | 2958.6 | 2950.6 | 2747.4 | 2249.1 | 1968.6 | 1.022 | 0.366775778 | 1.034287666 | 0.873704989 | 0.854498166 | 0.296021445 |
| A0A1U8CAZ6 | pulmonary surfactant-associated protein D isoform X3 OS=Mesocricetus auratus OX=10036 GN=Sftpd PE=3 SV=1 | 6.614 | 4 | 1 | 9 | 1 | 374 | 37.4 | 7.65 | 37.22 | 1 | 263.6 | 256.8 | 245 | 171.7 | 193.9 | 175.8 | 189.8 | 180.2 | 191.5 | 0.675 | 0.001264941 | 0.733603345 | 0.001004987 | 1.03712597 | 0.446993861 |
| A0A1U8CV72 | matrin-3 isoform X2 OS=Mesocricetus auratus OX=10036 GN=Matr3 PE=4 SV=1 | 4.553 | 3 | 1 | 2 | 1 | 793 | 88.2 | 5.76 | 11.51 | 1 | 39.3 | 0 | 0 | 33.5 | 0 | 0 | 28.5 | 0 | 0 | 0.852 | 0.916113278 | 0.72519084 | 0.835841998 | 0.850746269 | 0.915102076 |
| A0A1U7QNV9 | tumor-associated calcium signal transducer 2 OS=Mesocricetus auratus OX=10036 GN=Tacstd2 PE=3 SV=1 | 2.643 | 3 | 1 | 5 | 1 | 318 | 35.9 | 8.7 | 15.78 | 1 | 267.9 | 491.6 | 589.7 | 176.8 | 302.1 | 407 | 217 | 421.4 | 484.1 | 0.66 | 0.262157821 | 0.831974503 | 0.578305967 | 1.267073033 | 0.49403215 |
| Q9ERB9 | Peptidyl-prolyl cis-trans isomerase (Fragment) OS=Mesocricetus auratus OX=10036 PE=2 SV=1 | 35.642 | 81 | 6 | 66 | 1 | 70 | 7.8 | 8.18 | 186.51 | 6 | 155.3 | 0 | 56.6 | 63.8 | 0 | 22.3 | 120.3 | 0 | 43.1 | 0.402 | 0.462883445 | 0.771118452 | 0.793082255 | 1.897793264 | 0.563290513 |
| A0A1U8BXG1 | SEC14-like protein 4 OS=Mesocricetus auratus OX=10036 GN=LOC101836694 PE=4 SV=1 | 27.111 | 16 | 6 | 43 | 4 | 412 | 47 | 6.98 | 124.86 | 6 | 745.8 | 797.9 | 297.3 | 599.5 | 654.1 | 205.9 | 639.3 | 688.1 | 257.7 | 0.82 | 0.582308876 | 0.860999457 | 0.704801066 | 1.086056869 | 0.841375214 |
| A0A1U7R822 | Dynein light chain OS=Mesocricetus auratus OX=10036 GN=LOC101836744 PE=3 SV=1 | 16.23 | 27 | 5 | 22 | 5 | 89 | 10.4 | 7.4 | 69.59 | 5 | 343.9 | 268.5 | 389.9 | 320.8 | 239.2 | 325.7 | 375.8 | 872.2 | 412.2 | 0.933 | 0.440283471 | 1.656390302 | 0.301960296 | 1.874449588 | 0.245299686 |
| A0A1U7RGH9 | Histone H2A OS=Mesocricetus auratus OX=10036 GN=LOC101837114 PE=3 SV=1 | 25.751 | 31 | 5 | 55 | 3 | 128 | 13.5 | 10.58 | 144.45 | 5 | 438.4 | 297.7 | 467 | 693.8 | 441.8 | 737.1 | 352.4 | 256.3 | 388 | 1.583 | 0.12077642 | 0.828443188 | 0.356759945 | 0.532226197 | 0.069892775 |
| A0A1U8BIA6 | immunity-related GTPase family M protein 1-like isoform X1 OS=Mesocricetus auratus OX=10036 GN=LOC101837226 PE=3 SV=1 | 8.106 | 8 | 2 | 4 | 2 | 438 | 49.9 | 5.35 | 15.16 | 2 | 27 | 0 | 29.1 | 67.1 | 0 | 35.3 | 35.1 | 0 | 30 | 1.735 | 0.527018178 | 1.160427807 | 0.845494898 | 0.635742188 | 0.613604149 |
| A0A1U7QPZ7 | RNA transcription, translation and transport factor protein OS=Mesocricetus auratus OX=10036 GN=CUNH14orf166 PE=3 SV=1 | 2.708 | 3 | 1 | 10 | 1 | 244 | 28.1 | 6.65 | 29.63 | 1 | 61.5 | 71.7 | 81.9 | 71 | 75.8 | 74 | 54.1 | 73.5 | 70.7 | 1.039 | 0.780622423 | 0.921896792 | 0.543587054 | 0.898097826 | 0.340572571 |
| A0A1U7R3I2 | serine/threonine-protein phosphatase 2A 65 kDa regulatory subunit A alpha isoform OS=Mesocricetus auratus OX=10036 GN=Ppp2r1a PE=4 SV=1 | 23.62 | 15 | 7 | 23 | 7 | 589 | 65.3 | 5.11 | 65.71 | 7 | 81.5 | 407.7 | 320.1 | 74.7 | 387.4 | 309 | 74.9 | 393.8 | 284 | 0.965 | 0.929586505 | 0.930063017 | 0.895689171 | 0.976137985 | 0.965314328 |
| A0A1U7QIW2 | GTP-binding nuclear protein Ran OS=Mesocricetus auratus OX=10036 GN=LOC101837891 PE=3 SV=1 | 25.64 | 20 | 5 | 20 | 1 | 216 | 24.4 | 6.8 | 58.98 | 5 | 997.2 | 286.5 | 429.3 | 1129.4 | 263.5 | 465.1 | 932.2 | 250.2 | 421.5 | 1.083 | 0.894023627 | 0.936310566 | 0.908910757 | 0.863240043 | 0.812032794 |
| A0A1U8CVF2 | ATP synthase F1 subunit epsilon OS=Mesocricetus auratus OX=10036 GN=Atpif1 PE=3 SV=1 | 15.514 | 34 | 4 | 23 | 4 | 108 | 12.2 | 9.7 | 24.72 | 4 | 2019.4 | 1593.2 | 1644.6 | 1369.8 | 986.3 | 1091.7 | 1651.6 | 1208.4 | 1275.2 | 0.666 | 0.027890069 | 0.786578407 | 0.124085924 | 1.199373514 | 0.27238047 |
| A0A1U7RG56 | interferon-inducible GTPase 1-like OS=Mesocricetus auratus OX=10036 GN=LOC101838193 PE=3 SV=1 | 1.688 | 2 | 1 | 3 | 1 | 418 | 47.6 | 7.21 | 3.71 | 1 | 70.5 | 76.9 | 62.3 | 146.2 | 133.7 | 137.6 | 65.2 | 63 | 105.5 | 2.074 | 0.000274304 | 1.114449213 | 0.627570653 | 0.559760479 | 0.039610581 |
| A0A1U8D1U9 | GTP-binding protein Rheb OS=Mesocricetus auratus OX=10036 GN=Rheb PE=4 SV=1 | 1.979 | 10 | 1 | 2 | 1 | 79 | 8.6 | 6.02 | 5.11 | 1 | 30.3 | 0 | 29 | 39.4 | 0 | 38.3 | 27.7 | 0 | 26.3 | 1.309 | 0.727039822 | 0.910623946 | 0.901371217 | 0.694980695 | 0.645887451 |
| A0A1U8CG58 | 60S ribosomal protein L35 OS=Mesocricetus auratus OX=10036 GN=Rpl35 PE=3 SV=1 | 4.844 | 11 | 1 | 9 | 1 | 123 | 14.5 | 11.3 | 15.44 | 1 | 103 | 83 | 59.1 | 138.6 | 118.4 | 82.9 | 145.2 | 99.6 | 91 | 1.418 | 0.204695162 | 1.37005304 | 0.229708004 | 0.987937629 | 0.956243036 |
| A0A1U8BYF8 | cytochrome c oxidase subunit 4 isoform 1, mitochondrial OS=Mesocricetus auratus OX=10036 GN=LOC101839655 PE=3 SV=1 | 15.838 | 25 | 4 | 12 | 4 | 169 | 19.6 | 9.36 | 28.82 | 4 | 247.3 | 188.2 | 372.9 | 232.5 | 161.8 | 450.9 | 222 | 144.8 | 348.1 | 0.938 | 0.911674332 | 0.884339436 | 0.718404795 | 0.845835305 | 0.703733024 |
| A0A1U7R6S0 | splicing factor 3B subunit 2-like OS=Mesocricetus auratus OX=10036 GN=LOC101839719 PE=4 SV=1 | 5.914 | 5 | 2 | 12 | 2 | 315 | 36.4 | 8.6 | 11.72 | 2 | 44.2 | 74.8 | 37.5 | 37.8 | 85.5 | 44.1 | 43.2 | 83.9 | 32.2 | 1.143 | 0.857203827 | 1.017891374 | 0.964264749 | 0.951612903 | 0.907013872 |
| A0A1U7R5X4 | Riboflavin kinase OS=Mesocricetus auratus OX=10036 GN=Rfk PE=4 SV=1 | 3.733 | 7 | 1 | 2 | 1 | 155 | 17.3 | 7.05 | 3.11 | 1 | 61.1 | 63 | 0 | 75.8 | 94.4 | 0 | 92.5 | 78.2 | 0 | 1.363 | 0.689756941 | 1.375503626 | 0.68578382 | 1.00293772 | 0.996931955 |
| A0A1U7Q3R5 | Cytochrome b-c1 complex subunit Rieske, mitochondrial OS=Mesocricetus auratus OX=10036 GN=LOC101839974 PE=3 SV=1 | 4.097 | 6 | 2 | 8 | 2 | 273 | 29.4 | 8.41 | 5.1 | 2 | 840.7 | 242.6 | 508.4 | 643.1 | 778.4 | 409.7 | 1416.9 | 814.3 | 521.3 | 1.378 | 0.718871319 | 1.729283156 | 0.296831953 | 1.503112713 | 0.369149677 |
| A0A1U7QHC7 | Eukaryotic translation initiation factor 5A OS=Mesocricetus auratus OX=10036 GN=LOC101840102 PE=3 SV=1 | 21.583 | 36 | 5 | 34 | 5 | 154 | 16.9 | 5.4 | 119.92 | 5 | 394.1 | 254.7 | 609.4 | 489.2 | 315.6 | 751.5 | 386.3 | 249.2 | 604 | 1.241 | 0.577196604 | 0.985137498 | 0.967990912 | 0.796440275 | 0.554826236 |
| A0A1U7R6P8 | 1,4-alpha-glucan branching enzyme OS=Mesocricetus auratus OX=10036 GN=Gbe1 PE=3 SV=1 | 6.332 | 4 | 2 | 4 | 2 | 696 | 79.8 | 6.4 | 13.63 | 2 | 0 | 0 | 46.2 | 0 | 0 | 34.9 | 0 | 0 | 41.5 | 0.755 | 0.85545094 | 0.898268398 | 0.943345871 | 1.189111748 | 0.909156901 |
| P70686 | Glutathione S-transferase OS=Mesocricetus auratus OX=10036 GN=GSTYc PE=2 SV=1 | 52.788 | 43 | 12 | 97 | 8 | 221 | 25.5 | 9.04 | 210.29 | 12 | 4119.1 | 4700.3 | 5732.8 | 3171.6 | 3173.9 | 3704.3 | 5821 | 7596.6 | 6140.7 | 0.737 | 0.071822204 | 1.34400984 | 0.083314341 | 1.946138232 | 0.020240065 |
| A0A1U7RDX1 | histone H1.5 OS=Mesocricetus auratus OX=10036 GN=LOC101840609 PE=3 SV=1 | 43.922 | 32 | 9 | 49 | 2 | 221 | 22.4 | 10.92 | 95.65 | 9 | 310.7 | 345.8 | 397.9 | 215.2 | 247.1 | 334.5 | 251.7 | 240.8 | 318.8 | 0.704 | 0.128775592 | 0.769442337 | 0.082622509 | 1.018197791 | 0.916977576 |
| A0A1U7Q534 | interferon-induced transmembrane protein 3 isoform X1 OS=Mesocricetus auratus OX=10036 GN=LOC101840863 PE=3 SV=1 | 6.318 | 14 | 1 | 8 | 1 | 145 | 15.8 | 7.84 | 35.04 | 1 | 79.2 | 121.6 | 100.7 | 96.8 | 192.7 | 138.8 | 60.8 | 101.8 | 88.4 | 1.585 | 0.265474958 | 0.832504146 | 0.382920078 | 0.586037824 | 0.154892013 |
| A0A1U8CAE9 | Voltage-dependent anion-selective channel protein 1 OS=Mesocricetus auratus OX=10036 GN=LOC101841110 PE=3 SV=1 | 3.05 | 3 | 1 | 9 | 1 | 280 | 30.2 | 8.44 | 16.52 | 1 | 170.2 | 245.4 | 222.8 | 218.4 | 310.7 | 268.2 | 179 | 292.7 | 251.7 | 1.266 | 0.204362197 | 1.133145363 | 0.523277199 | 0.907312179 | 0.595646486 |
| A0A1U7QPC1 | prostaglandin-E(2) 9-reductase-like isoform X2 OS=Mesocricetus auratus OX=10036 GN=LOC101841267 PE=3 SV=1 | 36.789 | 28 | 8 | 36 | 8 | 351 | 40.3 | 7.15 | 75.46 | 8 | 441.1 | 577.2 | 487.4 | 360.9 | 499.4 | 419.8 | 393.1 | 517.7 | 422 | 0.865 | 0.254891171 | 0.885169689 | 0.353237722 | 1.041168659 | 0.765590836 |
| A0A1U7QU89 | OX-2 membrane glycoprotein isoform X1 OS=Mesocricetus auratus OX=10036 GN=LOC101841355 PE=4 SV=1 | 1.348 | 3 | 1 | 1 | 1 | 278 | 31.3 | 8.66 | 0 | 1 | 120.8 | 0 | 0 | 95.8 | 0 | 0 | 101.7 | 0 | 0 | 0.793 | 0.879434363 | 0.841887417 | 0.909720036 | 1.061586639 | 0.968347205 |
| A0A1U7QVT6 | 60S ribosomal protein L7a OS=Mesocricetus auratus OX=10036 GN=LOC101841383 PE=3 SV=1 | 24.084 | 26 | 7 | 29 | 7 | 266 | 30 | 10.55 | 65.05 | 7 | 545 | 647.7 | 753.6 | 484 | 573.8 | 716.2 | 426.7 | 530.7 | 766 | 0.888 | 0.560728554 | 0.885475004 | 0.56714696 | 0.971476888 | 0.896778609 |
| A0A1U8C3E7 | Cytochrome b-c1 complex subunit 6 OS=Mesocricetus auratus OX=10036 GN=LOC101841714 PE=3 SV=1 | 12.046 | 30 | 3 | 13 | 3 | 90 | 10.6 | 4.81 | 37.62 | 3 | 655.8 | 409.1 | 231.1 | 599.7 | 452.1 | 227.2 | 513.3 | 336.8 | 175 | 0.985 | 0.974115024 | 0.790972222 | 0.597889425 | 0.801485536 | 0.593157565 |
| A0A1U7QSF2 | cytochrome c oxidase subunit 7A2, mitochondrial OS=Mesocricetus auratus OX=10036 GN=LOC101841805 PE=3 SV=1 | 3.481 | 16 | 1 | 2 | 1 | 83 | 9.4 | 10.27 | 5.68 | 1 | 0 | 0 | 53.5 | 0 | 0 | 40 | 0 | 0 | 50.4 | 0.748 | 0.850451401 | 0.942056075 | 0.968386367 | 1.26 | 0.879815197 |
| A0A1U7R051 | myeloid cell nuclear differentiation antigen OS=Mesocricetus auratus OX=10036 GN=Mnda PE=3 SV=1 | 9.532 | 8 | 3 | 13 | 3 | 429 | 48.1 | 9.41 | 48.61 | 3 | 190.4 | 317.6 | 158.9 | 272.7 | 371.8 | 216.2 | 181.6 | 281.5 | 154.6 | 1.253 | 0.38650693 | 0.926225821 | 0.805048777 | 0.717671663 | 0.247760867 |
| A0A1U7QIZ4 | Eukaryotic translation initiation factor 3 subunit J OS=Mesocricetus auratus OX=10036 GN=Eif3j PE=3 SV=1 | 2.352 | 5 | 1 | 1 | 1 | 261 | 29.3 | 4.81 | 0 | 1 | 45.4 | 0 | 0 | 34.9 | 0 | 0 | 29.2 | 0 | 0 | 0.768 | 0.863992687 | 0.643171806 | 0.781460688 | 0.836676218 | 0.90653755 |
| A0A1U7QQ66 | actin, alpha skeletal muscle OS=Mesocricetus auratus OX=10036 GN=LOC101842364 PE=3 SV=1 | 21.478 | 31 | 4 | 46 | 1 | 150 | 16.5 | 7.81 | 97.44 | 4 | 47.1 | 105.2 | 94 | 43.1 | 92.9 | 79.3 | 36.6 | 71.2 | 78.3 | 0.899 | 0.679578716 | 0.755582623 | 0.417366117 | 0.86437529 | 0.64710512 |
| P70116 | Cytochrome b5 OS=Mesocricetus auratus OX=10036 GN=LOC101842475 PE=2 SV=1 | 29.45 | 37 | 6 | 31 | 6 | 171 | 19.2 | 7.12 | 111.22 | 6 | 288.8 | 306.8 | 370.8 | 289.8 | 275.1 | 359.2 | 294 | 272.8 | 346.8 | 0.969 | 0.71485496 | 0.945364238 | 0.62463112 | 0.988637593 | 0.92310841 |
| A0A1U7QFD2 | protein kinase C and casein kinase substrate in neurons protein 3 OS=Mesocricetus auratus OX=10036 GN=Pacsin3 PE=4 SV=1 | 2.684 | 2 | 1 | 5 | 1 | 427 | 48.7 | 6.2 | 16.01 | 1 | 122 | 83 | 139.9 | 121.7 | 67.8 | 113.9 | 95.6 | 77.4 | 143.3 | 0.817 | 0.59181339 | 0.917077414 | 0.731403873 | 1.042518128 | 0.87620238 |
| A0A1U7R8L4 | LDLR chaperone MESD OS=Mesocricetus auratus OX=10036 GN=Mesd PE=3 SV=1 | 6.171 | 9 | 1 | 9 | 1 | 226 | 25.1 | 6.33 | 29.29 | 1 | 87.4 | 0 | 0 | 88.5 | 0 | 0 | 81.6 | 0 | 0 | 1.012 | 0.993367406 | 0.933638444 | 0.963648281 | 0.922033898 | 0.957056891 |
| A0A1U8BUR8 | RNA-binding protein 14 isoform X1 OS=Mesocricetus auratus OX=10036 GN=LOC101843260 PE=4 SV=1 | 7.073 | 3 | 2 | 9 | 2 | 669 | 69.5 | 9.67 | 31.15 | 2 | 590 | 260.3 | 491.2 | 498.1 | 210.5 | 404.8 | 509.2 | 246.6 | 460.2 | 0.809 | 0.588710646 | 0.906448006 | 0.758325566 | 1.092150171 | 0.784516071 |
| A0A1U7R5N1 | CCT-beta OS=Mesocricetus auratus OX=10036 GN=Cct2 PE=3 SV=1 | 17.519 | 11 | 4 | 23 | 4 | 535 | 57.3 | 6.54 | 68.18 | 4 | 277.2 | 246.1 | 257.8 | 283.1 | 227.5 | 226.8 | 241.2 | 241 | 245.7 | 0.904 | 0.534704001 | 0.931890923 | 0.186739613 | 0.987116897 | 0.881011438 |
| P51581 | Cytochrome P450 2E1 OS=Mesocricetus auratus OX=10036 GN=CYP2E1 PE=2 SV=1 | 9.071 | 9 | 3 | 4 | 3 | 493 | 56.6 | 7.75 | 12.71 | 3 | 53.3 | 0 | 48 | 51.2 | 0 | 49.9 | 42.9 | 0 | 48.1 | 1 | 0.997908399 | 0.898321816 | 0.887651375 | 0.900098912 | 0.889453958 |
| A0A1U8BEB6 | alpha-1-inhibitor 3-like OS=Mesocricetus auratus OX=10036 GN=LOC101843823 PE=3 SV=1 | 22.739 | 6 | 7 | 19 | 6 | 1483 | 165.8 | 6.54 | 38.04 | 7 | 113.2 | 160.3 | 322.7 | 241.6 | 327.6 | 499.8 | 171.1 | 226 | 465.7 | 2.129 | 0.18861727 | 1.447165381 | 0.471155691 | 0.807109448 | 0.592624553 |
| A0A1U8BWE1 | Glutathione transferase OS=Mesocricetus auratus OX=10036 GN=LOC101843868 PE=3 SV=1 | 43.166 | 27 | 9 | 103 | 6 | 222 | 25.6 | 7.37 | 301.93 | 9 | 2962.4 | 1835.5 | 2889.5 | 2066.2 | 2253.7 | 1943.3 | 3047.2 | 3190.8 | 2707.3 | 0.702 | 0.321118294 | 1.163631397 | 0.373016537 | 1.428231575 | 0.009829512 |
| A0A1U8CMW2 | hereditary hemochromatosis protein homolog isoform X1 OS=Mesocricetus auratus OX=10036 GN=LOC101843916 PE=3 SV=1 | 4.732 | 6 | 2 | 6 | 2 | 335 | 37.8 | 6.98 | 18.23 | 2 | 284.3 | 275 | 113.3 | 273.1 | 273.3 | 108.7 | 219.8 | 232.6 | 73.7 | 0.96 | 0.943997192 | 0.782188522 | 0.552559678 | 0.803083499 | 0.59661799 |
| A0A1U8BJV1 | prothymosin alpha-like OS=Mesocricetus auratus OX=10036 GN=LOC101843929 PE=3 SV=1 | 33.974 | 15 | 5 | 19 | 5 | 117 | 13 | 4.01 | 59.37 | 5 | 307.7 | 222.5 | 168.1 | 317.6 | 216.3 | 193.4 | 271.2 | 191.6 | 169.1 | 1.032 | 0.870764002 | 0.904911929 | 0.688640412 | 0.868829919 | 0.554271281 |
| A0A1U7QEV1 | D-dopachrome decarboxylase OS=Mesocricetus auratus OX=10036 GN=LOC101844189 PE=3 SV=1 | 10.624 | 16 | 3 | 11 | 3 | 118 | 13.1 | 7.14 | 23.71 | 3 | 40.8 | 141.1 | 91.7 | 30.7 | 100.5 | 72.7 | 26.2 | 141.1 | 94.4 | 0.712 | 0.550894023 | 0.956505848 | 0.932847997 | 1.28347229 | 0.652689034 |
| A0A1U7RAC6 | 40S ribosomal protein S25 OS=Mesocricetus auratus OX=10036 GN=LOC101844694 PE=3 SV=1 | 10.525 | 27 | 4 | 13 | 1 | 125 | 13.7 | 10.05 | 34.96 | 4 | 391.2 | 378.6 | 161.3 | 405.5 | 353.2 | 199.5 | 365.6 | 337.2 | 170.1 | 0.935 | 0.930358975 | 0.937493288 | 0.850655371 | 0.910978919 | 0.759773297 |

**Supplementary Table 1: List of differentially expressed proteins identified across the 4dpi infected and 14dpi infected**

| Accession | Description | Sum PEP Score | Coverage [%] | # Peptides | # PSMs | # Unique Peptides | # AAs | MW [kDa] | calc. pI | Score Sequest HT: Sequest HT | # Peptides (by Search Engine): Sequest HT | Mock-infected (4dpi) Replicate_R1 | Mock-infected (4dpi) Replicate_R2 | Mock-infected (4dpi) Replicate_R3 | SARS-CoV-2-infected (4dpi) Replicate_R1 | SARS-CoV-2-infected (4dpi) Replicate_R2 | SARS-CoV-2-infected (4dpi) Replicate_R3 | SARS-CoV-2-(14dpi) Replicate_R1 | SARS-CoV-2-infected (14dpi) Replicate_R2 | SARS-CoV-2-infected (14dpi) Replicate_R3 | Abundance Ratio: SARS-CoV-2-infected (4dpi) / Mock-infected (4dpi) | P-Value: SARS-CoV-2-infected (4dpi) / Mock-infected (4dpi) | Abundance Ratio: SARS-CoV-2-infected (14dpi) / Mock-infected (4dpi) | P-Value: SARS-CoV-2-infected (14dpi) / Mock-infected (4dpi) | Abundance Ratio: SARS-CoV-2-infected (14dpi) / SARS-CoV-2-infected (4dpi) | P-Value: SARS-CoV-2-infected (14dpi) / SARS-CoV-2-infected (4dpi) |
|---|---|---|---|---|---|---|---|---|---|---|---|---|---|---|---|---|---|---|---|---|---|---|---|---|---|
| A0A1U8CUG0 | pregnancy zone protein-like OS=Mesocricetus auratus OX=10036 GN=LOC101844701 PE=3 SV=1 | 191.257 | 26 | 33 | 271 | 32 | 1481 | 165.2 | 6.42 | 959.6 | 33 | 4432.8 | 4199.9 | 3817 | 7353.1 | 6286.7 | 5664.9 | 4886.6 | 4311.7 | 4129.1 | 1.505 | 0.031672374 | 1.070499691 | 0.37360289 | 0.690370739 | 0.038850051 |
| A0A1U8CFS8 | Unspecific monooxygenase OS=Mesocricetus auratus OX=10036 GN=LOC101844759 PE=3 SV=1 | 15.127 | 8 | 4 | 21 | 3 | 491 | 56.2 | 7.09 | 89.82 | 4 | 390.8 | 442.3 | 370.9 | 354.5 | 369.1 | 351.9 | 368.9 | 489.5 | 363.7 | 0.949 | 0.175651367 | 1.015033223 | 0.904515003 | 1.136308694 | 0.356203139 |
| A0A1U8C2T0 | 60 kDa chaperonin OS=Mesocricetus auratus OX=10036 GN=LOC106021236 PE=3 SV=1 | 8.173 | 5 | 4 | 7 | 1 | 614 | 66.3 | 6.29 | 13.03 | 4 | 103.4 | 127.3 | 123.4 | 106.3 | 126.1 | 129.2 | 110.5 | 107 | 100.2 | 1.047 | 0.820303961 | 0.89720418 | 0.238017369 | 0.878595133 | 0.16724893 |
| A0A1U8C3E5 | cGMP-dependent protein kinase 1-like isoform X2 OS=Mesocricetus auratus OX=10036 GN=LOC106021271 PE=4 SV=1 | 1.533 | 6 | 1 | 1 | 1 | 212 | 24.2 | 6.37 | 0 | 1 | 0 | 95.1 | 0 | 0 | 61.9 | 0 | 0 | 98.8 | 0 | 0.651 | 0.786632535 | 1.038906414 | 0.979769023 | 1.596122779 | 0.770285307 |
| A0A1U8C4D7 | LOW QUALITY PROTEIN: transformer-2 protein homolog alpha-like OS=Mesocricetus auratus OX=10036 GN=LOC106021332 PE=4 SV=1 | 3.48 | 6 | 1 | 3 | 1 | 185 | 20.9 | 9.7 | 7.46 | 1 | 103.4 | 138.6 | 0 | 92.4 | 149.3 | 0 | 55.1 | 102.1 | 0 | 0.981 | 0.998754055 | 0.649586777 | 0.611933987 | 0.650393049 | 0.624119193 |
| A0A1U8C0S7 | Nucleoside diphosphate kinase OS=Mesocricetus auratus OX=10036 GN=LOC106021590 PE=3 SV=1 | 22.218 | 22 | 6 | 37 | 3 | 249 | 27.6 | 9.6 | 81.24 | 6 | 930 | 1217.3 | 1080.1 | 1374.8 | 1472.1 | 1607.4 | 1236 | 1242.8 | 1455.7 | 1.255 | 0.020162421 | 1.219092768 | 0.09996916 | 0.883303774 | 0.154497655 |
| A0A1U8CRU6 | spectrin alpha chain, erythrocytic 1-like OS=Mesocricetus auratus OX=10036 GN=LOC106022839 PE=4 SV=1 | 7.511 | 4 | 1 | 8 | 1 | 280 | 32.5 | 5.03 | 15.88 | 1 | 106.9 | 0 | 0 | 70.9 | 0 | 0 | 89.8 | 0 | 0 | 0.663 | 0.7948825 | 0.840037418 | 0.908591629 | 1.266572638 | 0.877217436 |
| A0A1U8D3Q7 | sulfide:quinone oxidoreductase, mitochondrial-like OS=Mesocricetus auratus OX=10036 GN=LOC106022852 PE=4 SV=1 | 5.317 | 10 | 1 | 3 | 1 | 147 | 15.6 | 9.14 | 8.75 | 1 | 42.4 | 0 | 50.5 | 27.7 | 0 | 33.4 | 26.2 | 0 | 45.6 | 0.657 | 0.606476965 | 0.772874058 | 0.749130116 | 1.17512275 | 0.842482789 |
| A0A1U7QAK6 | Lysyl oxidase homolog OS=Mesocricetus auratus OX=10036 GN=Loxl1 PE=3 SV=1 | 2.75 | 2 | 1 | 2 | 1 | 606 | 66.5 | 7.06 | 3.82 | 1 | 52.8 | 0 | 0 | 47.8 | 0 | 0 | 43 | 0 | 0 | 0.906 | 0.947433777 | 0.814393939 | 0.892795533 | 0.89958159 | 0.944110844 |
| A0A1U7Q7E8 | lipoma-preferred partner homolog OS=Mesocricetus auratus OX=10036 GN=LOC101828273 PE=4 SV=1 | 26.563 | 23 | 5 | 26 | 5 | 388 | 41.3 | 7.28 | 91.5 | 5 | 153.8 | 184 | 147.9 | 89.3 | 131.9 | 93.6 | 122 | 154.7 | 118.7 | 0.603 | 0.03319672 | 0.814082767 | 0.133714136 | 1.256035578 | 0.206691125 |
| A0A1U8BRI1 | leucine-rich repeat and calponin homology domain-containing protein 3 isoform X1 OS=Mesocricetus auratus OX=10036 GN=Lrch3 PE=4 SV=1 | 6.298 | 3 | 2 | 6 | 2 | 794 | 87.9 | 6.89 | 12.43 | 2 | 0 | 41.2 | 0 | 0 | 30.5 | 0 | 0 | 42 | 0 | 0.741 | 0.845675926 | 1.019417476 | 0.989802438 | 1.37704918 | 0.836495088 |
| A0A1U7QYS6 | leucine-rich repeat-containing protein 59 OS=Mesocricetus auratus OX=10036 GN=Lrrc59 PE=4 SV=1 | 25.95 | 25 | 5 | 18 | 5 | 307 | 35 | 9.57 | 53.67 | 5 | 283.5 | 171.6 | 310 | 192.6 | 103 | 212.9 | 224.9 | 126.6 | 261.4 | 0.687 | 0.193279425 | 0.801071755 | 0.434672824 | 1.205309735 | 0.544977338 |
| A0A1U8CDR4 | leucine-rich repeat flightless-interacting protein 1 isoform X14 OS=Mesocricetus auratus OX=10036 GN=Lrrfip1 PE=3 SV=1 | 1.552 | 2 | 1 | 1 | 1 | 805 | 88.4 | 4.87 | 0 | 1 | 32.8 | 0 | 0 | 43.4 | 0 | 0 | 35 | 0 | 0 | 1.325 | 0.855675747 | 1.067073171 | 0.965625258 | 0.806451613 | 0.887846265 |
| A0A1U7Q6I1 | lymphocyte-specific protein 1 isoform X1 OS=Mesocricetus auratus OX=10036 GN=Lsp1 PE=4 SV=1 | 7.258 | 8 | 2 | 4 | 2 | 330 | 36.6 | 4.72 | 13.12 | 2 | 0 | 123.5 | 0 | 0 | 110.7 | 0 | 0 | 108.9 | 0 | 0.896 | 0.942230481 | 0.881781377 | 0.933669705 | 0.983739837 | 0.991306741 |
| A0A1U7Q4A6 | Leukotriene A(4) hydrolase OS=Mesocricetus auratus OX=10036 GN=Lta4h PE=3 SV=1 | 20.296 | 9 | 5 | 18 | 5 | 611 | 69.1 | 6.29 | 54.75 | 5 | 280.5 | 328.6 | 224.8 | 281.1 | 328.5 | 210.1 | 258.2 | 336.8 | 220.9 | 1.002 | 0.922482224 | 0.978414678 | 0.901462416 | 0.995364158 | 0.980407785 |
| A0A1U7QDZ7 | leukotriene C4 synthase isoform X2 OS=Mesocricetus auratus OX=10036 GN=Ltc4s PE=3 SV=1 | 4.786 | 11 | 1 | 8 | 1 | 150 | 16.7 | 9.88 | 29.65 | 1 | 82 | 83.9 | 113.3 | 82.3 | 83.1 | 96.8 | 68.4 | 72.4 | 95.6 | 1.004 | 0.648903658 | 0.846704871 | 0.342708487 | 0.901601831 | 0.438092332 |
| A0A1U7Q4A8 | Lumican OS=Mesocricetus auratus OX=10036 GN=Lum PE=3 SV=1 | 35.338 | 31 | 8 | 71 | 8 | 338 | 38.2 | 6.54 | 228.01 | 8 | 937.9 | 888.4 | 712 | 854.2 | 693 | 626.6 | 811.5 | 897.6 | 608.5 | 0.88 | 0.275473648 | 0.913052043 | 0.541022863 | 1.06615144 | 0.684365058 |
| A0A1U7R588 | acyl-protein thioesterase 2 OS=Mesocricetus auratus OX=10036 GN=Lypla2 PE=3 SV=1 | 1.477 | 3 | 1 | 2 | 1 | 231 | 24.8 | 7.23 | 0 | 1 | 150.4 | 132.6 | 0 | 192.5 | 117.1 | 0 | 166.5 | 116 | 0 | 1.063 | 0.909823147 | 0.998233216 | 0.998173049 | 0.9124677 | 0.909552445 |
| A0A1U7QPM9 | Membrane-associated guanylate kinase inverted 3 OS=Mesocricetus auratus OX=10036 GN=Magi3 PE=3 SV=1 | 7.027 | 3 | 4 | 5 | 4 | 1469 | 160.9 | 7.77 | 8.1 | 4 | 149.4 | 47.4 | 48.4 | 103.2 | 42.9 | 49 | 131.5 | 25.1 | 22.8 | 0.802 | 0.695163391 | 0.731647635 | 0.679479234 | 0.919528447 | 0.905582897 |
| A0A1U7QQP6 | Alpha-mannosidase OS=Mesocricetus auratus OX=10036 GN=Man2a1 PE=3 SV=1 | 3.374 | 1 | 1 | 1 | 1 | 1148 | 131 | 7.65 | 4.79 | 1 | 0 | 0 | 47.9 | 0 | 0 | 54.4 | 0 | 0 | 45.5 | 1.136 | 0.932920832 | 0.949895616 | 0.972766198 | 0.836397059 | 0.906366726 |
| A0A1U7R6P4 | Amine oxidase OS=Mesocricetus auratus OX=10036 GN=Maob PE=3 SV=1 | 12.266 | 10 | 4 | 14 | 4 | 504 | 57 | 7.72 | 20.09 | 4 | 53.4 | 0 | 114.7 | 51.7 | 0 | 81.3 | 64.6 | 0 | 74.1 | 0.829 | 0.789786228 | 0.825104105 | 0.821976887 | 1.042857143 | 0.957186848 |
| A0A1U7QVL0 | dual specificity mitogen-activated protein kinase kinase 1 OS=Mesocricetus auratus OX=10036 GN=Map2k1 PE=3 SV=1 | 8.97 | 3 | 2 | 11 | 2 | 393 | 43.4 | 6.62 | 22.35 | 2 | 343.4 | 285.4 | 396.7 | 275.8 | 260.5 | 321.5 | 305.1 | 256.7 | 369.7 | 0.832 | 0.223041651 | 0.908337396 | 0.532096789 | 1.085917463 | 0.557319417 |
| A0A1U7R0S2 | dual specificity mitogen-activated protein kinase kinase 2 OS=Mesocricetus auratus OX=10036 GN=Map2k2 PE=3 SV=1 | 3.632 | 3 | 1 | 2 | 1 | 400 | 44.2 | 6.86 | 6.67 | 1 | 58.2 | 70.1 | 0 | 38 | 64.1 | 0 | 52.3 | 79.6 | 0 | 0.773 | 0.775308244 | 1.028059236 | 0.971760614 | 1.291870715 | 0.756892182 |
| A0A1U8C9I0 | Microtubule-associated protein OS=Mesocricetus auratus OX=10036 GN=Map4 PE=4 SV=1 | 39.076 | 20 | 9 | 25 | 9 | 1034 | 108.2 | 4.75 | 84.43 | 9 | 193.5 | 78.2 | 89.5 | 162.2 | 81.8 | 87.7 | 191.9 | 63.7 | 82 | 0.98 | 0.838507189 | 0.934662237 | 0.891880364 | 1.017787157 | 0.969381016 |
| A0A1U7QJT2 | mitogen-activated protein kinase kinase kinase kinase 4 isoform X21 OS=Mesocricetus auratus OX=10036 GN=Map4k4 PE=4 SV=1 | 7.151 | 2 | 3 | 13 | 3 | 1164 | 133.2 | 7.64 | 21.31 | 3 | 147.6 | 255 | 164.3 | 100.5 | 188.7 | 133.7 | 142.4 | 227.8 | 145.7 | 0.766 | 0.321948951 | 0.910037044 | 0.716541559 | 1.219910144 | 0.460373459 |
| A0A1U7QEZ9 | Mitogen-activated protein kinase OS=Mesocricetus auratus OX=10036 GN=Mapk1 PE=3 SV=1 | 14.071 | 16 | 5 | 10 | 3 | 358 | 41.2 | 6.98 | 23.9 | 5 | 191.3 | 126.5 | 131.9 | 170.7 | 117.3 | 130.4 | 178 | 113.7 | 141.5 | 0.982 | 0.712538563 | 0.963308873 | 0.853364278 | 1.035372849 | 0.850998697 |
| A0A1U7Q3C8 | Microtubule-associated protein RP/EB family member 2 OS=Mesocricetus auratus OX=10036 GN=Mapre2 PE=3 SV=1 | 10.895 | 9 | 3 | 12 | 3 | 326 | 36.9 | 5.38 | 34.64 | 3 | 178.1 | 392.6 | 238.2 | 186.5 | 360.2 | 199.9 | 145.6 | 361 | 205.1 | 0.926 | 0.818846592 | 0.879836815 | 0.738663587 | 0.953254755 | 0.89796728 |
| A0A1U7QZH4 | mitochondrial amidoxime reducing component 2 OS=Mesocricetus auratus OX=10036 GN=Marc2 PE=4 SV=1 | 7.796 | 7 | 3 | 15 | 3 | 338 | 38.2 | 8.78 | 30.16 | 3 | 172.4 | 183.5 | 89.8 | 132.8 | 131.1 | 63.8 | 144.9 | 162.8 | 96.6 | 0.741 | 0.354555079 | 0.907112407 | 0.720440005 | 1.233750381 | 0.44517966 |
| A0A1U7QL04 | S-adenosylmethionine synthase OS=Mesocricetus auratus OX=10036 GN=Mat2a PE=3 SV=1 | 9.002 | 10 | 3 | 10 | 3 | 395 | 43.7 | 6.48 | 18.08 | 3 | 181.3 | 214.7 | 160.1 | 175.4 | 206.3 | 200.6 | 160 | 196.7 | 135.9 | 1.071 | 0.666839182 | 0.885811904 | 0.424062311 | 0.845955693 | 0.231083378 |
| A0A1U8CIJ2 | Myelin basic protein OS=Mesocricetus auratus OX=10036 GN=Mbp PE=3 SV=1 | 10.981 | 10 | 3 | 8 | 3 | 328 | 35.6 | 9.77 | 17.23 | 3 | 0 | 0 | 106.2 | 0 | 0 | 109.1 | 0 | 0 | 100.9 | 1.028 | 0.985716361 | 0.950094162 | 0.972876709 | 0.924839597 | 0.958656816 |
| A0A1U8BXF0 | cell surface glycoprotein MUC18 isoform X2 OS=Mesocricetus auratus OX=10036 GN=Mcam PE=4 SV=1 | 6.011 | 3 | 2 | 12 | 2 | 607 | 66.9 | 5.81 | 38.42 | 2 | 129.2 | 76.1 | 105.5 | 96.8 | 70.9 | 81.4 | 120.1 | 72 | 90.4 | 0.885 | 0.317839074 | 0.908944659 | 0.673725326 | 1.134082698 | 0.533323517 |
| A0A1U7Q626 | Malate dehydrogenase OS=Mesocricetus auratus OX=10036 GN=Mdh1 PE=3 SV=1 | 34.183 | 32 | 9 | 42 | 9 | 334 | 36.5 | 6.93 | 106.69 | 9 | 342.4 | 215.4 | 214.3 | 353.3 | 250.7 | 202.5 | 277.1 | 195.2 | 216.3 | 0.971 | 0.861233685 | 0.891853387 | 0.608121504 | 0.853812771 | 0.49355419 |
| A0A1U7QJU2 | Malate dehydrogenase OS=Mesocricetus auratus OX=10036 GN=Mdh2 PE=3 SV=1 | 71.077 | 48 | 15 | 73 | 15 | 338 | 35.6 | 8.68 | 249 | 15 | 792.4 | 666.4 | 812.6 | 653.4 | 524.9 | 674.4 | 709.1 | 699.1 | 723 | 0.788 | 0.099742578 | 0.938275953 | 0.414733662 | 1.150321153 | 0.182935228 |
| A0A1U8CP78 | Methyl-CpG-binding protein 2 OS=Mesocricetus auratus OX=10036 GN=Mecp2 PE=4 SV=1 | 5.739 | 3 | 1 | 6 | 1 | 487 | 52.5 | 9.95 | 25.32 | 1 | 68.8 | 74.4 | 33.3 | 70.4 | 59.6 | 30.4 | 47.5 | 66.2 | 37.3 | 0.898 | 0.77521391 | 0.855524079 | 0.614724531 | 0.941396509 | 0.842089527 |
| A0A1U7QJC4 | methyltransferase-like protein 7A isoform X1 OS=Mesocricetus auratus OX=10036 GN=LOC101825898 PE=4 SV=1 | 30.721 | 20 | 5 | 18 | 5 | 244 | 28 | 8.51 | 71.31 | 5 | 88.5 | 102.9 | 102.2 | 87.4 | 94.6 | 100.7 | 119.5 | 113.7 | 118.4 | 0.92 | 0.582399507 | 1.197547684 | 0.04053746 | 1.243721259 | 0.014429827 |
| A0A1U7QKY6 | microfibrillar-associated protein 2 isoform X2 OS=Mesocricetus auratus OX=10036 GN=Mfap2 PE=3 SV=1 | 4.971 | 5 | 1 | 5 | 1 | 186 | 20.9 | 4.89 | 17.63 | 1 | 129.5 | 182.5 | 59.5 | 119.1 | 171.7 | 71.5 | 106.6 | 120.9 | 50.9 | 0.936 | 0.950057391 | 0.749394347 | 0.504973036 | 0.768423958 | 0.483720359 |
| A0A1U7QA78 | microfibril-associated glycoprotein 4 isoform X2 OS=Mesocricetus auratus OX=10036 GN=Mfap4 PE=4 SV=1 | 12.157 | 12 | 3 | 11 | 3 | 256 | 28.7 | 5.63 | 33.81 | 3 | 83.2 | 0 | 0 | 82.8 | 0 | 0 | 76.2 | 0 | 0 | 0.995 | 0.99744421 | 0.915865385 | 0.95352533 | 0.920289855 | 0.956060353 |
| A0A1U8CFQ1 | Lactadherin OS=Mesocricetus auratus OX=10036 GN=Mfge8 PE=4 SV=1 | 4.649 | 4 | 2 | 11 | 2 | 484 | 53.2 | 7.24 | 33.16 | 2 | 49.6 | 67.8 | 0 | 66 | 78.6 | 0 | 31.2 | 56.1 | 0 | 1.242 | 0.789478193 | 0.743611584 | 0.719680779 | 0.60373444 | 0.554740654 |
| A0A1U8BVK7 | F-actin monooxygenase OS=Mesocricetus auratus OX=10036 GN=Mical1 PE=3 SV=1 | 8.518 | 4 | 2 | 5 | 2 | 1055 | 116.8 | 6.11 | 18.25 | 2 | 81.4 | 48.1 | 45.8 | 83.7 | 267.3 | 52.3 | 71.4 | 30.7 | 37.9 | 1.538 | 0.374577464 | 0.798630918 | 0.527565655 | 0.347136127 | 0.319808366 |
| A0A1U8CBY9 | myeloperoxidase OS=Mesocricetus auratus OX=10036 GN=Mpo PE=4 SV=1 | 7.19 | 4 | 3 | 11 | 1 | 718 | 81.1 | 9.61 | 26.43 | 3 | 0 | 50.4 | 74.7 | 0 | 48.3 | 77 | 0 | 41.7 | 42.5 | 0.994 | 0.998409883 | 0.673061551 | 0.633536628 | 0.671987231 | 0.637228898 |
| A0A1U7QWV2 | 55 kDa erythrocyte membrane protein OS=Mesocricetus auratus OX=10036 GN=Mpp1 PE=4 SV=1 | 1.642 | 4 | 1 | 2 | 1 | 466 | 52.5 | 7.34 | 3.44 | 1 | 43.1 | 0 | 0 | 49.2 | 0 | 0 | 48.5 | 0 | 0 | 1.142 | 0.93025505 | 1.125290023 | 0.937722755 | 0.985772358 | 0.992401045 |
| A0A1U7QDM7 | MAGUK p55 subfamily member 2 isoform X1 OS=Mesocricetus auratus OX=10036 GN=Mpp2 PE=3 SV=1 | 4.898 | 4 | 2 | 2 | 2 | 569 | 63.2 | 6.21 | 5.84 | 2 | 0 | 0 | 28.7 | 0 | 0 | 22.3 | 0 | 0 | 37.2 | 0.774 | 0.869274686 | 1.296167247 | 0.86577054 | 1.668161435 | 0.752091757 |
| A0A1U7QFG9 | MAGUK p55 subfamily member 5 OS=Mesocricetus auratus OX=10036 GN=Mpp5 PE=3 SV=1 | 12.872 | 4 | 2 | 12 | 2 | 675 | 77.2 | 6.21 | 33.4 | 2 | 32.5 | 0 | 0 | 39.3 | 0 | 0 | 31.7 | 0 | 0 | 1.207 | 0.900579152 | 0.975384615 | 0.98678543 | 0.806615776 | 0.887948828 |
| A0A1U7QUH9 | moesin OS=Mesocricetus auratus OX=10036 GN=Msn PE=4 SV=1 | 277.427 | 62 | 55 | 495 | 44 | 577 | 67.7 | 6.54 | 1205.96 | 55 | 13591.6 | 12135.5 | 15139 | 11892.7 | 10481.9 | 12565.8 | 14599.9 | 12469.7 | 14803.8 | 0.864 | 0.144407859 | 1.024648792 | 0.784069213 | 1.198423601 | 0.077467703 |
| A0A1U7QCW2 | mitochondrial carrier homolog 2 isoform X1 OS=Mesocricetus auratus OX=10036 GN=Mtch2 PE=3 SV=1 | 10.096 | 10 | 2 | 14 | 2 | 303 | 33.3 | 8.21 | 52.59 | 2 | 125.7 | 137.5 | 124 | 106.6 | 113.6 | 108 | 119 | 106.3 | 132.7 | 0.826 | 0.026411467 | 0.924586777 | 0.342832187 | 1.090798294 | 0.321465638 |
| A0A1U7Q800 | Methylmalonyl-CoA isomerase OS=Mesocricetus auratus OX=10036 GN=Mut PE=3 SV=1 | 4.829 | 3 | 2 | 3 | 2 | 748 | 83 | 7.15 | 2.68 | 2 | 74.9 | 74.2 | 0 | 61.4 | 54.7 | 0 | 75.4 | 83.7 | 0 | 0.778 | 0.745928899 | 1.067069081 | 0.931496312 | 1.37037037 | 0.688137727 |
| A0A1U7Q577 | Major vault protein OS=Mesocricetus auratus OX=10036 GN=Mvp PE=4 SV=1 | 81.1 | 24 | 19 | 75 | 19 | 870 | 96.9 | 5.69 | 274.9 | 19 | 806.6 | 937.8 | 1496.3 | 670 | 820.5 | 1247.5 | 726.1 | 839.8 | 1376.9 | 0.831 | 0.574009934 | 0.908075416 | 0.750570364 | 1.074799123 | 0.809635381 |
| B2Z890 | Myxovirus resistance protein 2 OS=Mesocricetus auratus OX=10036 GN=Mx2 PE=2 SV=1 | 75.244 | 41 | 20 | 63 | 20 | 635 | 72.8 | 5.53 | 173.42 | 20 | 351.3 | 505 | 319.7 | 1724.2 | 1851.5 | 1315.6 | 561 | 549.8 | 396.5 | 4.218 | 0.009883163 | 1.281717687 | 0.230383978 | 0.308159385 | 0.013113163 |
| A0A1U7QZ98 | myeloid-associated differentiation marker OS=Mesocricetus auratus OX=10036 GN=Myadm PE=4 SV=1 | 1.572 | 6 | 1 | 1 | 1 | 317 | 35 | 8.31 | 2.52 | 1 | 0 | 56.6 | 0 | 0 | 42.5 | 0 | 0 | 36.2 | 0 | 0.75 | 0.852531563 | 0.639575972 | 0.779049984 | 0.851764706 | 0.915718521 |
| A0A1U7QA56 | myosin-10 OS=Mesocricetus auratus OX=10036 GN=Myh10 PE=3 SV=1 | 363.332 | 38 | 79 | 429 | 58 | 1997 | 231.2 | 5.53 | 1192.37 | 79 | 7751.5 | 8825.2 | 7824.4 | 5432.5 | 6281.5 | 5482.2 | 6613.9 | 7861.3 | 7242.4 | 0.712 | 0.006437437 | 0.89002545 | 0.148018304 | 1.262930182 | 0.032305032 |
| P13540 | Myosin-7 OS=Mesocricetus auratus OX=10036 GN=MYH7 PE=2 SV=2 | 64.004 | 10 | 16 | 49 | 8 | 1934 | 222.8 | 5.73 | 125.12 | 16 | 275.1 | 283.8 | 392.4 | 264.3 | 335.3 | 408.4 | 199.9 | 177.6 | 263.2 | 1.181 | 0.753546893 | 0.673499422 | 0.094821835 | 0.635615079 | 0.079001675 |
| A0A1U7Q3X8 | myosin-9 OS=Mesocricetus auratus OX=10036 GN=Myh9 PE=3 SV=1 | 529.861 | 43 | 97 | 612 | 81 | 1960 | 226.4 | 5.66 | 1915.26 | 97 | 10609.2 | 12808.6 | 12241.9 | 10886.5 | 11747.9 | 11315.2 | 13147.5 | 14636.6 | 13151.5 | 0.917 | 0.486922486 | 1.147951329 | 0.10528507 | 1.20577562 | 0.025532482 |
| A0A1U7QZN8 | myosin regulatory light chain 12B OS=Mesocricetus auratus OX=10036 GN=LOC101844224 PE=4 SV=1 | 19.716 | 22 | 5 | 28 | 3 | 172 | 19.8 | 4.84 | 71.69 | 5 | 1064.2 | 1086.1 | 1037.2 | 1014.2 | 1033.1 | 1223.9 | 1061 | 1117.6 | 1451.2 | 0.951 | 0.720175897 | 1.138760784 | 0.349302155 | 1.10962338 | 0.450851313 |
| A0A1U7R2Y7 | myosin regulatory light polypeptide 9 OS=Mesocricetus auratus OX=10036 GN=Myl9 PE=4 SV=1 | 15.038 | 22 | 4 | 16 | 2 | 172 | 19.8 | 4.92 | 29.42 | 4 | 137.6 | 150 | 262 | 120 | 103.3 | 182.3 | 78 | 138.8 | 219.9 | 0.696 | 0.369718124 | 0.794577875 | 0.545563937 | 1.076676529 | 0.840708003 |
| A0A1U7QRM9 | Myosin light chain kinase, smooth muscle OS=Mesocricetus auratus OX=10036 GN=Mylk PE=3 SV=1 | 38.526 | 7 | 10 | 39 | 9 | 1943 | 213.1 | 6.14 | 118.85 | 10 | 1196.5 | 739.6 | 883.1 | 940.2 | 597.1 | 668.7 | 1023.7 | 622.7 | 762 | 0.757 | 0.30091099 | 0.854284904 | 0.487498294 | 1.091749773 | 0.690327525 |
| A0A1U7R0K2 | unconventional myosin-Id isoform X1 OS=Mesocricetus auratus OX=10036 GN=Myo1d PE=3 SV=1 | 10.423 | 3 | 3 | 7 | 3 | 1006 | 116 | 9.38 | 20.14 | 3 | 38.1 | 0 | 91 | 44.3 | 0 | 71.3 | 25.7 | 0 | 76.8 | 0.955 | 0.900226865 | 0.793958172 | 0.811323029 | 0.886678201 | 0.893752842 |
| A0A1U7Q6T1 | GlcNAc kinase OS=Mesocricetus auratus OX=10036 GN=Nagk PE=3 SV=1 | 1.943 | 2 | 1 | 2 | 1 | 343 | 36.9 | 6.18 | 4.01 | 1 | 0 | 0 | 40.3 | 0 | 0 | 41.3 | 0 | 0 | 25.3 | 1.024 | 0.987003977 | 0.627791563 | 0.77114064 | 0.612590799 | 0.76092338 |
| A0A1U8CBW0 | nicotinamide phosphoribosyltransferase isoform X1 OS=Mesocricetus auratus OX=10036 GN=Nampt PE=3 SV=1 | 9.474 | 6 | 4 | 16 | 4 | 595 | 67 | 8.31 | 30.14 | 4 | 400.1 | 290.4 | 333.2 | 359.9 | 268.5 | 358.8 | 366.8 | 264.8 | 307.3 | 1.01 | 0.795896113 | 0.917163231 | 0.551614984 | 0.951073744 | 0.723061973 |
| A0A1U7Q4B3 | nucleosome assembly protein 1-like 1 isoform X2 OS=Mesocricetus auratus OX=10036 GN=Nap1l1 PE=3 SV=1 | 19.411 | 12 | 4 | 24 | 4 | 383 | 44.6 | 4.56 | 85.27 | 4 | 309.4 | 401.5 | 388.5 | 236.9 | 327.2 | 288.8 | 271.6 | 411.3 | 345.8 | 0.766 | 0.102821321 | 0.935692196 | 0.661729484 | 1.206120295 | 0.300191032 |
| A0A1U8CYN1 | alpha-soluble NSF attachment protein OS=Mesocricetus auratus OX=10036 GN=Napa PE=3 SV=1 | 16.017 | 17 | 3 | 11 | 3 | 295 | 33.2 | 5.45 | 52.36 | 3 | 66.8 | 37.3 | 0 | 67.7 | 49.6 | 0 | 56.7 | 38.2 | 0 | 1.162 | 0.882689929 | 0.911623439 | 0.910326871 | 0.809036658 | 0.790543583 |
| A0A1U7QP21 | Nuclear cap-binding protein subunit 2 OS=Mesocricetus auratus OX=10036 GN=Ncbp2 PE=3 SV=1 | 3.851 | 7 | 1 | 2 | 1 | 156 | 18 | 7.81 | 6.28 | 1 | 50.3 | 0 | 0 | 39.3 | 0 | 0 | 45 | 0 | 0 | 0.78 | 0.872012906 | 0.894632207 | 0.941223244 | 1.145038168 | 0.928660531 |

**Supplementary Table 1: List of differentially expressed proteins identified across the 4dpi infected and 14dpi infected**

| Accession | Description | Sum PEP Score | Coverage [%] | # Peptides | # PSMs | # Unique Peptides | # AAs | MW [kDa] | calc. pI | Score Sequest HT: Sequest HT | # Peptides (by Search Engine): Sequest HT | Mock-infected (4dpi) Replicate_R1 | Mock-infected (4dpi) Replicate_R2 | Mock-infected (4dpi) Replicate_R3 | SARS-CoV-2-infected (4dpi) Replicate_R1 | SARS-CoV-2-infected (4dpi) Replicate_R2 | SARS-CoV-2-infected (4dpi) Replicate_R3 | SARS-CoV-2-(14dpi) Replicate_R1 | SARS-CoV-2-infected (14dpi) Replicate_R2 | SARS-CoV-2-infected (14dpi) Replicate_R3 | Abundance Ratio: SARS-CoV-2-infected (4dpi) / Mock-infected (4dpi) | P-Value: SARS-CoV-2-infected (4dpi) / Mock-infected (4dpi) | Abundance Ratio: SARS-CoV-2-infected (14dpi) / Mock-infected (4dpi) | P-Value: SARS-CoV-2-infected (14dpi) / Mock-infected (4dpi) | Abundance Ratio: SARS-CoV-2-infected (14dpi) / SARS-CoV-2-infected (4dpi) | P-Value: SARS-CoV-2-infected (14dpi) / SARS-CoV-2-infected (4dpi) |
|---|---|---|---|---|---|---|---|---|---|---|---|---|---|---|---|---|---|---|---|---|---|---|---|---|---|
| A0A1U7Q3A1 | nck-associated protein 1 isoform X2 OS=Mesocricetus auratus OX=10036 GN=Nckap1 PE=4 SV=1 | 4.952 | 2 | 2 | 2 | 2 | 1128 | 128.7 | 6.62 | 6.84 | 2 | 0 | 63.9 | 0 | 0 | 60 | 0 | 0 | 49.3 | 0 | 0.939 | 0.966651909 | 0.771517997 | 0.865778597 | 0.821666667 | 0.897304321 |
| A0A1U7QMP9 | NADH dehydrogenase [ubiquinone] 1 alpha subcomplex subunit 10, mitochondrial OS=Mesocricetus auratus OX=10036 GN=Ndufa10 PE=3 SV=1 | 3.309 | 3 | 1 | 4 | 1 | 355 | 40.5 | 7.01 | 4.07 | 1 | 36.8 | 200.5 | 0 | 23.3 | 118.3 | 0 | 31.7 | 195.7 | 0 | 0.611 | 0.683529138 | 0.958280657 | 0.971382707 | 1.605932203 | 0.710578583 |
| A0A1U7QKJ7 | NADH dehydrogenase [ubiquinone] 1 alpha subcomplex subunit 12 OS=Mesocricetus auratus OX=10036 GN=Ndufa12 PE=3 SV=1 | 2.265 | 8 | 1 | 2 | 1 | 149 | 17.6 | 9.57 | 0 | 1 | 147.3 | 0 | 117.5 | 88.6 | 0 | 95.7 | 74.8 | 0 | 68.9 | 0.7 | 0.65139819 | 0.542673716 | 0.485297626 | 0.779706999 | 0.747308384 |
| A0A1U7QUD5 | Complex I-B16.6 OS=Mesocricetus auratus OX=10036 GN=Ndufa13 PE=3 SV=1 | 11.297 | 24 | 3 | 14 | 3 | 144 | 16.9 | 8.97 | 21.46 | 3 | 137.4 | 0 | 156 | 137.1 | 0 | 125.9 | 114.3 | 0 | 123.2 | 0.897 | 0.885445475 | 0.809475119 | 0.783396749 | 0.903041825 | 0.892852522 |
| A0A1U7QM94 | NADH dehydrogenase [ubiquinone] 1 alpha subcomplex subunit 2 OS=Mesocricetus auratus OX=10036 GN=Ndufa2 PE=3 SV=1 | 3.354 | 10 | 1 | 6 | 1 | 99 | 11 | 9.99 | 15.9 | 1 | 157.3 | 67.5 | 252.6 | 150.8 | 68.8 | 266.7 | 119.5 | 41.6 | 186.8 | 1.056 | 0.971648854 | 0.728739003 | 0.561557516 | 0.715402015 | 0.554879247 |
| A0A1U7QPC9 | Complex I subunit B13 OS=Mesocricetus auratus OX=10036 GN=Ndufa5 PE=3 SV=1 | 2.593 | 13 | 1 | 2 | 1 | 116 | 13.4 | 6.32 | 5.72 | 1 | 44.1 | 44.8 | 0 | 40.2 | 47 | 0 | 42.3 | 48 | 0 | 0.977 | 0.979617773 | 1.015748031 | 0.983480645 | 1.035550459 | 0.963252428 |
| A0A1U7RC90 | NADH dehydrogenase [ubiquinone] 1 alpha subcomplex subunit 9, mitochondrial OS=Mesocricetus auratus OX=10036 GN=Ndufa9 PE=4 SV=1 | 12.349 | 15 | 4 | 15 | 4 | 377 | 42.6 | 9.83 | 31.89 | 4 | 76 | 180.2 | 61.8 | 75.7 | 128.1 | 60.3 | 65.3 | 177.8 | 79.7 | 0.711 | 0.700653206 | 1.01509434 | 0.976665015 | 1.222264294 | 0.662845684 |
| A0A1U7QLT7 | Complex I-PDSW OS=Mesocricetus auratus OX=10036 GN=Ndufb10 PE=3 SV=1 | 10.534 | 28 | 3 | 18 | 3 | 176 | 20.9 | 8.7 | 48.5 | 3 | 98.8 | 155.7 | 128.2 | 76 | 129 | 103.6 | 101.3 | 136.4 | 124.8 | 0.808 | 0.333333433 | 0.947217141 | 0.749159304 | 1.174659754 | 0.392673502 |
| A0A1U7QBY9 | Complex I-B12 OS=Mesocricetus auratus OX=10036 GN=Ndufb3 PE=3 SV=1 | 1.909 | 8 | 1 | 5 | 1 | 106 | 11.9 | 9.01 | 12.62 | 1 | 318 | 526.8 | 214.1 | 233.4 | 426.2 | 154.2 | 269.9 | 432.2 | 184.6 | 0.727 | 0.541520583 | 0.837378412 | 0.651153937 | 1.089579749 | 0.834050185 |
| A0A1U7QTN8 | Complex I-B17 OS=Mesocricetus auratus OX=10036 GN=Ndufb6 PE=3 SV=1 | 10.629 | 27 | 2 | 8 | 2 | 128 | 15.4 | 9.66 | 14.95 | 2 | 110.2 | 210.8 | 147.9 | 109.2 | 191.3 | 126.8 | 103.8 | 219 | 153.8 | 0.874 | 0.737505845 | 1.016421412 | 0.956738755 | 1.115375614 | 0.714848323 |
| A0A1U7QNI9 | Complex I-B22 OS=Mesocricetus auratus OX=10036 GN=Ndufb9 PE=3 SV=1 | 6.376 | 13 | 2 | 5 | 2 | 179 | 21.9 | 7.49 | 2.25 | 2 | 0 | 186.5 | 334 | 0 | 101 | 190 | 0 | 190.1 | 362.4 | 0.555 | 0.538254046 | 1.061479347 | 0.94392691 | 1.89862543 | 0.513844679 |
| A0A1U7Q9N9 | NADH dehydrogenase [ubiquinone] 1 subunit C2 OS=Mesocricetus auratus OX=10036 GN=LOC101844122 PE=3 SV=1 | 5.492 | 10 | 2 | 3 | 2 | 120 | 14.1 | 8.16 | 7.49 | 2 | 161.4 | 164.3 | 0 | 110.5 | 132.9 | 0 | 138.5 | 202 | 0 | 0.744 | 0.709025449 | 1.045440589 | 0.95418135 | 1.3989318 | 0.680805004 |
| A0A1U7QPW6 | NADH-ubiquinone oxidoreductase 75 kDa subunit, mitochondrial OS=Mesocricetus auratus OX=10036 GN=Ndufs1 PE=3 SV=1 | 28.935 | 14 | 7 | 17 | 7 | 727 | 79.6 | 6 | 49.66 | 7 | 92 | 108.8 | 147 | 112.8 | 159.4 | 238.6 | 59.1 | 92.8 | 58.7 | 1.403 | 0.276380666 | 0.605520414 | 0.090234123 | 0.41229444 | 0.101731823 |
| A0A1U7Q371 | Complex I-30kD OS=Mesocricetus auratus OX=10036 GN=Ndufs3 PE=3 SV=1 | 21.053 | 25 | 6 | 13 | 6 | 263 | 30.1 | 7.53 | 29.04 | 6 | 135.4 | 375.4 | 150 | 110 | 296.4 | 118.3 | 94.7 | 260.6 | 110 | 0.79 | 0.670776052 | 0.704146489 | 0.53106478 | 0.886792453 | 0.818281977 |
| A0A1U7QDT8 | NADH dehydrogenase [ubiquinone] iron-sulfur protein 6, mitochondrial OS=Mesocricetus auratus OX=10036 GN=Ndufs6 PE=3 SV=1 | 11.173 | 33 | 3 | 9 | 3 | 116 | 13 | 8.92 | 23.11 | 3 | 39.4 | 60 | 112.2 | 40.3 | 62.4 | 97 | 48.3 | 62.5 | 108.1 | 0.865 | 0.891709538 | 1.034499055 | 0.93549463 | 1.096144216 | 0.806505223 |
| A0A1U8CRN3 | Complex I-20kD OS=Mesocricetus auratus OX=10036 GN=Ndufs7 PE=3 SV=1 | 5.32 | 9 | 2 | 6 | 2 | 219 | 24 | 9.76 | 6.98 | 2 | 144 | 42.3 | 69.8 | 145.3 | 37 | 59.6 | 138.4 | 44.4 | 76.3 | 1.009 | 0.921043085 | 1.011714174 | 0.981736843 | 1.071103762 | 0.900577 |
| A0A1U8CKE4 | nestin OS=Mesocricetus auratus OX=10036 GN=Nes PE=3 SV=1 | 32.31 | 10 | 9 | 26 | 9 | 2003 | 220.9 | 4.35 | 79.07 | 9 | 265.4 | 330.4 | 207.4 | 206.5 | 253.8 | 144.7 | 226.3 | 289.9 | 187.5 | 0.698 | 0.237868063 | 0.876120518 | 0.515340068 | 1.163140496 | 0.491294601 |
| A0A1U7QPE1 | nidogen-1 OS=Mesocricetus auratus OX=10036 GN=Nid1 PE=4 SV=1 | 85.158 | 16 | 15 | 91 | 15 | 1246 | 136.5 | 5.38 | 325.93 | 15 | 1263.4 | 1527.8 | 1925.2 | 1051.2 | 1242.7 | 1628.6 | 1082.9 | 1419.9 | 1701.1 | 0.846 | 0.361429829 | 0.891336613 | 0.55090004 | 1.071739962 | 0.722955717 |
| A0A1U7QC34 | Nucleoside diphosphate kinase OS=Mesocricetus auratus OX=10036 GN=LOC101838863 PE=3 SV=1 | 22.386 | 36 | 6 | 37 | 3 | 152 | 17.3 | 7.99 | 85.62 | 6 | 821.2 | 241.8 | 458.4 | 905.5 | 305.5 | 568.8 | 778.4 | 201.1 | 413.9 | 1.244 | 0.740214354 | 0.915866965 | 0.866829637 | 0.782896955 | 0.622772685 |
| A0A1U8CC67 | Nucleoside diphosphate kinase OS=Mesocricetus auratus OX=10036 GN=Nme4 PE=3 SV=1 | 1.851 | 7 | 1 | 1 | 1 | 123 | 13.5 | 6.3 | 2.76 | 1 | 0 | 52.7 | 0 | 0 | 52.4 | 0 | 0 | 44.5 | 0 | 0.994 | 0.996972454 | 0.844402277 | 0.911251508 | 0.849236641 | 0.914187449 |
| A0A1U7Q4X4 | N-myc-interactor OS=Mesocricetus auratus OX=10036 GN=Nmi PE=3 SV=1 | 1.866 | 5 | 1 | 1 | 1 | 302 | 34.2 | 5.33 | 3.01 | 1 | 0 | 0 | 117.3 | 0 | 0 | 86.3 | 0 | 0 | 114.8 | 0.736 | 0.84270648 | 0.978687127 | 0.988576934 | 1.330243337 | 0.853085788 |
| A0A1U7R7S2 | non-POU domain-containing octamer-binding protein OS=Mesocricetus auratus OX=10036 GN=Nono PE=4 SV=1 | 46.294 | 21 | 9 | 45 | 8 | 474 | 54.7 | 8.95 | 99.34 | 9 | 1141.4 | 1167.8 | 1332.8 | 1006.4 | 1062.5 | 1060.4 | 1155.7 | 1190.7 | 1296.9 | 0.796 | 0.093234249 | 1.000356947 | 0.99560201 | 1.164253987 | 0.040202935 |
| A0A1U8BX22 | nucleolar protein 56 OS=Mesocricetus auratus OX=10036 GN=Nop56 PE=3 SV=1 | 7.729 | 7 | 3 | 5 | 3 | 591 | 65.7 | 8.9 | 9.41 | 3 | 34 | 0 | 31.5 | 30.3 | 0 | 26.7 | 37.4 | 0 | 36.3 | 0.87 | 0.855028501 | 1.12519084 | 0.876210459 | 1.292982456 | 0.739743226 |
| A0A1U7RFY8 | Epididymal secretory protein E1 OS=Mesocricetus auratus OX=10036 GN=Npc2 PE=3 SV=1 | 14 | 34 | 4 | 13 | 4 | 149 | 16.3 | 7.97 | 28.82 | 4 | 332.6 | 249.5 | 140.3 | 447.1 | 298.4 | 222.2 | 393.5 | 280.9 | 198.9 | 1.41 | 0.398869304 | 1.208887043 | 0.560169801 | 0.902449106 | 0.735873497 |
| A0A1U8BZW3 | nuclear protein localization protein 4 homolog isoform X2 OS=Mesocricetus auratus OX=10036 GN=Nploc4 PE=3 SV=1 | 7.372 | 7 | 2 | 6 | 2 | 563 | 63.3 | 6.32 | 27.08 | 2 | 0 | 0 | 86.9 | 0 | 0 | 106.8 | 0 | 0 | 82.3 | 1.229 | 0.892345754 | 0.947065593 | 0.971188775 | 0.770599251 | 0.865191972 |
| A0A1U7QPR4 | Nucleophosmin OS=Mesocricetus auratus OX=10036 GN=Npm1 PE=3 SV=1 | 43.263 | 40 | 9 | 41 | 9 | 292 | 32.5 | 4.77 | 135.93 | 9 | 432.9 | 436.8 | 397 | 678.5 | 547.9 | 694.5 | 408.2 | 412.1 | 439.4 | 1.567 | 0.034870183 | 0.99447383 | 0.891686752 | 0.65578635 | 0.036747623 |
| A0A1U7QYV1 | nephronectin isoform X1 OS=Mesocricetus auratus OX=10036 GN=Npnt PE=3 SV=1 | 19.255 | 11 | 6 | 22 | 6 | 604 | 66.2 | 8.21 | 55 | 6 | 414.8 | 830.9 | 977.9 | 279.4 | 544.8 | 592.3 | 901.5 | 687.9 | 834.4 | 0.606 | 0.255648473 | 1.090034179 | 0.739495375 | 1.711118955 | 0.053448585 |
| A0A1U8BQN9 | neuroplastin isoform X1 OS=Mesocricetus auratus OX=10036 GN=Nptn PE=4 SV=1 | 11.335 | 10 | 4 | 4 | 4 | 397 | 44.5 | 8 | 10.36 | 4 | 151.6 | 202 | 0 | 112.7 | 141.6 | 0 | 137.5 | 164.8 | 0 | 0.722 | 0.682145576 | 0.854920814 | 0.840079732 | 1.188753441 | 0.8228601 |
| A0A1U7Q5Q7 | Vesicle-fusing ATPase OS=Mesocricetus auratus OX=10036 GN=Nsf PE=3 SV=1 | 15.277 | 6 | 5 | 15 | 5 | 744 | 82.5 | 6.95 | 31.05 | 5 | 240.4 | 243.5 | 84.8 | 207.2 | 190.5 | 66.5 | 226.6 | 230.3 | 95.7 | 0.784 | 0.639302381 | 0.971689819 | 0.941488717 | 1.190435157 | 0.66275819 |
| A0A1U7Q888 | NSFL1 cofactor p47 OS=Mesocricetus auratus OX=10036 GN=Nsfl1c PE=4 SV=1 | 18.648 | 19 | 5 | 9 | 5 | 370 | 40.7 | 5.15 | 37.78 | 5 | 65.1 | 0 | 230.2 | 72.3 | 0 | 165 | 84.8 | 0 | 232.4 | 0.892 | 0.829566794 | 1.074161869 | 0.943307449 | 1.336704593 | 0.766124947 |
| A0A1U7QVP5 | nucleobindin-2 OS=Mesocricetus auratus OX=10036 GN=Nucb2 PE=3 SV=1 | 25.966 | 24 | 7 | 26 | 7 | 420 | 50.2 | 5.21 | 95.83 | 7 | 409 | 244.2 | 171.5 | 374.8 | 244.9 | 171.7 | 364.8 | 208.3 | 166.9 | 1.003 | 0.909931752 | 0.897295986 | 0.775899023 | 0.935051807 | 0.849408385 |
| A0A1U7R2C6 | Nuclear distribution protein C homolog OS=Mesocricetus auratus OX=10036 GN=Nudc PE=3 SV=1 | 16.479 | 18 | 4 | 19 | 4 | 332 | 38.5 | 5.44 | 60.64 | 4 | 128.5 | 207.8 | 206.4 | 113.2 | 182 | 208.6 | 111 | 177.1 | 182.5 | 0.881 | 0.754272483 | 0.867145753 | 0.52903072 | 0.934100834 | 0.777804201 |
| A0A1U7R4E0 | Cleavage and polyadenylation specificity factor subunit 5 OS=Mesocricetus auratus OX=10036 GN=Nudt21 PE=3 SV=1 | 2.874 | 4 | 1 | 6 | 1 | 227 | 26.2 | 8.82 | 12.47 | 1 | 67.2 | 30.5 | 0 | 69.4 | 26 | 0 | 52.9 | 38 | 0 | 0.938 | 0.979510857 | 0.930399181 | 0.932315375 | 0.952830189 | 0.956328203 |
| A0A1U8C7K7 | nuclear mitotic apparatus protein 1 isoform X2 OS=Mesocricetus auratus OX=10036 GN=Numa1 PE=4 SV=1 | 15.558 | 3 | 5 | 18 | 5 | 2097 | 236.1 | 5.66 | 53.11 | 5 | 170.2 | 283 | 261.4 | 167.7 | 257.5 | 234.5 | 161.3 | 254.8 | 249.5 | 0.953 | 0.698880289 | 0.931430171 | 0.740656337 | 1.008943459 | 0.963678306 |
| A0A1U8BXK2 | protein numb homolog isoform X4 OS=Mesocricetus auratus OX=10036 GN=Numb PE=4 SV=1 | 5.69 | 5 | 2 | 4 | 2 | 596 | 64.7 | 8.47 | 14.96 | 2 | 106.8 | 32.1 | 0 | 91.8 | 33 | 0 | 117.3 | 27.9 | 0 | 0.941 | 0.915410021 | 1.045356371 | 0.966853762 | 1.163461538 | 0.886232501 |
| A0A1U7QEB0 | Ornithine--oxo-acid aminotransferase OS=Mesocricetus auratus OX=10036 GN=Oat PE=3 SV=1 | 1.976 | 3 | 1 | 1 | 1 | 439 | 48.3 | 7.03 | 2.5 | 1 | 0 | 0 | 74.7 | 0 | 0 | 59.6 | 0 | 0 | 65 | 0.798 | 0.882462066 | 0.870147256 | 0.926762524 | 1.090604027 | 0.954132239 |
| A0A1U8CLA2 | OCIA domain-containing protein 1 isoform X3 OS=Mesocricetus auratus OX=10036 GN=Ociad1 PE=4 SV=1 | 5.075 | 15 | 1 | 11 | 1 | 188 | 20.8 | 7.88 | 43.43 | 1 | 70.7 | 0 | 32.4 | 49.1 | 0 | 24.7 | 70.6 | 0 | 40.4 | 0.727 | 0.716856077 | 1.076624636 | 0.931801272 | 1.504065041 | 0.647432729 |
| A0A1U7RGH4 | 2-oxoglutarate dehydrogenase, mitochondrial OS=Mesocricetus auratus OX=10036 GN=Ogdh PE=3 SV=1 | 37.704 | 14 | 12 | 60 | 5 | 928 | 105.7 | 6.6 | 128.94 | 12 | 1194.5 | 1250.5 | 1490.9 | 962.1 | 985.4 | 1108.7 | 1113.5 | 1191.1 | 1351.3 | 0.805 | 0.064792155 | 0.928859981 | 0.464427549 | 1.196224069 | 0.085480574 |
| A0A1U7QF45 | mimecan OS=Mesocricetus auratus OX=10036 GN=Ogn PE=4 SV=1 | 19.062 | 24 | 5 | 25 | 5 | 212 | 24.1 | 5.5 | 58.39 | 5 | 58.4 | 160.3 | 147.6 | 55.3 | 158.3 | 135.4 | 50.3 | 98.4 | 107.5 | 0.988 | 0.903695406 | 0.699426699 | 0.38778399 | 0.734097421 | 0.449307157 |
| A0A1U7QUU5 | alpha-1-acid glycoprotein 1 OS=Mesocricetus auratus OX=10036 GN=LOC101825672 PE=3 SV=1 | 11.999 | 9 | 1 | 24 | 1 | 198 | 22.5 | 5.87 | 74.66 | 1 | 93.8 | 83.5 | 91.2 | 935.5 | 913 | 1020.2 | 0 | 0 | 0 | 10.957 | 0.001299416 | #DIV/0! | 0.001191753 | #DIV/0! | 0.001162822 |
| A0A1U7QAU1 | Protein disulfide-isomerase OS=Mesocricetus auratus OX=10036 GN=P4hb PE=3 SV=1 | 92.914 | 44 | 20 | 115 | 20 | 509 | 57 | 4.89 | 304.11 | 20 | 1809 | 1864.5 | 1679.1 | 1801.3 | 1898.7 | 1660.9 | 1747.1 | 1828.4 | 1663.3 | 1.018 | 0.976554566 | 0.978739304 | 0.630045588 | 0.977223974 | 0.655882989 |
| A0A1U7QMA4 | palladin isoform X2 OS=Mesocricetus auratus OX=10036 GN=Palld PE=4 SV=1 | 9.789 | 9 | 3 | 7 | 3 | 660 | 71.7 | 7.85 | 25.62 | 3 | 69.1 | 124.6 | 99.3 | 53.9 | 96.1 | 61.2 | 61.3 | 114.5 | 79.4 | 0.769 | 0.260165239 | 0.870989761 | 0.603643017 | 1.208333333 | 0.511835909 |
| A0A1U8C7F2 | Maillard deglycase OS=Mesocricetus auratus OX=10036 GN=Park7 PE=3 SV=1 | 32.937 | 39 | 7 | 36 | 7 | 189 | 19.9 | 6.79 | 100.78 | 7 | 727.1 | 827.1 | 927.6 | 485.7 | 557.2 | 662.8 | 660.1 | 740.6 | 870.1 | 0.715 | 0.029415353 | 0.914981062 | 0.45072633 | 1.331300932 | 0.079845067 |
| A0A1U7QK92 | alpha-parvin OS=Mesocricetus auratus OX=10036 GN=Parva PE=3 SV=1 | 38.039 | 31 | 10 | 29 | 10 | 372 | 42.2 | 5.95 | 61.35 | 10 | 620.2 | 415.6 | 589.3 | 366.2 | 233.7 | 318.9 | 647.6 | 438.4 | 665.1 | 0.59 | 0.044632872 | 1.07753369 | 0.686943458 | 1.905855464 | 0.042595337 |
| A0A1U7QFH5 | poly(RC)-binding protein 1 OS=Mesocricetus auratus OX=10036 GN=Pcbp1 PE=4 SV=1 | 20.607 | 18 | 5 | 22 | 3 | 356 | 37.5 | 7.09 | 69.1 | 5 | 111.6 | 135 | 97.4 | 89.9 | 114.9 | 90.8 | 100 | 82.2 | 62.6 | 0.805 | 0.308527008 | 0.711627907 | 0.098135642 | 0.828146143 | 0.284213773 |
| A0A1U8BWR9 | Proliferating cell nuclear antigen OS=Mesocricetus auratus OX=10036 GN=Pcna PE=3 SV=1 | 11.299 | 14 | 3 | 9 | 3 | 261 | 28.7 | 4.69 | 29.86 | 3 | 56.8 | 164.8 | 117.5 | 79.9 | 214.4 | 158.5 | 65 | 173.2 | 111.3 | 1.406 | 0.492437065 | 1.030669419 | 0.941337225 | 0.771863958 | 0.530869724 |
| A0A1U7Q674 | programmed cell death protein 6 isoform X2 OS=Mesocricetus auratus OX=10036 GN=Pdcd6 PE=4 SV=1 | 4.688 | 6 | 1 | 11 | 1 | 189 | 21.7 | 5.4 | 32.71 | 1 | 51.2 | 131.6 | 82.8 | 49.6 | 97.8 | 54 | 55.9 | 118.8 | 84.5 | 0.74 | 0.493391514 | 0.975903614 | 0.946244693 | 1.286991063 | 0.465105131 |
| A0A1U7QJ63 | programmed cell death 6-interacting protein isoform X2 OS=Mesocricetus auratus OX=10036 GN=Pdcd6ip PE=4 SV=1 | 23.665 | 10 | 9 | 33 | 9 | 868 | 95.9 | 6.39 | 53.82 | 9 | 883.1 | 585.3 | 919.3 | 808.3 | 499.4 | 820.7 | 734.1 | 547.8 | 822.8 | 0.858 | 0.593291989 | 0.881475897 | 0.520629882 | 0.988864875 | 0.955561147 |
| A0A1U8CUW9 | myomegalin isoform X9 OS=Mesocricetus auratus OX=10036 GN=LOC101838805 PE=4 SV=1 | 1.897 | 0 | 1 | 11 | 1 | 2488 | 279.2 | 5.71 | 29.54 | 1 | 252.4 | 560.7 | 153.6 | 1191.9 | 1893.2 | 1191.3 | 200.4 | 452.1 | 91.1 | 4.722 | 0.02461969 | 0.769214855 | 0.671652627 | 0.173884576 | 0.022659945 |
| A0A1U7QZ77 | Retinal rod rhodopsin-sensitive cGMP 3',5'-cyclic phosphodiesterase subunit delta OS=Mesocricetus auratus OX=10036 GN=Pde6d PE=3 SV=1 | 6.625 | 13 | 1 | 7 | 1 | 150 | 17.4 | 5.67 | 26.67 | 1 | 0 | 39.9 | 0 | 0 | 31.1 | 0 | 0 | 36.2 | 0 | 0.779 | 0.870829468 | 0.90726817 | 0.948571346 | 1.163987138 | 0.920153994 |
| A0A1U7QW04 | Pyruvate dehydrogenase E1 component subunit alpha OS=Mesocricetus auratus OX=10036 GN=Pdha1 PE=4 SV=1 | 11.244 | 12 | 3 | 8 | 3 | 390 | 43.3 | 8.19 | 36.98 | 3 | 59.9 | 184.6 | 186.8 | 62.1 | 143.2 | 142.1 | 37.6 | 151.2 | 181 | 0.776 | 0.609268729 | 0.857407837 | 0.75203014 | 1.064478987 | 0.892627482 |
| A0A1U7R1A6 | Pyruvate dehydrogenase E1 component subunit beta OS=Mesocricetus auratus OX=10036 GN=Pdhb PE=4 SV=1 | 13.025 | 11 | 3 | 16 | 3 | 360 | 39.2 | 6.65 | 50.12 | 3 | 32.9 | 38.8 | 34.6 | 30.6 | 34.9 | 30.4 | 33.2 | 25.3 | 26.1 | 0.877 | 0.206251919 | 0.795860771 | 0.085205019 | 0.882168926 | 0.280202659 |
| A0A1U7RH50 | Protein disulfide-isomerase A4 OS=Mesocricetus auratus OX=10036 GN=Pdia4 PE=3 SV=1 | 9.378 | 4 | 3 | 6 | 3 | 643 | 72.6 | 5.06 | 16.12 | 3 | 286.1 | 229.9 | 0 | 224.6 | 160.6 | 0 | 282.8 | 198.3 | 0 | 0.741 | 0.713647261 | 0.932364341 | 0.928143347 | 1.248961578 | 0.781059679 |
| A0A1U7RB20 | protein disulfide-isomerase A6 OS=Mesocricetus auratus OX=10036 GN=Pdia6 PE=3 SV=1 | 50.044 | 28 | 9 | 52 | 9 | 445 | 48.6 | 5.19 | 138.59 | 9 | 563.4 | 355.9 | 348.2 | 425 | 289.1 | 300.6 | 495.9 | 324.1 | 316.2 | 0.863 | 0.377108683 | 0.896410256 | 0.658785687 | 1.119739825 | 0.61097027 |
| A0A1U7Q5W8 | PDZ and LIM domain protein 1 OS=Mesocricetus auratus OX=10036 GN=Pdlim1 PE=4 SV=1 | 27.211 | 23 | 5 | 32 | 5 | 329 | 36 | 7.06 | 93.45 | 5 | 810.2 | 903.2 | 961.3 | 693.2 | 713.5 | 747 | 815.8 | 940.7 | 951.1 | 0.777 | 0.04588848 | 1.012300445 | 0.867893897 | 1.257185309 | 0.038783849 |
| A0A1U8BQ37 | PDZ and LIM domain protein 2 isoform X1 OS=Mesocricetus auratus OX=10036 GN=Pdlim2 PE=4 SV=1 | 26.665 | 14 | 4 | 22 | 4 | 346 | 37.2 | 8.73 | 82.31 | 4 | 202 | 281.1 | 199.1 | 185.7 | 274.3 | 157.8 | 187.8 | 229.7 | 220.8 | 0.937 | 0.654354263 | 0.93564937 | 0.657930184 | 1.03318226 | 0.868580957 |
| A0A1U7QFU5 | PDZ and LIM domain protein 3 isoform X2 OS=Mesocricetus auratus OX=10036 GN=Pdlim3 PE=4 SV=1 | 9.932 | 10 | 2 | 11 | 2 | 316 | 34.3 | 7.91 | 30.25 | 2 | 91.1 | 46.8 | 243.5 | 56.2 | 58 | 200.6 | 62.4 | 46.7 | 200.7 | 0.9 | 0.786460062 | 0.812270582 | 0.772877161 | 0.9841169 | 0.98173498 |
| A0A1U7R6H2 | PDZ and LIM domain protein 5 isoform X1 OS=Mesocricetus auratus OX=10036 GN=Pdlim5 PE=4 SV=1 | 7.338 | 4 | 2 | 10 | 2 | 593 | 63.5 | 8.44 | 26.71 | 2 | 182.1 | 316.4 | 178.4 | 169.6 | 312.6 | 167.6 | 168.3 | 296.3 | 172.9 | 0.94 | 0.89787276 | 0.94179347 | 0.842137424 | 0.981071099 | 0.951845474 |
| A0A1U8C5R2 | PDZ and LIM domain protein 7 isoform X3 OS=Mesocricetus auratus OX=10036 GN=Pdlim7 PE=4 SV=1 | 6.31 | 7 | 2 | 10 | 2 | 425 | 46.6 | 8.31 | 28.91 | 2 | 159.8 | 192.2 | 191.6 | 104.5 | 141.8 | 156.3 | 126.1 | 152.8 | 168.4 | 0.737 | 0.074203709 | 0.822847682 | 0.12239021 | 1.111028316 | 0.494719171 |
| A0A1U7QWC4 | Prefoldin subunit 4 OS=Mesocricetus auratus OX=10036 GN=Pfdn4 PE=3 SV=1 | 3.215 | 8 | 1 | 6 | 1 | 134 | 15.2 | 4.67 | 2.4 | 1 | 123.9 | 124 | 130.7 | 86.6 | 107 | 116.7 | 122.8 | 131.9 | 112.8 | 0.758 | 0.116455186 | 0.970681458 | 0.583848515 | 1.184337738 | 0.155853022 |
| A0A1U7RAW3 | prefoldin subunit 6 OS=Mesocricetus auratus OX=10036 GN=Pfdn6 PE=3 SV=1 | 4.777 | 8 | 1 | 6 | 1 | 127 | 14.4 | 8.88 | 19.03 | 1 | 159.4 | 49.4 | 72.8 | 123.9 | 50.3 | 66.6 | 177.6 | 56.2 | 73.8 | 0.954 | 0.754603076 | 1.092329545 | 0.872280512 | 1.277408638 | 0.64498789 |
| A0A1U7QER6 | ATP-dependent 6-phosphofructokinase OS=Mesocricetus auratus OX=10036 GN=Pfkl PE=3 SV=1 | 10.725 | 6 | 4 | 14 | 4 | 780 | 85.3 | 7.42 | 51.94 | 4 | 374.3 | 439 | 339.4 | 319.9 | 386.1 | 286.8 | 371.4 | 370.5 | 311.4 | 0.879 | 0.266140349 | 0.913767676 | 0.407565253 | 1.060938759 | 0.602256954 |
| A0A1U7QJS1 | Profilin OS=Mesocricetus auratus OX=10036 GN=Pfn1 PE=3 SV=1 | 44.294 | 46 | 6 | 62 | 6 | 140 | 15.1 | 8.62 | 230.81 | 6 | 288 | 286.8 | 0 | 388.1 | 264.4 | 0 | 264.3 | 275.4 | 0 | 1.115 | 0.870918758 | 0.938935282 | 0.933370184 | 0.827126437 | 0.80965071 |

**Supplementary Table 1: List of differentially expressed proteins identified across the 4dpi infected and 14dpi infected**

| Accession | Description | Sum PEP Score | Coverage [%] | # Peptides | # PSMs | # Unique Peptides | # AAs | MW [kDa] | calc. pI | Score Sequest HT: Sequest HT | # Peptides (by Search Engine): Sequest HT | Mock-infected (4dpi) Replicate_R1 | Mock-infected (4dpi) Replicate_R2 | Mock-infected (4dpi) Replicate_R3 | SARS-CoV-2-infected (4dpi) Replicate_R1 | SARS-CoV-2-infected (4dpi) Replicate_R2 | SARS-CoV-2-infected (4dpi) Replicate_R3 | SARS-CoV-2-(14dpi) Replicate_R1 | SARS-CoV-2-infected (14dpi) Replicate_R2 | SARS-CoV-2-infected (14dpi) Replicate_R3 | Abundance Ratio: SARS-CoV-2-infected (4dpi) / Mock-infected (4dpi) | P-Value: SARS-CoV-2-infected (4dpi) / Mock-infected (4dpi) | Abundance Ratio: SARS-CoV-2-infected (14dpi) / Mock-infected (4dpi) | P-Value: SARS-CoV-2-infected (14dpi) / Mock-infected (4dpi) | Abundance Ratio: SARS-CoV-2-infected (14dpi) / SARS-CoV-2-infected (4dpi) | P-Value: SARS-CoV-2-infected (14dpi) / SARS-CoV-2-infected (4dpi) |
|---|---|---|---|---|---|---|---|---|---|---|---|---|---|---|---|---|---|---|---|---|---|---|---|---|---|---|
| A0A1U7QRI5 | 6-phosphogluconate dehydrogenase, decarboxylating OS=Mesocricetus auratus OX=10036 GN=Pgd PE=3 SV=1 | 33.055 | 18 | 9 | 53 | 9 | 483 | 53.2 | 7.23 | 117.67 | 9 | 842 | 682 | 814.2 | 1214.4 | 861.5 | 1363.8 | 732.8 | 600.1 | 680.3 | 1.534 | 0.12135978 | 0.861004191 | 0.162985 | 0.5852836 | 0.077357669 |
| A0A1U7QEP0 | Phosphoglycerate kinase OS=Mesocricetus auratus OX=10036 GN=Pgk2 PE=3 SV=1 | 15.706 | 13 | 4 | 15 | 2 | 417 | 44.8 | 7.69 | 51.55 | 4 | 108 | 45.7 | 34.4 | 168.3 | 63.6 | 38.7 | 121.4 | 45 | 34.9 | 1.228 | 0.588351977 | 1.070175439 | 0.907839898 | 0.743902439 | 0.659724995 |
| A0A1U7RCY5 | 6-phosphogluconolactonase OS=Mesocricetus auratus OX=10036 GN=Pgls PE=3 SV=1 | 15.106 | 19 | 4 | 16 | 4 | 257 | 27.2 | 5.85 | 54.19 | 4 | 140.2 | 251.8 | 290 | 147.9 | 246.1 | 277.9 | 138.8 | 243.2 | 279.6 | 0.958 | 0.957701081 | 0.970087977 | 0.917497837 | 0.984670338 | 0.955300201 |
| A0A1U7QI58 | phosphoglucomutase-2 OS=Mesocricetus auratus OX=10036 GN=Pgm2 PE=3 SV=1 | 19.92 | 15 | 6 | 14 | 6 | 626 | 69.3 | 6.28 | 39.25 | 6 | 112.8 | 177.6 | 112.2 | 93.8 | 141.8 | 107.9 | 120.5 | 187.2 | 136.3 | 0.944 | 0.496368592 | 1.102831595 | 0.66539977 | 1.292576419 | 0.253067735 |
| A0A1U7QSE1 | Phosphoacetylglucosamine mutase OS=Mesocricetus auratus OX=10036 GN=Pgm3 PE=3 SV=1 | 8.269 | 6 | 3 | 13 | 3 | 542 | 59.5 | 5.9 | 25.16 | 3 | 235.7 | 253.4 | 315.2 | 198.2 | 192.2 | 278.9 | 197.9 | 198.3 | 244.4 | 0.841 | 0.291061798 | 0.796468979 | 0.141586903 | 0.957119378 | 0.783363116 |
| A0A1U7QB05 | phosphoglucomutase-like protein 5 OS=Mesocricetus auratus OX=10036 GN=Pgm5 PE=3 SV=1 | 22.272 | 14 | 7 | 20 | 7 | 567 | 62.2 | 7.18 | 52.27 | 7 | 353.3 | 772.7 | 358.2 | 278.7 | 640.5 | 288.5 | 334.2 | 688.6 | 343 | 0.814 | 0.641532189 | 0.920226385 | 0.838788903 | 1.130909994 | 0.767672687 |
| A0A1U7R899 | membrane-associated progesterone receptor component 1 OS=Mesocricetus auratus OX=10036 GN=Pgrmc1 PE=4 SV=1 | 9.467 | 16 | 3 | 8 | 3 | 195 | 21.6 | 4.65 | 23.88 | 3 | 317.9 | 213.9 | 349.2 | 244.9 | 166.8 | 247 | 275.1 | 178.9 | 294.2 | 0.77 | 0.214248249 | 0.849262202 | 0.461260869 | 1.135873691 | 0.541211612 |
| A0A1U7QCX1 | membrane-associated progesterone receptor component 2 OS=Mesocricetus auratus OX=10036 GN=Pgrmc2 PE=4 SV=1 | 4.362 | 15 | 2 | 3 | 2 | 220 | 23.4 | 5.07 | 4.52 | 2 | 121.5 | 0 | 179 | 100.5 | 0 | 144 | 115.2 | 0 | 156.5 | 0.815 | 0.7973893 | 0.904159734 | 0.89841736 | 1.111247444 | 0.893143065 |
| A0A1U7QNJ2 | Prohibitin OS=Mesocricetus auratus OX=10036 GN=Phb PE=3 SV=1 | 16.564 | 19 | 4 | 13 | 4 | 272 | 29.8 | 5.76 | 35.56 | 4 | 52.8 | 38.1 | 37.5 | 34 | 32.3 | 27.4 | 58.9 | 27.2 | 27.7 | 0.726 | 0.134060499 | 0.886292835 | 0.704670241 | 1.214514408 | 0.590411339 |
| A0A1U7Q3L1 | Prohibitin OS=Mesocricetus auratus OX=10036 GN=Phb2 PE=3 SV=1 | 15.218 | 25 | 5 | 14 | 5 | 299 | 33.3 | 9.83 | 33.19 | 5 | 88.5 | 86.2 | 0 | 75.9 | 75.3 | 0 | 76 | 66.4 | 0 | 0.865 | 0.848951457 | 0.81511162 | 0.789734012 | 0.941798942 | 0.936755101 |
| A0A1U7QIX3 | Non-specific serine/threonine protein kinase OS=Mesocricetus auratus OX=10036 GN=Pik3r4 PE=4 SV=1 | 2.036 | 1 | 1 | 2 | 1 | 1358 | 152.7 | 7.03 | 0 | 1 | 0 | 82.6 | 97.2 | 0 | 76.2 | 75.7 | 0 | 74 | 92.6 | 0.847 | 0.825576792 | 0.926585095 | 0.920536525 | 1.096774194 | 0.903587888 |
| A0A1U7QGS3 | Peptidyl-prolyl cis-trans isomerase OS=Mesocricetus auratus OX=10036 GN=Pin1 PE=4 SV=1 | 1.613 | 9 | 1 | 3 | 1 | 165 | 18.3 | 8.82 | 0 | 1 | 0 | 0 | 35.3 | 0 | 0 | 37.5 | 0 | 0 | 48.5 | 1.061 | 0.967980984 | 1.373937677 | 0.837568905 | 1.293333333 | 0.866849139 |
| A0A1U7Q896 | phosphatidylinositol transfer protein alpha isoform OS=Mesocricetus auratus OX=10036 GN=Pitpna PE=4 SV=1 | 4.559 | 8 | 2 | 2 | 1 | 270 | 31.7 | 6.55 | 2.73 | 2 | 0 | 111.8 | 49.2 | 0 | 77.1 | 55 | 0 | 100.4 | 51.7 | 0.878 | 0.821222761 | 0.944720497 | 0.948864472 | 1.151400454 | 0.866053097 |
| A0A1U7QTQ3 | phosphatidylinositol transfer protein beta isoform isoform X1 OS=Mesocricetus auratus OX=10036 GN=Pitpnb PE=4 SV=1 | 5.598 | 8 | 2 | 3 | 1 | 272 | 31.6 | 6.54 | 10.04 | 2 | 0 | 64.6 | 47.8 | 0 | 57.1 | 44.9 | 0 | 47.6 | 41.1 | 0.912 | 0.900430012 | 0.789145907 | 0.763556677 | 0.869607843 | 0.85600501 |
| A0A1U7R408 | phosphatidylinositol transfer protein beta isoform isoform X2 OS=Mesocricetus auratus OX=10036 GN=Pitpnb PE=4 SV=1 | 4.687 | 8 | 2 | 3 | 1 | 271 | 31.5 | 7.2 | 5.59 | 2 | 0 | 45.7 | 44 | 0 | 37.6 | 35.3 | 0 | 39.9 | 33.2 | 0.812 | 0.786495113 | 0.814938685 | 0.789999714 | 1.002743484 | 0.997114434 |
| A0A1U7QDU2 | Pitrilysin metalloproteinase 1 OS=Mesocricetus auratus OX=10036 GN=Pitrm1 PE=3 SV=1 | 2.865 | 2 | 2 | 3 | 2 | 1036 | 117.7 | 7.21 | 6.58 | 2 | 58.3 | 0 | 89.6 | 74.4 | 0 | 91.2 | 54.5 | 0 | 74.1 | 1.139 | 0.885358231 | 0.869506423 | 0.860837541 | 0.776570048 | 0.7482223 |
| A0A1U8C9I9 | Pyruvate kinase OS=Mesocricetus auratus OX=10036 GN=Pklr PE=3 SV=1 | 8.906 | 4 | 3 | 6 | 1 | 543 | 58.6 | 6.71 | 14.29 | 3 | 0 | 66.5 | 0 | 0 | 66.6 | 0 | 0 | 42.3 | 0 | 1.002 | 0.999203111 | 0.636090226 | 0.77671179 | 0.635135135 | 0.77607092 |
| A0A1U8BTR6 | HRAS-like suppressor 3 OS=Mesocricetus auratus OX=10036 GN=LOC101830643 PE=3 SV=1 | 2.837 | 8 | 1 | 1 | 1 | 162 | 17.7 | 8.57 | 3.36 | 1 | 60.4 | 0 | 0 | 83.3 | 0 | 0 | 40.8 | 0 | 0 | 1.38 | 0.835776124 | 0.675496689 | 0.803052954 | 0.489795918 | 0.678881054 |
| A0A1U7QW38 | phospholipase A-2-activating protein OS=Mesocricetus auratus OX=10036 GN=Plaa PE=3 SV=1 | 1.648 | 1 | 1 | 1 | 1 | 795 | 87.1 | 6.16 | 2.21 | 1 | 0 | 0 | 51.3 | 0 | 0 | 37.7 | 0 | 0 | 48 | 0.735 | 0.842168155 | 0.935672515 | 0.964796547 | 1.273209549 | 0.874614898 |
| A0A1U7QFP3 | Phospholipase B-like OS=Mesocricetus auratus OX=10036 GN=Plbd1 PE=3 SV=1 | 5.837 | 2 | 1 | 6 | 1 | 553 | 63.2 | 8.4 | 17.48 | 1 | 85.3 | 122.6 | 184 | 137.2 | 216.9 | 267.8 | 89.4 | 129.1 | 155.5 | 1.455 | 0.188239917 | 0.954325083 | 0.872610055 | 0.601382859 | 0.148906089 |
| A0A1U7QTX0 | Phospholipase B-like OS=Mesocricetus auratus OX=10036 GN=Plbd2 PE=3 SV=1 | 4.222 | 3 | 1 | 3 | 1 | 585 | 65.5 | 5.91 | 9.02 | 1 | 0 | 45.8 | 0 | 0 | 49.9 | 0 | 0 | 49.8 | 0 | 1.088 | 0.954655437 | 1.087336245 | 0.955710786 | 0.997995992 | 0.99893615 |
| A0A1U7R9D1 | Phosphoinositide phospholipase C OS=Mesocricetus auratus OX=10036 GN=Plcd1 PE=4 SV=1 | 29.872 | 7 | 5 | 19 | 5 | 782 | 89 | 6.8 | 60.59 | 5 | 172.2 | 0 | 213.5 | 134.2 | 0 | 141.8 | 136.9 | 0 | 239 | 0.719 | 0.673756637 | 0.974591652 | 0.974283476 | 1.361956522 | 0.712158907 |
| A0A1U7Q917 | pleckstrin homology domain-containing family A member 2 OS=Mesocricetus auratus OX=10036 GN=Plekha2 PE=4 SV=1 | 2.775 | 3 | 1 | 2 | 1 | 424 | 47.3 | 8.63 | 5.6 | 1 | 0 | 35.3 | 48.7 | 0 | 38 | 43.3 | 0 | 31.1 | 35.5 | 0.977 | 0.96614132 | 0.792857143 | 0.768426371 | 0.819188192 | 0.79532582 |
| A0A1U8CU45 | plasmalemma vesicle-associated protein OS=Mesocricetus auratus OX=10036 GN=Plvap PE=4 SV=1 | 8.275 | 7 | 2 | 11 | 2 | 440 | 49.7 | 8.53 | 32.78 | 2 | 190.1 | 222.6 | 0 | 167.2 | 218.5 | 0 | 136.7 | 153 | 0 | 0.929 | 0.929655095 | 0.701962685 | 0.656430868 | 0.751101893 | 0.717540952 |
| A0A1U8BML8 | protein PML isoform X1 OS=Mesocricetus auratus OX=10036 GN=Pml PE=4 SV=1 | 17.869 | 7 | 4 | 13 | 4 | 874 | 97.2 | 5.58 | 43.2 | 4 | 0 | 169.1 | 267.7 | 0 | 125.6 | 194.5 | 0 | 177.5 | 224.8 | 0.734 | 0.709890517 | 0.921016484 | 0.91727569 | 1.256794752 | 0.774071948 |
| A0A1U7RD08 | Phosphomannomutase OS=Mesocricetus auratus OX=10036 GN=Pmm2 PE=3 SV=1 | 6.067 | 10 | 3 | 13 | 3 | 242 | 27.6 | 6.57 | 22.42 | 3 | 218.3 | 770.6 | 149.2 | 647.2 | 700.1 | 145.6 | 755 | 760.3 | 158.8 | 0.909 | 0.677998814 | 1.470960373 | 0.558389514 | 1.121374506 | 0.832018003 |
| A0A1U8CYQ4 | Podocalyxin OS=Mesocricetus auratus OX=10036 GN=Podxl PE=3 SV=1 | 9.698 | 4 | 2 | 7 | 2 | 689 | 71.4 | 5.88 | 21.62 | 2 | 74.6 | 101.7 | 203.6 | 73.9 | 67.8 | 160.6 | 60.4 | 89.9 | 171.3 | 0.713 | 0.630025409 | 0.846538563 | 0.725086171 | 1.063843864 | 0.892581938 |
| A0A1U8CKP9 | DNA polymerase epsilon catalytic subunit OS=Mesocricetus auratus OX=10036 GN=Pole PE=3 SV=1 | 1.852 | 0 | 1 | 1 | 1 | 2192 | 251.1 | 6.18 | 1.68 | 1 | 0 | 0 | 89.5 | 0 | 0 | 77.8 | 0 | 0 | 75.4 | 0.869 | 0.926241314 | 0.842458101 | 0.910067814 | 0.969151671 | 0.983388641 |
| A0A1U7RA96 | serum paraoxonase/arylesterase 1 OS=Mesocricetus auratus OX=10036 GN=Pon1 PE=3 SV=1 | 2.648 | 5 | 1 | 3 | 1 | 355 | 39.8 | 5.73 | 5.39 | 1 | 0 | 0 | 72.8 | 0 | 0 | 72.5 | 0 | 0 | 81.8 | 0.995 | 0.997810072 | 1.123626374 | 0.938495432 | 1.128275862 | 0.936339605 |
| A0A1U7QNV3 | serum paraoxonase/arylesterase 2 OS=Mesocricetus auratus OX=10036 GN=Pon2 PE=3 SV=1 | 4.595 | 5 | 1 | 5 | 1 | 354 | 39.5 | 5.52 | 21.96 | 1 | 61.1 | 102.5 | 129 | 54.8 | 73.1 | 94.3 | 45.7 | 79.6 | 106.7 | 0.731 | 0.375076873 | 0.792891319 | 0.488728615 | 1.04410441 | 0.885191646 |
| A0A1U7R076 | serum paraoxonase/lactonase 3 OS=Mesocricetus auratus OX=10036 GN=Pon3 PE=3 SV=1 | 19.247 | 19 | 4 | 15 | 4 | 354 | 39.7 | 6.07 | 35.58 | 4 | 0 | 115.3 | 115.6 | 0 | 90.1 | 86.6 | 0 | 102.1 | 108.3 | 0.765 | 0.729466426 | 0.911216977 | 0.902022694 | 1.190718732 | 0.818812237 |
| A0A1U8C9X6 | NADPH--cytochrome P450 reductase OS=Mesocricetus auratus OX=10036 GN=LOC101843888 PE=3 SV=1 | 14.612 | 5 | 4 | 28 | 4 | 696 | 78.8 | 5.71 | 59.02 | 4 | 648.1 | 529.3 | 434.1 | 597.9 | 443.6 | 352.6 | 581 | 492.5 | 441.5 | 0.838 | 0.487385851 | 0.940117903 | 0.689938002 | 1.086722617 | 0.65653695 |
| A0A1U7QYR1 | periostin isoform X2 OS=Mesocricetus auratus OX=10036 GN=Postn PE=4 SV=1 | 86.491 | 27 | 18 | 96 | 18 | 780 | 86.7 | 7.39 | 267.02 | 18 | 2845 | 1974.1 | 2050.3 | 2216.6 | 1847.2 | 2110.7 | 2921.2 | 2417.4 | 3009.6 | 0.936 | 0.503079617 | 1.215273532 | 0.224424566 | 1.3520447 | 0.038151834 |
| A0A1U8CDA2 | Inorganic diphosphatase OS=Mesocricetus auratus OX=10036 GN=Ppa2 PE=3 SV=1 | 2.778 | 3 | 1 | 7 | 1 | 330 | 37.8 | 7.25 | 20.21 | 1 | 30.4 | 0 | 0 | 26.1 | 0 | 0 | 31.1 | 0 | 0 | 0.856 | 0.919814953 | 1.023026316 | 0.987929259 | 1.191570881 | 0.908097623 |
| A0A1U7Q8V7 | liprin-alpha-2 isoform X12 OS=Mesocricetus auratus OX=10036 GN=Ppfia2 PE=3 SV=1 | 1.52 | 2 | 1 | 3 | 1 | 1186 | 135.3 | 6.52 | 7.73 | 1 | 24.3 | 0 | 0 | 40.8 | 0 | 0 | 31.6 | 0 | 0 | 1.678 | 0.749485214 | 1.300411523 | 0.864162022 | 0.774509804 | 0.867686749 |
| A0A1U7QK19 | Peptidyl-prolyl cis-trans isomerase OS=Mesocricetus auratus OX=10036 GN=Ppib PE=3 SV=1 | 21.079 | 30 | 7 | 32 | 7 | 216 | 23.7 | 9.51 | 72.11 | 7 | 602.1 | 845.7 | 608 | 571.3 | 796.1 | 615.1 | 656.9 | 818.3 | 670 | 1.011 | 0.82878012 | 1.04348672 | 0.773042997 | 1.082068096 | 0.56541173 |
| A0A1U7QR01 | Protein-serine/threonine phosphatase OS=Mesocricetus auratus OX=10036 GN=Ppm1a PE=3 SV=1 | 5.434 | 7 | 2 | 2 | 2 | 382 | 42.3 | 5.29 | 5.21 | 2 | 0 | 68.7 | 0 | 0 | 51.2 | 0 | 0 | 43.8 | 0 | 0.746 | 0.848902403 | 0.637554585 | 0.777694212 | 0.85546875 | 0.917956504 |
| A0A1U7QNJ4 | protein phosphatase 1F OS=Mesocricetus auratus OX=10036 GN=Ppm1f PE=3 SV=1 | 22.945 | 13 | 5 | 25 | 5 | 448 | 49.4 | 5.2 | 60.32 | 5 | 27.7 | 29.9 | 160.8 | 25.2 | 35 | 125.4 | 29.4 | 34.4 | 156.8 | 0.979 | 0.851274144 | 1.01007326 | 0.990925355 | 1.188577586 | 0.835619858 |
| A0A1U7Q994 | protein phosphatase 1G OS=Mesocricetus auratus OX=10036 GN=Ppm1g PE=3 SV=1 | 12.331 | 7 | 3 | 13 | 3 | 547 | 59.3 | 4.34 | 30.78 | 3 | 339.7 | 525.9 | 287.2 | 270.7 | 413.4 | 236.3 | 265.2 | 446.2 | 233.6 | 0.786 | 0.443656567 | 0.819743234 | 0.519538532 | 1.02672751 | 0.928455621 |
| A0A1U7QKA8 | Protein phosphatase methylesterase 1 OS=Mesocricetus auratus OX=10036 GN=Ppme1 PE=3 SV=1 | 10.504 | 11 | 3 | 5 | 3 | 386 | 42.2 | 5.97 | 16.42 | 3 | 0 | 0 | 157.6 | 0 | 0 | 130.3 | 0 | 0 | 137.5 | 0.827 | 0.900458777 | 0.872461929 | 0.928141898 | 1.055257099 | 0.97150725 |
| A0A1U7RES6 | E3 ubiquitin-protein ligase PPP1R11 OS=Mesocricetus auratus OX=10036 GN=Ppp1r11 PE=4 SV=1 | 9.945 | 13 | 1 | 12 | 1 | 126 | 14 | 6.86 | 47.06 | 1 | 148.6 | 175.9 | 156.8 | 156.7 | 167.9 | 148.5 | 143.2 | 170.7 | 107.6 | 1.055 | 0.796724157 | 0.875753169 | 0.397739123 | 0.89093215 | 0.449877031 |
| A0A1U7Q8W0 | Protein phosphatase 1 regulatory subunit OS=Mesocricetus auratus OX=10036 GN=Ppp1r12a PE=4 SV=1 | 15.261 | 7 | 6 | 14 | 6 | 1031 | 115.1 | 5.47 | 25.3 | 6 | 529.8 | 111.4 | 468.5 | 361.7 | 84.1 | 332.8 | 438.9 | 98.7 | 399.7 | 0.71 | 0.526917385 | 0.844642696 | 0.751553282 | 1.203827382 | 0.723511354 |
| A0A1U7QXP4 | Serine/threonine-protein phosphatase 2A 55 kDa regulatory subunit B OS=Mesocricetus auratus OX=10036 GN=Ppp2r2a PE=3 SV=1 | 9.138 | 9 | 3 | 10 | 3 | 447 | 51.7 | 6.11 | 19.06 | 3 | 90.6 | 148.7 | 241.7 | 78.2 | 153.6 | 239.9 | 77.4 | 158.7 | 229.4 | 0.993 | 0.963797313 | 0.967775468 | 0.937758673 | 0.986856053 | 0.975835311 |
| A0A1U7R8U7 | Serine/threonine-protein phosphatase OS=Mesocricetus auratus OX=10036 GN=Ppp3ca PE=3 SV=1 | 1.603 | 2 | 1 | 1 | 1 | 521 | 58.6 | 5.86 | 0 | 1 | 0 | 0 | 68.1 | 0 | 0 | 62.7 | 0 | 0 | 70.4 | 0.921 | 0.956297616 | 1.033773862 | 0.98239178 | 1.122807018 | 0.938876509 |
| A0A1U7RHE5 | peroxiredoxin-2 OS=Mesocricetus auratus OX=10036 GN=Prdx2 PE=3 SV=1 | 43.988 | 31 | 8 | 76 | 8 | 198 | 21.8 | 5.41 | 237.18 | 8 | 1049 | 1363 | 1086.2 | 922 | 1164.7 | 954.5 | 1023.2 | 1325.9 | 1078.3 | 0.872 | 0.293663653 | 0.97976102 | 0.870624581 | 1.126989346 | 0.346747538 |
| A0A1U7RE88 | thioredoxin-dependent peroxide reductase, mitochondrial OS=Mesocricetus auratus OX=10036 GN=Prdx3 PE=4 SV=1 | 15.257 | 16 | 4 | 20 | 4 | 257 | 28.1 | 7.62 | 39.9 | 4 | 353.8 | 399.4 | 330.5 | 214.5 | 259.1 | 194.9 | 305.3 | 346.9 | 280.1 | 0.64 | 0.007639467 | 0.860293439 | 0.146861247 | 1.394614809 | 0.031938976 |
| A0A1U7QS22 | peroxiredoxin-4 OS=Mesocricetus auratus OX=10036 GN=Prdx4 PE=4 SV=1 | 7.185 | 7 | 2 | 19 | 1 | 274 | 31.1 | 7.15 | 55.02 | 2 | 43.7 | 47.3 | 57.5 | 44.8 | 53.5 | 59 | 43.3 | 50.8 | 54.5 | 1.096 | 0.642153777 | 1.000673401 | 0.995284174 | 0.944691672 | 0.613821716 |
| A0A1U7Q712 | peroxiredoxin-6 OS=Mesocricetus auratus OX=10036 GN=Prdx6 PE=3 SV=1 | 64.654 | 56 | 14 | 83 | 14 | 224 | 24.8 | 6.01 | 222.12 | 14 | 5864.8 | 3684 | 3027.7 | 1278.4 | 1636.9 | 1592.1 | 5366.2 | 3674.9 | 3087.2 | 0.433 | 0.085935649 | 0.964362104 | 0.898502191 | 2.690752984 | 0.061717366 |
| A0A1U7QJ85 | prolargin OS=Mesocricetus auratus OX=10036 GN=Prelp PE=4 SV=1 | 20.662 | 16 | 7 | 23 | 7 | 377 | 43.1 | 9.51 | 57.36 | 7 | 453.5 | 400.4 | 330.1 | 352 | 306.2 | 229.9 | 356.8 | 349.4 | 318.2 | 0.765 | 0.122254679 | 0.865202703 | 0.271972906 | 1.153473708 | 0.330297713 |
| A0A1U7R6F7 | cAMP-dependent protein kinase catalytic subunit beta OS=Mesocricetus auratus OX=10036 GN=Prkacb PE=3 SV=1 | 10.056 | 10 | 3 | 15 | 2 | 351 | 40.7 | 8.56 | 38.06 | 3 | 352.5 | 297.6 | 638.6 | 285.9 | 205.1 | 474.9 | 286.4 | 311.3 | 577.2 | 0.744 | 0.465573835 | 0.911693955 | 0.801194004 | 1.216378507 | 0.601144831 |
| A0A1U8BN84 | cAMP-dependent protein kinase type I-alpha regulatory subunit OS=Mesocricetus auratus OX=10036 GN=Prkar1a PE=3 SV=1 | 18.514 | 10 | 4 | 19 | 4 | 381 | 43 | 5.35 | 50.39 | 4 | 426 | 516.9 | 426.8 | 273.2 | 362.5 | 357.5 | 311.4 | 557.8 | 391.9 | 0.838 | 0.040001137 | 0.920712565 | 0.679836656 | 1.269734193 | 0.346505307 |
| A0A1U8CBY7 | cAMP-dependent protein kinase type I-beta regulatory subunit OS=Mesocricetus auratus OX=10036 GN=Prkar1b PE=3 SV=1 | 1.713 | 3 | 1 | 3 | 1 | 381 | 43.2 | 5.96 | 6.32 | 1 | 36.6 | 36.6 | 29 | 26.1 | 30.4 | 28.8 | 26 | 28.8 | 37.9 | 0.755 | 0.142647443 | 0.90704501 | 0.515307339 | 1.086752638 | 0.571724608 |
| A0A1U7QJ32 | cAMP-dependent protein kinase type II-alpha regulatory subunit OS=Mesocricetus auratus OX=10036 GN=Prkar2a PE=3 SV=1 | 21.058 | 21 | 6 | 20 | 6 | 402 | 45.5 | 4.93 | 60.67 | 6 | 105.2 | 62.6 | 45.9 | 77.8 | 62.8 | 40 | 91 | 47.8 | 41.5 | 0.893 | 0.628920588 | 0.84370613 | 0.661143614 | 0.99833887 | 0.996088847 |
| A0A1U7QLD2 | Protein kinase C OS=Mesocricetus auratus OX=10036 GN=Prkca PE=3 SV=1 | 9.052 | 6 | 2 | 12 | 2 | 672 | 76.7 | 7.05 | 9.49 | 2 | 517.2 | 443 | 437.6 | 372.4 | 322.7 | 333.5 | 488.6 | 443 | 412.7 | 0.72 | 0.022214639 | 0.961725569 | 0.626807722 | 1.30692203 | 0.021508764 |
| A0A1U8CW45 | Protein kinase C OS=Mesocricetus auratus OX=10036 GN=Prkcd PE=3 SV=1 | 4.293 | 3 | 2 | 4 | 2 | 700 | 80.1 | 8.21 | 10.96 | 2 | 38.7 | 0 | 0 | 47.7 | 0 | 0 | 37.6 | 0 | 0 | 1.234 | 0.890883585 | 0.971576227 | 0.984712385 | 0.788259958 | 0.876413567 |
| A0A1U7R791 | protein kinase C delta-binding protein OS=Mesocricetus auratus OX=10036 GN=Cavin3 PE=3 SV=1 | 5.357 | 4 | 1 | 8 | 1 | 264 | 27.9 | 6.16 | 34.61 | 1 | 280 | 265.1 | 180.2 | 294.6 | 260.3 | 176.2 | 239.1 | 220 | 179.5 | 0.994 | 0.969147779 | 0.880463257 | 0.474772269 | 0.87347832 | 0.491202461 |
| A0A1U7QNF6 | Protein kinase C OS=Mesocricetus auratus OX=10036 GN=Prkci PE=3 SV=1 | 2.07 | 1 | 1 | 4 | 1 | 596 | 68.1 | 5.85 | 10.74 | 1 | 44.7 | 176.4 | 0 | 35 | 136.6 | 0 | 50.6 | 160.6 | 0 | 0.779 | 0.818171384 | 0.955223881 | 0.965215622 | 1.230769231 | 0.843639749 |
| A0A1U8C607 | Glucosidase 2 subunit beta OS=Mesocricetus auratus OX=10036 GN=Prkcsh PE=4 SV=1 | 12.783 | 6 | 3 | 17 | 3 | 522 | 58.7 | 4.51 | 39.24 | 3 | 430.4 | 384.6 | 359.2 | 356.8 | 276.9 | 289.8 | 448.1 | 393.3 | 353.7 | 0.807 | 0.06295538 | 1.017799353 | 0.850038585 | 1.294098538 | 0.07085792 |
| A0A1U8BS80 | cGMP-dependent protein kinase 1 OS=Mesocricetus auratus OX=10036 GN=LOC101826743 PE=4 SV=1 | 6.538 | 7 | 2 | 9 | 2 | 283 | 32.4 | 7.83 | 17.51 | 2 | 106.3 | 242.5 | 143.3 | 86.4 | 219.8 | 134.6 | 108.8 | 204.1 | 99.9 | 0.887 | 0.77665938 | 0.838853891 | 0.642579177 | 0.936479129 | 0.864720328 |
| A0A1U7Q415 | pre-mRNA-processing factor 40 homolog B isoform X8 OS=Mesocricetus auratus OX=10036 GN=Prpf40b PE=4 SV=1 | 3.363 | 1 | 1 | 1 | 1 | 1000 | 113.2 | 6.98 | 3.11 | 1 | 0 | 141.2 | 0 | 0 | 93.4 | 0 | 0 | 118.6 | 0 | 0.662 | 0.793705118 | 0.839943343 | 0.908534206 | 1.269807281 | 0.875946451 |
| A0A1U8C2F7 | protein PRRC2C isoform X12 OS=Mesocricetus auratus OX=10036 GN=Prrc2c PE=4 SV=1 | 1.983 | 0 | 1 | 2 | 1 | 2742 | 300.3 | 9.2 | 2.33 | 1 | 0 | 63.4 | 57.4 | 0 | 45.7 | 57.5 | 0 | 47.2 | 49.4 | 0.85 | 0.837370226 | 0.799668874 | 0.771277213 | 0.936046512 | 0.930862965 |
| A0A1U7QV17 | periaxin OS=Mesocricetus auratus OX=10036 GN=Prx PE=4 SV=1 | 268.344 | 57 | 54 | 238 | 54 | 1362 | 144.1 | 8.6 | 717.91 | 54 | 1791.6 | 2389.8 | 2152.2 | 1382 | 1836.3 | 1456.5 | 1739 | 2416.6 | 2151.1 | 0.771 | 0.071535085 | 0.99575281 | 0.974447135 | 1.349084453 | 0.095106206 |
| A0A1U7R928 | PC4 and SFRS1-interacting protein isoform X1 OS=Mesocricetus auratus OX=10036 GN=Psip1 PE=3 SV=1 | 13.652 | 9 | 4 | 17 | 3 | 529 | 59.8 | 9.13 | 26.9 | 4 | 309.4 | 708.3 | 601.6 | 296.7 | 650.6 | 580.7 | 280.7 | 589.4 | 494 | 0.919 | 0.859371329 | 0.842401037 | 0.603231642 | 0.892735602 | 0.719712469 |

Supplementary Table 1: List of differentially expressed proteins identified across the 4dpi infected and 14dpi infected

| Accession | Description | Sum PEP Score | Coverage [%] | # Peptides | # PSMs | # Unique Peptides | # AAs | MW [kDa] | calc. pI | Score Sequest HT: Sequest HT | # Peptides (by Search Engine): Sequest HT | Mock-infected (4dpi) Replicate_R1 | Mock-infected (4dpi) Replicate_R2 | Mock-infected (4dpi) Replicate_R3 | SARS-CoV-2-infected (4dpi) Replicate_R1 | SARS-CoV-2-infected (4dpi) Replicate_R2 | SARS-CoV-2-infected (4dpi) Replicate_R3 | SARS-CoV-2-(14dpi) Replicate_R1 | SARS-CoV-2-infected (14dpi) Replicate_R2 | SARS-CoV-2-infected (14dpi) Replicate_R3 | Abundance Ratio: SARS-CoV-2-infected (4dpi) / Mock-infected (4dpi) | P-Value: SARS-CoV-2-infected (4dpi) / Mock-infected (4dpi) | Abundance Ratio: SARS-CoV-2-infected (14dpi) / Mock-infected (4dpi) | P-Value: SARS-CoV-2-infected (14dpi) / Mock-infected (4dpi) | Abundance Ratio: SARS-CoV-2-infected (14dpi) / SARS-CoV-2-infected (4dpi) | P-Value: SARS-CoV-2-infected (14dpi) / SARS-CoV-2-infected (4dpi) |
|---|---|---|---|---|---|---|---|---|---|---|---|---|---|---|---|---|---|---|---|---|---|---|---|---|---|
| A0A1U7QGL6 | Proteasome subunit alpha type OS=Mesocricetus auratus OX=10036 GN=Psma2 PE=3 SV=1 | 4.786 | 7 | 2 | 7 | 2 | 234 | 25.9 | 7.43 | 20.25 | 2 | 293.3 | 334.1 | 78.6 | 244.5 | 291.5 | 71.8 | 261.3 | 298.5 | 87.8 | 0.834 | 0.768349574 | 0.917280453 | 0.858890654 | 1.065482066 | 0.893636737 |
| A0A1U7QKR0 | Proteasome subunit alpha type OS=Mesocricetus auratus OX=10036 GN=Psma4 PE=3 SV=1 | 9.398 | 4 | 1 | 23 | 1 | 261 | 29.5 | 7.72 | 73.03 | 1 | 100.9 | 121.4 | 89.1 | 109.3 | 142.9 | 98.5 | 97.5 | 108.1 | 57.1 | 1.094 | 0.472911757 | 0.843609505 | 0.432192562 | 0.749073282 | 0.227168106 |
| A0A1U7R6M5 | Proteasome subunit alpha type OS=Mesocricetus auratus OX=10036 GN=Psma6 PE=3 SV=1 | 14.358 | 15 | 4 | 10 | 4 | 246 | 27.4 | 6.76 | 26.31 | 4 | 80.5 | 76.4 | 0 | 86.6 | 66 | 0 | 81.6 | 71.8 | 0 | 0.964 | 0.970938494 | 0.977692798 | 0.976163421 | 1.005242464 | 0.994544438 |
| A0A1U8CFN2 | Proteasome subunit beta OS=Mesocricetus auratus OX=10036 GN=Psmb1 PE=3 SV=1 | 10.196 | 15 | 3 | 8 | 3 | 240 | 26.3 | 7.8 | 29.63 | 3 | 150.9 | 265.1 | 157.5 | 126.8 | 222.8 | 148.9 | 138.4 | 262.9 | 142.8 | 0.841 | 0.624745315 | 0.948735833 | 0.867505104 | 1.091474423 | 0.778081053 |
| A0A1U7R248 | Proteasome subunit beta OS=Mesocricetus auratus OX=10036 GN=Psmb2 PE=3 SV=1 | 7.836 | 23 | 3 | 11 | 3 | 201 | 22.8 | 7.88 | 24.33 | 3 | 196.3 | 0 | 0 | 245.5 | 0 | 0 | 175.5 | 0 | 0 | 1.251 | 0.88355125 | 0.894039735 | 0.940876772 | 0.714867617 | 0.829081108 |
| A0A1U7RCL6 | Proteasome subunit beta OS=Mesocricetus auratus OX=10036 GN=Psmb4 PE=3 SV=1 | 2.385 | 3 | 1 | 9 | 1 | 264 | 29.1 | 5.47 | 20.94 | 1 | 219.8 | 110.5 | 421.3 | 190.9 | 83.5 | 320.3 | 199.9 | 88.8 | 345.1 | 0.792 | 0.671673354 | 0.843267696 | 0.755555243 | 1.065747436 | 0.90355167 |
| A0A1U7RHV7 | Proteasome subunit beta OS=Mesocricetus auratus OX=10036 GN=Psmb5 PE=3 SV=1 | 5.007 | 7 | 1 | 6 | 1 | 216 | 23.3 | 8.25 | 31.18 | 1 | 109.1 | 98.5 | 67.5 | 105.4 | 93.7 | 62 | 72.8 | 94.6 | 39.9 | 0.965 | 0.808204933 | 0.753544166 | 0.32938479 | 0.793948679 | 0.433224171 |
| A0A1U7QCV2 | 26S protease regulatory subunit 6A OS=Mesocricetus auratus OX=10036 GN=Psmc3 PE=3 SV=1 | 31.047 | 21 | 7 | 37 | 7 | 442 | 49.5 | 5.19 | 110.21 | 7 | 176.5 | 152.5 | 172.5 | 176.6 | 144.9 | 160.3 | 106.3 | 123.3 | 104.4 | 0.999 | 0.60827076 | 0.666001994 | 0.004869106 | 0.693233707 | 0.015081303 |
| A0A1U7QV15 | 26S proteasome AAA-ATPase subunit RPT3 OS=Mesocricetus auratus OX=10036 GN=Psmc4 PE=3 SV=1 | 28.607 | 24 | 7 | 50 | 7 | 418 | 47.4 | 5.21 | 130.46 | 7 | 484.3 | 705.4 | 727.1 | 415.7 | 515.9 | 604.5 | 456.1 | 696.8 | 707 | 0.797 | 0.259284692 | 0.970315109 | 0.874725904 | 1.210793568 | 0.34301858 |
| A0A1U7QLF6 | 26S protease regulatory subunit 8 OS=Mesocricetus auratus OX=10036 GN=Psmc5 PE=3 SV=1 | 26.086 | 23 | 6 | 30 | 6 | 416 | 46.6 | 7.88 | 93.49 | 6 | 208 | 53.4 | 186.8 | 185.4 | 75 | 157.2 | 152.4 | 57.5 | 134 | 0.915 | 0.871394671 | 0.767291388 | 0.578085339 | 0.823515326 | 0.607309417 |
| A0A1U7R4M1 | 26S protease regulatory subunit 10B OS=Mesocricetus auratus OX=10036 GN=Psmc6 PE=3 SV=1 | 10.958 | 9 | 3 | 13 | 3 | 403 | 45.8 | 7.78 | 35.74 | 3 | 522.4 | 420.7 | 238.3 | 393.8 | 274.5 | 186.3 | 405.6 | 310.3 | 139 | 0.652 | 0.353509292 | 0.723632978 | 0.393911746 | 1.000351041 | 0.99924141 |
| A0A1U7QZM8 | 26S proteasome non-ATPase regulatory subunit 1 OS=Mesocricetus auratus OX=10036 GN=Psmd1 PE=3 SV=1 | 8.524 | 2 | 3 | 13 | 3 | 953 | 105.8 | 5.39 | 34.43 | 3 | 663.3 | 554.2 | 555.2 | 530.2 | 494.3 | 485.9 | 624.1 | 532.1 | 593.4 | 0.861 | 0.124246809 | 0.98696903 | 0.873595208 | 1.158368644 | 0.0794273 |
| A0A1U7QC95 | 26S proteasome non-ATPase regulatory subunit 3 OS=Mesocricetus auratus OX=10036 GN=Psmd3 PE=3 SV=1 | 12.275 | 7 | 3 | 5 | 3 | 530 | 60.7 | 8.44 | 18.01 | 3 | 56.2 | 94.6 | 181.4 | 48.8 | 71.9 | 157 | 33.7 | 61.8 | 174.9 | 0.841 | 0.732630556 | 0.813967489 | 0.73587934 | 0.97371264 | 0.966524744 |
| A0A1U7R8Z7 | 26S proteasome non-ATPase regulatory subunit 5 OS=Mesocricetus auratus OX=10036 GN=Psmd5 PE=4 SV=1 | 10.591 | 7 | 2 | 4 | 2 | 504 | 56 | 5.27 | 11.56 | 2 | 0 | 0 | 43.8 | 0 | 0 | 36.6 | 0 | 0 | 37.3 | 0.836 | 0.905888861 | 0.851598174 | 0.915617754 | 1.019125683 | 0.989954196 |
| A0A1U7QPE5 | 26S proteasome non-ATPase regulatory subunit 6 OS=Mesocricetus auratus OX=10036 GN=Psmd6 PE=3 SV=1 | 11.661 | 12 | 4 | 7 | 4 | 389 | 45.5 | 5.6 | 19.39 | 4 | 168.6 | 305.1 | 64.4 | 144.9 | 223.5 | 59.4 | 151 | 185.5 | 66 | 0.79 | 0.688021167 | 0.74800223 | 0.60420718 | 0.940860215 | 0.894146451 |
| A0A1U7REP6 | proteasome activator complex subunit 2 OS=Mesocricetus auratus OX=10036 GN=Psme2 PE=3 SV=1 | 23.756 | 21 | 4 | 31 | 4 | 239 | 27.1 | 5.76 | 87.57 | 4 | 791.1 | 845.2 | 339 | 917.4 | 1322.8 | 811.3 | 815.5 | 1192.9 | 722.7 | 2.235 | 0.184128923 | 1.382625424 | 0.307953206 | 0.895002458 | 0.641138135 |
| A0A1U7R8K7 | Polypyrimidine tract-binding protein 1 OS=Mesocricetus auratus OX=10036 GN=Ptbp1 PE=4 SV=1 | 19.673 | 12 | 5 | 14 | 5 | 555 | 59.3 | 9.16 | 49.27 | 5 | 454.6 | 288.6 | 366.4 | 418.7 | 290.4 | 327.5 | 472.3 | 272 | 332.5 | 1.006 | 0.712454318 | 0.970439798 | 0.893235904 | 1.038780629 | 0.860024928 |
| A0A1U7QS96 | prostaglandin E synthase 3 OS=Mesocricetus auratus OX=10036 GN=Ptges3 PE=4 SV=1 | 9.605 | 17 | 2 | 15 | 2 | 160 | 18.7 | 4.55 | 50.47 | 2 | 236.1 | 246.9 | 230.2 | 197.5 | 207.2 | 218 | 199.6 | 214 | 207.6 | 0.873 | 0.018314777 | 0.871003926 | 0.009371725 | 0.997591135 | 0.948607453 |
| A0A1U7QPK8 | prostaglandin F2 receptor negative regulator OS=Mesocricetus auratus OX=10036 GN=Ptgfrn PE=4 SV=1 | 2.125 | 1 | 1 | 1 | 1 | 879 | 98.6 | 6.61 | 2.95 | 1 | 138.6 | 0 | 0 | 102.7 | 0 | 0 | 118.2 | 0 | 0 | 0.741 | 0.846123601 | 0.852813853 | 0.916353056 | 1.150925024 | 0.925998093 |
| A0A1U8CDC7 | Cyclooxygenase-1 OS=Mesocricetus auratus OX=10036 GN=Ptgs1 PE=3 SV=1 | 15.514 | 7 | 4 | 20 | 4 | 601 | 68.6 | 6.84 | 68.57 | 4 | 65.1 | 433.9 | 457.8 | 79.7 | 439.5 | 449.6 | 44 | 433.7 | 436.5 | 1.013 | 0.982952467 | 0.955476589 | 0.941584139 | 0.943641618 | 0.923636865 |
| A0A1U8CAD3 | Tyrosine-protein phosphatase non-receptor type OS=Mesocricetus auratus OX=10036 GN=Ptpn11 PE=3 SV=1 | 4.874 | 3 | 2 | 3 | 2 | 597 | 68.4 | 7.3 | 6.69 | 2 | 0 | 0 | 215.1 | 0 | 0 | 163.5 | 0 | 0 | 170.9 | 0.76 | 0.858474162 | 0.794514179 | 0.880358936 | 1.045259939 | 0.976541773 |
| A0A1U7QN74 | tyrosine-protein phosphatase non-receptor type 9 OS=Mesocricetus auratus OX=10036 GN=Ptpn9 PE=4 SV=1 | 3.469 | 2 | 1 | 1 | 1 | 593 | 68 | 8.15 | 2.88 | 1 | 0 | 0 | 89.5 | 0 | 0 | 112.5 | 0 | 0 | 94.4 | 1.257 | 0.881014377 | 1.054748603 | 0.971762043 | 0.839111111 | 0.908026052 |
| A0A1U7QP67 | polymerase I and transcript release factor OS=Mesocricetus auratus OX=10036 GN=Cavin1 PE=3 SV=1 | 101.699 | 34 | 18 | 157 | 18 | 391 | 43.7 | 5.45 | 465.7 | 18 | 6013.3 | 6156.3 | 5649.8 | 4396.2 | 4242.3 | 4436.2 | 6983.1 | 7199.8 | 5523.6 | 0.689 | 0.004049661 | 1.105901433 | 0.354973304 | 1.507223875 | 0.05066131 |
| A0A1U8C2Y9 | Aminoacyl-tRNA hydrolase OS=Mesocricetus auratus OX=10036 GN=Ptrh2 PE=4 SV=1 | 3.826 | 21 | 2 | 2 | 2 | 174 | 18.8 | 7.01 | 4.63 | 2 | 0 | 48.4 | 0 | 0 | 37 | 0 | 0 | 35.4 | 0 | 0.764 | 0.861266039 | 0.731404959 | 0.83989713 | 0.956756757 | 0.976573262 |
| A0A1U8CZB0 | transcriptional activator protein Pur-beta OS=Mesocricetus auratus OX=10036 GN=Purb PE=3 SV=1 | 25.268 | 42 | 5 | 21 | 4 | 297 | 31.6 | 5.69 | 37.46 | 5 | 78.8 | 138.5 | 38.6 | 56.2 | 83.5 | 39.6 | 57.2 | 98.7 | 37.2 | 0.603 | 0.48455244 | 0.754591637 | 0.579612368 | 1.076965979 | 0.84687844 |
| A0A1U7QLN5 | nectin-3 isoform X1 OS=Mesocricetus auratus OX=10036 GN=Nectin3 PE=4 SV=1 | 4.05 | 5 | 1 | 1 | 1 | 551 | 60.9 | 6.1 | 3.03 | 1 | 0 | 36.1 | 0 | 0 | 48.3 | 0 | 0 | 32 | 0 | 1.338 | 0.85028875 | 0.886426593 | 0.936409379 | 0.66252588 | 0.79440803 |
| A0A1U7QTV2 | paxillin OS=Mesocricetus auratus OX=10036 GN=Pxn PE=4 SV=1 | 24.631 | 8 | 2 | 19 | 2 | 589 | 64.4 | 6 | 70.7 | 2 | 225.4 | 191.7 | 168.8 | 186.8 | 139.4 | 135 | 255.6 | 233.9 | 209.6 | 0.829 | 0.149646775 | 1.193207032 | 0.15191155 | 1.515828274 | 0.02200863 |
| A0A1U7R1L4 | ras-related protein Rab-10 OS=Mesocricetus auratus OX=10036 GN=Rab10 PE=4 SV=1 | 26.898 | 21 | 4 | 14 | 2 | 200 | 22.6 | 8.38 | 47.41 | 4 | 0 | 110.3 | 0 | 0 | 117.1 | 0 | 0 | 96.1 | 0 | 1.062 | 0.968315657 | 0.871260199 | 0.927426074 | 0.820666097 | 0.896685344 |
| A0A1U7QY95 | ras-related protein Rab-11A OS=Mesocricetus auratus OX=10036 GN=Rab11a PE=4 SV=1 | 20.978 | 23 | 5 | 23 | 5 | 216 | 24.4 | 6.57 | 49.78 | 5 | 287.4 | 355.4 | 391.4 | 285.7 | 359.3 | 386.5 | 227.7 | 299 | 311.9 | 0.988 | 0.984249746 | 0.810868304 | 0.181690004 | 0.81299079 | 0.183726933 |
| A0A1U7QZN3 | ras-related protein Rab-12 OS=Mesocricetus auratus OX=10036 GN=Rab12 PE=4 SV=1 | 10.879 | 8 | 2 | 7 | 1 | 243 | 27.2 | 8.41 | 16.85 | 2 | 43.2 | 106.8 | 0 | 39 | 70.5 | 0 | 35.7 | 85 | 0 | 0.772 | 0.737216958 | 0.804666667 | 0.817985512 | 1.102283105 | 0.912890739 |
| A0A1U7Q629 | ras-related protein Rab-1A OS=Mesocricetus auratus OX=10036 GN=Rab1a PE=4 SV=1 | 24.079 | 19 | 4 | 15 | 1 | 205 | 22.6 | 5.72 | 44.08 | 4 | 82.1 | 78.3 | 68.8 | 95.4 | 70.6 | 71.3 | 76.6 | 60.8 | 57.4 | 1.036 | 0.78584169 | 0.84991274 | 0.192626336 | 0.820901812 | 0.238771912 |
| A0A1U7Q7J7 | ras-related protein Rab-2A OS=Mesocricetus auratus OX=10036 GN=Rab2a PE=4 SV=1 | 14.52 | 10 | 2 | 15 | 2 | 212 | 23.5 | 6.54 | 49.94 | 2 | 172.2 | 153.9 | 140.4 | 184.3 | 154.4 | 132.1 | 162.7 | 180.9 | 163.6 | 1.07 | 0.94014488 | 1.087245445 | 0.293906041 | 1.077315208 | 0.516761207 |
| A0A1U7R1E1 | ras-related protein Rab-35 isoform X1 OS=Mesocricetus auratus OX=10036 GN=Rab35 PE=4 SV=1 | 26.655 | 23 | 4 | 21 | 2 | 201 | 23 | 8.29 | 72.88 | 4 | 70.5 | 198.2 | 64.7 | 86 | 177.9 | 72.2 | 58.4 | 179.4 | 63 | 1.041 | 0.98772647 | 0.902219556 | 0.862602549 | 0.894971735 | 0.831286843 |
| A0A1U7R754 | ras-related protein Rab-5C OS=Mesocricetus auratus OX=10036 GN=Rab5c PE=4 SV=1 | 7.697 | 15 | 3 | 5 | 3 | 216 | 23.4 | 8.41 | 13.41 | 3 | 0 | 38.6 | 132.3 | 0 | 33.2 | 118.2 | 0 | 26.2 | 129.5 | 0.877 | 0.907928229 | 0.911059099 | 0.931928717 | 1.028401585 | 0.979709242 |
| A0A1U7REJ8 | ras-related protein Rab-8A OS=Mesocricetus auratus OX=10036 GN=Rab8a PE=4 SV=1 | 22.669 | 14 | 4 | 13 | 1 | 207 | 23.7 | 9.07 | 37.57 | 4 | 93.6 | 0 | 0 | 70.1 | 0 | 0 | 108.5 | 0 | 0 | 0.75 | 0.85127104 | 1.159188034 | 0.922291011 | 1.547788873 | 0.783411296 |
| A0A1U7QD35 | UV excision repair protein RAD23 OS=Mesocricetus auratus OX=10036 GN=Rad23b PE=3 SV=1 | 2.69 | 3 | 1 | 1 | 1 | 415 | 43.2 | 4.78 | 0 | 1 | 0 | 33.6 | 0 | 0 | 28.8 | 0 | 0 | 34.7 | 0 | 0.856 | 0.918965923 | 1.032738095 | 0.982922762 | 1.204861111 | 0.902425056 |
| A0A1U7QGM1 | Small monomeric GTPase OS=Mesocricetus auratus OX=10036 GN=Rala PE=3 SV=1 | 24.366 | 26 | 5 | 26 | 1 | 206 | 23.5 | 7.11 | 50.23 | 5 | 230.7 | 95.9 | 71.3 | 206.1 | 78.3 | 95.8 | 242.3 | 85.5 | 98.7 | 1.231 | 0.930822281 | 1.071877356 | 0.89902401 | 1.121778012 | 0.822398276 |
| A0A1U7QNG0 | ran-specific GTPase-activating protein OS=Mesocricetus auratus OX=10036 GN=Ranbp1 PE=4 SV=1 | 5.666 | 12 | 2 | 7 | 2 | 203 | 23.5 | 5.22 | 15.18 | 2 | 158.5 | 249.1 | 77.6 | 131.4 | 175.3 | 51.8 | 147 | 215.3 | 60.3 | 0.717 | 0.532148113 | 0.870981039 | 0.770578816 | 1.178800558 | 0.730296054 |
| A0A1U7R5R8 | ras-related protein Rap-1b OS=Mesocricetus auratus OX=10036 GN=Rap1b PE=3 SV=1 | 51.549 | 42 | 9 | 65 | 3 | 184 | 20.8 | 5.78 | 179.64 | 9 | 695.4 | 691.8 | 544.1 | 634 | 598.5 | 503.2 | 690 | 693.1 | 586.6 | 0.925 | 0.364427966 | 1.01988298 | 0.844923601 | 1.134815924 | 0.211924517 |
| A0A1U7QHV2 | histone-binding protein RBBP7 OS=Mesocricetus auratus OX=10036 GN=Rbbp7 PE=4 SV=1 | 12.501 | 14 | 5 | 22 | 2 | 425 | 47.8 | 5.05 | 55.61 | 5 | 200.1 | 145.2 | 175.2 | 177.9 | 154.9 | 171.8 | 189.7 | 139 | 179.3 | 0.934 | 0.781191909 | 0.97598463 | 0.859996334 | 1.00673801 | 0.951152307 |
| A0A1U8CKN4 | RNA-binding protein 3 isoform X2 OS=Mesocricetus auratus OX=10036 GN=Rbm3 PE=4 SV=1 | 5.008 | 17 | 2 | 5 | 2 | 155 | 16.8 | 7.58 | 10.19 | 2 | 61.4 | 76.2 | 112.5 | 72.3 | 73.8 | 106.8 | 79.7 | 81.9 | 115.2 | 0.968 | 0.9631697 | 1.106757297 | 0.6661329 | 1.094503756 | 0.646327161 |
| A0A1U7QR19 | RNA-binding protein 8A OS=Mesocricetus auratus OX=10036 GN=Rbm8a PE=3 SV=1 | 9.616 | 16 | 2 | 11 | 2 | 174 | 19.9 | 5.72 | 48.32 | 2 | 41.6 | 0 | 0 | 33.6 | 0 | 0 | 37.5 | 0 | 0 | 0.808 | 0.888621127 | 0.901442308 | 0.945193305 | 1.116071429 | 0.942022458 |
| A0A1U7QF74 | reticulocalbin-1 OS=Mesocricetus auratus OX=10036 GN=Rcn1 PE=4 SV=1 | 17.292 | 26 | 5 | 16 | 5 | 325 | 38.1 | 4.82 | 55.88 | 5 | 257.2 | 219.6 | 268 | 219.2 | 213.2 | 241.3 | 233 | 214.5 | 280.6 | 0.971 | 0.251258346 | 0.977577873 | 0.832609014 | 1.080748107 | 0.465813874 |
| A0A1U7QAI8 | reticulocalbin-2 OS=Mesocricetus auratus OX=10036 GN=Rcn2 PE=4 SV=1 | 23.411 | 23 | 5 | 13 | 5 | 352 | 40.8 | 4.56 | 47.91 | 5 | 67.9 | 111.8 | 0 | 78.7 | 111.2 | 0 | 56.5 | 80 | 0 | 1.074 | 0.9450337 | 0.759599332 | 0.740270572 | 0.718799368 | 0.68631486 |
| A0A1U7QG33 | Receptor expression-enhancing protein OS=Mesocricetus auratus OX=10036 GN=Reep5 PE=3 SV=1 | 12.122 | 21 | 5 | 12 | 5 | 189 | 21.5 | 8.12 | 8.96 | 5 | 125.4 | 207.4 | 167.5 | 120.9 | 150.5 | 132.5 | 117.1 | 210.9 | 169.8 | 0.749 | 0.307005531 | 0.995002998 | 0.982669738 | 1.232483288 | 0.369727543 |
| A0A1U7QBD1 | transcription factor p65 isoform X1 OS=Mesocricetus auratus OX=10036 GN=Rela PE=4 SV=1 | 1.654 | 2 | 1 | 2 | 1 | 549 | 60.2 | 5.87 | 5.01 | 1 | 0 | 40.3 | 35.6 | 0 | 30.5 | 24.2 | 0 | 30.7 | 38.8 | 0.718 | 0.679051881 | 0.915678524 | 0.908173955 | 1.270566728 | 0.760145098 |
| A0A1U7R2G4 | GTP-binding protein OS=Mesocricetus auratus OX=10036 GN=Rem1 PE=3 SV=1 | 1.506 | 4 | 1 | 1 | 1 | 297 | 33.2 | 9.26 | 0 | 1 | 85.3 | 0 | 0 | 115.4 | 0 | 0 | 67 | 0 | 0 | 1.353 | 0.844946879 | 0.785463072 | 0.874644279 | 0.580589255 | 0.739396135 |
| A0A1U8C7T7 | Protein RER1 OS=Mesocricetus auratus OX=10036 GN=Rer1 PE=3 SV=1 | 1.865 | 4 | 1 | 1 | 1 | 196 | 23 | 9.51 | 3.37 | 1 | 143.4 | 0 | 0 | 118.8 | 0 | 0 | 126.9 | 0 | 0 | 0.829 | 0.901490461 | 0.884937238 | 0.935532121 | 1.068181818 | 0.965077183 |
| A0A1U7QG12 | regulator of G-protein signaling protein-like OS=Mesocricetus auratus OX=10036 GN=Rgsl1 PE=4 SV=1 | 1.392 | 1 | 1 | 1 | 1 | 1112 | 129.8 | 9.17 | 0 | 1 | 0 | 0 | 60.5 | 0 | 0 | 29.9 | 0 | 0 | 76.8 | 0.493 | 0.68178644 | 1.269421488 | 0.876097782 | 2.568561873 | 0.614818321 |
| A0A1U7QPN2 | rho-related GTP-binding protein RhoC OS=Mesocricetus auratus OX=10036 GN=Rhoc PE=4 SV=1 | 9.269 | 17 | 3 | 10 | 1 | 193 | 22 | 6.58 | 15.88 | 3 | 177.9 | 298.7 | 150.3 | 101.6 | 182.8 | 84 | 143.9 | 242.9 | 119.3 | 0.571 | 0.20117037 | 0.80730579 | 0.534854873 | 1.373778502 | 0.399830286 |
| A0A1U7QVP9 | rho-related GTP-binding protein RhoG OS=Mesocricetus auratus OX=10036 GN=Rhog PE=4 SV=1 | 2.661 | 5 | 1 | 2 | 1 | 191 | 21.3 | 8.12 | 0 | 1 | 0 | 0 | 229 | 0 | 0 | 184.2 | 0 | 0 | 233.3 | 0.804 | 0.886542805 | 1.018777293 | 0.990135451 | 1.266558089 | 0.877223163 |
| A0A1U8D0Z2 | E3 ubiquitin-protein ligase RING1 OS=Mesocricetus auratus OX=10036 GN=Ring1 PE=4 SV=1 | 1.932 | 4 | 1 | 4 | 1 | 362 | 37.4 | 6.13 | 2.49 | 1 | 99.1 | 84.4 | 0 | 118.1 | 94.5 | 0 | 97.7 | 84.4 | 0 | 1.155 | 0.848352543 | 0.992370572 | 0.99194779 | 0.8565381 | 0.840612795 |
| A0A1U8C3T2 | Ribonuclease inhibitor OS=Mesocricetus auratus OX=10036 GN=Rnh1 PE=4 SV=1 | 13.415 | 10 | 4 | 9 | 4 | 456 | 49.9 | 4.81 | 24.73 | 4 | 138.7 | 164.8 | 133.6 | 124.9 | 142.4 | 136 | 110.6 | 163 | 145.2 | 0.9 | 0.377665234 | 0.95813315 | 0.756897048 | 1.038432928 | 0.775245357 |
| A0A1U7QRZ7 | aminopeptidase B OS=Mesocricetus auratus OX=10036 GN=Rnpep PE=3 SV=1 | 28.724 | 12 | 7 | 34 | 7 | 650 | 73 | 5.6 | 97.65 | 7 | 186.6 | 410.6 | 168.1 | 163.3 | 361.4 | 155.4 | 194.5 | 431 | 161.2 | 0.875 | 0.796739332 | 1.02796289 | 0.953643878 | 1.156741656 | 0.760343649 |
| A0A1U8CYJ6 | Protein XRP2 OS=Mesocricetus auratus OX=10036 GN=Rp2 PE=3 SV=1 | 3.687 | 3 | 1 | 3 | 1 | 350 | 39.7 | 5.33 | 7.95 | 1 | 93.2 | 177.2 | 0 | 82 | 157.4 | 0 | 97.2 | 194.1 | 0 | 0.884 | 0.887402445 | 1.077292899 | 0.931300991 | 1.21679198 | 0.822759389 |
| A0A1U7RAY9 | Ribosomal protein OS=Mesocricetus auratus OX=10036 GN=Rpl10a PE=3 SV=1 | 25.433 | 30 | 8 | 27 | 8 | 217 | 24.8 | 9.94 | 63.75 | 8 | 522.6 | 540.1 | 381.5 | 616 | 574 | 471.3 | 459.7 | 502.4 | 380.9 | 1.179 | 0.336381406 | 0.929926603 | 0.615733103 | 0.808403058 | 0.132497679 |
| A0A1U7QWV4 | 60S ribosomal protein L12 OS=Mesocricetus auratus OX=10036 GN=Rpl12 PE=3 SV=1 | 15.212 | 25 | 3 | 22 | 3 | 165 | 17.8 | 9.42 | 92.83 | 3 | 532.8 | 234.6 | 261.7 | 472 | 208.8 | 226.4 | 452.9 | 219 | 249.1 | 0.865 | 0.766271394 | 0.894956758 | 0.780284503 | 1.01521164 | 0.969331633 |
| A0A1U7RIR0 | 60S ribosomal protein L13a OS=Mesocricetus auratus OX=10036 GN=Rpl13a PE=3 SV=1 | 10.777 | 14 | 4 | 24 | 4 | 203 | 23.4 | 11.11 | 57.99 | 4 | 1161.3 | 1194.9 | 1330.9 | 1033.2 | 1121.1 | 1228.4 | 1082.6 | 1164.6 | 1223.7 | 0.93 | 0.256592688 | 0.941363131 | 0.33958325 | 1.026073846 | 0.696864887 |
| A0A1U7R751 | 60S ribosomal protein L14 OS=Mesocricetus auratus OX=10036 GN=Rpl14 PE=3 SV=1 | 11.947 | 10 | 2 | 28 | 2 | 208 | 22.6 | 11 | 79.85 | 2 | 440.3 | 279 | 969.1 | 373.5 | 246.7 | 734 | 342.4 | 209 | 793.7 | 0.848 | 0.68661375 | 0.796671405 | 0.697550013 | 0.993280165 | 0.9901028 |
| A0A1U7QD10 | 60S ribosomal protein L17 OS=Mesocricetus auratus OX=10036 GN=LOC101829440 PE=3 SV=1 | 20.084 | 10 | 2 | 21 | 2 | 184 | 21.4 | 10.18 | 72.53 | 2 | 28.9 | 0 | 0 | 43.8 | 0 | 0 | 23.1 | 0 | 0 | 1.519 | 0.792599291 | 0.799307958 | 0.883372556 | 0.52739726 | 0.703764299 |
| A0A1U7RD92 | 60S ribosomal protein L18 OS=Mesocricetus auratus OX=10036 GN=Rpl18 PE=3 SV=1 | 8.147 | 12 | 2 | 5 | 2 | 188 | 21.6 | 11.78 | 15.17 | 2 | 42 | 74.7 | 56.7 | 38.7 | 94.6 | 74.6 | 36 | 58.3 | 37.8 | 0.95 | 0.58314454 | 0.761822376 | 0.314554936 | 0.635401635 | 0.259974109 |
| A0A1U7R0M5 | 60S ribosomal protein L21 OS=Mesocricetus auratus OX=10036 GN=Rpl21 PE=3 SV=1 | 2.33 | 4 | 1 | 5 | 1 | 160 | 18.6 | 10.49 | 11.05 | 1 | 178.9 | 113.1 | 198 | 198.4 | 109 | 191.6 | 200.1 | 121.6 | 183.1 | 1.075 | 0.941766364 | 1.030204082 | 0.894960696 | 1.011623246 | 0.961267871 |
| A0A1U7QW59 | 60S ribosomal protein L23 OS=Mesocricetus auratus OX=10036 GN=Rpl23 PE=3 SV=1 | 4.453 | 11 | 2 | 3 | 2 | 140 | 14.9 | 10.51 | 6.13 | 2 | 72.2 | 62.6 | 66.5 | 55.9 | 57.6 | 62.8 | 90.4 | 43.1 | 45 | 0.944 | 0.079817973 | 0.886736215 | 0.673731225 | 1.012478729 | 0.966642726 |
| A0A1U7QE09 | 60S ribosomal protein L23a OS=Mesocricetus auratus OX=10036 GN=Rpl23a PE=3 SV=1 | 20.311 | 35 | 5 | 25 | 5 | 156 | 17.7 | 10.45 | 60.6 | 5 | 782.2 | 554.1 | 770.5 | 828.2 | 587.3 | 839.8 | 760.3 | 569 | 774.8 | 1.088 | 0.678371448 | 0.998718436 | 0.99322036 | 0.932957921 | 0.659416661 |
| A0A1U7QUA9 | 60S ribosomal protein L24 OS=Mesocricetus auratus OX=10036 GN=Rpl24 PE=3 SV=1 | 5.007 | 11 | 2 | 6 | 2 | 157 | 17.8 | 11.25 | 15.42 | 2 | 0 | 0 | 30.7 | 0 | 0 | 40.9 | 0 | 0 | 39.6 | 1.33 | 0.852355877 | 1.28990228 | 0.868159943 | 0.968215159 | 0.982876505 |
| A0A1U7QHJ7 | 60S ribosomal protein L26 OS=Mesocricetus auratus OX=10036 GN=Rpl26 PE=3 SV=1 | 15.569 | 23 | 5 | 25 | 5 | 145 | 17.3 | 10.48 | 45.36 | 5 | 773.5 | 654.2 | 696.6 | 789.9 | 619.7 | 668.7 | 724.6 | 659.5 | 702.4 | 1.021 | 0.816745848 | 0.982205903 | 0.771700755 | 1.003945532 | 0.963363928 |
| A0A1U7QDR3 | 60S ribosomal protein L27 OS=Mesocricetus auratus OX=10036 GN=Rpl27 PE=3 SV=1 | 11.667 | 22 | 4 | 22 | 4 | 136 | 15.7 | 10.49 | 46.45 | 4 | 913.2 | 1016.1 | 715.9 | 956.2 | 1094.6 | 815 | 815.1 | 923.7 | 740.6 | 1.047 | 0.57175143 | 0.937320429 | 0.625384622 | 0.865168539 | 0.263610017 |

Supplementary Table 1: List of differentially expressed proteins identified across the 4dpi infected and 14dpi infected

| Accession | Description | Sum PEP Score | Coverage [%] | # Peptides | # PSMs | # Unique Peptides | # AAs | MW [kDa] | calc. pI | Score Sequest HT: Sequest HT | # Peptides (by Search Engine): Sequest HT | Mock-infected (4dpi) Replicate_R1 | Mock-infected (4dpi) Replicate_R2 | Mock-infected (4dpi) Replicate_R3 | SARS-CoV-2-infected (4dpi) Replicate_R1 | SARS-CoV-2-infected (4dpi) Replicate_R2 | SARS-CoV-2-infected (4dpi) Replicate_R3 | SARS-CoV-2-(14dpi) Replicate_R1 | SARS-CoV-2-infected (14dpi) Replicate_R2 | SARS-CoV-2-infected (14dpi) Replicate_R3 | Abundance Ratio: SARS-CoV-2-infected (4dpi) / Mock-infected (4dpi) | P-Value: SARS-CoV-2-infected (4dpi) / Mock-infected (4dpi) | Abundance Ratio: SARS-CoV-2-infected (14dpi) / Mock-infected (4dpi) | P-Value: SARS-CoV-2-infected (14dpi) / Mock-infected (4dpi) | Abundance Ratio: SARS-CoV-2-infected (14dpi) / SARS-CoV-2-infected (4dpi) | P-Value: SARS-CoV-2-infected (14dpi) / SARS-CoV-2-infected (4dpi) |
|---|---|---|---|---|---|---|---|---|---|---|---|---|---|---|---|---|---|---|---|---|---|---|---|---|---|---|
| A0A1U7R985 | 60S ribosomal protein L28 OS=Mesocricetus auratus OX=10036 GN=LOC101823529 PE=3 SV=1 | 6.279 | 15 | 2 | 4 | 2 | 137 | 15.7 | 12.02 | 9.54 | 2 | 449.6 | 0 | 299.8 | 377.1 | 0 | 290.4 | 318.8 | 0 | 239 | 0.901 | 0.883457011 | 0.744328796 | 0.717369566 | 0.835655431 | 0.81845702 |
| A0A1U8BTJ2 | 60S ribosomal protein L3 OS=Mesocricetus auratus OX=10036 GN=Rpl3 PE=3 SV=1 | 27.848 | 16 | 6 | 24 | 6 | 403 | 46.1 | 10.21 | 52.37 | 6 | 636.9 | 454.1 | 583 | 576.7 | 376.3 | 474.9 | 540.4 | 377.1 | 509.7 | 0.829 | 0.359561799 | 0.852568698 | 0.327984915 | 0.99950977 | 0.997716303 |
| A0A1U7RA38 | 60S ribosomal protein L34 OS=Mesocricetus auratus OX=10036 GN=Rpl34 PE=3 SV=1 | 3.103 | 7 | 1 | 3 | 1 | 117 | 13.3 | 11.47 | 10.21 | 1 | 100.9 | 69.5 | 90.6 | 97.3 | 80.8 | 108.6 | 96.4 | 68.4 | 92.1 | 1.133 | 0.524183406 | 0.984291188 | 0.919478431 | 0.896058598 | 0.450081159 |
| A0A1U7QXT6 | 60S ribosomal protein L36 OS=Mesocricetus auratus OX=10036 GN=Rpl36 PE=3 SV=1 | 2.802 | 10 | 1 | 3 | 1 | 105 | 12.2 | 11.59 | 10.2 | 1 | 40.9 | 58.7 | 52.2 | 40.3 | 54.8 | 56.5 | 33.1 | 46 | 39.3 | 0.989 | 0.993162043 | 0.77997365 | 0.164125251 | 0.781002639 | 0.163129111 |
| A0A1U7QA51 | 60S ribosomal protein L9 OS=Mesocricetus auratus OX=10036 GN=Rpl9 PE=3 SV=1 | 20.791 | 28 | 5 | 22 | 5 | 192 | 21.8 | 9.95 | 53.22 | 5 | 566 | 453.2 | 338.9 | 558.4 | 477 | 365.6 | 551.6 | 436.3 | 319.9 | 1.053 | 0.876398947 | 0.962962963 | 0.866626331 | 0.933476089 | 0.740055578 |
| A0A1U7QK56 | 60S acidic ribosomal protein P1 OS=Mesocricetus auratus OX=10036 GN=Rplp1 PE=3 SV=1 | 13.837 | 14 | 1 | 30 | 1 | 114 | 11.5 | 4.32 | 128.99 | 1 | 121.8 | 62.1 | 35.4 | 171 | 80.3 | 35.4 | 114.4 | 67.5 | 34.5 | 1.404 | 0.664016953 | 0.986776106 | 0.97899599 | 0.754795954 | 0.64429729 |
| A0A1U8CQN0 | 60S acidic ribosomal protein P2 isoform X2 OS=Mesocricetus auratus OX=10036 GN=Rplp2 PE=3 SV=1 | 26.143 | 32 | 3 | 35 | 3 | 114 | 11.6 | 4.54 | 137.07 | 3 | 233 | 335.4 | 304.5 | 350.9 | 522 | 449.4 | 244.2 | 331.2 | 387.1 | 1.486 | 0.07417398 | 1.102646351 | 0.595510308 | 0.727898359 | 0.139596272 |
| A0A1U7QR04 | Dolichyl-diphosphooligosaccharide--protein glycosyltransferase subunit 1 OS=Mesocricetus auratus OX=10036 GN=Rpn1 PE=3 SV=1 | 23.797 | 20 | 7 | 37 | 7 | 606 | 68.3 | 6.47 | 89.57 | 7 | 468.3 | 690.2 | 338.5 | 523.6 | 703.2 | 430.4 | 413.4 | 713.8 | 335.2 | 1.118 | 0.703905147 | 0.976887108 | 0.944111501 | 0.882452329 | 0.670643429 |
| A0A1U7QS58 | 40S ribosomal protein S10 OS=Mesocricetus auratus OX=10036 GN=Rps10 PE=3 SV=1 | 15.523 | 24 | 3 | 12 | 3 | 165 | 18.9 | 10.15 | 37.32 | 3 | 104.6 | 0 | 0 | 93.8 | 0 | 0 | 100.3 | 0 | 0 | 0.897 | 0.94246002 | 0.958891013 | 0.977752509 | 1.069296375 | 0.964526844 |
| A0A1U7R979 | 40S ribosomal protein S11 OS=Mesocricetus auratus OX=10036 GN=Rps11 PE=3 SV=1 | 10.545 | 34 | 5 | 11 | 5 | 158 | 18.4 | 10.3 | 15.63 | 5 | 289.6 | 181.7 | 241.6 | 289.4 | 187.7 | 250.7 | 252.5 | 153.5 | 236.8 | 0.999 | 0.913706089 | 0.901669238 | 0.621915481 | 0.883209673 | 0.543179584 |
| A0A1U7QGY7 | 40S ribosomal protein S12 OS=Mesocricetus auratus OX=10036 GN=Rps12 PE=3 SV=1 | 9.614 | 21 | 3 | 18 | 3 | 132 | 14.5 | 7.21 | 53.92 | 3 | 618.5 | 476.9 | 522.1 | 742 | 542.3 | 643.6 | 538.1 | 399.5 | 514 | 1.137 | 0.226373359 | 0.897434312 | 0.40719239 | 0.752943617 | 0.097117098 |
| A0A1U7QYC6 | 40S ribosomal protein S13 OS=Mesocricetus auratus OX=10036 GN=Rps13 PE=3 SV=1 | 27.42 | 32 | 6 | 35 | 6 | 151 | 17.2 | 10.54 | 105.43 | 6 | 394.7 | 165.3 | 290.6 | 450.5 | 146 | 296.8 | 442.5 | 179.9 | 274.2 | 0.994 | 0.903835228 | 1.054079473 | 0.88735156 | 1.003694168 | 0.992940184 |
| A0A1U7Q287 | 40S ribosomal protein S15 OS=Mesocricetus auratus OX=10036 GN=Rps15 PE=3 SV=1 | 2.809 | 8 | 1 | 3 | 1 | 145 | 17 | 10.39 | 4.45 | 1 | 0 | 25.6 | 34.8 | 0 | 43.4 | 31.5 | 0 | 25.9 | 38.8 | 1.239 | 0.786205709 | 1.071192053 | 0.93055676 | 0.863818425 | 0.85355617 |
| A0A1U7QBT3 | 40S ribosomal protein S15a OS=Mesocricetus auratus OX=10036 GN=Rps15a PE=3 SV=1 | 10.142 | 23 | 3 | 11 | 3 | 130 | 14.8 | 10.13 | 26.09 | 3 | 189.6 | 192.9 | 318.1 | 216.2 | 202.3 | 320.2 | 173.2 | 163.7 | 262.8 | 1.006 | 0.832834576 | 0.855980588 | 0.561224509 | 0.811831596 | 0.397391227 |
| A0A1U7QUG8 | 40S ribosomal protein S16 OS=Mesocricetus auratus OX=10036 GN=Rps16 PE=3 SV=1 | 13.253 | 8 | 2 | 24 | 2 | 146 | 16.4 | 10.21 | 57.44 | 2 | 164.3 | 79.6 | 68.1 | 168.5 | 103 | 82.4 | 147.9 | 74.2 | 62.1 | 1.209 | 0.744498702 | 0.910897436 | 0.830334545 | 0.80305171 | 0.567331347 |
| A0A1U8CSH8 | 40S ribosomal protein S19 OS=Mesocricetus auratus OX=10036 GN=Rps19 PE=3 SV=1 | 30.843 | 53 | 9 | 40 | 9 | 145 | 16.1 | 10.32 | 92.24 | 9 | 732.3 | 749.1 | 578.4 | 744.4 | 704.8 | 560.8 | 706.6 | 684.2 | 508.8 | 0.97 | 0.841600936 | 0.922225459 | 0.554889463 | 0.945074627 | 0.683518793 |
| A0A1U7QCJ6 | 40S ribosomal protein S25 OS=Mesocricetus auratus OX=10036 GN=Rps25 PE=3 SV=1 | 15.503 | 30 | 4 | 19 | 1 | 125 | 13.8 | 10.17 | 59.08 | 4 | 192 | 166.9 | 109.3 | 186 | 180.9 | 111.3 | 222.3 | 163.9 | 131.2 | 1.022 | 0.927361155 | 1.105083298 | 0.674028149 | 1.081974069 | 0.734642708 |
| A0A1U8C9B9 | 40S ribosomal protein S27 OS=Mesocricetus auratus OX=10036 GN=Rps27 PE=3 SV=1 | 9.564 | 31 | 3 | 11 | 3 | 84 | 9.5 | 9.45 | 29.63 | 3 | 363.8 | 208.7 | 367.1 | 303.4 | 169.4 | 343.6 | 303.1 | 180.3 | 293.4 | 0.841 | 0.609382572 | 0.826734781 | 0.456960256 | 0.951494366 | 0.851481284 |
| A0A1U7R7Y3 | 40S ribosomal protein S28 OS=Mesocricetus auratus OX=10036 GN=Rps28 PE=3 SV=1 | 3.389 | 13 | 1 | 3 | 1 | 69 | 7.8 | 10.7 | 9.87 | 1 | 68.1 | 89.2 | 64.8 | 76.5 | 111.4 | 79.1 | 68.2 | 66.2 | 66.6 | 1.22 | 0.339893203 | 0.904997749 | 0.454763539 | 0.752808989 | 0.188733478 |
| A0A1U7Q9Q4 | 40S ribosomal protein S3 OS=Mesocricetus auratus OX=10036 GN=Rps3 PE=3 SV=1 | 38.977 | 40 | 8 | 38 | 8 | 243 | 26.7 | 9.66 | 121.52 | 8 | 409.8 | 297 | 364.5 | 527.3 | 363.9 | 455.7 | 409.4 | 268.2 | 314.3 | 1.225 | 0.194218932 | 0.925884439 | 0.644692746 | 0.736431806 | 0.134548318 |
| A0A1U7R570 | 40S ribosomal protein S4 OS=Mesocricetus auratus OX=10036 GN=Rps4x PE=3 SV=1 | 40.525 | 32 | 8 | 54 | 8 | 263 | 29.6 | 10.15 | 130.14 | 8 | 1953.6 | 2180.3 | 1922.7 | 1671 | 1933.6 | 1609.6 | 1891.7 | 2094.8 | 1889.3 | 0.837 | 0.096575129 | 0.970148268 | 0.600879741 | 1.126884278 | 0.1503574 |
| A0A1U8BX98 | Non-specific serine/threonine protein kinase OS=Mesocricetus auratus OX=10036 GN=Rps6ka3 PE=3 SV=1 | 13.428 | 9 | 3 | 9 | 3 | 718 | 81.3 | 7.15 | 21.34 | 3 | 0 | 110.7 | 0 | 0 | 96.6 | 0 | 0 | 127.8 | 0 | 0.873 | 0.928241199 | 1.154471545 | 0.924402711 | 1.322981366 | 0.855745397 |
| A0A1U7QH16 | 40S ribosomal protein S8 OS=Mesocricetus auratus OX=10036 GN=Rps8 PE=3 SV=1 | 13.24 | 22 | 4 | 13 | 4 | 208 | 24.2 | 10.32 | 31.5 | 4 | 202.1 | 207.3 | 224.6 | 206.1 | 204 | 231.9 | 191.7 | 202.7 | 217.9 | 0.984 | 0.825185945 | 0.965772871 | 0.517647872 | 0.953738318 | 0.448265614 |
| A0A1U7RAP6 | ras-related protein R-Ras OS=Mesocricetus auratus OX=10036 GN=Rras PE=4 SV=1 | 37.135 | 43 | 8 | 35 | 6 | 218 | 23.8 | 7.34 | 98.39 | 8 | 223.6 | 186.5 | 203.6 | 195.1 | 162.1 | 201.4 | 216.5 | 180.8 | 220.5 | 0.989 | 0.322003847 | 1.006680789 | 0.938284925 | 1.105979234 | 0.323657763 |
| A0A1U7QN56 | ras-related protein R-Ras2 OS=Mesocricetus auratus OX=10036 GN=Rras2 PE=4 SV=1 | 16.356 | 19 | 3 | 12 | 1 | 204 | 23.3 | 6.01 | 42.89 | 3 | 52.3 | 34.4 | 89.2 | 66 | 42.3 | 98.4 | 67.5 | 46.7 | 100 | 1.23 | 0.677200207 | 1.217737351 | 0.598919396 | 1.03628447 | 0.916782276 |
| A0A1U7QHJ6 | Ribonucleoside-diphosphate reductase OS=Mesocricetus auratus OX=10036 GN=Rrm1 PE=3 SV=1 | 2.055 | 1 | 1 | 1 | 1 | 792 | 90.2 | 6.9 | 0 | 1 | 47.3 | 0 | 0 | 31.9 | 0 | 0 | 35.6 | 0 | 0 | 0.674 | 0.802335556 | 0.752642706 | 0.853666711 | 1.115987461 | 0.942061826 |
| A0A1U7QXM5 | ras suppressor protein 1 isoform X1 OS=Mesocricetus auratus OX=10036 GN=Rsu1 PE=4 SV=1 | 39.329 | 36 | 8 | 47 | 8 | 277 | 31.5 | 8.63 | 134.96 | 8 | 1152.1 | 758.8 | 667.2 | 966.5 | 849.6 | 561.1 | 1069.7 | 1126.3 | 570.3 | 0.858 | 0.744803613 | 1.072999496 | 0.799704526 | 1.16367996 | 0.58096346 |
| A0A1U8BPZ6 | rhotekin-2 isoform X1 OS=Mesocricetus auratus OX=10036 GN=Rtkn2 PE=4 SV=1 | 4.138 | 4 | 2 | 4 | 2 | 600 | 67.3 | 8.06 | 7.37 | 2 | 28.9 | 32.3 | 108.7 | 33 | 61.5 | 126.1 | 26.2 | 22.4 | 97.4 | 1.333 | 0.678883089 | 0.859329017 | 0.834337581 | 0.661831369 | 0.536661834 |
| A0A1U7QE52 | Reticulon OS=Mesocricetus auratus OX=10036 GN=Rtn4 PE=4 SV=1 | 36.36 | 38 | 8 | 25 | 8 | 362 | 38.9 | 4.69 | 81.45 | 8 | 601.7 | 198.7 | 207 | 481.1 | 191.1 | 154.6 | 531.6 | 172.6 | 172.1 | 0.762 | 0.739803695 | 0.869863014 | 0.81920864 | 1.059869376 | 0.92202701 |
| A0A1U7Q6V2 | RuvB-like helicase OS=Mesocricetus auratus OX=10036 GN=Ruvbl1 PE=3 SV=1 | 8.938 | 11 | 4 | 5 | 4 | 456 | 50.2 | 6.42 | 7.31 | 4 | 149.2 | 0 | 47.5 | 129.3 | 0 | 47.2 | 146.5 | 0 | 57.2 | 0.928 | 0.913292753 | 1.035587189 | 0.971447467 | 1.154107649 | 0.881345328 |
| A0A1U7RDA2 | RuvB-like helicase OS=Mesocricetus auratus OX=10036 GN=Ruvbl2 PE=3 SV=1 | 3.708 | 2 | 1 | 2 | 1 | 463 | 51.1 | 5.64 | 4.29 | 1 | 56 | 0 | 44.4 | 58.4 | 0 | 36.5 | 54.2 | 0 | 35.4 | 0.927 | 0.943041268 | 0.892430279 | 0.884811547 | 0.944151739 | 0.94320221 |
| A0A1U7QW51 | Calpactin I light chain OS=Mesocricetus auratus OX=10036 GN=S100a10 PE=4 SV=1 | 40.54 | 35 | 4 | 82 | 4 | 97 | 11.3 | 6.77 | 164.38 | 4 | 1015.8 | 1069.1 | 1157.7 | 1663.9 | 1895.6 | 1836.9 | 1575.5 | 1899.2 | 1874.1 | 1.74 | 0.002168206 | 1.649540492 | 0.012097497 | 0.991179305 | 0.906023851 |
| A0A1U7R695 | Protein S100 OS=Mesocricetus auratus OX=10036 GN=S100a11 PE=3 SV=1 | 3.766 | 23 | 2 | 3 | 2 | 100 | 11.2 | 7.12 | 0 | 2 | 0 | 141.7 | 0 | 0 | 187.3 | 0 | 0 | 117.5 | 0 | 1.322 | 0.856177831 | 0.829216655 | 0.901961036 | 0.627335825 | 0.770834525 |
| A0A1U7R3I8 | Protein S100 OS=Mesocricetus auratus OX=10036 GN=S100a6 PE=3 SV=1 | 4.299 | 11 | 2 | 6 | 2 | 126 | 13.9 | 6.16 | 14.61 | 2 | 215.6 | 229.5 | 280.5 | 245.6 | 222.8 | 299.9 | 162.5 | 196.7 | 205.2 | 1.139 | 0.662500756 | 0.77783903 | 0.095780437 | 0.734608877 | 0.077064284 |
| A0A1U7R3M2 | Protein S100 OS=Mesocricetus auratus OX=10036 GN=LOC101828213 PE=3 SV=1 | 16.574 | 36 | 4 | 13 | 4 | 90 | 10.3 | 5.71 | 32.23 | 4 | 188.7 | 382.5 | 164.2 | 307.4 | 528.9 | 308.4 | 160.1 | 328.9 | 124.5 | 1.383 | 0.248255589 | 0.834239869 | 0.686451867 | 0.535948283 | 0.143563189 |
| A0A1U7QZ92 | phosphatidylinositide phosphatase SAC1 OS=Mesocricetus auratus OX=10036 GN=Sacm1l PE=4 SV=1 | 4.37 | 4 | 2 | 6 | 2 | 586 | 66.9 | 7.4 | 10.35 | 2 | 90.9 | 77.2 | 0 | 59.9 | 64.5 | 0 | 89.4 | 69.5 | 0 | 0.743 | 0.701255666 | 0.945270672 | 0.941375558 | 1.27733119 | 0.754289617 |
| A0A1U7QUT4 | SUMO-activating enzyme subunit 1 isoform X1 OS=Mesocricetus auratus OX=10036 GN=Sae1 PE=3 SV=1 | 7.5 | 6 | 1 | 6 | 1 | 349 | 38.4 | 5.36 | 33.63 | 1 | 31.2 | 0 | 0 | 35.5 | 0 | 0 | 38.1 | 0 | 0 | 1.139 | 0.931949225 | 1.221153846 | 0.895590285 | 1.073239437 | 0.962585154 |
| A0A1U7QQD5 | GTP-binding protein SAR1a OS=Mesocricetus auratus OX=10036 GN=Sar1a PE=3 SV=1 | 10.176 | 17 | 2 | 5 | 2 | 198 | 22.4 | 6.68 | 14.41 | 2 | 0 | 36.5 | 0 | 0 | 43.7 | 0 | 0 | 34.9 | 0 | 1.196 | 0.905658493 | 0.956164384 | 0.976245536 | 0.798627002 | 0.88294503 |
| A0A1U8CKY3 | sarcosine dehydrogenase, mitochondrial OS=Mesocricetus auratus OX=10036 GN=Sardh PE=3 SV=1 | 3.644 | 1 | 1 | 1 | 1 | 919 | 101.7 | 7.18 | 2.87 | 1 | 120.3 | 0 | 0 | 112.3 | 0 | 0 | 134.4 | 0 | 0 | 0.933 | 0.963569815 | 1.117206983 | 0.941490422 | 1.196794301 | 0.905857655 |
| A0A1U8CJD0 | SAP domain-containing ribonucleoprotein isoform X1 OS=Mesocricetus auratus OX=10036 GN=Sarnp PE=4 SV=1 | 1.958 | 8 | 1 | 2 | 1 | 213 | 23.4 | 6.34 | 2.59 | 1 | 43.2 | 50.5 | 0 | 37.5 | 42.2 | 0 | 39.3 | 41.7 | 0 | 0.852 | 0.83260722 | 0.864461046 | 0.848603856 | 1.016311167 | 0.982897169 |
| A0A1U8BYX9 | U4/U6.U5 tri-snRNP-associated protein 1 isoform X2 OS=Mesocricetus auratus OX=10036 GN=Sart1 PE=3 SV=1 | 1.973 | 2 | 1 | 1 | 1 | 672 | 75.5 | 5.26 | 2.87 | 1 | 0 | 0 | 62.7 | 0 | 0 | 49.8 | 0 | 0 | 52.5 | 0.794 | 0.880197866 | 0.837320574 | 0.906931716 | 1.054216867 | 0.972028628 |
| I1WYE0 | Scavenger receptor class B member 2 OS=Mesocricetus auratus OX=10036 GN=Scarb2 PE=2 SV=1 | 4.346 | 3 | 1 | 4 | 1 | 478 | 53.8 | 5.16 | 11.78 | 1 | 126.7 | 150 | 0 | 112 | 129.3 | 0 | 107.1 | 179.4 | 0 | 0.873 | 0.857953502 | 1.03541742 | 0.964998117 | 1.18731869 | 0.831302849 |
| A0A1U7QZ82 | selenocysteine lyase isoform X1 OS=Mesocricetus auratus OX=10036 GN=Scly PE=4 SV=1 | 8.884 | 8 | 3 | 7 | 3 | 456 | 50.1 | 6.99 | 17.56 | 3 | 85.1 | 106.8 | 93.8 | 67.5 | 80.2 | 79.5 | 83.1 | 110.2 | 88.8 | 0.751 | 0.070550102 | 0.98739937 | 0.913921989 | 1.241637324 | 0.143305239 |
| A0A1U8BWA0 | non-specific lipid-transfer protein isoform X1 OS=Mesocricetus auratus OX=10036 GN=Scp2 PE=3 SV=1 | 5.335 | 2 | 1 | 5 | 1 | 547 | 59 | 7.59 | 14.89 | 1 | 43.1 | 250.3 | 86.2 | 41.5 | 261.1 | 89 | 34.9 | 215.7 | 73.6 | 1.033 | 0.967352862 | 0.854056902 | 0.836375579 | 0.827885598 | 0.80819674 |
| A0A1U7Q7V2 | Succinate dehydrogenase [ubiquinone] flavoprotein subunit, mitochondrial OS=Mesocricetus auratus OX=10036 GN=Sdha PE=3 SV=1 | 15.469 | 9 | 5 | 24 | 5 | 662 | 72.4 | 7.27 | 43.15 | 5 | 359.7 | 306.3 | 463.8 | 367.5 | 296 | 459.4 | 289 | 289.5 | 436.4 | 0.991 | 0.973928151 | 0.898300584 | 0.600413945 | 0.903820465 | 0.62524129 |
| A0A1U8CE56 | Succinate dehydrogenase [ubiquinone] iron-sulfur subunit, mitochondrial OS=Mesocricetus auratus OX=10036 GN=Sdhb PE=3 SV=1 | 15.377 | 19 | 5 | 30 | 5 | 280 | 31.6 | 8.72 | 44.19 | 5 | 1058.5 | 679.2 | 780.5 | 919.7 | 597.1 | 761.8 | 1131.3 | 707.8 | 796.2 | 0.942 | 0.616182911 | 1.046501469 | 0.831517997 | 1.156543492 | 0.500218047 |
| A0A1U7QX40 | serum deprivation-response protein OS=Mesocricetus auratus OX=10036 GN=Cavin2 PE=3 SV=1 | 59.693 | 32 | 14 | 86 | 14 | 415 | 46.2 | 5.2 | 263.94 | 14 | 1837.2 | 2509.4 | 2333.1 | 1631.6 | 2143.8 | 1988.3 | 1630.5 | 2209.9 | 2707 | 0.888 | 0.296915874 | 0.980193721 | 0.911893741 | 1.135971685 | 0.506891839 |
| A0A1U7QKR5 | septin-11 isoform X2 OS=Mesocricetus auratus OX=10036 GN=Sept11 PE=3 SV=1 | 13.309 | 9 | 4 | 9 | 1 | 425 | 48.9 | 6.81 | 23.89 | 4 | 34 | 0 | 42.9 | 30.6 | 0 | 38.8 | 32 | 0 | 40.3 | 0.902 | 0.894063127 | 0.940182055 | 0.936006562 | 1.041786744 | 0.957483666 |
| A0A1U7QXL5 | Selenide, water dikinase OS=Mesocricetus auratus OX=10036 GN=Sephs1 PE=4 SV=1 | 4.649 | 4 | 1 | 3 | 1 | 392 | 42.9 | 5.97 | 5.61 | 1 | 87.5 | 67.2 | 74.3 | 54.4 | 39.2 | 63.1 | 82.1 | 73.8 | 62.5 | 0.718 | 0.059865388 | 0.95371179 | 0.689643825 | 1.393746011 | 0.087109565 |
| A0A1U8C353 | Septin-2 OS=Mesocricetus auratus OX=10036 GN=Sept2 PE=3 SV=1 | 50.927 | 34 | 9 | 50 | 9 | 361 | 41.6 | 6.86 | 179.28 | 9 | 905.6 | 778.7 | 404.6 | 784.5 | 680.5 | 394.5 | 786.9 | 685.4 | 392.3 | 0.898 | 0.709546202 | 0.892622912 | 0.716938315 | 1.002742673 | 0.992324362 |
| A0A1U7QVY1 | septin-4 isoform X1 OS=Mesocricetus auratus OX=10036 GN=Sept4 PE=3 SV=1 | 11.66 | 9 | 4 | 7 | 3 | 478 | 54.9 | 5.87 | 19.64 | 4 | 276 | 67.9 | 230.2 | 166.3 | 51.4 | 165.8 | 199.7 | 48.7 | 185.9 | 0.636 | 0.447445463 | 0.756488417 | 0.590982967 | 1.132464146 | 0.797400682 |
| A0A1U7R0K4 | Septin-7 OS=Mesocricetus auratus OX=10036 GN=Sept7 PE=3 SV=1 | 36.504 | 16 | 8 | 44 | 7 | 532 | 60.6 | 9.16 | 121.23 | 8 | 999.5 | 697.9 | 984.8 | 757 | 497.1 | 690 | 866.9 | 673.9 | 958.5 | 0.712 | 0.124776326 | 0.931809708 | 0.661995382 | 1.285582017 | 0.181646514 |
| A0A1U8BZY0 | septin-9 isoform X1 OS=Mesocricetus auratus OX=10036 GN=Sept9 PE=3 SV=1 | 13.022 | 11 | 5 | 12 | 5 | 584 | 65.5 | 8.95 | 24.33 | 5 | 368.1 | 103 | 119.2 | 248.1 | 96.9 | 99.9 | 319.1 | 79.3 | 111.8 | 0.936 | 0.656719468 | 0.864306285 | 0.826547423 | 1.146774556 | 0.822806215 |
| A0A1U7QKY4 | plasminogen activator inhibitor 1 RNA-binding protein OS=Mesocricetus auratus OX=10036 GN=Serbp1 PE=4 SV=1 | 35.858 | 21 | 9 | 30 | 9 | 402 | 44.2 | 8.29 | 69.25 | 9 | 992.2 | 1247.9 | 1214.6 | 902.3 | 1157 | 1141.1 | 742.8 | 996.1 | 979.6 | 0.927 | 0.502029658 | 0.786899007 | 0.098993949 | 0.849425072 | 0.238718494 |
| A0A1U8BK25 | alpha-1-antitrypsin isoform X1 OS=Mesocricetus auratus OX=10036 GN=Serpina1 PE=3 SV=1 | 45.87 | 22 | 10 | 56 | 10 | 412 | 45.7 | 6.1 | 184.65 | 10 | 594.3 | 750 | 994.4 | 1147.9 | 1322.4 | 1758.6 | 561.9 | 693.6 | 933.4 | 1.924 | 0.0526066 | 0.935947321 | 0.769703057 | 0.517605051 | 0.043272385 |
| A0A1U8CNG4 | serpin B8 OS=Mesocricetus auratus OX=10036 GN=Serpinb8 PE=3 SV=1 | 14.143 | 9 | 2 | 16 | 2 | 374 | 42.5 | 6.54 | 43.06 | 2 | 0 | 0 | 49.7 | 0 | 0 | 25.5 | 0 | 0 | 59 | 0.513 | 0.694248482 | 1.187122736 | 0.91001589 | 2.31372549 | 0.641627848 |
| A0A1U7QNI5 | heparin cofactor 2 OS=Mesocricetus auratus OX=10036 GN=Serpind1 PE=3 SV=1 | 1.411 | 2 | 1 | 1 | 1 | 479 | 54.4 | 6.6 | 0 | 1 | 0 | 100.4 | 0 | 0 | 97 | 0 | 0 | 78.1 | 0 | 0.966 | 0.981737566 | 0.777888446 | 0.869837662 | 0.805154639 | 0.887035579 |
| A0A1U8BM09 | alpha-2-antiplasmin OS=Mesocricetus auratus OX=10036 GN=Serpinf2 PE=3 SV=1 | 13.707 | 11 | 3 | 17 | 3 | 405 | 45.5 | 5.62 | 47.79 | 3 | 207.1 | 210.5 | 172.3 | 366.9 | 361.8 | 303.3 | 155.5 | 189.9 | 159.6 | 1.76 | 0.00654324 | 0.856077301 | 0.159114168 | 0.489341085 | 0.004472889 |
| A0A1U8CDV4 | plasma protease C1 inhibitor OS=Mesocricetus auratus OX=10036 GN=Serping1 PE=3 SV=1 | 8.605 | 7 | 3 | 8 | 3 | 554 | 60.9 | 6.42 | 18.32 | 3 | 49.8 | 64.9 | 28.6 | 80.3 | 88.1 | 53.6 | 63.9 | 60.3 | 22.3 | 1.543 | 0.151647729 | 1.022330775 | 0.95304365 | 0.65990991 | 0.214952552 |
| A0A1U7QK99 | Collagen-binding protein OS=Mesocricetus auratus OX=10036 GN=Serpinh1 PE=3 SV=1 | 28.044 | 20 | 6 | 21 | 6 | 417 | 46.7 | 8.68 | 67.65 | 6 | 72.3 | 100.3 | 61.2 | 47.3 | 84.3 | 59.9 | 61.4 | 112.5 | 50 | 0.979 | 0.425910136 | 0.957656116 | 0.891751927 | 1.169190601 | 0.656620584 |
| A0A1U8D3U0 | Meiosis protein 5 homolog OS=Mesocricetus auratus OX=10036 GN=Sfr1 PE=3 SV=1 | 9.204 | 12 | 3 | 7 | 3 | 444 | 49.1 | 5.02 | 22.3 | 3 | 120.7 | 105.7 | 131.9 | 74.2 | 92 | 97.3 | 119.5 | 118.5 | 114.1 | 0.66 | 0.037876177 | 0.982696065 | 0.813197744 | 1.336242884 | 0.045190739 |
| A0A1U7Q2N8 | Sidoreflexin OS=Mesocricetus auratus OX=10036 GN=Sfxn3 PE=3 SV=1 | 16.096 | 15 | 4 | 19 | 4 | 321 | 35.4 | 9.42 | 48.93 | 4 | 556.4 | 381.1 | 408 | 478.2 | 310 | 310.1 | 526 | 301.3 | 341.6 | 0.814 | 0.351404417 | 0.868747677 | 0.542316973 | 1.064281162 | 0.805112897 |
| Q60538 | Beta-sarcoglycan OS=Mesocricetus auratus OX=10036 GN=SGCB PE=2 SV=1 | 1.702 | 4 | 1 | 2 | 1 | 320 | 34.8 | 8.76 | 6.52 | 1 | 80.5 | 77.8 | 0 | 60.8 | 55.9 | 0 | 42.5 | 51.9 | 0 | 0.736 | 0.696125689 | 0.596336071 | 0.535457566 | 0.80891174 | 0.783260003 |
| A0A1U7Q6I5 | sphingosine-1-phosphate lyase 1 OS=Mesocricetus auratus OX=10036 GN=Sgpl1 PE=3 SV=1 | 2.755 | 1 | 1 | 3 | 1 | 568 | 64 | 9.2 | 4.34 | 1 | 157.6 | 165.2 | 167.6 | 144.5 | 143.9 | 162.7 | 163.5 | 128.7 | 143.9 | 0.907 | 0.155487659 | 0.889274062 | 0.20805212 | 0.966747949 | 0.698049114 |
| A0A1U7QIF2 | SH3 domain-binding glutamic acid-rich-like protein OS=Mesocricetus auratus OX=10036 GN=Sh3bgrl3 PE=3 SV=1 | 3.873 | 20 | 1 | 2 | 1 | 93 | 10.5 | 4.93 | 7.74 | 1 | 37 | 0 | 0 | 51.9 | 0 | 0 | 49 | 0 | 0 | 1.404 | 0.827795374 | 1.324324324 | 0.855251791 | 0.944123314 | 0.969544386 |
| A0A1U7QXU4 | endophilin-A2 OS=Mesocricetus auratus OX=10036 GN=Sh3gl1 PE=3 SV=1 | 10.67 | 6 | 2 | 14 | 2 | 368 | 41.5 | 5.72 | 41.27 | 2 | 322.4 | 379.7 | 485.1 | 368.4 | 450.3 | 455.9 | 336.8 | 387.4 | 484.6 | 1.03 | 0.632775499 | 1.01819407 | 0.916447581 | 0.948375961 | 0.696942416 |
| A0A1U7QV54 | Serine hydroxymethyltransferase OS=Mesocricetus auratus OX=10036 GN=Shmt2 PE=3 SV=1 | 6.054 | 5 | 2 | 4 | 2 | 501 | 55.5 | 8.4 | 10.87 | 2 | 31.9 | 30 | 0 | 49 | 36.4 | 0 | 27.2 | 37 | 0 | 1.364 | 0.687652486 | 1.037156704 | 0.962053002 | 0.75175644 | 0.721787252 |

**Supplementary Table 1: List of differentially expressed proteins identified across the 4dpi infected and 14dpi infected**

| Accession | Description | Sum PEP Score | Coverage [%] | # Peptides | # PSMs | # Unique Peptides | # AAs | MW [kDa] | calc. pI | Score Sequest HT: Sequest HT | # Peptides (by Search Engine): Sequest HT | Mock-infected (4dpi) Replicate_R1 | Mock-infected (4dpi) Replicate_R2 | Mock-infected (4dpi) Replicate_R3 | SARS-CoV-2-infected (4dpi) Replicate_R1 | SARS-CoV-2-infected (4dpi) Replicate_R2 | SARS-CoV-2-infected (4dpi) Replicate_R3 | SARS-CoV-2-(14dpi) Replicate_R1 | SARS-CoV-2-infected (14dpi) Replicate_R2 | SARS-CoV-2-infected (14dpi) Replicate_R3 | Abundance Ratio: SARS-CoV-2-infected (4dpi) / Mock-infected (4dpi) | P-Value: SARS-CoV-2-infected (4dpi) / Mock-infected (4dpi) | Abundance Ratio: SARS-CoV-2-infected (14dpi) / Mock-infected (4dpi) | P-Value: SARS-CoV-2-infected (14dpi) / Mock-infected (4dpi) | Abundance Ratio: SARS-CoV-2-infected (14dpi) / SARS-CoV-2-infected (4dpi) | P-Value: SARS-CoV-2-infected (14dpi) / SARS-CoV-2-infected (4dpi) |
|---|---|---|---|---|---|---|---|---|---|---|---|---|---|---|---|---|---|---|---|---|---|---|---|---|---|
| A0A1U7QLB4 | solute carrier family 12 member 4 isoform X2 OS=Mesocricetus auratus OX=10036 GN=Slc12a4 PE=3 SV=1 | 1.819 | 2 | 1 | 1 | 1 | 1085 | 120.6 | 6.76 | 3.2 | 1 | 37.2 | 0 | 0 | 35.5 | 0 | 0 | 33.9 | 0 | 0 | 0.954 | 0.975213482 | 0.911290323 | 0.950893651 | 0.954929577 | 0.975561801 |
| A0A1U7R1I8 | tricarboxylate transport protein, mitochondrial OS=Mesocricetus auratus OX=10036 GN=Slc25a1 PE=3 SV=1 | 10.392 | 10 | 3 | 15 | 3 | 311 | 34 | 9.86 | 35.18 | 3 | 121.3 | 113.1 | 152.1 | 109.7 | 102.6 | 142.6 | 98.3 | 97.2 | 136.4 | 0.904 | 0.571494132 | 0.858732212 | 0.357972363 | 0.935193012 | 0.68939746 |
| A0A1U7R553 | long-chain fatty acid transport protein 1 isoform X1 OS=Mesocricetus auratus OX=10036 GN=Slc27a1 PE=3 SV=1 | 3.212 | 1 | 1 | 3 | 1 | 843 | 90.9 | 9.31 | 10.71 | 1 | 102.4 | 75.9 | 57.4 | 73.1 | 72.2 | 51 | 89.3 | 83.9 | 65.1 | 0.743 | 0.441350317 | 1.011030972 | 0.957335152 | 1.213958227 | 0.245394303 |
| A0A1U7R690 | zinc transporter SLC39A7 OS=Mesocricetus auratus OX=10036 GN=Slc39a7 PE=4 SV=1 | 2.516 | 3 | 1 | 1 | 1 | 479 | 50.1 | 6.84 | 0 | 1 | 0 | 57.9 | 0 | 0 | 45.4 | 0 | 0 | 59 | 0 | 0.784 | 0.873787598 | 1.018998273 | 0.990020476 | 1.299559471 | 0.864484277 |
| A0A1U7QDP3 | Anion exchange protein OS=Mesocricetus auratus OX=10036 GN=Slc4a1 PE=3 SV=1 | 30.527 | 11 | 9 | 54 | 9 | 919 | 102.4 | 5.26 | 105.86 | 9 | 1330.9 | 1481.2 | 1578.7 | 1370 | 1501.3 | 1586.3 | 1280 | 1411.8 | 1570.1 | 1.014 | 0.827582966 | 0.970643163 | 0.717814887 | 0.956097452 | 0.569979463 |
| A0A1U7QDE6 | Na(+)/H(+) exchange regulatory cofactor NHE-RF OS=Mesocricetus auratus OX=10036 GN=Slc9a3r1 PE=4 SV=1 | 14.211 | 9 | 3 | 17 | 3 | 354 | 38.4 | 6.02 | 63.31 | 3 | 369.1 | 221.5 | 232.2 | 355.9 | 232.1 | 269.5 | 405.3 | 257.9 | 252 | 0.964 | 0.857133097 | 1.112299465 | 0.678844024 | 1.06728863 | 0.773645184 |
| A0A1U7QV91 | Na(+)/H(+) exchange regulatory cofactor NHE-RF OS=Mesocricetus auratus OX=10036 GN=Slc9a3r2 PE=4 SV=1 | 28.572 | 27 | 8 | 29 | 8 | 337 | 37.6 | 7.62 | 93.82 | 8 | 313.7 | 341.3 | 829.2 | 250.3 | 272.4 | 674.7 | 295.4 | 330.1 | 739.3 | 0.798 | 0.682982812 | 0.919552621 | 0.865375257 | 1.139802906 | 0.792428215 |
| A0A1U7QZ27 | sarcolemmal membrane-associated protein isoform X5 OS=Mesocricetus auratus OX=10036 GN=Slmap PE=4 SV=1 | 1.45 | 1 | 1 | 1 | 1 | 821 | 94.1 | 5.22 | 1.77 | 1 | 76.9 | 0 | 0 | 68.2 | 0 | 0 | 65.3 | 0 | 0 | 0.888 | 0.936668027 | 0.849154746 | 0.914137803 | 0.957478006 | 0.976972042 |
| A0A1U7R271 | Structural maintenance of chromosomes protein OS=Mesocricetus auratus OX=10036 GN=Smc1a PE=3 SV=1 | 17.759 | 6 | 6 | 13 | 6 | 1233 | 143.1 | 7.64 | 33.69 | 6 | 290.4 | 484.7 | 450.6 | 248.4 | 398.7 | 393.7 | 272.4 | 419.1 | 399.8 | 0.823 | 0.472891908 | 0.890348372 | 0.587018922 | 1.048520369 | 0.815225083 |
| A0A1U7QCR4 | Structural maintenance of chromosomes protein OS=Mesocricetus auratus OX=10036 GN=Smc3 PE=3 SV=1 | 9.124 | 3 | 3 | 3 | 3 | 1217 | 141.5 | 7.18 | 9.85 | 3 | 0 | 0 | 129.9 | 0 | 0 | 126.1 | 0 | 0 | 118.3 | 0.97 | 0.984259847 | 0.910700539 | 0.950553629 | 0.93814433 | 0.966189069 |
| A0A1U7Q4L2 | U5 small nuclear ribonucleoprotein 200 kDa helicase OS=Mesocricetus auratus OX=10036 GN=Snrnp200 PE=3 SV=1 | 7.513 | 3 | 3 | 7 | 3 | 2136 | 244.4 | 6.06 | 4.97 | 3 | 32.5 | 0 | 0 | 34 | 0 | 0 | 29.4 | 0 | 0 | 1.045 | 0.97608944 | 0.904615385 | 0.947035262 | 0.864705882 | 0.923509879 |
| A0A1U8CV31 | U1 small nuclear ribonucleoprotein 70 kDa OS=Mesocricetus auratus OX=10036 GN=Snrnp70 PE=4 SV=1 | 8.869 | 9 | 3 | 9 | 3 | 444 | 51.7 | 9.94 | 28.33 | 3 | 366.5 | 182.5 | 64.2 | 329.9 | 149.1 | 63.7 | 362.1 | 170.1 | 41.4 | 0.9 | 0.85181556 | 0.935420744 | 0.922920901 | 1.056937535 | 0.936792468 |
| A0A1U8BNJ8 | small nuclear ribonucleoprotein-associated proteins B and B' OS=Mesocricetus auratus OX=10036 GN=Snrpb PE=4 SV=1 | 7.144 | 20 | 3 | 18 | 3 | 83 | 9.7 | 9.25 | 43.9 | 3 | 371 | 469 | 315.2 | 397.8 | 513.7 | 352.7 | 344.1 | 443.6 | 316.2 | 1.101 | 0.609957275 | 0.955592105 | 0.787672683 | 0.873200443 | 0.436681959 |
| A0A1U7RFU3 | Small nuclear ribonucleoprotein Sm D2 OS=Mesocricetus auratus OX=10036 GN=Snrpd2 PE=3 SV=1 | 2.995 | 7 | 1 | 2 | 1 | 118 | 13.5 | 9.91 | 4.67 | 1 | 29.8 | 38.4 | 0 | 36.9 | 34.9 | 0 | 21.5 | 22 | 0 | 1.06 | 0.946169349 | 0.637829912 | 0.586357089 | 0.605849582 | 0.544851032 |
| A0A1U7QQ84 | Small nuclear ribonucleoprotein Sm D3 OS=Mesocricetus auratus OX=10036 GN=Snrpd3 PE=3 SV=1 | 8.214 | 15 | 2 | 15 | 2 | 126 | 13.9 | 10.32 | 40.44 | 2 | 102.4 | 84 | 64.8 | 110.5 | 76.3 | 83.2 | 101.3 | 64.8 | 58.2 | 1.086 | 0.698749902 | 0.892914013 | 0.631703216 | 0.830740741 | 0.423434072 |
| A0A1U7QNB4 | Small nuclear ribonucleoprotein G OS=Mesocricetus auratus OX=10036 GN=Snrpg PE=3 SV=1 | 2.081 | 9 | 1 | 2 | 1 | 76 | 8.5 | 8.88 | 6.16 | 1 | 0 | 358 | 291.2 | 0 | 325.8 | 212.7 | 0 | 277 | 256.3 | 0.815 | 0.812659948 | 0.821472582 | 0.798862835 | 0.990343547 | 0.990049407 |
| A0A1U7QZ16 | SNW domain-containing protein 1 OS=Mesocricetus auratus OX=10036 GN=Snw1 PE=3 SV=1 | 6.126 | 4 | 2 | 7 | 2 | 536 | 61.6 | 9.52 | 15.55 | 2 | 51.7 | 343.3 | 271.7 | 37.8 | 252.7 | 253.8 | 33.6 | 282 | 236.9 | 0.772 | 0.737828984 | 0.828708565 | 0.760214274 | 1.015065221 | 0.980456711 |
| A0A1U7QMY6 | Sorting nexin-1 OS=Mesocricetus auratus OX=10036 GN=Snx1 PE=3 SV=1 | 7.045 | 4 | 2 | 14 | 1 | 522 | 59 | 5.19 | 17.94 | 2 | 225.8 | 281.5 | 205.3 | 226.6 | 247.3 | 199 | 206.5 | 252.3 | 214.3 | 0.969 | 0.651359764 | 0.944569183 | 0.653719428 | 1.000297221 | 0.997487011 |
| A0A1U7QL55 | sorting nexin-12 OS=Mesocricetus auratus OX=10036 GN=Snx12 PE=4 SV=1 | 1.761 | 7 | 1 | 1 | 1 | 166 | 19.2 | 7.25 | 0 | 1 | 0 | 0 | 99.9 | 0 | 0 | 72.1 | 0 | 0 | 81.5 | 0.722 | 0.833573314 | 0.815815816 | 0.893678798 | 1.13037448 | 0.935370222 |
| A0A1U7QG80 | Sorting nexin-3 OS=Mesocricetus auratus OX=10036 GN=Snx3 PE=3 SV=1 | 17.864 | 44 | 5 | 12 | 5 | 162 | 18.8 | 8.66 | 33.55 | 5 | 81 | 63.6 | 105.8 | 70 | 53.5 | 103.6 | 66.4 | 64.1 | 85.7 | 0.979 | 0.706660258 | 0.86341853 | 0.473531566 | 0.952003523 | 0.838300083 |
| A0A1U7QH68 | Superoxide dismutase [Cu-Zn] OS=Mesocricetus auratus OX=10036 GN=Sod1 PE=3 SV=1 | 35.245 | 47 | 8 | 41 | 8 | 154 | 15.8 | 6.51 | 126.99 | 8 | 342.7 | 474 | 648.6 | 383.9 | 468.7 | 701.9 | 318.8 | 451.4 | 655.9 | 1.082 | 0.8303031 | 0.973247799 | 0.926037903 | 0.917401094 | 0.769662343 |
| A0A1U7QH60 | Superoxide dismutase OS=Mesocricetus auratus OX=10036 GN=Sod2 PE=3 SV=1 | 18.846 | 14 | 3 | 39 | 3 | 222 | 24.4 | 8.31 | 86.83 | 3 | 422.6 | 375.4 | 263.7 | 1427.2 | 2374.3 | 1475.4 | 152.8 | 198.8 | 103.7 | 5.493 | 0.042059088 | 0.428840539 | 0.030270485 | 0.086281718 | 0.033927658 |
| A0A1U8BUL8 | Osteonectin OS=Mesocricetus auratus OX=10036 GN=Sparc PE=3 SV=1 | 12.283 | 13 | 4 | 8 | 4 | 307 | 34.9 | 4.84 | 22.52 | 4 | 136.2 | 132.7 | 66.8 | 126.9 | 89.9 | 52.8 | 142.4 | 124.4 | 62.7 | 0.683 | 0.517840598 | 0.981531129 | 0.953143852 | 1.222181009 | 0.569792154 |
| A0A1U7R0Z1 | Microsomal signal peptidase 12 kDa subunit OS=Mesocricetus auratus OX=10036 GN=Spcs1 PE=3 SV=1 | 4.693 | 8 | 1 | 6 | 1 | 162 | 17.7 | 9.55 | 14.79 | 1 | 81.8 | 182.2 | 37.5 | 97.1 | 180.1 | 57.5 | 92.8 | 207.3 | 34.9 | 0.988 | 0.853264351 | 1.111111111 | 0.874670622 | 1.000896325 | 0.998802224 |
| A0A1U7QKA5 | Microsomal signal peptidase 25 kDa subunit OS=Mesocricetus auratus OX=10036 GN=Spcs2 PE=3 SV=1 | 6.848 | 13 | 2 | 9 | 2 | 226 | 25 | 8.57 | 30.95 | 2 | 57.5 | 72.3 | 0 | 52.9 | 79 | 0 | 56.6 | 56.2 | 0 | 1.002 | 0.983616419 | 0.869029276 | 0.854718814 | 0.855193328 | 0.842177201 |
| A0A1U7Q6F7 | Cytospin-A OS=Mesocricetus auratus OX=10036 GN=Specc1l PE=3 SV=1 | 15.5 | 4 | 4 | 11 | 4 | 1118 | 124.4 | 5.64 | 29.39 | 4 | 314.4 | 120.5 | 154.3 | 257.8 | 107.9 | 144.4 | 263.1 | 85.4 | 111.2 | 0.895 | 0.743910396 | 0.780210455 | 0.62471749 | 0.901195844 | 0.826185484 |
| A0A1U7QH09 | SPRY domain-containing protein 3 OS=Mesocricetus auratus OX=10036 GN=Spryd3 PE=4 SV=1 | 2.378 | 2 | 1 | 12 | 1 | 441 | 49.6 | 6.16 | 26.24 | 1 | 342.3 | 311.5 | 348.8 | 623.7 | 560 | 654.4 | 431.9 | 362 | 436.1 | 1.875 | 0.004232799 | 1.226810293 | 0.068835945 | 0.669169251 | 0.005600448 |
| A0A1U7R2R1 | spectrin alpha chain, erythrocytic 1 OS=Mesocricetus auratus OX=10036 GN=Spta1 PE=4 SV=1 | 94.519 | 14 | 22 | 84 | 21 | 2138 | 247.3 | 5.15 | 256.27 | 22 | 1374.1 | 1294.8 | 1234.5 | 1166.3 | 1047.1 | 981.8 | 1110.7 | 1035.4 | 985.7 | 0.809 | 0.028205243 | 0.802326177 | 0.009343669 | 0.980157737 | 0.763898184 |
| A0A1U7R454 | spectrin alpha chain, non-erythrocytic 1 isoform X3 OS=Mesocricetus auratus OX=10036 GN=Sptan1 PE=3 SV=1 | 372.4 | 32 | 80 | 404 | 79 | 2477 | 285.1 | 5.33 | 1116.12 | 80 | 8811.6 | 9166.3 | 9267.2 | 7746 | 7492.5 | 8430 | 9830.9 | 9429.8 | 10572.9 | 0.831 | 0.033148725 | 1.09500791 | 0.108415125 | 1.260477005 | 0.009970204 |
| A0A1U7QBK2 | Spectrin beta chain OS=Mesocricetus auratus OX=10036 GN=Sptbn1 PE=3 SV=1 | 332.661 | 29 | 67 | 388 | 65 | 2363 | 274.1 | 5.59 | 1196.07 | 67 | 7757.7 | 7463.1 | 6945.8 | 6631.6 | 6667.4 | 5881.3 | 7313.1 | 7646.5 | 6490.7 | 0.847 | 0.046712987 | 0.967685617 | 0.601556313 | 1.118350599 | 0.157983779 |
| A0A1U7QIV4 | sulfide:quinone oxidoreductase, mitochondrial OS=Mesocricetus auratus OX=10036 GN=Sqor PE=4 SV=1 | 37.372 | 26 | 6 | 31 | 6 | 352 | 39 | 9.35 | 97.62 | 6 | 328.3 | 505 | 376.5 | 259.5 | 370.7 | 270.6 | 334 | 496.9 | 320.3 | 0.758 | 0.190216154 | 0.951562242 | 0.813371119 | 1.277975133 | 0.291896828 |
| A0A1U7QRN8 | Tyrosine-protein kinase OS=Mesocricetus auratus OX=10036 GN=Src PE=3 SV=1 | 10.144 | 6 | 4 | 18 | 1 | 536 | 59.9 | 7.42 | 39.81 | 4 | 43.6 | 44.4 | 0 | 36.3 | 34.9 | 0 | 34.5 | 34.7 | 0 | 0.809 | 0.782043874 | 0.786363636 | 0.754758489 | 0.971910112 | 0.969805771 |
| A0A1U7QYZ8 | sorcin isoform X2 OS=Mesocricetus auratus OX=10036 GN=Sri PE=4 SV=1 | 1.838 | 5 | 1 | 3 | 1 | 198 | 21.7 | 5.59 | 5.76 | 1 | 0 | 215.5 | 105.5 | 0 | 222.4 | 110.2 | 0 | 190.4 | 96.2 | 1.038 | 0.967576407 | 0.892834891 | 0.896911979 | 0.861695731 | 0.865056056 |
| A0A1U8CKL9 | Arsenite-resistance protein 2 OS=Mesocricetus auratus OX=10036 GN=Srrt PE=3 SV=1 | 5.345 | 4 | 3 | 4 | 3 | 875 | 100.2 | 5.96 | 3.33 | 3 | 0 | 100 | 46.2 | 0 | 80.8 | 44.4 | 0 | 92.8 | 39.1 | 0.881 | 0.860130463 | 0.902188782 | 0.909745491 | 1.053514377 | 0.953081839 |
| A0A1U7QYP9 | Serine/arginine-rich splicing factor 1 OS=Mesocricetus auratus OX=10036 GN=Srsf1 PE=3 SV=1 | 14.251 | 19 | 5 | 21 | 4 | 248 | 27.7 | 10.36 | 47.99 | 5 | 566 | 737.4 | 645.2 | 374.8 | 481 | 493.7 | 354.8 | 467.2 | 443.2 | 0.652 | 0.035974916 | 0.649286667 | 0.023969456 | 0.937532419 | 0.610440049 |
| E7CYH4 | Arginine/serine-rich 3 splicing factor (Fragment) OS=Mesocricetus auratus OX=10036 GN=SFRS3 PE=2 SV=1 | 5.159 | 25 | 2 | 13 | 2 | 122 | 14.5 | 11.85 | 31.08 | 2 | 491 | 551.5 | 478.5 | 380.8 | 431.9 | 373.2 | 352.3 | 402.5 | 363.9 | 0.776 | 0.019895265 | 0.735502959 | 0.010961794 | 0.943334177 | 0.403048543 |
| A0A1U7Q8F4 | serine/arginine-rich splicing factor 5 isoform X2 OS=Mesocricetus auratus OX=10036 GN=Srsf5 PE=3 SV=1 | 2.012 | 3 | 1 | 1 | 1 | 269 | 30.9 | 11.56 | 0 | 1 | 53.8 | 0 | 0 | 46.2 | 0 | 0 | 53.7 | 0 | 0 | 0.859 | 0.919925354 | 0.998141264 | 0.99901334 | 1.162337662 | 0.920887172 |
| A0A1U7Q9C0 | serine/arginine-rich splicing factor 7 isoform X1 OS=Mesocricetus auratus OX=10036 GN=Srsf7 PE=4 SV=1 | 7.231 | 14 | 3 | 15 | 3 | 238 | 27.4 | 11.84 | 32.07 | 3 | 415.5 | 546.6 | 618.3 | 371.5 | 431.6 | 599.2 | 291 | 459.3 | 464.9 | 0.88 | 0.547737811 | 0.768919261 | 0.213505303 | 0.866576339 | 0.522601786 |
| A0A1U7QHF7 | serine/arginine-rich splicing factor 9 OS=Mesocricetus auratus OX=10036 GN=Srsf9 PE=4 SV=1 | 4.848 | 8 | 2 | 9 | 1 | 221 | 25.6 | 8.65 | 25.44 | 2 | 36.1 | 0 | 0 | 40.5 | 0 | 0 | 26.4 | 0 | 0 | 1.123 | 0.939306377 | 0.731301939 | 0.839829996 | 0.651851852 | 0.787273673 |
| A0A1U8CR29 | lupus La protein OS=Mesocricetus auratus OX=10036 GN=Ssb PE=4 SV=1 | 18.268 | 18 | 7 | 13 | 7 | 406 | 46.8 | 9.69 | 32.68 | 7 | 256.1 | 505.9 | 368.3 | 263.5 | 491.2 | 320.3 | 229 | 442.4 | 337.4 | 1.029 | 0.862064412 | 0.892506414 | 0.692149749 | 0.938418605 | 0.822493775 |
| A0A1U7R4L2 | Single-stranded DNA-binding protein, mitochondrial OS=Mesocricetus auratus OX=10036 GN=Ssbp1 PE=3 SV=1 | 1.567 | 9 | 1 | 1 | 1 | 148 | 17.2 | 9.64 | 2.38 | 1 | 64.7 | 0 | 0 | 54.6 | 0 | 0 | 41.9 | 0 | 0 | 0.844 | 0.91094276 | 0.647604328 | 0.784430292 | 0.767399267 | 0.863146387 |
| A0A1U8CBA9 | Stathmin OS=Mesocricetus auratus OX=10036 GN=Stmn1 PE=3 SV=1 | 4.131 | 11 | 2 | 7 | 2 | 149 | 17.3 | 5.97 | 6.78 | 2 | 269.6 | 189.4 | 376.9 | 300.4 | 204.5 | 409.9 | 288.3 | 183.9 | 378.4 | 1.114 | 0.760226325 | 1.017585836 | 0.953020171 | 0.929820726 | 0.806403521 |
| A0A1U7RF35 | Dolichyl-diphosphooligosaccharide--protein glycotransferase OS=Mesocricetus auratus OX=10036 GN=Stt3a PE=3 SV=1 | 12.724 | 6 | 5 | 25 | 5 | 705 | 80.5 | 8.1 | 61.01 | 5 | 728.9 | 718.1 | 550.4 | 604.4 | 622.1 | 473.4 | 742 | 608.6 | 518.5 | 0.86 | 0.256298787 | 0.935766496 | 0.648783476 | 1.099535267 | 0.523710242 |
| A0A1U7RFF3 | syntaxin-binding protein 1 isoform X2 OS=Mesocricetus auratus OX=10036 GN=Stxbp1 PE=3 SV=1 | 16.245 | 9 | 4 | 9 | 4 | 594 | 67.5 | 6.84 | 25.92 | 4 | 0 | 0 | 36.5 | 0 | 0 | 41.3 | 0 | 0 | 32.9 | 1.131 | 0.934848114 | 0.901369863 | 0.945151193 | 0.796610169 | 0.881677711 |
| A0A1U7QL90 | syntaxin-binding protein 6 OS=Mesocricetus auratus OX=10036 GN=Stxbp6 PE=4 SV=1 | 2.304 | 4 | 1 | 3 | 1 | 210 | 23.7 | 9.19 | 7.92 | 1 | 0 | 40.4 | 0 | 0 | 35.6 | 0 | 0 | 35.2 | 0 | 0.881 | 0.933319059 | 0.871287129 | 0.927442122 | 0.988764045 | 0.994007791 |
| A0A1U7QZR6 | Activated RNA polymerase II transcriptional coactivator p15 OS=Mesocricetus auratus OX=10036 GN=Sub1 PE=3 SV=1 | 7.884 | 12 | 2 | 8 | 2 | 127 | 14.4 | 9.6 | 20.99 | 2 | 244.9 | 128.5 | 168.5 | 209.1 | 109.2 | 142.5 | 156 | 98.2 | 108.6 | 0.846 | 0.581376115 | 0.669496217 | 0.218449491 | 0.787326389 | 0.405867096 |
| A0A1U7QF78 | Succinate--CoA ligase [ADP-forming] subunit beta, mitochondrial OS=Mesocricetus auratus OX=10036 GN=Sucla2 PE=3 SV=1 | 7.346 | 6 | 3 | 5 | 3 | 463 | 50.1 | 7.68 | 4.21 | 3 | 0 | 0 | 59.6 | 0 | 0 | 68.8 | 0 | 0 | 59.8 | 1.155 | 0.924451706 | 1.003355705 | 0.998223356 | 0.869186047 | 0.926188967 |
| A0A1U7QF52 | protein SGT1 homolog OS=Mesocricetus auratus OX=10036 GN=Sugt1 PE=3 SV=1 | 4.528 | 11 | 2 | 2 | 2 | 336 | 38.1 | 5.3 | 3.34 | 2 | 0 | 0 | 74.3 | 0 | 0 | 95.5 | 0 | 0 | 51.2 | 1.286 | 0.869915266 | 0.68909825 | 0.812084331 | 0.536125654 | 0.709584846 |
| A0A1U8C6W7 | transcription factor SPT20 homolog isoform X1 OS=Mesocricetus auratus OX=10036 GN=Supt20h PE=3 SV=1 | 1.722 | 1 | 1 | 1 | 1 | 858 | 93.5 | 8.18 | 2.38 | 1 | 44.7 | 0 | 0 | 32.7 | 0 | 0 | 42.2 | 0 | 0 | 0.732 | 0.839987489 | 0.944071588 | 0.969515438 | 1.290519878 | 0.867923596 |
| A0A1U8C7S0 | Phosphoinositide 5-phosphatase OS=Mesocricetus auratus OX=10036 GN=Synj1 PE=3 SV=1 | 3.314 | 1 | 1 | 3 | 1 | 1555 | 170.2 | 6.89 | 10.59 | 1 | 0 | 0 | 52.1 | 0 | 0 | 44.7 | 0 | 0 | 41.3 | 0.857 | 0.919461439 | 0.792706334 | 0.879220062 | 0.92393736 | 0.95814276 |
| A0A1U7R1X8 | synemin isoform X2 OS=Mesocricetus auratus OX=10036 GN=Synm PE=3 SV=1 | 27.454 | 7 | 6 | 21 | 6 | 1258 | 139.9 | 5.02 | 53.51 | 6 | 107.8 | 0 | 0 | 103.9 | 0 | 0 | 74.4 | 0 | 0 | 0.964 | 0.980467891 | 0.690166976 | 0.812792332 | 0.716073147 | 0.829872146 |
| A0A1U8BXX5 | Transgelin OS=Mesocricetus auratus OX=10036 GN=Tagln PE=3 SV=1 | 78.504 | 68 | 15 | 88 | 15 | 201 | 22.6 | 9 | 210.29 | 15 | 2581.2 | 2829.2 | 2488.2 | 1591.2 | 2199.5 | 1679 | 1813.1 | 2469.4 | 1833.7 | 0.702 | 0.031774151 | 0.774339756 | 0.092621081 | 1.11819661 | 0.495283709 |
| A0A1U7QPL5 | Transgelin OS=Mesocricetus auratus OX=10036 GN=Tagln2 PE=3 SV=1 | 87.501 | 69 | 14 | 105 | 13 | 199 | 22.4 | 8.25 | 296.63 | 14 | 1560.6 | 1646.8 | 1338.3 | 1875.3 | 1265.4 | 1643.9 | 1832.9 | 1675.6 | 1758.7 | 0.924 | 0.717316175 | 1.158721429 | 0.102985707 | 1.100865276 | 0.463752013 |
| A0A1U7R3G6 | tapasin OS=Mesocricetus auratus OX=10036 GN=Tapbp PE=4 SV=1 | 3.668 | 3 | 1 | 1 | 1 | 456 | 49 | 7.62 | 3.61 | 1 | 70.1 | 0 | 0 | 57 | 0 | 0 | 56.9 | 0 | 0 | 0.814 | 0.892006 | 0.811697575 | 0.891118232 | 0.998245614 | 0.99906878 |
| A0A1U7Q6F4 | Tubulin-specific chaperone A OS=Mesocricetus auratus OX=10036 GN=Tbca PE=3 SV=1 | 2.151 | 12 | 1 | 2 | 1 | 108 | 12.6 | 5.44 | 0 | 1 | 0 | 104.7 | 48.3 | 0 | 71.4 | 35.4 | 0 | 95.6 | 51.1 | 0.707 | 0.698338725 | 0.958823529 | 0.961593136 | 1.373595506 | 0.720701511 |
| A0A1U7R8N1 | tubulin-folding cofactor B OS=Mesocricetus auratus OX=10036 GN=Tbcb PE=4 SV=1 | 9.386 | 14 | 2 | 8 | 2 | 242 | 27 | 5 | 24.45 | 2 | 81.3 | 55.7 | 93.6 | 92 | 70.1 | 110.3 | 69.9 | 65.4 | 90.5 | 1.258 | 0.436051496 | 0.979184735 | 0.912530996 | 0.828928047 | 0.336586198 |
| A0A1U8BDZ2 | T-complex protein 1 subunit alpha OS=Mesocricetus auratus OX=10036 GN=Tcp1 PE=3 SV=1 | 4.853 | 11 | 2 | 7 | 2 | 285 | 30.6 | 7.91 | 16.03 | 2 | 207.6 | 211.7 | 94.1 | 151.3 | 150.5 | 74 | 180.8 | 204.4 | 82.4 | 0.711 | 0.385479438 | 0.910790806 | 0.790201256 | 1.244278872 | 0.540946272 |
| A0A1U8CFH0 | Very-long-chain enoyl-CoA reductase OS=Mesocricetus auratus OX=10036 GN=Tecr PE=3 SV=1 | 5.439 | 9 | 3 | 15 | 3 | 291 | 34 | 9.57 | 32.93 | 3 | 104.4 | 121.6 | 0 | 105.5 | 118.3 | 0 | 79.7 | 101.8 | 0 | 0.992 | 0.989695136 | 0.803097345 | 0.777702536 | 0.810991957 | 0.786581004 |
| A0A1U7QM87 | teneurin-1 isoform X2 OS=Mesocricetus auratus OX=10036 GN=Tenm1 PE=3 SV=1 | 1.539 | 0 | 1 | 1 | 1 | 2731 | 305.5 | 6.33 | 2.48 | 1 | 0 | 0 | 318.8 | 0 | 0 | 161.9 | 0 | 0 | 178.7 | 0.508 | 0.69077991 | 0.560539523 | 0.725927703 | 1.103767758 | 0.947830326 |
| A0A1U8BSW4 | serotransferrin OS=Mesocricetus auratus OX=10036 GN=LOC101828182 PE=3 SV=1 | 275.073 | 54 | 43 | 370 | 42 | 693 | 76.1 | 7.33 | 1155.92 | 43 | 6388.9 | 6426.7 | 6609.3 | 13669.4 | 15253.2 | 14400.2 | 6377.9 | 6480.9 | 6241.7 | 2.179 | 0.002767407 | 0.983299785 | 0.327876369 | 0.440887939 | 0.002674368 |
| A0A1U7QGF1 | Transforming growth factor-beta-induced protein ig-h3 OS=Mesocricetus auratus OX=10036 GN=Tgfbi PE=4 SV=1 | 24.638 | 9 | 5 | 23 | 5 | 683 | 74.6 | 6.9 | 72.48 | 5 | 101.5 | 94.4 | 100.1 | 110.8 | 90.1 | 104.4 | 91.8 | 91.4 | 93.1 | 1.091 | 0.671661516 | 0.933445946 | 0.087016999 | 0.905011464 | 0.254401197 |
| A0A1U7R7M7 | protein-glutamine gamma-glutamyltransferase 2 OS=Mesocricetus auratus OX=10036 GN=Tgm2 PE=3 SV=1 | 120.658 | 31 | 22 | 159 | 22 | 686 | 77 | 5.08 | 480.12 | 22 | 2536.4 | 3450.4 | 3291.5 | 2397.9 | 3343 | 3343 | 2352.3 | 3489.6 | 3460.8 | 0.969 | 0.885689763 | 1.002629792 | 0.987045581 | 1.024086571 | 0.888922489 |
| A0A1U7QHW5 | Thrombomodulin OS=Mesocricetus auratus OX=10036 GN=Thbd PE=4 SV=1 | 7.097 | 3 | 1 | 11 | 1 | 580 | 62.9 | 4.68 | 35.93 | 1 | 121.7 | 52 | 77.2 | 97.9 | 41.3 | 51.8 | 131.5 | 65.9 | 95.5 | 0.805 | 0.498365258 | 1.167397369 | 0.641581005 | 1.533507853 | 0.257730386 |
| A0A1U7Q315 | thrombospondin-1 isoform X1 OS=Mesocricetus auratus OX=10036 GN=Thbs1 PE=3 SV=1 | 16.493 | 5 | 5 | 19 | 5 | 1171 | 129.6 | 4.91 | 56.68 | 5 | 160.1 | 177 | 179.4 | 172 | 205.6 | 233.4 | 161.2 | 148.3 | 190.4 | 1.258 | 0.211245764 | 0.9678606 | 0.71719989 | 0.818166939 | 0.171069779 |

Supplementary Table 1: List of differentially expressed proteins identified across the 4dpi infected and 14dpi infected

| Accession | Description | Sum PEP Score | Coverage [%] | # Peptides | # PSMs | # Unique Peptides | # AAs | MW [kDa] | calc. pI | Score Sequest HT: Sequest HT | # Peptides (by Search Engine): Sequest HT | Mock-infected (4dpi) Replicate_R1 | Mock-infected (4dpi) Replicate_R2 | Mock-infected (4dpi) Replicate_R3 | SARS-CoV-2-infected (4dpi) Replicate_R1 | SARS-CoV-2-infected (4dpi) Replicate_R2 | SARS-CoV-2-infected (4dpi) Replicate_R3 | SARS-CoV-2-(14dpi) Replicate_R1 | SARS-CoV-2-infected (14dpi) Replicate_R2 | SARS-CoV-2-infected (14dpi) Replicate_R3 | Abundance Ratio: SARS-CoV-2-infected (4dpi) / Mock-infected (4dpi) | P-Value: SARS-CoV-2-infected (4dpi) / Mock-infected (4dpi) | Abundance Ratio: SARS-CoV-2-infected (14dpi) / Mock-infected (4dpi) | P-Value: SARS-CoV-2-infected (14dpi) / Mock-infected (4dpi) | Abundance Ratio: SARS-CoV-2-infected (14dpi) / SARS-CoV-2-infected (4dpi) | P-Value: SARS-CoV-2-infected (14dpi) / SARS-CoV-2-infected (4dpi) |
|---|---|---|---|---|---|---|---|---|---|---|---|---|---|---|---|---|---|---|---|---|---|---|---|---|---|
| A0A1U7R439 | protein THEM6 OS=Mesocricetus auratus OX=10036 GN=Them6 PE=4 SV=1 | 3.819 | 7 | 2 | 3 | 2 | 208 | 23.8 | 9.35 | 4.48 | 2 | 106.2 | 0 | 0 | 130.6 | 0 | 0 | 95.6 | 0 | 0 | 1.229 | 0.892034536 | 0.900188324 | 0.944463996 | 0.732006126 | 0.840288828 |
| A0A1U8CIM1 | thyroid hormone receptor-associated protein 3 OS=Mesocricetus auratus OX=10036 GN=Thrap3 PE=3 SV=1 | 10.556 | 3 | 2 | 3 | 2 | 952 | 108.3 | 10.17 | 8.28 | 2 | 0 | 96.9 | 0 | 0 | 97.6 | 0 | 0 | 54.4 | 0 | 1.007 | 0.996182771 | 0.561403509 | 0.726507405 | 0.557377049 | 0.72380646 |
| A0A1U7RBK3 | Mitochondrial import inner membrane translocase subunit TIM44 OS=Mesocricetus auratus OX=10036 GN=Timm44 PE=3 SV=1 | 3.328 | 5 | 2 | 2 | 2 | 452 | 51 | 8.46 | 2.51 | 2 | 0 | 0 | 58.9 | 0 | 0 | 87.8 | 0 | 0 | 60.7 | 1.49 | 0.799954573 | 1.030560272 | 0.984041079 | 0.691343964 | 0.813571764 |
| A0A1U8CU64 | tight junction protein ZO-1 isoform X7 OS=Mesocricetus auratus OX=10036 GN=Tjp1 PE=4 SV=1 | 124.207 | 18 | 24 | 124 | 1 | 1782 | 198.9 | 6.79 | 418.12 | 24 | 35.8 | 80.3 | 51.1 | 33.7 | 70.3 | 58.2 | 31.9 | 71.9 | 44 | 1.125 | 0.926435167 | 0.883971292 | 0.732448801 | 0.911220715 | 0.779319973 |
| A0A1U7RDZ8 | transketolase OS=Mesocricetus auratus OX=10036 GN=Tkt PE=4 SV=1 | 38.469 | 18 | 10 | 51 | 10 | 623 | 67.6 | 7.49 | 111.45 | 10 | 780 | 824.9 | 1172.4 | 761.6 | 811.5 | 1094.8 | 764.1 | 881.7 | 1081.3 | 0.934 | 0.832933143 | 0.981924891 | 0.919494526 | 1.022189737 | 0.894095962 |
| A0A1U7R141 | talin-1 OS=Mesocricetus auratus OX=10036 GN=Tln1 PE=4 SV=1 | 286.994 | 27 | 68 | 329 | 63 | 2541 | 269.5 | 6.18 | 1024.56 | 68 | 8537 | 8214.6 | 7953.5 | 7243.4 | 6887.6 | 6520.3 | 7735.3 | 7288.6 | 7083.7 | 0.83 | 0.008184613 | 0.894859766 | 0.028424063 | 1.070518563 | 0.162906381 |
| A0A1U7QJZ7 | talin-2 isoform X2 OS=Mesocricetus auratus OX=10036 GN=Tln2 PE=4 SV=1 | 23.18 | 2 | 6 | 30 | 1 | 2544 | 271.7 | 5.57 | 60.54 | 6 | 0 | 38.1 | 0 | 0 | 29.1 | 0 | 0 | 26.3 | 0 | 0.764 | 0.860828064 | 0.690288714 | 0.812872964 | 0.903780069 | 0.94655085 |
| A0A1U7R4U9 | Transmembrane 9 superfamily member OS=Mesocricetus auratus OX=10036 GN=Tm9sf2 PE=3 SV=1 | 6.335 | 2 | 1 | 10 | 1 | 661 | 75.4 | 7.44 | 29.27 | 1 | 122.2 | 204.7 | 116.1 | 113.2 | 200.8 | 133.3 | 109.4 | 211.3 | 88.4 | 0.946 | 0.972427706 | 0.923476298 | 0.824519273 | 0.914598703 | 0.798392567 |
| A0A1U7Q4M1 | Transmembrane 9 superfamily member OS=Mesocricetus auratus OX=10036 GN=Tm9sf3 PE=3 SV=1 | 2.039 | 4 | 1 | 1 | 1 | 587 | 67.5 | 7.21 | 0 | 1 | 0 | 0 | 40.7 | 0 | 0 | 23.5 | 0 | 0 | 48 | 0.577 | 0.737248956 | 1.179361179 | 0.913386874 | 2.042553191 | 0.678741427 |
| A0A1U7QME7 | transmembrane emp24 domain-containing protein 7 OS=Mesocricetus auratus OX=10036 GN=Tmed7 PE=3 SV=1 | 3.742 | 4 | 1 | 7 | 1 | 225 | 25.2 | 6.2 | 27.24 | 1 | 273.4 | 229.5 | 188.7 | 264 | 230.3 | 183.1 | 267.7 | 233.3 | 169.9 | 0.97 | 0.895685675 | 0.970069404 | 0.863759296 | 0.990404488 | 0.95624605 |
| A0A1U7QGD6 | transmembrane emp24 domain-containing protein 9 OS=Mesocricetus auratus OX=10036 GN=Tmed9 PE=3 SV=1 | 8.161 | 11 | 2 | 8 | 2 | 235 | 27.2 | 7.58 | 3.47 | 2 | 332.7 | 250.7 | 228.9 | 379.1 | 290.8 | 287.3 | 296.7 | 256.7 | 225 | 1.16 | 0.330179002 | 0.95826665 | 0.782090557 | 0.813205182 | 0.186681747 |
| A0A1U7Q615 | Hyaluronoglucosaminidase OS=Mesocricetus auratus OX=10036 GN=Tmem2 PE=3 SV=1 | 7.964 | 2 | 2 | 9 | 2 | 1383 | 153.7 | 7.65 | 40.47 | 2 | 528.5 | 369.4 | 141.6 | 411.7 | 287.5 | 106.3 | 468.9 | 341.3 | 131.2 | 0.778 | 0.616057507 | 0.905627706 | 0.837554987 | 1.168715084 | 0.7498134 |
| A0A1U7R109 | tropomodulin-1 OS=Mesocricetus auratus OX=10036 GN=Tmod1 PE=4 SV=1 | 5.223 | 10 | 2 | 2 | 2 | 359 | 40.4 | 5.1 | 7.43 | 2 | 0 | 53.9 | 0 | 0 | 47.1 | 0 | 0 | 52.5 | 0 | 0.874 | 0.928962182 | 0.974025974 | 0.986046746 | 1.114649682 | 0.942689533 |
| A0A1U7QT87 | Thymosin beta OS=Mesocricetus auratus OX=10036 GN=Tmsb4x PE=3 SV=1 | 21.525 | 57 | 7 | 37 | 5 | 44 | 5.1 | 5.06 | 44.94 | 7 | 238.7 | 274.6 | 161 | 572.3 | 890.3 | 850.4 | 509.3 | 823.6 | 817.4 | 3.243 | 0.022771561 | 3.188936675 | 0.031885548 | 0.929658452 | 0.725799354 |
| A0A1U7QFR6 | 182 kDa tankyrase-1-binding protein OS=Mesocricetus auratus OX=10036 GN=Tnks1bp1 PE=4 SV=1 | 47.393 | 16 | 13 | 35 | 13 | 1696 | 178.4 | 4.88 | 107.01 | 13 | 385.9 | 211.8 | 558.7 | 334.9 | 168.3 | 475.3 | 366.6 | 228.6 | 509 | 0.868 | 0.680843887 | 0.95485991 | 0.899309878 | 1.128461932 | 0.744926796 |
| A0A1U7RDR7 | troponin C, skeletal muscle OS=Mesocricetus auratus OX=10036 GN=Tnnc2 PE=4 SV=1 | 1.508 | 4 | 1 | 3 | 1 | 160 | 18.1 | 4.2 | 1.88 | 1 | 154.4 | 207.8 | 191 | 169 | 198.4 | 177 | 118.2 | 202.9 | 181.4 | 0.955 | 0.880781346 | 0.90835141 | 0.607814424 | 0.923034533 | 0.646374051 |
| A0A1U7Q6E2 | transportin-1 OS=Mesocricetus auratus OX=10036 GN=Tnpo1 PE=4 SV=1 | 4.725 | 3 | 2 | 4 | 2 | 890 | 101.2 | 4.96 | 9.41 | 2 | 0 | 0 | 41.5 | 0 | 0 | 42.5 | 0 | 0 | 44.3 | 1.024 | 0.987375168 | 1.06746988 | 0.965429064 | 1.042352941 | 0.978015705 |
| P86216 | Triosephosphate isomerase OS=Mesocricetus auratus OX=10036 GN=TPI1 PE=1 SV=2 | 65.369 | 53 | 11 | 80 | 11 | 249 | 26.8 | 6.55 | 252.19 | 11 | 1232.9 | 1553.7 | 1549.2 | 1063.3 | 1290.4 | 1273.8 | 1112 | 1519.5 | 1542 | 0.822 | 0.150059353 | 0.962567462 | 0.774320439 | 1.150516885 | 0.331591526 |
| A0A1U7QBL7 | tropomyosin alpha-1 chain isoform X17 OS=Mesocricetus auratus OX=10036 GN=Tpm1 PE=3 SV=1 | 91.588 | 54 | 20 | 133 | 1 | 248 | 28.7 | 4.78 | 302.02 | 20 | 549.9 | 354.5 | 248.7 | 171 | 105.1 | 67.2 | 492.8 | 292.9 | 228.8 | 0.311 | 0.079456164 | 0.879802272 | 0.717279919 | 2.955141276 | 0.092072735 |
| A0A1U7QVG1 | tropomyosin alpha-1 chain isoform X6 OS=Mesocricetus auratus OX=10036 GN=Tpm1 PE=3 SV=1 | 88.664 | 55 | 21 | 132 | 2 | 284 | 32.6 | 4.73 | 286.92 | 21 | 748.4 | 1199.5 | 780 | 1264.5 | 1333.1 | 1614.4 | 1210.7 | 1486.8 | 1451.5 | 1.642 | 0.056954577 | 1.520950181 | 0.061703696 | 0.985042735 | 0.88654236 |
| A0A1U8C992 | tropomyosin beta chain isoform X3 OS=Mesocricetus auratus OX=10036 GN=Tpm2 PE=3 SV=1 | 79.383 | 47 | 17 | 124 | 2 | 284 | 33 | 4.69 | 302.57 | 17 | 282.6 | 97.4 | 26.6 | 234.8 | 72.1 | 40 | 222.8 | 81.9 | 34 | 1.014 | 0.848387153 | 0.833005411 | 0.824425424 | 0.976362064 | 0.975236546 |
| A0A1U7QJJ2 | tropomyosin alpha-3 chain isoform X3 OS=Mesocricetus auratus OX=10036 GN=Tpm3 PE=3 SV=1 | 102.63 | 60 | 21 | 178 | 2 | 248 | 29 | 4.78 | 388.94 | 21 | 865.1 | 624 | 594.5 | 984.9 | 667.3 | 638.4 | 1052.8 | 765.5 | 725.7 | 1.07 | 0.650036709 | 1.220963717 | 0.318008627 | 1.110626037 | 0.606936153 |
| A0A1U7QZG6 | Tubulin polymerization-promoting protein family member 3 OS=Mesocricetus auratus OX=10036 GN=Tppp3 PE=3 SV=1 | 43.495 | 41 | 8 | 91 | 8 | 176 | 19 | 9.11 | 215.52 | 8 | 4928.3 | 4482.2 | 5139.2 | 3190.4 | 2467.4 | 2948.3 | 4439.1 | 3701.3 | 4444.8 | 0.567 | 0.002397955 | 0.864980034 | 0.109136981 | 1.462358095 | 0.015885301 |
| A0A1U7QS36 | Nucleoprotein TPR OS=Mesocricetus auratus OX=10036 GN=Tpr PE=3 SV=1 | 18.443 | 2 | 4 | 13 | 4 | 2417 | 272.4 | 5.02 | 28.76 | 4 | 0 | 0 | 354.5 | 0 | 0 | 301 | 0 | 0 | 349.2 | 0.849 | 0.914094436 | 0.985049365 | 0.992012038 | 1.16013289 | 0.921869342 |
| A0A1U7QF68 | translationally-controlled tumor protein OS=Mesocricetus auratus OX=10036 GN=Tpt1 PE=3 SV=1 | 19.193 | 18 | 4 | 33 | 4 | 172 | 19.5 | 4.86 | 89.07 | 4 | 930.5 | 1220.3 | 850.2 | 1021.5 | 1234.2 | 851.6 | 881.7 | 1108 | 783.2 | 1.096 | 0.833280947 | 0.923992003 | 0.634934856 | 0.892382454 | 0.490201154 |
| A0A1U8CV59 | Trafficking protein particle complex subunit OS=Mesocricetus auratus OX=10036 GN=Trappc3 PE=3 SV=1 | 5.794 | 11 | 2 | 12 | 2 | 180 | 20.3 | 4.96 | 32.04 | 2 | 130.8 | 74.7 | 33.6 | 142.4 | 84.1 | 43.2 | 101.7 | 70.9 | 27.7 | 1.127 | 0.812555096 | 0.837724801 | 0.734685173 | 0.742677049 | 0.557136421 |
| A0A1U8BQR0 | CCA tRNA nucleotidyltransferase 1, mitochondrial OS=Mesocricetus auratus OX=10036 GN=Trnt1 PE=3 SV=1 | 1.77 | 2 | 1 | 1 | 1 | 434 | 50 | 8.48 | 1.91 | 1 | 0 | 0 | 123.1 | 0 | 0 | 92.2 | 0 | 0 | 76.4 | 0.749 | 0.851305248 | 0.620633631 | 0.766331051 | 0.828633406 | 0.901602177 |
| A0A1U7QB17 | transient receptor potential cation channel subfamily M member 3 isoform X14 OS=Mesocricetus auratus OX=10036 GN=Trpm3 PE=4 SV=1 | 3.712 | 1 | 2 | 12 | 2 | 1668 | 190.2 | 7.27 | 19.04 | 2 | 294.5 | 816.6 | 278.6 | 296.9 | 957.9 | 4504.7 | 211.4 | 711.2 | 558.5 | 2.238 | 0.38087181 | 1.065769591 | 0.901391859 | 0.257157739 | 0.388985583 |
| A0A1U8CML8 | tetraspanin-8 OS=Mesocricetus auratus OX=10036 GN=Tspan8 PE=4 SV=1 | 25.543 | 10 | 2 | 10 | 2 | 163 | 17.7 | 4.75 | 42.98 | 2 | 366.4 | 129.4 | 125.5 | 238.6 | 146.1 | 130.6 | 386.5 | 172.2 | 158.6 | 1.041 | 0.713183845 | 1.154514727 | 0.782981708 | 1.392004657 | 0.471434598 |
| A0A1U7Q653 | Tubulin alpha chain OS=Mesocricetus auratus OX=10036 GN=Tuba4a PE=3 SV=1 | 126.748 | 48 | 21 | 266 | 6 | 448 | 49.9 | 5.06 | 911.14 | 21 | 3021.9 | 3767.9 | 3889.1 | 4038.7 | 4043.8 | 4101.2 | 3454.9 | 3816.6 | 3984.2 | 1.073 | 0.204968581 | 1.054013054 | 0.579903566 | 0.923832662 | 0.18435008 |
| A0A1U7QRM6 | Tubulin beta chain OS=Mesocricetus auratus OX=10036 GN=Tubb3 PE=3 SV=1 | 95.036 | 36 | 13 | 153 | 3 | 450 | 50.4 | 4.93 | 593.96 | 13 | 362.5 | 250.2 | 419.2 | 327.5 | 214.4 | 463.7 | 323.7 | 195.1 | 362.7 | 0.857 | 0.925609441 | 0.854249443 | 0.518652997 | 0.87659109 | 0.665705418 |
| A0A1U7R1I6 | Tubulin beta chain OS=Mesocricetus auratus OX=10036 GN=Tubb4b PE=3 SV=1 | 135.468 | 49 | 19 | 213 | 2 | 445 | 49.8 | 4.89 | 812.92 | 19 | 2169.8 | 1460.2 | 1984.1 | 2386.5 | 2625.7 | 2190.3 | 2475.4 | 2727.9 | 2382 | 1.1 | 0.114428249 | 1.35111594 | 0.0717399 | 1.053148212 | 0.478684104 |
| A0A1U7Q587 | Elongation factor Tu OS=Mesocricetus auratus OX=10036 GN=Tufm PE=3 SV=1 | 46.925 | 27 | 9 | 48 | 9 | 452 | 49.6 | 7.56 | 140.75 | 9 | 403.2 | 177.8 | 445 | 367.6 | 150.4 | 383.4 | 339.5 | 152.2 | 425.1 | 0.846 | 0.729665767 | 0.893567251 | 0.768742286 | 1.017084535 | 0.965089916 |
| A0A1U7QAR2 | twinfilin-2 OS=Mesocricetus auratus OX=10036 GN=Twf2 PE=3 SV=1 | 10.836 | 11 | 3 | 15 | 3 | 349 | 39.3 | 6.8 | 30.45 | 3 | 43.2 | 41.3 | 52.5 | 48.1 | 41.5 | 55 | 42.1 | 31.1 | 33.3 | 1.011 | 0.652686948 | 0.777372263 | 0.102820251 | 0.736514523 | 0.070467488 |
| A0A1U7Q7V1 | Thioredoxin domain-containing protein 12 OS=Mesocricetus auratus OX=10036 GN=Txndc12 PE=4 SV=1 | 8.742 | 14 | 2 | 9 | 2 | 170 | 19 | 5.4 | 21.44 | 2 | 0 | 30.5 | 0 | 0 | 38.2 | 0 | 0 | 29.3 | 0 | 1.252 | 0.882820884 | 0.960655738 | 0.978725783 | 0.767015707 | 0.862900949 |
| A0A1U7Q8U8 | thioredoxin domain-containing protein 5 OS=Mesocricetus auratus OX=10036 GN=Txndc5 PE=3 SV=1 | 23.103 | 14 | 4 | 29 | 4 | 417 | 46.2 | 5.8 | 101.53 | 4 | 132.6 | 70.7 | 223.6 | 115.8 | 49.6 | 169 | 151 | 100.4 | 255.3 | 0.756 | 0.614429849 | 1.186929023 | 0.697486602 | 1.515251196 | 0.376043291 |
| A0A1U7QIL4 | thioredoxin-like protein 1 isoform X1 OS=Mesocricetus auratus OX=10036 GN=Txnl1 PE=4 SV=1 | 15.972 | 24 | 4 | 17 | 4 | 289 | 32.2 | 4.96 | 27.32 | 4 | 71.5 | 56.4 | 97.9 | 67.1 | 48.3 | 72.9 | 57.6 | 63.3 | 108.5 | 0.857 | 0.438441228 | 1.015943313 | 0.955589842 | 1.21826872 | 0.499386453 |
| A0A1U7QYQ1 | Thioredoxin-disulfide reductase OS=Mesocricetus auratus OX=10036 GN=Txnrd1 PE=3 SV=1 | 12.447 | 8 | 4 | 14 | 4 | 610 | 66.7 | 7.3 | 29.43 | 4 | 270.6 | 291.6 | 465.9 | 243 | 238.1 | 367.5 | 236.9 | 274.9 | 409.7 | 0.789 | 0.475092479 | 0.896313588 | 0.684545392 | 1.085906198 | 0.737405702 |
| A0A1U7QQ69 | UDP-N-acetylhexosamine pyrophosphorylase-like protein 1 OS=Mesocricetus auratus OX=10036 GN=Uap1l1 PE=4 SV=1 | 5.479 | 4 | 2 | 2 | 2 | 507 | 56.4 | 5.86 | 4.75 | 2 | 0 | 153.3 | 0 | 0 | 107 | 0 | 0 | 124.3 | 0 | 0.698 | 0.817959054 | 0.810828441 | 0.890576931 | 1.161682243 | 0.921178885 |
| A0A1U8CLZ3 | Ubiquitin-activating enzyme E1 OS=Mesocricetus auratus OX=10036 GN=Uba1 PE=3 SV=1 | 30.691 | 12 | 9 | 24 | 9 | 1058 | 117.7 | 5.6 | 51.5 | 9 | 341.8 | 280.6 | 342.7 | 269 | 270.4 | 283.4 | 287 | 309.1 | 316.4 | 0.884 | 0.141537273 | 0.945497876 | 0.495802064 | 1.109017987 | 0.057440851 |
| A0A1U7R573 | 60S ribosomal protein L40 OS=Mesocricetus auratus OX=10036 GN=Uba52 PE=3 SV=1 | 45.44 | 56 | 9 | 90 | 9 | 128 | 14.7 | 9.83 | 173.03 | 9 | 1467.7 | 1155.7 | 1198.9 | 1658.8 | 1337.4 | 1559.1 | 1790.1 | 1460.7 | 1311.7 | 1.13 | 0.147302574 | 1.193653036 | 0.232699416 | 1.001580576 | 0.989520975 |
| A0A1U7Q231 | SUMO-conjugating enzyme UBC9 OS=Mesocricetus auratus OX=10036 GN=Ube2i PE=3 SV=1 | 10.926 | 15 | 2 | 12 | 2 | 158 | 18.1 | 8.32 | 35.97 | 2 | 153.7 | 162.6 | 186.8 | 176 | 190.9 | 190.7 | 154.6 | 142.7 | 200.1 | 1.021 | 0.200698311 | 0.988670244 | 0.930343519 | 0.892037303 | 0.370914641 |
| A0A1U7QD93 | ubiquitin/ISG15-conjugating enzyme E2 L6 isoform X2 OS=Mesocricetus auratus OX=10036 GN=Ube2l6 PE=3 SV=1 | 2.805 | 10 | 1 | 3 | 1 | 153 | 17.8 | 5.58 | 5.18 | 1 | 52.4 | 0 | 37.8 | 101.7 | 0 | 69.4 | 47 | 0 | 31 | 1.887 | 0.48337618 | 0.864745011 | 0.854899118 | 0.455873758 | 0.420896716 |
| A0A1U7RG31 | NEDD8-conjugating enzyme Ubc12 OS=Mesocricetus auratus OX=10036 GN=Ube2m PE=3 SV=1 | 3.709 | 10 | 2 | 2 | 2 | 183 | 20.9 | 7.69 | 2.41 | 2 | 0 | 0 | 43.7 | 0 | 0 | 40.9 | 0 | 0 | 30.9 | 0.935 | 0.964939931 | 0.707093822 | 0.823970935 | 0.755501222 | 0.855508877 |
| A0A1U7QI67 | ubiquitin-conjugating enzyme E2 N OS=Mesocricetus auratus OX=10036 GN=Ube2n PE=3 SV=1 | 13.998 | 20 | 4 | 19 | 1 | 152 | 17.2 | 6.18 | 44.17 | 4 | 847.9 | 618.8 | 830.4 | 679.7 | 532.5 | 698.8 | 773.9 | 590.7 | 689.8 | 0.861 | 0.234971272 | 0.894345044 | 0.427482181 | 1.075039246 | 0.55649838 |
| A0A1U7QB87 | nucleolar transcription factor 1 isoform X1 OS=Mesocricetus auratus OX=10036 GN=Ubtf PE=4 SV=1 | 4.936 | 3 | 2 | 4 | 2 | 764 | 89.5 | 5.81 | 13.24 | 2 | 84.2 | 78.8 | 57.6 | 80.4 | 82.4 | 52.3 | 60.8 | 60.3 | 71.8 | 0.966 | 0.892087832 | 0.874433364 | 0.382218648 | 0.89679219 | 0.536161648 |
| A0A1U7RI62 | Ubiquitin-fold modifier-conjugating enzyme 1 OS=Mesocricetus auratus OX=10036 GN=Ufc1 PE=3 SV=1 | 3.301 | 6 | 1 | 5 | 1 | 167 | 19.5 | 7.4 | 8.55 | 1 | 104 | 140.8 | 300.8 | 97.4 | 134.7 | 250 | 84.2 | 159.8 | 282.9 | 0.831 | 0.795063257 | 0.965725806 | 0.94420144 | 1.092926779 | 0.850219601 |
| A0A1U8BNU7 | UTP--glucose-1-phosphate uridylyltransferase OS=Mesocricetus auratus OX=10036 GN=Ugp2 PE=3 SV=1 | 13.51 | 10 | 5 | 15 | 5 | 497 | 55.7 | 7.33 | 37.43 | 5 | 543.6 | 623.3 | 850.7 | 438.4 | 546 | 760.3 | 515 | 579.1 | 797 | 0.809 | 0.528569045 | 0.937301745 | 0.753846687 | 1.083911274 | 0.721403424 |
| A0A1U7R961 | UDP-glucuronosyltransferase OS=Mesocricetus auratus OX=10036 GN=LOC101841486 PE=3 SV=1 | 35.964 | 14 | 5 | 37 | 3 | 533 | 60.7 | 8.66 | 149.33 | 5 | 813 | 661.2 | 1093.2 | 545.5 | 417.3 | 754.6 | 817 | 644.9 | 1095.8 | 0.671 | 0.15610203 | 0.996221859 | 0.986706855 | 1.48928613 | 0.168658541 |
| A0A1U8BJY7 | General vesicular transport factor p115 OS=Mesocricetus auratus OX=10036 GN=Uso1 PE=3 SV=1 | 18.036 | 6 | 4 | 8 | 4 | 960 | 107.1 | 5.05 | 26.11 | 4 | 170 | 0 | 0 | 137.9 | 0 | 0 | 155 | 0 | 0 | 0.811 | 0.890793723 | 0.911764706 | 0.951167016 | 1.124002901 | 0.938320429 |
| A0A1U8CR18 | Ubiquitin carboxyl-terminal hydrolase OS=Mesocricetus auratus OX=10036 GN=Usp14 PE=3 SV=1 | 21.574 | 14 | 5 | 21 | 5 | 493 | 55.9 | 5.24 | 62.44 | 5 | 251.6 | 167.9 | 243.3 | 197.8 | 133 | 192.7 | 206.9 | 127 | 182 | 0.786 | 0.245221563 | 0.778364514 | 0.241743702 | 0.98548233 | 0.93975044 |
| A0A1U7Q8G4 | Ubiquitin carboxyl-terminal hydrolase OS=Mesocricetus auratus OX=10036 GN=Usp5 PE=3 SV=1 | 25.061 | 7 | 5 | 27 | 5 | 858 | 95.7 | 5.01 | 79.33 | 5 | 159.6 | 138.6 | 0 | 130.2 | 117.3 | 0 | 134.2 | 113.7 | 0 | 0.831 | 0.808065877 | 0.831321261 | 0.81008781 | 1.001616162 | 0.998299347 |
| A0A1U8CLB1 | Ubiquitinyl hydrolase 1 OS=Mesocricetus auratus OX=10036 GN=Usp9x PE=3 SV=1 | 4.414 | 1 | 2 | 2 | 2 | 2558 | 290.7 | 5.87 | 3.15 | 2 | 30.3 | 0 | 0 | 31.9 | 0 | 0 | 28.1 | 0 | 0 | 1.05 | 0.972737053 | 0.927392739 | 0.960109246 | 0.880877743 | 0.933135544 |
| A0A1U7Q3J7 | utrophin isoform X2 OS=Mesocricetus auratus OX=10036 GN=Utrn PE=4 SV=1 | 75.928 | 8 | 23 | 73 | 23 | 3430 | 391.8 | 5.31 | 193.85 | 23 | 685.6 | 745.3 | 381.7 | 541.5 | 576.2 | 289.8 | 579.9 | 679.4 | 326 | 0.773 | 0.404637514 | 0.874600022 | 0.64879905 | 1.126323268 | 0.691459259 |
| A0A1U8CIN2 | Vesicle-associated membrane protein 3 OS=Mesocricetus auratus OX=10036 GN=Vamp3 PE=2 SV=1 | 10.115 | 41 | 3 | 9 | 3 | 104 | 11.5 | 8.5 | 19.6 | 3 | 300.9 | 261.3 | 197.8 | 228.3 | 213.3 | 170.6 | 259.3 | 261.5 | 162 | 0.816 | 0.244911869 | 0.898421053 | 0.594091686 | 1.11532179 | 0.570326302 |
| A0A1U7QCH5 | vesicle-associated membrane protein 8 OS=Mesocricetus auratus OX=10036 GN=Vamp8 PE=3 SV=1 | 2.212 | 8 | 1 | 4 | 1 | 101 | 11.4 | 8.19 | 15.18 | 1 | 254.9 | 373.6 | 225.3 | 217.2 | 292.8 | 180.9 | 172 | 322.7 | 170.1 | 0.803 | 0.392191099 | 0.778636683 | 0.406520599 | 0.962223187 | 0.893413289 |
| A0A1U7RE56 | Valyl-tRNA synthetase OS=Mesocricetus auratus OX=10036 GN=Vars PE=3 SV=1 | 14.06 | 4 | 4 | 10 | 4 | 1264 | 140 | 7.61 | 28.51 | 4 | 0 | 171.8 | 0 | 0 | 199.1 | 0 | 0 | 132.8 | 0 | 1.159 | 0.922417098 | 0.772991851 | 0.866719019 | 0.667001507 | 0.797394256 |
| A0A1U7QY73 | Metavinculin OS=Mesocricetus auratus OX=10036 GN=Vcl PE=3 SV=1 | 175.918 | 37 | 40 | 193 | 40 | 1134 | 123.7 | 5.66 | 534.05 | 40 | 4422 | 5827.5 | 5330 | 3639.8 | 4376 | 4104.6 | 4051.2 | 5135.9 | 4756.8 | 0.823 | 0.088523037 | 0.895015886 | 0.357014018 | 1.15044883 | 0.198017211 |
| A0A1U7QTL0 | 15S Mg(2+)-ATPase p97 subunit OS=Mesocricetus auratus OX=10036 GN=Vcp PE=3 SV=1 | 118.982 | 32 | 20 | 139 | 20 | 806 | 89.3 | 5.26 | 445.07 | 20 | 1793.9 | 1957.5 | 1997.9 | 1456.3 | 1619.5 | 1625.5 | 1715.7 | 1620.4 | 1679.9 | 0.827 | 0.014244745 | 0.872454038 | 0.042532329 | 1.066938932 | 0.190932441 |
| A0A1U7R165 | Outer mitochondrial membrane protein porin 2 OS=Mesocricetus auratus OX=10036 GN=Vdac2 PE=3 SV=1 | 19.35 | 23 | 6 | 41 | 6 | 295 | 31.8 | 7.74 | 79.45 | 6 | 610.1 | 667.6 | 776.6 | 809.3 | 837.9 | 896.3 | 677.3 | 784 | 824.6 | 1.327 | 0.059094084 | 1.112739133 | 0.305667174 | 0.898722233 | 0.183601277 |
| A0A1U7Q905 | Voltage-dependent anion-selective channel protein 3 OS=Mesocricetus auratus OX=10036 GN=Vdac3 PE=3 SV=1 | 26.267 | 17 | 5 | 30 | 5 | 284 | 30.8 | 8.66 | 59.68 | 5 | 0 | 81.7 | 0 | 0 | 70.3 | 0 | 0 | 60.7 | 0 | 0.861 | 0.920965262 | 0.742962056 | 0.847407966 | 0.86344239 | 0.922752617 |
| A0A1U7QL88 | Vimentin OS=Mesocricetus auratus OX=10036 GN=Vim PE=3 SV=1 | 287.742 | 71 | 47 | 461 | 40 | 466 | 53.8 | 5.12 | 1344.9 | 47 | 9655.1 | 9472.7 | 9692.4 | 8632.5 | 8561.1 | 9199.9 | 9537 | 9173 | 10083.5 | 0.904 | 0.045504198 | 0.999073566 | 0.976659066 | 1.090931479 | 0.078577205 |
| A0A1U7QLL3 | Vacuolar protein sorting-associated protein 16 homolog OS=Mesocricetus auratus OX=10036 GN=Vps16 PE=3 SV=1 | 4.021 | 2 | 1 | 1 | 1 | 839 | 94.8 | 6.98 | 4.21 | 1 | 39.4 | 0 | 0 | 27.9 | 0 | 0 | 41 | 0 | 0 | 0.709 | 0.824647601 | 1.040609137 | 0.978902023 | 1.46953405 | 0.806371348 |

**Supplementary Table 1: List of differentially expressed proteins identified across the 4dpi infected and 14dpi infected**

| Accession | Description | Sum PEP Score | Coverage [%] | # Peptides | # PSMs | # Unique Peptides | # AAs | MW [kDa] | calc. pI | Score Sequest HT: Sequest HT | # Peptides (by Search Engine): Sequest HT | Mock-infected (4dpi) Replicate_R1 | Mock-infected (4dpi) Replicate_R2 | Mock-infected (4dpi) Replicate_R3 | SARS-CoV-2-infected (4dpi) Replicate_R1 | SARS-CoV-2-infected (4dpi) Replicate_R2 | SARS-CoV-2-infected (4dpi) Replicate_R3 | SARS-CoV-2-(14dpi) Replicate_R1 | SARS-CoV-2-infected (14dpi) Replicate_R2 | SARS-CoV-2-infected (14dpi) Replicate_R3 | Abundance Ratio: SARS-CoV-2-infected (4dpi) / Mock-infected (4dpi) | P-Value: SARS-CoV-2-infected (4dpi) / Mock-infected (4dpi) | Abundance Ratio: SARS-CoV-2-infected (14dpi) / Mock-infected (4dpi) | P-Value: SARS-CoV-2-infected (14dpi) / Mock-infected (4dpi) | Abundance Ratio: SARS-CoV-2-infected (14dpi) / SARS-CoV-2-infected (4dpi) | P-Value: SARS-CoV-2-infected (14dpi) / SARS-CoV-2-infected (4dpi) |
|---|---|---|---|---|---|---|---|---|---|---|---|---|---|---|---|---|---|---|---|---|---|---|---|---|---|
| A0A1U7Q6H6 | vacuolar protein sorting-associated protein 26A OS=Mesocricetus auratus OX=10036 GN=Vps26a PE=3 SV=1 | 7.257 | 15 | 2 | 4 | 2 | 327 | 38.1 | 6.57 | 13.51 | 2 | 62.7 | 82.7 | 68.2 | 51.7 | 88.1 | 66.1 | 49.2 | 73.7 | 57 | 1.05 | 0.845601358 | 0.842228464 | 0.298959651 | 0.873725109 | 0.540548273 |
| A0A1U7R5B8 | Vacuolar protein sorting-associated protein 35 OS=Mesocricetus auratus OX=10036 GN=Vps35 PE=3 SV=1 | 22.811 | 11 | 6 | 20 | 6 | 796 | 91.6 | 5.44 | 52 | 6 | 48.4 | 0 | 0 | 23.3 | 0 | 0 | 39 | 0 | 0 | 0.481 | 0.673380266 | 0.805785124 | 0.887429756 | 1.673819742 | 0.750728332 |
| A0A1U7QY02 | vitronectin OS=Mesocricetus auratus OX=10036 GN=Vtn PE=4 SV=1 | 23.842 | 15 | 5 | 18 | 5 | 486 | 55.5 | 5.73 | 57.41 | 5 | 152.9 | 264.7 | 214.4 | 192.9 | 445.4 | 295.5 | 146.8 | 226.3 | 167.9 | 1.483 | 0.305404905 | 0.856012658 | 0.495278725 | 0.579353181 | 0.209490117 |
| A0A1U7QJG0 | WD repeat-containing protein 1 OS=Mesocricetus auratus OX=10036 GN=Wdr1 PE=4 SV=1 | 42.613 | 20 | 10 | 43 | 10 | 606 | 66.3 | 6.6 | 108.4 | 10 | 1249.1 | 1754.4 | 1048.5 | 1064.7 | 1357.7 | 1149.8 | 2409.7 | 1662.7 | 2088.9 | 0.953 | 0.538139165 | 1.520557749 | 0.080145859 | 1.724791445 | 0.04264746 |
| A0A1U8BY60 | WD repeat-containing protein 61 OS=Mesocricetus auratus OX=10036 GN=Wdr61 PE=4 SV=1 | 2.684 | 3 | 1 | 3 | 1 | 305 | 33.6 | 5.47 | 7.5 | 1 | 85 | 76.5 | 69.8 | 101.5 | 67.3 | 54.2 | 92.6 | 60.5 | 71.1 | 1.194 | 0.866166781 | 0.969303934 | 0.83569172 | 1.005381166 | 0.98247804 |
| A0A1U7QV01 | Xanthine dehydrogenase OS=Mesocricetus auratus OX=10036 GN=Xdh PE=3 SV=1 | 39.959 | 11 | 10 | 50 | 10 | 1332 | 146.1 | 7.27 | 151.69 | 10 | 528.3 | 478.5 | 396.6 | 821.5 | 678.2 | 644 | 509.6 | 493.7 | 378.3 | 1.46 | 0.024946809 | 0.984466296 | 0.903805249 | 0.644493166 | 0.023157386 |
| A0A1U7QQ50 | xaa-Pro aminopeptidase 2 OS=Mesocricetus auratus OX=10036 GN=Xpnpep2 PE=3 SV=1 | 19.385 | 8 | 3 | 13 | 3 | 674 | 76 | 6.3 | 55.56 | 3 | 59.5 | 81.3 | 56.7 | 77.9 | 116.3 | 76.7 | 61.2 | 79.6 | 51.4 | 1.43 | 0.197313194 | 0.973164557 | 0.883854211 | 0.709486896 | 0.176629596 |
| A0A1U7QPJ1 | exportin-1 OS=Mesocricetus auratus OX=10036 GN=Xpo1 PE=3 SV=1 | 8.377 | 6 | 4 | 5 | 4 | 1071 | 122.9 | 6.07 | 7.14 | 4 | 0 | 0 | 86.5 | 0 | 0 | 73.7 | 0 | 0 | 70.6 | 0.852 | 0.91587486 | 0.816184971 | 0.893907977 | 0.957937585 | 0.977226002 |
| A0A1U7QVC7 | synaptobrevin homolog YKT6 OS=Mesocricetus auratus OX=10036 GN=Ykt6 PE=3 SV=1 | 4.243 | 6 | 1 | 1 | 1 | 198 | 22.3 | 6.79 | 3.78 | 1 | 0 | 0 | 45 | 0 | 0 | 32.4 | 0 | 0 | 38.2 | 0.721 | 0.832446108 | 0.848888889 | 0.913976612 | 1.179012346 | 0.913539091 |
| A0A1U7RKI4 | LOW QUALITY PROTEIN: YTH domain-containing family protein 3 OS=Mesocricetus auratus OX=10036 GN=Ythdf3 PE=4 SV=1 | 2.497 | 2 | 1 | 1 | 1 | 587 | 64.1 | 8.9 | 2.85 | 1 | 0 | 0 | 72.4 | 0 | 0 | 65.6 | 0 | 0 | 81.2 | 0.905 | 0.947882239 | 1.121546961 | 0.939463235 | 1.237804878 | 0.8887394 |
| A0A1U7QJL9 | 14-3-3 protein epsilon isoform X1 OS=Mesocricetus auratus OX=10036 GN=Ywhae PE=3 SV=1 | 75.801 | 50 | 14 | 93 | 12 | 255 | 29.2 | 4.74 | 294.16 | 14 | 1297.5 | 1163.1 | 1130.4 | 1230.1 | 997.7 | 1114.4 | 1192.2 | 1095.9 | 1128.3 | 0.97 | 0.384797099 | 0.951378446 | 0.390103239 | 1.022200945 | 0.758852068 |
| A0A1U7R3S5 | 14-3-3 protein gamma OS=Mesocricetus auratus OX=10036 GN=Ywhag PE=3 SV=1 | 41.545 | 36 | 9 | 60 | 5 | 247 | 28.3 | 4.89 | 116.02 | 9 | 836.1 | 440.2 | 878.1 | 699 | 348 | 809.4 | 799.6 | 433.4 | 951.4 | 0.814 | 0.640601196 | 1.013924991 | 0.963910166 | 1.176686059 | 0.626115795 |
| A0A1U8CFP2 | zyxin isoform X1 OS=Mesocricetus auratus OX=10036 GN=Zyx PE=4 SV=1 | 12.666 | 10 | 4 | 11 | 4 | 574 | 61.5 | 6.47 | 28.91 | 4 | 399.4 | 137.7 | 234.9 | 270.2 | 84.7 | 153.4 | 401 | 124.9 | 246.9 | 0.631 | 0.406325869 | 1.001036269 | 0.998190567 | 1.520361991 | 0.419157564 |
| A0A1U7R5X7 | filamin-A OS=Mesocricetus auratus OX=10036 GN=LOC101842052 PE=3 SV=2 | 357.692 | 34 | 76 | 371 | 37 | 2620 | 278.6 | 6.04 | 1210.33 | 76 | 8429.1 | 9491.9 | 8012.6 | 7232 | 7911.6 | 7272.7 | 8148.2 | 8823 | 8143.5 | 0.908 | 0.099259381 | 0.968423204 | 0.619869374 | 1.12037669 | 0.046278539 |
| A0A3Q0CDD9 | plectin isoform X23 OS=Mesocricetus auratus OX=10036 GN=LOC101836251 PE=3 SV=1 | 349.498 | 22 | 90 | 408 | 90 | 4529 | 515.6 | 5.78 | 1116.74 | 90 | 9119 | 10785.2 | 10024.9 | 7522.5 | 8250.5 | 8137.4 | 7776.9 | 9488 | 8786.5 | 0.777 | 0.035851137 | 0.870437133 | 0.135140045 | 1.089542626 | 0.288090384 |
| A0A3Q0D2E6 | myosin-11 OS=Mesocricetus auratus OX=10036 GN=Myh11 PE=3 SV=1 | 266.416 | 35 | 62 | 286 | 46 | 1945 | 223.7 | 5.52 | 772.35 | 62 | 5175 | 4424.4 | 4323.5 | 3496.4 | 2971.9 | 2835 | 3826 | 3337 | 3201.1 | 0.672 | 0.012087936 | 0.744392332 | 0.02704872 | 1.114024056 | 0.270818291 |
| A0A3Q0DF64 | LOW QUALITY PROTEIN: myosin-14 OS=Mesocricetus auratus OX=10036 GN=Myh14 PE=3 SV=1 | 250.309 | 34 | 58 | 283 | 50 | 2006 | 229.1 | 5.53 | 832.55 | 58 | 4715.4 | 4053.2 | 3783 | 3296.9 | 2888.8 | 2756.3 | 3780.7 | 3309 | 3307.5 | 0.713 | 0.029183338 | 0.828356544 | 0.104757728 | 1.162737643 | 0.098898827 |
| A0A3Q0CD83 | alpha-actinin-4 OS=Mesocricetus auratus OX=10036 GN=Actn4 PE=3 SV=1 | 198.02 | 42 | 35 | 228 | 21 | 933 | 107.1 | 5.39 | 729.45 | 35 | 4012.6 | 4099.5 | 3678.8 | 4643.7 | 4755.6 | 3495.7 | 4837.4 | 5080.5 | 4523.4 | 0.953 | 0.462023683 | 1.224783519 | 0.014198286 | 1.119914696 | 0.330978926 |
| A0A3Q0D8D0 | basement membrane-specific heparan sulfate proteoglycan core protein OS=Mesocricetus auratus OX=10036 GN=Hspg2 PE=4 SV=1 | 186.468 | 16 | 38 | 213 | 28 | 3626 | 388.6 | 6.44 | 752.79 | 38 | 3811 | 3927.7 | 3945.4 | 3659.6 | 3664.4 | 3697.6 | 3404.7 | 3568.4 | 3561.6 | 0.96 | 0.027192782 | 0.901626997 | 0.005740462 | 0.955823111 | 0.087082132 |
| P02094 | Hemoglobin subunit beta OS=Mesocricetus auratus OX=10036 GN=HBB PE=1 SV=2 | 185.705 | 91 | 16 | 633 | 4 | 147 | 16 | 7.75 | 2412.22 | 16 | 6543.1 | 6777.6 | 6313 | 18152.5 | 21695.7 | 16180.2 | 6106.1 | 5242.5 | 5223.2 | 3.083 | 0.01669925 | 0.844048753 | 0.05458892 | 0.295775 | 0.012603916 |
| A0A3Q0CM76 | cytoplasmic dynein 1 heavy chain 1 OS=Mesocricetus auratus OX=10036 GN=Dync1h1 PE=3 SV=1 | 181.51 | 14 | 52 | 198 | 52 | 4645 | 532 | 6.4 | 531.54 | 52 | 3998 | 3995.3 | 3750 | 3484.9 | 3402.6 | 3211.8 | 3196.9 | 3493.8 | 3195.5 | 0.856 | 0.008968134 | 0.841858762 | 0.009375933 | 0.978899528 | 0.609619718 |
| A0A3Q0CU68 | heat shock protein HSP 90-beta OS=Mesocricetus auratus OX=10036 GN=Hsp90ab1 PE=3 SV=1 | 165.43 | 35 | 34 | 204 | 26 | 811 | 92.5 | 5.3 | 590 | 34 | 4640.2 | 4844.6 | 5767.7 | 5293.8 | 5314 | 6050.6 | 4416.4 | 4999.7 | 5650.2 | 1.097 | 0.339787586 | 0.987792165 | 0.906693653 | 0.904426596 | 0.296466964 |
| A0A1U8CT86 | selenium-binding protein 1 OS=Mesocricetus auratus OX=10036 GN=LOC106023276 PE=3 SV=2 | 161.64 | 48 | 22 | 277 | 22 | 472 | 52.4 | 6.47 | 902.71 | 22 | 4285 | 3512.5 | 3920.3 | 3115.9 | 2695.3 | 2937.6 | 4177.7 | 3478.7 | 3695.3 | 0.749 | 0.028403651 | 0.968756934 | 0.708808872 | 1.297515088 | 0.031952026 |
| A0A3Q0D994 | C3-beta-c OS=Mesocricetus auratus OX=10036 GN=C3 PE=4 SV=1 | 154.706 | 27 | 34 | 168 | 34 | 1661 | 185.7 | 6.39 | 522.58 | 34 | 2193.4 | 2238.5 | 2086 | 3986.4 | 4047.6 | 3807.3 | 2147.2 | 1924.2 | 1996 | 1.808 | 0.000112629 | 0.930882646 | 0.141991916 | 0.512393065 | 4.15384E-05 |
| A0A1U7RJY0 | Calreticulin OS=Mesocricetus auratus OX=10036 GN=LOC101838124 PE=3 SV=2 | 148.904 | 52 | 22 | 154 | 22 | 416 | 48.1 | 4.49 | 455.13 | 22 | 2040.2 | 2108.6 | 1648.4 | 2344.9 | 2366.1 | 1882.3 | 2355.8 | 2494.4 | 1948.5 | 1.142 | 0.281848685 | 1.172755813 | 0.201170625 | 1.031152837 | 0.778466501 |
| A0A3Q0CBH3 | Alpha-actinin cytoskeletal isoform OS=Mesocricetus auratus OX=10036 GN=LOC101825458 PE=3 SV=1 | 141.286 | 34 | 29 | 171 | 14 | 892 | 102.9 | 5.41 | 509.87 | 29 | 1858.2 | 1469.4 | 1644.6 | 1716 | 1445 | 1581.2 | 1787.9 | 1426.4 | 1693.3 | 0.929 | 0.608880573 | 0.987007763 | 0.896927904 | 1.034878327 | 0.702861054 |
| A0A3Q0CNB5 | SEC14-like protein 3 OS=Mesocricetus auratus OX=10036 GN=LOC106021050 PE=4 SV=1 | 138.881 | 50 | 18 | 189 | 15 | 400 | 45.9 | 5.92 | 704.2 | 18 | 1114.2 | 1034.4 | 878.8 | 982.3 | 958 | 791.9 | 1101.9 | 1148.5 | 964.2 | 0.901 | 0.34349448 | 1.061835238 | 0.521552009 | 1.176561013 | 0.120154549 |
| C0HJG9 | Annexin A2 (Fragments) OS=Mesocricetus auratus OX=10036 PE=1 SV=1 | 132.649 | 69 | 20 | 210 | 1 | 199 | 22.4 | 4.93 | 771.29 | 20 | 282.3 | 447.5 | 177.6 | 205.6 | 367.5 | 154.4 | 262.4 | 404.3 | 154.1 | 0.728 | 0.587508286 | 0.904562486 | 0.800513552 | 1.128247423 | 0.764333756 |
| A0A3Q0DDN2 | microtubule-actin cross-linking factor 1 isoform X3 OS=Mesocricetus auratus OX=10036 GN=Macf1 PE=3 SV=1 | 121.841 | 6 | 32 | 114 | 32 | 7445 | 843.2 | 5.41 | 341.43 | 32 | 794.3 | 813.2 | 807.7 | 547.3 | 618.2 | 678.5 | 701.5 | 661.2 | 657.4 | 0.76 | 0.035011778 | 0.836411063 | 0.005350317 | 1.095498915 | 0.257979758 |
| A0A1U7RB44 | elongation factor 2 OS=Mesocricetus auratus OX=10036 GN=Eef2 PE=3 SV=2 | 121.378 | 34 | 23 | 165 | 22 | 858 | 95.2 | 6.83 | 468.81 | 23 | 1575.2 | 2039.9 | 2271.3 | 1863.9 | 2324.7 | 2617.3 | 1472.1 | 1994.7 | 2254.4 | 1.182 | 0.364902154 | 0.971935309 | 0.86686956 | 0.840623577 | 0.318916318 |
| E3WI46 | ATP synthase subunit beta OS=Mesocricetus auratus OX=10036 GN=atp5b PE=2 SV=1 | 121.247 | 48 | 19 | 166 | 19 | 529 | 56.3 | 5.34 | 472.82 | 19 | 1804 | 1155.2 | 1579.2 | 1506.2 | 944 | 1237.9 | 1768.5 | 1140 | 1535.4 | 0.835 | 0.321821903 | 0.979177684 | 0.91086818 | 1.204929367 | 0.362653593 |
| A0A1U8CIQ4 | tight junction protein ZO-1 isoform X5 OS=Mesocricetus auratus OX=10036 GN=Tjp1 PE=4 SV=2 | 120.846 | 17 | 24 | 120 | 1 | 1812 | 202.3 | 7.01 | 385.79 | 24 | 2417.4 | 1806.5 | 2158.3 | 1878.5 | 1426.5 | 1645.2 | 2190.4 | 1739.5 | 2054.5 | 0.777 | 0.101740872 | 0.937670396 | 0.584348608 | 1.208920852 | 0.138629135 |
| A0A3Q0CIH4 | ezrin OS=Mesocricetus auratus OX=10036 GN=Ezr PE=4 SV=1 | 118.512 | 39 | 27 | 187 | 17 | 586 | 69.3 | 6.2 | 490.27 | 27 | 1821.8 | 2566.3 | 2425.3 | 1631.9 | 2172.3 | 2212.9 | 1436 | 2066.9 | 1959.9 | 0.887 | 0.42135261 | 0.801772977 | 0.209805936 | 0.907879211 | 0.5318631 |
| P01945 | Hemoglobin subunit alpha OS=Mesocricetus auratus OX=10036 GN=HBA PE=1 SV=1 | 114.004 | 96 | 12 | 823 | 10 | 141 | 15.2 | 8.22 | 1914.42 | 12 | 20211.6 | 19002.2 | 19195.5 | 20740.7 | 19522.5 | 20462 | 22159 | 22449 | 25643.8 | 1.066 | 0.21601017 | 1.202750247 | 0.059297034 | 1.156880504 | 0.092605903 |
| A0A3Q0D5K3 | 2-phospho-D-glycerate hydro-lyase OS=Mesocricetus auratus OX=10036 GN=Eno1 PE=3 SV=1 | 111.816 | 44 | 19 | 125 | 19 | 434 | 47.2 | 6.57 | 371.5 | 19 | 1818.3 | 2151.9 | 1879.6 | 2464.5 | 2441.5 | 2580 | 2307.4 | 2133.5 | 2482.5 | 1.135 | 0.021010891 | 1.183527642 | 0.067525247 | 0.92484638 | 0.195274073 |
| A0A3Q0CFI9 | tropomyosin alpha-4 chain isoform X2 OS=Mesocricetus auratus OX=10036 GN=Tpm4 PE=3 SV=1 | 111.599 | 53 | 23 | 180 | 9 | 229 | 26.4 | 4.79 | 401.79 | 23 | 3985.3 | 4139.1 | 4325.5 | 5599.6 | 4854.5 | 6483.8 | 4957 | 4541.7 | 5367.8 | 1.499 | 0.080694322 | 1.194105977 | 0.061462887 | 0.877706209 | 0.283088379 |
| A0A3Q0CPN1 | Pyruvate kinase OS=Mesocricetus auratus OX=10036 GN=Pkm PE=3 SV=1 | 110.959 | 46 | 22 | 107 | 20 | 552 | 60.1 | 7.49 | 314.03 | 22 | 987.5 | 1659 | 1929.4 | 1077.6 | 2783.5 | 1934 | 2595.4 | 2580.7 | 1670.8 | 1.511 | 0.522369636 | 1.496295811 | 0.142468166 | 1.181498162 | 0.583905621 |
| A0A3Q0CQS3 | tensin-1 isoform X1 OS=Mesocricetus auratus OX=10036 GN=Tns1 PE=3 SV=1 | 108.866 | 19 | 23 | 126 | 21 | 1759 | 187.8 | 7.84 | 429.46 | 23 | 2032.2 | 2009.6 | 2280.8 | 1805.5 | 1909 | 2105.4 | 1745.5 | 1622.9 | 2105.5 | 0.901 | 0.246762397 | 0.865767248 | 0.184817415 | 0.940548807 | 0.54080718 |
| A0A1U8CKN9 | Tubulin alpha chain OS=Mesocricetus auratus OX=10036 GN=LOC101831620 PE=3 SV=2 | 106.606 | 43 | 19 | 191 | 3 | 450 | 50 | 5.1 | 604.05 | 19 | 50.3 | 0 | 15.6 | 75.5 | 0 | 40.2 | 57.2 | 0 | 30.2 | 1.965 | 0.567794517 | 1.326251897 | 0.763415996 | 0.755401901 | 0.74883491 |
| A0A3Q0CPL0 | Protein disulfide-isomerase OS=Mesocricetus auratus OX=10036 GN=Pdia3 PE=3 SV=1 | 106.491 | 42 | 20 | 158 | 20 | 505 | 56.7 | 6.49 | 461.14 | 20 | 3629.1 | 3755.4 | 3790.5 | 3600.9 | 3887.2 | 4133.8 | 3850.9 | 4224 | 4254.1 | 1.035 | 0.439585605 | 1.103266219 | 0.082879514 | 1.060842031 | 0.308390088 |
| A0A3Q0CPH8 | radixin OS=Mesocricetus auratus OX=10036 GN=Rdx PE=4 SV=1 | 104.855 | 37 | 27 | 165 | 16 | 604 | 71 | 6.6 | 388.54 | 27 | 1990.4 | 1317 | 1370.3 | 1574.6 | 1112.1 | 1111.7 | 1697.1 | 1156.4 | 1096.3 | 0.791 | 0.337610556 | 0.844389337 | 0.448284196 | 1.039858888 | 0.84760748 |
| A0A3Q0CKS1 | unconventional myosin-Ic OS=Mesocricetus auratus OX=10036 GN=Myo1c PE=3 SV=1 | 101.443 | 27 | 26 | 122 | 19 | 1021 | 116.5 | 9.22 | 312.71 | 26 | 2175.2 | 2236.9 | 1999.6 | 1716.6 | 1890.7 | 1731.2 | 2052.7 | 2074 | 1873.8 | 0.806 | 0.01856078 | 0.935867243 | 0.224628557 | 1.12400487 | 0.060311149 |
| A0A3Q0CUU3 | 14-3-3 protein zeta/delta OS=Mesocricetus auratus OX=10036 GN=Ywhaz PE=3 SV=1 | 98.366 | 55 | 13 | 129 | 10 | 245 | 27.7 | 4.79 | 396.98 | 13 | 2126.7 | 2500.8 | 2221.3 | 3568 | 2600.5 | 2561.5 | 3834.9 | 4451.2 | 3098.5 | 1.212 | 0.189031752 | 1.662276603 | 0.051534052 | 1.304077892 | 0.160535141 |
| A0A3Q0CEV2 | hemoglobin subunit epsilon-Y2 OS=Mesocricetus auratus OX=10036 GN=LOC101841152 PE=3 SV=1 | 82.978 | 47 | 10 | 234 | 2 | 147 | 16 | 8.65 | 807.43 | 10 | 139 | 313.5 | 424 | 197 | 442.9 | 615.5 | 114.6 | 264.1 | 394.8 | 1.418 | 0.444779788 | 0.882487165 | 0.781821816 | 0.616138283 | 0.341185595 |
| A0A3Q0CUP9 | Spectrin beta chain OS=Mesocricetus auratus OX=10036 GN=Sptb PE=3 SV=1 | 78.96 | 11 | 21 | 70 | 19 | 2094 | 240.7 | 5.58 | 247.69 | 21 | 1280 | 893.2 | 956.9 | 1040.4 | 812.9 | 804.3 | 1080.8 | 832.1 | 749.7 | 0.841 | 0.340735544 | 0.850643749 | 0.375275328 | 1.001881397 | 0.990117744 |
| A0A3Q0CG55 | myosin-2 OS=Mesocricetus auratus OX=10036 GN=LOC101822998 PE=3 SV=1 | 74.098 | 14 | 19 | 68 | 3 | 1684 | 193.3 | 6.19 | 190.33 | 19 | 969.5 | 1063.6 | 957.9 | 2924.2 | 3431.8 | 2948.7 | 593.5 | 681.6 | 586.3 | 3.128 | 0.004691942 | 0.622333668 | 0.001190557 | 0.200049437 | 0.003451897 |
| A0A1U7RE28 | peptidyl-prolyl cis-trans isomerase A OS=Mesocricetus auratus OX=10036 GN=Ppia PE=4 SV=2 | 73.419 | 30 | 13 | 127 | 2 | 320 | 35.6 | 6.33 | 355.69 | 13 | 3542.2 | 3399.5 | 3707.3 | 2832.3 | 2594.8 | 2942.8 | 4026.5 | 3919.9 | 4431.3 | 0.794 | 0.005322258 | 1.162334492 | 0.045048867 | 1.478834873 | 0.003427087 |
| A0A3Q0DB38 | A-kinase anchor protein 2 isoform X5 OS=Mesocricetus auratus OX=10036 GN=Akap2 PE=3 SV=1 | 73.142 | 16 | 15 | 79 | 15 | 1020 | 111 | 4.96 | 206.97 | 15 | 2248.9 | 2305.7 | 1901.2 | 1702.5 | 1682.8 | 1474.7 | 2008.4 | 1917.7 | 1631.6 | 0.776 | 0.032036486 | 0.860884786 | 0.153742558 | 1.143559671 | 0.172132195 |
| A0A3Q0CVQ2 | laminin subunit alpha-4 OS=Mesocricetus auratus OX=10036 GN=Lama4 PE=4 SV=1 | 71.319 | 11 | 14 | 65 | 14 | 1817 | 201.7 | 6.23 | 251.39 | 14 | 297.4 | 323.3 | 455.3 | 258.1 | 264.9 | 405.7 | 196.9 | 230.4 | 365.9 | 0.853 | 0.513664416 | 0.737174721 | 0.25590917 | 0.854097125 | 0.557263622 |
| A0A1U7Q607 | tight junction protein ZO-2 isoform X2 OS=Mesocricetus auratus OX=10036 GN=Tjp2 PE=4 SV=2 | 69.793 | 18 | 17 | 71 | 17 | 1110 | 125.2 | 6.06 | 239.23 | 17 | 1496.2 | 1371.7 | 1323.8 | 1199.1 | 1171.3 | 1182.8 | 1171.4 | 1224.9 | 1179.9 | 0.839 | 0.050529184 | 0.853162201 | 0.046385903 | 1.006473038 | 0.706690755 |
| A0A3Q0CM78 | LOW QUALITY PROTEIN: heat shock protein HSP 90-alpha OS=Mesocricetus auratus OX=10036 GN=Hsp90aa1 PE=3 SV=1 | 68.542 | 22 | 15 | 74 | 8 | 553 | 64.8 | 5.26 | 243.3 | 15 | 1642 | 1829.6 | 1524.5 | 2078.7 | 1762.1 | 1550.3 | 1852.3 | 1743.7 | 1584.4 | 1.017 | 0.508600085 | 1.036888773 | 0.630833641 | 0.960917067 | 0.710922507 |
| A0A3Q0D9S4 | caldesmon OS=Mesocricetus auratus OX=10036 GN=Cald1 PE=4 SV=1 | 67.195 | 32 | 18 | 74 | 18 | 533 | 61.3 | 6.93 | 175.52 | 18 | 1597.5 | 1745.5 | 1621.9 | 1123 | 1343.7 | 1227.2 | 1247.7 | 1368.6 | 1272.8 | 0.757 | 0.007485989 | 0.783318899 | 0.004225119 | 1.052843878 | 0.438139623 |
| A0A3Q0DA60 | 14-3-3 protein theta OS=Mesocricetus auratus OX=10036 GN=Ywhaq PE=3 SV=1 | 66.714 | 52 | 12 | 85 | 7 | 245 | 27.7 | 4.78 | 236.15 | 12 | 380.3 | 426.2 | 1071.8 | 1087.8 | 1483.7 | 1996.7 | 3313.2 | 1802 | 2564.9 | 2.731 | 0.061732894 | 4.088856945 | 0.029366711 | 1.681209229 | 0.126593832 |
| A0A3Q0DAC7 | Nucleolin OS=Mesocricetus auratus OX=10036 GN=Ncl PE=4 SV=1 | 66.438 | 23 | 12 | 58 | 12 | 699 | 75.7 | 4.68 | 149.79 | 12 | 707.8 | 555.4 | 640.2 | 778.9 | 567.8 | 702.3 | 629.6 | 481 | 587.2 | 1.081 | 0.560404593 | 0.891982768 | 0.333920112 | 0.828599317 | 0.205024743 |
| A0A1U7RGT0 | basement membrane-specific heparan sulfate proteoglycan core protein-like OS=Mesocricetus auratus OX=10036 GN=LOC101823263 PE=4 SV=2 | 65.505 | 10 | 15 | 90 | 5 | 2010 | 213.6 | 7.2 | 279.94 | 15 | 780.8 | 1034.9 | 934.8 | 688.5 | 948.7 | 809.6 | 634.6 | 812.1 | 736.8 | 0.917 | 0.391284973 | 0.793855663 | 0.112099896 | 0.89239006 | 0.396432434 |
| A0A3Q0CMN7 | LOW QUALITY PROTEIN: myosin heavy chain, skeletal muscle, adult OS=Mesocricetus auratus OX=10036 GN=LOC101822732 PE=3 SV=1 | 64.884 | 13 | 17 | 61 | 4 | 1549 | 177.5 | 5.88 | 164.18 | 17 | 104.4 | 44.8 | 75.6 | 561.5 | 267 | 353.5 | 27.9 | 0 | 15.3 | 5.249 | 0.062515765 | 0.288256228 | 0.053857581 | 0.054822335 | 0.048043196 |
| A0A3Q0D4N8 | Calcium-activated neutral proteinase 2 OS=Mesocricetus auratus OX=10036 GN=Capn2 PE=3 SV=1 | 64.639 | 23 | 16 | 53 | 16 | 727 | 82.8 | 5.15 | 158.66 | 16 | 639.5 | 703.1 | 510.7 | 493 | 561 | 352.1 | 555.8 | 607.1 | 522.5 | 0.798 | 0.149600786 | 0.909404845 | 0.437259328 | 1.198634521 | 0.266684419 |
| D7RXW0 | Tubulin alpha chain (Fragment) OS=Mesocricetus auratus OX=10036 PE=2 SV=1 | 63.037 | 46 | 11 | 139 | 4 | 178 | 19.6 | 5.52 | 465.32 | 11 | 138.5 | 128.2 | 160.4 | 121.6 | 161 | 147.4 | 162.1 | 181.2 | 171.6 | 0.919 | 0.951672631 | 1.205572465 | 0.070668227 | 1.19744186 | 0.118213271 |
| A0A3Q0CX13 | hemoglobin subunit beta OS=Mesocricetus auratus OX=10036 GN=LOC101833678 PE=3 SV=1 | 62.593 | 44 | 8 | 232 | 1 | 147 | 16.1 | 8.09 | 831.95 | 8 | 325.3 | 301.3 | 346.1 | 311 | 293.5 | 281.5 | 436.3 | 379.5 | 425.6 | 0.846 | 0.146929337 | 1.27624139 | 0.017121739 | 1.401128668 | 0.009626996 |
| A0A3Q0D493 | Epoxide hydrolase OS=Mesocricetus auratus OX=10036 GN=Ephx1 PE=3 SV=1 | 60.522 | 40 | 15 | 78 | 15 | 455 | 52.5 | 8.37 | 238.06 | 15 | 1767.8 | 1389 | 1801.6 | 1433 | 1142.4 | 1527.1 | 1438.6 | 1255.3 | 1556.6 | 0.848 | 0.181138968 | 0.857232172 | 0.221575433 | 1.036075564 | 0.752350437 |
| A0A1U7Q957 | T1-TrpRS OS=Mesocricetus auratus OX=10036 GN=Wars PE=3 SV=2 | 59.869 | 40 | 14 | 57 | 14 | 475 | 53.5 | 6.61 | 149.82 | 14 | 385.9 | 387.7 | 361.4 | 851.4 | 773.3 | 848.8 | 520.5 | 548.7 | 559.5 | 1.994 | 0.00146687 | 1.434977974 | 0.000538656 | 0.658459673 | 0.002867643 |
| A0A3Q0D4N0 | catenin delta-1 isoform X10 OS=Mesocricetus auratus OX=10036 GN=Ctnnd1 PE=3 SV=1 | 57.496 | 16 | 13 | 60 | 13 | 956 | 106.8 | 6.77 | 187.41 | 13 | 284.5 | 227.4 | 421.2 | 214.2 | 163.5 | 359.4 | 245.8 | 202.2 | 381.4 | 0.853 | 0.471089409 | 0.888865073 | 0.683823398 | 1.125220459 | 0.719367317 |
| A0A3Q0D7D7 | actin-related protein 3 OS=Mesocricetus auratus OX=10036 GN=Actr3 PE=3 SV=1 | 54.781 | 30 | 12 | 62 | 12 | 418 | 47.3 | 5.88 | 167.66 | 12 | 714.7 | 848.3 | 794 | 705.7 | 832.3 | 732.9 | 740 | 868.6 | 861.2 | 0.923 | 0.62715747 | 1.047857446 | 0.545327773 | 1.087586419 | 0.307813073 |
| A0A3Q0CTG2 | peroxiredoxin-1 OS=Mesocricetus auratus OX=10036 GN=Prdx1 PE=3 SV=1 | 54.428 | 35 | 11 | 81 | 10 | 235 | 25.8 | 7.99 | 165.33 | 11 | 2575.6 | 2676.8 | 3015.8 | 2708.2 | 2396.1 | 2293.6 | 3454.1 | 3321.7 | 3845.9 | 0.761 | 0.187207815 | 1.284644784 | 0.019976555 | 1.435772314 | 0.006781778 |
| A0A1U8CRG0 | myosin-4 OS=Mesocricetus auratus OX=10036 GN=LOC101823448 PE=3 SV=2 | 54.334 | 12 | 17 | 55 | 1 | 1768 | 202.8 | 6.02 | 157.04 | 17 | 118.6 | 53.4 | 76.4 | 314.7 | 60.4 | 100.2 | 106.1 | 42.7 | 64.5 | 1.311 | 0.441242676 | 0.858695652 | 0.683357648 | 0.448769198 | 0.384638534 |
| A0A3Q0CY55 | lactotransferrin OS=Mesocricetus auratus OX=10036 GN=Ltf PE=3 SV=1 | 54.15 | 14 | 8 | 64 | 8 | 707 | 77.7 | 8.07 | 197.06 | 8 | 960.2 | 1250.7 | 969.7 | 1135.3 | 1741.6 | 1086.6 | 827.1 | 997.1 | 835.6 | 1.285 | 0.346825004 | 0.83625731 | 0.207283308 | 0.671073546 | 0.168581117 |

**Supplementary Table 1: List of differentially expressed proteins identified across the 4dpi infected and 14dpi infected**

| Accession | Description | Sum PEP Score | Coverage [%] | # Peptides | # PSMs | # Unique Peptides | # AAs | MW [kDa] | calc. pI | Score Sequest HT: Sequest HT | # Peptides (by Search Engine): Sequest HT | Mock-infected (4dpi) Replicate_R1 | Mock-infected (4dpi) Replicate_R2 | Mock-infected (4dpi) Replicate_R3 | SARS-CoV-2-infected (4dpi) Replicate_R1 | SARS-CoV-2-infected (4dpi) Replicate_R2 | SARS-CoV-2-infected (4dpi) Replicate_R3 | SARS-CoV-2-(14dpi) Replicate_R1 | SARS-CoV-2-infected (14dpi) Replicate_R2 | SARS-CoV-2-infected (14dpi) Replicate_R3 | Abundance Ratio: SARS-CoV-2-infected (4dpi) / Mock-infected (4dpi) | P-Value: SARS-CoV-2-infected (4dpi) / Mock-infected (4dpi) | Abundance Ratio: SARS-CoV-2-infected (14dpi) / Mock-infected (4dpi) | P-Value: SARS-CoV-2-infected (14dpi) / Mock-infected (4dpi) | Abundance Ratio: SARS-CoV-2-infected (14dpi) / SARS-CoV-2-infected (4dpi) | P-Value: SARS-CoV-2-infected (14dpi) / SARS-CoV-2-infected (4dpi) |
|---|---|---|---|---|---|---|---|---|---|---|---|---|---|---|---|---|---|---|---|---|---|---|---|---|---|---|
| A0A3Q0D153 | LIM and SH3 domain protein 1 OS=Mesocricetus auratus OX=10036 GN=Lasp1 PE=4 SV=1 | 53.599 | 55 | 11 | 54 | 11 | 262 | 29.9 | 6.98 | 153.72 | 11 | 1276 | 1111.8 | 1365.2 | 1480.1 | 1714.8 | 1418.8 | 1576.7 | 1965.8 | 1556.1 | 1.186 | 0.072321566 | 1.358539835 | 0.057361037 | 1.105100028 | 0.37923185 |
| A0A3Q0CII5 | myoferlin isoform X1 OS=Mesocricetus auratus OX=10036 GN=Myof PE=3 SV=1 | 53.589 | 10 | 13 | 38 | 13 | 2080 | 236.5 | 6.2 | 117.31 | 13 | 70 | 41.8 | 411.2 | 60.2 | 35.4 | 327.2 | 58.2 | 29.9 | 326.5 | 0.86 | 0.836402701 | 0.792734226 | 0.82403649 | 0.980605487 | 0.984573571 |
| A0A3Q0CJZ8 | LIM domain only protein 7 isoform X5 OS=Mesocricetus auratus OX=10036 GN=Lmo7 PE=4 SV=1 | 51.431 | 10 | 15 | 58 | 15 | 1748 | 198.1 | 6.25 | 122.91 | 15 | 1583 | 1094.2 | 806 | 1300 | 855 | 718.1 | 1537.9 | 1267.6 | 808.3 | 0.871 | 0.519771403 | 1.037494258 | 0.895499663 | 1.257805158 | 0.423369674 |
| Q8VD96 | Uteroglobin OS=Mesocricetus auratus OX=10036 GN=SCGB1A1 PE=3 SV=1 | 51.385 | 46 | 6 | 93 | 6 | 96 | 10.5 | 4.96 | 317.76 | 6 | 3208.8 | 3034.5 | 3146.6 | 957 | 882.4 | 970.7 | 2891.1 | 2639.4 | 2931.2 | 0.291 | 3.36348E-05 | 0.901149107 | 0.056409536 | 3.01117398 | 0.001127722 |
| A0A3Q0CNL7 | Chloride intracellular channel protein OS=Mesocricetus auratus OX=10036 GN=Clic5 PE=3 SV=1 | 50.705 | 25 | 5 | 71 | 5 | 251 | 28.3 | 6.27 | 318.54 | 5 | 494.6 | 501.4 | 634.8 | 311.7 | 277.1 | 377.7 | 527.5 | 498.5 | 699.7 | 0.63 | 0.020684131 | 1.058192298 | 0.706386679 | 1.785514744 | 0.03887085 |
| A0A3Q0D0I4 | pulmonary surfactant-associated protein B isoform X3 OS=Mesocricetus auratus OX=10036 GN=Sftpb PE=4 SV=1 | 50.507 | 20 | 8 | 73 | 8 | 358 | 39.9 | 6.58 | 210.31 | 8 | 1425.8 | 1229.4 | 1428.2 | 1035.6 | 946.5 | 1092.7 | 1663 | 1575.3 | 1709.2 | 0.77 | 0.01780546 | 1.211612872 | 0.028517693 | 1.609047743 | 0.00043388 |
| A0A3Q0DGT5 | proteasome activator complex subunit 1 OS=Mesocricetus auratus OX=10036 GN=Psme1 PE=3 SV=1 | 49.233 | 30 | 9 | 47 | 9 | 308 | 34.7 | 6.71 | 139.46 | 9 | 1249.5 | 895 | 299.8 | 1247.6 | 975.8 | 498.8 | 1140.3 | 875.6 | 381.5 | 1.095 | 0.806661552 | 0.980812503 | 0.967103619 | 0.88068474 | 0.746069331 |
| A0A3Q0DCP7 | Integrin beta OS=Mesocricetus auratus OX=10036 GN=Itgb1 PE=3 SV=1 | 49.23 | 20 | 10 | 60 | 10 | 658 | 72.5 | 6.55 | 208.44 | 10 | 353 | 169.8 | 389.1 | 290.6 | 124.5 | 321.4 | 334.5 | 162.4 | 342.2 | 0.823 | 0.557403392 | 0.920166685 | 0.800491849 | 1.139307536 | 0.707253998 |
| A0A1U7RF88 | Hemopexin OS=Mesocricetus auratus OX=10036 GN=Hpx PE=3 SV=2 | 49.043 | 31 | 13 | 61 | 13 | 461 | 51.2 | 8.06 | 154.12 | 13 | 514 | 719 | 678.1 | 1214.3 | 2051.6 | 2558.4 | 376.8 | 634.6 | 461.2 | 3.096 | 0.076136168 | 0.770550992 | 0.21408166 | 0.252837251 | 0.061038654 |
| A0A3Q0CRA4 | tropomodulin-3 OS=Mesocricetus auratus OX=10036 GN=Tmod3 PE=4 SV=1 | 48.939 | 32 | 9 | 42 | 9 | 352 | 39.6 | 5.08 | 144.54 | 9 | 99.1 | 170.8 | 40 | 111.7 | 159.3 | 39.5 | 85.5 | 171.4 | 41.4 | 1.053 | 0.997083462 | 0.962568571 | 0.946085657 | 0.960708535 | 0.941076456 |
| A0A3Q0CLE6 | Transaldolase OS=Mesocricetus auratus OX=10036 GN=Taldo1 PE=3 SV=1 | 48.767 | 26 | 10 | 81 | 10 | 337 | 37.5 | 6.71 | 212.23 | 10 | 1990.9 | 1583.3 | 1962.8 | 2527.7 | 2083.7 | 2452.1 | 1865 | 1481.7 | 1838.1 | 1.249 | 0.055411303 | 0.936391548 | 0.550751168 | 0.73402704 | 0.027922746 |
| A0A3Q0CJ34 | Phosphoglycerate kinase OS=Mesocricetus auratus OX=10036 GN=LOC110340293 PE=3 SV=1 | 48.648 | 32 | 12 | 61 | 10 | 417 | 44.5 | 6.99 | 153.84 | 12 | 2020.8 | 1234.7 | 1312 | 2234.8 | 1481.2 | 1901.3 | 2034.8 | 1361.8 | 1576.3 | 1.207 | 0.352135507 | 1.088757526 | 0.694989324 | 0.885282965 | 0.506985631 |
| A0A3Q0CVB3 | Fructose-bisphosphate aldolase OS=Mesocricetus auratus OX=10036 GN=Aldoa PE=3 SV=1 | 46.42 | 22 | 11 | 66 | 11 | 418 | 45.1 | 8.24 | 163.77 | 11 | 2184 | 2721.8 | 2371.3 | 2653.4 | 2857.7 | 2334.3 | 2093.6 | 2284.3 | 1954.3 | 1.036 | 0.436224708 | 0.87015432 | 0.177799595 | 0.807122645 | 0.0592196 |
| A0A3Q0CF45 | NADH-cytochrome b5 reductase OS=Mesocricetus auratus OX=10036 GN=LOC101842045 PE=3 SV=1 | 46.369 | 41 | 7 | 62 | 7 | 278 | 31.5 | 8.15 | 209.68 | 7 | 1629.3 | 1633.6 | 1577.1 | 1097.8 | 1190.3 | 1168.5 | 1670.1 | 1576 | 1640 | 0.729 | 0.000390593 | 1.009524793 | 0.670784193 | 1.413556674 | 0.000267144 |
| A0A3Q0D272 | ras-related protein Rap-1A OS=Mesocricetus auratus OX=10036 GN=Rap1a PE=3 SV=1 | 46.229 | 38 | 8 | 57 | 2 | 184 | 21 | 6.67 | 154.92 | 8 | 67.6 | 54.6 | 0 | 60.2 | 66.4 | 0 | 63.5 | 89.5 | 0 | 1.04 | 0.962882022 | 1.252045827 | 0.776665752 | 1.208530806 | 0.809069059 |
| A0A3Q0D9A3 | F-actin-capping protein subunit beta OS=Mesocricetus auratus OX=10036 GN=Capzb PE=3 SV=1 | 46.074 | 34 | 8 | 55 | 8 | 301 | 33.7 | 6.43 | 162.93 | 8 | 1259.2 | 1226.3 | 1487 | 1033.9 | 867.1 | 1161.6 | 1229.7 | 1248.8 | 1564.7 | 0.821 | 0.062443066 | 1.017797357 | 0.871518359 | 1.320185463 | 0.080730774 |
| A0A1U8CJA8 | nidogen-2 OS=Mesocricetus auratus OX=10036 GN=Nid2 PE=4 SV=2 | 45.223 | 9 | 9 | 31 | 9 | 1389 | 151.6 | 5.47 | 102.25 | 9 | 602.8 | 481.4 | 430.3 | 508.9 | 345 | 312.6 | 536.8 | 436 | 405.4 | 0.726 | 0.219934954 | 0.910003301 | 0.523801093 | 1.181483069 | 0.394117757 |
| A0A3Q0D147 | heterogeneous nuclear ribonucleoprotein A3 OS=Mesocricetus auratus OX=10036 GN=Hnrnpa3 PE=4 SV=1 | 44.968 | 22 | 11 | 49 | 10 | 379 | 39.6 | 9.01 | 122.3 | 11 | 1043.1 | 1408.2 | 908.5 | 988.4 | 1447.1 | 1372.2 | 1054.6 | 1230.9 | 1319.7 | 1.028 | 0.508970709 | 1.073040062 | 0.660240555 | 0.946818289 | 0.70412469 |
| A0A3Q0CL86 | Histone H2A OS=Mesocricetus auratus OX=10036 GN=LOC110340248 PE=3 SV=1 | 43.78 | 46 | 8 | 126 | 1 | 130 | 14.1 | 11.05 | 288.63 | 8 | 1753.7 | 761.4 | 767.5 | 1561.9 | 1699.3 | 658.9 | 1583.5 | 1440.5 | 516.6 | 0.859 | 0.67069665 | 1.078596235 | 0.863604852 | 0.903191245 | 0.799980835 |
| A0A3Q0D3Q7 | rho GTPase-activating protein 1 OS=Mesocricetus auratus OX=10036 GN=Arhgap1 PE=4 SV=1 | 43.24 | 19 | 7 | 57 | 7 | 439 | 50.5 | 6.48 | 169.32 | 7 | 306.7 | 276.4 | 613.2 | 239.1 | 240.4 | 443.9 | 343.3 | 338.9 | 640.2 | 0.842 | 0.521052098 | 1.105408342 | 0.788746758 | 1.432098765 | 0.339944917 |
| P51640 | Glyceraldehyde-3-phosphate dehydrogenase (Fragment) OS=Mesocricetus auratus OX=10036 GN=GAPDH PE=2 SV=1 | 42.925 | 25 | 9 | 65 | 4 | 312 | 33.5 | 8.85 | 168.46 | 9 | 1265.9 | 1144.9 | 1153.5 | 1229.5 | 1118.4 | 1063.6 | 1489 | 2242 | 1395.5 | 0.977 | 0.462654616 | 1.438290828 | 0.189183187 | 1.502711417 | 0.162700128 |
| A0A3Q0CEQ0 | LOW QUALITY PROTEIN: uncharacterized protein LOC101825376 OS=Mesocricetus auratus OX=10036 GN=LOC101825376 PE=3 SV=1 | 42.866 | 34 | 10 | 132 | 2 | 317 | 36 | 11.25 | 242.64 | 10 | 5185.3 | 7078.8 | 7645.1 | 6148.5 | 7814.4 | 6557.5 | 8663.8 | 7367.6 | 6867.9 | 0.858 | 0.832917698 | 1.150186848 | 0.343528335 | 1.115928539 | 0.340575633 |
| A0A3Q0CDV3 | Plasminogen OS=Mesocricetus auratus OX=10036 GN=Plg PE=3 SV=1 | 42.811 | 19 | 13 | 47 | 13 | 808 | 90.1 | 6.83 | 126.88 | 13 | 1197.9 | 788.9 | 547.3 | 1653.8 | 1234.8 | 801.5 | 1135.8 | 761.4 | 553.7 | 1.565 | 0.286813873 | 0.967167831 | 0.918712115 | 0.664182542 | 0.24782446 |
| A0A1U8BXQ8 | rho GDP-dissociation inhibitor 2 OS=Mesocricetus auratus OX=10036 GN=Arhgdib PE=3 SV=2 | 42.381 | 53 | 8 | 38 | 8 | 200 | 22.9 | 5.05 | 104.12 | 8 | 404.9 | 328.2 | 268.9 | 333.3 | 346.7 | 310 | 401.1 | 324.9 | 269.8 | 1.056 | 0.929807542 | 0.993812375 | 0.971705446 | 1.005858586 | 0.964906644 |
| A0A3Q0CL99 | vinexin isoform X1 OS=Mesocricetus auratus OX=10036 GN=Sorbs3 PE=4 SV=1 | 42.251 | 20 | 9 | 52 | 9 | 735 | 82.3 | 9.23 | 180.97 | 9 | 1247.1 | 615.4 | 923.3 | 919.6 | 545 | 764.6 | 1073 | 531.8 | 818.6 | 0.857 | 0.441755692 | 0.869911695 | 0.642043205 | 1.087116454 | 0.752911079 |
| A0A3Q0DEE1 | Histone H2B OS=Mesocricetus auratus OX=10036 GN=LOC101843256 PE=3 SV=1 | 42.234 | 59 | 10 | 127 | 4 | 126 | 13.9 | 10.32 | 318.99 | 10 | 4992.2 | 4890.3 | 3720 | 4750.1 | 5070 | 4329 | 6370.6 | 5187.6 | 4196.3 | 1.037 | 0.71894453 | 1.158206212 | 0.40090201 | 1.113463047 | 0.490870521 |
| A0A1U7R9P7 | L-lactate dehydrogenase OS=Mesocricetus auratus OX=10036 GN=LOC101835564 PE=3 SV=2 | 42.185 | 53 | 8 | 35 | 5 | 267 | 29.4 | 6.68 | 108.62 | 8 | 270.6 | 199.4 | 196.9 | 258.2 | 207.8 | 218.8 | 241.7 | 165.9 | 178.5 | 1.034 | 0.846675357 | 0.878842405 | 0.468605275 | 0.855870327 | 0.31484058 |
| A0A3Q0D5J7 | phosphatidylethanolamine-binding protein 1 OS=Mesocricetus auratus OX=10036 GN=Pebp1 PE=3 SV=1 | 41.941 | 45 | 7 | 40 | 7 | 187 | 20.9 | 7.11 | 165.96 | 7 | 478.6 | 356.1 | 547.8 | 462.5 | 295.4 | 475.3 | 451.8 | 342.8 | 557.9 | 0.967 | 0.570481407 | 0.978300181 | 0.910663165 | 1.096740188 | 0.664130386 |
| A0A3Q0D6W2 | myosin light polypeptide 6 isoform X1 OS=Mesocricetus auratus OX=10036 GN=Myl6 PE=4 SV=1 | 40.931 | 44 | 7 | 44 | 7 | 156 | 17.3 | 4.55 | 143.75 | 7 | 542.9 | 568.8 | 586.5 | 487 | 560.1 | 910.4 | 578 | 633.4 | 986.4 | 0.923 | 0.576846201 | 1.294193852 | 0.322347884 | 1.122758621 | 0.683961676 |
| A0A3Q0CPK0 | Angiotensin-converting enzyme OS=Mesocricetus auratus OX=10036 GN=LOC101824864 PE=3 SV=1 | 40.915 | 13 | 15 | 48 | 15 | 1259 | 145.7 | 7.2 | 114.06 | 15 | 715.1 | 890.4 | 810.5 | 659.5 | 871.6 | 737.2 | 672.1 | 899.2 | 852.3 | 0.922 | 0.572964841 | 1.003145695 | 0.97797776 | 1.068465371 | 0.607404519 |
| A0A1U7R9U5 | T-complex protein 1 subunit zeta OS=Mesocricetus auratus OX=10036 GN=LOC101831920 PE=3 SV=2 | 40.752 | 17 | 9 | 39 | 9 | 531 | 57.9 | 7.08 | 105.22 | 9 | 587.3 | 487.5 | 882.2 | 575 | 595.9 | 925.3 | 515.5 | 437.9 | 847.9 | 1.105 | 0.791345486 | 0.920439448 | 0.778895197 | 0.859316859 | 0.593058739 |
| A0A3Q0CCM9 | Heterogeneous nuclear ribonucleoproteins A2/B1 OS=Mesocricetus auratus OX=10036 GN=Hnrnpa2b1 PE=4 SV=1 | 40.477 | 30 | 10 | 50 | 9 | 313 | 33.9 | 9.03 | 95.12 | 10 | 1371.9 | 1449.3 | 1350.7 | 1225 | 1300.5 | 1080 | 1232.6 | 1325.6 | 1290.5 | 0.893 | 0.082315597 | 0.922529303 | 0.056625421 | 1.067452503 | 0.34035776 |
| A0A3Q0CH40 | serine protease inhibitor A3N-like OS=Mesocricetus auratus OX=10036 GN=LOC110340613 PE=3 SV=1 | 40.212 | 16 | 9 | 44 | 6 | 522 | 58.1 | 6.54 | 151.03 | 9 | 368.9 | 649 | 952 | 584.5 | 795.2 | 1436 | 329.7 | 566.8 | 451.6 | 1.468 | 0.417161302 | 0.684349459 | 0.34684064 | 0.4787797 | 0.190186993 |
| A0A3Q0CM14 | Elongation factor 1-gamma OS=Mesocricetus auratus OX=10036 GN=Eef1g PE=4 SV=1 | 37.866 | 24 | 10 | 51 | 10 | 437 | 50 | 6.95 | 143.63 | 10 | 1235.3 | 1202.5 | 1097.6 | 1130.4 | 1111.3 | 1441.3 | 1299.6 | 1262.8 | 1682.8 | 0.915 | 0.701184808 | 1.200769361 | 0.214122667 | 1.152647298 | 0.339196536 |
| P86206 | 60 kDa heat shock protein, mitochondrial (Fragments) OS=Mesocricetus auratus OX=10036 GN=HSPD1 PE=1 SV=1 | 37.623 | 36 | 8 | 22 | 3 | 271 | 29 | 4.82 | 39.83 | 8 | 239.7 | 51.3 | 0 | 190.7 | 48 | 0 | 173.9 | 39.5 | 0 | 0.863 | 0.86044024 | 0.733333333 | 0.78916145 | 0.894009217 | 0.918915939 |
| A0A1U7QW31 | Calgranulin-B OS=Mesocricetus auratus OX=10036 GN=LOC101830989 PE=3 SV=2 | 37.395 | 43 | 6 | 46 | 6 | 133 | 15.1 | 8.65 | 113.44 | 6 | 606.2 | 576 | 555.7 | 1051.1 | 919.9 | 917.2 | 220.3 | 303.1 | 211.7 | 1.704 | 0.007785516 | 0.42298176 | 0.002122233 | 0.254518385 | 0.000401033 |
| A0A3Q0D5V9 | S-(hydroxymethyl)glutathione dehydrogenase OS=Mesocricetus auratus OX=10036 GN=LOC101844855 PE=3 SV=1 | 37.111 | 21 | 8 | 45 | 8 | 374 | 39.7 | 7.49 | 160.54 | 8 | 1197.2 | 1357.2 | 1239.7 | 1046.6 | 1169.6 | 922.7 | 1103.9 | 1365.1 | 1247.1 | 0.874 | 0.072538401 | 0.979441765 | 0.788122762 | 1.183886075 | 0.137777944 |
| A0A1U7R5S8 | Glutathione transferase OS=Mesocricetus auratus OX=10036 GN=LOC101822730 PE=3 SV=2 | 36.418 | 33 | 10 | 53 | 8 | 222 | 25.7 | 8.85 | 128.07 | 10 | 1889.1 | 2249.2 | 2018.6 | 1340.9 | 1506.7 | 1701.9 | 1812.2 | 2225.1 | 2489.6 | 0.67 | 0.022470016 | 1.060095178 | 0.618825349 | 1.434641169 | 0.058802429 |
| A0A3Q0CGE1 | Heat shock 70 kDa protein 4 OS=Mesocricetus auratus OX=10036 GN=Hspa4 PE=3 SV=1 | 36.408 | 15 | 10 | 44 | 10 | 840 | 93.9 | 5.19 | 89.47 | 10 | 326.2 | 233.5 | 448.2 | 288.2 | 176.4 | 387.3 | 331 | 283.4 | 488.8 | 0.838 | 0.582417702 | 1.094553031 | 0.735952599 | 1.294987675 | 0.390034973 |
| A0A3Q0D6L5 | splicing factor, proline- and glutamine-rich OS=Mesocricetus auratus OX=10036 GN=Sfpq PE=4 SV=1 | 36.227 | 14 | 7 | 35 | 6 | 703 | 75.7 | 9.44 | 104.01 | 7 | 586.5 | 571.2 | 388.5 | 581.1 | 507.4 | 387.6 | 566.3 | 564.4 | 375.8 | 0.888 | 0.797187301 | 0.97432415 | 0.889796579 | 1.020594811 | 0.910593531 |
| A0A3Q0D2U7 | 40S ribosomal protein S3a OS=Mesocricetus auratus OX=10036 GN=LOC110342775 PE=3 SV=1 | 36.023 | 36 | 10 | 48 | 10 | 264 | 29.9 | 9.73 | 134.45 | 10 | 1256.2 | 1124 | 1213.5 | 1156.9 | 1087 | 1111.8 | 1206 | 1197.7 | 1309.8 | 0.921 | 0.168377948 | 1.033336116 | 0.493960124 | 1.106624549 | 0.059723807 |
| A0A3Q0CGP8 | band 4.1-like protein 2 isoform X9 OS=Mesocricetus auratus OX=10036 GN=Epb41l2 PE=4 SV=1 | 35.63 | 15 | 10 | 36 | 10 | 1017 | 113.3 | 5.4 | 76.95 | 10 | 359.4 | 183.5 | 271.7 | 258.6 | 133.9 | 179.6 | 301.3 | 143.7 | 261.8 | 0.72 | 0.272046474 | 0.867665112 | 0.632181385 | 1.235448348 | 0.496624247 |
| A0A1U8BS70 | aldo-keto reductase family 1 member C13-like OS=Mesocricetus auratus OX=10036 GN=LOC101838853 PE=3 SV=2 | 35.309 | 29 | 7 | 45 | 7 | 323 | 36.9 | 6.54 | 147.89 | 7 | 460.8 | 322.6 | 386.6 | 435.7 | 302.1 | 411.8 | 453.1 | 325.9 | 395.8 | 1.065 | 0.911301438 | 1.004102564 | 0.977908839 | 1.021920668 | 0.886459331 |
| A0A3Q0D8P8 | laminin subunit alpha-3 OS=Mesocricetus auratus OX=10036 GN=Lama3 PE=4 SV=1 | 34.923 | 4 | 12 | 27 | 12 | 3337 | 365.5 | 6.73 | 73.31 | 12 | 275.7 | 623.6 | 442.5 | 229.5 | 658.6 | 431.9 | 254.3 | 499.6 | 350 | 1.044 | 0.965942272 | 0.82270085 | 0.558495859 | 0.836287879 | 0.647194038 |
| A0A3Q0CEZ5 | Phosphoglycerate mutase OS=Mesocricetus auratus OX=10036 GN=Pgam1 PE=3 SV=1 | 34.761 | 17 | 4 | 44 | 4 | 254 | 28.8 | 7.18 | 157.86 | 4 | 340.6 | 288.5 | 265.5 | 382.9 | 319.3 | 297.5 | 332.5 | 343 | 305.8 | 1.121 | 0.36098574 | 1.096914822 | 0.330129688 | 0.981594478 | 0.841488105 |
| A0A3Q0D2S5 | T-complex protein 1 subunit gamma OS=Mesocricetus auratus OX=10036 GN=Cct3 PE=3 SV=1 | 34.589 | 22 | 9 | 36 | 9 | 545 | 60.6 | 6.64 | 97.77 | 9 | 286.6 | 211.7 | 395.6 | 280.2 | 183.4 | 358 | 265.5 | 222.8 | 387.4 | 0.912 | 0.759460869 | 0.979639781 | 0.937512571 | 1.065847128 | 0.810959604 |
| A0A3Q0CXD0 | LIM and calponin homology domains-containing protein 1 isoform X5 OS=Mesocricetus auratus OX=10036 GN=Limch1 PE=4 SV=1 | 34.15 | 13 | 10 | 35 | 10 | 958 | 107.3 | 5.54 | 95.3 | 10 | 869.1 | 1569.7 | 1499.8 | 500.2 | 952.2 | 971.2 | 730.1 | 1197 | 1265.6 | 0.576 | 0.144533517 | 0.810617986 | 0.426983328 | 1.317337845 | 0.32426324 |
| A0A3Q0DAZ8 | 14-3-3 protein beta/alpha OS=Mesocricetus auratus OX=10036 GN=Ywhab PE=3 SV=1 | 33.709 | 31 | 7 | 60 | 2 | 208 | 23.8 | 5.26 | 151.26 | 7 | 156.9 | 43.6 | 143.2 | 131.4 | 33 | 108.4 | 163.1 | 36.5 | 142.6 | 0.757 | 0.638531106 | 0.995635729 | 0.992934583 | 1.254398827 | 0.664510534 |
| A0A3Q0CK34 | RNA helicase OS=Mesocricetus auratus OX=10036 GN=LOC101834606 PE=3 SV=1 | 33.232 | 17 | 8 | 42 | 5 | 407 | 46.3 | 8 | 97.46 | 8 | 294.5 | 251.6 | 247.5 | 298.4 | 241.2 | 242.8 | 246.1 | 263 | 257.3 | 0.986 | 0.884616901 | 0.965725806 | 0.615373295 | 0.979550102 | 0.806831266 |
| A0A3Q0CVK0 | Adenine phosphoribosyltransferase OS=Mesocricetus auratus OX=10036 GN=Aprt PE=3 SV=1 | 32.925 | 66 | 9 | 45 | 9 | 180 | 19.5 | 7.25 | 91.74 | 9 | 683.7 | 720.3 | 658.4 | 773.9 | 782.3 | 793.3 | 639.7 | 640.2 | 650.2 | 1.205 | 0.025029913 | 0.935851435 | 0.128952158 | 0.821493935 | 0.000119532 |
| A0A3Q0CS80 | Antithrombin-III OS=Mesocricetus auratus OX=10036 GN=Serpinc1 PE=3 SV=1 | 32.144 | 23 | 8 | 20 | 6 | 465 | 52.4 | 7.28 | 56.42 | 8 | 497.6 | 261.7 | 338.2 | 582.5 | 311 | 388.8 | 434.2 | 239.6 | 341.9 | 1.171 | 0.594672006 | 0.92546697 | 0.776136044 | 0.792092334 | 0.423043634 |
| A0A3Q0DA40 | periplakin OS=Mesocricetus auratus OX=10036 GN=Ppl PE=3 SV=1 | 32.027 | 9 | 12 | 24 | 12 | 1756 | 204.1 | 5.64 | 44.49 | 12 | 314.8 | 344.1 | 583.1 | 272.6 | 316.3 | 522.6 | 347 | 338.9 | 553.8 | 0.896 | 0.72401599 | 0.998148148 | 0.994797915 | 1.115339631 | 0.703294262 |
| A0A3Q0D6Q5 | transcriptional activator protein Pur-alpha OS=Mesocricetus auratus OX=10036 GN=Pura PE=3 SV=1 | 31.957 | 21 | 5 | 36 | 4 | 346 | 38.1 | 6.44 | 90.56 | 5 | 693.4 | 1026.4 | 739 | 511.1 | 667.3 | 510.7 | 625.8 | 1070.8 | 763.1 | 0.691 | 0.116788892 | 1.000366032 | 0.998663762 | 1.456219288 | 0.180585318 |
| A0A1U8BV28 | RNA helicase OS=Mesocricetus auratus OX=10036 GN=Eif4a1 PE=3 SV=2 | 31.722 | 21 | 9 | 53 | 8 | 406 | 46.1 | 5.48 | 126.46 | 9 | 1266.8 | 1045.6 | 1068.7 | 1329.2 | 1072.8 | 1142.6 | 1203.9 | 1124.8 | 1104 | 1.049 | 0.627671795 | 1.015261305 | 0.837915639 | 0.968430853 | 0.685498658 |
| A0A3Q0DBG9 | Eukaryotic translation initiation factor 3 subunit A OS=Mesocricetus auratus OX=10036 GN=Eif3a PE=3 SV=1 | 31.225 | 13 | 12 | 47 | 12 | 1043 | 125.2 | 6.98 | 97.09 | 12 | 1150 | 1706.2 | 1668.2 | 996.1 | 1498 | 1487.6 | 1046.2 | 1438.4 | 1459.9 | 0.866 | 0.500106593 | 0.871828309 | 0.440766903 | 0.990657257 | 0.956543538 |
| A0A3Q0CBC6 | ATP synthase subunit d, mitochondrial OS=Mesocricetus auratus OX=10036 GN=Atp5h PE=3 SV=1 | 31.145 | 53 | 6 | 46 | 6 | 156 | 18.2 | 5.44 | 165.68 | 6 | 654.1 | 649.2 | 607.5 | 533.1 | 528.8 | 512.1 | 534.1 | 555.8 | 594.3 | 0.843 | 0.008229251 | 0.881410927 | 0.031678778 | 1.070012706 | 0.16192876 |
| A0A1U7QWZ8 | Tyrosine-protein kinase OS=Mesocricetus auratus OX=10036 GN=Lyn PE=3 SV=2 | 30.854 | 19 | 10 | 40 | 7 | 512 | 58.6 | 7.37 | 100.3 | 10 | 694.9 | 957.1 | 595.2 | 722 | 940.1 | 642.5 | 606.4 | 805.4 | 475.4 | 1.039 | 0.898026846 | 0.839800641 | 0.453285433 | 0.818883971 | 0.34783863 |
| A0A1U8CZF4 | H-2 class I histocompatibility antigen, Q10 alpha chain-like OS=Mesocricetus auratus OX=10036 GN=LOC101823249 PE=3 SV=2 | 30.4 | 20 | 8 | 32 | 8 | 399 | 45.1 | 6.74 | 87.74 | 8 | 545.1 | 681.9 | 384.9 | 687.4 | 817.3 | 497.7 | 629.6 | 781.9 | 408.6 | 1.211 | 0.36163009 | 1.12916434 | 0.643327535 | 0.908959249 | 0.69257621 |
| A0A3Q0DAS2 | complement component receptor 1-like protein OS=Mesocricetus auratus OX=10036 GN=Cr1l PE=4 SV=1 | 29.314 | 10 | 6 | 52 | 6 | 499 | 55.1 | 8 | 111.51 | 6 | 699.8 | 845.3 | 759.6 | 583 | 711.3 | 1079 | 593.1 | 708.8 | 1153 | 1.19 | 0.894234671 | 1.065171172 | 0.799966039 | 1.034382505 | 0.910230979 |
| A0A3Q0CTU7 | alcohol dehydrogenase [NADP(+)] OS=Mesocricetus auratus OX=10036 GN=Akr1a1 PE=3 SV=1 | 29.179 | 22 | 7 | 31 | 7 | 324 | 36.5 | 7.14 | 89.19 | 7 | 289.4 | 621.6 | 558.4 | 257.8 | 470 | 476.8 | 276.6 | 635.6 | 537.1 | 0.891 | 0.522094255 | 0.986320947 | 0.966018882 | 1.203137971 | 0.566094781 |
| A0A3Q0DB92 | 75 kDa glucose-regulated protein OS=Mesocricetus auratus OX=10036 GN=Hspa9 PE=3 SV=1 | 29.138 | 8 | 6 | 35 | 6 | 679 | 73.6 | 6.07 | 55.17 | 6 | 586.1 | 592.9 | 420.9 | 480.9 | 442.4 | 349.7 | 544 | 535.6 | 403.3 | 0.821 | 0.195037484 | 0.926870429 | 0.619742791 | 1.164886096 | 0.309357193 |

**Supplementary Table 1: List of differentially expressed proteins identified across the 4dpi infected and 14dpi infected**

| Accession | Description | Sum PEP Score | Coverage [%] | # Peptides | # PSMs | # Unique Peptides | # AAs | MW [kDa] | calc. pI | Score Sequest HT: Sequest HT | # Peptides (by Search Engine): Sequest HT | Mock-infected (4dpi) Replicate_R1 | Mock-infected (4dpi) Replicate_R2 | Mock-infected (4dpi) Replicate_R3 | SARS-CoV-2-infected (4dpi) Replicate_R1 | SARS-CoV-2-infected (4dpi) Replicate_R2 | SARS-CoV-2-infected (4dpi) Replicate_R3 | SARS-CoV-2-(14dpi) Replicate_R1 | SARS-CoV-2-infected (14dpi) Replicate_R2 | SARS-CoV-2-infected (14dpi) Replicate_R3 | Abundance Ratio: SARS-CoV-2-infected (4dpi) / Mock-infected (4dpi) | P-Value: SARS-CoV-2-infected (4dpi) / Mock-infected (4dpi) | Abundance Ratio: SARS-CoV-2-infected (14dpi) / Mock-infected (4dpi) | P-Value: SARS-CoV-2-infected (14dpi) / Mock-infected (4dpi) | Abundance Ratio: SARS-CoV-2-infected (14dpi) / SARS-CoV-2-infected (4dpi) | P-Value: SARS-CoV-2-infected (14dpi) / SARS-CoV-2-infected (4dpi) |
|---|---|---|---|---|---|---|---|---|---|---|---|---|---|---|---|---|---|---|---|---|---|---|---|---|---|---|
| A0A1U7RA66 | ras-related protein Rab-1B OS=Mesocricetus auratus OX=10036 GN=LOC101839987 PE=4 SV=2 | 28.91 | 27 | 6 | 19 | 3 | 201 | 22.2 | 5.73 | 50.82 | 6 | 348.2 | 541.9 | 485.9 | 312.6 | 493.1 | 397.8 | 322.7 | 514.4 | 472 | 0.877 | 0.500493503 | 0.951380814 | 0.79865396 | 1.08774408 | 0.675767506 |
| A0A3Q0D1L2 | Seryl-tRNA synthetase OS=Mesocricetus auratus OX=10036 GN=Sars PE=3 SV=1 | 28.016 | 19 | 6 | 32 | 6 | 536 | 61.3 | 6.86 | 69.83 | 6 | 566.6 | 249.1 | 133.6 | 421.1 | 199.1 | 92.3 | 533.5 | 231.8 | 150.1 | 0.799 | 0.652846466 | 0.964289476 | 0.951428538 | 1.28477193 | 0.679255124 |
| A0A3Q0CPW9 | synaptic vesicle membrane protein VAT-1 homolog OS=Mesocricetus auratus OX=10036 GN=Vat1 PE=4 SV=1 | 27.954 | 23 | 6 | 29 | 6 | 364 | 37.5 | 6.32 | 78.91 | 6 | 90.6 | 257.8 | 123.5 | 49.7 | 194.3 | 95.5 | 88 | 247.2 | 125.4 | 0.672 | 0.544881218 | 0.976054249 | 0.959779732 | 1.356701031 | 0.564467932 |
| A0A3Q0CWH6 | glutathione S-transferase A2-like OS=Mesocricetus auratus OX=10036 GN=LOC110341562 PE=4 SV=1 | 27.742 | 40 | 8 | 70 | 5 | 137 | 15.7 | 9.28 | 148.33 | 8 | 2166.7 | 1794.1 | 1262.3 | 1153.7 | 1079.2 | 790 | 3718.2 | 3528.5 | 3300.4 | 0.602 | 0.091663559 | 2.01931803 | 0.01045985 | 3.48906679 | 0.000111788 |
| A0A3Q0DCV8 | ADP/ATP translocase 2 OS=Mesocricetus auratus OX=10036 GN=Slc25a5 PE=3 SV=1 | 27.68 | 30 | 8 | 39 | 8 | 298 | 32.9 | 9.74 | 93.61 | 8 | 1093.7 | 1099.7 | 1018 | 1094.6 | 1170.1 | 887.6 | 1137.3 | 1313.8 | 1078.2 | 0.888 | 0.841400966 | 1.098991094 | 0.270156812 | 1.119595216 | 0.319500446 |
| A0A3Q0DBB3 | nebulette OS=Mesocricetus auratus OX=10036 GN=Nebl PE=4 SV=1 | 27.564 | 6 | 6 | 24 | 6 | 970 | 111.5 | 7.97 | 80.06 | 6 | 862.6 | 906.9 | 606.2 | 247.8 | 569.3 | 433.5 | 787.4 | 739.2 | 542.3 | 0.678 | 0.046981956 | 0.870859115 | 0.444342248 | 1.654325924 | 0.08776441 |
| A0A3Q0D861 | Serine/threonine-protein phosphatase OS=Mesocricetus auratus OX=10036 GN=Ppp1cc PE=3 SV=1 | 27.549 | 20 | 6 | 45 | 6 | 337 | 38.5 | 6.13 | 152.71 | 6 | 452 | 375.1 | 443.9 | 445.7 | 433.4 | 429.5 | 422.8 | 311 | 409.3 | 1.155 | 0.661146843 | 0.899370574 | 0.382544817 | 0.873528962 | 0.256585603 |
| A0A3Q0D951 | 26S proteasome AAA-ATPase subunit RPT1 OS=Mesocricetus auratus OX=10036 GN=Psmc2 PE=3 SV=1 | 27.422 | 30 | 8 | 27 | 8 | 433 | 48.6 | 5.95 | 61.53 | 8 | 248.2 | 235.1 | 262.2 | 188.5 | 198.6 | 244.7 | 225.9 | 221.3 | 283.4 | 0.847 | 0.146835784 | 0.980013414 | 0.83393035 | 1.156378601 | 0.281954868 |
| A0A3Q0CFR4 | niban-like protein 1 OS=Mesocricetus auratus OX=10036 GN=Fam129b PE=3 SV=1 | 27.316 | 11 | 7 | 30 | 7 | 748 | 84.6 | 6.01 | 70.56 | 7 | 252.5 | 100.9 | 433.5 | 248.3 | 91.7 | 392.2 | 233.1 | 81.9 | 397.2 | 0.93 | 0.894908577 | 0.90507053 | 0.860013318 | 0.972685059 | 0.960274396 |
| A0A3Q0D6C3 | prolow-density lipoprotein receptor-related protein 1 OS=Mesocricetus auratus OX=10036 GN=Lrp1 PE=3 SV=1 | 27.28 | 3 | 9 | 22 | 8 | 4544 | 504.1 | 5.38 | 64.5 | 9 | 495.9 | 322.5 | 423.6 | 418.2 | 284.8 | 336.3 | 420.7 | 325.8 | 1294.2 | 0.883 | 0.35108031 | 1.643075684 | 0.479773196 | 1.963533147 | 0.392043461 |
| A0A3Q0CVT5 | Calcium-transporting ATPase OS=Mesocricetus auratus OX=10036 GN=Atp2b4 PE=3 SV=1 | 26.348 | 9 | 7 | 18 | 5 | 1266 | 140.2 | 6.21 | 57.37 | 7 | 198.5 | 79.9 | 141.7 | 532.5 | 86 | 135 | 551.2 | 67.1 | 136.8 | 1.077 | 0.51734056 | 1.797429184 | 0.539783799 | 1.002123424 | 0.998067433 |
| A0A3Q0CF55 | protein phosphatase 1 regulatory subunit 14A OS=Mesocricetus auratus OX=10036 GN=Ppp1r14a PE=3 SV=1 | 26.086 | 31 | 3 | 20 | 3 | 147 | 16.7 | 8.02 | 102.63 | 3 | 0 | 32.2 | 0 | 0 | 27.9 | 0 | 0 | 45.9 | 0 | 0.865 | 0.924559726 | 1.425465839 | 0.82030081 | 1.64516129 | 0.757730023 |
| A0A3Q0DH51 | Kinesin-like protein OS=Mesocricetus auratus OX=10036 GN=Kif5b PE=3 SV=1 | 26.006 | 11 | 7 | 25 | 7 | 963 | 109.5 | 6.44 | 70.92 | 7 | 250.7 | 155.5 | 105 | 211.3 | 133.1 | 105.6 | 232.5 | 155.6 | 106.7 | 0.843 | 0.722296926 | 0.967918623 | 0.92737016 | 1.099555556 | 0.773398601 |
| A0A1U7QMX9 | Inorganic diphosphatase OS=Mesocricetus auratus OX=10036 GN=Ppa1 PE=3 SV=2 | 25.999 | 33 | 7 | 45 | 7 | 289 | 32.7 | 5.59 | 118.01 | 7 | 1152.3 | 1119.9 | 1225.8 | 1572 | 1180.2 | 1361 | 1609.4 | 1277.8 | 1420.3 | 1.364 | 0.206116687 | 1.231417953 | 0.095048063 | 1.04723816 | 0.685735776 |
| A0A3Q0CUL4 | 60S ribosomal protein L4 OS=Mesocricetus auratus OX=10036 GN=Rpl4 PE=3 SV=1 | 25.936 | 9 | 4 | 35 | 4 | 418 | 46.9 | 11 | 81.94 | 4 | 355.7 | 437.8 | 239.8 | 369 | 457.2 | 231 | 370.7 | 407.7 | 235.1 | 1.037 | 0.931798363 | 0.980838092 | 0.936494322 | 0.958664397 | 0.871410253 |
| A0A3Q0CLR6 | 60S ribosomal protein L7 OS=Mesocricetus auratus OX=10036 GN=Rpl7 PE=3 SV=1 | 25.828 | 26 | 6 | 30 | 6 | 248 | 29.1 | 10.71 | 99.17 | 6 | 51.7 | 89.6 | 170.2 | 35.3 | 88.9 | 190.4 | 55.9 | 82.2 | 147 | 1.119 | 0.986542144 | 0.915248796 | 0.852472321 | 0.906230134 | 0.863577375 |
| P86231 | 2-oxoglutarate dehydrogenase, mitochondrial (Fragments) OS=Mesocricetus auratus OX=10036 GN=OGDH PE=1 SV=1 | 25.415 | 37 | 8 | 47 | 1 | 180 | 21.1 | 9.94 | 98.1 | 8 | 0 | 0 | 33.3 | 0 | 0 | 28.2 | 0 | 0 | 37.3 | 0.847 | 0.912737437 | 1.12012012 | 0.940128622 | 1.322695035 | 0.855850743 |
| A0A3Q0D1C4 | MICOS complex subunit MIC60 OS=Mesocricetus auratus OX=10036 GN=Immt PE=3 SV=1 | 25.407 | 13 | 7 | 15 | 7 | 741 | 82.2 | 6.34 | 36.16 | 7 | 76.2 | 47.8 | 103.7 | 72.1 | 30.7 | 84.3 | 51.7 | 31.9 | 88.9 | 0.708 | 0.586010497 | 0.757575758 | 0.472671324 | 0.921966863 | 0.844674986 |
| A0A1U7QSR7 | Annexin OS=Mesocricetus auratus OX=10036 GN=Anxa11 PE=3 SV=2 | 25.039 | 12 | 6 | 28 | 6 | 498 | 53.6 | 7.65 | 91.23 | 6 | 367.9 | 694 | 321.8 | 285.5 | 535.4 | 263.7 | 339.3 | 639 | 299.5 | 0.776 | 0.535110862 | 0.923466069 | 0.835012424 | 1.178130186 | 0.666248661 |
| A0A1U8BJA9 | 26S protease regulatory subunit 4 OS=Mesocricetus auratus OX=10036 GN=Psmc1 PE=3 SV=2 | 24.488 | 24 | 8 | 21 | 8 | 440 | 49.3 | 6.35 | 56.35 | 8 | 50.2 | 224.2 | 222.7 | 47.8 | 227.1 | 231.1 | 39.5 | 181.1 | 222.7 | 0.953 | 0.973395143 | 0.891772279 | 0.833755081 | 0.876086957 | 0.811530275 |
| A0A3Q0D978 | 60 kDa chaperonin OS=Mesocricetus auratus OX=10036 GN=Hspd1 PE=3 SV=1 | 24.446 | 24 | 6 | 17 | 1 | 391 | 41.8 | 8.5 | 44.08 | 6 | 35.7 | 0 | 50.9 | 26.6 | 0 | 39.2 | 28 | 0 | 45.9 | 0.757 | 0.734863247 | 0.85334873 | 0.843998355 | 1.123100304 | 0.886039057 |
| A0A3Q0C9A0 | elongation factor 1-delta OS=Mesocricetus auratus OX=10036 GN=Eef1d PE=3 SV=1 | 24.314 | 12 | 5 | 7 | 5 | 649 | 71.9 | 7.03 | 27.35 | 5 | 112 | 0 | 0 | 99.3 | 0 | 0 | 121.5 | 0 | 0 | 0.887 | 0.936515653 | 1.084821429 | 0.956929442 | 1.223564955 | 0.894589904 |
| A0A3Q0CMB2 | serine protease inhibitor A3N-like isoform X2 OS=Mesocricetus auratus OX=10036 GN=LOC101835835 PE=3 SV=1 | 24.238 | 19 | 7 | 26 | 4 | 420 | 47.5 | 8.38 | 63.75 | 7 | 190.5 | 98.1 | 284.5 | 295.1 | 128.7 | 268.7 | 121.8 | 76.9 | 233.8 | 1.042 | 0.621877018 | 0.754667597 | 0.547118342 | 0.624548736 | 0.281587411 |
| A0A3Q0CDF0 | chromobox protein homolog 3 isoform X1 OS=Mesocricetus auratus OX=10036 GN=Cbx3 PE=4 SV=1 | 24.095 | 30 | 4 | 18 | 3 | 183 | 20.8 | 5.33 | 54.7 | 4 | 218 | 162.7 | 261.6 | 254.1 | 185.3 | 268 | 214.3 | 157.3 | 302.2 | 1.165 | 0.602314224 | 1.049042504 | 0.848058168 | 0.95250212 | 0.8336335 |
| A0A3Q0CXU9 | collagen alpha-1(IV) chain OS=Mesocricetus auratus OX=10036 GN=Col4a1 PE=4 SV=1 | 24.087 | 4 | 5 | 24 | 5 | 1669 | 160.9 | 8.02 | 94.47 | 5 | 328.9 | 395.9 | 194.4 | 289.4 | 315.8 | 175.5 | 269.2 | 342.3 | 172.1 | 0.903 | 0.565711862 | 0.852480418 | 0.590044462 | 1.003714615 | 0.988934661 |
| A0A3Q0CE42 | serpin B6 OS=Mesocricetus auratus OX=10036 GN=LOC101842215 PE=3 SV=1 | 24.021 | 12 | 5 | 28 | 5 | 379 | 43 | 5.66 | 74.24 | 5 | 517 | 407.7 | 537.6 | 436.4 | 333.9 | 461 | 503.9 | 397.8 | 547.1 | 0.844 | 0.241421268 | 0.990767968 | 0.943785856 | 1.176642573 | 0.287615277 |
| A0A1U7Q9W9 | Histidine--tRNA ligase, cytoplasmic OS=Mesocricetus auratus OX=10036 GN=Hars PE=3 SV=2 | 23.986 | 8 | 5 | 27 | 5 | 509 | 57.5 | 6.07 | 87.74 | 5 | 702.4 | 597.9 | 707.7 | 633.6 | 506.5 | 669 | 712.5 | 542.9 | 644.6 | 0.904 | 0.343152221 | 0.946215139 | 0.588995896 | 1.050245979 | 0.68633252 |
| A0A3Q0CF07 | 60S ribosomal protein L10 OS=Mesocricetus auratus OX=10036 GN=Rpl10 PE=4 SV=1 | 23.124 | 34 | 5 | 34 | 5 | 214 | 24.6 | 10.08 | 81.45 | 5 | 548.2 | 1175.1 | 1197.1 | 1697.6 | 1752.8 | 1757.5 | 919.7 | 1513.9 | 1064 | 1.492 | 0.068638367 | 1.197644158 | 0.527974439 | 0.671595077 | 0.08434228 |
| A0A3Q0CNC7 | ribosome-binding protein 1 OS=Mesocricetus auratus OX=10036 GN=Rrbp1 PE=4 SV=1 | 22.881 | 8 | 7 | 21 | 7 | 993 | 110.6 | 5.8 | 37.1 | 7 | 863.5 | 500.8 | 746 | 752.4 | 448.7 | 675.7 | 887.1 | 516.9 | 697.8 | 0.871 | 0.609698236 | 0.995972137 | 0.985939653 | 1.11988491 | 0.622388514 |
| A0A1U7R6Z8 | 14-3-3 protein eta OS=Mesocricetus auratus OX=10036 GN=Ywhah PE=3 SV=2 | 22.83 | 33 | 8 | 32 | 4 | 246 | 28.2 | 4.89 | 29.43 | 8 | 176.8 | 194.4 | 105.3 | 172.7 | 177.6 | 112.4 | 153.4 | 190.2 | 96.9 | 0.977 | 0.900416671 | 0.924449108 | 0.770565422 | 0.952020748 | 0.840326442 |
| A0A3Q0DCP2 | afadin OS=Mesocricetus auratus OX=10036 GN=Afdn PE=4 SV=1 | 22.573 | 5 | 7 | 28 | 7 | 1916 | 216.4 | 6.48 | 68.27 | 7 | 700.9 | 454 | 710.5 | 510.8 | 346.8 | 587.8 | 646.4 | 462 | 670.5 | 0.738 | 0.273749389 | 0.953629248 | 0.801059408 | 1.230731977 | 0.315519594 |
| A0A3Q0CF03 | Sulfurtransferase OS=Mesocricetus auratus OX=10036 GN=Tst PE=4 SV=1 | 22.517 | 29 | 5 | 15 | 5 | 297 | 33.3 | 6.95 | 56.54 | 5 | 31.2 | 0 | 49.7 | 27.2 | 0 | 34.2 | 30.9 | 0 | 41.4 | 0.774 | 0.736139471 | 0.893695921 | 0.888106975 | 1.17752443 | 0.834076251 |
| A0A3Q0CZC0 | chromobox protein homolog 1 OS=Mesocricetus auratus OX=10036 GN=Cbx1 PE=4 SV=1 | 22.412 | 23 | 4 | 29 | 3 | 185 | 21.4 | 4.93 | 57.42 | 4 | 535.4 | 471.5 | 534.5 | 438.3 | 389.2 | 399 | 634 | 547.3 | 617.8 | 0.804 | 0.019046511 | 1.167185675 | 0.068031029 | 1.46685691 | 0.007120137 |
| A0A3Q0CSY2 | ES1 protein homolog, mitochondrial OS=Mesocricetus auratus OX=10036 GN=CUNH21orf33 PE=4 SV=1 | 22.208 | 28 | 6 | 16 | 6 | 266 | 27.8 | 8.28 | 73.32 | 6 | 80.3 | 372.5 | 222.5 | 63.7 | 318.5 | 228.7 | 59.2 | 351.2 | 161.1 | 1.028 | 0.858249475 | 0.846290538 | 0.787690042 | 0.935504993 | 0.913576959 |
| A0A3Q0CDJ7 | APC-binding protein EB1 OS=Mesocricetus auratus OX=10036 GN=Mapre1 PE=3 SV=1 | 22.143 | 26 | 7 | 29 | 7 | 268 | 30 | 5.22 | 77.51 | 7 | 741.9 | 550.8 | 783.6 | 701.1 | 506.7 | 704.2 | 713.6 | 525.6 | 813.7 | 0.92 | 0.602640153 | 0.988729952 | 0.947322473 | 1.073692469 | 0.684080959 |
| A0A1U8D291 | Calcium-activated neutral proteinase 1 OS=Mesocricetus auratus OX=10036 GN=Capn1 PE=3 SV=2 | 22.015 | 9 | 5 | 15 | 5 | 713 | 81.6 | 5.66 | 56.36 | 5 | 236.8 | 122.9 | 135 | 201.9 | 108 | 124.2 | 214.9 | 100.6 | 118.4 | 0.879 | 0.686195398 | 0.877097231 | 0.709510005 | 0.999539277 | 0.998911718 |
| A0A3Q0D8D4 | far upstream element-binding protein 2 OS=Mesocricetus auratus OX=10036 GN=Khsrp PE=4 SV=1 | 21.822 | 8 | 6 | 37 | 4 | 748 | 76.7 | 7.33 | 55.1 | 6 | 982.9 | 873.9 | 705.8 | 1035.9 | 917.8 | 771.1 | 983.4 | 837.9 | 705.3 | 1.05 | 0.65226784 | 0.985951768 | 0.921081067 | 0.927260716 | 0.583739387 |
| A0A3Q0CY20 | collagen alpha-5(VI) chain OS=Mesocricetus auratus OX=10036 GN=LOC101844826 PE=4 SV=1 | 21.614 | 3 | 7 | 28 | 7 | 2431 | 266.3 | 6.37 | 60.75 | 7 | 175.8 | 0 | 334.3 | 183.3 | 0 | 343.2 | 149.8 | 0 | 286.5 | 1.035 | 0.970384709 | 0.855322486 | 0.856222878 | 0.828679962 | 0.827667954 |
| A0A3Q0CPB3 | RNA-binding protein 25 OS=Mesocricetus auratus OX=10036 GN=Rbm25 PE=4 SV=1 | 21.572 | 6 | 5 | 15 | 5 | 918 | 108.1 | 7.71 | 39.01 | 5 | 141.8 | 0 | 0 | 142.1 | 0 | 0 | 130.5 | 0 | 0 | 1.002 | 0.998879191 | 0.920310296 | 0.956072042 | 0.918367347 | 0.954960057 |
| A0A1U7RJ54 | Purine nucleoside phosphorylase OS=Mesocricetus auratus OX=10036 GN=Pnp PE=3 SV=2 | 21.5 | 25 | 6 | 16 | 6 | 291 | 32.3 | 6.68 | 47.24 | 6 | 270.7 | 163.7 | 97.2 | 244.5 | 147.7 | 85.5 | 234.2 | 121.9 | 76.5 | 0.887 | 0.806186706 | 0.813769752 | 0.657230739 | 0.905589282 | 0.830623852 |
| A0A3Q0CQL3 | dihydropyrimidinase-related protein 3 OS=Mesocricetus auratus OX=10036 GN=Dpysl3 PE=3 SV=1 | 21.48 | 11 | 6 | 27 | 3 | 683 | 73.7 | 6.35 | 62.95 | 6 | 125.6 | 126 | 194.6 | 134.3 | 112.2 | 144.5 | 117.1 | 139.4 | 174.3 | 1.006 | 0.518484103 | 0.965486329 | 0.865858957 | 1.101790281 | 0.536229491 |
| A0A3Q0D849 | voltage-dependent calcium channel subunit alpha-2/delta-1 OS=Mesocricetus auratus OX=10036 GN=Cacna2d1 PE=3 SV=1 | 21.238 | 5 | 4 | 20 | 4 | 1110 | 125.2 | 5.38 | 51.82 | 4 | 358.9 | 237.2 | 258.2 | 235.9 | 145.5 | 172.1 | 327.4 | 199.1 | 250.1 | 0.667 | 0.102727665 | 0.909048344 | 0.650235875 | 1.403071364 | 0.187937959 |
| A0A3Q0DB32 | 15-oxoprostaglandin 13-reductase OS=Mesocricetus auratus OX=10036 GN=Ptgr1 PE=3 SV=1 | 21.092 | 21 | 6 | 27 | 6 | 329 | 35.8 | 7.72 | 77.7 | 6 | 419.9 | 570.1 | 476.1 | 367.5 | 469.2 | 410.4 | 392.8 | 507.1 | 463.2 | 0.854 | 0.248437942 | 0.929745584 | 0.568765057 | 1.093015797 | 0.43426184 |
| A0A3Q0CZW3 | choline transporter-like protein 2 OS=Mesocricetus auratus OX=10036 GN=Slc44a2 PE=3 SV=1 | 20.977 | 9 | 5 | 22 | 5 | 705 | 80.2 | 8.18 | 68.91 | 5 | 353.5 | 149 | 190.6 | 296.5 | 137.1 | 151.1 | 376.8 | 168.7 | 208.6 | 0.855 | 0.677892053 | 1.088010388 | 0.830899398 | 1.289721225 | 0.528878431 |
| A0A3Q0DEB6 | Nucleobindin-1 OS=Mesocricetus auratus OX=10036 GN=Nucb1 PE=3 SV=1 | 20.966 | 17 | 6 | 19 | 6 | 457 | 53.2 | 5.14 | 55.63 | 6 | 144 | 298.7 | 429.8 | 116.2 | 300 | 400.6 | 125.9 | 275.3 | 391.7 | 0.888 | 0.881884433 | 0.908767908 | 0.825749494 | 0.970739471 | 0.947365567 |
| A0A1U7R731 | cytochrome c1, heme protein, mitochondrial OS=Mesocricetus auratus OX=10036 GN=LOC101832091 PE=3 SV=2 | 20.911 | 17 | 4 | 18 | 4 | 326 | 35.4 | 9.16 | 47.6 | 4 | 318.1 | 381.3 | 476.3 | 298.1 | 289.2 | 358 | 288.3 | 336.8 | 477.6 | 0.752 | 0.23262938 | 0.937909331 | 0.75647297 | 1.166507987 | 0.4608315 |
| A0A3Q0DET6 | Deoxynucleoside triphosphate triphosphohydrolase SAMHD1 OS=Mesocricetus auratus OX=10036 GN=Samhd1 PE=3 SV=1 | 20.895 | 11 | 7 | 13 | 7 | 664 | 76.7 | 8.12 | 29.75 | 7 | 99.9 | 108.5 | 158.1 | 156.1 | 154.2 | 292.9 | 83.2 | 107.5 | 194.2 | 1.528 | 0.221599856 | 1.050204638 | 0.882612298 | 0.638096817 | 0.276206662 |
| A0A3Q0CK05 | protein kinase C and casein kinase substrate in neurons protein 2 OS=Mesocricetus auratus OX=10036 GN=Pacsin2 PE=3 SV=1 | 20.78 | 20 | 6 | 14 | 6 | 488 | 56 | 5.14 | 40.78 | 6 | 118.5 | 160.8 | 61.6 | 138.5 | 184.5 | 72.1 | 96.1 | 146.2 | 52.9 | 1.147 | 0.699395737 | 0.865943092 | 0.718820915 | 0.74715262 | 0.476840559 |
| A0A3Q0CQK6 | Post-proline cleaving enzyme OS=Mesocricetus auratus OX=10036 GN=Prep PE=3 SV=1 | 20.682 | 8 | 4 | 14 | 4 | 710 | 80.4 | 5.78 | 47.63 | 4 | 40.1 | 36.1 | 115.8 | 48.6 | 36 | 116.7 | 29 | 28.6 | 110.1 | 0.996 | 0.935629894 | 0.8734375 | 0.839593006 | 0.833084948 | 0.776796429 |
| A0A1U7QGY2 | receptor of activated protein C kinase 1 OS=Mesocricetus auratus OX=10036 GN=Rack1 PE=4 SV=2 | 20.596 | 16 | 5 | 26 | 5 | 317 | 35.1 | 7.69 | 76.95 | 5 | 591 | 734.5 | 647.5 | 586.4 | 712.5 | 648.2 | 572.9 | 729.6 | 634.3 | 0.998 | 0.883804696 | 0.981652306 | 0.854796242 | 0.994710082 | 0.9560243 |
| A0A3Q0D567 | Band 4.1 OS=Mesocricetus auratus OX=10036 GN=Epb41 PE=4 SV=1 | 20.583 | 6 | 5 | 20 | 5 | 1021 | 113.3 | 5.33 | 54.98 | 5 | 416.3 | 174.3 | 405 | 324.8 | 140.8 | 319.2 | 382 | 180.9 | 414.7 | 0.78 | 0.520858613 | 0.98192045 | 0.958188737 | 1.245667686 | 0.536392189 |
| A0A3Q0D5D6 | intercellular adhesion molecule 1 isoform X1 OS=Mesocricetus auratus OX=10036 GN=Icam1 PE=3 SV=1 | 20.543 | 11 | 6 | 36 | 6 | 606 | 66.1 | 5.81 | 63.37 | 6 | 683.2 | 862.8 | 849.8 | 679.2 | 836.3 | 907.1 | 723.6 | 832.3 | 912.2 | 0.969 | 0.924780255 | 1.030177811 | 0.777112528 | 1.018781474 | 0.870017675 |
| A0A3Q0CUQ5 | Alpha-1,4 glucan phosphorylase OS=Mesocricetus auratus OX=10036 GN=Pygl PE=3 SV=1 | 20.542 | 9 | 8 | 20 | 6 | 920 | 105.2 | 7.06 | 42.57 | 8 | 241.9 | 187.4 | 189.1 | 217 | 167.8 | 181.7 | 211.2 | 191.5 | 198.2 | 0.895 | 0.497399928 | 0.971701164 | 0.781161277 | 1.060723742 | 0.526262976 |
| A0A1U8BYQ6 | platelet endothelial cell adhesion molecule OS=Mesocricetus auratus OX=10036 GN=Pecam1 PE=4 SV=2 | 20.447 | 10 | 6 | 17 | 6 | 735 | 81.9 | 7.64 | 30.55 | 6 | 148.9 | 177.9 | 130.5 | 134.7 | 154 | 116.4 | 159.6 | 153 | 129.7 | 0.905 | 0.380594855 | 0.967198775 | 0.779345707 | 1.091829178 | 0.431645565 |
| A0A3Q0DC82 | hydroxyacyl-coenzyme A dehydrogenase, mitochondrial OS=Mesocricetus auratus OX=10036 GN=Hadh PE=3 SV=1 | 20.32 | 18 | 8 | 40 | 8 | 314 | 34.3 | 8.82 | 70.34 | 8 | 2254.3 | 1757.9 | 2273.8 | 1643.7 | 1299.8 | 1501.2 | 2346.4 | 1304.7 | 1867.9 | 0.66 | 0.046806394 | 0.877982819 | 0.510282487 | 1.241703602 | 0.357757145 |
| A0A1U8C350 | Peroxiredoxin-5 OS=Mesocricetus auratus OX=10036 GN=Prdx5 PE=3 SV=2 | 20.141 | 29 | 5 | 11 | 5 | 246 | 25.4 | 8 | 34.71 | 5 | 22.6 | 0 | 28.2 | 29.1 | 0 | 30.9 | 25.4 | 0 | 28.9 | 1.188 | 0.828087222 | 1.068897638 | 0.930354906 | 0.905 | 0.895204418 |
| A0A3Q0CEC6 | ankyrin-1 isoform X10 OS=Mesocricetus auratus OX=10036 GN=Ank1 PE=4 SV=1 | 19.983 | 4 | 4 | 12 | 4 | 1942 | 213.4 | 6.74 | 43.58 | 4 | 147.4 | 61.7 | 122.2 | 168.1 | 61.2 | 82.2 | 130.1 | 55 | 124.7 | 0.992 | 0.881576121 | 0.935104135 | 0.848174818 | 0.994542536 | 0.989604961 |
| A0A3Q0CXQ5 | Proteasome subunit alpha type OS=Mesocricetus auratus OX=10036 GN=Psma7 PE=3 SV=1 | 19.952 | 24 | 4 | 20 | 4 | 248 | 27.9 | 8.46 | 76.71 | 4 | 239.8 | 159.6 | 215.6 | 270.9 | 200.9 | 285.3 | 220.2 | 185.9 | 250.1 | 1.259 | 0.250925617 | 1.06699187 | 0.673538716 | 0.866728305 | 0.358304824 |
| A0A3Q0CGB1 | guanine nucleotide-binding protein G(I)/G(S)/G(T) subunit beta-1 OS=Mesocricetus auratus OX=10036 GN=LOC101843384 PE=3 SV=1 | 19.719 | 14 | 4 | 27 | 1 | 340 | 37.4 | 6 | 82.76 | 4 | 356.9 | 274.1 | 521.4 | 317.3 | 182.9 | 465.8 | 311.8 | 248.5 | 383.4 | 0.667 | 0.600652988 | 0.818899688 | 0.459695835 | 0.976915114 | 0.939948285 |
| O35533 | Carboxylic ester hydrolase OS=Mesocricetus auratus OX=10036 PE=2 SV=1 | 19.604 | 11 | 4 | 13 | 2 | 559 | 62.1 | 5.72 | 42.32 | 4 | 0 | 45.8 | 0 | 0 | 32.7 | 0 | 0 | 37.8 | 0 | 0.712 | 0.828494355 | 0.825327511 | 0.89956535 | 1.155963303 | 0.923733579 |
| A0A3Q0D6L3 | ras-related C3 botulinum toxin substrate 1 isoform X1 OS=Mesocricetus auratus OX=10036 GN=Rac1 PE=4 SV=1 | 19.53 | 29 | 6 | 38 | 2 | 211 | 23.4 | 8.78 | 64.01 | 6 | 1914.4 | 1670.1 | 1935.8 | 1661.6 | 1453.4 | 1730.7 | 1946.1 | 2739.1 | 2042.4 | 0.868 | 0.132324405 | 1.218701882 | 0.243827339 | 1.388364942 | 0.117253416 |
| A0A3Q0CS56 | Small monomeric GTPase OS=Mesocricetus auratus OX=10036 GN=Ralb PE=3 SV=1 | 19.062 | 26 | 5 | 23 | 1 | 206 | 23.3 | 6.62 | 45.21 | 5 | 40 | 94.9 | 0 | 37.1 | 72.9 | 0 | 40 | 81.1 | 0 | 0.844 | 0.823150008 | 0.897702001 | 0.904960877 | 1.100909091 | 0.912164866 |
| A0A1U8D230 | Junction plakoglobin OS=Mesocricetus auratus OX=10036 GN=Jup PE=3 SV=1 | 18.869 | 9 | 6 | 16 | 5 | 745 | 81.8 | 6.14 | 42.27 | 6 | 57.2 | 50.7 | 102 | 62.3 | 21.1 | 85.6 | 60.5 | 42.3 | 104.5 | 0.655 | 0.612587367 | 0.987613149 | 0.97352083 | 1.226627219 | 0.653862515 |
| A0A3Q0DGK2 | 40S ribosomal protein S18 OS=Mesocricetus auratus OX=10036 GN=Rps18 PE=3 SV=1 | 18.821 | 28 | 7 | 44 | 7 | 152 | 17.7 | 10.99 | 89.43 | 7 | 1446.7 | 1060.1 | 1795.1 | 1390.4 | 1051.7 | 1685.3 | 1959.8 | 2203.5 | 1469.5 | 0.961 | 0.845950623 | 1.309374927 | 0.216688882 | 1.364733246 | 0.152801157 |
| A0A3Q0CVY4 | complement factor H OS=Mesocricetus auratus OX=10036 GN=Cfh PE=4 SV=1 | 18.778 | 5 | 6 | 25 | 6 | 1186 | 134 | 6.81 | 85.47 | 6 | 578.6 | 514.5 | 247.1 | 851.8 | 703.8 | 376.9 | 456 | 383.9 | 145.2 | 1.472 | 0.323602114 | 0.735039546 | 0.440578094 | 0.509754204 | 0.145056693 |

Supplementary Table 1: List of differentially expressed proteins identified across the 4dpi infected and 14dpi infected

| Accession | Description | Sum PEP Score | Coverage [%] | # Peptides | # PSMs | # Unique Peptides | # AAs | MW [kDa] | calc. pI | Score Sequest HT: Sequest HT | # Peptides (by Search Engine): Sequest HT | Mock-infected (4dpi) Replicate_R1 | Mock-infected (4dpi) Replicate_R2 | Mock-infected (4dpi) Replicate_R3 | SARS-CoV-2-infected (4dpi) Replicate_R1 | SARS-CoV-2-infected (4dpi) Replicate_R2 | SARS-CoV-2-infected (4dpi) Replicate_R3 | SARS-CoV-2-(14dpi) Replicate_R1 | SARS-CoV-2-infected (14dpi) Replicate_R2 | SARS-CoV-2-infected (14dpi) Replicate_R3 | Abundance Ratio: SARS-CoV-2-infected (4dpi) / Mock-infected (4dpi) | P-Value: SARS-CoV-2-infected (4dpi) / Mock-infected (4dpi) | Abundance Ratio: SARS-CoV-2-infected (14dpi) / Mock-infected (4dpi) | P-Value: SARS-CoV-2-infected (14dpi) / Mock-infected (4dpi) | Abundance Ratio: SARS-CoV-2-infected (14dpi) / SARS-CoV-2-infected (4dpi) | P-Value: SARS-CoV-2-infected (14dpi) / SARS-CoV-2-infected (4dpi) |
|---|---|---|---|---|---|---|---|---|---|---|---|---|---|---|---|---|---|---|---|---|---|---|---|---|---|
| A0A3Q0DC70 | Proton-translocating NAD(P)(+) transhydrogenase OS=Mesocricetus auratus OX=10036 GN=Nnt PE=3 SV=1 | 18.729 | 7 | 6 | 19 | 6 | 1086 | 113.9 | 7.33 | 50.73 | 6 | 729.5 | 498.9 | 267.9 | 567.8 | 390 | 218.9 | 602.1 | 453.1 | 240.4 | 0.778 | 0.560719181 | 0.865869144 | 0.71444877 | 1.101045296 | 0.798778712 |
| A0A3Q0CQI8 | 116 kDa U5 small nuclear ribonucleoprotein component OS=Mesocricetus auratus OX=10036 GN=Eftud2 PE=3 SV=1 | 18.6 | 9 | 5 | 13 | 4 | 971 | 109.3 | 5 | 28.68 | 5 | 55.4 | 98.3 | 0 | 36.1 | 63.3 | 0 | 48.3 | 80.9 | 0 | 0.648 | 0.625699585 | 0.840598569 | 0.836063374 | 1.299798793 | 0.756602511 |
| A0A3Q0CET2 | cytochrome P450 2B15-like OS=Mesocricetus auratus OX=10036 GN=LOC110339371 PE=3 SV=1 | 18.433 | 36 | 5 | 22 | 3 | 111 | 13 | 8.43 | 58.18 | 5 | 0 | 0 | 114 | 0 | 0 | 121.9 | 0 | 0 | 93.1 | 1.069 | 0.964525921 | 0.816666667 | 0.894206936 | 0.763740771 | 0.860803217 |
| A0A3Q0D971 | splicing factor 3B subunit 1 OS=Mesocricetus auratus OX=10036 GN=Sf3b1 PE=3 SV=1 | 18.416 | 3 | 3 | 20 | 3 | 1304 | 145.7 | 7.09 | 72.48 | 3 | 70.6 | 46.6 | 0 | 64.6 | 43.1 | 0 | 64.6 | 28.6 | 0 | 0.92 | 0.915777704 | 0.795221843 | 0.788751547 | 0.86536676 | 0.864883649 |
| A0A3Q0DCZ1 | Tetraspanin OS=Mesocricetus auratus OX=10036 GN=Cd9 PE=3 SV=1 | 18.369 | 15 | 4 | 15 | 4 | 226 | 25.2 | 7.58 | 42.99 | 4 | 695.5 | 683.1 | 357 | 614.9 | 487.1 | 288 | 722.1 | 528.4 | 416.3 | 0.713 | 0.475297005 | 0.96035953 | 0.880143619 | 1.199136691 | 0.518639906 |
| A0A3Q0D5C6 | sialic acid synthase OS=Mesocricetus auratus OX=10036 GN=Nans PE=4 SV=1 | 18.307 | 8 | 4 | 22 | 4 | 359 | 40 | 7.06 | 50.95 | 4 | 87.4 | 156.8 | 210.6 | 71.9 | 126.4 | 152 | 86 | 140.5 | 210.9 | 0.726 | 0.467571739 | 0.961741425 | 0.914569834 | 1.248644019 | 0.543762113 |
| A0A1U8C662 | 60S ribosomal protein L13 OS=Mesocricetus auratus OX=10036 GN=Rpl13 PE=3 SV=2 | 18.162 | 21 | 5 | 21 | 5 | 211 | 24.3 | 11.59 | 51.88 | 5 | 242.8 | 467.2 | 315.9 | 188.5 | 392.9 | 229.5 | 259.3 | 443.1 | 290.5 | 0.752 | 0.474714297 | 0.967833122 | 0.905752349 | 1.224441978 | 0.512321675 |
| A0A1U7R5I7 | ceruloplasmin OS=Mesocricetus auratus OX=10036 GN=Cp PE=3 SV=2 | 18.092 | 8 | 5 | 9 | 5 | 1092 | 124.9 | 6.02 | 25.36 | 5 | 91.3 | 87.2 | 34.9 | 109.8 | 172.8 | 85.1 | 92.8 | 131.5 | 24.2 | 1.734 | 0.189641826 | 1.16447985 | 0.766788042 | 0.675822682 | 0.387130872 |
| A0A3Q0CLW2 | septin-8 isoform X1 OS=Mesocricetus auratus OX=10036 GN=Sept8 PE=3 SV=1 | 18.004 | 13 | 6 | 11 | 3 | 423 | 49.5 | 6.3 | 25.85 | 6 | 50.6 | 94.3 | 403 | 33.1 | 90.7 | 329.8 | 38.8 | 73.1 | 404.4 | 0.961 | 0.837569407 | 0.942325242 | 0.950951966 | 1.138227513 | 0.894759183 |
| A0A3Q0CSM9 | neuroblast differentiation-associated protein AHNAK OS=Mesocricetus auratus OX=10036 GN=Ahnak PE=4 SV=1 | 17.691 | 30 | 2 | 20 | 2 | 119 | 13.2 | 5.43 | 117.33 | 2 | 92.7 | 144.7 | 164.2 | 109.7 | 141.6 | 182.9 | 95 | 157.4 | 194.9 | 1.067 | 0.736114529 | 1.113794821 | 0.696710219 | 1.030170428 | 0.90989539 |
| A0A3Q0CEI8 | LOW QUALITY PROTEIN: uncharacterized protein LOC101834332 OS=Mesocricetus auratus OX=10036 GN=LOC101834332 PE=3 SV=1 | 17.439 | 5 | 5 | 29 | 3 | 929 | 105.5 | 7.09 | 61.71 | 5 | 150.6 | 347.8 | 144.1 | 209.5 | 309.5 | 470.3 | 737.4 | 532.5 | 724.7 | 1.768 | 0.317962905 | 3.104435798 | 0.008727536 | 2.016173052 | 0.030086161 |
| A0A3Q0CHF6 | Dynamin GTPase OS=Mesocricetus auratus OX=10036 GN=Dnm1 PE=3 SV=1 | 17.367 | 8 | 6 | 19 | 2 | 851 | 96 | 7.01 | 42.49 | 6 | 336.4 | 293.7 | 294.5 | 333.9 | 277.4 | 292.3 | 284.9 | 265.7 | 297.7 | 0.945 | 0.766911083 | 0.917477828 | 0.217282637 | 0.938800354 | 0.407641186 |
| A0A3Q0D389 | Protein transport protein SEC23 OS=Mesocricetus auratus OX=10036 GN=Sec23a PE=3 SV=1 | 16.959 | 7 | 4 | 16 | 4 | 765 | 86.1 | 7.08 | 47.24 | 4 | 150 | 287.8 | 180.6 | 127 | 244 | 131 | 163.7 | 260.6 | 168.6 | 0.848 | 0.531707788 | 0.958764554 | 0.879408191 | 1.181075697 | 0.575733734 |
| A0A3Q0CV25 | Aminopeptidase OS=Mesocricetus auratus OX=10036 GN=Npepps PE=3 SV=1 | 16.912 | 7 | 5 | 25 | 5 | 920 | 103.3 | 5.62 | 58.12 | 5 | 307.5 | 437.3 | 305.1 | 268.7 | 398.3 | 294.4 | 298.8 | 404.1 | 276.2 | 0.911 | 0.643370206 | 0.932565006 | 0.708994278 | 1.018410651 | 0.920997907 |
| A0A3Q0CN72 | Aldehyde dehydrogenase OS=Mesocricetus auratus OX=10036 GN=Aldh3a2 PE=3 SV=1 | 16.897 | 13 | 5 | 18 | 5 | 484 | 54 | 8.06 | 38.79 | 5 | 0 | 0 | 133.7 | 0 | 0 | 130.2 | 0 | 0 | 133.5 | 0.973 | 0.985935758 | 0.998504114 | 0.999206093 | 1.025345622 | 0.986729075 |
| A0A1U8BI71 | alpha-centractin OS=Mesocricetus auratus OX=10036 GN=Actr1a PE=3 SV=2 | 16.709 | 20 | 5 | 14 | 5 | 376 | 42.6 | 6.64 | 29.37 | 5 | 205.5 | 72 | 57.9 | 198.9 | 61.6 | 48.5 | 157.9 | 56.3 | 47.4 | 0.968 | 0.902236407 | 0.779964222 | 0.699119263 | 0.846601942 | 0.805585981 |
| A0A3Q0CMS7 | pre-mRNA-processing-splicing factor 8 OS=Mesocricetus auratus OX=10036 GN=Prpf8 PE=4 SV=1 | 16.687 | 3 | 6 | 18 | 6 | 2335 | 273.5 | 8.84 | 44.97 | 6 | 309.4 | 399.7 | 446 | 280.6 | 384.5 | 340.2 | 291.8 | 364.9 | 454.1 | 0.962 | 0.379811766 | 0.961648342 | 0.822977992 | 1.104943798 | 0.567995068 |
| A0A3Q0CYQ4 | ubiquitin-conjugating enzyme E2 variant 1 OS=Mesocricetus auratus OX=10036 GN=Ube2v1 PE=4 SV=1 | 16.649 | 33 | 4 | 23 | 4 | 147 | 16.3 | 7.96 | 75.05 | 4 | 342.9 | 335.9 | 195.3 | 330.7 | 287.5 | 182.5 | 340.2 | 308.2 | 195.9 | 0.953 | 0.726558532 | 0.965907791 | 0.886009042 | 1.054452354 | 0.826325292 |
| A0A1U7RIX6 | Proteasome subunit alpha type OS=Mesocricetus auratus OX=10036 GN=LOC101822549 PE=3 SV=2 | 16.476 | 17 | 4 | 15 | 4 | 263 | 29.5 | 6.46 | 39.31 | 4 | 417.4 | 175.1 | 200.4 | 371.1 | 146.4 | 188.1 | 356.2 | 177.2 | 199.5 | 0.939 | 0.792340256 | 0.924328415 | 0.844938526 | 1.038690476 | 0.923730877 |
| A0A3Q0CUC8 | Non-specific serine/threonine protein kinase OS=Mesocricetus auratus OX=10036 GN=Pak2 PE=4 SV=1 | 16.291 | 12 | 4 | 17 | 4 | 524 | 57.9 | 5.77 | 51.48 | 4 | 293.9 | 372.4 | 245 | 246.5 | 292.5 | 208.7 | 325.8 | 359.9 | 253.5 | 0.786 | 0.295881506 | 1.030615604 | 0.857778174 | 1.256118764 | 0.187086088 |
| A0A1U7QVD8 | drebrin-like protein OS=Mesocricetus auratus OX=10036 GN=Dbnl PE=3 SV=2 | 16.285 | 15 | 4 | 22 | 4 | 434 | 48.5 | 4.89 | 75.57 | 4 | 113.1 | 49.5 | 0 | 80.2 | 44.6 | 0 | 80.3 | 33.8 | 0 | 0.799 | 0.770809576 | 0.701722017 | 0.709992951 | 0.914262821 | 0.918804309 |
| A0A3Q0CV89 | liprin-beta-1 OS=Mesocricetus auratus OX=10036 GN=Ppfibp1 PE=3 SV=1 | 16.164 | 5 | 4 | 18 | 4 | 1032 | 116 | 5.67 | 48.78 | 4 | 56.6 | 51.6 | 85.8 | 35.6 | 46.3 | 95.8 | 49.9 | 39.8 | 85.7 | 0.898 | 0.814980294 | 0.904123711 | 0.742720096 | 0.987056837 | 0.97532614 |
| A0A3Q0D7V9 | Coatomer subunit beta' OS=Mesocricetus auratus OX=10036 GN=Copb2 PE=3 SV=1 | 16.161 | 10 | 6 | 14 | 6 | 905 | 102.4 | 5.31 | 30.95 | 6 | 331.1 | 287.4 | 235.6 | 315.4 | 245.4 | 212.1 | 296.6 | 264.8 | 233.4 | 0.952 | 0.546400364 | 0.930570191 | 0.587017209 | 1.028334843 | 0.849170353 |
| A0A3Q0CRE4 | Hexokinase OS=Mesocricetus auratus OX=10036 GN=Hk1 PE=3 SV=1 | 16.08 | 5 | 4 | 18 | 4 | 916 | 101.9 | 6.67 | 46.58 | 4 | 0 | 82.2 | 102.6 | 0 | 80.7 | 90.4 | 0 | 68.7 | 97.9 | 0.93 | 0.91960297 | 0.901515152 | 0.893987191 | 0.973699591 | 0.9724216 |
| A0A3Q0CV00 | microsomal glutathione S-transferase 1 OS=Mesocricetus auratus OX=10036 GN=Mgst1 PE=3 SV=1 | 16.076 | 12 | 3 | 12 | 3 | 155 | 17.5 | 9.42 | 35.07 | 3 | 86.8 | 142 | 110.9 | 61.4 | 99.1 | 80.6 | 67.2 | 110.1 | 81.4 | 0.708 | 0.173842222 | 0.761554313 | 0.258936841 | 1.072998756 | 0.742912293 |
| A0A1U7R6S5 | collagen alpha-2(IV) chain OS=Mesocricetus auratus OX=10036 GN=LOC101843198 PE=4 SV=2 | 15.983 | 17 | 4 | 14 | 4 | 358 | 37.4 | 8.09 | 34.87 | 4 | 127.9 | 84 | 237.4 | 106.9 | 74.6 | 190.3 | 101.3 | 91.7 | 196.7 | 0.836 | 0.676437351 | 0.86734921 | 0.744794789 | 1.048144164 | 0.90722044 |
| A0A3Q0CK07 | keratin, type II cytoskeletal 4 OS=Mesocricetus auratus OX=10036 GN=Krt4 PE=3 SV=1 | 15.872 | 7 | 5 | 42 | 1 | 543 | 58 | 7.66 | 77.65 | 5 | 82.1 | 0 | 110.8 | 73 | 0 | 92.4 | 67.9 | 0 | 88 | 0.861 | 0.843715685 | 0.808190772 | 0.787008005 | 0.942563482 | 0.938776058 |
| A0A3Q0CF98 | high mobility group protein B1-like OS=Mesocricetus auratus OX=10036 GN=LOC101834153 PE=4 SV=1 | 15.828 | 10 | 4 | 32 | 4 | 195 | 22.1 | 5.76 | 24.6 | 4 | 2409.4 | 1746.8 | 2387.4 | 1775.3 | 1299.5 | 1754.1 | 2298.8 | 1804.8 | 2506.9 | 0.744 | 0.106253162 | 1.01022373 | 0.944493949 | 1.368945308 | 0.089559473 |
| A0A1U7R5B2 | 26S proteasome non-ATPase regulatory subunit 8 OS=Mesocricetus auratus OX=10036 GN=Psmd8 PE=3 SV=2 | 15.771 | 15 | 4 | 8 | 4 | 352 | 39.6 | 9.54 | 23.68 | 4 | 51.3 | 63.5 | 55 | 41.9 | 58.9 | 48 | 37.9 | 58.3 | 47.1 | 0.928 | 0.323860069 | 0.84393404 | 0.283651987 | 0.963037634 | 0.82414922 |
| A0A3Q0CMK8 | poly(RC)-binding protein 2 OS=Mesocricetus auratus OX=10036 GN=Pcbp2 PE=4 SV=1 | 15.608 | 20 | 5 | 11 | 3 | 362 | 38.2 | 6.79 | 29.14 | 5 | 0 | 27.3 | 0 | 0 | 36.8 | 0 | 0 | 29.7 | 0 | 1.348 | 0.846685649 | 1.087912088 | 0.955432208 | 0.807065217 | 0.888229564 |
| A0A3Q0CRY9 | Succinate--CoA ligase [GDP-forming] subunit beta, mitochondrial OS=Mesocricetus auratus OX=10036 GN=Suclg2 PE=3 SV=1 | 15.554 | 13 | 5 | 21 | 5 | 434 | 47.1 | 7.74 | 35.96 | 5 | 366.2 | 314.7 | 192.7 | 342.1 | 228.7 | 202.1 | 330.3 | 279.8 | 174.2 | 0.934 | 0.643493556 | 0.897779304 | 0.68868867 | 1.014749644 | 0.954743977 |
| A0A3Q0D3Y9 | Threonyl-tRNA synthetase OS=Mesocricetus auratus OX=10036 GN=Tars PE=3 SV=1 | 15.403 | 8 | 5 | 10 | 5 | 681 | 78.5 | 7.47 | 27.23 | 5 | 97.7 | 88.7 | 35.4 | 105.8 | 105.1 | 48.9 | 118.5 | 75.5 | 38.6 | 1.185 | 0.664330621 | 1.048692516 | 0.910966871 | 0.89530408 | 0.776726899 |
| A0A3Q0CHK6 | Complex I-49kD OS=Mesocricetus auratus OX=10036 GN=Ndufs2 PE=3 SV=1 | 15.384 | 14 | 4 | 11 | 4 | 463 | 52.5 | 7.33 | 29.89 | 4 | 0 | 86.4 | 0 | 0 | 53.5 | 0 | 0 | 94.3 | 0 | 0.619 | 0.765376087 | 1.091435185 | 0.953731636 | 1.762616822 | 0.730491317 |
| A0A3Q0CQG7 | Actin-related protein 2 OS=Mesocricetus auratus OX=10036 GN=Actr2 PE=3 SV=1 | 15.267 | 13 | 5 | 19 | 5 | 427 | 48.7 | 6.74 | 39 | 5 | 452.2 | 611.4 | 329.7 | 420.6 | 545.2 | 308.8 | 350.2 | 449.7 | 308.3 | 0.936 | 0.729304038 | 0.79537788 | 0.376502034 | 0.869449239 | 0.534094479 |
| A0A3Q0D6H8 | Aspartate--tRNA ligase, cytoplasmic OS=Mesocricetus auratus OX=10036 GN=Dars PE=3 SV=1 | 15.002 | 11 | 4 | 15 | 4 | 501 | 57 | 6.65 | 54.79 | 4 | 229.7 | 206.1 | 194.4 | 193.8 | 155.4 | 152.1 | 195 | 154.4 | 173.9 | 0.783 | 0.068014804 | 0.830371311 | 0.086176737 | 1.043885897 | 0.701697931 |
| A0A1U7R965 | ras-related protein Rab-7a OS=Mesocricetus auratus OX=10036 GN=LOC101839102 PE=4 SV=2 | 14.862 | 12 | 2 | 7 | 2 | 207 | 23.5 | 6.7 | 27.74 | 2 | 70 | 82.5 | 61.8 | 69.4 | 63.8 | 52.4 | 43.6 | 66.9 | 47.1 | 0.858 | 0.290701066 | 0.735417639 | 0.117830634 | 0.849137931 | 0.356152189 |
| A0A3Q0CFS5 | Adenylate kinase isoenzyme 1 OS=Mesocricetus auratus OX=10036 GN=Ak1 PE=3 SV=1 | 14.74 | 16 | 3 | 26 | 3 | 210 | 23.4 | 7.05 | 62.06 | 3 | 662.2 | 711.1 | 899.4 | 498.3 | 569.7 | 673.7 | 559.5 | 613.3 | 831.3 | 0.752 | 0.123908101 | 0.881814582 | 0.462748839 | 1.150657404 | 0.429819519 |
| A0A3Q0CG49 | cAMP-dependent protein kinase catalytic subunit alpha OS=Mesocricetus auratus OX=10036 GN=Prkaca PE=3 SV=1 | 14.697 | 10 | 4 | 24 | 3 | 351 | 40.6 | 8.79 | 62.36 | 4 | 910.2 | 923.2 | 1214.9 | 603.2 | 563.6 | 819.1 | 918.5 | 904.2 | 1270.1 | 0.663 | 0.052450967 | 1.014598301 | 0.928783321 | 1.557379526 | 0.071249084 |
| A0A3Q0D7A4 | integrin alpha-1 OS=Mesocricetus auratus OX=10036 GN=Itga1 PE=3 SV=1 | 14.625 | 3 | 4 | 10 | 4 | 1179 | 131 | 6.14 | 25.48 | 4 | 73.9 | 148.5 | 97.4 | 62.3 | 112.7 | 85.2 | 67.7 | 119.1 | 89.4 | 0.813 | 0.499611154 | 0.86366479 | 0.617463617 | 1.061491161 | 0.810621529 |
| A0A3Q0D509 | 60S acidic ribosomal protein P0 OS=Mesocricetus auratus OX=10036 GN=Rplp0 PE=3 SV=1 | 14.573 | 7 | 2 | 25 | 2 | 318 | 34.3 | 6.25 | 89.45 | 2 | 195.1 | 158.3 | 220.9 | 196.3 | 171 | 248.9 | 174.1 | 135.8 | 221 | 1.006 | 0.659309156 | 0.924429741 | 0.663140663 | 0.861570919 | 0.446125083 |
| A0A1U7QK94 | Cytochrome c oxidase polypeptide Va OS=Mesocricetus auratus OX=10036 GN=LOC101843749 PE=3 SV=2 | 14.396 | 27 | 4 | 20 | 4 | 146 | 16.2 | 6.54 | 41.96 | 4 | 554.7 | 885.6 | 1000.2 | 609.1 | 890.1 | 1049.6 | 513.7 | 881.5 | 1014.5 | 1.035 | 0.855213335 | 0.987379635 | 0.961680648 | 0.945425298 | 0.826174846 |
| A0A3Q0CR59 | Unconventional myosin-6 OS=Mesocricetus auratus OX=10036 GN=Myo6 PE=3 SV=1 | 14.316 | 5 | 6 | 11 | 6 | 1294 | 149.2 | 8.41 | 15.65 | 6 | 447.5 | 173.3 | 115 | 338.2 | 155.8 | 103.1 | 350.1 | 168.1 | 111.8 | 0.823 | 0.732026684 | 0.856210927 | 0.793712215 | 1.055099648 | 0.918935731 |
| A0A3Q0CPM0 | stress-induced-phosphoprotein 1 isoform X1 OS=Mesocricetus auratus OX=10036 GN=Stip1 PE=4 SV=1 | 14.242 | 8 | 4 | 9 | 4 | 547 | 62.9 | 6.98 | 26.35 | 4 | 69.9 | 53.8 | 0 | 64.8 | 55.6 | 0 | 44.6 | 32.1 | 0 | 0.98 | 0.97181969 | 0.620048504 | 0.570261178 | 0.637043189 | 0.584638507 |
| A0A3Q0CT51 | 26S proteasome non-ATPase regulatory subunit 2 OS=Mesocricetus auratus OX=10036 GN=Psmd2 PE=3 SV=1 | 14.16 | 7 | 4 | 10 | 4 | 908 | 100.1 | 5.17 | 31.28 | 4 | 0 | 0 | 229.4 | 0 | 0 | 213.2 | 0 | 0 | 210.7 | 0.929 | 0.961238023 | 0.918482999 | 0.9550263 | 0.988273921 | 0.993744884 |
| A0A3Q0CY50 | ras-related protein Rab-6A OS=Mesocricetus auratus OX=10036 GN=Rab6a PE=4 SV=1 | 14.114 | 15 | 3 | 7 | 2 | 208 | 23.5 | 5.54 | 21.59 | 3 | 246.4 | 0 | 96.5 | 264.9 | 0 | 74.5 | 219.2 | 0 | 73.1 | 0.911 | 0.99179498 | 0.852435112 | 0.869681645 | 0.861225692 | 0.885236395 |
| A0A1U7R6E4 | protein SET OS=Mesocricetus auratus OX=10036 GN=LOC101831694 PE=3 SV=2 | 14.067 | 15 | 4 | 11 | 4 | 289 | 33.3 | 4.34 | 38.66 | 4 | 233.9 | 45.3 | 49.1 | 198.5 | 59.6 | 66.6 | 226.2 | 43.2 | 46.8 | 0.999 | 0.988366643 | 0.963143466 | 0.965142341 | 0.97382199 | 0.971974225 |
| A0A3Q0CWH5 | E3 ubiquitin-protein ligase OS=Mesocricetus auratus OX=10036 GN=Nedd4 PE=4 SV=1 | 14.059 | 5 | 4 | 19 | 4 | 888 | 102.3 | 5.34 | 42.25 | 4 | 0 | 0 | 36.7 | 0 | 0 | 28.6 | 0 | 0 | 32 | 0.779 | 0.870729661 | 0.871934605 | 0.927827876 | 1.118881119 | 0.940707274 |
| A0A3Q0D3H9 | 60S ribosomal protein L5 isoform X3 OS=Mesocricetus auratus OX=10036 GN=Rpl5 PE=3 SV=1 | 14.02 | 12 | 3 | 25 | 3 | 296 | 34.4 | 9.7 | 57.5 | 3 | 778.1 | 1086.1 | 832.2 | 814.1 | 1068.8 | 910 | 845.9 | 1017.5 | 901.1 | 1.093 | 0.803499835 | 1.025255897 | 0.84618621 | 0.989831358 | 0.921808217 |
| A0A3Q0CTE4 | inhibitor of carbonic anhydrase OS=Mesocricetus auratus OX=10036 GN=LOC101828453 PE=3 SV=1 | 13.83 | 7 | 4 | 21 | 4 | 704 | 76.8 | 6.74 | 65.91 | 4 | 236.8 | 127.3 | 247.9 | 385.8 | 199.6 | 308.5 | 292.8 | 134.4 | 233.3 | 1.629 | 0.236632853 | 1.079248366 | 0.801728822 | 0.738896968 | 0.336512043 |
| A0A1U7QQ48 | UMP-CMP kinase OS=Mesocricetus auratus OX=10036 GN=Cmpk1 PE=3 SV=2 | 13.815 | 8 | 2 | 16 | 2 | 227 | 25.7 | 7.91 | 45.78 | 2 | 200.6 | 213.5 | 282.8 | 174 | 184.5 | 278.1 | 211.2 | 239.6 | 330.6 | 0.864 | 0.657180639 | 1.121251256 | 0.561659394 | 1.227458373 | 0.379918646 |
| A0A3Q0CVN3 | Tyrosine-protein kinase OS=Mesocricetus auratus OX=10036 GN=Fyn PE=3 SV=1 | 13.683 | 7 | 5 | 21 | 2 | 534 | 60.1 | 6 | 43.91 | 5 | 99.2 | 42.1 | 33.3 | 59.4 | 42.5 | 41.1 | 90.8 | 39.8 | 23.2 | 1.009 | 0.666315164 | 0.880870561 | 0.822733024 | 1.075524476 | 0.87861766 |
| A0A1U7QMP4 | 26S proteasome non-ATPase regulatory subunit 11 OS=Mesocricetus auratus OX=10036 GN=Psmd11 PE=3 SV=2 | 13.599 | 9 | 3 | 11 | 3 | 422 | 47.4 | 6.48 | 33.88 | 3 | 119.7 | 69.6 | 46.2 | 141.1 | 73.1 | 60 | 105 | 80.6 | 54.1 | 1.179 | 0.717754563 | 1.017834395 | 0.960266637 | 0.874179431 | 0.717572058 |
| A0A3Q0CXF4 | NADH dehydrogenase [ubiquinone] flavoprotein 2, mitochondrial OS=Mesocricetus auratus OX=10036 GN=Ndufv2 PE=3 SV=1 | 13.43 | 12 | 3 | 20 | 3 | 248 | 27.3 | 7.71 | 55.19 | 3 | 1093.2 | 418.4 | 1023.7 | 671.7 | 309.8 | 612.6 | 686.3 | 317.1 | 623 | 0.598 | 0.284788942 | 0.641501992 | 0.299246999 | 1.020262217 | 0.949538263 |
| A0A3Q0DCV9 | nexilin isoform X3 OS=Mesocricetus auratus OX=10036 GN=Nexn PE=4 SV=1 | 13.371 | 5 | 5 | 16 | 5 | 658 | 78.4 | 5.27 | 35.35 | 5 | 253.4 | 152.6 | 365.2 | 163 | 124.2 | 313.6 | 195.6 | 138.5 | 291.4 | 0.859 | 0.537507977 | 0.811073651 | 0.560144079 | 1.041111851 | 0.915943004 |
| A0A3Q0D4T8 | Coatomer subunit alpha OS=Mesocricetus auratus OX=10036 GN=Copa PE=4 SV=1 | 13.37 | 5 | 4 | 14 | 4 | 1224 | 138.3 | 7.65 | 29.15 | 4 | 186.1 | 147.9 | 138.5 | 154.7 | 138.3 | 115.4 | 177.2 | 108.4 | 117.6 | 0.885 | 0.315503768 | 0.853333333 | 0.431249848 | 0.987267385 | 0.947767639 |
| A0A1U7RIK2 | cullin-associated NEDD8-dissociated protein 1 OS=Mesocricetus auratus OX=10036 GN=LOC101840342 PE=3 SV=2 | 13.257 | 4 | 4 | 12 | 4 | 1230 | 136.3 | 5.78 | 33.25 | 4 | 45.1 | 37 | 105.9 | 43.8 | 29.7 | 82 | 37.1 | 34.7 | 105.8 | 0.803 | 0.708440792 | 0.944680851 | 0.918659885 | 1.142122186 | 0.807653066 |
| A0A3Q0CJ20 | actin-related protein 2/3 complex subunit 4 OS=Mesocricetus auratus OX=10036 GN=Arpc4 PE=3 SV=1 | 13.033 | 15 | 4 | 21 | 4 | 242 | 27.5 | 9.47 | 56.73 | 4 | 648.2 | 718.8 | 534.5 | 669.5 | 741.1 | 484.1 | 721.6 | 813.7 | 599.4 | 1.033 | 0.981951787 | 1.122640021 | 0.398189308 | 1.12666913 | 0.464372785 |
| A0A1U7R659 | transforming protein RhoA OS=Mesocricetus auratus OX=10036 GN=Rhoa PE=4 SV=2 | 12.963 | 19 | 5 | 17 | 3 | 193 | 21.8 | 6.1 | 15.88 | 5 | 624.4 | 624.1 | 606.9 | 483.6 | 350.3 | 386.1 | 619.1 | 625.6 | 633.6 | 0.577 | 0.031360613 | 1.012342352 | 0.35085765 | 1.539590164 | 0.030235338 |
| A0A3Q0CYX1 | 15-hydroxyprostaglandin dehydrogenase [NAD(+)] OS=Mesocricetus auratus OX=10036 GN=Hpgd PE=3 SV=1 | 12.767 | 10 | 2 | 13 | 2 | 266 | 29 | 6.54 | 39.32 | 2 | 223.7 | 0 | 0 | 195 | 0 | 0 | 217 | 0 | 0 | 0.872 | 0.927690017 | 0.970049173 | 0.983879013 | 1.112820513 | 0.943549323 |
| A0A1U8CHN2 | Coronin OS=Mesocricetus auratus OX=10036 GN=Coro1c PE=3 SV=1 | 12.713 | 15 | 5 | 12 | 5 | 474 | 53.2 | 6.93 | 17.13 | 5 | 52.4 | 73.5 | 165.8 | 45.1 | 62.2 | 122.7 | 58.3 | 83.3 | 158.2 | 0.847 | 0.65357331 | 1.027768255 | 0.956039795 | 1.303478261 | 0.57659391 |
| A0A1U7RIJ7 | protein transport protein Sec31A OS=Mesocricetus auratus OX=10036 GN=LOC101838460 PE=3 SV=2 | 12.476 | 4 | 5 | 24 | 5 | 1231 | 133.7 | 6.65 | 63.54 | 5 | 339.8 | 521.2 | 422.2 | 335.9 | 518.3 | 384.1 | 342.9 | 482.4 | 420.2 | 0.91 | 0.852892435 | 0.970620324 | 0.859189485 | 1.005814423 | 0.973622281 |
| A0A3Q0CH68 | keratin, type I cytoskeletal 42 isoform X2 OS=Mesocricetus auratus OX=10036 GN=LOC101827825 PE=3 SV=1 | 12.434 | 9 | 5 | 17 | 2 | 442 | 49.1 | 5.25 | 37.41 | 5 | 0 | 162.6 | 71 | 0 | 132.7 | 30.3 | 0 | 118.2 | 35.3 | 0.59 | 0.723428675 | 0.657106164 | 0.674488367 | 0.941717791 | 0.955508953 |
| A0A3Q0CNF9 | S-phase kinase-associated protein 1 OS=Mesocricetus auratus OX=10036 GN=Skp1 PE=3 SV=1 | 12.357 | 25 | 3 | 7 | 3 | 163 | 18.6 | 4.54 | 8.29 | 3 | 72.2 | 106.2 | 38.2 | 71.3 | 103.5 | 36.6 | 65.4 | 118.1 | 38.6 | 0.987 | 0.9528363 | 1.025392428 | 0.955042625 | 1.050614948 | 0.912177109 |

Supplementary Table 1: List of differentially expressed proteins identified across the 4dpi infected and 14dpi infected

| Accession | Description | Sum PEP Score | Coverage [%] | # Peptides | # PSMs | # Unique Peptides | # AAs | MW [kDa] | calc. pI | Score Sequest HT: Sequest HT | # Peptides (by Search Engine): Sequest HT | Mock-infected (4dpi) Replicate_R1 | Mock-infected (4dpi) Replicate_R2 | Mock-infected (4dpi) Replicate_R3 | SARS-CoV-2-infected (4dpi) Replicate_R1 | SARS-CoV-2-infected (4dpi) Replicate_R2 | SARS-CoV-2-infected (4dpi) Replicate_R3 | SARS-CoV-2-(14dpi) Replicate_R1 | SARS-CoV-2-infected (14dpi) Replicate_R2 | SARS-CoV-2-infected (14dpi) Replicate_R3 | Abundance Ratio: SARS-CoV-2-infected (4dpi) / Mock-infected (4dpi) | P-Value: SARS-CoV-2-infected (4dpi) / Mock-infected (4dpi) | Abundance Ratio: SARS-CoV-2-infected (14dpi) / Mock-infected (4dpi) | P-Value: SARS-CoV-2-infected (14dpi) / Mock-infected (4dpi) | Abundance Ratio: SARS-CoV-2-infected (14dpi) / SARS-CoV-2-infected (4dpi) | P-Value: SARS-CoV-2-infected (14dpi) / SARS-CoV-2-infected (4dpi) |
|---|---|---|---|---|---|---|---|---|---|---|---|---|---|---|---|---|---|---|---|---|---|---|---|---|---|---|
| A0A3Q0D6F0 | Adenylate kinase 2, mitochondrial OS=Mesocricetus auratus OX=10036 GN=Ak2 PE=3 SV=1 | 12.354 | 24 | 3 | 23 | 3 | 232 | 25.5 | 7.42 | 79.64 | 3 | 118 | 95.3 | 126.5 | 121.4 | 109.1 | 118.6 | 135.5 | 75.1 | 126.3 | 0.938 | 0.780128074 | 0.991465568 | 0.966196499 | 0.965052993 | 0.850255156 |
| A0A1U7R8F8 | transcription intermediary factor 1-beta OS=Mesocricetus auratus OX=10036 GN=Trim28 PE=4 SV=2 | 12.201 | 6 | 4 | 8 | 4 | 833 | 88.7 | 5.72 | 27.1 | 4 | 44.6 | 0 | 0 | 37.9 | 0 | 0 | 30.7 | 0 | 0 | 0.85 | 0.914514216 | 0.688340807 | 0.811582396 | 0.810026385 | 0.890077128 |
| A0A3Q0CV02 | Proteasome subunit alpha type OS=Mesocricetus auratus OX=10036 GN=Psma3 PE=3 SV=1 | 12.148 | 17 | 4 | 16 | 4 | 255 | 28.4 | 5.44 | 12.48 | 4 | 653.1 | 599.6 | 593.3 | 574.5 | 553.9 | 520.6 | 577.4 | 484.3 | 519.6 | 0.877 | 0.058057406 | 0.856608884 | 0.063067588 | 0.958944815 | 0.520606715 |
| A0A1U8BXE1 | 40S ribosomal protein S6 OS=Mesocricetus auratus OX=10036 GN=Rps6 PE=3 SV=2 | 11.856 | 14 | 4 | 8 | 4 | 249 | 28.7 | 10.84 | 20.25 | 4 | 111 | 113.3 | 241 | 118 | 111.4 | 257.2 | 94 | 106.1 | 204.1 | 1.044 | 0.917152117 | 0.868686869 | 0.732156429 | 0.830661734 | 0.667566036 |
| A0A3Q0CE39 | kinectin OS=Mesocricetus auratus OX=10036 GN=Ktn1 PE=4 SV=1 | 11.791 | 3 | 3 | 5 | 3 | 1350 | 154.6 | 5.9 | 11.11 | 3 | 0 | 109.1 | 0 | 0 | 89.6 | 0 | 0 | 113.6 | 0 | 0.821 | 0.897055826 | 1.041246563 | 0.978577853 | 1.267857143 | 0.876712128 |
| A0A1U7QJ36 | cytochrome b-c1 complex subunit 1, mitochondrial OS=Mesocricetus auratus OX=10036 GN=LOC101825065 PE=4 SV=2 | 11.67 | 6 | 3 | 11 | 3 | 480 | 52.7 | 6.25 | 36.62 | 3 | 237 | 535.4 | 222.5 | 206.6 | 483.3 | 207 | 253.6 | 464.2 | 214.4 | 0.872 | 0.823932532 | 0.936978591 | 0.878857881 | 1.039357788 | 0.927015035 |
| A0A1U8BSZ2 | 26S proteasome non-ATPase regulatory subunit 13 OS=Mesocricetus auratus OX=10036 GN=Psmd13 PE=3 SV=2 | 11.626 | 8 | 3 | 5 | 3 | 376 | 42.8 | 5.83 | 16.17 | 3 | 0 | 47 | 45.8 | 0 | 37.6 | 45.5 | 0 | 28.3 | 34 | 0.89 | 0.884554453 | 0.671336207 | 0.619303088 | 0.749699158 | 0.714266469 |
| A0A3Q0CE75 | tenascin-X OS=Mesocricetus auratus OX=10036 GN=Tnxb PE=3 SV=1 | 11.573 | 2 | 3 | 11 | 3 | 3624 | 392.3 | 6.13 | 38.91 | 3 | 56.7 | 94.3 | 139.4 | 50.9 | 93.7 | 83.9 | 42.7 | 79.1 | 135.5 | 0.987 | 0.501747154 | 0.886019284 | 0.775097105 | 1.126039387 | 0.77037039 |
| A0A3Q0CEA8 | vasodilator-stimulated phosphoprotein isoform X2 OS=Mesocricetus auratus OX=10036 GN=Vasp PE=4 SV=1 | 11.531 | 9 | 3 | 10 | 2 | 380 | 39.3 | 8.97 | 23.04 | 3 | 102.6 | 55.9 | 226.3 | 71 | 37.8 | 195.6 | 90.9 | 47.4 | 204.6 | 0.751 | 0.721132602 | 0.891112266 | 0.849835705 | 1.126478318 | 0.857659876 |
| A0A1U8C678 | alpha-1B-glycoprotein OS=Mesocricetus auratus OX=10036 GN=A1bg PE=4 SV=2 | 11.465 | 6 | 2 | 15 | 2 | 513 | 57.6 | 6.37 | 44.41 | 2 | 179.1 | 90.7 | 134.1 | 251 | 124.1 | 161.2 | 167.9 | 85.8 | 151 | 1.369 | 0.393965263 | 1.001980688 | 0.994405186 | 0.754614954 | 0.394612091 |
| A0A3Q0CTA3 | LIM and senescent cell antigen-like-containing domain protein OS=Mesocricetus auratus OX=10036 GN=Lims1 PE=4 SV=1 | 11.427 | 12 | 4 | 12 | 4 | 362 | 41.6 | 8.05 | 27.37 | 4 | 326.2 | 223.4 | 195.7 | 304.9 | 208.2 | 177 | 291.7 | 191.3 | 191.9 | 0.932 | 0.756065448 | 0.905541393 | 0.675007111 | 0.977974207 | 0.925659884 |
| A0A1U7QU32 | protein FAM49A OS=Mesocricetus auratus OX=10036 GN=Fam49a PE=3 SV=2 | 11.385 | 15 | 3 | 7 | 3 | 323 | 37.3 | 6.01 | 17.37 | 3 | 47.3 | 53.9 | 0 | 40.4 | 53.5 | 0 | 30.3 | 40.9 | 0 | 0.921 | 0.922181296 | 0.703557312 | 0.660149239 | 0.758253461 | 0.728683161 |
| A0A1U7RAI7 | T-complex protein 1 subunit alpha OS=Mesocricetus auratus OX=10036 GN=LOC101830320 PE=3 SV=2 | 11.367 | 5 | 3 | 22 | 3 | 466 | 51.3 | 8.1 | 82.3 | 3 | 512.2 | 289.9 | 329.5 | 314.5 | 205.6 | 231.5 | 360.8 | 226.7 | 267.8 | 0.709 | 0.197837027 | 0.75583245 | 0.323651825 | 1.137972326 | 0.540062197 |
| A0A3Q0D655 | Ras-related protein Rab-14 OS=Mesocricetus auratus OX=10036 GN=Rab14 PE=3 SV=1 | 11.247 | 13 | 2 | 5 | 1 | 187 | 21 | 5.63 | 18.19 | 2 | 0 | 0 | 40.6 | 0 | 0 | 37.1 | 0 | 0 | 53.5 | 0.914 | 0.952334611 | 1.31773399 | 0.85768179 | 1.442048518 | 0.814970927 |
| A0A3Q0DDS2 | von Willebrand factor OS=Mesocricetus auratus OX=10036 GN=Vwf PE=4 SV=1 | 11.232 | 2 | 3 | 9 | 3 | 2813 | 308.4 | 5.53 | 20.7 | 3 | 0 | 0 | 53.4 | 0 | 0 | 53.9 | 0 | 0 | 60.3 | 1.009 | 0.995057626 | 1.129213483 | 0.935906221 | 1.118738404 | 0.940773978 |
| A0A3Q0CJH2 | Glutathione reductase OS=Mesocricetus auratus OX=10036 GN=Gsr PE=3 SV=1 | 11.228 | 14 | 4 | 4 | 4 | 503 | 54 | 7.56 | 12.23 | 4 | 0 | 0 | 169.7 | 0 | 0 | 143.2 | 0 | 0 | 141.9 | 0.844 | 0.910910608 | 0.836181497 | 0.906234794 | 0.990921788 | 0.995163715 |
| A0A3Q0CXS8 | Asparagine--tRNA ligase OS=Mesocricetus auratus OX=10036 GN=Nars PE=3 SV=1 | 11.178 | 9 | 4 | 6 | 4 | 559 | 64.1 | 5.96 | 16.25 | 4 | 0 | 96.2 | 90.6 | 0 | 67 | 50.9 | 0 | 99.3 | 105.2 | 0.626 | 0.575054428 | 1.094753747 | 0.904637556 | 1.73452078 | 0.515586592 |
| A0A3Q0D7G9 | NAD(P)H-hydrate epimerase OS=Mesocricetus auratus OX=10036 GN=Naxe PE=3 SV=1 | 11.019 | 15 | 3 | 8 | 3 | 282 | 30.8 | 7.69 | 22.24 | 3 | 0 | 37.5 | 0 | 0 | 37.8 | 0 | 0 | 21 | 0 | 1.008 | 0.99577434 | 0.56 | 0.725565741 | 0.555555556 | 0.722585179 |
| A0A3Q0DD63 | cytoskeleton-associated protein 4 OS=Mesocricetus auratus OX=10036 GN=Ckap4 PE=4 SV=1 | 10.866 | 9 | 5 | 13 | 5 | 574 | 63.5 | 5.69 | 24.57 | 5 | 256.7 | 224.8 | 287.8 | 249.2 | 247.9 | 221.4 | 229.8 | 221 | 275.4 | 0.769 | 0.466991876 | 0.943975042 | 0.593601248 | 1.010716771 | 0.901636031 |
| A0A3Q0CQ96 | Signal transducer and activator of transcription OS=Mesocricetus auratus OX=10036 GN=Stat3 PE=3 SV=1 | 10.842 | 5 | 3 | 6 | 3 | 769 | 87.9 | 6.3 | 15.76 | 3 | 0 | 123.4 | 106 | 0 | 111.2 | 87.5 | 0 | 110.6 | 110.8 | 0.863 | 0.85174306 | 0.965126417 | 0.962551459 | 1.114242577 | 0.887205755 |
| A0A1U7QN73 | cullin-3 OS=Mesocricetus auratus OX=10036 GN=Cul3 PE=3 SV=2 | 10.8 | 4 | 2 | 12 | 2 | 768 | 88.9 | 8.48 | 39.3 | 2 | 268.9 | 259 | 283 | 337.9 | 301.4 | 341.1 | 280 | 260.7 | 303 | 1.164 | 0.028364835 | 1.040448884 | 0.490925788 | 0.860567115 | 0.061347383 |
| A0A3Q0D9W8 | carboxypeptidase M OS=Mesocricetus auratus OX=10036 GN=Cpm PE=3 SV=1 | 10.734 | 7 | 3 | 16 | 3 | 443 | 50.5 | 8.21 | 36.17 | 3 | 237.3 | 228.3 | 200.9 | 206.2 | 182.6 | 156.6 | 172.1 | 149.6 | 117.6 | 0.802 | 0.093338148 | 0.659114779 | 0.021235919 | 0.80546388 | 0.173420273 |
| A0A1U7R5M2 | Coatomer subunit gamma OS=Mesocricetus auratus OX=10036 GN=LOC101835139 PE=3 SV=2 | 10.649 | 4 | 3 | 12 | 3 | 874 | 97.5 | 5.39 | 35.95 | 3 | 292.9 | 201.8 | 674.7 | 246.8 | 180.5 | 600.1 | 199.6 | 176.5 | 562.4 | 0.894 | 0.820089024 | 0.802548315 | 0.708442142 | 0.913470897 | 0.877560221 |
| A0A3Q0CWL9 | carbonyl reductase [NADPH] 3 OS=Mesocricetus auratus OX=10036 GN=LOC106021442 PE=3 SV=1 | 10.62 | 13 | 3 | 8 | 3 | 277 | 30.6 | 5.6 | 18.45 | 3 | 378.3 | 168.3 | 195.8 | 345.3 | 153.4 | 165 | 347.5 | 164.8 | 163.1 | 0.913 | 0.786511965 | 0.909752155 | 0.816147221 | 1.017628447 | 0.966468991 |
| A0A3Q0CXE4 | phosphoglucomutase-1 isoform X1 OS=Mesocricetus auratus OX=10036 GN=Pgm1 PE=3 SV=1 | 10.495 | 6 | 3 | 11 | 3 | 580 | 63.6 | 5.73 | 24.65 | 3 | 138.2 | 160.5 | 83.6 | 111 | 157.3 | 62.3 | 124.1 | 150 | 74.1 | 0.803 | 0.655242576 | 0.910803034 | 0.739655586 | 1.05323654 | 0.876503249 |
| A0A1U7QP95 | histidine-rich glycoprotein OS=Mesocricetus auratus OX=10036 GN=Hrg PE=4 SV=2 | 10.37 | 5 | 3 | 14 | 3 | 555 | 62.2 | 7.53 | 33.32 | 3 | 439.9 | 448.2 | 459.7 | 548.7 | 551.7 | 615.1 | 544.6 | 510.3 | 518.5 | 1.231 | 0.02365176 | 1.167383885 | 0.006951635 | 0.917167007 | 0.147096455 |
| P86225 | Isocitrate dehydrogenase [NAD] subunit alpha, mitochondrial (Fragments) OS=Mesocricetus auratus OX=10036 GN=IDH3A PE=1 SV=1 | 10.174 | 25 | 3 | 10 | 1 | 134 | 14.9 | 8.97 | 26.88 | 3 | 124.1 | 92.3 | 61.7 | 102.4 | 89.2 | 66.6 | 122.8 | 113.7 | 72.7 | 0.966 | 0.769723469 | 1.111830277 | 0.684974735 | 1.197521301 | 0.419407191 |
| A0A6F8PBD7 | Lysosomal associated membrane protein 1 OS=Mesocricetus auratus OX=10036 GN=Lamp1 PE=2 SV=1 | 10.131 | 6 | 2 | 8 | 2 | 408 | 44 | 7.2 | 33.44 | 2 | 93.1 | 50.3 | 31 | 112.3 | 40.7 | 38.5 | 85.8 | 48 | 38.4 | 0.873 | 0.861047859 | 0.987385321 | 0.976545216 | 0.89921671 | 0.833275051 |
| A0A3Q0D5J5 | glycogenin-1 OS=Mesocricetus auratus OX=10036 GN=Gyg1 PE=4 SV=1 | 10.126 | 9 | 3 | 9 | 3 | 394 | 43.6 | 5.86 | 17.56 | 3 | 183.9 | 209.5 | 85.7 | 144.2 | 193.4 | 53.1 | 145.8 | 173.8 | 66.7 | 0.735 | 0.625527906 | 0.806303486 | 0.56686804 | 0.988738162 | 0.97897782 |
| A0A3Q0CLN3 | phosphate carrier protein, mitochondrial OS=Mesocricetus auratus OX=10036 GN=Slc25a3 PE=3 SV=1 | 10.05 | 8 | 3 | 24 | 3 | 357 | 39.5 | 9.26 | 57.17 | 3 | 915.8 | 911.7 | 1036.7 | 1007.2 | 1014.4 | 1136.5 | 955.2 | 984.8 | 1132.2 | 1.1 | 0.17026197 | 1.072620627 | 0.372211072 | 0.972800101 | 0.700668625 |
| A0A3Q0D7U0 | huntingtin-interacting protein 1 isoform X4 OS=Mesocricetus auratus OX=10036 GN=Hip1 PE=3 SV=1 | 9.865 | 4 | 3 | 6 | 3 | 1006 | 112.5 | 5.58 | 18.11 | 3 | 54.1 | 0 | 0 | 52.5 | 0 | 0 | 44.8 | 0 | 0 | 0.97 | 0.984084305 | 0.828096118 | 0.901271467 | 0.853333333 | 0.916667056 |
| A0A3Q0DF81 | supervillin isoform X8 OS=Mesocricetus auratus OX=10036 GN=Svil PE=3 SV=1 | 9.857 | 1 | 2 | 8 | 2 | 1782 | 199.9 | 6.44 | 24.47 | 2 | 123.5 | 117.9 | 94.8 | 116.1 | 91.3 | 72.1 | 97.2 | 106.6 | 86 | 0.774 | 0.296683652 | 0.861986913 | 0.228020915 | 1.036851521 | 0.823670201 |
| A0A1U7R946 | tetraspanin-8-like OS=Mesocricetus auratus OX=10036 GN=LOC101832001 PE=3 SV=2 | 9.796 | 9 | 2 | 13 | 2 | 163 | 17.8 | 4.93 | 42.32 | 2 | 280.5 | 171.6 | 262.2 | 288.1 | 169.2 | 249.9 | 343 | 173.1 | 338.5 | 0.953 | 0.963497313 | 1.196416072 | 0.521211817 | 1.208427602 | 0.505100012 |
| A0A3Q0D4J1 | Staphylococcal nuclease domain-containing protein OS=Mesocricetus auratus OX=10036 GN=Snd1 PE=4 SV=1 | 9.778 | 7 | 4 | 8 | 4 | 910 | 102 | 7.28 | 19.08 | 4 | 0 | 0 | 62.3 | 0 | 0 | 64.3 | 0 | 0 | 69.4 | 1.032 | 0.983248763 | 1.113964687 | 0.943011306 | 1.079315708 | 0.959609138 |
| A0A3Q0DB47 | Cytoplasmic FMR1-interacting protein OS=Mesocricetus auratus OX=10036 GN=Cyfip1 PE=3 SV=1 | 9.741 | 3 | 2 | 13 | 2 | 1295 | 149.2 | 6.81 | 51.07 | 2 | 85 | 108.6 | 36 | 73.7 | 95.3 | 27.1 | 90.7 | 117.4 | 31.8 | 0.878 | 0.723113493 | 1.044860627 | 0.922543951 | 1.223355431 | 0.675958289 |
| A0A3Q0D826 | Actin-related protein 2/3 complex subunit 3 OS=Mesocricetus auratus OX=10036 GN=Arpc3 PE=3 SV=1 | 9.741 | 18 | 3 | 12 | 3 | 170 | 19.6 | 8.6 | 26.03 | 3 | 84.3 | 0 | 79.3 | 89.4 | 0 | 76.1 | 55.3 | 0 | 72.9 | 1.009 | 0.987822013 | 0.783618582 | 0.753972758 | 0.774622356 | 0.744512407 |
| A0A3Q0D7E5 | farnesyl pyrophosphate synthase OS=Mesocricetus auratus OX=10036 GN=Fdps PE=3 SV=1 | 9.686 | 12 | 3 | 6 | 3 | 413 | 47.5 | 5.82 | 12.08 | 3 | 57 | 97.8 | 0 | 46.2 | 97.8 | 0 | 61.8 | 109 | 0 | 0.901 | 0.932660694 | 1.103359173 | 0.90610298 | 1.186111111 | 0.843379232 |
| A0A3Q0CPK5 | splicing factor 3B subunit 2 OS=Mesocricetus auratus OX=10036 GN=Sf3b2 PE=4 SV=1 | 9.561 | 6 | 1 | 9 | 1 | 347 | 38.9 | 5.08 | 30.77 | 1 | 36.2 | 29.4 | 0 | 26.1 | 38.9 | 0 | 28.8 | 33.7 | 0 | 0.976 | 0.990596489 | 0.952743902 | 0.94938321 | 0.961538462 | 0.95982546 |
| A0A1U8CVE7 | 28S ribosomal protein S36, mitochondrial OS=Mesocricetus auratus OX=10036 GN=LOC101835666 PE=4 SV=2 | 9.494 | 33 | 2 | 5 | 2 | 103 | 11.2 | 10.2 | 15.43 | 2 | 59.7 | 65.5 | 152.8 | 49.9 | 61.1 | 140.9 | 61.4 | 52.2 | 134.6 | 0.846 | 0.844455968 | 0.892805755 | 0.81557739 | 0.985311632 | 0.976137643 |
| A0A1U8BQK4 | interferon-induced protein with tetratricopeptide repeats 1-like OS=Mesocricetus auratus OX=10036 GN=LOC106020791 PE=4 SV=2 | 9.429 | 8 | 2 | 8 | 2 | 265 | 30.9 | 8.47 | 28.52 | 2 | 48.7 | 0 | 0 | 565.3 | 1038.8 | 234.4 | 62.2 | 33.3 | 52.9 | 11.598 | 0.124328836 | 1.015742642 | 0.166795275 | 0.080717977 | 0.137023918 |
| A0A3Q0CTD3 | Cytoplasmic protein OS=Mesocricetus auratus OX=10036 GN=Nck1 PE=4 SV=1 | 9.359 | 8 | 3 | 19 | 3 | 377 | 42.8 | 6.67 | 56.95 | 3 | 448.5 | 314.5 | 452 | 396.2 | 291.6 | 391.8 | 408.6 | 327.7 | 404.1 | 0.868 | 0.473749649 | 0.938600823 | 0.665094947 | 1.056317155 | 0.664061092 |
| A0A3Q0CKE4 | guanine nucleotide-binding protein subunit alpha-14 isoform X1 OS=Mesocricetus auratus OX=10036 GN=Gna14 PE=3 SV=1 | 9.298 | 7 | 2 | 8 | 1 | 355 | 41.4 | 6.3 | 20.26 | 2 | 102.8 | 0 | 52.6 | 112.6 | 0 | 66.1 | 122 | 0 | 52.4 | 1.173 | 0.868942593 | 1.122265122 | 0.897653477 | 0.975937325 | 0.977673675 |
| A0A3Q0CHA0 | endoglin OS=Mesocricetus auratus OX=10036 GN=Eng PE=4 SV=1 | 9.264 | 5 | 2 | 7 | 2 | 653 | 70 | 6.18 | 16.98 | 2 | 112.7 | 0 | 41.6 | 80.1 | 0 | 23.3 | 85.5 | 0 | 34 | 0.631 | 0.69947724 | 0.774465327 | 0.793408226 | 1.155705996 | 0.883606041 |
| A0A3Q0CLC2 | myosin phosphatase Rho-interacting protein OS=Mesocricetus auratus OX=10036 GN=Mprip PE=4 SV=1 | 9.23 | 3 | 4 | 11 | 4 | 2424 | 274.3 | 5.71 | 20.43 | 4 | 95.6 | 48.2 | 97.1 | 80.8 | 32.5 | 76.5 | 100.2 | 45.7 | 97.5 | 0.713 | 0.487062622 | 1.01037775 | 0.973895961 | 1.282402529 | 0.490344874 |
| A0A3Q0D2X1 | guanylate-binding protein 6 OS=Mesocricetus auratus OX=10036 GN=LOC101837799 PE=3 SV=1 | 9.22 | 7 | 4 | 10 | 2 | 620 | 70.5 | 6.2 | 29.79 | 4 | 352.4 | 252.8 | 249.9 | 375.2 | 244.3 | 256.6 | 285.3 | 248.4 | 255.2 | 1.065 | 0.902733278 | 0.922582154 | 0.587710435 | 0.900467983 | 0.562886256 |
| A0A3Q0CN90 | alpha-adducin OS=Mesocricetus auratus OX=10036 GN=Add1 PE=3 SV=1 | 9.191 | 9 | 3 | 11 | 3 | 781 | 85.6 | 5.81 | 40.26 | 3 | 200.9 | 151.3 | 128.1 | 163.5 | 125.6 | 127.3 | 172.2 | 115.8 | 89.1 | 0.879 | 0.449621958 | 0.785134291 | 0.351434202 | 0.905619597 | 0.666097877 |
| A0A3Q0CLT5 | BAG family molecular chaperone regulator 3 OS=Mesocricetus auratus OX=10036 GN=Bag3 PE=4 SV=1 | 9.141 | 9 | 3 | 10 | 3 | 548 | 58.2 | 7.4 | 28.8 | 3 | 222.5 | 259.1 | 218.1 | 214.2 | 252.2 | 225.6 | 182.4 | 189.3 | 176.8 | 0.973 | 0.888698838 | 0.783907389 | 0.051685415 | 0.792630058 | 0.040675112 |
| A0A1U7QQ37 | Clathrin light chain OS=Mesocricetus auratus OX=10036 GN=Cltb PE=3 SV=2 | 9.097 | 16 | 4 | 11 | 4 | 229 | 25.2 | 4.63 | 21.82 | 4 | 44.6 | 152.4 | 232.4 | 37.6 | 131.1 | 207.1 | 32.4 | 109.1 | 145.2 | 0.861 | 0.819382469 | 0.667675827 | 0.505150984 | 0.762905801 | 0.64581025 |
| A0A3Q0CJY5 | hsc70-interacting protein OS=Mesocricetus auratus OX=10036 GN=St13 PE=3 SV=1 | 9.054 | 6 | 2 | 13 | 2 | 368 | 41.1 | 5.31 | 20.56 | 2 | 196.2 | 250.8 | 98 | 116.8 | 129.7 | 64.1 | 192.8 | 271.6 | 82.5 | 0.596 | 0.216332411 | 1.003486239 | 0.993303102 | 1.760785576 | 0.285552042 |
| A0A3Q0CG52 | H-2 class I histocompatibility antigen, K-K alpha chain OS=Mesocricetus auratus OX=10036 GN=LOC101837574 PE=3 SV=1 | 9.046 | 23 | 3 | 8 | 3 | 120 | 14 | 6.79 | 23 | 3 | 219.6 | 83.1 | 242.9 | 368.5 | 144 | 396.4 | 217.9 | 107.2 | 238.5 | 1.678 | 0.280052719 | 1.032991202 | 0.930406803 | 0.620090219 | 0.290209007 |
| A0A3Q0D882 | 60S ribosomal protein L6 OS=Mesocricetus auratus OX=10036 GN=Rpl6 PE=3 SV=1 | 8.991 | 13 | 3 | 15 | 3 | 242 | 27.2 | 10.68 | 38.37 | 3 | 82.1 | 59 | 53.9 | 72.4 | 55.9 | 53.9 | 92.3 | 847.5 | 60.5 | 0.881 | 0.707244493 | 5.12974359 | 0.406204758 | 5.490120746 | 0.400140571 |
| A0A3Q0DA11 | Low molecular weight cytosolic acid phosphatase OS=Mesocricetus auratus OX=10036 GN=Acp1 PE=3 SV=1 | 8.936 | 8 | 2 | 9 | 2 | 158 | 18.1 | 6.32 | 31.87 | 2 | 305.1 | 234.6 | 116.9 | 224 | 178.9 | 79.5 | 223.8 | 156.3 | 102.5 | 0.763 | 0.453321832 | 0.734998477 | 0.431824108 | 1.000414594 | 0.999097181 |
| A0A3Q0D2X8 | synaptopodin-2 OS=Mesocricetus auratus OX=10036 GN=Synpo2 PE=4 SV=1 | 8.926 | 5 | 3 | 4 | 3 | 1261 | 136 | 7.96 | 10.93 | 3 | 38.4 | 34.2 | 0 | 30 | 34.6 | 0 | 37 | 32 | 0 | 0.889 | 0.878048997 | 0.950413223 | 0.94649043 | 1.068111455 | 0.930850337 |
| A0A1U7QE42 | clathrin interactor 1 isoform X2 OS=Mesocricetus auratus OX=10036 GN=Clint1 PE=4 SV=2 | 8.898 | 7 | 3 | 5 | 3 | 623 | 68 | 6.42 | 12.94 | 3 | 27.3 | 36.8 | 42.9 | 38.1 | 32 | 45.7 | 30.5 | 36.2 | 37.4 | 1.03 | 0.652286618 | 0.972897196 | 0.860055649 | 0.898963731 | 0.448477362 |
| P81468 | Neutrophil defensin 4 OS=Mesocricetus auratus OX=10036 PE=1 SV=1 | 8.829 | 45 | 1 | 17 | 1 | 33 | 3.8 | 8.29 | 52.93 | 1 | 66.3 | 20.5 | 105.5 | 124.8 | 49.5 | 214 | 89.4 | 33.1 | 147.5 | 2.04 | 0.309422875 | 1.404056162 | 0.565969393 | 0.695338656 | 0.537397329 |
| A0A3Q0CS88 | Dynamin GTPase OS=Mesocricetus auratus OX=10036 GN=Dnm3 PE=3 SV=1 | 8.797 | 4 | 3 | 12 | 1 | 670 | 76 | 8.24 | 24.89 | 3 | 75.8 | 0 | 0 | 55.9 | 0 | 0 | 58.2 | 0 | 0 | 0.738 | 0.843841938 | 0.767810026 | 0.86340917 | 1.041144902 | 0.978629539 |
| A0A3Q0CZY2 | beta-2-syntrophin OS=Mesocricetus auratus OX=10036 GN=Sntb2 PE=3 SV=1 | 8.779 | 8 | 2 | 10 | 2 | 522 | 56.4 | 8.57 | 31.18 | 2 | 123.5 | 99.1 | 134.3 | 107 | 94.4 | 121.5 | 117.3 | 82.7 | 111.5 | 0.866 | 0.436899785 | 0.8727935 | 0.368079236 | 0.964694952 | 0.78985865 |
| A0A3Q0CL24 | Tripeptidyl-peptidase 2 OS=Mesocricetus auratus OX=10036 GN=Tpp2 PE=4 SV=1 | 8.657 | 3 | 2 | 7 | 2 | 1262 | 139.5 | 6.54 | 31.58 | 2 | 0 | 0 | 110.1 | 0 | 0 | 91.9 | 0 | 0 | 57.2 | 0.834 | 0.905324894 | 0.519527702 | 0.698529021 | 0.622415669 | 0.767528759 |
| A0A1U8CSV7 | Isocitrate dehydrogenase [NAD] subunit, mitochondrial OS=Mesocricetus auratus OX=10036 GN=LOC101843465 PE=3 SV=2 | 8.634 | 12 | 3 | 9 | 1 | 288 | 31.4 | 6.24 | 18.57 | 3 | 81.3 | 176 | 183.6 | 58.4 | 175.6 | 151.2 | 60.6 | 124.2 | 134.3 | 0.828 | 0.721739987 | 0.723746881 | 0.375633105 | 0.828400831 | 0.635888416 |
| A0A3Q0CUI3 | transformer-2 protein homolog beta OS=Mesocricetus auratus OX=10036 GN=Tra2b PE=4 SV=1 | 8.615 | 7 | 2 | 14 | 2 | 288 | 33.6 | 11.25 | 39.74 | 2 | 539.5 | 874.9 | 893.8 | 502.5 | 746.9 | 806.6 | 249.2 | 470.8 | 470.5 | 0.854 | 0.601593182 | 0.515769864 | 0.062797467 | 0.579036965 | 0.075321277 |
| A0A3Q0CGZ9 | unconventional myosin-Ib isoform X3 OS=Mesocricetus auratus OX=10036 GN=LOC106022830 PE=3 SV=1 | 8.612 | 4 | 4 | 6 | 4 | 1080 | 125.1 | 9.07 | 11.03 | 4 | 83.2 | 0 | 0 | 56 | 0 | 0 | 52 | 0 | 0 | 0.673 | 0.80144245 | 0.625 | 0.769265333 | 0.928571429 | 0.960778658 |
| A0A3Q0D8R7 | pulmonary surfactant-associated protein A OS=Mesocricetus auratus OX=10036 GN=LOC101835350 PE=4 SV=1 | 8.552 | 10 | 2 | 15 | 2 | 248 | 26.3 | 5.47 | 12.69 | 2 | 551.2 | 744.6 | 586 | 557.2 | 748.6 | 597.7 | 550.8 | 635.6 | 629.5 | 1.011 | 0.934955503 | 0.964980338 | 0.760787501 | 0.953979511 | 0.682286388 |
| A0A3Q0CVF3 | Mitogen-activated protein kinase OS=Mesocricetus auratus OX=10036 GN=Mapk3 PE=3 SV=1 | 8.543 | 13 | 4 | 4 | 2 | 346 | 40.1 | 7.47 | 7.89 | 4 | 128.9 | 189.8 | 133.1 | 106 | 150.9 | 95.8 | 107.6 | 183.7 | 109 | 0.822 | 0.272973277 | 0.88601151 | 0.620581073 | 1.134958889 | 0.631901543 |

**Supplementary Table 1: List of differentially expressed proteins identified across the 4dpi infected and 14dpi infected**

| Accession | Description | Sum PEP Score | Coverage [%] | # Peptides | # PSMs | # Unique Peptides | # AAs | MW [kDa] | calc. pI | Score Sequest HT: Sequest HT | # Peptides (by Search Engine): Sequest HT | Mock-infected (4dpi) Replicate_R1 | Mock-infected (4dpi) Replicate_R2 | Mock-infected (4dpi) Replicate_R3 | SARS-CoV-2-infected (4dpi) Replicate_R1 | SARS-CoV-2-infected (4dpi) Replicate_R2 | SARS-CoV-2-infected (4dpi) Replicate_R3 | SARS-CoV-2-(14dpi) Replicate_R1 | SARS-CoV-2-infected (14dpi) Replicate_R2 | SARS-CoV-2-infected (14dpi) Replicate_R3 | Abundance Ratio: SARS-CoV-2-infected (4dpi) / Mock-infected (4dpi) | P-Value: SARS-CoV-2-infected (4dpi) / Mock-infected (4dpi) | Abundance Ratio: SARS-CoV-2-infected (14dpi) / Mock-infected (4dpi) | P-Value: SARS-CoV-2-infected (14dpi) / Mock-infected (4dpi) | Abundance Ratio: SARS-CoV-2-infected (14dpi) / SARS-CoV-2-infected (4dpi) | P-Value: SARS-CoV-2-infected (14dpi) / SARS-CoV-2-infected (4dpi) |
|---|---|---|---|---|---|---|---|---|---|---|---|---|---|---|---|---|---|---|---|---|---|---|---|---|---|---|
| A0A1U8CCX6 | SUMO-activating enzyme subunit 2 OS=Mesocricetus auratus OX=10036 GN=Uba2 PE=3 SV=2 | 8.516 | 5 | 3 | 7 | 3 | 656 | 72.3 | 5.25 | 14.86 | 3 | 202.3 | 0 | 40.4 | 185.1 | 0 | 36 | 213.2 | 0 | 37.3 | 0.902 | 0.935728335 | 1.032138443 | 0.978397952 | 1.132971506 | 0.915646414 |
| A0A3Q0CMB3 | D-beta-hydroxybutyrate dehydrogenase, mitochondrial OS=Mesocricetus auratus OX=10036 GN=Bdh1 PE=4 SV=1 | 8.439 | 14 | 3 | 17 | 2 | 206 | 23.4 | 7.44 | 40.17 | 3 | 238.8 | 339.5 | 251.3 | 228.4 | 333.5 | 251.6 | 234 | 357 | 268.7 | 1.001 | 0.91071971 | 1.036282546 | 0.846170537 | 1.056791641 | 0.767228782 |
| A0A3Q0CUP1 | Non-specific serine/threonine protein kinase OS=Mesocricetus auratus OX=10036 GN=Stk3 PE=3 SV=1 | 8.421 | 9 | 4 | 6 | 4 | 491 | 55.9 | 5.15 | 9.74 | 4 | 91.3 | 0 | 63.3 | 75.5 | 0 | 55.8 | 86.8 | 0 | 56.1 | 0.854 | 0.836583858 | 0.924320828 | 0.921318448 | 1.088347296 | 0.915041753 |
| A0A3Q0CK22 | TRIO and F-actin-binding protein isoform X2 OS=Mesocricetus auratus OX=10036 GN=Triobp PE=4 SV=1 | 8.384 | 11 | 2 | 9 | 2 | 402 | 44.7 | 8.5 | 24.6 | 2 | 41.1 | 0 | 0 | 31.3 | 0 | 0 | 33.4 | 0 | 0 | 0.762 | 0.859402435 | 0.812652068 | 0.891712332 | 1.067092652 | 0.965615621 |
| A0A3Q0CMX8 | Synaptosomal-associated protein OS=Mesocricetus auratus OX=10036 GN=Snap23 PE=3 SV=1 | 8.346 | 7 | 2 | 8 | 2 | 210 | 23.2 | 4.93 | 24.5 | 2 | 143 | 50 | 0 | 110.9 | 35.8 | 0 | 96.4 | 30.4 | 0 | 0.745 | 0.786712685 | 0.656994819 | 0.68841069 | 0.864349012 | 0.885871802 |
| A0A3Q0CG14 | Phospholipid scramblase OS=Mesocricetus auratus OX=10036 GN=Plscr3 PE=3 SV=1 | 8.307 | 13 | 2 | 7 | 2 | 241 | 25.7 | 7.15 | 21.51 | 2 | 226.8 | 121.4 | 78.7 | 173.7 | 94.9 | 63.7 | 173.3 | 126.8 | 73.7 | 0.766 | 0.598529213 | 0.875614898 | 0.755887853 | 1.12488715 | 0.767044988 |
| A0A3Q0D7K0 | Vacuolar protein sorting-associated protein 29 OS=Mesocricetus auratus OX=10036 GN=Vps29 PE=3 SV=1 | 8.288 | 13 | 2 | 6 | 2 | 182 | 20.5 | 6.79 | 24.72 | 2 | 128.8 | 183.8 | 165.8 | 135.1 | 226.6 | 174.3 | 104.3 | 130.4 | 112.3 | 1.051 | 0.576392588 | 0.725334448 | 0.096368841 | 0.64738806 | 0.131519999 |
| A0A3Q0CUA1 | transforming growth factor beta-1-induced transcript 1 protein isoform X3 OS=Mesocricetus auratus OX=10036 GN=Tgfb1i1 PE=3 SV=1 | 8.19 | 6 | 2 | 11 | 2 | 444 | 48.3 | 7.23 | 30.63 | 2 | 93.2 | 83.4 | 35.6 | 75.8 | 76.7 | 31.9 | 73.1 | 55.2 | 30.2 | 0.874 | 0.709868997 | 0.746936852 | 0.461062873 | 0.859544469 | 0.678791132 |
| A0A3Q0CIN0 | asporin OS=Mesocricetus auratus OX=10036 GN=Aspn PE=3 SV=1 | 8.166 | 9 | 3 | 9 | 3 | 373 | 42.6 | 8.44 | 14.53 | 3 | 151.1 | 268.7 | 382.8 | 152 | 236 | 321 | 171.3 | 282.5 | 391.4 | 0.878 | 0.72710626 | 1.053077498 | 0.885145683 | 1.192101551 | 0.603082427 |
| A0A3Q0D8C4 | Heterochromatin protein 1-binding protein 3 OS=Mesocricetus auratus OX=10036 GN=Hp1bp3 PE=4 SV=1 | 8.086 | 6 | 3 | 9 | 3 | 521 | 57.8 | 9.64 | 9.79 | 3 | 0 | 0 | 28.9 | 0 | 0 | 32.2 | 0 | 0 | 43.5 | 1.114 | 0.942906918 | 1.505190311 | 0.795637402 | 1.350931677 | 0.845636096 |
| A0A3Q0CWS8 | intersectin-1 OS=Mesocricetus auratus OX=10036 GN=Itsn1 PE=4 SV=1 | 8.082 | 1 | 1 | 2 | 1 | 1714 | 193.7 | 7.91 | 9.42 | 1 | 189.1 | 0 | 0 | 145.3 | 0 | 0 | 160 | 0 | 0 | 0.768 | 0.863771485 | 0.846113168 | 0.912291755 | 1.101169993 | 0.949066739 |
| A0A3Q0D874 | Beta-glucuronidase OS=Mesocricetus auratus OX=10036 GN=Gusb PE=3 SV=1 | 8.028 | 4 | 3 | 12 | 3 | 648 | 74.6 | 6.68 | 31.71 | 3 | 232.9 | 104.4 | 238.7 | 246.3 | 118.1 | 241.8 | 218.9 | 113.1 | 243.5 | 1.032 | 0.876356537 | 0.999131944 | 0.997894613 | 0.949356648 | 0.868551779 |
| A0A3Q0CDT6 | methylcrotonoyl-CoA carboxylase beta chain, mitochondrial OS=Mesocricetus auratus OX=10036 GN=Mccc2 PE=4 SV=1 | 8.015 | 6 | 2 | 9 | 2 | 563 | 61.4 | 8.53 | 31.75 | 2 | 94 | 49.2 | 31.4 | 81.1 | 39.7 | 40 | 74.1 | 41.5 | 28.8 | 1.012 | 0.852974555 | 0.827033219 | 0.686233124 | 0.89800995 | 0.79066287 |
| A0A1U8C2G6 | golgin subfamily B member 1 isoform X2 OS=Mesocricetus auratus OX=10036 GN=Golgb1 PE=4 SV=2 | 7.966 | 1 | 2 | 5 | 2 | 3189 | 365.5 | 5.01 | 13.99 | 2 | 77.7 | 0 | 0 | 97.6 | 0 | 0 | 61.3 | 0 | 0 | 1.255 | 0.881361003 | 0.788931789 | 0.876838112 | 0.62807377 | 0.771330189 |
| A0A1U8BZY7 | Ribosomal protein L15 OS=Mesocricetus auratus OX=10036 GN=LOC101838491 PE=3 SV=2 | 7.947 | 9 | 2 | 6 | 2 | 204 | 24.1 | 11.58 | 11.63 | 2 | 87.3 | 179.8 | 87.3 | 94.2 | 182.8 | 110.6 | 93.2 | 199.7 | 105.3 | 1.021 | 0.801363905 | 1.123589165 | 0.765231941 | 1.027347781 | 0.939032541 |
| A0A1U8BTW3 | histidine triad nucleotide-binding protein 1 OS=Mesocricetus auratus OX=10036 GN=Hint1 PE=4 SV=2 | 7.87 | 17 | 2 | 14 | 2 | 126 | 13.8 | 6.64 | 33.65 | 2 | 574.3 | 486 | 384.3 | 523.2 | 468 | 381.8 | 557.6 | 570.4 | 396.5 | 0.921 | 0.746755302 | 1.055309428 | 0.751125439 | 1.110342316 | 0.510779933 |
| A0A3Q0D152 | Butyryl-CoA dehydrogenase OS=Mesocricetus auratus OX=10036 GN=Ivd PE=3 SV=1 | 7.592 | 5 | 2 | 7 | 2 | 422 | 46.1 | 7.88 | 15.98 | 2 | 77.2 | 110.4 | 295.3 | 61.6 | 98.6 | 206.4 | 87.7 | 126.3 | 261.1 | 0.699 | 0.659628154 | 0.983847587 | 0.977369564 | 1.295962902 | 0.624842583 |
| A0A3Q0CN22 | RNA helicase OS=Mesocricetus auratus OX=10036 GN=Dhx15 PE=4 SV=1 | 7.589 | 4 | 2 | 9 | 2 | 660 | 74.5 | 7.31 | 32.36 | 2 | 83.4 | 174.3 | 57.6 | 94.3 | 149.8 | 61.3 | 74.7 | 151.8 | 59.5 | 0.86 | 0.943892568 | 0.907072629 | 0.84093368 | 0.936476752 | 0.87487628 |
| A0A3Q0DDN3 | quinone oxidoreductase OS=Mesocricetus auratus OX=10036 GN=Cryz PE=3 SV=1 | 7.586 | 6 | 2 | 11 | 2 | 409 | 43.8 | 8.68 | 29.01 | 2 | 443.5 | 155.3 | 444.4 | 319.3 | 107.4 | 331.8 | 361.4 | 133.3 | 402.4 | 0.703 | 0.478574664 | 0.859950153 | 0.722336156 | 1.182729071 | 0.698852054 |
| A0A3Q0CQF6 | Ubiquitin carboxyl-terminal hydrolase OS=Mesocricetus auratus OX=10036 GN=Uchl3 PE=3 SV=1 | 7.555 | 12 | 3 | 3 | 3 | 294 | 33.2 | 6.68 | 6.88 | 3 | 48.7 | 0 | 96.6 | 43.4 | 0 | 113.3 | 31.1 | 0 | 107.1 | 1.022 | 0.934257289 | 0.951135582 | 0.958111494 | 0.881940013 | 0.899488055 |
| A0A3Q0D540 | calcium-binding mitochondrial carrier protein Aralar1 OS=Mesocricetus auratus OX=10036 GN=Slc25a12 PE=3 SV=1 | 7.533 | 6 | 3 | 4 | 3 | 678 | 74.7 | 8.1 | 9.25 | 3 | 36.8 | 0 | 0 | 37.9 | 0 | 0 | 23.6 | 0 | 0 | 1.029 | 0.984385102 | 0.641304348 | 0.78020889 | 0.622691293 | 0.767713988 |
| A0A3Q0D9H4 | U2 snRNP auxiliary factor large subunit OS=Mesocricetus auratus OX=10036 GN=U2af2 PE=3 SV=1 | 7.488 | 9 | 2 | 7 | 2 | 475 | 53.5 | 9.09 | 27 | 2 | 544.7 | 363.8 | 151.1 | 407.4 | 306.2 | 121.6 | 392.9 | 212.5 | 120.6 | 0.842 | 0.626674599 | 0.685164213 | 0.473439761 | 0.869252874 | 0.768900652 |
| A0A3Q0CYN6 | protein NOXP20 isoform X2 OS=Mesocricetus auratus OX=10036 GN=Fam114a1 PE=3 SV=1 | 7.47 | 2 | 1 | 4 | 1 | 442 | 48 | 5.03 | 11.11 | 1 | 60.3 | 187 | 159.5 | 33.9 | 125.7 | 139.1 | 58 | 179.4 | 146.9 | 0.729 | 0.517573787 | 0.944690265 | 0.894107451 | 1.286575159 | 0.592483946 |
| A0A3Q0CTE1 | Arginine--tRNA ligase, cytoplasmic OS=Mesocricetus auratus OX=10036 GN=Rars PE=3 SV=1 | 7.443 | 5 | 3 | 10 | 3 | 661 | 75.6 | 6.8 | 6.42 | 3 | 97 | 60.9 | 76.1 | 119.9 | 97.4 | 97 | 99.1 | 67.4 | 76.3 | 1.293 | 0.113723478 | 1.037606838 | 0.845388562 | 0.77251034 | 0.123522707 |
| A0A1U8CB90 | E3 ubiquitin/ISG15 ligase TRIM25 OS=Mesocricetus auratus OX=10036 GN=Trim25 PE=4 SV=2 | 7.43 | 3 | 2 | 9 | 2 | 637 | 72.2 | 8.25 | 24.96 | 2 | 48.2 | 0 | 31.5 | 51.7 | 0 | 34.6 | 40.6 | 0 | 33.7 | 1.086 | 0.920727814 | 0.932245922 | 0.928746774 | 0.860950174 | 0.849440236 |
| A0A3Q0CMP4 | voltage-dependent anion-selective channel protein 1 OS=Mesocricetus auratus OX=10036 GN=Vdac1 PE=3 SV=1 | 7.343 | 12 | 1 | 5 | 1 | 165 | 18.4 | 7.93 | 26.61 | 1 | 29.3 | 83.3 | 0 | 27.7 | 61.2 | 0 | 36.4 | 63.7 | 0 | 0.833 | 0.80734378 | 0.888987567 | 0.898691051 | 1.125984252 | 0.89096486 |
| A0A1U7RJ95 | Guanine nucleotide-binding protein subunit gamma OS=Mesocricetus auratus OX=10036 GN=Gng12 PE=3 SV=2 | 7.343 | 26 | 1 | 10 | 1 | 72 | 8 | 8.97 | 41.94 | 1 | 40 | 131.4 | 100.1 | 37.1 | 113.5 | 82.5 | 46.6 | 139.4 | 122.7 | 0.863 | 0.732302071 | 1.137016575 | 0.767504862 | 1.324324324 | 0.526485645 |
| A0A1U7QU54 | Myoglobin OS=Mesocricetus auratus OX=10036 GN=Mb PE=3 SV=2 | 7.271 | 14 | 2 | 6 | 2 | 159 | 17.6 | 7.21 | 6.84 | 2 | 45.1 | 34.9 | 30.6 | 156.7 | 125.8 | 106.1 | 56.9 | 31.9 | 31.8 | 3.471 | 0.017894003 | 1.090415913 | 0.746207777 | 0.310344828 | 0.011570931 |
| A0A1U7Q5D3 | WAS/WASL-interacting protein family member 1 OS=Mesocricetus auratus OX=10036 GN=Wipf1 PE=4 SV=2 | 7.259 | 8 | 2 | 10 | 2 | 494 | 50.2 | 11.53 | 32.45 | 2 | 0 | 0 | 55.5 | 0 | 0 | 50.6 | 0 | 0 | 34.4 | 0.912 | 0.951136483 | 0.61981982 | 0.765784026 | 0.679841897 | 0.805942182 |
| A0A3Q0D013 | guanylate-binding protein 6-like OS=Mesocricetus auratus OX=10036 GN=LOC101823391 PE=3 SV=1 | 7.237 | 5 | 3 | 6 | 1 | 642 | 73.3 | 5.87 | 15.02 | 3 | 0 | 67.4 | 0 | 0 | 88.3 | 0 | 0 | 55.5 | 0 | 1.309 | 0.860525026 | 0.823442136 | 0.898401616 | 0.628539071 | 0.771642701 |
| A0A3Q0CJ83 | serpin B9 OS=Mesocricetus auratus OX=10036 GN=LOC106020742 PE=3 SV=1 | 7.206 | 10 | 3 | 6 | 2 | 335 | 39 | 6.44 | 11.79 | 3 | 0 | 344.5 | 271.9 | 0 | 321.6 | 203 | 0 | 228.4 | 211.7 | 0.835 | 0.838651156 | 0.713984426 | 0.672718511 | 0.838924895 | 0.825511786 |
| A0A3Q0D7Q3 | Calcium-transporting ATPase OS=Mesocricetus auratus OX=10036 GN=Atp2a2 PE=3 SV=1 | 7.204 | 3 | 2 | 6 | 1 | 824 | 90.5 | 5.62 | 20.04 | 2 | 0 | 40 | 57.1 | 0 | 42.6 | 40.2 | 0 | 37.1 | 49 | 0.866 | 0.838352142 | 0.886714727 | 0.878319413 | 1.039855072 | 0.959224411 |
| A0A1U7R2S5 | Transcription elongation factor SPT5 OS=Mesocricetus auratus OX=10036 GN=Supt5h PE=3 SV=2 | 7.135 | 2 | 1 | 5 | 1 | 1086 | 120.6 | 5.14 | 18 | 1 | 0 | 0 | 40.7 | 0 | 0 | 55 | 0 | 0 | 42.6 | 1.351 | 0.845486903 | 1.046683047 | 0.975821856 | 0.774545455 | 0.867709467 |
| A0A3Q0D1N6 | KH domain-containing, RNA-binding, signal transduction-associated protein 1 OS=Mesocricetus auratus OX=10036 GN=Khdrbs1 PE=3 SV=1 | 7.117 | 7 | 2 | 8 | 2 | 347 | 38.7 | 6.47 | 19.55 | 2 | 82.6 | 164.6 | 81.2 | 91.2 | 184.1 | 72.6 | 88 | 193.8 | 77.2 | 1.119 | 0.890410803 | 1.09317905 | 0.837288476 | 1.03190572 | 0.945372792 |
| A0A1U7RH54 | splicing factor 3A subunit 3-like OS=Mesocricetus auratus OX=10036 GN=LOC101833484 PE=4 SV=2 | 7.09 | 22 | 2 | 4 | 2 | 134 | 16 | 4.98 | 11.99 | 2 | 0 | 35.6 | 80.7 | 0 | 34.9 | 56.6 | 0 | 33 | 96.7 | 0.829 | 0.788309981 | 1.115219261 | 0.909331945 | 1.417486339 | 0.722342134 |
| A0A3Q0CNF3 | RNA-binding protein EWS OS=Mesocricetus auratus OX=10036 GN=Ewsr1 PE=3 SV=1 | 7.061 | 2 | 1 | 12 | 1 | 656 | 68.5 | 9.33 | 41.25 | 1 | 395.4 | 142.9 | 201.6 | 445.3 | 147.3 | 206.6 | 362.2 | 131.9 | 212.6 | 1.126 | 0.876164522 | 0.955129071 | 0.918775207 | 0.884259259 | 0.800135558 |
| A0A3Q0CVD3 | Guanylate cyclase OS=Mesocricetus auratus OX=10036 GN=Gucy1a3 PE=3 SV=1 | 7.008 | 5 | 3 | 8 | 3 | 691 | 77.5 | 7.65 | 23.13 | 3 | 164.1 | 0 | 141.9 | 126.3 | 0 | 117.7 | 127.2 | 0 | 122.2 | 0.799 | 0.769224024 | 0.81503268 | 0.790135883 | 1.022131148 | 0.976818142 |
| A0A3Q0CFA5 | osteoclast-stimulating factor 1 isoform X2 OS=Mesocricetus auratus OX=10036 GN=Ostf1 PE=4 SV=1 | 6.996 | 15 | 3 | 12 | 3 | 181 | 20.2 | 5.05 | 26.17 | 3 | 0 | 123.5 | 254.7 | 0 | 119.7 | 277.4 | 0 | 142.4 | 254.5 | 1.027 | 0.956664322 | 1.049444738 | 0.955112094 | 0.999496349 | 0.999541391 |
| A0A3Q0CJL7 | RNA helicase OS=Mesocricetus auratus OX=10036 GN=Ddx42 PE=4 SV=1 | 6.992 | 2 | 1 | 4 | 1 | 929 | 101.8 | 6.98 | 12.62 | 1 | 0 | 49.7 | 39.4 | 0 | 32.5 | 60.9 | 0 | 44.5 | 25.2 | 1.006 | 0.953791114 | 0.782267116 | 0.761705707 | 0.746252677 | 0.737051186 |
| A0A3Q0D281 | SWI/SNF complex subunit SMARCC2 OS=Mesocricetus auratus OX=10036 GN=Smarcc2 PE=4 SV=1 | 6.814 | 4 | 2 | 2 | 2 | 1243 | 135.5 | 5.64 | 9.06 | 2 | 41.6 | 42.5 | 0 | 31 | 31.1 | 0 | 30.5 | 35.5 | 0 | 0.738 | 0.697273441 | 0.784780024 | 0.753573305 | 1.062801932 | 0.935856183 |
| A0A3Q0CQJ9 | peroxisomal multifunctional enzyme type 2 OS=Mesocricetus auratus OX=10036 GN=Hsd17b4 PE=4 SV=1 | 6.767 | 3 | 2 | 7 | 2 | 770 | 83.2 | 8.73 | 24.47 | 2 | 193 | 96 | 97.9 | 156.4 | 73.3 | 69.6 | 163.4 | 87.7 | 66.5 | 0.81 | 0.532695538 | 0.820883949 | 0.623534319 | 1.061142666 | 0.888506033 |
| A0A3Q0CQJ3 | Sorting nexin-2 OS=Mesocricetus auratus OX=10036 GN=Snx2 PE=3 SV=1 | 6.748 | 5 | 2 | 9 | 1 | 523 | 58.9 | 5.05 | 19.17 | 2 | 0 | 0 | 42 | 0 | 0 | 44.5 | 0 | 0 | 36.4 | 1.06 | 0.969374741 | 0.866666667 | 0.924683571 | 0.817977528 | 0.895020012 |
| A0A3Q0D8G1 | Endothelin-converting enzyme 1 OS=Mesocricetus auratus OX=10036 GN=Ece1 PE=3 SV=1 | 6.714 | 4 | 2 | 6 | 2 | 769 | 86.9 | 5.9 | 13.13 | 2 | 29.8 | 224.7 | 140.3 | 34.9 | 215.9 | 142 | 29.8 | 157.2 | 151.5 | 0.961 | 0.993517407 | 0.85739615 | 0.803198676 | 0.861761711 | 0.801041714 |
| A0A3Q0CKF9 | probable ATP-dependent RNA helicase DDX17 isoform X1 OS=Mesocricetus auratus OX=10036 GN=LOC110340397 PE=4 SV=1 | 6.707 | 9 | 2 | 5 | 2 | 270 | 29.1 | 9.26 | 16.3 | 2 | 66.3 | 0 | 56.6 | 49.7 | 0 | 25.4 | 60.3 | 0 | 62.3 | 0.58 | 0.564963816 | 0.997558991 | 0.997420348 | 1.632490013 | 0.564261965 |
| A0A3Q0CSF3 | Calcyclin-binding protein OS=Mesocricetus auratus OX=10036 GN=Cacybp PE=4 SV=1 | 6.671 | 15 | 2 | 2 | 2 | 229 | 26.5 | 7.87 | 7.34 | 2 | 0 | 99.6 | 0 | 0 | 66.9 | 0 | 0 | 103.2 | 0 | 0.671 | 0.800516718 | 1.036144578 | 0.98117859 | 1.542600897 | 0.78486656 |
| A0A1U7R233 | importin-5 OS=Mesocricetus auratus OX=10036 GN=Ipo5 PE=4 SV=2 | 6.661 | 3 | 2 | 3 | 2 | 1097 | 123.6 | 4.92 | 6 | 2 | 0 | 107.3 | 43.2 | 0 | 85.6 | 53.5 | 0 | 116.8 | 30.2 | 0.994 | 0.928977649 | 0.976744186 | 0.981348823 | 1.056793674 | 0.954394565 |
| A0A3Q0CRK9 | ATP synthase peripheral stalk subunit OSCP OS=Mesocricetus auratus OX=10036 GN=Atp5o PE=3 SV=1 | 6.587 | 6 | 2 | 10 | 2 | 213 | 23.2 | 9.96 | 15.55 | 2 | 124.9 | 51.2 | 0 | 72.3 | 32.8 | 0 | 104.7 | 47.4 | 0 | 0.609 | 0.608944888 | 0.863713799 | 0.87395352 | 1.447193149 | 0.694683372 |
| A0A3Q0CFL5 | eukaryotic translation initiation factor 4B OS=Mesocricetus auratus OX=10036 GN=Eif4b PE=4 SV=1 | 6.552 | 4 | 2 | 3 | 2 | 611 | 69 | 5.67 | 10.77 | 2 | 109.6 | 39.9 | 178.4 | 138 | 26.4 | 178.9 | 100.7 | 37 | 161.6 | 1.03 | 0.936655194 | 0.912778286 | 0.868010268 | 0.871832217 | 0.813651477 |
| A0A3Q0CFV4 | UV excision repair protein RAD23 OS=Mesocricetus auratus OX=10036 GN=Rad23a PE=3 SV=1 | 6.471 | 4 | 1 | 9 | 1 | 390 | 42.5 | 4.54 | 32.62 | 1 | 44.8 | 83.4 | 90.3 | 46.3 | 88.4 | 104.2 | 47.7 | 98 | 104.4 | 1.06 | 0.776555652 | 1.144622426 | 0.669909935 | 1.04688154 | 0.888084225 |
| A0A1U7QWN6 | 26S proteasome non-ATPase regulatory subunit 9 OS=Mesocricetus auratus OX=10036 GN=Psmd9 PE=3 SV=2 | 6.385 | 10 | 2 | 4 | 2 | 222 | 24.5 | 6.65 | 8.42 | 2 | 106.4 | 0 | 0 | 100.2 | 0 | 0 | 99.2 | 0 | 0 | 0.941 | 0.968203126 | 0.932330827 | 0.96290899 | 0.99001996 | 0.994680897 |
| A0A3Q0CLK7 | PRKC apoptosis WT1 regulator protein OS=Mesocricetus auratus OX=10036 GN=Pawr PE=4 SV=1 | 6.323 | 8 | 2 | 4 | 2 | 288 | 32.5 | 4.79 | 10.29 | 2 | 0 | 278.5 | 242 | 0 | 192.2 | 166.7 | 0 | 194.7 | 178.4 | 0.689 | 0.641714811 | 0.716810759 | 0.673316277 | 1.039565339 | 0.959094034 |
| A0A3Q0CZ40 | xaa-Pro aminopeptidase 1 isoform X4 OS=Mesocricetus auratus OX=10036 GN=Xpnpep1 PE=3 SV=1 | 6.313 | 5 | 2 | 8 | 2 | 492 | 55 | 5.72 | 27.71 | 2 | 60.5 | 231.2 | 96.6 | 83.5 | 288.9 | 102.3 | 55.2 | 164 | 65.5 | 1.379 | 0.748795455 | 0.733195982 | 0.613793777 | 0.599747209 | 0.455182068 |
| A0A3Q0CUR5 | amphiphysin isoform X1 OS=Mesocricetus auratus OX=10036 GN=Amph PE=4 SV=1 | 6.309 | 4 | 3 | 12 | 2 | 866 | 94.3 | 4.6 | 33.56 | 3 | 338 | 300 | 554.6 | 284.5 | 238.4 | 493.1 | 325.6 | 263 | 479.6 | 0.842 | 0.625401394 | 0.895690089 | 0.706330178 | 1.051377953 | 0.872379134 |
| A0A1U8BU27 | Polyadenylate-binding protein OS=Mesocricetus auratus OX=10036 GN=Pabpc1 PE=3 SV=2 | 6.305 | 3 | 2 | 5 | 1 | 635 | 70.5 | 9.52 | 0 | 2 | 79.2 | 78.5 | 0 | 93.5 | 125.6 | 0 | 86 | 90.9 | 0 | 1.375 | 0.681566506 | 1.121750159 | 0.879312263 | 0.807393884 | 0.784238415 |
| A0A3Q0DBW5 | beta-catenin-like protein 1 isoform X1 OS=Mesocricetus auratus OX=10036 GN=Ctnnbl1 PE=4 SV=1 | 6.202 | 6 | 2 | 5 | 2 | 567 | 65.4 | 5.21 | 17.24 | 2 | 142.7 | 57.9 | 80.2 | 124.1 | 49 | 68.7 | 117.7 | 50.1 | 65.8 | 0.845 | 0.721181612 | 0.831908832 | 0.655472773 | 0.966087676 | 0.932660651 |
| A0A3Q0D3A4 | immunoglobulin lambda-1 light chain-like isoform X4 OS=Mesocricetus auratus OX=10036 GN=LOC101841371 PE=4 SV=1 | 6.194 | 18 | 3 | 8 | 3 | 232 | 25.1 | 5.21 | 7.49 | 3 | 225 | 248.1 | 262.2 | 343.8 | 318.5 | 341.2 | 246.2 | 288.2 | 307.2 | 1.284 | 0.003572745 | 1.144566843 | 0.182781437 | 0.838664674 | 0.078406835 |
| A0A3Q0CUT6 | V-type proton ATPase subunit C OS=Mesocricetus auratus OX=10036 GN=Atp6v1c1 PE=3 SV=1 | 6.126 | 7 | 2 | 5 | 2 | 382 | 43.9 | 7.46 | 15.08 | 2 | 73.9 | 63.4 | 158.9 | 60.9 | 55.9 | 154.7 | 67.1 | 64.6 | 161.6 | 0.902 | 0.86108289 | 0.990209318 | 0.983516219 | 1.080294659 | 0.880320286 |
| A0A3Q0C9B3 | 60 kDa poly(U)-binding-splicing factor OS=Mesocricetus auratus OX=10036 GN=Puf60 PE=3 SV=1 | 6.085 | 5 | 2 | 4 | 2 | 563 | 60.1 | 5.29 | 8.88 | 2 | 30.2 | 0 | 38.7 | 35.6 | 0 | 40.6 | 26.4 | 0 | 38.5 | 1.112 | 0.895328782 | 0.941944848 | 0.938899824 | 0.851706037 | 0.836629092 |
| A0A3Q0DDC2 | rho GTPase-activating protein 45 isoform X1 OS=Mesocricetus auratus OX=10036 GN=Arhgap45 PE=4 SV=1 | 6.017 | 2 | 2 | 4 | 2 | 1122 | 122.9 | 5.96 | 12.03 | 2 | 83.7 | 41.2 | 32.3 | 92.8 | 57.1 | 42.3 | 73.9 | 41 | 33.2 | 1.208 | 0.621318839 | 0.942111959 | 0.888152831 | 0.770551509 | 0.49415578 |
| A0A3Q0CNR5 | ubiquitin-like modifier-activating enzyme 6 OS=Mesocricetus auratus OX=10036 GN=Uba6 PE=3 SV=1 | 5.981 | 5 | 2 | 3 | 2 | 895 | 100.6 | 6.48 | 7.21 | 2 | 59.3 | 0 | 37.7 | 47.8 | 0 | 36.5 | 58.9 | 0 | 38.5 | 0.884 | 0.86048203 | 1.004123711 | 0.995912235 | 1.15539739 | 0.855859394 |
| A0A1U8CWG1 | serine/threonine-protein kinase 38 OS=Mesocricetus auratus OX=10036 GN=Stk38 PE=4 SV=2 | 5.977 | 6 | 2 | 2 | 2 | 465 | 54.1 | 7.34 | 4.64 | 2 | 0 | 56.9 | 0 | 0 | 41.3 | 0 | 0 | 62 | 0 | 0.726 | 0.836262747 | 1.089630931 | 0.954601717 | 1.501210654 | 0.796812383 |
| A0A3Q0D7Q4 | F-actin-capping protein subunit alpha OS=Mesocricetus auratus OX=10036 GN=Capza2 PE=3 SV=1 | 5.945 | 8 | 2 | 10 | 2 | 227 | 26 | 5.45 | 14.94 | 2 | 48 | 130 | 170.6 | 32.1 | 127.7 | 158.7 | 36.4 | 136 | 156.3 | 0.93 | 0.857658585 | 0.942914515 | 0.904104064 | 1.032025118 | 0.952059866 |

**Supplementary Table 1: List of differentially expressed proteins identified across the 4dpi infected and 14dpi infected**

| Accession | Description | Sum PEP Score | Coverage [%] | # Peptides | # PSMs | # Unique Peptides | # AAs | MW [kDa] | calc. pI | Score Sequest HT: Sequest HT | # Peptides (by Search Engine): Sequest HT | Mock-infected (4dpi) Replicate_R1 | Mock-infected (4dpi) Replicate_R2 | Mock-infected (4dpi) Replicate_R3 | SARS-CoV-2-infected (4dpi) Replicate_R1 | SARS-CoV-2-infected (4dpi) Replicate_R2 | SARS-CoV-2-infected (4dpi) Replicate_R3 | SARS-CoV-2-(14dpi) Replicate_R1 | SARS-CoV-2-infected (14dpi) Replicate_R2 | SARS-CoV-2-infected (14dpi) Replicate_R3 | Abundance Ratio: SARS-CoV-2-infected (4dpi) / Mock-infected (4dpi) | P-Value: SARS-CoV-2-infected (4dpi) / Mock-infected (4dpi) | Abundance Ratio: SARS-CoV-2-infected (14dpi) / Mock-infected (4dpi) | P-Value: SARS-CoV-2-infected (14dpi) / Mock-infected (4dpi) | Abundance Ratio: SARS-CoV-2-infected (14dpi) / SARS-CoV-2-infected (4dpi) | P-Value: SARS-CoV-2-infected (14dpi) / SARS-CoV-2-infected (4dpi) |
|---|---|---|---|---|---|---|---|---|---|---|---|---|---|---|---|---|---|---|---|---|---|---|---|---|---|
| A0A3Q0CQA1 | splicing factor 3A subunit 1 isoform X1 OS=Mesocricetus auratus OX=10036 GN=Sf3a1 PE=4 SV=1 | 5.929 | 4 | 2 | 4 | 2 | 799 | 89.3 | 5.29 | 9.22 | 2 | 0 | 101.1 | 39.1 | 0 | 90.3 | 31.5 | 0 | 99.7 | 44.3 | 0.848 | 0.884436515 | 1.027104137 | 0.976951331 | 1.18226601 | 0.859313545 |
| A0A3Q0D833 | Eukaryotic translation initiation factor 4H OS=Mesocricetus auratus OX=10036 GN=Eif4h PE=4 SV=1 | 5.911 | 9 | 2 | 8 | 2 | 248 | 27.3 | 7.23 | 21.8 | 2 | 57.7 | 137.9 | 45 | 68.5 | 151.3 | 35.8 | 46.6 | 87.1 | 33.7 | 1.024 | 0.917063938 | 0.695760599 | 0.514154475 | 0.654929577 | 0.497936683 |
| A0A3Q0CZ93 | Paraspeckle component 1 OS=Mesocricetus auratus OX=10036 GN=Pspc1 PE=3 SV=1 | 5.792 | 3 | 1 | 4 | 1 | 522 | 58.7 | 6.67 | 12.94 | 1 | 56 | 0 | 39.2 | 58.3 | 0 | 43.4 | 49.8 | 0 | 38.4 | 1.073 | 0.932712527 | 0.926470588 | 0.922128594 | 0.867256637 | 0.855119026 |
| A0A1U7RJT6 | activator of 90 kDa heat shock protein ATPase homolog 1 OS=Mesocricetus auratus OX=10036 GN=LOC101841164 PE=3 SV=2 | 5.751 | 4 | 1 | 2 | 1 | 338 | 38 | 5.43 | 7.75 | 1 | 0 | 0 | 87.5 | 0 | 0 | 99.3 | 0 | 0 | 78.4 | 1.135 | 0.93330727 | 0.896 | 0.942022458 | 0.789526687 | 0.877213895 |
| P62848 | 40S ribosomal protein S24 OS=Mesocricetus auratus OX=10036 GN=RPS24 PE=3 SV=1 | 5.727 | 9 | 2 | 3 | 2 | 133 | 15.4 | 10.78 | 9.81 | 2 | 61.8 | 56.1 | 92.7 | 52.1 | 56.7 | 67 | 50.9 | 61.4 | 83.3 | 1.01 | 0.421598104 | 0.928774929 | 0.753661787 | 1.112627986 | 0.577304872 |
| A0A3Q0CPZ2 | Sorting nexin OS=Mesocricetus auratus OX=10036 GN=Snx5 PE=3 SV=1 | 5.71 | 5 | 2 | 5 | 2 | 404 | 46.8 | 6.62 | 7.02 | 2 | 38.1 | 35.6 | 44.4 | 41.2 | 31.7 | 54.3 | 29.2 | 28.2 | 42.6 | 1.023 | 0.700074215 | 0.846740051 | 0.336257582 | 0.786163522 | 0.328337678 |
| A0A1U8BQV2 | Coatomer subunit delta OS=Mesocricetus auratus OX=10036 GN=Arcn1 PE=3 SV=2 | 5.704 | 6 | 3 | 6 | 3 | 511 | 57.2 | 6.21 | 4.26 | 3 | 230.6 | 459.4 | 192.1 | 187.3 | 373.5 | 148.9 | 208.4 | 400.6 | 147.7 | 0.813 | 0.625249929 | 0.857839247 | 0.730358062 | 1.066225166 | 0.886593173 |
| A0A3Q0CRB5 | Troponin T, fast skeletal muscle OS=Mesocricetus auratus OX=10036 GN=Tnnt3 PE=3 SV=1 | 5.665 | 10 | 2 | 6 | 2 | 239 | 28.3 | 9.36 | 16.01 | 2 | 134.1 | 65.5 | 134.3 | 215.8 | 282.8 | 445.7 | 82.1 | 35.3 | 77.9 | 3.398 | 0.084185609 | 0.58490566 | 0.177870639 | 0.206819867 | 0.06126346 |
| A0A1U8BJK2 | Ena/VASP-like protein OS=Mesocricetus auratus OX=10036 GN=Evl PE=3 SV=2 | 5.637 | 7 | 2 | 3 | 2 | 418 | 44.7 | 8.85 | 5.74 | 2 | 0 | 41.7 | 0 | 0 | 38.9 | 0 | 0 | 30.2 | 0 | 0.933 | 0.963204723 | 0.724220624 | 0.835207855 | 0.776349614 | 0.868858534 |
| A0A3Q0DAV2 | myc box-dependent-interacting protein 1 OS=Mesocricetus auratus OX=10036 GN=Bin1 PE=4 SV=1 | 5.604 | 3 | 2 | 9 | 1 | 603 | 66.3 | 5.45 | 19.91 | 2 | 178.1 | 0 | 0 | 115.6 | 0 | 0 | 148.8 | 0 | 0 | 0.649 | 0.785414085 | 0.835485682 | 0.905808791 | 1.287197232 | 0.869197202 |
| A0A3Q0CL92 | Platelet-activating factor acetylhydrolase IB subunit alpha OS=Mesocricetus auratus OX=10036 GN=Pafah1b1 PE=3 SV=1 | 5.504 | 4 | 2 | 5 | 1 | 410 | 46.6 | 7.37 | 13.73 | 2 | 0 | 0 | 51.2 | 0 | 0 | 58 | 0 | 0 | 37.4 | 1.132 | 0.934246936 | 0.73046875 | 0.839286917 | 0.644827586 | 0.782570263 |
| A0A1U8CMU8 | elongin-B OS=Mesocricetus auratus OX=10036 GN=Elob PE=4 SV=2 | 5.446 | 14 | 2 | 4 | 2 | 149 | 16.7 | 9.19 | 10.18 | 2 | 75 | 0 | 289.4 | 100.2 | 0 | 258.1 | 95.9 | 0 | 276.7 | 1.092 | 0.986725145 | 1.022502744 | 0.982742203 | 1.039910689 | 0.967689636 |
| A0A1U8BQY9 | actin-like protein 6A OS=Mesocricetus auratus OX=10036 GN=Actl6a PE=3 SV=2 | 5.331 | 3 | 1 | 10 | 1 | 429 | 47.4 | 5.6 | 35.27 | 1 | 0 | 32.9 | 53.8 | 0 | 36.8 | 48.7 | 0 | 31.7 | 49.4 | 1.007 | 0.986018149 | 0.935409458 | 0.934426126 | 0.948538012 | 0.946615665 |
| A0A3Q0DIG1 | Importin subunit alpha OS=Mesocricetus auratus OX=10036 GN=Kpna4 PE=3 SV=1 | 5.315 | 6 | 2 | 5 | 2 | 521 | 57.9 | 4.98 | 8.48 | 2 | 41.1 | 41.3 | 0 | 36.3 | 35.7 | 0 | 42.5 | 47.7 | 0 | 0.873 | 0.858664184 | 1.094660194 | 0.904866457 | 1.252777778 | 0.769679788 |
| A0A3Q0DFP8 | RNA-binding motif protein, X chromosome isoform X1 OS=Mesocricetus auratus OX=10036 GN=Rbmx PE=4 SV=1 | 5.279 | 4 | 2 | 8 | 2 | 414 | 45.1 | 10.02 | 19.54 | 2 | 150.5 | 117.8 | 396.3 | 147.3 | 127.7 | 377 | 147.6 | 100.1 | 355.1 | 0.979 | 0.973521628 | 0.907011736 | 0.869679285 | 0.924539877 | 0.890631772 |
| A0A3Q0D0S9 | Non-specific serine/threonine protein kinase OS=Mesocricetus auratus OX=10036 GN=Slk PE=4 SV=1 | 5.277 | 2 | 2 | 2 | 2 | 1192 | 136.4 | 5.11 | 2.29 | 2 | 102.5 | 0 | 0 | 74 | 0 | 0 | 85.7 | 0 | 0 | 0.722 | 0.833723503 | 0.836097561 | 0.906183417 | 1.158108108 | 0.922773518 |
| A0A3Q0DAQ0 | LEM domain-containing protein 2 OS=Mesocricetus auratus OX=10036 GN=Lemd2 PE=4 SV=1 | 5.275 | 4 | 2 | 10 | 2 | 501 | 56.3 | 8.81 | 26.77 | 2 | 601.5 | 614.2 | 462.8 | 535.9 | 536.3 | 417.3 | 434.4 | 498.2 | 386 | 0.891 | 0.373304101 | 0.785582365 | 0.119092194 | 0.885263511 | 0.330616196 |
| A0A3Q0D353 | Dynamin-1-like protein OS=Mesocricetus auratus OX=10036 GN=Dnm1l PE=3 SV=1 | 5.19 | 4 | 2 | 2 | 2 | 736 | 81.8 | 6.81 | 4.87 | 2 | 45.1 | 0 | 0 | 28.6 | 0 | 0 | 38.8 | 0 | 0 | 0.634 | 0.775407344 | 0.860310421 | 0.92087228 | 1.356643357 | 0.843612135 |
| A0A3Q0DB58 | cell division cycle protein 16 homolog isoform X2 OS=Mesocricetus auratus OX=10036 GN=Cdc16 PE=4 SV=1 | 5.16 | 5 | 2 | 6 | 2 | 553 | 63.1 | 5.35 | 20.02 | 2 | 359.5 | 139.6 | 349.9 | 236.3 | 106.4 | 270.1 | 320.3 | 107.5 | 286.8 | 0.696 | 0.42430306 | 0.841696113 | 0.670004088 | 1.166122715 | 0.704376942 |
| A0A3Q0DCI2 | trafficking protein particle complex subunit 8 isoform X2 OS=Mesocricetus auratus OX=10036 GN=Trappc8 PE=4 SV=1 | 5.151 | 2 | 2 | 4 | 2 | 1429 | 160 | 6.84 | 10.61 | 2 | 190.3 | 0 | 0 | 141.8 | 0 | 0 | 158.1 | 0 | 0 | 0.745 | 0.848818224 | 0.830793484 | 0.902930474 | 1.114950635 | 0.942548239 |
| A0A3Q0CJR0 | SH3 and multiple ankyrin repeat domains protein 3 OS=Mesocricetus auratus OX=10036 GN=Shank3 PE=3 SV=1 | 5.124 | 1 | 2 | 4 | 2 | 1811 | 194.2 | 9.29 | 6.06 | 2 | 0 | 71 | 324.7 | 0 | 55.5 | 211.4 | 0 | 61.6 | 260.4 | 0.713 | 0.735526633 | 0.813747789 | 0.855411376 | 1.206444361 | 0.864774355 |
| A0A3Q0CS69 | transmembrane protein 43 OS=Mesocricetus auratus OX=10036 GN=Tmem43 PE=3 SV=1 | 5.115 | 12 | 2 | 5 | 2 | 194 | 21.6 | 6.8 | 16.17 | 2 | 0 | 69.5 | 84.1 | 0 | 42.7 | 84.8 | 0 | 64.2 | 76.1 | 0.787 | 0.819346461 | 0.913411458 | 0.905622667 | 1.100392157 | 0.906271228 |
| A0A1U7RCY0 | Transmembrane 9 superfamily member OS=Mesocricetus auratus OX=10036 GN=Tm9sf4 PE=3 SV=2 | 5.108 | 5 | 2 | 3 | 2 | 643 | 74.7 | 7.08 | 8.01 | 2 | 91.7 | 0 | 0 | 79.3 | 0 | 0 | 78.3 | 0 | 0 | 0.865 | 0.923552148 | 0.85387132 | 0.916992111 | 0.98738966 | 0.993270237 |
| A0A3Q0CNT5 | LOW QUALITY PROTEIN: casein kinase II subunit alpha OS=Mesocricetus auratus OX=10036 GN=Csnk2a1 PE=3 SV=1 | 4.983 | 4 | 1 | 2 | 1 | 387 | 44.4 | 7.47 | 8.15 | 1 | 174.7 | 0 | 158.8 | 157.6 | 0 | 169 | 193.4 | 0 | 172.9 | 0.98 | 0.977890012 | 1.098350825 | 0.901498916 | 1.121555419 | 0.879823034 |
| A0A3Q0CDC7 | Tyrosine-protein phosphatase non-receptor type OS=Mesocricetus auratus OX=10036 GN=Ptpn6 PE=3 SV=1 | 4.953 | 4 | 2 | 8 | 2 | 595 | 67.6 | 7.94 | 9.89 | 2 | 83.7 | 39.5 | 37.9 | 85.2 | 62.2 | 55.6 | 102.9 | 47.4 | 54 | 1.346 | 0.47839011 | 1.268156425 | 0.566865672 | 1.006403941 | 0.983813902 |
| A0A3Q0CI19 | serine/threonine-protein kinase OSR1-like isoform X1 OS=Mesocricetus auratus OX=10036 GN=LOC106022793 PE=4 SV=1 | 4.947 | 7 | 1 | 3 | 1 | 153 | 17.5 | 8.94 | 10.67 | 1 | 39.2 | 0 | 0 | 29.7 | 0 | 0 | 37 | 0 | 0 | 0.759 | 0.856893801 | 0.943877551 | 0.969406831 | 1.245791246 | 0.885501096 |
| A0A3Q0CV77 | lysine--tRNA ligase OS=Mesocricetus auratus OX=10036 GN=Kars PE=4 SV=1 | 4.821 | 13 | 2 | 4 | 2 | 241 | 27 | 4.88 | 2.22 | 2 | 0 | 39.9 | 0 | 0 | 53.5 | 0 | 0 | 41.1 | 0 | 1.341 | 0.849242286 | 1.030075188 | 0.984290514 | 0.768224299 | 0.863674139 |
| A0A3Q0D557 | Histone deacetylase OS=Mesocricetus auratus OX=10036 GN=Hdac1 PE=3 SV=1 | 4.763 | 4 | 2 | 3 | 2 | 482 | 55 | 5.54 | 7.53 | 2 | 80.7 | 0 | 71.9 | 71.5 | 0 | 68 | 64.8 | 0 | 70.6 | 0.916 | 0.905621327 | 0.887287025 | 0.874887174 | 0.970609319 | 0.968435681 |
| A0A3Q0D7X3 | sorting nexin-6 OS=Mesocricetus auratus OX=10036 GN=Snx6 PE=3 SV=1 | 4.734 | 6 | 2 | 4 | 2 | 341 | 39 | 6.11 | 0 | 2 | 0 | 156.4 | 99.5 | 0 | 141.5 | 87.8 | 0 | 151.7 | 104.6 | 0.894 | 0.892509224 | 1.001563111 | 0.998437995 | 1.117749673 | 0.889734526 |
| A0A3Q0CYQ8 | cytosolic purine 5'-nucleotidase OS=Mesocricetus auratus OX=10036 GN=Nt5c2 PE=3 SV=1 | 4.648 | 7 | 2 | 2 | 2 | 370 | 43 | 6.05 | 2.46 | 2 | 0 | 0 | 46.2 | 0 | 0 | 95.4 | 0 | 0 | 36.7 | 2.065 | 0.67526044 | 0.794372294 | 0.8802696 | 0.384696017 | 0.611981615 |
| A0A3Q0CE29 | 60S ribosomal protein L18a OS=Mesocricetus auratus OX=10036 GN=Rpl18a PE=4 SV=1 | 4.615 | 19 | 2 | 7 | 2 | 84 | 10 | 11.17 | 19.41 | 2 | 227.7 | 81.3 | 375 | 220.2 | 81.5 | 390.1 | 221.3 | 99.7 | 319.9 | 1.04 | 0.984162124 | 0.936988304 | 0.899251836 | 0.926423822 | 0.885251722 |
| A0A3Q0CX56 | Phosphorylase b kinase regulatory subunit OS=Mesocricetus auratus OX=10036 GN=Phka2 PE=3 SV=1 | 4.605 | 2 | 2 | 2 | 2 | 1235 | 138.4 | 6.48 | 5.87 | 2 | 238.5 | 0 | 0 | 219.4 | 0 | 0 | 197.6 | 0 | 0 | 0.92 | 0.955846615 | 0.82851153 | 0.901527172 | 0.900638104 | 0.944725675 |
| A0A3Q0D827 | ubiquitin-conjugating enzyme E2 J1 OS=Mesocricetus auratus OX=10036 GN=Ube2j1 PE=4 SV=1 | 4.588 | 5 | 1 | 1 | 1 | 278 | 30.8 | 8.16 | 3.91 | 1 | 0 | 39.4 | 0 | 0 | 34.9 | 0 | 0 | 31.7 | 0 | 0.887 | 0.936032662 | 0.804568528 | 0.886668999 | 0.908309456 | 0.949173338 |
| A0A1U7QXV8 | Vesicle-fusing ATPase OS=Mesocricetus auratus OX=10036 GN=LOC101831032 PE=3 SV=2 | 4.569 | 4 | 2 | 7 | 2 | 485 | 54.2 | 9.14 | 16.66 | 2 | 186.2 | 95.3 | 56.8 | 186.3 | 83.6 | 66.1 | 153.7 | 84.3 | 56.1 | 1 | 0.989281239 | 0.869346734 | 0.775680949 | 0.875297619 | 0.783786531 |
| A0A3Q0CGK2 | Methionine aminopeptidase 2 OS=Mesocricetus auratus OX=10036 GN=Metap2 PE=3 SV=1 | 4.548 | 3 | 1 | 4 | 1 | 478 | 52.9 | 5.82 | 15.36 | 1 | 88.3 | 120.8 | 62.5 | 74.8 | 119.1 | 53.2 | 109.7 | 109.7 | 54.4 | 0.986 | 0.766884127 | 1.008100147 | 0.978001681 | 1.10805342 | 0.756172171 |
| A0A3Q0DEV8 | dynein light chain roadblock-type 1 isoform X3 OS=Mesocricetus auratus OX=10036 GN=Dynlrb1 PE=3 SV=1 | 4.539 | 19 | 1 | 3 | 1 | 63 | 7.4 | 7.25 | 8.47 | 1 | 46.3 | 0 | 36.4 | 32.5 | 0 | 31.6 | 41.4 | 0 | 26.8 | 0.781 | 0.744636324 | 0.824667473 | 0.807840307 | 1.063962559 | 0.936729314 |
| A0A3Q0DBT4 | SH3 domain-binding glutamic acid-rich-like protein OS=Mesocricetus auratus OX=10036 GN=Sh3bgrl PE=3 SV=1 | 4.534 | 7 | 1 | 6 | 1 | 146 | 16 | 7.83 | 19.81 | 1 | 161.3 | 215 | 320.5 | 266.1 | 325 | 460.9 | 98.6 | 136 | 184.3 | 1.523 | 0.189058582 | 0.601176808 | 0.177126148 | 0.398193916 | 0.0506411 |
| A0A3Q0D7P5 | protein NipSnap homolog 2 isoform X2 OS=Mesocricetus auratus OX=10036 GN=Nipsnap2 PE=4 SV=1 | 4.494 | 5 | 1 | 4 | 1 | 225 | 26.1 | 9.04 | 4.43 | 1 | 105.3 | 41.7 | 76.9 | 96.7 | 47.6 | 66.8 | 94.2 | 41.2 | 82.8 | 0.888 | 0.864074482 | 0.974542206 | 0.941851879 | 1.033633349 | 0.917831806 |
| A0A3Q0CI06 | transmembrane emp24 domain-containing protein 4 OS=Mesocricetus auratus OX=10036 GN=LOC101825465 PE=3 SV=1 | 4.48 | 20 | 2 | 3 | 2 | 131 | 14.8 | 7.11 | 0 | 2 | 59.8 | 90.7 | 0 | 45.2 | 72.7 | 0 | 54.5 | 107.9 | 0 | 0.779 | 0.766206438 | 1.079069767 | 0.927644003 | 1.377438507 | 0.716421253 |
| A0A3Q0CCZ5 | F-BAR domain only protein 2 OS=Mesocricetus auratus OX=10036 GN=Fcho2 PE=3 SV=1 | 4.472 | 3 | 1 | 5 | 1 | 809 | 88.7 | 6.89 | 14.25 | 1 | 36.6 | 33 | 0 | 38 | 40.9 | 0 | 33.8 | 29 | 0 | 1.135 | 0.868763425 | 0.902298851 | 0.892385215 | 0.795944233 | 0.76722264 |
| A0A3Q0CLQ6 | GTPase HRas isoform X2 OS=Mesocricetus auratus OX=10036 GN=Hras PE=4 SV=1 | 4.46 | 12 | 2 | 4 | 1 | 170 | 18.8 | 5 | 7.67 | 2 | 0 | 132.2 | 336.2 | 0 | 109.6 | 261.5 | 0 | 97.7 | 347.7 | 0.803 | 0.806937328 | 0.95089667 | 0.95965668 | 1.200215575 | 0.857172686 |
| A0A3Q0CFQ0 | Protein quaking OS=Mesocricetus auratus OX=10036 GN=Qki PE=4 SV=1 | 4.453 | 6 | 1 | 9 | 1 | 287 | 31.6 | 9.11 | 25.18 | 1 | 163.9 | 30.4 | 44.1 | 135.7 | 22.8 | 26.9 | 131.2 | 44 | 40.8 | 0.751 | 0.769458206 | 0.906040268 | 0.892972893 | 1.165048544 | 0.840515425 |
| A0A1U8CFT4 | protein LSM14 homolog A OS=Mesocricetus auratus OX=10036 GN=Lsm14a PE=3 SV=2 | 4.431 | 5 | 2 | 7 | 2 | 465 | 50.7 | 9.6 | 14.72 | 2 | 485.2 | 667.7 | 548.1 | 364 | 436.5 | 348.4 | 348.1 | 446.2 | 501.1 | 0.671 | 0.055561932 | 0.761552028 | 0.126906902 | 1.127513274 | 0.413971335 |
| A0A3Q0CFS7 | 2-oxoisovalerate dehydrogenase subunit alpha OS=Mesocricetus auratus OX=10036 GN=Bckdha PE=3 SV=1 | 4.387 | 2 | 1 | 5 | 1 | 417 | 47.5 | 7.09 | 12.7 | 1 | 78 | 74.7 | 76.5 | 55.9 | 49.2 | 46.3 | 55.3 | 58.5 | 76.1 | 0.659 | 0.006841577 | 0.828534031 | 0.177593726 | 1.254293263 | 0.175317234 |
| A0A3Q0CK75 | beta-parvin OS=Mesocricetus auratus OX=10036 GN=Parvb PE=3 SV=1 | 4.383 | 3 | 1 | 3 | 1 | 366 | 41.7 | 6.43 | 8.86 | 1 | 60 | 55.6 | 57.7 | 36.6 | 38.1 | 53.5 | 55.9 | 56.6 | 50.9 | 0.699 | 0.101468883 | 0.94287363 | 0.215437674 | 1.274570983 | 0.152054193 |
| A0A3Q0CWU9 | ataxin-2-like protein isoform X6 OS=Mesocricetus auratus OX=10036 GN=Atxn2l PE=3 SV=1 | 4.372 | 2 | 1 | 2 | 1 | 1064 | 111.9 | 8.69 | 11.88 | 1 | 0 | 0 | 68.5 | 0 | 0 | 55.2 | 0 | 0 | 75.8 | 0.806 | 0.887463692 | 1.106569343 | 0.946500878 | 1.373188406 | 0.837828097 |
| A0A1U7RA50 | 3-hydroxy-3-methylglutarate-CoA lyase OS=Mesocricetus auratus OX=10036 GN=Hmgcl PE=3 SV=2 | 4.29 | 4 | 1 | 3 | 1 | 325 | 34.2 | 8.41 | 8.26 | 1 | 0 | 0 | 103.8 | 0 | 0 | 84.4 | 0 | 0 | 83.4 | 0.813 | 0.891992323 | 0.803468208 | 0.885980431 | 0.988151659 | 0.993679282 |
| A0A3Q0D0T5 | striatin-3 OS=Mesocricetus auratus OX=10036 GN=Strn3 PE=3 SV=1 | 4.265 | 4 | 2 | 7 | 2 | 796 | 87 | 5.29 | 18.47 | 2 | 381.9 | 322.9 | 377.3 | 311.2 | 273.5 | 311.3 | 336.5 | 275.4 | 319.7 | 0.815 | 0.061286778 | 0.860918584 | 0.129103404 | 1.039732143 | 0.623826607 |
| A0A1U7R741 | Ribosome maturation protein SBDS OS=Mesocricetus auratus OX=10036 GN=LOC101837582 PE=3 SV=2 | 4.256 | 8 | 2 | 5 | 2 | 237 | 27.3 | 9.23 | 2.22 | 2 | 47 | 135.3 | 123.8 | 40.8 | 107.1 | 101 | 51.7 | 130.2 | 126.2 | 0.816 | 0.61547936 | 1.006533812 | 0.986736609 | 1.237846525 | 0.584786817 |
| A0A3Q0DBL8 | Calcium/calmodulin-dependent protein kinase OS=Mesocricetus auratus OX=10036 GN=Camk2d PE=3 SV=1 | 4.206 | 2 | 1 | 2 | 1 | 503 | 56 | 6.49 | 6.34 | 1 | 44.4 | 0 | 35.2 | 35.8 | 0 | 44.1 | 46.5 | 0 | 33.2 | 1.004 | 0.996080417 | 1.001256281 | 0.998707743 | 0.997496871 | 0.997415548 |
| A0A3Q0CF74 | Fibulin-1 OS=Mesocricetus auratus OX=10036 GN=Fbln1 PE=3 SV=1 | 4.156 | 3 | 2 | 5 | 2 | 704 | 77.9 | 5.1 | 14.33 | 2 | 208.1 | 75.7 | 65.5 | 176.3 | 71.3 | 66 | 182.7 | 79.8 | 58.8 | 0.948 | 0.848785109 | 0.919839679 | 0.883748727 | 1.024553571 | 0.963355951 |
| A0A3Q0CVN6 | myristoylated alanine-rich C-kinase substrate OS=Mesocricetus auratus OX=10036 GN=Marcks PE=3 SV=1 | 4.132 | 7 | 2 | 2 | 2 | 317 | 30.2 | 4.35 | 1.65 | 2 | 77 | 0 | 0 | 109.7 | 0 | 0 | 74.1 | 0 | 0 | 1.425 | 0.8205576 | 0.962337662 | 0.979651878 | 0.675478578 | 0.803040904 |
| A0A3Q0CF38 | rho guanine nucleotide exchange factor 1 OS=Mesocricetus auratus OX=10036 GN=Arhgef1 PE=4 SV=1 | 4.126 | 2 | 1 | 2 | 1 | 968 | 108.9 | 7.33 | 8.24 | 1 | 86.8 | 0 | 127.4 | 70 | 0 | 117.8 | 108 | 0 | 120.9 | 0.863 | 0.87098699 | 1.068627451 | 0.931654274 | 1.21884984 | 0.803069335 |
| A0A1U8BPT2 | Peptidyl-prolyl cis-trans isomerase OS=Mesocricetus auratus OX=10036 GN=Ppil3 PE=3 SV=2 | 4.094 | 7 | 1 | 6 | 1 | 158 | 17.5 | 8.66 | 15.76 | 1 | 139.5 | 134 | 188.2 | 141.1 | 122.1 | 160.8 | 132 | 151.8 | 149 | 0.966 | 0.578675045 | 0.937405241 | 0.641552334 | 1.020754717 | 0.832567182 |
| A0A3Q0CQ14 | THO complex subunit 4 OS=Mesocricetus auratus OX=10036 GN=Alyref PE=4 SV=1 | 3.95 | 4 | 1 | 4 | 1 | 255 | 27 | 11.15 | 18.05 | 1 | 228 | 121.3 | 131.2 | 222.3 | 131.4 | 111.6 | 166.8 | 119.5 | 86.3 | 0.975 | 0.92129047 | 0.775442248 | 0.438659699 | 0.800773694 | 0.501153995 |
| A0A3Q0D3W2 | interferon-activable protein 203-like OS=Mesocricetus auratus OX=10036 GN=LOC101842177 PE=3 SV=1 | 3.94 | 3 | 1 | 1 | 1 | 336 | 38.2 | 9.73 | 3.63 | 1 | 0 | 0 | 26.9 | 0 | 0 | 90.3 | 0 | 0 | 24.8 | 3.355 | 0.561104739 | 0.921933086 | 0.956999329 | 0.274640089 | 0.54828893 |
| A0A3Q0DAQ4 | Exocyst complex component Sec8 OS=Mesocricetus auratus OX=10036 GN=Exoc4 PE=3 SV=1 | 3.938 | 1 | 1 | 2 | 1 | 975 | 110.7 | 6.58 | 5.33 | 1 | 0 | 66.6 | 109.3 | 0 | 67.6 | 104.6 | 0 | 67.6 | 79.5 | 0.985 | 0.979053698 | 0.836270608 | 0.824098621 | 0.854239257 | 0.842561545 |
| A0A3Q0CK73 | Phenylalanyl-tRNA synthetase beta subunit OS=Mesocricetus auratus OX=10036 GN=Farsb PE=3 SV=1 | 3.886 | 4 | 2 | 2 | 2 | 589 | 65.9 | 7.21 | 3.45 | 2 | 0 | 130.8 | 0 | 0 | 113.2 | 0 | 0 | 80.8 | 0 | 0.866 | 0.923951572 | 0.617737003 | 0.76438384 | 0.713780919 | 0.82836771 |
| A0A3Q0D180 | Epithelial cell adhesion molecule OS=Mesocricetus auratus OX=10036 GN=Epcam PE=3 SV=1 | 3.876 | 4 | 1 | 4 | 1 | 318 | 35.5 | 6.38 | 8.25 | 1 | 0 | 82.2 | 0 | 0 | 65.6 | 0 | 0 | 74.9 | 0 | 0.797 | 0.882584841 | 0.911192214 | 0.950837102 | 1.141768293 | 0.930147103 |
| A0A3Q0D5U4 | Carboxylic ester hydrolase OS=Mesocricetus auratus OX=10036 GN=LOC101825337 PE=3 SV=1 | 3.869 | 3 | 1 | 5 | 1 | 427 | 47.2 | 5.83 | 14.63 | 1 | 190.8 | 243 | 309.1 | 150.2 | 148.9 | 184.3 | 144.9 | 234.6 | 277.6 | 0.693 | 0.115625471 | 0.884506663 | 0.611790664 | 1.359329748 | 0.273799912 |
| A0A3Q0DBM1 | nuclear autoantigen Sp-100-like OS=Mesocricetus auratus OX=10036 GN=LOC106022168 PE=4 SV=1 | 3.852 | 8 | 1 | 5 | 1 | 189 | 21.4 | 5.02 | 18.45 | 1 | 42.5 | 77.9 | 35 | 90.4 | 191.5 | 59.6 | 33.2 | 55.7 | 29.4 | 2.185 | 0.255775843 | 0.761261261 | 0.479542812 | 0.346412884 | 0.198903506 |
| A0A3Q0D5F1 | protein canopy homolog 2 OS=Mesocricetus auratus OX=10036 GN=Cnpy2 PE=3 SV=1 | 3.831 | 8 | 1 | 4 | 1 | 193 | 22.1 | 5.26 | 14.31 | 1 | 43.5 | 0 | 0 | 34 | 0 | 0 | 34.6 | 0 | 0 | 0.782 | 0.872201442 | 0.795402299 | 0.880917944 | 1.017647059 | 0.990723912 |
| A0A3Q0CFI6 | ATPase Asna1 OS=Mesocricetus auratus OX=10036 GN=Asna1 PE=3 SV=1 | 3.805 | 5 | 2 | 2 | 2 | 348 | 38.8 | 4.91 | 3.17 | 2 | 29.2 | 0 | 41.6 | 25.3 | 0 | 32.8 | 29.9 | 0 | 34.9 | 0.827 | 0.802907639 | 0.915254237 | 0.909207782 | 1.115318417 | 0.886941455 |
| A0A1U7Q3D8 | protein disulfide-isomerase TMX3 OS=Mesocricetus auratus OX=10036 GN=Tmx3 PE=4 SV=2 | 3.766 | 4 | 2 | 4 | 2 | 486 | 54.8 | 5.22 | 7.16 | 2 | 41.6 | 51.8 | 50.8 | 46.1 | 58.3 | 54.6 | 45.6 | 50.2 | 40.5 | 1.124 | 0.367723877 | 0.945214979 | 0.573003122 | 0.857232704 | 0.177559831 |

Supplementary Table 1: List of differentially expressed proteins identified across the 4dpi infected and 14dpi infected

| Accession | Description | Sum PEP Score | Coverage [%] | # Peptides | # PSMs | # Unique Peptides | # AAs | MW [kDa] | calc. pI | Score Sequest HT: Sequest HT | # Peptides (by Search Engine): Sequest HT | Mock-infected (4dpi) Replicate_R1 | Mock-infected (4dpi) Replicate_R2 | Mock-infected (4dpi) Replicate_R3 | SARS-CoV-2-infected (4dpi) Replicate_R1 | SARS-CoV-2-infected (4dpi) Replicate_R2 | SARS-CoV-2-infected (4dpi) Replicate_R3 | SARS-CoV-2-(14dpi) Replicate_R1 | SARS-CoV-2-infected (14dpi) Replicate_R2 | SARS-CoV-2-infected (14dpi) Replicate_R3 | Abundance Ratio: SARS-CoV-2-infected (4dpi) / Mock-infected (4dpi) | P-Value: SARS-CoV-2-infected (4dpi) / Mock-infected (4dpi) | Abundance Ratio: SARS-CoV-2-infected (14dpi) / Mock-infected (4dpi) | P-Value: SARS-CoV-2-infected (14dpi) / Mock-infected (4dpi) | Abundance Ratio: SARS-CoV-2-infected (14dpi) / SARS-CoV-2-infected (4dpi) | P-Value: SARS-CoV-2-infected (14dpi) / SARS-CoV-2-infected (4dpi) |
|---|---|---|---|---|---|---|---|---|---|---|---|---|---|---|---|---|---|---|---|---|---|---|---|---|---|---|
| A0A1U7QMI0 | Peroxidase OS=Mesocricetus auratus OX=10036 GN=Pxdn PE=4 SV=2 | 3.762 | 2 | 2 | 3 | 2 | 1451 | 162.2 | 7.08 | 1.94 | 2 | 0 | 78.3 | 0 | 0 | 79.1 | 0 | 0 | 79.8 | 0 | 1.01 | 0.994609258 | 1.019157088 | 0.989937861 | 1.008849558 | 0.995327593 |
| A0A3Q0D0V7 | Small acidic protein OS=Mesocricetus auratus OX=10036 GN=CUNH11orf58 PE=3 SV=1 | 3.752 | 5 | 1 | 7 | 1 | 152 | 17 | 4.67 | 19.49 | 1 | 361.8 | 343.8 | 532.1 | 314 | 253.8 | 483.4 | 320.9 | 366.5 | 539.8 | 0.909 | 0.533661576 | 0.991516523 | 0.970763038 | 1.167427702 | 0.573266139 |
| A0A3Q0D7W2 | pre-B-cell leukemia transcription factor-interacting protein 1 isoform X2 OS=Mesocricetus auratus OX=10036 GN=Pbxip1 PE=4 SV=1 | 3.728 | 3 | 1 | 3 | 1 | 784 | 87.8 | 5.36 | 4.68 | 1 | 48 | 40.9 | 0 | 52.6 | 38 | 0 | 40.7 | 36.4 | 0 | 1.009 | 0.980391358 | 0.867266592 | 0.852114124 | 0.850993377 | 0.835870873 |
| A0A3Q0CH57 | Complex I-MNLL OS=Mesocricetus auratus OX=10036 GN=Ndufb1 PE=3 SV=1 | 3.692 | 19 | 1 | 4 | 1 | 58 | 7 | 8.21 | 2.4 | 1 | 149.5 | 87.7 | 103.2 | 151 | 78.7 | 79.9 | 145.2 | 86.7 | 100.1 | 0.954 | 0.752488158 | 0.975323149 | 0.918330386 | 1.072351421 | 0.815127812 |
| A0A3Q0CGX8 | Nuclear cap-binding protein subunit 3 OS=Mesocricetus auratus OX=10036 GN=Ncbp3 PE=3 SV=1 | 3.666 | 7 | 1 | 1 | 1 | 258 | 29.2 | 9.42 | 4.45 | 1 | 0 | 0 | 41.6 | 0 | 0 | 29.3 | 0 | 0 | 36.8 | 0.705 | 0.822148355 | 0.884615385 | 0.935342401 | 1.255972696 | 0.88141704 |
| A0A3Q0DBP5 | collagen alpha-4(IV) chain OS=Mesocricetus auratus OX=10036 GN=Col4a4 PE=4 SV=1 | 3.607 | 1 | 1 | 3 | 1 | 1706 | 166.4 | 8.31 | 8.41 | 1 | 42.6 | 54.8 | 58.5 | 36.9 | 47.5 | 47.1 | 36.4 | 47.9 | 48.4 | 0.861 | 0.248330623 | 0.851186658 | 0.282831523 | 1.009125475 | 0.942801013 |
| A0A3Q0DG87 | collagen alpha-1(V) chain isoform X4 OS=Mesocricetus auratus OX=10036 GN=Col5a1 PE=4 SV=1 | 3.577 | 0 | 1 | 3 | 1 | 1796 | 179.1 | 4.97 | 9.41 | 1 | 85.8 | 79.5 | 73.2 | 57.5 | 43.7 | 47.5 | 72.9 | 56.9 | 69.6 | 0.645 | 0.005750033 | 0.8360587 | 0.104432765 | 1.340954943 | 0.058758035 |
| A0A3Q0CFQ5 | Calpain inhibitor OS=Mesocricetus auratus OX=10036 GN=Cast PE=3 SV=1 | 3.518 | 3 | 1 | 3 | 1 | 755 | 81.4 | 5.19 | 11.27 | 1 | 0 | 77.9 | 174.8 | 0 | 81.8 | 184.7 | 0 | 106.6 | 206.2 | 1.053 | 0.953148183 | 1.237831421 | 0.810544615 | 1.173733583 | 0.85652462 |
| A0A3Q0CG89 | golgin subfamily A member 2 OS=Mesocricetus auratus OX=10036 GN=LOC101831552 PE=4 SV=1 | 3.493 | 2 | 2 | 2 | 2 | 1030 | 116.5 | 5.02 | 3.9 | 2 | 0 | 0 | 117.6 | 0 | 0 | 120.7 | 0 | 0 | 91.3 | 1.026 | 0.986204778 | 0.776360544 | 0.868865492 | 0.756420878 | 0.856100961 |
| A0A1U7QEG5 | septin-1 OS=Mesocricetus auratus OX=10036 GN=Sept1 PE=3 SV=2 | 3.486 | 5 | 2 | 2 | 1 | 390 | 44.5 | 6.87 | 3.37 | 2 | 0 | 0 | 84.7 | 0 | 0 | 72.6 | 0 | 0 | 61.5 | 0.858 | 0.918965923 | 0.72609209 | 0.836430822 | 0.847107438 | 0.912895678 |
| A0A1U8D286 | scaffold attachment factor B1 isoform X1 OS=Mesocricetus auratus OX=10036 GN=LOC101830764 PE=4 SV=2 | 3.455 | 2 | 1 | 11 | 1 | 611 | 69.9 | 9.73 | 30.29 | 1 | 479.8 | 510.2 | 403.8 | 577.4 | 652.4 | 411.5 | 370.1 | 425.1 | 324.5 | 1.215 | 0.373266078 | 0.803343378 | 0.101152102 | 0.682203132 | 0.120446179 |
| A0A3Q0DDP9 | lysophosphatidylcholine acyltransferase 1 OS=Mesocricetus auratus OX=10036 GN=Lpcat1 PE=4 SV=1 | 3.446 | 4 | 2 | 2 | 2 | 534 | 59.9 | 6.11 | 4.6 | 2 | 56.7 | 0 | 0 | 56.7 | 0 | 0 | 34.6 | 0 | 0 | 0.999 | 1 | 0.610229277 | 0.759335069 | 0.610229277 | 0.759335069 |
| A0A3Q0CSJ6 | Tyrosine-protein phosphatase non-receptor type OS=Mesocricetus auratus OX=10036 GN=Ptpn1 PE=3 SV=1 | 3.438 | 6 | 1 | 2 | 1 | 359 | 40.9 | 5.81 | 8.5 | 1 | 40 | 0 | 74.9 | 42.9 | 0 | 50.2 | 38.9 | 0 | 72.1 | 0.848 | 0.800287586 | 0.966057441 | 0.967557408 | 1.19226638 | 0.830975652 |
| A0A3Q0CFW7 | Calcium-transporting ATPase OS=Mesocricetus auratus OX=10036 GN=Atp2a3 PE=3 SV=1 | 3.421 | 2 | 2 | 3 | 1 | 944 | 103.3 | 6.46 | 2.37 | 2 | 0 | 195 | 152.4 | 0 | 181.6 | 146.8 | 0 | 154 | 145.9 | 0.947 | 0.941620306 | 0.863270006 | 0.848369247 | 0.913215591 | 0.90517563 |
| A0A3Q0CZR3 | dipeptidyl peptidase 8 OS=Mesocricetus auratus OX=10036 GN=Dpp8 PE=4 SV=1 | 3.389 | 1 | 1 | 1 | 1 | 892 | 102.3 | 5.68 | 3.34 | 1 | 0 | 0 | 55.8 | 0 | 0 | 44 | 0 | 0 | 48.5 | 0.788 | 0.876584528 | 0.869175627 | 0.926182747 | 1.102272727 | 0.948541457 |
| A0A3Q0CJ32 | protein unc-13 homolog D OS=Mesocricetus auratus OX=10036 GN=Unc13d PE=4 SV=1 | 3.346 | 2 | 1 | 3 | 1 | 836 | 95.1 | 6.84 | 9.3 | 1 | 57.3 | 45.7 | 40.7 | 33.5 | 38.1 | 28.8 | 46.6 | 41 | 43.8 | 0.704 | 0.079481871 | 0.91440501 | 0.498301194 | 1.30876494 | 0.040199002 |
| A0A3Q0D6M7 | Procollagen-lysine 5-dioxygenase OS=Mesocricetus auratus OX=10036 GN=Plod3 PE=4 SV=1 | 3.335 | 2 | 1 | 2 | 1 | 742 | 85.1 | 6.54 | 3.21 | 1 | 41 | 0 | 0 | 37.4 | 0 | 0 | 17.8 | 0 | 0 | 0.912 | 0.951414946 | 0.434146341 | 0.642863288 | 0.475935829 | 0.669806907 |
| A0A1U8CL93 | poly(ADP-ribose) glycohydrolase ARH3 isoform X2 OS=Mesocricetus auratus OX=10036 GN=Adprhl2 PE=3 SV=2 | 3.324 | 4 | 1 | 1 | 1 | 308 | 32.1 | 5.81 | 3.06 | 1 | 54.9 | 0 | 0 | 50 | 0 | 0 | 61.3 | 0 | 0 | 0.911 | 0.950580313 | 1.116575592 | 0.941786161 | 1.226 | 0.893582473 |
| A0A3Q0DGR4 | endophilin-B2 isoform X10 OS=Mesocricetus auratus OX=10036 GN=Sh3glb2 PE=4 SV=1 | 3.305 | 3 | 1 | 2 | 1 | 398 | 43.9 | 6.28 | 6.33 | 1 | 0 | 43.1 | 0 | 0 | 41.5 | 0 | 0 | 31.9 | 0 | 0.963 | 0.979948549 | 0.740139211 | 0.845577251 | 0.768674699 | 0.863962146 |
| A0A1U7QWN9 | Tripeptidyl aminopeptidase OS=Mesocricetus auratus OX=10036 GN=Tpp1 PE=4 SV=2 | 3.281 | 2 | 1 | 5 | 1 | 585 | 63.8 | 6.44 | 11 | 1 | 62.1 | 164 | 189.5 | 69.5 | 191.8 | 206.4 | 55.3 | 203.9 | 200 | 1.169 | 0.78078331 | 1.104908566 | 0.828066602 | 0.981825957 | 0.967539337 |
| A0A3Q0D5S8 | Guanine nucleotide-binding protein subunit gamma OS=Mesocricetus auratus OX=10036 GN=Gng5 PE=3 SV=1 | 3.27 | 21 | 2 | 8 | 2 | 68 | 7.3 | 9.85 | 7.5 | 2 | 601.6 | 320 | 731.9 | 433.5 | 253.4 | 508.1 | 550.2 | 234.6 | 746.5 | 0.721 | 0.356557106 | 0.92609616 | 0.843057037 | 1.281422594 | 0.551035464 |
| A0A3Q0DAD2 | regulation of nuclear pre-mRNA domain-containing protein 1B isoform X2 OS=Mesocricetus auratus OX=10036 GN=Rprd1b PE=4 SV=1 | 3.262 | 3 | 1 | 2 | 1 | 294 | 33.4 | 5.77 | 6.14 | 1 | 0 | 212 | 160.1 | 0 | 181 | 149.7 | 0 | 152.7 | 162.3 | 0.894 | 0.878739553 | 0.846546627 | 0.82959959 | 0.952524947 | 0.948889833 |
| A0A3Q0CG72 | epsin-2 OS=Mesocricetus auratus OX=10036 GN=Epn2 PE=3 SV=1 | 3.225 | 4 | 2 | 3 | 2 | 649 | 69.1 | 7.65 | 6.1 | 2 | 0 | 104 | 0 | 0 | 115.3 | 0 | 0 | 87.9 | 0 | 1.109 | 0.945514382 | 0.845192308 | 0.911732023 | 0.762359063 | 0.859917041 |
| A0A3Q0CTJ0 | nardilysin OS=Mesocricetus auratus OX=10036 GN=Nrdc PE=4 SV=1 | 3.204 | 1 | 1 | 5 | 1 | 1224 | 139.5 | 4.94 | 15.37 | 1 | 0 | 79.7 | 98.4 | 0 | 87.5 | 101.1 | 0 | 61.2 | 98.5 | 1.061 | 0.94008302 | 0.896687254 | 0.890065616 | 0.846765642 | 0.832872403 |
| A0A3Q0D9M8 | integrin alpha-8 OS=Mesocricetus auratus OX=10036 GN=Itga8 PE=3 SV=1 | 3.2 | 1 | 1 | 7 | 1 | 968 | 106.9 | 6.42 | 26.77 | 1 | 188.9 | 136.8 | 108.4 | 244.6 | 184 | 166.7 | 215.1 | 109.6 | 123.1 | 1.459 | 0.182612268 | 1.031559548 | 0.916563959 | 0.752225769 | 0.300087513 |
| A0A1U7QZN9 | U2 small nuclear ribonucleoprotein A' isoform X1 OS=Mesocricetus auratus OX=10036 GN=Snrpa1 PE=4 SV=2 | 3.139 | 4 | 1 | 4 | 1 | 255 | 28.3 | 8.62 | 6.93 | 1 | 59.5 | 68.7 | 91.5 | 61.4 | 54.4 | 98.4 | 45.2 | 48.9 | 66.6 | 1.032 | 0.918164893 | 0.73145198 | 0.173325053 | 0.750233427 | 0.327369242 |
| A0A3Q0CGW3 | 60S ribosomal protein L27a OS=Mesocricetus auratus OX=10036 GN=LOC110339759 PE=3 SV=1 | 3.101 | 8 | 1 | 5 | 1 | 158 | 17.6 | 10.98 | 18.15 | 1 | 48 | 74.7 | 62.6 | 45.1 | 57.6 | 52.8 | 41.8 | 82.4 | 53.7 | 0.914 | 0.33263397 | 0.96006476 | 0.87289418 | 1.144051447 | 0.604965659 |
| A0A3Q0CWJ7 | syntaxin-binding protein 3 OS=Mesocricetus auratus OX=10036 GN=Stxbp3 PE=3 SV=1 | 3.07 | 3 | 1 | 1 | 1 | 593 | 68 | 7.78 | 4.08 | 1 | 57.8 | 0 | 0 | 55.9 | 0 | 0 | 42.1 | 0 | 0 | 0.967 | 0.982281425 | 0.728373702 | 0.837920443 | 0.75313059 | 0.853981321 |
| A0A1U8BX04 | Peptidylprolyl isomerase OS=Mesocricetus auratus OX=10036 GN=Fkbp1a PE=4 SV=2 | 3.028 | 9 | 1 | 6 | 1 | 108 | 11.9 | 8.16 | 0 | 1 | 165.7 | 174.4 | 211.2 | 121.7 | 122.6 | 188.5 | 203.6 | 204.7 | 216.6 | 0.735 | 0.218158123 | 1.13350263 | 0.214810392 | 1.443853974 | 0.09671973 |
| A0A3Q0CLI1 | endoplasmic reticulum metallopeptidase 1 OS=Mesocricetus auratus OX=10036 GN=Ermp1 PE=4 SV=1 | 2.84 | 1 | 1 | 2 | 1 | 900 | 100 | 7.96 | 6.92 | 1 | 62.5 | 0 | 0 | 40.9 | 0 | 0 | 54.5 | 0 | 0 | 0.655 | 0.788978309 | 0.872 | 0.927866826 | 1.332518337 | 0.852257399 |
| A0A3Q0CVT6 | 2-arachidonoylglycerol hydrolase ABHD12 OS=Mesocricetus auratus OX=10036 GN=Abhd12 PE=3 SV=1 | 2.786 | 7 | 2 | 3 | 2 | 335 | 37.6 | 8.34 | 3.36 | 2 | 0 | 207.9 | 0 | 0 | 153.3 | 0 | 0 | 192.3 | 0 | 0.737 | 0.843781322 | 0.924963925 | 0.958727621 | 1.254403131 | 0.882043413 |
| A0A3Q0DDK3 | Neuropilin OS=Mesocricetus auratus OX=10036 GN=Nrp1 PE=3 SV=1 | 2.765 | 2 | 1 | 1 | 1 | 923 | 103 | 6.07 | 4.76 | 1 | 0 | 0 | 32.4 | 0 | 0 | 36.2 | 0 | 0 | 23.9 | 1.118 | 0.941454384 | 0.737654321 | 0.843963644 | 0.660220994 | 0.792868914 |
| A0A3Q0C9P0 | Eukaryotic translation initiation factor 3 subunit K OS=Mesocricetus auratus OX=10036 GN=Eif3k PE=3 SV=1 | 2.757 | 7 | 1 | 2 | 1 | 218 | 25.1 | 4.93 | 5.67 | 1 | 84 | 0 | 95.8 | 74.7 | 0 | 91.2 | 91.7 | 0 | 84.8 | 0.92 | 0.915888227 | 0.981646274 | 0.980443028 | 1.063893912 | 0.935001757 |
| A0A3Q0CHL2 | Small nuclear ribonucleoprotein E OS=Mesocricetus auratus OX=10036 GN=Snrpe PE=3 SV=1 | 2.736 | 12 | 1 | 8 | 1 | 92 | 10.8 | 9.44 | 18.3 | 1 | 30.1 | 0 | 0 | 40.9 | 0 | 0 | 30.9 | 0 | 0 | 1.36 | 0.842850164 | 1.026578073 | 0.986092476 | 0.755501222 | 0.855508877 |
| A0A1U7R651 | dehydrogenase/reductase SDR family member 7 OS=Mesocricetus auratus OX=10036 GN=LOC101828166 PE=3 SV=2 | 2.733 | 2 | 1 | 1 | 1 | 486 | 54.3 | 9.63 | 2.14 | 1 | 0 | 38.3 | 0 | 0 | 40.1 | 0 | 0 | 25.7 | 0 | 1.046 | 0.975662983 | 0.671018277 | 0.800071454 | 0.640897756 | 0.779936292 |
| A0A1U7Q429 | ATP synthase lipid-binding protein OS=Mesocricetus auratus OX=10036 GN=Atp5g2 PE=3 SV=2 | 2.712 | 5 | 1 | 4 | 1 | 141 | 14.9 | 9.99 | 6.82 | 1 | 105 | 70.1 | 70.9 | 85 | 64.5 | 66.6 | 69.5 | 66.3 | 57.2 | 0.92 | 0.503019891 | 0.784552846 | 0.260686031 | 0.893105044 | 0.375596953 |
| A0A3Q0DBH1 | 3'(2'),5'-bisphosphate nucleotidase 1 isoform X2 OS=Mesocricetus auratus OX=10036 GN=Bpnt1 PE=3 SV=1 | 2.683 | 3 | 1 | 2 | 1 | 308 | 33.2 | 5.87 | 3.69 | 1 | 0 | 0 | 26.5 | 0 | 0 | 33.5 | 0 | 0 | 28.4 | 1.264 | 0.878172211 | 1.071698113 | 0.96334317 | 0.847761194 | 0.913292526 |
| A0A3Q0D6B0 | dedicator of cytokinesis protein 9 OS=Mesocricetus auratus OX=10036 GN=Dock9 PE=3 SV=1 | 2.679 | 1 | 1 | 1 | 1 | 2079 | 236.5 | 8.12 | 3.2 | 1 | 41.7 | 0 | 0 | 44.3 | 0 | 0 | 44.2 | 0 | 0 | 1.062 | 0.967967474 | 1.059952038 | 0.969161246 | 0.997742664 | 0.998801516 |
| A0A3Q0D5S6 | dnaJ homolog subfamily C member 3 OS=Mesocricetus auratus OX=10036 GN=Dnajc3 PE=4 SV=1 | 2.622 | 5 | 1 | 3 | 1 | 514 | 58.9 | 5.83 | 8.11 | 1 | 0 | 0 | 56.8 | 0 | 0 | 61.6 | 0 | 0 | 52.4 | 1.084 | 0.957082026 | 0.922535211 | 0.957343054 | 0.850649351 | 0.915043389 |
| A0A3Q0DAB4 | RNA-binding protein 39 OS=Mesocricetus auratus OX=10036 GN=Rbm39 PE=4 SV=1 | 2.607 | 2 | 1 | 4 | 1 | 496 | 55.6 | 10.46 | 9.23 | 1 | 100.8 | 60.5 | 42.9 | 112 | 61.6 | 45.7 | 68.4 | 35.8 | 38.4 | 1.105 | 0.857892256 | 0.698334966 | 0.375308263 | 0.650250798 | 0.338933641 |
| A0A1U8D2N2 | heat shock 70 kDa protein 4L-like OS=Mesocricetus auratus OX=10036 GN=LOC101831648 PE=3 SV=2 | 2.597 | 1 | 1 | 6 | 1 | 659 | 74.3 | 5.77 | 15.43 | 1 | 167.1 | 260.8 | 328 | 116.2 | 170.6 | 231.1 | 166 | 194.3 | 305.2 | 0.698 | 0.245279143 | 0.880407461 | 0.658083688 | 1.284997104 | 0.415811681 |
| A0A3Q0CDJ1 | ELKS/Rab6-interacting/CAST family member 1 OS=Mesocricetus auratus OX=10036 GN=Erc1 PE=4 SV=1 | 2.572 | 1 | 1 | 1 | 1 | 1123 | 128.6 | 5.91 | 0 | 1 | 0 | 27.1 | 0 | 0 | 28.1 | 0 | 0 | 31.1 | 0 | 1.037 | 0.980792451 | 1.147601476 | 0.927499036 | 1.106761566 | 0.946409812 |
| A0A3Q0D688 | cytochrome c, testis-specific OS=Mesocricetus auratus OX=10036 GN=LOC110342638 PE=3 SV=1 | 2.556 | 7 | 1 | 6 | 1 | 105 | 11.7 | 9.54 | 10.5 | 1 | 106.6 | 89.2 | 85.6 | 135.1 | 120.9 | 111.7 | 137.2 | 97.4 | 93.1 | 1.355 | 0.037767672 | 1.164534471 | 0.396120182 | 0.891215665 | 0.457721299 |
| A0A3Q0D158 | Vesicle-associated membrane protein-associated protein A OS=Mesocricetus auratus OX=10036 GN=Vapa PE=3 SV=1 | 2.548 | 4 | 1 | 2 | 1 | 260 | 29.3 | 8.29 | 5.53 | 1 | 0 | 44.2 | 38.5 | 0 | 31.1 | 33.3 | 0 | 40.4 | 39.3 | 0.78 | 0.746826589 | 0.963724305 | 0.960995558 | 1.23757764 | 0.780887013 |
| A0A1U7R5E3 | Protein-synthesizing GTPase OS=Mesocricetus auratus OX=10036 GN=Eif2s3 PE=4 SV=2 | 2.536 | 2 | 1 | 2 | 1 | 472 | 51.1 | 8.4 | 2.54 | 1 | 0 | 82.7 | 75.3 | 0 | 66.1 | 72.5 | 0 | 81.5 | 70.3 | 0.877 | 0.863101931 | 0.960759494 | 0.957822695 | 1.095238095 | 0.904617199 |
| A0A3Q0CFT2 | Oxysterol-binding protein OS=Mesocricetus auratus OX=10036 GN=Osbp PE=3 SV=1 | 2.519 | 2 | 1 | 1 | 1 | 682 | 77.7 | 6.83 | 2.68 | 1 | 0 | 0 | 69 | 0 | 0 | 68.1 | 0 | 0 | 46.4 | 0.987 | 0.99303759 | 0.672463768 | 0.801034187 | 0.681350954 | 0.806944728 |
| A0A1U8CNI4 | 1,4-beta-N-acetylmuramidase C OS=Mesocricetus auratus OX=10036 GN=LOC101832219 PE=3 SV=2 | 2.506 | 5 | 1 | 3 | 1 | 169 | 18.9 | 9.13 | 8.51 | 1 | 33.9 | 21.8 | 33.9 | 70.1 | 53.9 | 52.8 | 38.7 | 43.1 | 46.5 | 2.105 | 0.016502231 | 1.431919643 | 0.064868989 | 0.725678733 | 0.086208629 |
| A0A3Q0D1W0 | ATPase family AAA domain-containing protein 5 OS=Mesocricetus auratus OX=10036 GN=Atad5 PE=4 SV=1 | 2.463 | 1 | 1 | 13 | 1 | 1836 | 205.2 | 9.01 | 35.66 | 1 | 1763.7 | 836.9 | 1302.2 | 711.6 | 255.8 | 638.8 | 908.2 | 448.2 | 650.5 | 0.403 | 0.084440287 | 0.51422056 | 0.126862016 | 1.249470801 | 0.529479045 |
| A0A3Q0D5X5 | UBX domain-containing protein 4 OS=Mesocricetus auratus OX=10036 GN=Ubxn4 PE=4 SV=1 | 2.437 | 1 | 1 | 3 | 1 | 572 | 63.2 | 8.57 | 2.66 | 1 | 171.4 | 77.4 | 0 | 166.3 | 67.2 | 0 | 160.9 | 66.4 | 0 | 0.918 | 0.944790602 | 0.913585209 | 0.921249904 | 0.973447537 | 0.976931504 |
| A0A3Q0CAG6 | DEP domain-containing protein 1A isoform X2 OS=Mesocricetus auratus OX=10036 GN=Depdc1 PE=4 SV=1 | 2.429 | 1 | 1 | 1 | 1 | 741 | 84.9 | 8.46 | 2.13 | 1 | 203.3 | 0 | 0 | 268 | 0 | 0 | 193.7 | 0 | 0 | 1.318 | 0.857491235 | 0.952779144 | 0.974369172 | 0.722761194 | 0.834253451 |
| A0A3Q0DCZ3 | high mobility group protein B1 OS=Mesocricetus auratus OX=10036 GN=Hmgb1 PE=4 SV=1 | 2.406 | 5 | 1 | 6 | 1 | 155 | 17.5 | 6.25 | 0 | 1 | 437.3 | 326.1 | 247.4 | 575.4 | 265.7 | 595.5 | 732.9 | 1052 | 735.6 | 1.698 | 0.322625253 | 2.49356945 | 0.024267858 | 1.754489768 | 0.074145906 |
| A0A3Q0CM60 | poly [ADP-ribose] polymerase 14 OS=Mesocricetus auratus OX=10036 GN=Parp14 PE=4 SV=1 | 2.391 | 2 | 1 | 2 | 1 | 478 | 53 | 5.99 | 5.79 | 1 | 0 | 136.1 | 0 | 0 | 180 | 0 | 0 | 144.3 | 0 | 1.322 | 0.855901562 | 1.060249816 | 0.969012845 | 0.801666667 | 0.884851984 |
| A0A3Q0CWM0 | mitochondrial import receptor subunit TOM70 OS=Mesocricetus auratus OX=10036 GN=Tomm70 PE=4 SV=1 | 2.387 | 1 | 1 | 1 | 1 | 611 | 67.4 | 7.52 | 0 | 1 | 0 | 62.3 | 0 | 0 | 56.7 | 0 | 0 | 43.4 | 0 | 0.909 | 0.950214359 | 0.696629213 | 0.817067972 | 0.765432099 | 0.861887057 |
| A0A3Q0D402 | very-long-chain 3-oxoacyl-CoA reductase OS=Mesocricetus auratus OX=10036 GN=Hsd17b12 PE=3 SV=1 | 2.318 | 3 | 1 | 3 | 1 | 312 | 34.9 | 9.52 | 8.04 | 1 | 44.2 | 32.1 | 48 | 28.6 | 33.4 | 32.3 | 42.5 | 33.5 | 26.1 | 0.824 | 0.164001417 | 0.821399839 | 0.334073954 | 1.08271474 | 0.64502412 |
| A0A3Q0D4B0 | Signal sequence receptor subunit gamma OS=Mesocricetus auratus OX=10036 GN=Ssr3 PE=3 SV=1 | 2.292 | 7 | 1 | 1 | 1 | 256 | 29.7 | 9.77 | 0 | 1 | 0 | 0 | 49.2 | 0 | 0 | 47.9 | 0 | 0 | 38.3 | 0.973 | 0.985802557 | 0.778455285 | 0.870198107 | 0.799582463 | 0.883544846 |
| A0A1U7QEV0 | metaxin-2 isoform X1 OS=Mesocricetus auratus OX=10036 GN=Mtx2 PE=3 SV=2 | 2.288 | 4 | 1 | 3 | 1 | 263 | 29.7 | 5.82 | 8.3 | 1 | 146.5 | 130.9 | 122.6 | 119.6 | 109.2 | 100.8 | 113.5 | 114.1 | 109.3 | 0.834 | 0.060858017 | 0.84225 | 0.08926595 | 1.022148058 | 0.703329414 |
| A0A1U7QCC8 | dynein heavy chain 8, axonemal OS=Mesocricetus auratus OX=10036 GN=Dnah8 PE=3 SV=2 | 2.265 | 0 | 1 | 5 | 1 | 4735 | 541.9 | 5.99 | 1.74 | 1 | 570 | 101.1 | 0 | 384 | 75.5 | 0 | 556.5 | 2827.1 | 0 | 0.71 | 0.757543774 | 5.041871554 | 0.406152952 | 7.363656148 | 0.376645695 |
| A0A3Q0D6X5 | 40S ribosomal protein S20 OS=Mesocricetus auratus OX=10036 GN=LOC110342821 PE=3 SV=1 | 2.262 | 10 | 1 | 3 | 1 | 90 | 10.2 | 8.75 | 0 | 1 | 0 | 84.3 | 55 | 0 | 73.3 | 51.4 | 0 | 66.8 | 65.2 | 0.902 | 0.889675149 | 0.947595118 | 0.944949609 | 1.058540497 | 0.941055844 |
| A0A3Q0D1Q5 | Mitochondrial Rho GTPase OS=Mesocricetus auratus OX=10036 GN=Rhot1 PE=3 SV=1 | 2.243 | 1 | 1 | 1 | 1 | 672 | 76.9 | 7.06 | 2.21 | 1 | 41.3 | 0 | 0 | 40.3 | 0 | 0 | 30.6 | 0 | 0 | 0.977 | 0.987003977 | 0.740920097 | 0.846083935 | 0.759305211 | 0.857956051 |
| A0A3Q0D9E2 | 3-phosphoinositide-dependent protein kinase 1 OS=Mesocricetus auratus OX=10036 GN=Pdpk1 PE=3 SV=1 | 2.224 | 2 | 1 | 4 | 1 | 532 | 61 | 7.2 | 11.52 | 1 | 34.7 | 0 | 0 | 26.1 | 0 | 0 | 23.6 | 0 | 0 | 0.75 | 0.853356253 | 0.680115274 | 0.806123834 | 0.904214559 | 0.94680286 |
| A0A3Q0CCH5 | maltase-glucoamylase, intestinal OS=Mesocricetus auratus OX=10036 GN=LOC101844514 PE=3 SV=1 | 2.198 | 1 | 1 | 8 | 1 | 2515 | 285.9 | 5.2 | 12.72 | 1 | 16.5 | 29.2 | 26.1 | 27.8 | 47 | 39.9 | 37 | 31.9 | 30 | 1.691 | 0.111945678 | 1.377437326 | 0.126958479 | 0.862249346 | 0.4537836 |
| A0A3Q0CMT9 | coiled-coil domain-containing protein 92 OS=Mesocricetus auratus OX=10036 GN=LOC101829958 PE=4 SV=1 | 2.177 | 3 | 1 | 3 | 1 | 263 | 30.1 | 8.07 | 7.19 | 1 | 0 | 330.2 | 403.2 | 0 | 248.2 | 318.5 | 0 | 322.5 | 347.5 | 0.771 | 0.742615197 | 0.913553313 | 0.905495879 | 1.182283395 | 0.8274859 |

Supplementary Table 1: List of differentially expressed proteins identified across the 4dpi infected and 14dpi infected

| Accession | Description | Sum PEP Score | Coverage [%] | # Peptides | # PSMs | # Unique Peptides | # AAs | MW [kDa] | calc. pI | Score Sequest HT: Sequest HT | # Peptides (by Search Engine): Sequest HT | Mock-infected (4dpi) Replicate_R1 | Mock-infected (4dpi) Replicate_R2 | Mock-infected (4dpi) Replicate_R3 | SARS-CoV-2-infected (4dpi) Replicate_R1 | SARS-CoV-2-infected (4dpi) Replicate_R2 | SARS-CoV-2-infected (4dpi) Replicate_R3 | SARS-CoV-2-(14dpi) Replicate_R1 | SARS-CoV-2-infected (14dpi) Replicate_R2 | SARS-CoV-2-infected (14dpi) Replicate_R3 | Abundance Ratio: SARS-CoV-2-infected (4dpi) / Mock-infected (4dpi) | P-Value: SARS-CoV-2-infected (4dpi) / Mock-infected (4dpi) | Abundance Ratio: SARS-CoV-2-infected (14dpi) / Mock-infected (4dpi) | P-Value: SARS-CoV-2-infected (14dpi) / Mock-infected (4dpi) | Abundance Ratio: SARS-CoV-2-infected (14dpi) / SARS-CoV-2-infected (4dpi) | P-Value: SARS-CoV-2-infected (14dpi) / SARS-CoV-2-infected (4dpi) |
|---|---|---|---|---|---|---|---|---|---|---|---|---|---|---|---|---|---|---|---|---|---|---|---|---|---|
| A0A3Q0CE90 | Isocitrate dehydrogenase [NAD] subunit, mitochondrial OS=Mesocricetus auratus OX=10036 GN=Idh3g PE=3 SV=1 | 2.173 | 2 | 1 | 1 | 1 | 389 | 42.4 | 8.78 | 2.07 | 1 | 0 | 30.8 | 0 | 0 | 25.2 | 0 | 0 | 31.4 | 0 | 0.819 | 0.89514666 | 1.019480519 | 0.989769655 | 1.246031746 | 0.885404054 |
| A0A3Q0CQB6 | merlin isoform X3 OS=Mesocricetus auratus OX=10036 GN=Nf2 PE=4 SV=1 | 2.129 | 2 | 1 | 1 | 1 | 622 | 72.4 | 6.83 | 2.28 | 1 | 0 | 55.7 | 0 | 0 | 55.7 | 0 | 0 | 45.3 | 0 | 1 | 1 | 0.813285458 | 0.892106359 | 0.813285458 | 0.892106359 |
| A0A3Q0CUN6 | Serine/threonine-protein phosphatase 2A 56 kDa regulatory subunit OS=Mesocricetus auratus OX=10036 GN=Ppp2r5e PE=3 SV=1 | 2.128 | 1 | 1 | 1 | 1 | 467 | 54.7 | 6.95 | 0 | 1 | 0 | 0 | 140.4 | 0 | 0 | 115 | 0 | 0 | 140.5 | 0.819 | 0.895708422 | 1.000712251 | 0.999622407 | 1.22173913 | 0.895347186 |
| A0A3Q0DD86 | Cactin OS=Mesocricetus auratus OX=10036 GN=Cactin PE=3 SV=1 | 2.123 | 1 | 1 | 7 | 1 | 800 | 93 | 9.74 | 14.97 | 1 | 295.5 | 358.4 | 1050.7 | 169.6 | 153 | 493.6 | 1919.1 | 371.5 | 937.5 | 0.47 | 0.352014018 | 1.893758066 | 0.393720384 | 3.955035531 | 0.212707334 |
| A0A3Q0CYM2 | Delta-aminolevulinic acid dehydratase OS=Mesocricetus auratus OX=10036 GN=Alad PE=3 SV=1 | 2.105 | 4 | 1 | 1 | 1 | 330 | 36.1 | 6.98 | 2.57 | 1 | 0 | 53 | 0 | 0 | 40.7 | 0 | 0 | 33.9 | 0 | 0.769 | 0.863482411 | 0.639622642 | 0.779081281 | 0.832923833 | 0.904238488 |
| A0A3Q0D488 | protein enabled homolog isoform X3 OS=Mesocricetus auratus OX=10036 GN=Enah PE=3 SV=1 | 2.104 | 1 | 1 | 4 | 1 | 795 | 85.9 | 7.83 | 11.93 | 1 | 93.7 | 90 | 214.2 | 81 | 114.8 | 233.4 | 77 | 68.8 | 154.6 | 0.876 | 0.873927792 | 0.754963559 | 0.549164937 | 0.699906803 | 0.478271949 |
| A0A3Q0DF55 | napsin-A OS=Mesocricetus auratus OX=10036 GN=Napsa PE=3 SV=1 | 2.102 | 2 | 1 | 6 | 1 | 400 | 43 | 8.24 | 14.71 | 1 | 173.1 | 189.6 | 175.5 | 153.2 | 192.7 | 183.6 | 143 | 168.2 | 143.5 | 0.888 | 0.839223718 | 0.844853214 | 0.057682229 | 0.858734655 | 0.170374148 |
| A0A3Q0C9D0 | Nicotinate phosphoribosyltransferase OS=Mesocricetus auratus OX=10036 GN=Naprt PE=3 SV=1 | 2.038 | 1 | 1 | 2 | 1 | 514 | 55 | 7.78 | 4.01 | 1 | 0 | 33.6 | 71.8 | 0 | 31.5 | 57.7 | 0 | 46.1 | 84 | 0.867 | 0.849559142 | 1.234345351 | 0.809585587 | 1.458520179 | 0.670531228 |
| A0A3Q0CMK9 | Kinesin light chain OS=Mesocricetus auratus OX=10036 GN=Klc1 PE=3 SV=1 | 1.989 | 2 | 1 | 3 | 1 | 528 | 59.4 | 6.44 | 7.66 | 1 | 130.9 | 120.8 | 94.2 | 129.7 | 123 | 89.8 | 128.7 | 111.7 | 79.9 | 0.991 | 0.948554137 | 0.925990171 | 0.661920022 | 0.935182482 | 0.715586744 |
| A0A3Q0DD58 | isochorismatase domain-containing protein 2A OS=Mesocricetus auratus OX=10036 GN=LOC101839073 PE=3 SV=1 | 1.988 | 5 | 1 | 1 | 1 | 187 | 20.7 | 9.04 | 2.12 | 1 | 0 | 0 | 129.3 | 0 | 0 | 88.5 | 0 | 0 | 139.4 | 0.685 | 0.809005306 | 1.078112916 | 0.960196685 | 1.575141243 | 0.775888771 |
| A0A3Q0D155 | Ribosomal protein L19 OS=Mesocricetus auratus OX=10036 GN=Rpl19 PE=3 SV=1 | 1.979 | 8 | 1 | 3 | 1 | 119 | 14.2 | 12.04 | 0 | 1 | 76.9 | 96.9 | 0 | 52.6 | 95.8 | 0 | 70.7 | 109.2 | 0 | 0.823 | 0.844633682 | 1.035097814 | 0.964996537 | 1.212264151 | 0.81644213 |
| A0A3Q0CUT3 | Non-specific serine/threonine protein kinase OS=Mesocricetus auratus OX=10036 GN=Stk10 PE=3 SV=1 | 1.969 | 1 | 1 | 2 | 1 | 1000 | 115.5 | 7.47 | 5.47 | 1 | 405 | 0 | 221 | 413.2 | 0 | 256.8 | 316.4 | 0 | 189.6 | 1.089 | 0.934619972 | 0.808306709 | 0.802149275 | 0.755223881 | 0.737713029 |
| A0A1U7QT64 | ribonucleoprotein PTB-binding 1 OS=Mesocricetus auratus OX=10036 GN=Raver1 PE=4 SV=2 | 1.961 | 2 | 1 | 3 | 1 | 744 | 78.6 | 8.66 | 5.6 | 1 | 0 | 45.3 | 42.3 | 0 | 44.9 | 54.4 | 0 | 45.6 | 40.4 | 1.129 | 0.869559555 | 0.98173516 | 0.980522244 | 0.866062437 | 0.851113411 |
| A0A3Q0CZW8 | anillin isoform X1 OS=Mesocricetus auratus OX=10036 GN=Anln PE=4 SV=1 | 1.948 | 1 | 1 | 12 | 1 | 1142 | 125.5 | 7.46 | 0 | 1 | 186.9 | 36.1 | 660.1 | 972.1 | 522.9 | 603.1 | 742.8 | 428.5 | 450.1 | 3.918 | 0.16399856 | 1.836032159 | 0.330892961 | 0.772794433 | 0.410970236 |
| A0A3Q0CI27 | Integrin beta OS=Mesocricetus auratus OX=10036 GN=Itgb6 PE=3 SV=1 | 1.938 | 1 | 1 | 7 | 1 | 787 | 85.6 | 5.26 | 20.51 | 1 | 347.3 | 625.1 | 400.9 | 112.3 | 198 | 118.6 | 331.5 | 609.4 | 416.3 | 0.303 | 0.054495116 | 0.988276414 | 0.966000771 | 3.16437398 | 0.051995725 |
| A0A1U7QYI0 | short transient receptor potential channel 6 OS=Mesocricetus auratus OX=10036 GN=Trpc6 PE=4 SV=2 | 1.934 | 1 | 1 | 1 | 1 | 929 | 106.2 | 6.83 | 2.61 | 1 | 0 | 89.6 | 0 | 0 | 60.5 | 0 | 0 | 84.3 | 0 | 0.675 | 0.802870994 | 0.940848214 | 0.967708731 | 1.39338843 | 0.830922614 |
| A0A1U8BSQ8 | Small ubiquitin-related modifier OS=Mesocricetus auratus OX=10036 GN=Sumo1 PE=3 SV=2 | 1.924 | 7 | 1 | 2 | 1 | 101 | 11.6 | 5.82 | 1.93 | 1 | 56.2 | 0 | 47.6 | 59.7 | 0 | 41 | 39.7 | 0 | 33.9 | 0.956 | 0.968792721 | 0.709055877 | 0.665362761 | 0.730883813 | 0.698861697 |
| A0A3Q0CMN9 | centrosomal protein of 152 kDa OS=Mesocricetus auratus OX=10036 GN=Cep152 PE=4 SV=1 | 1.869 | 1 | 1 | 1 | 1 | 1743 | 197.1 | 5.71 | 2.67 | 1 | 101 | 0 | 0 | 45.1 | 0 | 0 | 99.6 | 0 | 0 | 0.446 | 0.650778731 | 0.986138614 | 0.992598004 | 2.208425721 | 0.654813379 |
| A0A3Q0CW18 | uncharacterized protein LOC110341366 OS=Mesocricetus auratus OX=10036 GN=LOC110341366 PE=3 SV=1 | 1.836 | 4 | 1 | 4 | 1 | 191 | 20.8 | 9.23 | 9.19 | 1 | 234.9 | 98.2 | 312.6 | 252 | 105.9 | 358.2 | 229 | 98.7 | 294.5 | 1.146 | 0.819754582 | 0.963605389 | 0.931100179 | 0.868873062 | 0.754404508 |
| A0A3Q0D4M9 | cytoskeleton-associated protein 5 OS=Mesocricetus auratus OX=10036 GN=Ckap5 PE=4 SV=1 | 1.829 | 1 | 1 | 1 | 1 | 2032 | 225.5 | 7.8 | 4.06 | 1 | 0 | 0 | 95.4 | 0 | 0 | 125.9 | 0 | 0 | 83.8 | 1.32 | 0.85695245 | 0.878406709 | 0.931672842 | 0.665607625 | 0.796464584 |
| A0A3Q0CGJ5 | ankyrin repeat and sterile alpha motif domain-containing protein 1B OS=Mesocricetus auratus OX=10036 GN=Anks1b PE=4 SV=1 | 1.811 | 1 | 1 | 2 | 1 | 1305 | 144.1 | 6.28 | 5.33 | 1 | 76.6 | 0 | 40.8 | 55 | 0 | 32.3 | 121.3 | 0 | 46.8 | 0.754 | 0.733435173 | 1.431856899 | 0.709578845 | 1.925544101 | 0.540622727 |
| A0A3Q0CPU2 | coiled-coil domain-containing protein 91 OS=Mesocricetus auratus OX=10036 GN=Ccdc91 PE=4 SV=1 | 1.806 | 2 | 1 | 2 | 1 | 442 | 49.8 | 5.06 | 5.76 | 1 | 146 | 0 | 172.8 | 145.5 | 0 | 138.2 | 126.2 | 0 | 151.6 | 0.893 | 0.87819976 | 0.871392723 | 0.857447982 | 0.979203384 | 0.977862108 |
| A0A3Q0D004 | DNA (cytosine-5)-methyltransferase OS=Mesocricetus auratus OX=10036 GN=Dnmt1 PE=3 SV=1 | 1.803 | 1 | 1 | 1 | 1 | 1617 | 182.7 | 8.13 | 3.49 | 1 | 22.1 | 0 | 0 | 42.6 | 0 | 0 | 17.7 | 0 | 0 | 1.926 | 0.698031841 | 0.800904977 | 0.884374483 | 0.415492958 | 0.631073362 |
| A0A3Q0D4J2 | protein RCC2 OS=Mesocricetus auratus OX=10036 GN=Rcc2 PE=4 SV=1 | 1.782 | 2 | 1 | 1 | 1 | 520 | 55.9 | 8.79 | 2.06 | 1 | 0 | 0 | 40 | 0 | 0 | 35.2 | 0 | 0 | 30.4 | 0.878 | 0.93261631 | 0.76 | 0.858402484 | 0.863636364 | 0.922868922 |
| A0A3Q0D1L1 | FERM, RhoGEF and pleckstrin domain-containing protein 1 OS=Mesocricetus auratus OX=10036 GN=Farp1 PE=4 SV=1 | 1.746 | 1 | 1 | 2 | 1 | 1069 | 120.8 | 8.24 | 4.23 | 1 | 72.7 | 114 | 0 | 42.8 | 90.9 | 0 | 79.5 | 119.6 | 0 | 0.685 | 0.699571071 | 1.066416711 | 0.936098852 | 1.489154824 | 0.647283216 |
| A0A3Q0D230 | schlafen family member 12-like OS=Mesocricetus auratus OX=10036 GN=LOC101844930 PE=4 SV=1 | 1.727 | 2 | 1 | 3 | 1 | 576 | 65.5 | 7.75 | 0 | 1 | 36.8 | 0 | 0 | 19.5 | 0 | 0 | 80.7 | 0 | 0 | 0.53 | 0.705426038 | 2.192934783 | 0.656875511 | 4.138461538 | 0.530723633 |
| A0A3Q0CSJ7 | 60S ribosomal protein L35a OS=Mesocricetus auratus OX=10036 GN=Rpl35a PE=3 SV=1 | 1.724 | 7 | 1 | 3 | 1 | 110 | 12.5 | 10.89 | 6.63 | 1 | 57.6 | 81.3 | 75.5 | 68.5 | 88.9 | 67.9 | 81.8 | 94.5 | 89.6 | 1.094 | 0.732876344 | 1.240205224 | 0.12215345 | 1.180204172 | 0.180557937 |
| A0A3Q0D1M4 | ubiquitin recognition factor in ER-associated degradation protein 1 isoform X2 OS=Mesocricetus auratus OX=10036 GN=Ufd1 PE=3 SV=1 | 1.711 | 3 | 1 | 1 | 1 | 302 | 33.8 | 6.7 | 2.17 | 1 | 0 | 118.6 | 0 | 0 | 114.4 | 0 | 0 | 96.3 | 0 | 0.965 | 0.980888005 | 0.811973019 | 0.891289713 | 0.841783217 | 0.909656519 |
| A0A1U8C2S8 | kinesin-like protein KIF13B isoform X3 OS=Mesocricetus auratus OX=10036 GN=Kif13b PE=3 SV=2 | 1.708 | 1 | 1 | 1 | 1 | 1501 | 169 | 5.86 | 2.35 | 1 | 0 | 0 | 38.8 | 0 | 0 | 38.1 | 0 | 0 | 41.8 | 0.982 | 0.990346024 | 1.077319588 | 0.960584634 | 1.097112861 | 0.951004842 |
| A0A3Q0D9E3 | filamin-binding LIM protein 1 OS=Mesocricetus auratus OX=10036 GN=Fblim1 PE=4 SV=1 | 1.698 | 3 | 1 | 1 | 1 | 377 | 41.2 | 6.43 | 2.07 | 1 | 0 | 41.3 | 0 | 0 | 35 | 0 | 0 | 28.2 | 0 | 0.848 | 0.913108276 | 0.682808717 | 0.807912764 | 0.805714286 | 0.887385476 |
| A0A3Q0D359 | interferon-stimulated gene 20 kDa protein isoform X3 OS=Mesocricetus auratus OX=10036 GN=Isg20 PE=4 SV=1 | 1.695 | 4 | 1 | 2 | 1 | 183 | 20.6 | 9.47 | 7.27 | 1 | 0 | 149.1 | 65.8 | 0 | 130.7 | 68.4 | 0 | 115.3 | 49.2 | 0.955 | 0.931284353 | 0.765472313 | 0.774426555 | 0.826217981 | 0.83038374 |
| A0A1U8C475 | vesicle-associated membrane protein 5 OS=Mesocricetus auratus OX=10036 GN=Vamp5 PE=4 SV=2 | 1.695 | 3 | 1 | 2 | 1 | 244 | 27.3 | 9.19 | 0 | 1 | 52.7 | 0 | 63.3 | 38.1 | 0 | 39 | 52.3 | 0 | 68.6 | 0.668 | 0.613613281 | 1.042241379 | 0.957029997 | 1.568093385 | 0.587135499 |
| A0A3Q0D8A0 | kinetochore-associated protein 1 OS=Mesocricetus auratus OX=10036 GN=Kntc1 PE=4 SV=1 | 1.594 | 0 | 1 | 3 | 1 | 2208 | 250.5 | 5.85 | 8.41 | 1 | 0 | 0 | 41.5 | 0 | 0 | 26.7 | 0 | 0 | 23.9 | 0.645 | 0.78159586 | 0.575903614 | 0.736246064 | 0.895131086 | 0.94151485 |
| A0A1U8CW59 | Branched-chain-amino-acid aminotransferase OS=Mesocricetus auratus OX=10036 GN=Bcat2 PE=3 SV=2 | 1.589 | 2 | 1 | 4 | 1 | 393 | 44.4 | 8.22 | 7.29 | 1 | 36.5 | 0 | 0 | 33.2 | 0 | 0 | 29.7 | 0 | 0 | 0.912 | 0.949912355 | 0.81369863 | 0.8923633 | 0.894578313 | 0.941191734 |
| A0A3Q0DC52 | beta-1,3-glucosyltransferase OS=Mesocricetus auratus OX=10036 GN=B3glct PE=4 SV=1 | 1.565 | 2 | 1 | 1 | 1 | 449 | 51 | 6.83 | 2.81 | 1 | 0 | 0 | 138.6 | 0 | 0 | 115.5 | 0 | 0 | 113.7 | 0.833 | 0.904489688 | 0.82034632 | 0.896487432 | 0.984415584 | 0.991670804 |
| A0A1U8BYI1 | gamma-soluble NSF attachment protein OS=Mesocricetus auratus OX=10036 GN=Napg PE=3 SV=2 | 1.562 | 4 | 1 | 1 | 1 | 312 | 34.7 | 5.41 | 0 | 1 | 0 | 0 | 41.6 | 0 | 0 | 37.1 | 0 | 0 | 40.3 | 0.891 | 0.939581215 | 0.96875 | 0.983169042 | 1.086253369 | 0.956235127 |
| A0A3Q0DGZ7 | hydroxysteroid dehydrogenase-like protein 2 OS=Mesocricetus auratus OX=10036 GN=Hsdl2 PE=4 SV=1 | 1.559 | 3 | 1 | 2 | 1 | 444 | 48.2 | 7.96 | 0 | 1 | 0 | 0 | 31.6 | 0 | 0 | 39.1 | 0 | 0 | 26.8 | 1.236 | 0.888928122 | 0.848101266 | 0.913498882 | 0.685421995 | 0.809647041 |
| A0A3Q0CIM0 | Acetyltransferase component of pyruvate dehydrogenase complex OS=Mesocricetus auratus OX=10036 GN=Dlat PE=3 SV=1 | 1.558 | 1 | 1 | 2 | 1 | 644 | 68.2 | 8.25 | 2.23 | 1 | 0 | 59.7 | 59.1 | 0 | 67.4 | 58.6 | 0 | 47.9 | 67.4 | 1.058 | 0.937981582 | 0.970538721 | 0.968939922 | 0.915079365 | 0.908465534 |
| A0A1U7QXH9 | protein TFG OS=Mesocricetus auratus OX=10036 GN=Tfg PE=4 SV=2 | 1.55 | 2 | 1 | 1 | 1 | 399 | 43.2 | 5.1 | 0 | 1 | 0 | 0 | 38.5 | 0 | 0 | 35.8 | 0 | 0 | 31.7 | 0.932 | 0.961515407 | 0.823376623 | 0.898361151 | 0.88547486 | 0.935848914 |
| QQV30662.1:1-7096 | ORF1ab polyprotein [organism=Severe acute respiratory syndrome coronavirus 2] [isolate=SARS-CoV-2/human/USA/CA-QDX-3078/2021] | 1.549 | 0 | 1 | 8 | 1 | 7096 | 793.6 | 6.73 | 20.43 | 1 | 122.9 | 57.8 | 135.2 | 337.4 | 156.4 | 374.4 | 125.4 | 54 | 130.9 | 2.77 | 0.098742272 | 0.982272871 | 0.959444899 | 0.357406128 | 0.096284245 |
| A0A1U8BT74 | Integrin beta OS=Mesocricetus auratus OX=10036 GN=Itgb2 PE=3 SV=2 | 1.529 | 1 | 1 | 2 | 1 | 769 | 84.7 | 7.39 | 4.18 | 1 | 0 | 0 | 45.8 | 0 | 0 | 45.3 | 0 | 0 | 31.3 | 0.99 | 0.994178797 | 0.683406114 | 0.808309344 | 0.690949227 | 0.813310393 |
| A0A1U8BRR2 | sorbin and SH3 domain-containing protein 2 isoform X14 OS=Mesocricetus auratus OX=10036 GN=Sorbs2 PE=4 SV=2 | 1.492 | 2 | 1 | 1 | 1 | 771 | 85.3 | 9.14 | 3.05 | 1 | 0 | 0 | 49.2 | 0 | 0 | 43.7 | 0 | 0 | 56.9 | 0.888 | 0.937459233 | 1.156504065 | 0.9234913 | 1.302059497 | 0.863539677 |
| A0A3Q0CLU3 | RING-type E3 ubiquitin transferase OS=Mesocricetus auratus OX=10036 GN=Rc3h1 PE=4 SV=1 | 1.471 | 2 | 1 | 1 | 1 | 1129 | 125.4 | 7.18 | 0 | 1 | 23.9 | 0 | 0 | 53.1 | 0 | 0 | 30.9 | 0 | 0 | 2.219 | 0.653064875 | 1.292887029 | 0.86701934 | 0.581920904 | 0.740291578 |
| A0A3Q0DFP7 | proteasome-associated protein ECM29 homolog OS=Mesocricetus auratus OX=10036 GN=Kiaa0368 PE=4 SV=1 | 1.465 | 0 | 1 | 1 | 1 | 1845 | 204.1 | 7.25 | 1.77 | 1 | 0 | 102.5 | 0 | 0 | 97.4 | 0 | 0 | 77.4 | 0 | 0.951 | 0.972960042 | 0.755121951 | 0.855264615 | 0.794661191 | 0.880451491 |
| A0A3Q0D5Q8 | polymeric immunoglobulin receptor isoform X2 OS=Mesocricetus auratus OX=10036 GN=Pigr PE=4 SV=1 | 1.446 | 1 | 1 | 4 | 1 | 777 | 85.3 | 5.77 | 10.76 | 1 | 202.1 | 396.8 | 350 | 170.6 | 317.3 | 329.1 | 184.1 | 327.3 | 309.3 | 0.8 | 0.602420695 | 0.864896196 | 0.596350223 | 1.004528764 | 0.986415104 |
| A0A3Q0CJY3 | RNA 3'-terminal phosphate cyclase-like protein OS=Mesocricetus auratus OX=10036 GN=Rcl1 PE=3 SV=1 | 1.442 | 2 | 1 | 1 | 1 | 373 | 40.9 | 9.17 | 2.42 | 1 | 0 | 0 | 0 | 565.7 | 0 | 0 | 523.6 | 0 | 0 | 100 | 0.422649731 | #DIV/0! | 0.422649731 | 0.925578929 | 0.959077748 |
| A0A3Q0D8E2 | dapper homolog 3-like OS=Mesocricetus auratus OX=10036 GN=LOC110343426 PE=4 SV=1 | 1.439 | 3 | 1 | 1 | 1 | 271 | 28.5 | 12.04 | 1.85 | 1 | 0 | 169.4 | 0 | 0 | 227.4 | 0 | 0 | 169.7 | 0 | 1.342 | 0.848691056 | 1.001770956 | 0.99906164 | 0.746262093 | 0.849544937 |
| A0A3Q0CZD2 | centromere protein J OS=Mesocricetus auratus OX=10036 GN=Cenpj PE=3 SV=1 | 1.434 | 1 | 1 | 3 | 1 | 1337 | 151.6 | 6.61 | 5.55 | 1 | 100.3 | 99.8 | 95.3 | 135.2 | 113.3 | 129.7 | 140.7 | 115.6 | 126.8 | 1.321 | 0.045676316 | 1.296885579 | 0.050715355 | 1.012956108 | 0.875748386 |
| A0A3Q0CB37 | zinc finger protein 185 isoform X4 OS=Mesocricetus auratus OX=10036 GN=Znf185 PE=4 SV=1 | 1.419 | 2 | 1 | 1 | 1 | 579 | 62.3 | 5.38 | 0 | 1 | 0 | 0 | 45.7 | 0 | 0 | 36.5 | 0 | 0 | 25.6 | 0.799 | 0.882982763 | 0.560175055 | 0.725683181 | 0.701369863 | 0.820198421 |
| A0A3Q0CW22 | Anion exchange protein OS=Mesocricetus auratus OX=10036 GN=Slc4a7 PE=3 SV=1 | 1.402 | 1 | 1 | 3 | 1 | 1095 | 123.1 | 6.58 | 7.01 | 1 | 230.9 | 145.9 | 156.2 | 262 | 214.8 | 226 | 239.8 | 201.2 | 209.1 | 1.447 | 0.157459476 | 1.219699812 | 0.282011341 | 0.925014229 | 0.397306461 |
| A0A3Q0D220 | kelch-like protein 23 OS=Mesocricetus auratus OX=10036 GN=Klhl23 PE=4 SV=1 | 1.387 | 2 | 1 | 2 | 1 | 558 | 63.8 | 5.54 | 5.13 | 1 | 0 | 132.2 | 0 | 0 | 155 | 0 | 0 | 100.5 | 0 | 1.172 | 0.916407086 | 0.7602118 | 0.858538543 | 0.648387097 | 0.784954467 |
| A0A3Q0C9R8 | DNA repair protein XRCC1 OS=Mesocricetus auratus OX=10036 GN=Xrcc1 PE=4 SV=1 | 1.367 | 1 | 1 | 1 | 1 | 620 | 67.5 | 6.43 | 0 | 1 | 98.6 | 0 | 0 | 84.1 | 0 | 0 | 88.2 | 0 | 0 | 0.853 | 0.91643 | 0.8945233 | 0.9411596 | 1.0487515 | 0.9747774 |
| A0A3Q0CR11 | Phosphodiesterase OS=Mesocricetus auratus OX=10036 GN=Pde9a PE=3 SV=1 | 1.358 | 1 | 1 | 3 | 1 | 542 | 62.3 | 6.48 | 0 | 1 | 0 | 0 | 344.6 | 0 | 0 | 246.5 | 0 | 0 | 272.9 | 0.715 | 0.8293794 | 0.7919327 | 0.8787323 | 1.1070994 | 0.9462498 |
| A0A3Q0CEI2 | unconventional myosin-IXb OS=Mesocricetus auratus OX=10036 GN=Myo9b PE=3 SV=1 | 1.351 | 1 | 1 | 2 | 1 | 1681 | 188.2 | 8.88 | 0 | 1 | 46.9 | 51.3 | 0 | 37.1 | 49.9 | 0 | 34.9 | 34.8 | 0 | 0.878 | 0.8747461 | 0.709776 | 0.663831 | 0.8011494 | 0.7768743 |
| A0A1U7RFM2 | AP-1 complex-associated regulatory protein OS=Mesocricetus auratus OX=10036 GN=Ap1ar PE=4 SV=2 | 1.343 | 4 | 1 | 1 | 1 | 295 | 33.7 | 4.77 | 2.27 | 1 | 0 | 0 | 52.5 | 0 | 0 | 51.7 | 0 | 0 | 42.9 | 0.985 | 0.9918573 | 0.8171429 | 0.8945024 | 0.8297872 | 0.902312 |
| A0A3Q0DAY1 | complement C5 OS=Mesocricetus auratus OX=10036 GN=LOC101837460 PE=4 SV=1 | 1.341 | 2 | 1 | 2 | 1 | 562 | 63.3 | 5.31 | 2.56 | 1 | 44.4 | 0 | 50.5 | 61.7 | 0 | 64.3 | 33.6 | 0 | 31.9 | 1.329 | 0.7155675 | 0.6902002 | 0.6416524 | 0.5198413 | 0.4569344 |
| A0A3Q0DAT1 | zinc finger protein 76 isoform X4 OS=Mesocricetus auratus OX=10036 GN=Znf76 PE=4 SV=1 | 1.312 | 2 | 1 | 1 | 1 | 534 | 58.1 | 6.42 | 2.42 | 1 | 0 | 64.4 | 0 | 0 | 57.9 | 0 | 0 | 61.1 | 0 | 0.898 | 0.943812 | 0.9487578 | 0.9721325 | 1.0552677 | 0.9715019 |
| A0A3Q0D4P2 | regulator of nonsense transcripts 2 OS=Mesocricetus auratus OX=10036 GN=Upf2 PE=4 SV=1 | 1.31 | 1 | 1 | 1 | 1 | 1267 | 147 | 5.62 | 2.53 | 1 | 0 | 0 | 48.6 | 0 | 0 | 59.9 | 0 | 0 | 37.5 | 1.233 | 0.8909032 | 0.7716049 | 0.8658341 | 0.6260434 | 0.7699663 |
| A0A1U7RJ41 | MAP7 domain-containing protein 1-like isoform X2 OS=Mesocricetus auratus OX=10036 GN=LOC101822374 PE=3 SV=2 | 1.299 | 3 | 1 | 2 | 1 | 426 | 47.3 | 8.95 | 0 | 1 | 159.4 | 0 | 0 | 128.8 | 0 | 0 | 121.2 | 0 | 0 | 0.808 | 0.888832 | 0.7603513 | 0.8586282 | 0.9409938 | 0.9677904 |
| A0A3Q0CY91 | acidic amino acid decarboxylase GADL1 OS=Mesocricetus auratus OX=10036 GN=Gadl1 PE=3 SV=1 | 1.296 | 2 | 1 | 5 | 1 | 523 | 59.2 | 6.84 | 0 | 1 | 261.8 | 357.8 | 373.5 | 207.9 | 319.2 | 295.2 | 198.3 | 280.7 | 291.7 | 0.79 | 0.3065354 | 0.7760548 | 0.1819943 | 0.9372492 | 0.7212489 |
