## Supplementary table 2 for "Quantitative proteomics of hamster lung tissues infected with SARS-CoV-2 reveal host-factors having implication in the disease pathogenesis and severity"

**Supplementary Table 2: Top GO-Biological process of proteins significantly upregulated and downregulated in infected (4dpi) lung tissues as compared to mock-infected (4dpi) using Enrichr.**

| Term | p-value | Adjusted p-value | Odds Ratio | Combined Score | Genes | Regulation |
|---|---|---|---|---|---|---|
| platelet degranulation (GO:0002576) | 4.76E-15 | 2.58E-12 | 75.71700992 | 2497.074896 | LGALS3BP;FGB;FGA;ORM1;TF;TMSB4X;FGG;SERPINF2;ALB;APOA1 | |
| regulated exocytosis (GO:0045055) | 2.9E-14 | 7.85E-12 | 62.47321991 | 1947.487298 | LGALS3BP;FGB;FGA;ORM1;TF;TMSB4X;FGG;SERPINF2;ALB;APOA1 | |
| positive regulation of substrate adhesion-dependent cell spreading (GO:1900026) | 7E-10 | 0.000000127 | 161.8912338 | 3412.546035 | FGB;FGA;FGG;APOA1;S100A10 | |
| regulation of substrate adhesion-dependent cell spreading (GO:1900024) | 6.4E-09 | 0.000000867 | 98.8640873 | 1865.308378 | FGB;FGA;FGG;APOA1;S100A10 | |
| fibrinolysis (GO:0042730) | 0.000000011 | 0.00000119 | 229.3678161 | 4202.87193 | FGB;FGA;FGG;SERPINF2 | |
| positive regulation of cell-substrate adhesion (GO:0010811) | 2.19E-08 | 0.00000181 | 75.68389058 | 1334.808942 | FGB;FGA;FGG;APOA1;S100A10 | |
| regulation of heterotypic cell-cell adhesion (GO:0034114) | 2.34E-08 | 0.00000181 | 183.4666667 | 3223.733837 | FGB;FGA;FGG;APOA1 | |
| positive regulation of cell morphogenesis involved in differentiation (GO:0010770) | 3.83E-08 | 0.0000026 | 67.09568733 | 1145.765041 | FGB;FGA;FGG;APOA1;S100A10 | Upregulated |
| cellular protein metabolic process (GO:0044267) | 6.08E-08 | 0.00000366 | 15.38842105 | 255.6874559 | C3;C4A;FGA;TF;WARS;FGG;ALB;CTSH;APOA1 | |
| platelet aggregation (GO:0070527) | 0.000000243 | 0.000012 | 94.82996433 | 1444.319374 | FGB;FGA;FGG;HBB | |
| positive regulation of cell-cell adhesion (GO:0022409) | 0.000000243 | 0.000012 | 94.82996433 | 1444.319374 | FGB;FGA;FGG;SERPINF2 | |
| cellular macromolecular complex assembly (GO:0034622) | 0.000000275 | 0.0000124 | 91.66436782 | 1384.738598 | FGB;FGA;FGG;PDE4DIP | |
| zymogen activation (GO:0031638) | 0.000000348 | 0.0000135 | 85.92672414 | 1277.712529 | FGB;FGA;FGG;CTSH | |
| negative regulation of blood coagulation (GO:0030195) | 0.000000348 | 0.0000135 | 85.92672414 | 1277.712529 | FGB;FGA;FGG;SERPINF2 | |
| homotypic cell-cell adhesion (GO:0034109) | 0.000000436 | 0.0000157 | 80.86409736 | 1184.373928 | FGB;FGA;FGG;HBB | |
| wound healing, spreading of cells (GO:0044319) | 0.000128 | 0.031754842 | 147.8888889 | 1325.54864 | COL5A1;RHOA | |
| collagen fibril organization (GO:0030199) | 0.000272 | 0.033775447 | 98.54814815 | 808.9130319 | COL1A2;COL5A1 | |
| protein complex subunit organization (GO:0071822) | 0.000659 | 0.054468982 | 61.82945736 | 452.8970859 | COL1A2;COL5A1 | |
| actin cytoskeleton reorganization (GO:0031532) | 0.001208274 | 0.068655974 | 45.0259887 | 302.5099237 | PARVA;RHOA | |
| Rho protein signal transduction (GO:0007266) | 0.00167838 | 0.068655974 | 37.92952381 | 242.3668496 | COL1A2;RHOA | |
| transforming growth factor beta receptor signaling pathway (GO:0007179) | 0.001867701 | 0.068655974 | 35.87207207 | 225.3859127 | COL1A2;RHOA | |
| negative regulation of kinase activity (GO:0033673) | 0.00266358 | 0.068655974 | 29.80374532 | 176.6791137 | PRDX3;GNAQ | |
| skin development (GO:0043588) | 0.002838851 | 0.068655974 | 28.82753623 | 169.0549316 | COL1A2;COL5A1 | Downregulated |
| cellular response to transforming growth factor beta stimulus (GO:0071560) | 0.003205288 | 0.068655974 | 27.05442177 | 155.3722787 | COL1A2;RHOA | |
| mRNA 5'-splice site recognition (GO:0000395) | 0.005089741 | 0.068655974 | 249.725 | 1318.679916 | SRSF1 | |
| response to chemokine (GO:1990868) | 0.005089741 | 0.068655974 | 249.725 | 1318.679916 | RHOA | |
| cellular response to chemokine (GO:1990869) | 0.005089741 | 0.068655974 | 249.725 | 1318.679916 | RHOA | |
| collagen biosynthetic process (GO:0032964) | 0.005089741 | 0.068655974 | 249.725 | 1318.679916 | COL5A1 | |
| negative regulation of cell size (GO:0045792) | 0.005089741 | 0.068655974 | 249.725 | 1318.679916 | RHOA | |
| inner ear receptor stereocilium organization (GO:0060122) | 0.005935662 | 0.068655974 | 208.09375 | 1066.850192 | CLIC5 | |
