## Supplementary table 3 for "Quantitative proteomics of hamster lung tissues infected with SARS-CoV-2 reveal host-factors having implication in the disease pathogenesis and severity"

**Supplementary Table 3: Top GO-Biological process among of proteins significantly upregulated and downregulated in infected (14dpi) lung tissues as compared to mock-infected (4dpi) using Enrichr.**

| Term | p-value | Adjusted p-value | Odds Ratio | Combined Score | Genes | Regulation |
|---|---|---|---|---|---|---|
| positive regulation of substrate adhesion-dependent cell spreading (GO:1900026) | 6.28E-05 | 0.008492914 | 228.1828571 | 2207.749648 | FGB;S100A10 | Upregulated |
| regulation of substrate adhesion-dependent cell spreading (GO:1900024) | 1.46E-04 | 0.008492914 | 146.1684982 | 1290.681803 | FGB;S100A10 | |
| positive regulation of blood vessel endothelial cell migration (GO:0043536) | 1.76E-04 | 0.008492914 | 132.5448505 | 1145.535735 | TMSB4X;HMGB1 | |
| pattern recognition receptor signaling pathway (GO:0002221) | 2.01E-04 | 0.008492914 | 123.8819876 | 1054.586314 | FGB;HMGB1 | |
| regulation of blood vessel endothelial cell migration (GO:0043535) | 2.18E-04 | 0.008492914 | 118.7083333 | 1000.80642 | TMSB4X;HMGB1 | |
| positive regulation of cell-substrate adhesion (GO:0010811) | 2.36E-04 | 0.008492914 | 113.9485714 | 951.703254 | FGB;S100A10 | |
| positive regulation of cell morphogenesis involved in differentiation (GO:0010770) | 2.94E-04 | 0.009062158 | 101.7091837 | 827.2024469 | FGB;S100A10 | |
| positive regulation of endothelial cell migration (GO:0010595) | 4.28E-04 | 0.011552295 | 83.71008403 | 649.3146812 | TMSB4X;HMGB1 | |
| toll-like receptor signaling pathway (GO:0002224) | 6.45E-04 | 0.015483336 | 67.71088435 | 497.4071883 | FGB;HMGB1 | |
| platelet degranulation (GO:0002576) | 0.001334166 | 0.027464838 | 46.53161593 | 308.0136431 | FGB;TMSB4X | |
| regulated exocytosis (GO:0045055) | 0.001892465 | 0.027464838 | 38.83561644 | 243.4944567 | FGB;TMSB4X | |
| positive regulation of ATP biosynthetic process (GO:2001171) | 0.002697256 | 0.027464838 | 499.65 | 2955.689739 | TMSB4X | |
| membrane raft assembly (GO:0001765) | 0.002697256 | 0.027464838 | 499.65 | 2955.689739 | S100A10 | |
| T-helper 1 type immune response (GO:0042088) | 0.003146172 | 0.027464838 | 416.3541667 | 2398.853181 | HMGB1 | |
| regulation of toll-like receptor 9 signaling pathway (GO:0034163) | 0.003146172 | 0.027464838 | 416.3541667 | 2398.853181 | HMGB1 | |
| antimicrobial humoral immune response mediated by antimicrobial peptide (GO:0061[illegible] | 2.09E-04 | 0.027846683 | 121.2401216 | 1027.072421 | S100A9;CAMP | Downregulated |
| mRNA 5'-splice site recognition (GO:0000395) | 0.002697256 | 0.049263118 | 499.65 | 2955.689739 | SRSF1 | |
| oxygen homeostasis (GO:0032364) | 0.002697256 | 0.049263118 | 499.65 | 2955.689739 | SOD2 | |
| negative regulation of systemic arterial blood pressure (GO:0003085) | 0.002697256 | 0.049263118 | 499.65 | 2955.689739 | SOD2 | |
| leukocyte aggregation (GO:0070486) | 0.003146172 | 0.049263118 | 416.3541667 | 2398.853181 | S100A9 | |
| gas homeostasis (GO:0033483) | 0.003146172 | 0.049263118 | 416.3541667 | 2398.853181 | SOD2 | |
| removal of superoxide radicals (GO:0019430) | 0.004043466 | 0.049263118 | 312.234375 | 1720.615303 | SOD2 | |
| vasodilation (GO:0042311) | 0.004043466 | 0.049263118 | 312.234375 | 1720.615303 | SOD2 | |
| cellular response to superoxide (GO:0071451) | 0.004043466 | 0.049263118 | 312.234375 | 1720.615303 | SOD2 | |
| defense response to bacterium (GO:0042742) | 0.004923101 | 0.049263118 | 23.61267185 | 125.4734102 | S100A9;CAMP | |
| alternative mRNA splicing, via spliceosome (GO:0000380) | 0.005388063 | 0.049263118 | 227.0454545 | 1185.987693 | SRSF1 | |
| phospholipase C-activating dopamine receptor signaling pathway (GO:0060158) | 0.005835903 | 0.049263118 | 208.1145833 | 1070.484451 | GNAQ | |
| phototransduction, visible light (GO:0007603) | 0.006731046 | 0.049263118 | 178.3660714 | 892.01312 | GNAQ | |
| G-protein coupled acetylcholine receptor signaling pathway (GO:0007213) | 0.00717835 | 0.049263118 | 166.4666667 | 821.7936241 | GNAQ | |
| detection of visible light (GO:0009584) | 0.00717835 | 0.049263118 | 166.4666667 | 821.7936241 | GNAQ | |
